# Pan-cancer whole genome analyses of metastatic solid tumors

**DOI:** 10.1101/415133

**Authors:** Peter Priestley, Jonathan Baber, Martijn P. Lolkema, Neeltje Steeghs, Ewart de Bruijn, Charles Shale, Korneel Duyvesteyn, Susan Haidari, Arne van Hoeck, Wendy Onstenk, Paul Roepman, Mircea Voda, Haiko J. Bloemendal, Vivianne C.G. Tjan-Heijnen, Carla M.L. van Herpen, Mariette Labots, Petronella O. Witteveen, Egbert F. Smit, Stefan Sleijfer, Emile E. Voest, Edwin Cuppen

**Affiliations:** Hartwig Medical Foundation, Science Park 408, Amsterdam, The Netherlands; Hartwig Medical Foundation Australia, Sydney, Australia; Center for Personalized Cancer Treatment, The Netherlands; Erasmus MC Cancer Institute, Doctor Molewaterplein 40, Rotterdam, The Netherlands; Netherlands Cancer Institute/Antoni van Leeuwenhoekhuis, Plesmanlaan 121, Amsterdam, The Netherlands; Center for Molecular Medicine and Oncode Institute, University Medical Center Utrecht, Heidelberglaan 100, Utrecht, The Netherlands; Meander Medisch Centrum, Maatweg 3, Amersfoort, The Netherlands; Maastricht University Medical Center, P. Debyelaan 25, Maastricht, The Netherlands; Radboud University Medical Center, Geert Grooteplein Zuid 10, Nijmegen, The Netherlands; VU Medical Center, De Boelelaan 1117, Amsterdam, The Netherlands; Cancer Center, University Medical Center Utrecht, Heidelberglaan 100, Utrecht, The Netherlands

## Abstract

Metastatic cancer is one of the major causes of death and is associated with poor treatment efficiency. A better understanding of the characteristics of late stage cancer is required to help tailor personalised treatment, reduce overtreatment and improve outcomes. Here we describe the largest pan-cancer study of metastatic solid tumor genomes, including 2,520 whole genome-sequenced tumor-normal pairs, analyzed at a median depth of 106x and 38x respectively, and surveying over 70 million somatic variants. Metastatic lesions were found to be very diverse, with mutation characteristics reflecting those of the primary tumor types, although with high rates of whole genome duplication events (56%). Metastatic lesions are relatively homogeneous with the vast majority (96%) of driver mutations being clonal and up to 80% of tumor suppressor genes bi-allelically inactivated through different mutational mechanisms. For 62% of all patients, genetic variants that may be associated with outcome of approved or experimental therapies were detected. These actionable events were distributed across various mutation types underlining the importance of comprehensive genomic tumor profiling for cancer precision medicine.

---

Metastatic cancer is one of the leading causes of death globally and is a major burden for society despite the availability of an increasing number of (targeted) drugs. Health care costs associated with treatment of metastatic disease are increasing rapidly due to the high cost of novel targeted treatments and immunotherapy, while many patients do not benefit from these approaches with inevitable adverse effects for most patients. Metastatic cancer therefore poses a major challenge for society to balance between individual and societal treatment responsibilities. Since cancer genomes evolve over time, both in the highly heterogeneous primary tumor mass and as disseminated metastatic cells^1,2^, a better understanding of metastatic cancer genomes is crucial to further improve on tailoring treatment for late stage cancers.

In recent years, several large-scale whole genome sequencing (WGS) analysis efforts such as TCGA and ICGC have yielded valuable insights in the diversity of the molecular processes driving different types of adult^3,4^ and pediatric^5,6^ cancer and have fueled the promises of genome-driven oncology care^7^. However, most analyses were done on primary tumor material whereas metastatic cancers, which cause the bulk of the disease burden and 90% of all cancer deaths, have been less comprehensively studied at the whole genome level, with previous efforts focusing on tumor-specific cohorts^8–10^ or at a targeted gene panel^11^ or exome level^12^.

Here we describe the first large-scale pan-cancer whole-genome landscape of metastatic cancers based on the Hartwig Medical Foundation (HMF) cohort of 2,520 paired tumor and normal genomes from 2,399 patients, collected prospectively in 41 hospitals in the Netherlands (Supplementary Table 1, Extended Data Fig. 1). All samples were paired with standardized clinical information (Supplementary Table 2). The sample distribution over age and primary tumor types broadly reflects solid cancer incidence in the Western world, including rare cancers (Fig. 1a-b).

**Figure 1:**
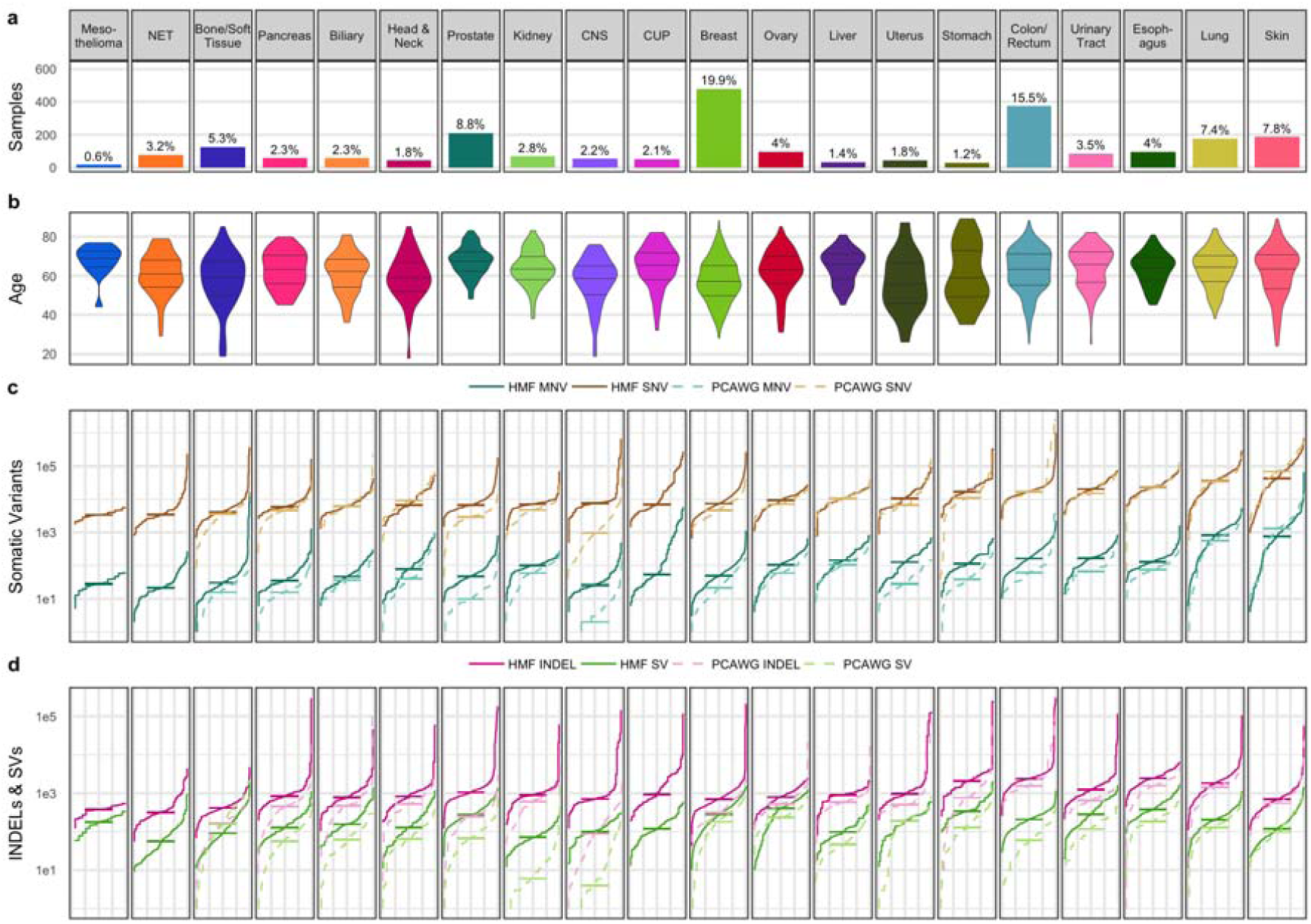
Mutational load of metastatic cancer per tumor type. a) The number of samples of each tumor type cohort with more than 10 samples. Tumor types are ranked from lowest to highest overall mutation burden (TMB). b) Violin plot showing age distribution of each tumor type with 25th, 50th and 75th percentiles marked. c)-d) cumulative distribution function plot (individual samples were ranked independently for each variant type) of mutational load for each tumor type for SNV and MNV (c) and INDEL and SV (d). The median for each cohort is indicated with a vertical line. Dotted lines indicate the mutational loads in primary cancers from the PCAWG cohort^14^.

The cohort has been analyzed with paired-end whole genome sequencing with a median depth of 106x for tumor samples and 38x for the blood control (Extended Data Fig. 1). Sequencing data were analyzed for all types of somatic variants using an optimized bioinformatic pipeline based on open source tools (Methods, Supplementary Information). We identified a total of 59,472,629 single nucleotide variants (SNVs), 839,126 multiple nucleotide variants (MNVs), 9,598,205 insertions and deletions (INDELs) and 653,452 structural variants (SVs) (Supplementary Table 2).

## Mutational landscape of metastatic cancer

We analysed the tumor mutational burden (TMB) of each class of variants per cancer type based on tissue of origin (Fig. 1, Supplementary Table 2). In line with previous studies on primary cancers^13,14^, we found extensive variation in mutational load of up to 3 orders of magnitude both within and across cancer types.

The median SNV counts per sample were highest in skin, predominantly consisting of melanoma (44k) and lung (36k) tumors with ten-fold higher SNV counts than sarcomas (4.1k), neuroendocrine tumors (NET) (3.5k) and mesotheliomas (3.4k). SNVs were mapped to COSMIC mutational signatures and were found to broadly match the patterns described in previous cancer cohorts per cancer type (Extended Data Figs. 2, 3)^13^. However, a number of broad spectrum signatures particularly S3, S8, S9, and S16 as well as some more specific signature (e.g. S17 in specific tumor types) appear to be overrepresented in our cohort. These observations may indicate enrichment of tumors deficient in specific DNA repair processes (S3) or increased hypermutation processes (S9) among advanced cancers or reflect mutagenic effects of previous treatments (Extended Data Fig. 3).

The variation for MNVs was even greater with lung (median=821) and skin (median=764) tumors having five times the median MNV counts of any other tumor type. This can be explained by the well-known mutational impact of UV radiation (CC->TT MNV) and smoking (CC->AA MNV) mutational signatures, respectively (Extended Data Fig. 2). Although only di-nucleotide substitutions are typically reported as MNVs, 10.7% of the MNVs involve three nucleotides and 0.6% had four or more nucleotides affected.

INDEL counts were typically ten-fold lower than SNVs, with a lower relative rate for skin and lung cancers (Fig. 1d, Extended Data Fig. 2). Genome-wide analysis of INDELs at microsatellite loci identified 60 samples with microsatellite instability (MSI) (Supplementary Table 2), representing 2.5% of all tumors. The highest rates of MSI were observed in central nervous system (CNS) (9.4%), uterus (9.1%) and prostate (6.1%) tumors. For metastatic colorectal cancer lesions we found an MSI frequency of only 4.0%, which is lower than reported for primary colorectal cancer, and in line with better prognosis for patients with localized MSI colorectal cancer, which less often metastasizes^15^. Remarkably, 67% of all INDELs in the entire cohort were found in the 60 MSI samples, and 85% of all INDELs in the cohort were found in microsatellites or short tandem repeats. Only 0.33% of INDELs (32k, ∼1% of non-microsatellite INDELs) were found in coding sequences, of which the majority (88%) had a predicted high impact by affecting the open reading frame of the gene.

The median rate of SVs across the cohort was 193 per tumor, with the highest median counts observed in ovary (412) and esophageal (372) tumors, and the lowest in kidney tumors (71) and NET (56) (Fig. 1d). Simple deletions were the most commonly observed SV subtype (33% of all SVs) and were the most prevalent in every cancer type except stomach and esophageal tumors which were highly enriched in translocations (Extended Data Fig. 2).

To gain insight into the overall genomic differences between primary and metastatic cancer, we compared the TMB of the HMF cohort against the PCAWG dataset^14^ which is the largest comparable whole genome sequenced tumor cohort available to date and which has 95% of biopsies taken from treatment-naive primary tumors. SNV mutational load does not, in general, appear to be indicative for disease progression as it is not significantly different in this study compared to PCAWG for the majority of cancer types (Fig. 1c). Prostate cancer and breast cancer are clear exceptions with structurally higher mutational loads (q<1e^−10^, Mann-Whitney Test) potentially reflecting relevant tumor biology and is, for prostate, consistent with other reports^10,16^. CNS tumors also have a higher mutational load but this may be explained by the different age distributions of the cohorts.

In contrast, INDEL, MNV, and SV mutational loads are significantly higher across nearly all cancer types analyzed (Fig. 1d). This is most notable for prostate cancer where we observe a more than four-fold increased rate of each of MNV, INDEL & SV. Whilst these observations may represent the advancement of disease and higher rate of certain mutational processes in metastatic cancers, they are also partially due to differences in sequencing depth and bioinformatic analysis pipelines (Extended Data Figs. 4, 5, Supplementary Information).

## Copy number alteration landscape

Copy number alterations (CNAs) are important hallmarks of tumorigenesis^17^. Pan-cancer, the most highly amplified regions in our metastatic cancer cohort contain the established oncogenes such as EGFR, CCNE1, CCND1 and MDM2 (Fig. 2). Chromosomal arms 1q, 5p, 8q and 20q are also highly enriched in moderate amplification across the cohort each affecting >20% of all samples. For the amplifications of 5p and 8q this is likely related to the common amplification targets of TERT and MYC, respectively. However, the targets of the amplifications on 1q, predominantly found in breast cancers (>50% of samples) and amplifications on 20q, predominantly found in colorectal cancers (>65% of samples), are less clear.

**Figure 2:**
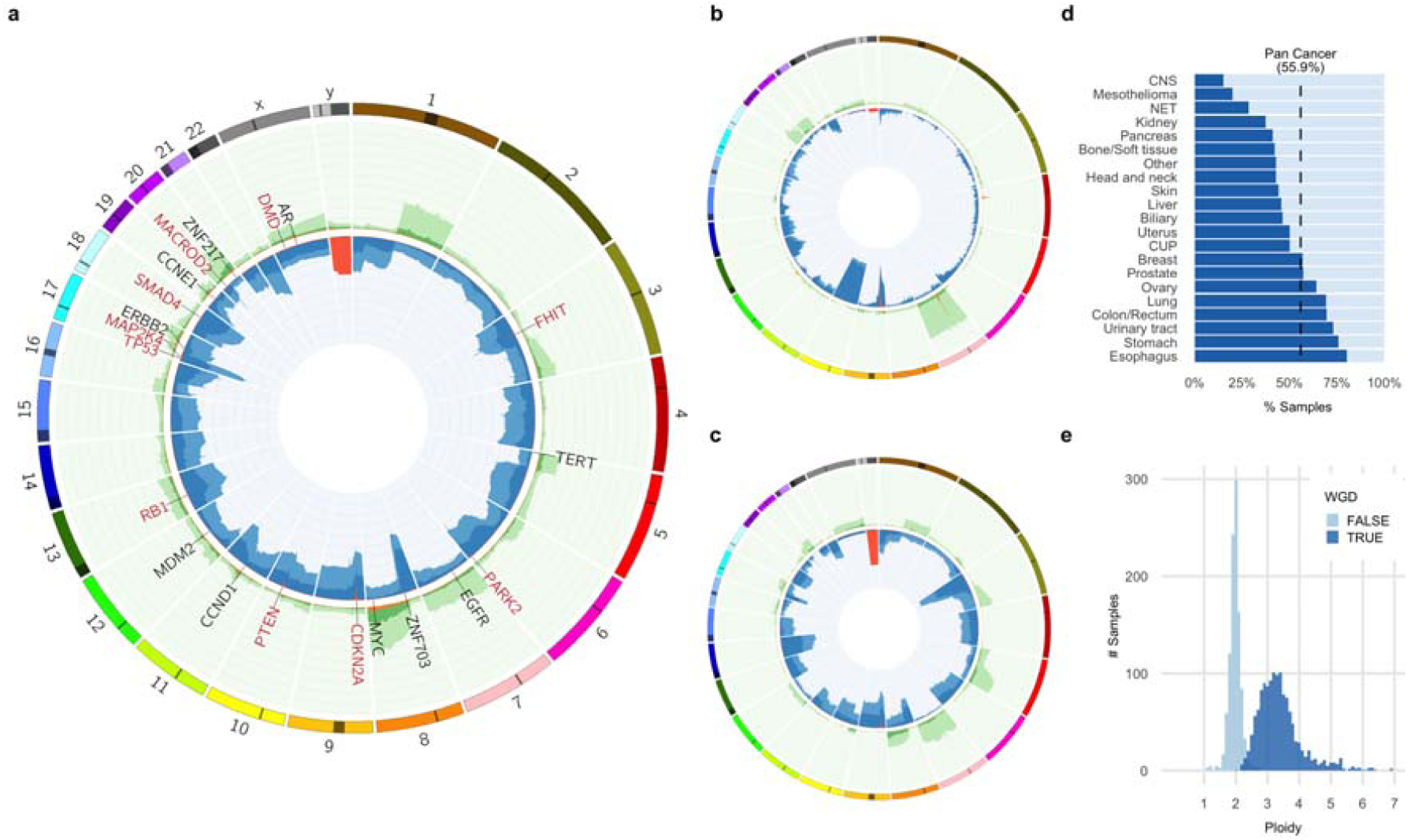
Copy number landscape of metastatic cancer. Proportion of samples with amplification and deletion events by genomic position per cohort – pan-cancer (a), central nervous system (CNS) (b) and kidney (c). The inner ring shows the % of tumors with homozygous deletion (orange), LOH and significant loss (copy number < 0.6x sample ploidy – dark blue) and near copy neutral LOH (light blue). Outer ring shows % of tumors with high level amplification (>3x sample ploidy – orange), moderate amplification (>2x sample ploidy – dark green) and low level amplification (>1.4x amplification – light green). The scale on both rings is 0-100% and inverted for the inner ring. The most frequently observed high-level gene amplifications (black text) and homozygous deletions (red text) are shown for pan-cancer. d) Proportion of tumors with a whole genome duplication (WGD) event (dark blue) grouped by tumor type. e) Sample ploidy distribution over the complete cohort for samples with and without WGD.

We identified some intriguing patterns of recurrent loss of heterozygosity (LOH) caused by CNAs. Overall an average of 23% of the autosomal DNA per tumor has LOH. Unsurprisingly, TP53 has the highest LOH recurrence at 67% of samples. Many of the other LOH peaks are also explained by well-known tumor suppressor genes (TSG). However, several clear LOH peaks are observed which cannot easily be explained by known TSG selection, such as one on 8p (57% of samples). 8p LOH has previously been linked to lipid metabolism and drug response^18^, although involvement of individual genes has not been established.

There are remarkable differences in LOH between cancer types (Fig. 2, Supplementary Image File 1). For instance, we observed LOH events on the 3p arm in 90% of kidney samples^19^ and LOH of the complete chromosome 10 in 72% of CNS tumors (predominantly glioblastoma multiforme^20^). Even in the case of the TP53 region on chromosome 17, different tumor types display clearly different patterns of LOH. Ovarian cancers exhibit LOH of the full chromosome 17 in 75% of samples whereas in prostate cancer, which also has 70% LOH for TP53, this is nearly always caused by highly focal deletions.

Unlike LOH events, homozygous deletions are nearly always restricted to small chromosomal regions. Not a single example was found in which a complete autosomal arm was homozygously deleted. Homozygous deletions of genes are surprisingly rare as well: we found only a mean of 2.0 instances per tumor where one or multiple consecutive genes are fully or partially homozygously deleted. In 46% of these events a putative TSG was deleted. The scarcity of passenger homozygous deletions underlines the fact that despite widespread copy number alterations in metastatic tumors, the vast majority of genes or gross chromosomal organization likely remain essential for tumor cell survival. Chromosome Y loss, which has been described anecdotally for various tumor types^21,22^, is a special case and is deleted in 36% of all male tumor genomes but varies strongly between tumor types from 5% to 68% for CNS and biliary tumors respectively (Extended Data Fig. 6).

An extreme form of copy number change can be caused by whole genome duplication (WGD). We found WGD events in 56% of all samples ranging between 15% in CNS to 80% in esophageal tumors (Fig. 2). This is much higher than reported previously for primary tumors (25%-37%)^23,24^ and also higher than estimated from panel-based sequencing analyses of advanced tumors (30%)^25^. Ploidy levels, in combination with accurate tumor purity information, are essential for correct interpretation of the measured raw SNV and INDEL frequencies, e.g. to discriminate bi-allelic inactivation of TSG from heterozygous events which are more likely to be passengers or to determine (sub)clonality. Hence determining the WGD status of a tumor is highly relevant for diagnostic applications. Furthermore, WGD has previously been found to correlate with a greater incidence of cancer recurrence for ovarian cancer^24^ and has been associated with poor prognosis across cancer types, independently of established clinical prognostic factors^25^.

## Significantly mutated genes

To identify significantly mutated genes (SMGs) potentially specific for metastatic cancer, we used the dNdScv approach^26^ with strict cutoffs (q<0.01) for the pan-cancer and tumor-type specific datasets. In addition to reproducing previous results on cancer drivers, a few novel genes were identified (Extended Data Fig. 8, Supplementary Table 3). In the pan-cancer analyses we found only a single novel SMG, which was not either present in the curated COSMIC Cancer Gene Census or found by Martincorena et al^26^. This gene, MLK4 (q=2e^−4^), is a mixed lineage kinase that regulates the JNK,P38 and ERK signaling pathways and has been reported to inhibit tumorigenesis in colorectal cancer^27^. In addition, in our tumor type-specific analyses, which for several tumor types is limited by the number of samples, we identified a novel metastatic breast cancer-specific SMG – ZFPM1 (also known as Friend of GATA1 (FOG1), q=8e^−5^), a zinc-finger transcription factor protein without clear links with cancer. Nonetheless, we found six unique frameshift variants (all in a context of biallelic inactivation) and three nonsense variants, which suggests a driver role for this gene in metastatic breast cancer.

Our cohort also lends support to some prior SMG findings. In particular, eight significantly mutated putative TSG in the HMF cohort were also found by Martincorena et al^26^ – GPS2 (pan-cancer, q=1e^−5^ & breast, q=2e^−3^), SOX9 (colorectal & pan-cancer, q=0), TGIF1 (pan-cancer, q=3e^−3^ & colorectal q=6e^−3^), ZFP36L1 (urinary tract q=3e^−4^, pan-cancer q=9e^−4^) and ZFP36L2 (colorectal & pan-cancer, q=0), HLA-B (lymphoid, q=5e^−5^), MGA (pan-cancer, q=4e^−3^), KMT2B (skin, q=3e^−3^) and RARG (urinary tract 8e^−4^). None of these genes are currently included in the COSMIC Cancer Gene Census^28^. ZFP36L1 and ZFP36L2 are of particular interest as these genes are zinc-finger proteins that normally play a repressive regulatory role in cell proliferation, presumably through a cyclin D dependent and p53 independent pathway^29^. ZFP36L2 is also independently found as a significantly deleted gene in our cohort, most prominently in colorectal and prostate cancers.

We also searched for genes that were significantly amplified or deleted (Supplementary Table 4). CDKN2A and PTEN were the most significantly deleted genes overall, but many of the top genes involved common fragile sites (CFS) particularly FHIT and DMD, deleted in 5% and 4% of samples, respectively. The role of CFS in tumorigenesis is unclear and aberrations affecting these genes are frequently treated as passenger mutations reflecting localized genomic instability^30^. However, the uneven distribution of the deletions across cancer types may indicate that some of these could be genuine tumor-type specific cancer drivers. For example, we find deletions in DMD to be highly enriched in esophageal tumors (deleted in 38% of samples, whilst SV burden in this tumortype is only about 2-fold higher than average), GIST (Gastro-Intestinal Stromal Tumors; 24%) and pancreatic neuroendocrine tumors (panNET; 41%), which is consistent with a recent study that indicated DMD as a TSG in cancers with myogenic programs^31^. However, tissue type-specific gene expression and differences in origins of replication may also contribute to the observed patterns^30^. We also identified several significantly deleted genes not reported previously, including MLLT4 (n=13) and PARD3 (n=9).

Unlike homozygous deletions, amplification peaks tend to be broad and often encompass large number of genes, making identification of the amplification target challenging. However, SRY-related high-mobility group box 4 gene (SOX4, 6p22.3) stands out as a significantly amplified single gene peak (26 amplifications) and is highly enriched in urinary tract cancers (19% of samples highly amplified). SOX4 is known to be over-expressed in prostate, hepatocellular, lung, bladder and medulloblastoma cancers with poor prognostic features and advanced disease status and is a modulator of the PI3K/Akt signaling^32^.

Also notable was a broad amplification peak of 10 genes around ZMIZ1 at 10q22.3 (n=32) which has not previously been reported. ZMIZ1 is a member of the Protein Inhibitor of Activated STAT (PIAS)-like family of coregulators and is a direct and selective cofactor of Notch1 in T-cell development and leukemia^33^. CDX2, previously identified as an amplified lineage-survival oncogene in colorectal cancer^34^, is also highly amplified in our cohort with 20 out of 22 amplified samples found in colorectal cancer, representing 5.4% of all colorectal samples.

## Driver mutation catalog

We created a comprehensive catalog of all cancer driver mutations across all samples in our cohort and all variant classes similar as described previously in primary tumors^35^ (N. Lopez, personal communication). To do this, we combined our SMG discovery efforts with those from Martincorena et al.^26^ and a panel of well known cancer genes (Cosmic Curated Genes)^36^, and added gene fusions, TERT promoter mutations and germline predisposition variants. Accounting for the proportion of SNV and INDELs estimated by dNdScv to be passengers, we found 13,384 somatic driver events among the 20,071 identified mutations in the combined gene panel (Supplementary Table 5) together with 189 germline predisposition variants (Supplementary Table 6). The somatic drivers include 7,400 coding mutation, 615 non-coding point mutation drivers, 2,700 homozygous deletions (25% of which are in common fragile sites), 2,392 focal amplifications and 276 fusion events.

For the cohort as a whole, 55% of point mutations in the gene panel driver catalog were predicted to be genuine driver events. To facilitate analysis of variants of unknown significance (VUS) at a per patient level, we calculated a sample-specific likelihood for each point mutation to be a driver taking into account the TMB of the sample as well as the biallelic inactivation status of the gene for TSG and hotspot positions in oncogenes. Predictions of pathogenic variant overlap with known biology, e.g. clustering of benign missense variants in the 3’ half of the APC gene (Supplementary Image File 2) fits with the absence of FAP-causing germline variants in this part of the gene^37^.

Overall, the catalog matches previous inventories of cancer drivers. TP53 (52% of samples), CDKN2A (21%), PIK3CA (16%), APC (15%), KRAS (15%), PTEN (13%) and TERT (12%) were the most common driver genes together making up 26% of all the driver mutations in the catalog (Fig. 3). However, all of the ten most prevalent driver genes in our cohort were reported at a higher rate than for primary cancers^38^, which may reflect the more advanced disease state. AR and ESR1 in particular are more prevalent, with driver mutations in 44% of prostate and 16% of breast cancers, respectively. Both genes are linked to resistance to hormonal therapy, a common treatment for these tumor types, and have been previously reported as enriched in advanced metastatic cancer^11^ but are identified at higher rates in this study.

**Figure 3:**
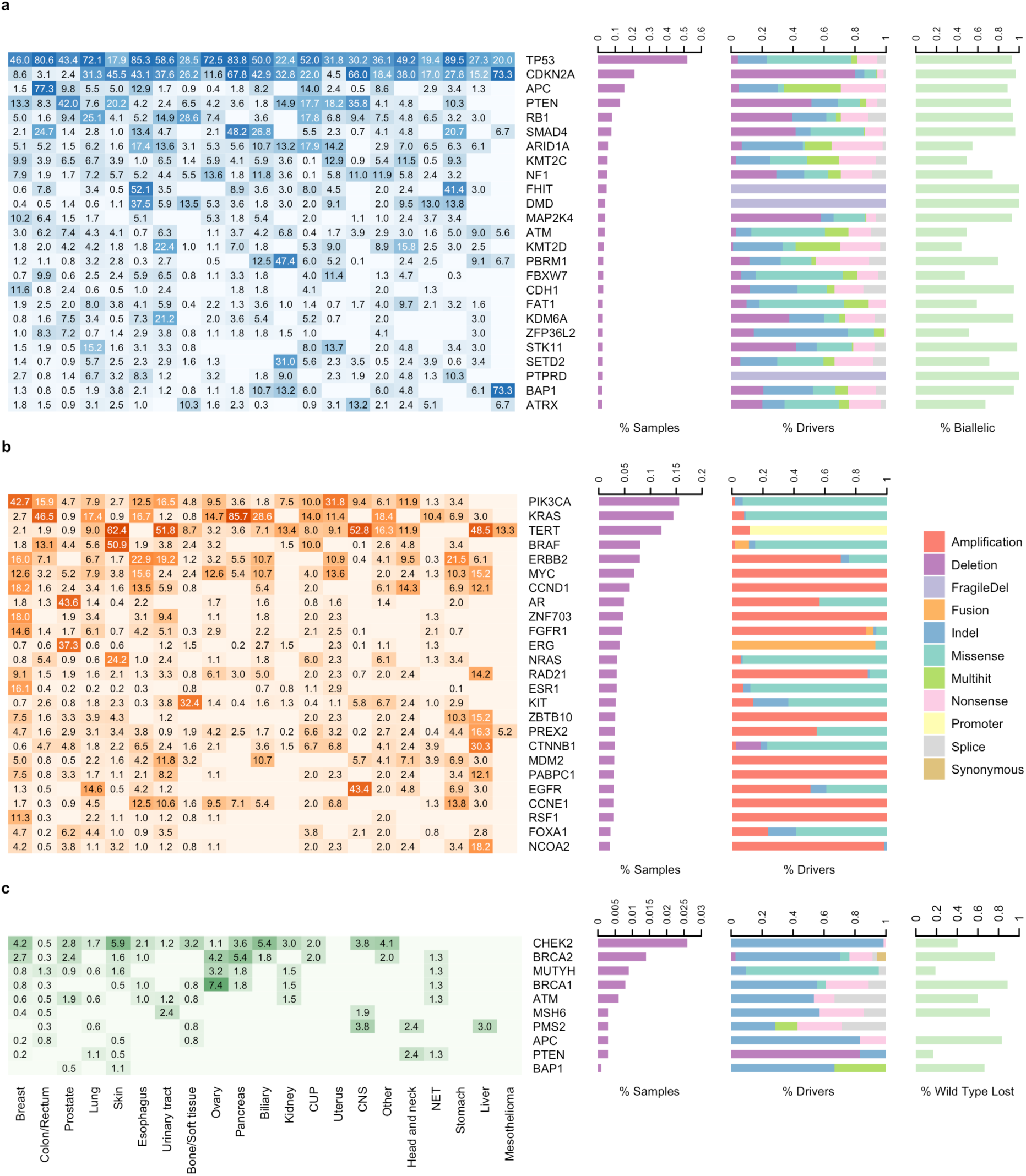
Most prevalent driver genes in metastatic cancer. Most prevalent somatically mutated TSG (a), oncogenes (b), and germline predisposition variants (c). From left to right, the heatmap shows the % of samples in each cancer type which are found to have each gene mutated; absolute bar chart shows the pan-cancer % of samples with the given gene mutated; relative bar chart shows the breakdown by type of alteration. For TSG, the % of samples with a driver in which the gene is found biallelically inactivated, and for germline predisposition variants the % of samples with loss of wild type in the tumor are also shown.

Looking at a per patient level, the mean number of total driver events per patient was 5.7, with the highest rate in urinary tract tumors (mean=8.0) and the lowest in NET (mean=2.8) (Fig. 4). Esophageal and stomach tumors also had elevated driver counts, largely due to a much higher rate of deletions in CFS genes (mean=1.6 for both stomach and esophageal) compared to other cancer types (pan-cancer mean=0.3). Fragile sites aside, the differential rates of drivers between cancer types in each variant class do correlate with the relative mutational load (Extended Data Fig. 6), with the exception of skin cancers, which have a lower than expected number of SNV drivers.

**Figure 4:**
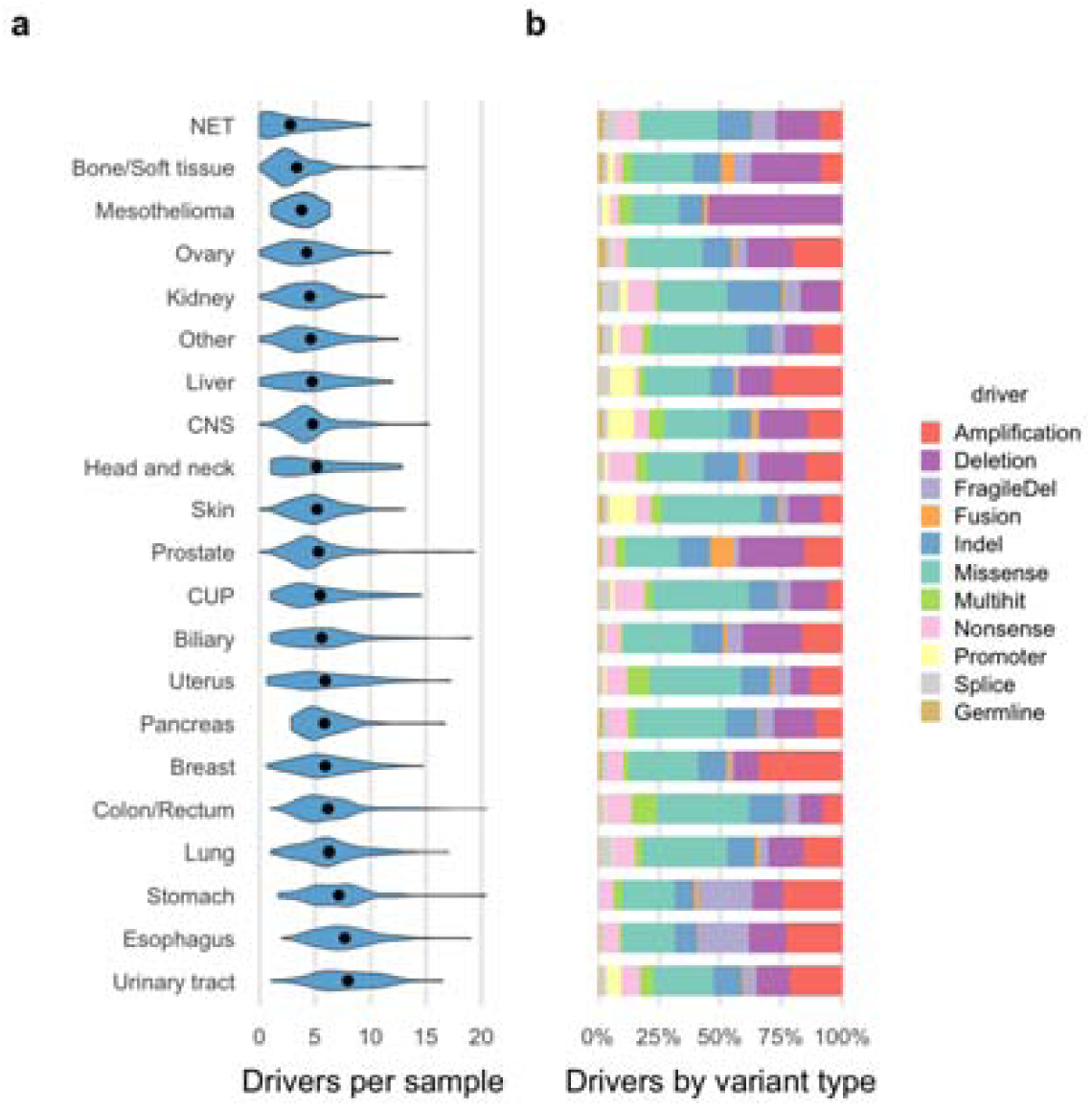
Number of drivers and type of mutation per sample by tumor type. a) Violin plot showing the distribution of the number of drivers per sample grouped by tumor type. Black dots indicate the mean values for each tumor type. b) Relative bar chart showing the breakdown per cancer type of the type of alteration.

In 98.6% of all samples at least one somatic driver mutation or germline predisposition variant was found. Of the 34 samples with no identified driver, 18 were NET of the small intestine (representing 49% of all patients of this subtype). This likely indicates that small intestine NETs have a distinct set of drivers that are not captured yet in any of the cancer gene resources used and are also not prevalent enough in our relatively small NET cohort to be detected as significant. Alternatively, NET tumors could be mainly driven by epigenetic mechanisms not detected by WGS^39^.

The number of amplified driver genes varied significantly between cancer types (Extended Data Fig. 7) with highly elevated rates per sample in breast cancer (mean = 2.1), esophageal (mean=1.8, urinary tract and stomach (both mean=1.7) cancers and nearly no amplification drivers in kidney cancer (mean=0.1) and none in the mesothelioma cohort. In tumor types with high rates of amplifications, these amplifications are generally found across a broad spectrum of oncogenes, suggesting there are mutagenic processes active in these tissues that favor amplifications, rather than tissue-specific selection of individual driver genes. AR and EGFR are notable exceptions, with highly selective amplifications in prostate, and in CNS and lung cancers, respectively, in line with previous reports^20,40,41^. Intriguingly, we also found two-fold more amplification drivers in samples with WGD despite amplifications being defined as relative to the average sample ploidy.

The 189 germline variants identified in 29 cancer predisposition genes (present in 7.9% of the cohort) consisted of 8 deletions and 181 point mutations (Fig. 3c, Supplementary Table 6). The top 5 affected genes were the well-known germline drivers CHEK2, BRCA2, MUTYH, BRCA1 and ATM, and together contain nearly 80% of the observed predisposition variants. The corresponding wild type alleles were found to be lost in the tumor sample in more than half of the cases, either by LOH or somatic point mutation, indicating a high penetrance for these variants, particularly in BRCA1 (89% of cases), APC (83%) and BRCA2 (79%).

The 276 fusions consisted of 168 in-frame coding fusions, 90 cis-activating fusions involving repositioning of regulatory elements in 5’ genic regions, and 18 in-frame intragenic deletions where one or more exons were deleted (Supplementary Table 7). ERG (n=88), BRAF (n=17), ERBB4 (n=16), ALK (n=12), NRG1 (9 samples) and ETV4 (n=7) were the most commonly observed 3’ partners together making up more than half of the fusions. 76 of the 89 ERG fusions were TMPRSS2-ERG affecting 36% of all prostate cancer samples in the cohort. 146 fusion pairs were not previously recorded in CGI, OncoKb, COSMIC or CIViC^36,42–44^. A novel recurrent KMT2A-BCOR fusion was observed (in sarcoma and stomach cancer) and there were also 3 recurrent novel localized fusions resulting from adjacent gene pairs: YWHAE-CRK (n=2), FGFR2-ATE1 (n=2) and BCR-GNAZ (n=2).

Only promoter mutations in TERT were included in the study due to the current lack of robust evidence for other recurrent oncogenic non-coding mutations^45^. A total of 257 variants were found at 5 known recurrent variant hotspots^11^ and included in the driver catalog.

We found that 71% of somatic driver point mutations in oncogenes occur at or within 5 nucleotides of already known pathogenic mutational hotspots. In the six most prevalent oncogenes (KRAS, PIK3CA, BRAF, NRAS, TERT & ESR1) the rate was 97% (Extended Data Fig. 9). Furthermore, in many of the key oncogenes, we document several likely activating but non-canonical variants near known mutational hotspots, particularly in-frame INDELs. Despite in-frame INDELs being exceptionally rare overall (mean=1.7 per tumor), we found an excess in known oncogenes including PIK3CA (n=18), ERBB2 (n=10) and BRAF (n=8) frequently occurring at or near known hotspots^46^ (Extended Data Fig. 9). Notably, many of the in-frame INDELs are enriched in specific tumor types. For instance, all 18 KIT in-frame INDELs were found in sarcomas, 6 out of 8 MUC6 in-frame INDELSs in esophageal tumors, and 6 of 10 ERBB2 in-frame INDELs in lung tumors. Finally, we identified 10 in-frame INDELs in FOXA1, which are highly enriched in prostate cancer (7 of 10 cases) and clustered in two locations that were not previously associated with pathogenic mutations^47^. In CTNNB1 we identified an interesting novel recurrent in-frame deletion of the complete exon 3 in 12 samples, 9 of which are colorectal cancers. Surprisingly, these deletions were homozygous but suspected to be activating as CTNNB1 normally acts as an oncogene in the WNT/beta-catenin pathway and none of these nine colorectal samples had any APC driver mutations.

For tumor supressor genes (TSG), our results strongly support the Knudson two-hit hypothesis^48^ with 80% of all TSG drivers found to have biallelic inactivation by genetic alterations (Fig. 3; i.e. either by homozygous deletion (32%), multiple somatic point mutations (7%), or a point mutation in combination with LOH (41%)). This rate is the highest observed in any large-scale cancer WGS study. For many key tumor suppressor genes the biallelic inactivation rate is almost 100% (e.g. TP53 (93%), CDKN2A (97%), RB1 (94%), PTEN (92%) and SMAD4 (96%)), suggesting that biallelic inactivation of these genes is a strong requirement for metastatic cancer. Other prominent TSGs, however, have lower biallelic rates, including ARID1A (55%), KMT2C (49%) and ATM (49%). It is unclear whether we systematically missed the second hit in these cases, as this could potentially be mediated through non-mutational epigenetic mechanisms^49^, or if these genes impact on tumorigenesis via a haploinsufficiency mechanism^50^.

We examined the pairwise co-occurrence of driver gene mutations per cancer type and found ten combinations of genes that were significantly mutually exclusively mutated, and ten combinations of genes which were significantly co-occurrently mutated (excluding pairs of genes on the same chromosome which are frequently co-amplified or co-deleted) (Extended Data Fig. 10). The 20 significant findings include previously reported co-occurrence of mutated DAX|MEN1 in pancreatic NET (q=7e^−4^), and CDH1|SPOP in prostate tumors (q=5e^−4^), as well as negative associations of mutated genes within the same signal transduction pathway such as KRAS|BRAF (q=4e^−4^) and KRAS|NRAS (q=0.008) in colorectal cancer, BRAF|NRAS in skin cancer (q=6e^−12^), CDKN2A|RB1 in lung cancer (q=8e^−5^) and APC|CTNNB1 in colorectal cancer (q=3e^−6^). APC is also strongly negatively correlated with both BRAF (q=9e^−5^) and RNF43 (q=4e^−6^) which together are characteristic of the serrated molecular subtype of colorectal cancers^51^. We also found that SMAD2|SMAD3 are highly positively correlated in colorectal cancer (q=0.02), mirroring a result reported previously in a large cohort of colorectal cancers^52^.

In breast cancer, we found a number of significant novel relationships, including a positive relationship for GATA3|VMP1(q=6e^−5^) and FOXA1|PIK3CA (q=3e^−3^), and negative relationships for ESR1|TP53 (q=9e^−4^) and GATA3|TP53 (q=5e^−5^), which will need further validation and experimental follow-up to understand underlying biology.

## Clonality of variants

To obtain insight into ongoing tumor evolution dynamics, we examined the clonality of all variants. Surprisingly, only 6.5% of all SNV, MNV & INDELs across the cohort and just 3.6% of the driver point mutations were found to be subclonal (Extended Data Fig. 11). The low proportion of samples with subclonal variants could be partially due to the detection limits of the sequencing approach (sequencing depth, bioinformatic analysis settings), particularly for low purity samples (Extended Data Fig. 11d). However, even for samples with purities higher than 80% the total proportion of subclonal variants only reaches 10.6%. Furthermore, sensitized detection of variants at hotspot positions in cancer genes showed that our analysis pipeline detected over 96% of variants with allele frequencies of > 3%. Although the cohort contains some samples with (very) high fractions of subclonal variants, overall the metastatic tumor samples are relatively homogeneous without the presence of multiple diverged major subclones. Low intratumor heterogeneity may be in part attributed to the fact that nearly all biopsies were obtained by a core needle biopsy, which results in highly localized sampling, but is nevertheless much lower compared to previous observations in primary cancers^2^.

In the 117 patients with independently collected repeat biopsies from the same patient (Supplementary Table 8) we found 11% of all SNVs to be subclonal. Whilst 71% of clonal variants were shared between biopsies, only 29% of the subclonal variants were shared. Although we can not exclude the presence of larger amounts of lower frequency subclonal variants, our results suggest a model where individual metastatic lesions are dominated by a single clone at any one point in time and that more limited tumor evolution and subclonal selection takes places after distant metastatic seeding. This contrasts with observations in primary tumors, where larger degrees of subclonality and multiple major subclones are more frequently observed^2,53^, but supports other recent studies which demonstrate minimal driver gene heterogeneity in metastases^8,54^.

## Clinical actionability

We analyzed opportunities for biomarker-based treatment for all patients by mapping driver events to three clinical annotation databases: CGI^44^, CIViC^42^ and OncoKB^43^. In 1,480 patients (62%) at least one ‘actionable’ event was identified (Supplementary Table 9). Whilst these numbers are in line with results from primary tumors^35^, longitudinal studies will be required to conclude if genomic analyses for therapeutic guidance should be repeated when a patient experiences progressive disease. Half of the patients with an actionable event (31% of total) contained a biomarker with a predicted sensitivity to a drug at level A (approved anti-cancer drugs) and lacked any known resistance biomarkers for the same drug (Fig. 5a). In 18% of patients the suggested therapy was a registered indication, while in 13% of cases it was outside the labeled indication. In a further 31% of patients a level B (experimental therapy) biomarker was identified. The actionable biomarkers spanned all variant classes including 1,815 SNVs, 48 MNVs, 190 indels, 745 CNAs, 69 fusion genes and 60 patients with microsatellite instability (Fig. 5b).

**Figure 5:**
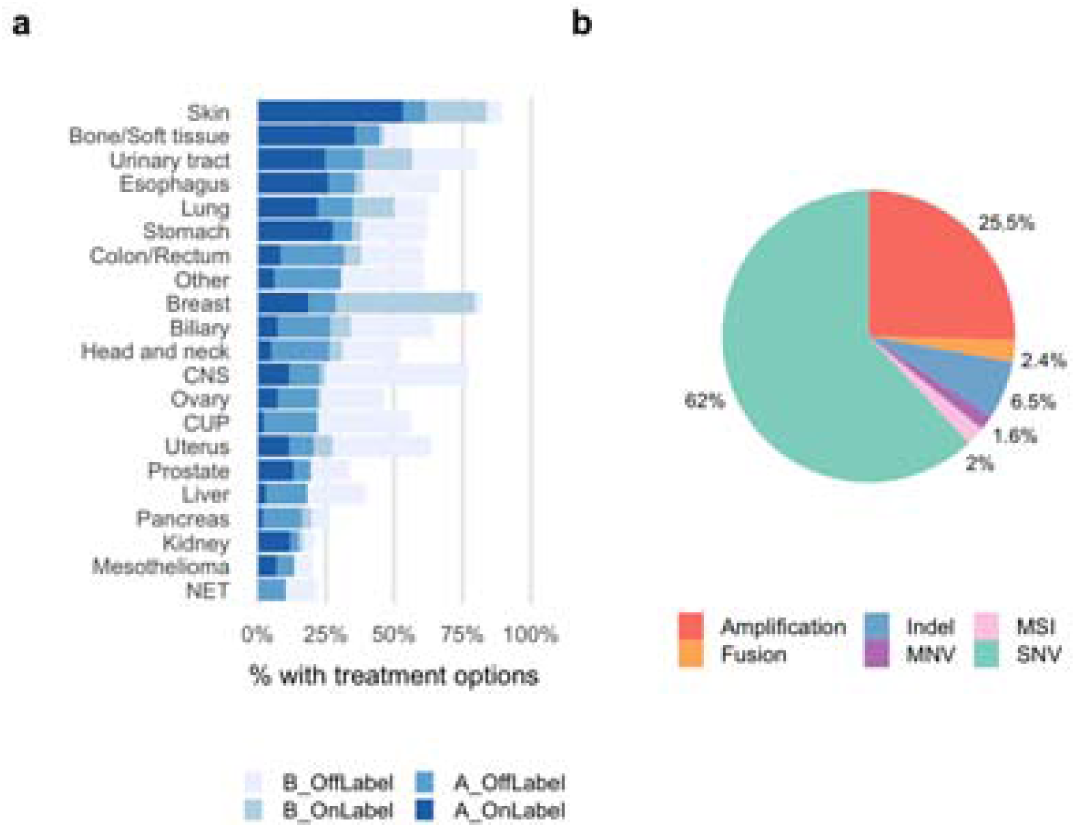
Actionability. a) Percentage of samples in each cancer type with an actionable mutation based on data in CGI, CIViC and OncoKB knowledgebases. Level ‘A’ represents presence of biomarkers with either an approved therapy or guidelines and level B represents biomarkers having strong biological evidence or clinical trials indicating they are actionable. On label indicates treatment registered by federal authorities for that tumor type, while off-label indicates a registration for other tumor types. b) Break down of the actionable variants by variant type.

Tumor mutation burden is an important emerging biomarker for response to immune checkpoint inhibitor therapy^55^ as it is a proxy for the amount of neo-antigens in the tumor cells. For NSCLC it has been shown in at least 2 subgroup analyses of large phase III trials that both PFS and OS are significantly improved with first line immunotherapy as compared to chemotherapy for patients whose tumors have a TMB >10 mutations per Mb^56,57^. Although various clinical studies based on this parameter are currently emerging, TMB was not yet included in the above actionability analysis. However, when applying the same cut-off to all samples in our cohort, an overall 18% of patients would qualify, varying from 0% for liver, mesothelioma and ovarian cancer patients to more than 50% for lung and skin cancer patients (Extended Data Fig. 6).

## Discussion

Genomic testing of tumors faces numerous challenges in meeting clinical needs, including the interpretation of variants of unknown significance (VUS), the steadily expanding universe of actionable genes, often with an increasingly small fraction of patients affected, and the development of advanced genome-derived biomarkers such as tumor mutational load, DNA repair status and mutational signatures. Our results demonstrate in several ways that WGS analyses of metastatic cancer can provide novel and relevant insights and be instrumental in addressing some of these key challenges in cancer precision medicine.

First, our systematic and large-scale pan-cancer analyses on metastatic cancer tissue allowed for the identification of several novel (cancer type-specific) cancer drivers and mutation hotspots. Second, the driver catalog analyses can be used to mitigate the problem of VUS interpretation^35^ both by leveraging previously identified pathogenic mutations (accounting for more than 2/3rds of oncogenic point-mutation drivers) and by careful analysis of the biallelic inactivation of putative TSG which accounts for over 80% of TSG drivers in metastatic cancer. Third, we demonstrate the importance of accounting for all types of variants, including large scale genomic rearrangements (via fusions and copy number alteration events), which account for more than half of all drivers, but also activating MNV and INDELs which we have shown are commonly found in many key oncogenes. Fourth, we have shown that using WGS, even with very strict variant calling criteria, we could find driver variants in more than 98% of all metastatic tumors, including putatively actionable events in a clinical and experimental setting for up to 62% of patients.

Although we did not find metastatic tumor genomes to be fundamentally different to primary tumors in terms of the mutational landscape or genes driving advanced tumorigenesis, we described characteristics that could contribute to therapy responsiveness or resistance in individual patients. In particular we showed that WGD is a more pervasive element of tumorigenesis than previously understood affecting over half of all metastatic cancers. We also found metastatic lesions to be less heterogeneous than reported for primary tumors, although the limited sequencing depth of ∼100x does not allow for drawing conclusions regarding low-frequency subclonal variants.

It should be noted that differences between WGS cohorts should be interpreted with some caution as inevitable differences between experimental and computational approaches may impact on observations and can only be addressed in longitudinal studies including the different stages of disease. Furthermore, the HMF cohort includes a mix of treatment-naive metastatic patients and patients who have undergone (extensive) prior systemic treatments. While this may impact on specific tumor characteristics, it also provides opportunities for studying treatment response and resistance as this data is recorded in the studies.

Finally, the resource described here is a valuable complementary resource to comparable whole genome sequencing-based resources of primary tumors in advancing fundamental and translational cancer research. All non-privacy sensitive data can be browsed through a local interface (database.hartwigmedicalfoundation.nl) developed by ICGC^58^ and all other data is made freely available for scientific research by a controlled access mechanism (see www.hartwigmedicalfoundation.nl/en for details).

## Online methods

A detailed description of methods and validations is available as Supplementary Information.

### Sample collection

Patients with advanced cancer not curable by local treatment options and being candidates for any type of systemic treatment and any line of treatment were included as part of the CPCT-02 (NCT01855477) and DRUP (NCT02925234) clinical studies, which were approved by the medical ethical committees (METC) of the University Medical Center Utrecht and the Netherlands Cancer Institute, respectively. A total of 41 academic, teaching and general hospitals across the Netherlands participated in these studies and collected material and clinical data by standardized protocols^59^. Patients have given explicit consent for whole genome sequencing and data sharing for cancer research purposes. Core needle biopsies were sampled from the metastatic lesion, or when considered not feasible or not safe, from the primary tumor site and frozen in liquid nitrogen. A single 6 micron section was collected for hematoxylin-eosin (HE) staining and estimation of tumor cellularity by an experienced pathologist and 25 sections of 20 micron were collected in a tube for DNA isolation. In parallel, a tube of blood was collected. Left-over material (biopsy, DNA) was stored in biobanks associated with the studies at the University Medical Center Utrecht and the Netherlands Cancer Institute.

### Whole genome sequencing and variant calling

DNA was isolated from biopsies (>30% tumor cellularity) and blood according to supplier’s protocols (Qiagen) using the DSP DNA Midi kit for blood and QIAsymphony DSP DNA Mini kit for tissue. A total of 50-200 ng of DNA (sheared to average fragment length of 450nt) was used as input for TruSeq Nano LT library preparation (Illumina). Barcoded libraries were sequenced as pools on HiSeqX generating 2 × 150 read pairs using standard settings (Illumina). BCL output was converted using bcl2fastq tool (Illumina, versions 2.17 to 2.20) using default parameters. Reads were mapped to the reference genome GRCH37 using BWA-mem v0.7.5a^60^, duplicates were marked for filtering and INDELs were realigned using GATK v3.4.46 IndelRealigner^61^. GATK HaplotypeCaller v3.4.46^62^ was run to call germline variants in the reference sample. For somatic SNV and INDEL variant calling, GATK BQSR^63^ was applied to recalibrate base qualities. SNV & INDEL somatic variants were called using Strelka v1.0.14^64^ with optimized settings and post-calling filtering. Structural Variants were called using Manta(v1.0.3)^65^ with default parameters followed by additional filtering to improve precision using an internally built tool (Breakpoint-Inspector v1.5). To assess the impact of sequencing depth on variant calling sensitivity, we downsampled the BAMS of 10 samples at random by 50% and sequenced two samples twice to double the normal sequencing depth and reran the identical somatic variant calling pipeline.

### Purity, ploidy and copy number calling

Copy number calling and sample purity determination was performed using PURPLE (PURity & PLoidy Estimator), which combines B-allele frequency (BAF), read depth and structural variants to estimate the purity of a tumor sample and determine the copy number and minor allele ploidy for every base in the genome. The purity and ploidy estimates and copy number profile obtained from PURPLE were validated on in silico simulated tumor purities, by DNA fluorescence in situ hybridisation (FISH) and by comparison with an alternative tool (ASCAT^66^). ASCAT was run on GC corrected data using default parameters except for gamma which was set to 1 which is recommended for massively parallel sequencing data. We implement a simple heuristic that determines if Whole Genome Duplication has occurred: Major allele Ploidy >1.5 on at least 50% of at least 11 autosomes as the number of duplicated autosomes per sample (ie. the number of autosomes which satisfy the above rule) follows a bimodal distribution with 95% of samples have either <= 6 or > =15 autosomes duplicated.

### Sample selection for downstream analyses

Following copy number calling, samples were filtered out based on absence of somatic variants, purity <20%, and GC biases, yielding a high-quality data set of 2,520 samples. Where multiple biopsies exist for a single patient, the highest purity sample was used for downstream analyses (resulting in 2,399 samples).

### Mutational Signature analysis

Mutational signatures were determined by fitting SNV counts per 96 tri-nucleotide context to the 30 COSMIC signatures^28^ using the mutationalPatterns package^67^. Residuals were calculated as the sum of the absolute difference between observed and fitted across the 96 buckets. Signatures with <5% overall contribution to a sample or absolute fitted mutational load <300 variants were excluded from the summary plot (Extended Data Fig. 4a).

### Germline predisposition variant calling

We searched for pathogenic germline variants (SNVs, INDELs and CNAs) in a broad list of 152 germline predisposition genes curated by Huang et al^68^, using GATK HaplotypeCaller^62^ output from each sample. For each variant identified we assessed the genotype in the germline (HET or HOM), whether there was a 2nd somatic hit in the tumor, and whether the wild type or the variant itself was lost by a CNA. We observed that for the variants in many of the 152 predisposition genes that a loss of wild type in the tumor via LOH was lower than the average rate of LOH across the cohort and that fewer than 5% of observed variants had a 2nd somatic hit in the same gene. Moreover, in many of these genes the ALT variant was lost via LOH as frequently as the wild type, suggesting that a significant portion of the 566 variants may be passengers. For our downstream analysis and driver catalog, we therefore restricted our analysis to a more conservative ‘High Confidence’ list including only the 25 cancer related genes in the ACMG secondary findings reporting guidelines (v2.0)^69^, together with 4 curated genes (CDKN2A, CHEK2, BAP1 & ATM), selected because these are the only additional genes from the larger list of 152 genes with a significantly elevated proportion of called germline variants with loss of wild type in the tumor sample.

### Clonality and biallelic status of point mutations

The ploidy of each variant is calculated by adjusting the observed VAF by the purity and then multiplying by the local copy number to work out the absolute number of chromatids that contain the variant. We mark a mutation as biallelic (i.e. no wild type remaining) if Variant Ploidy > Local Copy Number – 0.5. For each variant we also determine a probability that it is subclonal. This is achieved via a two-step process involving fitting the somatic ploidies for each sample into a set of clonal and subclonal peaks and calculating the probability that each individual variant belongs to each peak. Subclonal counts are calculated as the total density of the subclonal peaks for each sample. Subclonal driver counts are calculated as the sum across the driver catalog of subclonal probability * driver likelihood.

### MSI status determination

To determine the MSI status we used the method described by the MSISeq tool^70^ and counted the number of INDELS per million bases occuring in homopolymers of 5 or more bases or dinucleotide, trinucleotide and tetranucleotide sequences of repeat count 4 or more. MSIseq score of >4 were considered microsatellite instable (MSI).

### Significantly mutated driver genes

We used Ensembl^71^ v89.37 as a basis for gene definitions and have taken the union of Entrez identifiable genes and protein coding genes as our base panel (25,963 genes of which 20,083 genes are protein coding). Pan cancer and at an individual cancer level we tested the normalised dNdS rates using dNdScv^26^ against a null hypothesis that dNdS=1 for each variant subtype. To identify SMGs in our cohort we used a strict significance cutoff of q<0.01.

To search for significantly amplified and deleted genes we first calculated the minimum exonic copy number per gene. For amplifications, we searched for all the genes with high level amplifications only (defined as minimum Exonic Copy number > 3 * sample ploidy). For deletions, we searched for all the genes in each sample with either full or partial gene homozygous deletions (defined as minimum exonic copy number < 0.5) excluding the Y chromosome. We then searched separately for amplifications and deletions, on a per chromosome basis, for the most significant focal peaks, using an iterative GISTIC-like peel off method^72^. Most of the deletion peaks resolve clearly to a single target gene reflecting the fact that homozygous deletions are highly focal, but for amplifications this is not the case and the majority of our peaks have 10 or more candidates. We therefore annotated the peaks, to choose a single putative target gene using an objective set of automated curation rules. Finally, filtering was applied to yield highly significant deletions and amplifications.

Homozygous deletions were also annotated as common fragile site (CFS) based on their genomic characteristics, including a strong enrichment in long genes (>500,000 bases) and a high rate (>30%) of deletions between 20 kb and 1 mb^30^.

### Somatic driver catalog construction

We created a catalog of driver variants in our cohort across all variant types on a per patient basis. This was done in a similar incremental manner to Sabarinathan et al^35^ (N. Lopez, personal communication) whereby we first calculated the number of drivers in a broad panel of known and significantly mutated genes across the full cohort, and then assigned the drivers for each gene to individual patients by ranking and prioritising each of the observed variants. Key points of difference in this study were both the prioritisation mechanism used and our choice to ascribe each mutation a probability of being a driver rather than a binary cutoff based on absolute ranking.

The four steps to create the catalog are: 1) Create a panel of driver genes for point mutations using significantly mutated genes and known drivers using the union of Martincorena significantly mutated genes^26^ (filtered to significance of q<0.01), HMF significantly mutated genes (q<0.01) at global level or at cancer type level and Cosmic Curated Genes^28^ (v83). 2) Determine TSG or Oncogene status of each significantly mutated gene using a logistic regression classification model (trained using Cosmic annotation). 3) Add mutations from all variant classes to the catalog when meeting any of the following criteria i) all missense and in-frame indels for panel oncogenes, ii) all non synonymous and essential splice point mutations for tumor suppressor genes, iii) all high level amplifications for significantly amplified target genes and panel oncogenes, iv) all homozygous deletions for significantly deleted target genes and panel TSG, v) all known or promiscuous inframe gene fusions, and vi) recurrent TERT promoter mutations. 4) Calculate a per sample driver likelihood (between 0 and 1) for each mutation in the catalog to ensure that only likely pathogenic and excess mutations (based on dNdS) are used for determining the number of drivers. All driver mutation counts reported at a per cancer type or sample level refer to the sum of driver likelihoods for that cancer type or sample. To examine the co-occurence of drivers, the driver-gene catalog was filtered to exclude fusions and any driver with a driver likelihood of < 0.5. Separately for each cancer type, every pair of driver genes was tested to see whether they co-occur more or less frequently than expected if they were independent using Fisher’s Exact Test. The results were adjusted to a FDR using the number of gene-pair comparison being tested in each cancer type cohort. Gene pairs with a positive correlation which were on the same chromosome were excluded from the analysis as they are frequently co-amplified or deleted by chance.

### Actionability analysis

To determine clinical actionability of the variants observed in each sample, we compared all variants with three external clinical annotation databases (OncoKB^43^, CGI^44^ and CIViC^42^) that were mapped to a common data model as defined by https://civicdb.org/help/evidence/evidence-levels. Here, we considered only A and B level variants. This classification of actionable events roughly corresponds to the recently proposed ESMO Scale for Clinical Actionability of molecular Targets (ESCAT)^73^ as follows: ESCAT I-A+B (for A on-label) and I-C (for A off-label) and ESCAT II-A+B (for B on-label) and III-A (for B off-label). For each actionable mutation, it was also determined to be either on-label (ie. evidence supports treatment in that specific cancer type) or off-label (evidence exists in another cancer type). To do this, we annotated both the patient cancer types and the database cancer types with relevant DOIDs, using the disease ontology database^74^. For each actionable mutation in each sample, we aggregated all the mapped evidence that was available supporting both on-label and off-label treatments at A or B evidence level. Treatments that also had evidence supporting resistance based on other biomarkers in the sample at the same or higher evidence level were excluded as non-actionable. Samples classified as MSI in our driver catalog were also mapped as actionable at level A evidence based on clinical annotation in the OncoKb database. For each sample we reported the highest level of actionability, ranked first by evidence level and then by on-label vs off-label.

## Supporting information

Detailed methods

Supplementary Image 1

Supplementary Image 2

Supplementary Image 3

## Data availability

All data described in this study is freely available from the Hartwig Medical Foundation for academic research within the constraints of the consent given by the patients. Standardized procedures and request forms can be found at https://www.hartwigmedicalfoundation.nl/en. All bioinformatic analysis tools and scripts used are available at https://github.com/hartwigmedical/.

## Supplementary Information

Supplementary Information is linked to the online version of the paper at www.nature.com/nature.

## Acknowledgements

We thank the Hartwig Foundation and Barcode for Life for financial support of clinical studies and WGS analyses. Implementation of the data portal was supported by a grant from KWF Kankerbestrijding (HMF2017-8225, GENONCO). We are particularly grateful to all patients, nurses and medical specialists for their essential contributions making this study possible and Hans van Snellenberg (Hartwig Medical Foundation) for operational management. We would like to thank Stefan Willems, Wendy de Leng, Alexander Hoischen and Winand Dinjens for support with pathology assessments and mutation validations and Jeroen de Ridder, Wigard Kloosterman and Harmen van de Werken for critically reading the manuscript.

## Author contributions

EFS, SF, EEV and EC designed the study. HJK, VCGT, CMLvH, ML, POW, MPL, NS, EFS, SS, EEV contributed patient material, MPL and NS supervised clinical studies and EdB supervised WGS data generation. PP, JB, KD, SH, AvH, WO, PR, CS, and MV structured and analyzed data. PP, JB and EC wrote the manuscript. All authors provided input for improvement of the manuscript.

## Author information

Reprints and permissions information is available at www.nature.com/reprints. The authors declare the following competing interests: EEV is board member of the Hartwig Medical Foundation.

## Data availability

All data described in this study is available through Hartwig Medical Foundation (www.hartwigmedicalfoundation.nl/en/applying-for-data/).

## Code availability

All bioinformatic analysis tools and scripts used are available at https://github.com/hartwigmedical/.

**Supplementary Image File 1: Copy Number profile per cancer types**

Circos plots showing the proportion of samples with amplification and deletion events by genomic position per cancer type. The inner ring shows the % of tumors with homozygous deletion (red), LOH and significant loss (copy number < 0.6x sample ploidy – dark blue) and near copy neutral LOH (light blue). The outer ring shows the % of tumors with high level amplification (>3x sample ploidy – orange), moderate amplification (>2x sample ploidy – dark green) and low level amplification (>1.4x amplification – light green). Scales on both rings are 0-100% and inverted for the inner ring. The most frequently observed high level gene amplifications (black text) and homozygous deletions (red text) are labelled.

**Supplementary Image File 2: Coding mutation profiles by tumor suppressor driver gene**

Location and driver classification of all coding mutations (SNVs and indels) in tumor suppressor genes (TSG) in the driver catalog. The lollipops on the chart show the location (coding sequence coordinates) and count of mutations for all candidate drivers. The height of lollipop represents the total count of each individual variant in the cohort (log scale). The height of the solid line represents the sum of driver likelihoods for that variant, ie. the proportion that are expected to be drivers. (Partially) dotted lines hence indicate variants for which driver role is uncertain. Variants are unshaded if all instances of that variant are monoallelic single hits with no LOH. The right column chart shows the estimated number of drivers (calculated as the sum of driver likelihoods) and passenger variants in each gene by cancer type.

**Supplementary Image File 3: Coding mutation profiles by oncogene driver gene**

Location and driver classification of all coding mutations (SNVs and indels) in oncogenes (a) and tumor suppressor genes (TSG) (b) in the driver catalog. The lollipops on the chart show the location (coding sequence coordinates) and count of mutations for all candidate drivers. The height of lollipop represents the total count of each individual variant in the cohort (log scale). The height of the solid line represents the sum of driver likelihoods for that variant, ie. the proportion that are expected to be drivers. (Partially) dotted lines hence indicate variants for which driver role is uncertain. The right column chart shows the estimated number of drivers (calculated as the sum of driver likelihoods) and passenger variants in each gene by cancer type.

**Supplementary Information:** Detailed description of methods, parameters and validation results

**Supplementary Table 1:** Overview of contributing organizations and local principal investigators

**Supplementary Table 2:** Overview of cohort and sample characteristics

**Supplementary Table 3:** Pan-cancer and cancer type-specific dNdScv results

**Supplementary Table 4:** Recurring amplifications (a) and deletions (b) and associated target genes

**Supplementary Table 5:** Somatic driver catalog

**Supplementary Table 6:** Germline driver catalog

**Supplementary Table 7:** Gene Fusions

**Supplementary Table 8:** Overview of patients with multiple biopsies

**Supplementary Table 9:** Actionable mutations

---

**Extended Data Figure 1:**
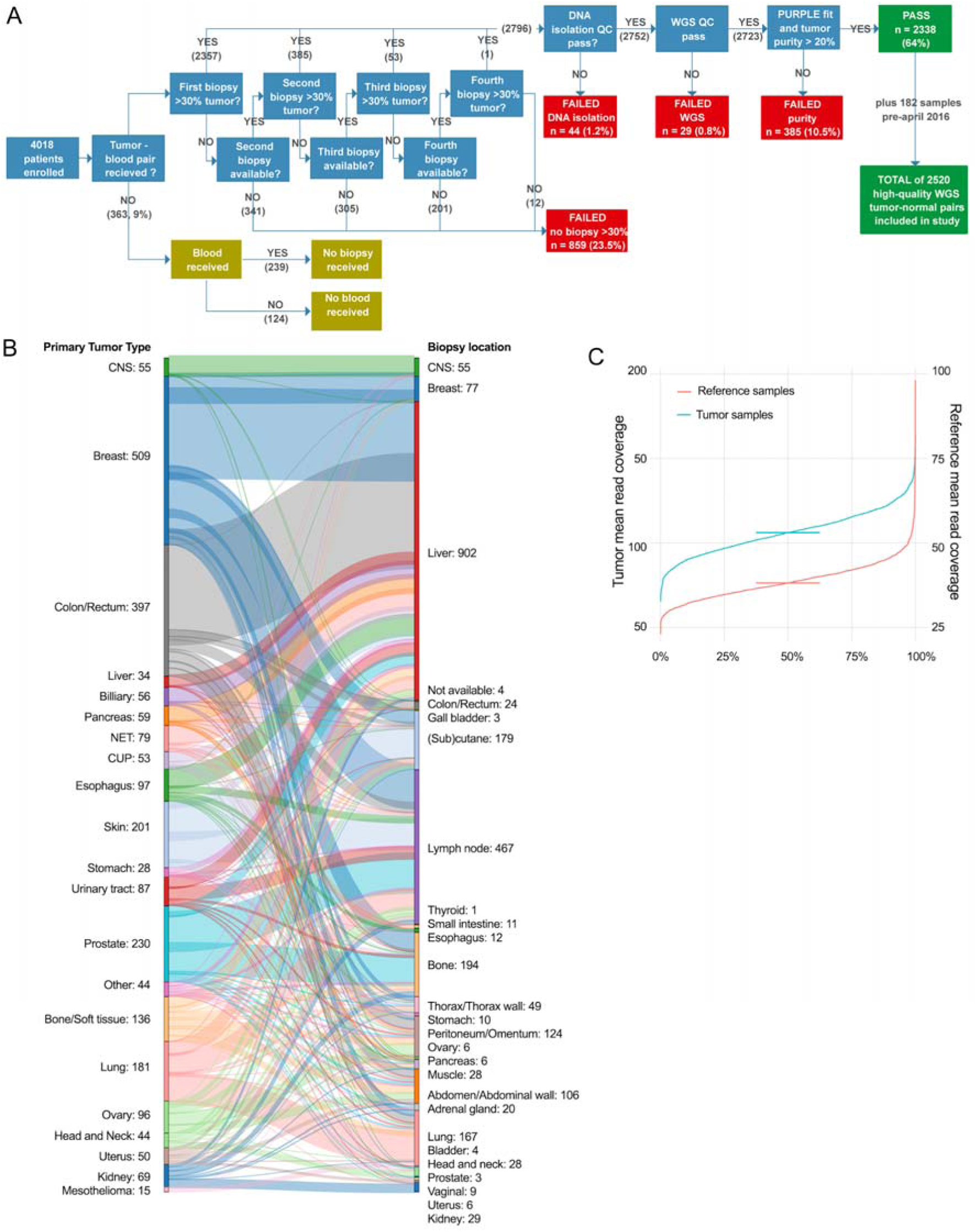
Hartwig sample workflow, biopsy locations and sequence coverage. a) Sample workflow from patient to high-quality WGS data. A total of 4,018 patients were enrolled in the study between April 2016 and April 2018. For 9% of patients no blood and/or biopsy material was obtained, mostly because conditions of patients prohibited further study participation. Up to 4 fresh-frozen biopsies per patient were received, which were sequentially analyzed to identify a biopsy with more than 30% tumor cellularity as determined by routine histology assessment. For 859 patients no suitable biopsy was obtained and 2,796 patients were further processed for WGS. 44 and 29 samples failed in either DNA isolation or library preparation and raw WGS data quality QC, respectively. For an additional 385 samples the WGS data was of good quality, but the tumor purity determination based on WGS data (PURPLE) was less than 20% making reliable and comprehensive somatic variant calling and were therefore excluded. Eventually, 2,338 tumor-normal sample pairs with high-quality WGS data were obtained, which were supplemented with 182 pairs from pre-April 2016, adding up to 2,520 tumor normal pairs that were included in this study. b) Breakdown of cohort by biopsy location. Tumor biopsies were taken from a broad range of locations. Primary tumor type is shown on the left and the biopsy location on the right. c) Distribution of sample sequencing depth for tumor and blood reference.

**Extended Data Figure 2:**
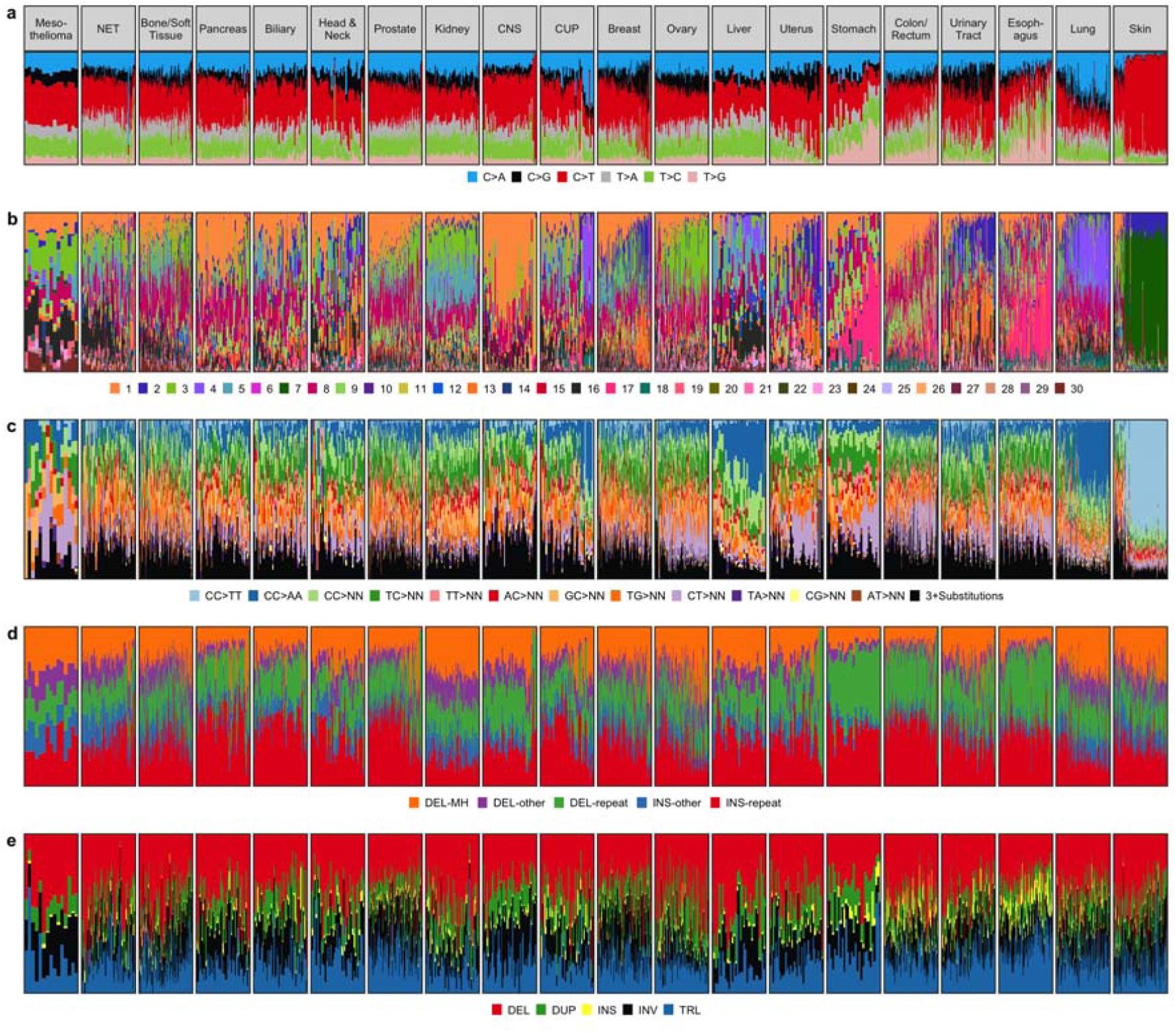
Mutational context distribution per tumor type. Variant subtype, mutational context or signature per individual sample for each of Single Nucleotide Variant (SNV) (a), SNV by COSMIC Signature (b), Multi Nucleotide Variant (MNV) (c), INsertion/DELetion (INDEL) (d), Structural Variant (SV) (e). Each column chart is ranked within tumor type by mutational load from low to high in that variant class. MNVs are classified by the dinucleotide substitution with NN referring to any dinucleotide combination. SVs are classified by type: INV = inversion, DEL = deletion, DUP = tandem duplication, TRL = translocation, INS = insertion. Highly characteristic known patterns can be discerned, for example the high rates of C>T SNVs, CC>TT MNVs and Cosmic S18 for skin tumors, high rates of C>A SNVs and Cosmic S4 for lung tumors.

**Extended Data Figure 3:**
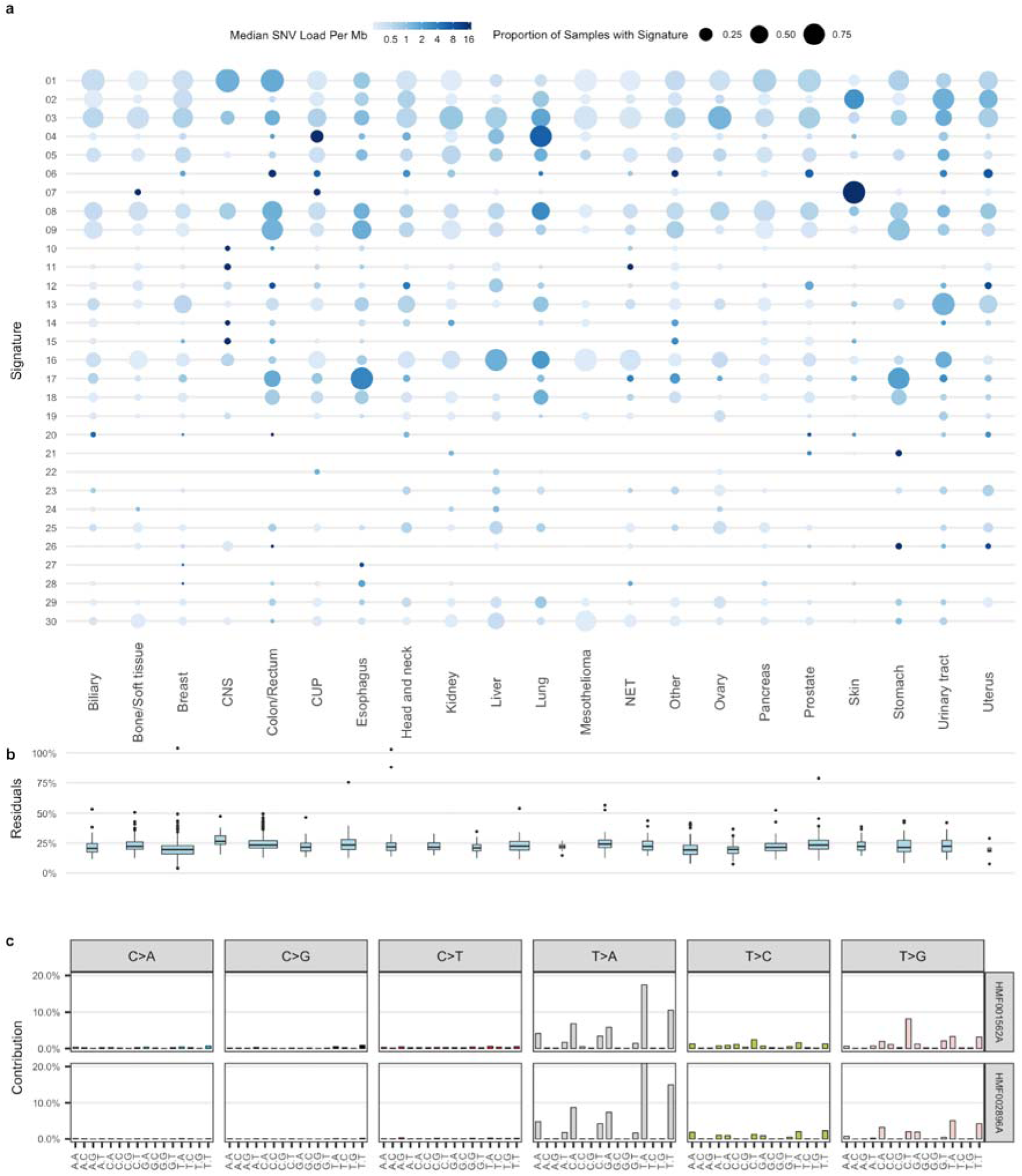
SNV Mutational Signatures. a) Prevalence and median mutational load of fitted cosmic SNV mutational signature per cancer type. The observed distribution largely reflects the patterns as observed from primary cancers^13^. b) Box-and-whisker plot of relative residuals in fits per cancer type (sum of absolute difference between fitted and actual divided by total mutational load) c) Proportion of variants by 96 trinucleotide mutational context for 2 selected samples with high residuals and high mutational load. Upper panel and lower panel represent the highest outlier for Breast (HMF002896) and Esophagus (HMF001562), respectively from panel b. Both of these samples were previously treated with an experimental drug, SYD985 a Duocarmycin-Based HER2-Targeting Antibody-Drug conjugate^75^.

**Extended Data Figure 4:**
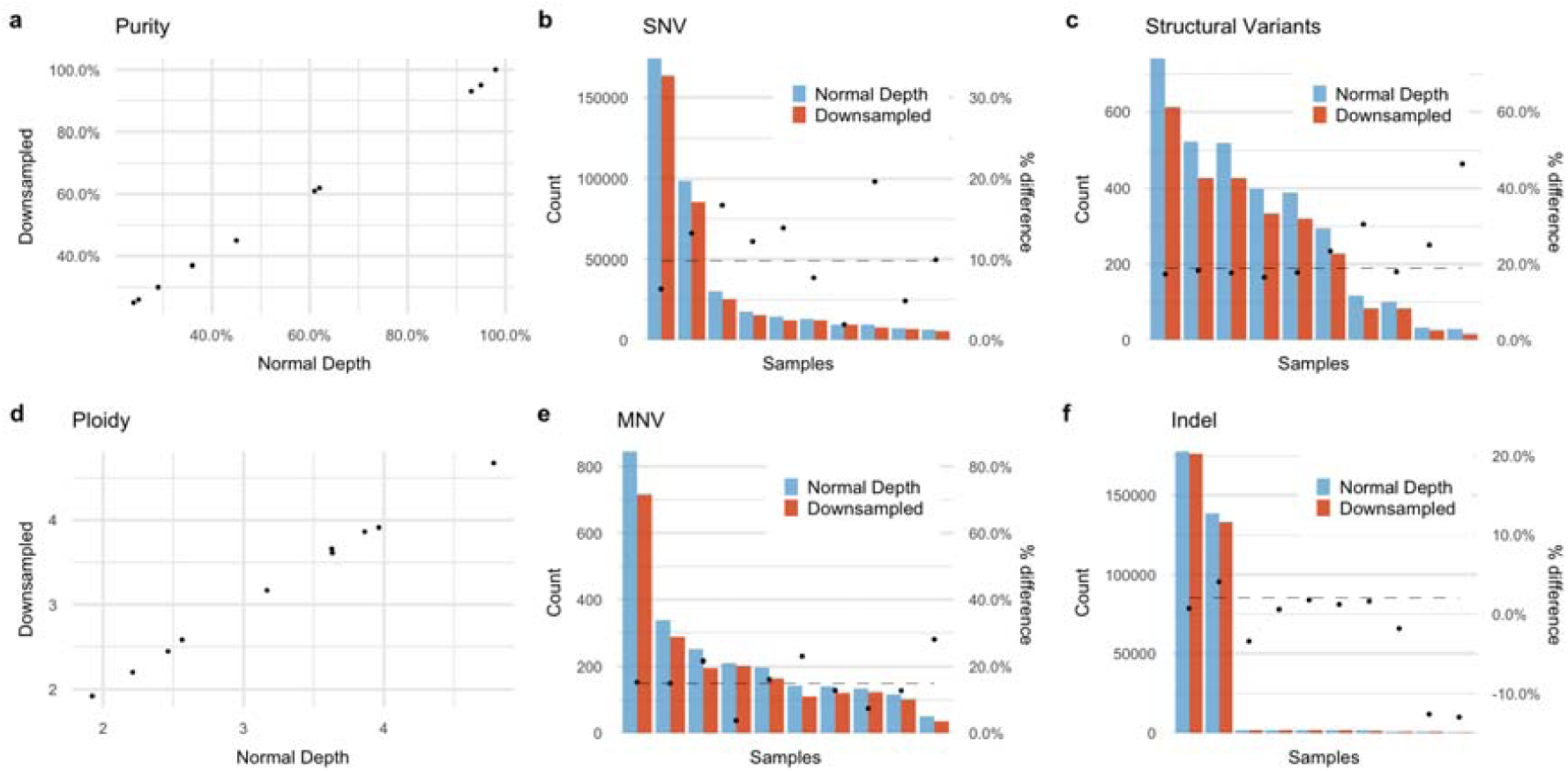
Impact of sequencing depth on variant calling. Comparison of variant calling of 10 randomly selected samples at normal depth and 50% downsampled (∼50x, similar as the mean coverage for the PCAWG study^14^) for purity (a), SNV counts (b), SV counts (c), Ploidy (d), MNV counts (e) and INDEL counts (f). Decreasing coverage results to an average decrease in sensitivity of 10% for SNV, 2% for INDEL, 15% for MNV and 19% for SVs.

**Extended Data Figure 5:**
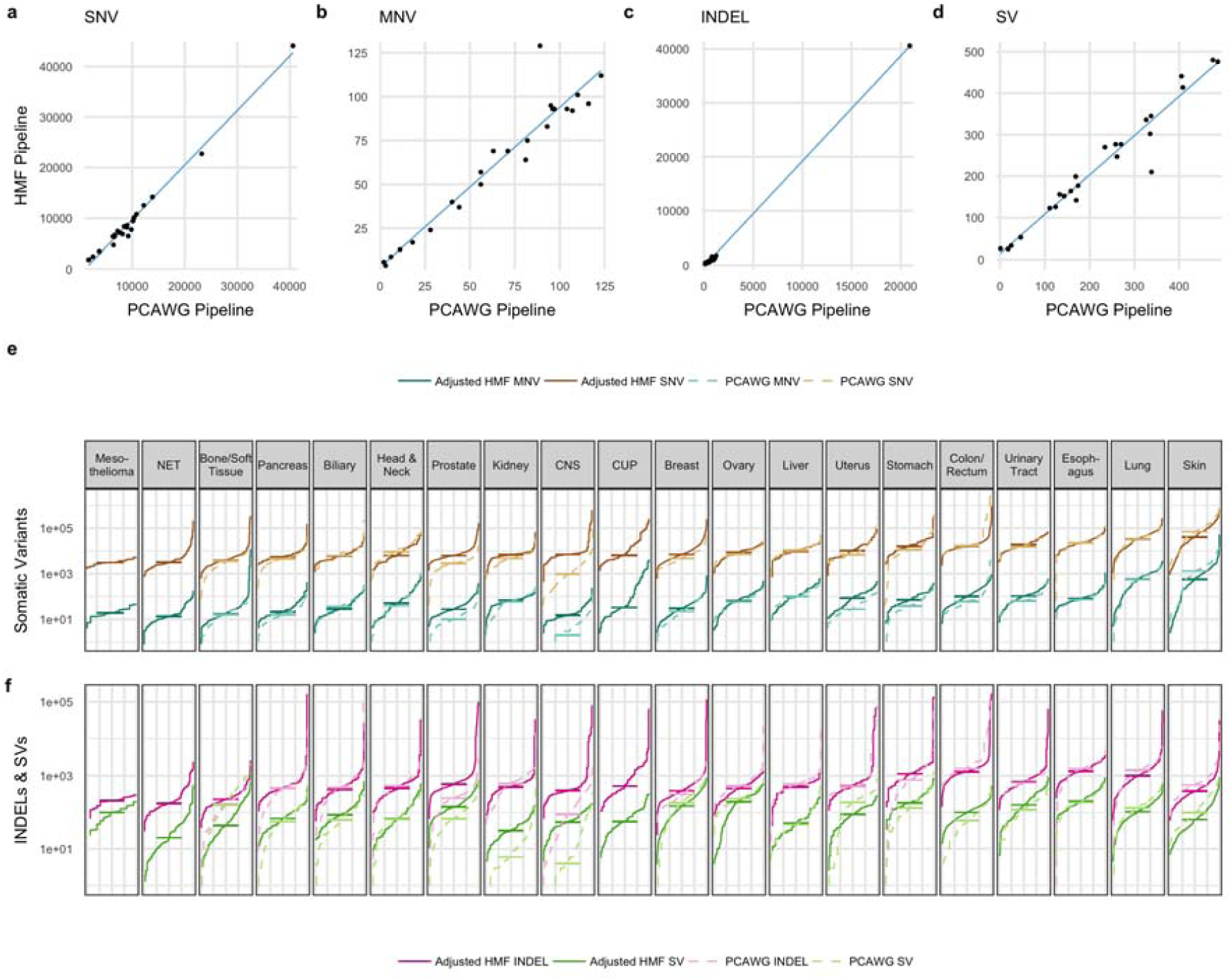
Impact of bioinformatic analysis pipeline on variant calling. Comparison of observed mutational count per sample for SNV (a), MNV (b), indel (c), and SV (d) on 24 patient samples analyzed by PCAWG pipeline and HMF pipeline. The PCAWG pipeline was found to have a 43% lower sensitivity for INDELs (which is based on a consensus calling), 18% lower for SVs (based on a different algorithm) and 6% lower for MNVs (only includes MNVs involving 2 nt), with nearly the same sensitivity for SNVs. Cumulative distribution function plot for each tumor type of coverage and pipeline-adjusted mutational load for SNV and MNV (e) and INDEL and SV (f). Mutational loads as shown in Figure 1 of the main manuscript were adjusted for the sensitivity effects caused by differences in sequencing depth coverage (Extended Data Fig. 4) and analysis pipeline differences (this figure, panels a-d). After this correction, the TMB between primary and metastatic cohorts across all variant types are much more comparable (e, f), indicating that technical differences do contribute to the reported mutational load differences between primary and metastatic tumors. Prostate cancer is the most notable exception with approximately 2x the TMB in all variant classes, although more subtle differences, potentially driven by biology, can be observed for other tumor and mutation types as well. For cancer types which are comparable to the PCAWG cohort, the equivalent PCAWG numbers are shown by dotted lines. The median for each cohort is displayed by a horizontal line.

**Extended Data Figure 6:**
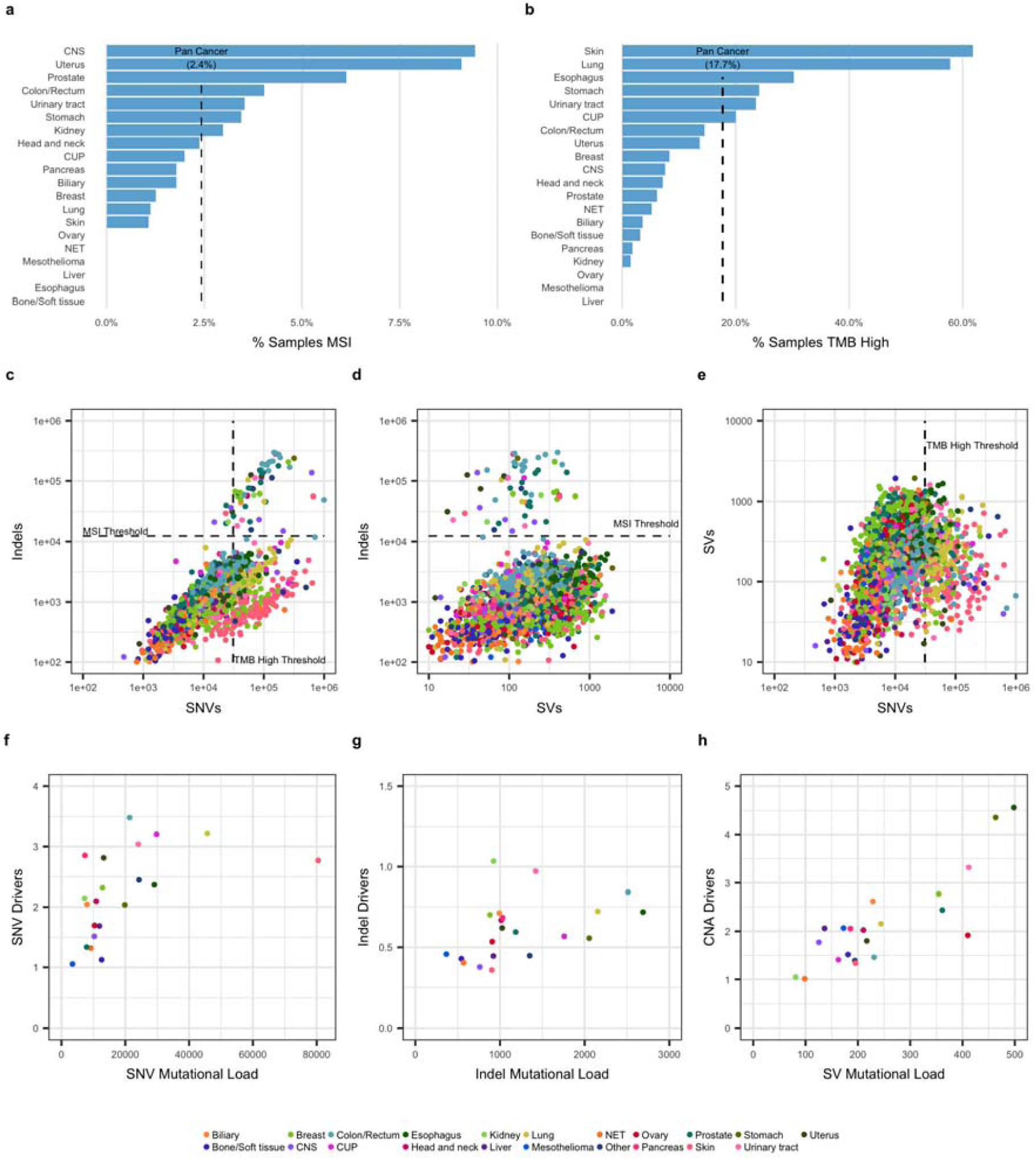
Mutational load, genome wide analyses and drivers. a) Proportion of samples by cancer type classified as microsatellite instable (MSISeq score > 4) b) Proportion of samples with a high mutational burden (TMB > 10 SNV / Mb) c)-e) Scatter plot of mutational load per sample for INDEL vs SNV (c), INDEL vs SV (d), and SV vs SNV (e). MSI (MSISeq score > 4) and ‘high TMB’ (>10 SNV/ MB) thresholds are indicated. f)-h) Mean mutational load vs driver rate for SNV (f), INDEL (g) and SV (h) grouped by cancer type. MSI samples were excluded.

**Extended Data Figure 7:**
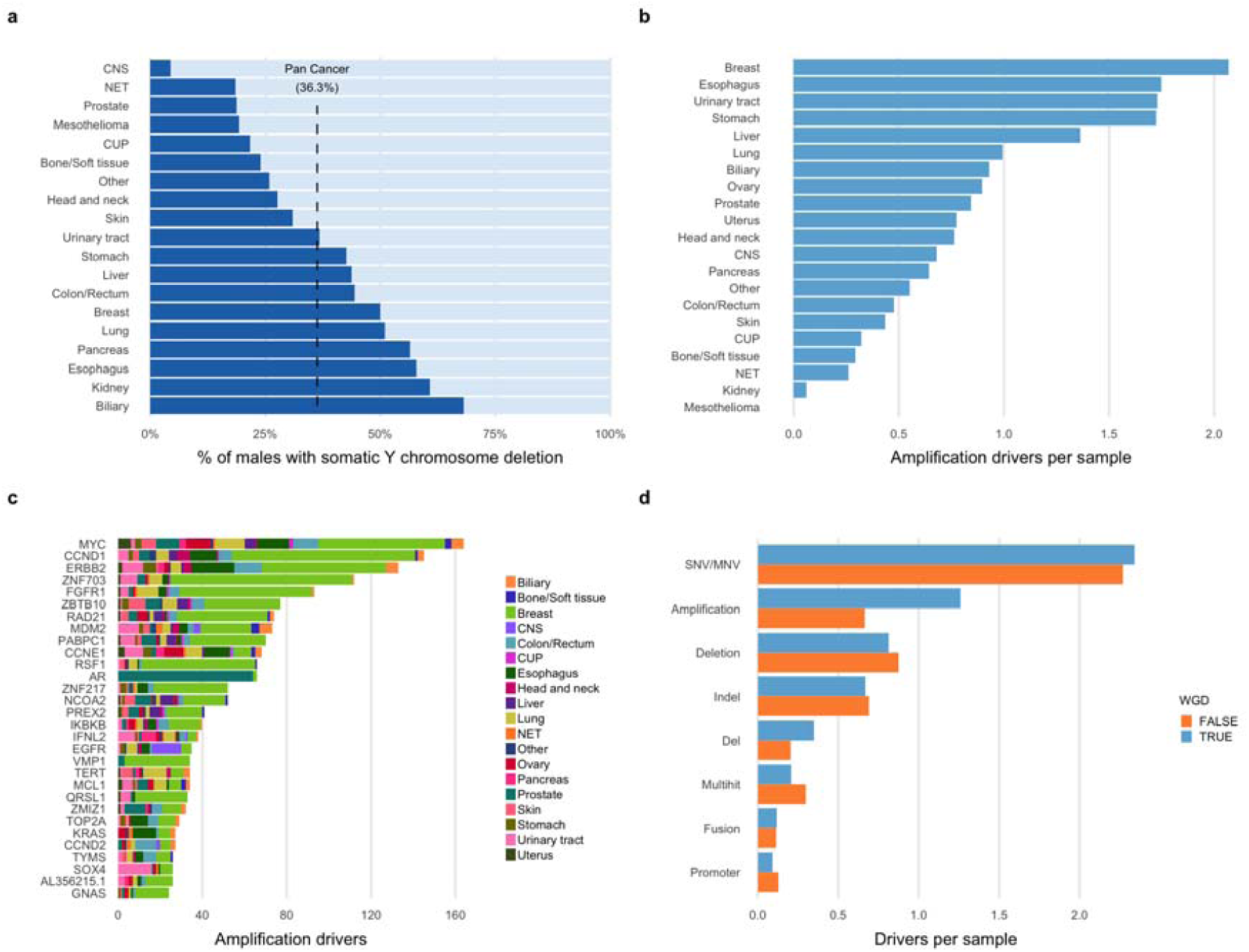
Somatic Y chromosome loss and driver amplifications. a) Proportion of Male tumors with somatic loss of >50% of Y chromosome (dark blue) grouped by tumor type. b) Mean rate of amplification drivers per cancer type. c) Breakdown of the number of amplification drivers per gene by cancer type. d) Mean rate of drivers per variant type for samples with and without WGD.

**Extended Data Figure 8:**
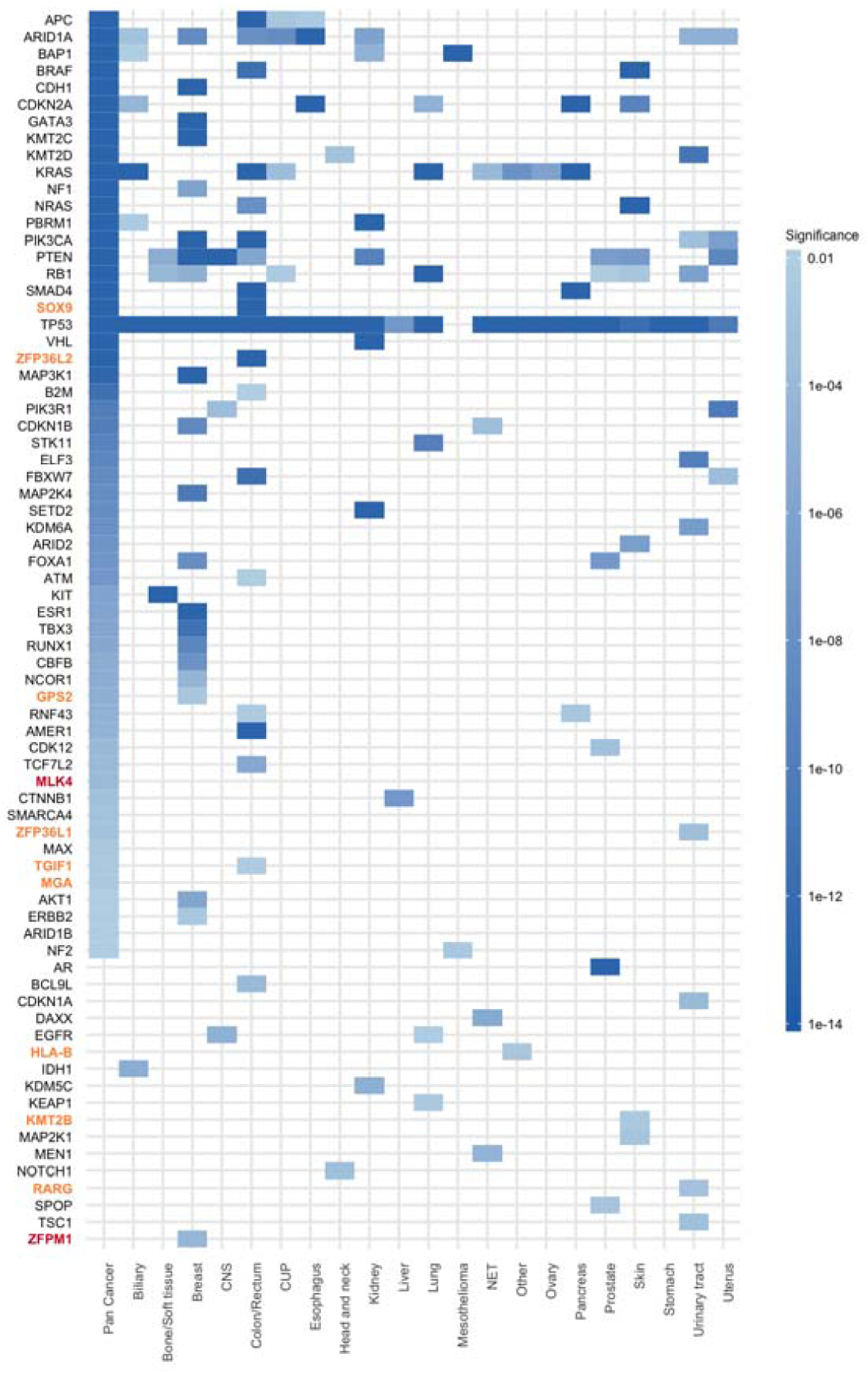
Significantly mutated genes. Tile chart showing genes found to be significantly mutated per cancer type cohort and pan-cancer using dNdScv. Gene names marked in orange are also significant in Martincorena et al^26^, but not found in COSMIC curated or census. Gene names marked in red are novel in this study.

**Extended Data Figure 9:**
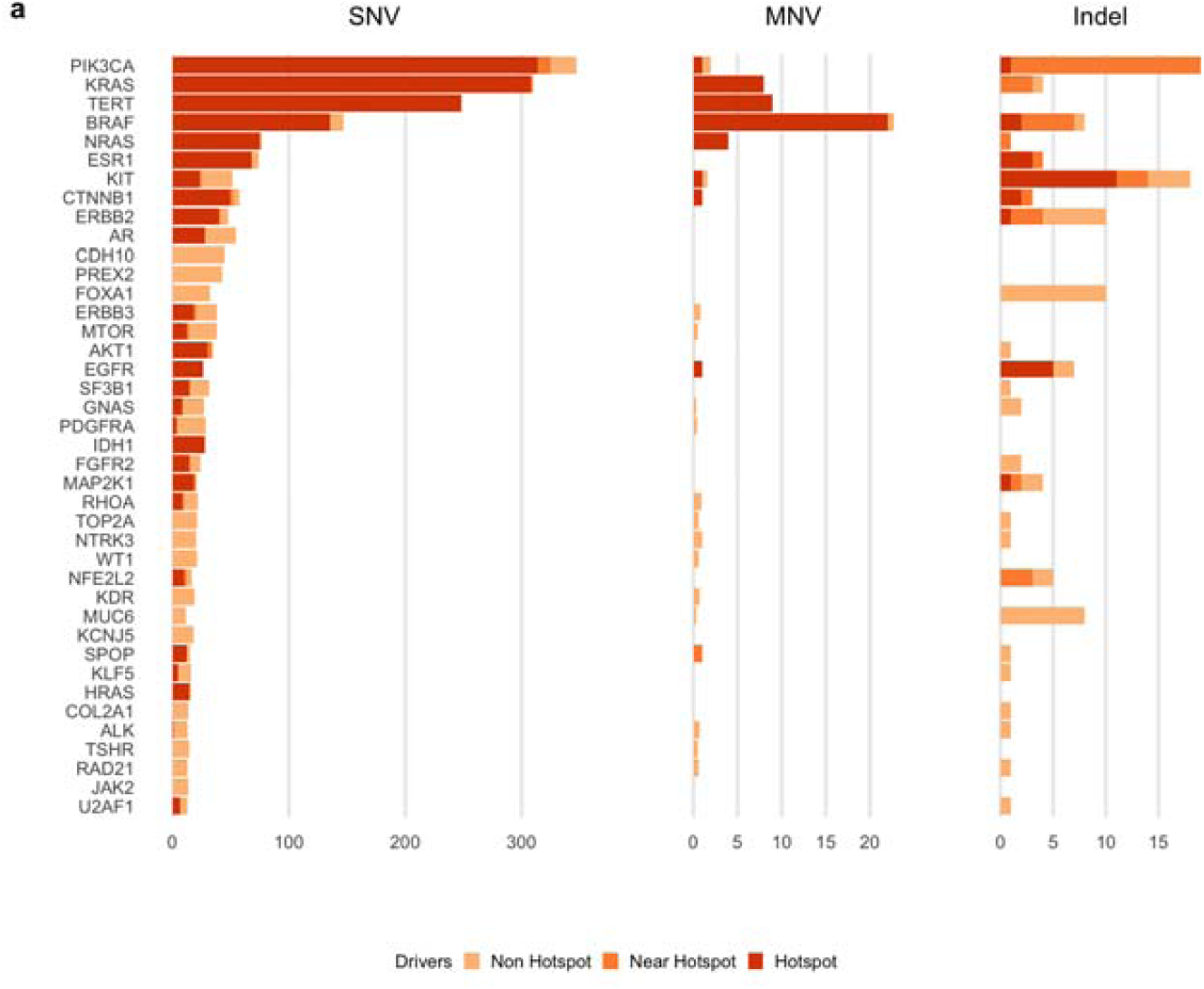
Oncogenic Hotspots. Count of driver point mutations by variant type. Known pathogenic mutations curated from external databases are categorized as hotspot mutations. Mutations within 5 bases of a known pathogenic mutation are shown as near hotspot and all other mutations are shown as non-hotspot.

**Extended Data Figure 10:**
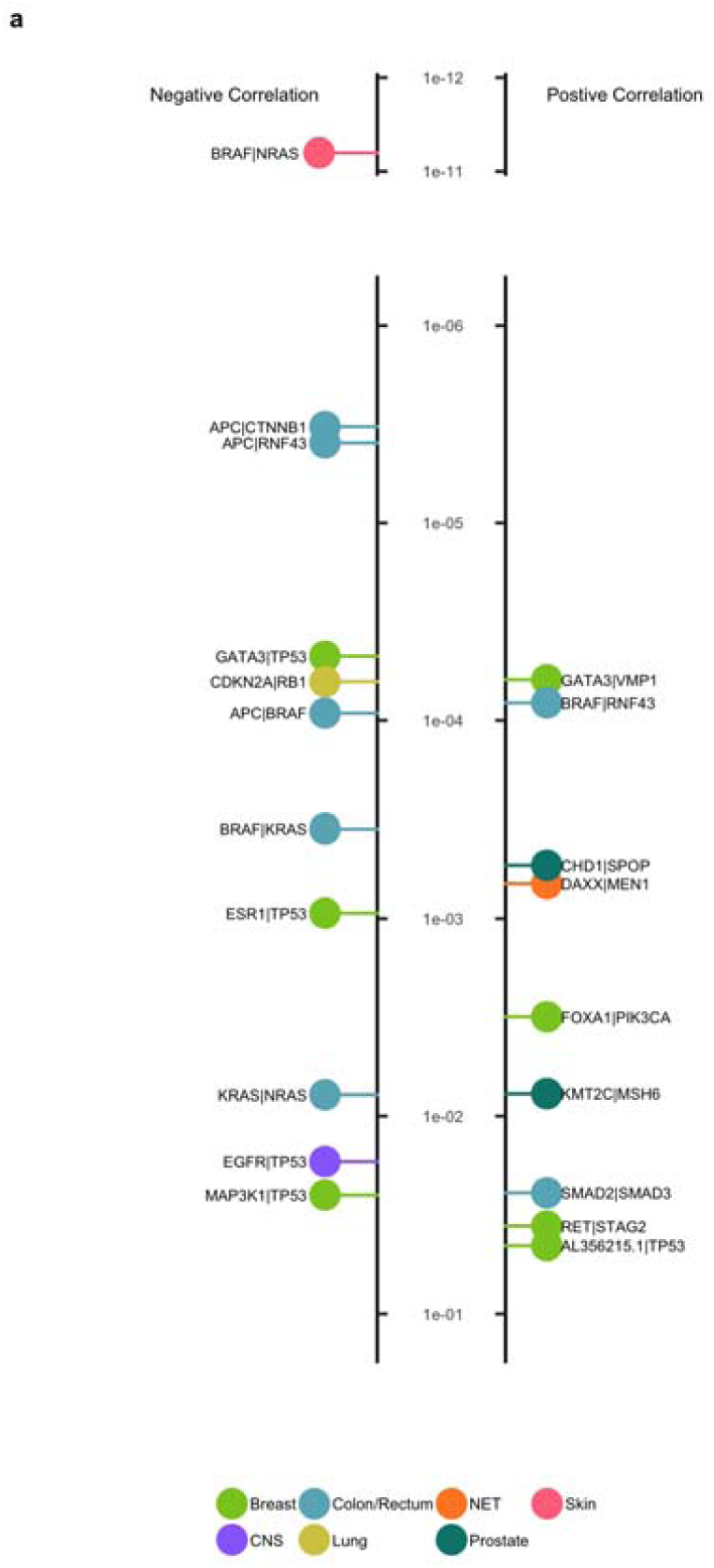
Driver co-occurrence. a) Mutated driver gene pairs which are significantly positively (on the right) or negatively (on the left) correlated in individual tumor types sorted by q-value. The color indicates the tumor type as depicted below the chart.

**Extended Data Fig 11:**
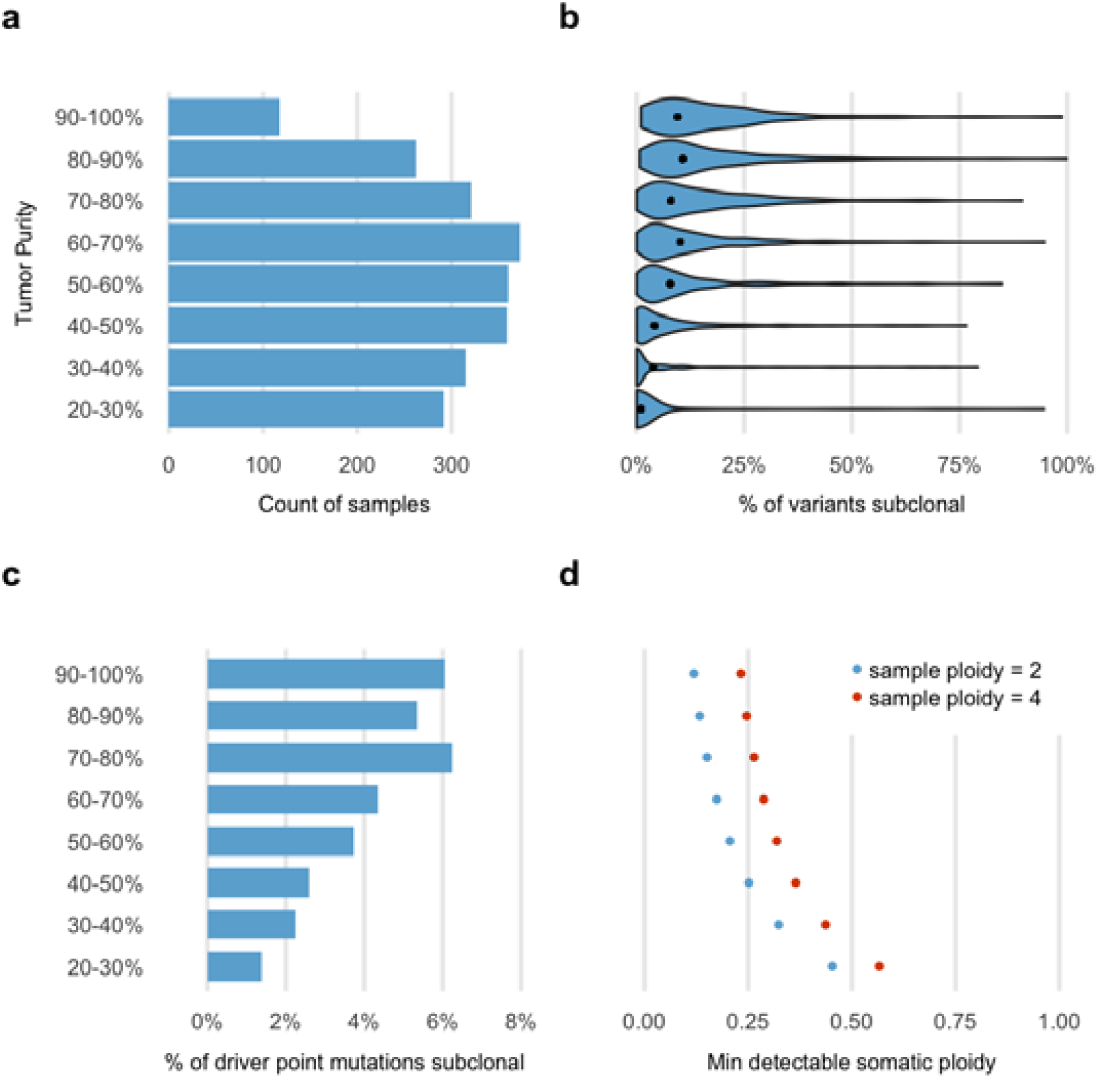
Subclonality of somatic variants. a) Count of samples per tumor purity bucket. b) Violin plot showing the percentage of point mutations which are subclonal in each purity bucket per sample. Black dots indicate the mean for each bucket. c) Percentage of driver point mutations that are subclonal in each purity bucket. d) Approximate somatic ploidy detection cutoff of the HMF pipeline at median 106x depth coverage for each purity bucket and for sample ploidy 2 and 4. Subclonal variants with cellular fraction less than this cutoff are unlikely to be detected by our pipeline analyses.

