## Supplementary material for "Pan-cancer whole genome analyses of metastatic solid tumors": Detailed methods

***Content***

1. Sample collection 2
2. Sequencing workflow 2
3. Somatic point mutation calling 2
4. Validation of somatic point mutation calling 3
5. Mutational Signature analysis 6
6. Somatic structural variant calling 6
7. Identification of gene fusions 6
8. Validation of gene fusions 7
9. Purity, ploidy and copy number calling 8
10. Validation of purity, ploidy and copy number output 11
11. Sample filtering based on copy number output 13
12. Impact of sequencing depth coverage and analysis pipeline on somatic variant calling sensitivity 13
13. Germline predisposition variant calling 15
14. Clonality and biallelic status of point mutations 17
15. WGD status determination 18
16. MSI status determination 19
17. Holistic gene panel for driver discovery 20
18. Significantly mutated driver genes discovery 20
19. Significantly amplified & deleted driver gene discovery 20
20. Fragile site annotation 22
21. Somatic driver catalog construction 23
22. Driver co-occurrence analysis 25
23. Actionability analysis 26
24. Data availability 30
25. References 30

### 1. Sample collection

Patients with advanced cancer not curable by local treatment options and being candidates for any type of systemic treatment and any line of treatment were included as part of the CPCT-02 (NCT01855477) and DRUP (NCT02925234) clinical studies, which were approved by the medical ethical committees (METC) of the University Medical Center Utrecht and the Netherlands Cancer Institute, respectively. A total of 41 academic, teaching and general hospitals across the Netherlands participated in these studies and collected material and clinical data by standardized protocols[1]. Patients have given explicit consent for whole genome sequencing and data sharing for cancer research purposes. Clinical data, including primary tumor type, biopsy location, gender and birth year were collected in electronic case record forms and stored in a central database.

Core needle biopsies were sampled from the metastatic lesion, or when considered not feasible or not safe, from the primary tumor site when still in situ. One to four biopsies were collected (average of 2.1 per patient) and frozen in liquid nitrogen directly after sampling and further processed at a central pathology tissue facility. Frozen biopsies were mounted on a microtome in water droplets for optimal preservation of all types of biomolecules (DNA, RNA and proteins) for subsequent and future omics-based analyses. A single 6 micron section was collected for hematoxylin-eosin (HE) staining and estimation of tumor cellularity by an experienced pathologist. Subsequently, 25 sections of 20 micron, containing an estimated 25,000 to 500,000 cells, were collected in a tube for DNA isolation. In parallel, a tube of blood was collected in CellSave (Menarini-Silicon Biosystems) tubes, which was shipped by room temperature to the central sequencing facility at the Hartwig Medical Foundation. Left-over material (biopsy, DNA) after sample processing was stored in biobanks associated with the studies at the University Medical Center Utrecht and the Netherlands Cancer Institute.

### 2. Sequencing workflow

DNA was isolated from biopsy and blood on an automated setup (QiaSymphony) according to supplier's protocols (Qiagen) using the DSP DNA Midi kit for blood and QIAsymphony DSP DNA Mini kit for tissue and quantified (Qubit). Before starting DNA isolation from tissue, the biopsy was dissolved in 100 microliter Nuclease-free water by using the Qiagen TissueLyzer and split in two equal fractions for parallel automated DNA and RNA isolation (QiaSymphony). Typically, DNA yield for the tissue biopsy ranged between 50 and 5,000 ng. A total of 50 - 200 ng of DNA was used as input for TruSeq Nano LT library preparation (Illumina), which was performed on an automated liquid handling platform (Beckman Coulter). DNA was sheared using sonication (Covaris) to average fragment lengths of 450 nt. Barcoded libraries were sequenced as pools (blood control 1 lane equivalent, tumor 3 lane equivalents) on HiSeqX (V2.5 reagents) generating 2 x 150 read pairs using standard settings (Illumina).

BCL output from the HiSeqX platform was converted using bcl2fastq tool (Illumina, versions 2.17 to 2.20 have been used) using default parameters. Reads were mapped to the reference genome GRCH37 using BWA-mem v0.7.5a[2], duplicates were marked for filtering and INDELs were realigned using GATK v3.4.46 IndelRealigner[3]. GATK HaplotypeCaller v3.4.46[4] was run to call germline variants in the reference sample. For somatic SNV and INDEL variant calling, GATK BQSR[5] is also applied to recalibrate base qualities.

### 3. Somatic point mutation calling

We called SNV & INDEL somatic variants using Strelka v1.0.14[6] with the following optimisations:

- **Preservation of known variants:** From the raw Strelka output we marked all known pathogenic variants from external databases such that these would be preserved from all subsequent filtering. The list of pathogenic variants used was the union of:
  - Point mutations in CIViC[7] with level C evidence or higher (download = 01-mar-2018)
  - Somatic variants from CGI[8] (update: 17-jan-2018)
  - Oncogenic or likelyOncogenic variants from OncoKb[9] (download = 01-mar-2018); http://oncokb.org/api/v1/utils/allAnnotatedVariants.txt)
  - TERT promoter variants at genomic coordinates: 5:1295242, 5:1295228, 5:1295250
- **Modified quality score filtering**
  - We split variants into high confidence (HC) and low confidence (LC) regions using the NA12878 GIABv3.2.2 high confidence region definitions[10], based on the observation that we produce far higher rates of false positives variant calls in LC regions
  - Set quality score cutoffs for SNV & INDEL to 10 for HC regions and 20 for LC regions (default = 15 for SNV, 30 for INDEL)
  - Added an additional quality filter to tighten filtering for low allelic frequency variants: quality score * allele frequency > 1.3
- **Improved repeat sensitivity:** Switched off the default Strelka repeat filter to improve indel calling in microsatellites and short repeats.
- **Panel of normals (PON) to remove germline leakage:** Filtered out any variants which were found by GATK haplotypecaller in more than 5 samples in a germline PON consisting of 2000 of our reference blood samples. PON available at (https://resources.hartwigmedicalfoundation.nl/)
- **PON to remove strelka-specific artefacts:** Filtered any variant which was supported by 2 or more reads in strelka in the reference sample in at least 4 patients in our cohort. PON available at (https://resources.hartwigmedicalfoundation.nl/)
- **Removal of INDELS near a PON filtered INDEL -** Regions of complex haplotype alterations are often called as multiple long indels, which can make it more difficult to construct an effective PON, and sometimes we find residual artefacts at these locations. Hence we also filter inserts or deletes which are 3 bases or longer where there is a PON-filtered INDEL of 3 bases or longer within 10 bases in the same sample.
- **MNV Correction** - Variants occurring on consecutive positions, or 1 base apart were considered potential multi nucleotide variants (MNVs). The BAM files were re-examined, and the variants were merged into a single MNV if greater than 80% of the reads with a mapping quality score of at least 10 and which are neither unmapped, duplicated, secondary, nor supplementary containing any of the individual variants also contained the other variants of the potential MNV. The attributes of the resulting MNV variant were determined by picking the minimum values from the individual variants forming the MNV. MNVs were marked as PON filtered only if both individual variants were PON filtered.

The settings and tools for this optimized HMF pipeline are available at https://github.com/hartwigmedical/.

### *4. Validation of somatic point mutation calling*

We performed three separate analyses to validate our somatic variant calling pipeline as follows:

#### 4.1. Validation of somatic precision and sensitivity pipeline on a known benchmark

We tested the default Strelka and HMF optimized settings on a GIAB mix-in sample (ref = NA24385; tumor = 70% NA24385 and 30% NA12878) to test sensitivity at a realistic purity and on a null tumor (ref = NA12878, tumor = NA12878) to test precision. The results of this analysis are as follows:

| Configuration | SNV sensitivity | SNV false positive / genome | INDEL sensitivity | Indel false positive / genome |
|---|---|---|---|---|
| Strelka default | 93% | 3500 | 24% | 41 |
| Optimized HMF pipeline | 96% | 109 | 77% | 27 |

**4.2. External independent validation of SNV and INDEL calling precision on real samples**
We performed external validation of a set of single nucleotide variants (SNV) and short insertion/deletions (indels) that have been detected by Whole Genome Sequencing (WGS) using the single molecule Molecular Inversion Probe (smMIP) technology[11]. SNV and short indels variants were semi-randomly selected from 30 patient samples. The first selection was to include every variant that was reported in a panel of 114 'actionable' cancer genes as used in the routine CPCT-02 study analysis. This way, a total of 82 variants (67 SNVs, 15 indels) were selected in 45 genes. The second selection involved random sampling adding up to a total of 256 coding and non-coding variants from the same 30 patient samples.

A custom smMIP panel was designed to cover the selected variants. For 45 variants (17.6%) no smMIP design was possible, all of which were intergenic variants. For the other 211 variants probes could successfully be designed. Analysis of the smMIP sequencing data indicated that for 17 of the 211 variants (8.1%) the smMIP sequencing data was of insufficient quality (mostly due to repeat stretches), while the WGS data seemed sufficiently reliable for accurate calling (confirmed by visual inspection of the read data), including 3 coding variants (*RB1, ERBB4* and *BRCA2*) and 14 intergenic regions. The retrospective investigation of the WGS data indicated that for another three variants (1.4%) the smMIP as well as the WGS data was of insufficient quality due to large homopolymer stretches.

In total 192 variants could be successfully sequenced and analyzed using the smMIP and could be used for confirmation of the WGS findings. 189 SNVs and indel variants (98.4%) were confirmed by smMIP sequencing, indicating a very high accuracy of WGS-derived variant calling results. All three variants that could not be confirmed by smMIP were from intergenic regions, including 1 variant that showed a mixed double-variant (chr3:75887550_G>T/C) and for which both technologies had difficulties in accurately calling the genotype. For the remaining 2 variants (chr8:106533360_106533361insAC, chr12:125662751_125662752insA), it remains unclear if these could not be detected by smMIP or were falsely called by WGS, as they fall in repetitive genomic stretches.

38 out of the 211 orthogonally validated variants by smSMP analysis were INDELS. From these 8 could not be validated because no smSMP data was obtained although manual inspection showed reliable WGS data and variant calls. An additional three did not validate, but also looked suspicious in the WGS data upon manual inspection. Two variants were not validated by smSMP. So when considering only INDEL variants for which independent smMIP data was available (27 variants), 2 (7.4%) were not validated.

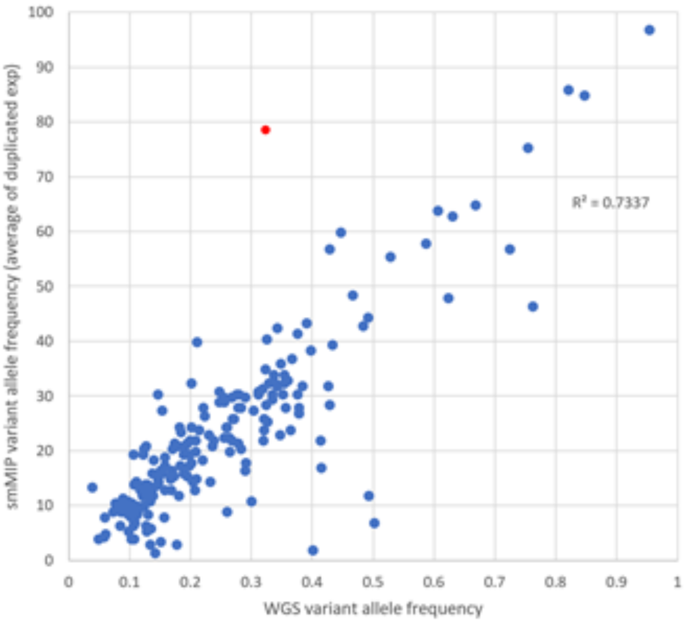


The 189 successfully confirmed variants showed a good linear correlation in variant allele frequency between WGS and smMIP sequencing (average of duplicates) with an $R^2$ of 0.733. This result indicated that WGS, with its lower read depth (on average between 100-110x) than smMIP and without a read-barcoding system, is accurate in quantitatively determining the variant frequency at frequencies above 5%. One variant (ch19:55276095C>T, indicated in red in the figure above) showed a large deviation in variant frequency, which was likely due to the much lower than expected coverage of the variant, both in the WGS (37 reads) as well as in the smMIP data (28 and 35 reads).

**4.3. Validation of somatic variant calling sensitivity by reanalysis of known hotspots.**

To validate somatic calling sensitivity and performance limitations of our pipeline on real samples, we built a customised tool, SAGE (https://github.com/hartwigmedical/hmftools/tree/master/sage) to reanalyse all 10,211 known pathogenic hotspot variants in the coding region of the genome (sourced from CIVIC, OncoKb and CGI as described above). These locations have a much higher prior likelihood of finding a variant in cancer samples.

SAGE searches for each hotspot in the tumor BAM files directly and calls a variant if the sum of read base qualities supporting the ALT > 100, effectively equating to 3 high quality reads of support. Our standard somatic pipeline typically requires 6 or more reads support to call a variant. For the purposes of this validation we excluded from SAGE a small number of variants in high repeat contexts (repeat count >=8) and in regions with very high tumor copy number (tumor read depth > 300) as both these contexts can cause low VAF artefacts which we want to avoid in a sensitivity test.

We evaluated on a randomly selected 1247 samples with the following results

| Hotspot variants found in standard somatic pipeline | Additional variants found by SAGE | % variants missed by somatic pipeline |
|---|---|---|
| 1160 | 37 | 3.1% |

Of the 37 additional variants found by SAGE but not in our standard somatic pipeline, 27 (2.3%) were found to have been missed by Strelka due to low read count in the tumor (all with only 3 to 6 reads supporting the ALT allele), 8 (0.7%) due to insufficient coverage in the reference sample, and 2 (0.2%) for unknown reason.

Overall this analysis suggests that we capture more than 96% of all variants with 3 or more reads of support in the tumor (equivalent to ~3% VAF).

### *5. Mutational Signature analysis*

Mutational signatures were determined by fitting SNV counts per 96 tri-nucleotide context to the 30 COSMIC signatures[12] using the mutationalPatterns package[13]. Signatures with <5% overall contribution to a sample or absolute fitted muational load <300 variants were excluded from the plot (Extended Data Fig. 3). Residuals were calculated as the sum of the absolute difference between observed and fitted across the 96 buckets. A small number of samples had extremely high residuals suggesting potential novel mutational signatures. In particular, 2 of the outlier samples (HMF002896 and HMF001562) also had a very high mutational load and strikingly similar mutational signatures dominated by T>A transversions (Extended Data Fig. 3C).

### *6. Somatic structural variant calling*

Structural Variants were called using Manta(v1.0.3)[14] with default parameters. We then re-examined each breakpoint, calculated variant allele frequencies for each break end and applied seven additional filters to the Manta output to improve precision using an internally built tool called ‘Breakpoint-Inspector’ (BPI, https://github.com/hartwigmedical/hmftools/tree/master/break-point-inspector) v1.5. Two main types of filters are applied by BPI:

- **Evidence of variant in reference sample** - Variants are filtered out if we can find any evidence of paired read support, split read support or soft clipping concordance (5+ bases at exact breakpoint) in the matching blood sample.
- **Inadequate support for variant in tumor sample** - For all inversions and translocations and for long deletions and tandem duplications (>1000 bases between breakpoints) we require at least 1 read with paired read support. For short deletions and duplications (<1000 bases between breakpoints) we require at least 1 read with split read support. In both cases at least one of those reads must be anchored with at least 30 bases at each breakpoint. We also require the minimum read coverage across each breakpoint in the tumor to be > 10 depth.

Each breakend was annotated with it’s position in all transcripts from ‘KNOWN’ genes in Ensembl v89.37[15]. Each gene was marked as disrupted if there was at least one structural variant that impacted on the canonical transcript.

### *7. Identification of gene fusions*

For each structural variant, every combination of annotated overlapping transcripts from each breakend was tested to see if it could potentially form an intronic inframe fusion. A list of 411 curated known fusion pairs was sourced by taking the union of known fusions from the following external databases:

- Cosmic curated fusions[12] (v83)
- OncoKb[9] (download = 01-mar-2018)
- CGI[8] (update: 17-jan-2018)
- CIViC[7] (download = 01-mar-2018)

We then also created a list of promiscuous fusion partners using the following rules

- **3' promiscuous:** Any gene which appears on the 3' side in more than 3 of the curated fusion pairs OR appears at least once on the 3' side and is marked as promiscuous in either OncoKb, CGI or CIVIC
- **5' promiscuous:** Any gene which appears on the 5' side in more than 3 of the curated fusion pairs OR appears at least once on the 5' side and is marked as promiscuous in either OncoKb, CGI or CIVIC

For each promiscuous partner we also curated a list of essential domains that must be preserved to form a viable fusion partner.

Finally, we report an intronic inframe fusion if the following conditions are met

- Matches an exact fusion from the curated list OR is intergenic and matches 5' promiscuous OR matches 3' promiscuous gene
- Curated domains are preserved
- Does not involve the 3'UTR region of either gene
- For intragenic fusions, must start and end in coding regions of the gene
- 3' partner is a protein coding gene and the transcript does not result in nonsense mediated decay

#### *8. Validation of gene fusions*

Whole transcriptome analysis (RNA-seq) of 60 samples with identified fusions was used to validate our gene fusion calling pipeline.

RNA was isolated from the same biopsy material as used for DNA isolation using an automated setup (QiaSymphony) using the QIAsymphony RNA kit (#931636, Qiagen) according to supplier's protocols. RNA was quantified using Qubit RNA HS Assay Kit (Thermo Fisher). Typically, RNA yield for the tissue biopsy ranged between 500 and 5,000 ng. 100 ng of total RNA was used as input for KAPA RNA HyperPrep Kit with RiboErase (HMR) (#KR1351, Roche) and TruSeq DNA CD Indexes 96 Indexes (#PN 20015949, Illumina) performed on an automated liquid handling platform (Beckman Coulter). The standard protocol used involved 240 sec 85 degrees Celcius fragmentation and 15 PCR cycles. Each sample was subsequently sequenced in a multiplexed setup with 2x75 bp reads on a NextSeq 500/550 using the High Output Kit v2 (Illumina, #FC-404-2002), targeting 50M raw reads per sample. BCL output from the NextSeq500 platform was converted using Illumina bcl2fastq tool (versions 2.17 to 2.20 have been used) using default parameters.

STAR-Fusion[16] was used with default settings to call fusion transcripts from the RNA. 37 out of 63 fusions were readily identified independently in the RNA. Manual inspection of the expected chimeric junctions for the remaining 22 fusions revealed RNA support for a further 6 fusions (4 of which were TMPRSS2-ERG), although below the threshold to be called automatically in the RNA with the settings used. Overall, 68% of the tested fusions were thus independently validated by the RNA analysis. The full results are summarised below:

| Total fusions tested | Transcript fusion found by STAR-Fusion | Read support in RNA but not called by STAR-Fusion | No evidence of fusion transcript in RNA |
|---|---|---|---|
| 63 | 37 (59%) | 6 (10%) | 16 (32%) |

### *9. Purity, ploidy and copy number calling*

Accurate copy number calling is closely linked with correct sample purity determination. Currently, there is not a clear consensus in the community for a preferred tool for this purpose. We tested several tools (freeC, CANVAS and Sequenza) on the COLO829 benchmark, but none of them provided a correct fit[17]. Therefore we developed PURPLE (PURity & PLoidy Estimator) as an alternative.

PURPLE combines B-allele frequency (BAF), read depth and structural variants to estimate the purity and copy number profile of a tumor sample and follows a similar purity fitting methodology to several other popular tools such as ASCAT, Sequenza and CANVAS, only with a different optimisation function to determine the best fit.

The main advantages of PURPLE (v2.14) for the purposes of this study are:

- extensive attention to removal of artefacts by filtering of inputs (see below sections 7.1, 7.2 and 7.3) and smoothing of output to avoid false positive copy number calling (section 7.5)
- integrated SV and copy number calling allow single base accuracy of copy number calls and accurately call each individual variant as heterozygous or homozygous as well as the detection of partial loss of genes

There are five key steps in the PURPLE pipeline:

**1. Calculate BAF in tumor at high confidence heterozygous germline loci**
We determine the BAF of the tumor sample by finding heterozygous locations in the reference sample from a panel of 796,447 common germline heterozygous SNP locations. To ensure that we only capture heterozygous points, we filter the panel to only loci with allelic frequencies in the reference sample between 40% and 60% and with depth between 50% and 150% of the reference sample genome wide average. Typically, this yields 140k-200k heterozygous germline variants per patient. We then calculate the allelic frequency of corresponding locations in the tumor.

**2. Determine read depth ratios for tumor and reference genomes**
The raw read counts per 1,000 base window for both normal and tumor samples, by counting the number of alignment starts in the respective bam files with a mapping quality score of at least 10 that is neither unmapped, duplicated, secondary, nor supplementary. Windows with a GC content less than 0.2 or greater than 0.6 or with an average mappability below 0.85 are excluded from further analysis.

Next we apply a GC normalization to calculate the read ratios. We divide the read count of each window by the median read count of all windows sharing the same GC content then normalise further to the ratio of the median to mean read count of all windows.

Finally, the reference sample ratios have a further ‘diploid’ normalization applied to them to remove megabase scale GC biases. This normalization assumes that the median ratio of each 10Mb window (minimum 1Mb readable) should be diploid for autosomes and haploid for sex chromosomes in males in the germline sample.

**3. Segmentation**
We segment the genome into regions of uniform copy number by combining segments generated from the read ratios for both tumor and reference sample, from the BAF points with structural variant breakpoints derived from Manta & BPI. Read ratios and BAF points are segmented independently using the Bioconductor copynumber package[18] which uses a piecewise constant fit (PCF) algorithm (with

custom settings gamma = 100, k =1). These segment breaks are then combined with the structural variants breaks according to the following rules:

1. Every structural variant break starts a new segment, as does chromosome starts, ends and centromeres. This is regardless of if they are distinguishable from existing segments or not.
2. Ratio and BAF segment breaks are only included if they are distinguishable from an existing segment.
3. To be distinguishable, a break must be at least one complete mappable read depth window away from an existing segment.

Once the segments have been established we map our observations to them. In each segment we take the median BAF of the tumor sample and the median read ratio of the tumor and reference samples. We also record the number of BAF points within the segment as the BAFCount.

A reference sample copy number status is determined at this this stage based on the observed copy number ratio in the reference sample, either 'DIPLOID' (0.8<= read depth ratio<=1.2), 'HETEROZYGOUS_DELETION' (0.1<=ratio<0.8), 'HOMOZYGOUS_DELETION' (ratio<0.1),'AMPLIFICATION'(1.2<ratio<=2.2)or 'NOISE' (ratio>2.2). The purity fitting and smoothing steps below use only the DIPLOID germline segments.

**4. Purity Fitting**
Next we jointly fit tumor purity and sample ploidy (expressed as a normalisation factor) according to the following principles:

1. The absolute copy number of each segment should be close to an integer ploidy
2. The BAF of each segment should be close to a % implied by integer major and minor allele ploidies.
3. Higher ploidies have more degenerate fits but are less biologically plausible and should be penalised
4. Segments are weighted by the count of BAF observations which is treated as a proxy for confidence of BAF and read depth ratio inputs.
5. Segments with lower observed BAFs have more degenerate fits and are weighted less in the fit

For any given tumor purity and sample ploidy we calculate the score by first modelling the major and minor allele ploidy of each segment and minimising the deviation between the observed and modelled values according to the following formulas:

ModelDeviation = abs(ObservedRatio - ModelRatio) + abs(ObservedBaf - ModelBaf)
ModelBaf = (tumorPurity * (segmentMinorPloidy - 1) + 1) / (tumorPurity * (segmentPloidy - 2) + 2)
ModelRatio = sampleNormFactor + (segmentPloidy - 2) * tumorPurity * sampleNormFactor / 2d;

Once modelled, each segment is given a ploidy penalty:

PloidyPenalty = 1 +min(SingleEventDistance, WholeGenomeDoublingDistance);
WholeGenomeDoublingDistance = 1 + abs(segmentMajorAllele - 2) +abs(segmentMinorAllele - 2);
SingleEventDistance = abs(segmentMajorAllele - 1) + abs(segmentMinorAllele - 1);

Summing up over all the segments generates a score for each tumor purity / sample ploidy combination from which we can select the minimum:

$$FitedPurityScore = \frac{1}{TotalBafCount}\sum_{i=1}^{n} PloidyPenalty_i \times ModelDeviation_i \times BafCount_i \times ObservedBaf_i$$

If a sample has a fitted purity solution which is >98.5% diploid and a score within 10% of the best fitted score, the sample is designated as highly diploid and a fit is determined by the highest vaf somatic ploidy peak.

Given a fitted purity and sample ploidy we are then able to determine the purity adjusted copy number and BAF of each segment in the tumor genome from the unadjusted read ratios and BAFs respectively.

**5. Smoothing**

Since the segmentation algorithm is highly sensitive, and there is a significant amount of noise in the read depth in whole genome sequencing, many adjacent segments created above will have a similar copy number and BAF profile and can be combined and averaged to form a larger, smoothed, region.

We apply a number of rules to merge adjacent regions to create a smooth copy number profile.

1. Never merge a segment break created from a structural variant break end.
2. Use the count of BAF points as a proxy for confidence or weight in the region. Note that some segments may have a BAF count of 0.
3. Merge segments where the difference in BAF and copy number is within tolerances.
4. BAF tolerance is linear between 0.03 and 0.35 dependent on BAF count.
5. Copy number tolerance is linear between 0.3 and 0.7 dependent on BAF count. The tolerance also increases linearly as purity of the tumor sample decreases below 20%.
6. Start from most confident segment and smooth outwards until we reach a segment outside of tolerance. Move on to next highest unsmoothed section.
7. It is possible to merge in (multiple) segments that would otherwise be outside of tolerances if:
    a. The total dubious region is sufficiently small (<30k bases or <50k bases if approaching centromere); and
    b. The dubious region does not end because of a structural variant; and
    c. The dubious region ends at a centromere, telomere or a segment that is within tolerances.
8. When the entire short arm of a chromosome is lacking copy number information (generally on chromosome 13, 14, 15, 21 or 22), the copy number of the long arm is extended to the short arm.
9. Any remaining unknown segments are given the expected copy number of their associated chromosome, i.e. 2 for autosomes and female allosomes, 1 for male allosomes.

Where clusters of SVs exist which are closer together than our read depth ratio window resolution of 1,000 bases, the segments in between will not have any copy number information associated with them. To resolve this, we infer the ploidy from the surrounding copy number regions. The outermost segment of any SV cluster will be associated with a structural variant with a ploidy that can be determined from the adjacent copy number region and the VAF of the SV. We use this ploidy and the orientation of structural variant to calculate the change in copy number across the SV and hence the copy number of the outermost unknown segment. We repeat this process iteratively and infer the copy number of all regions within a cluster.

Once region smoothing is complete, it is possible there will be regions of unknown BAF, if no BAF points were present in a copy number region. We infer this BAF by assuming that they share their minor allele ploidy with their neighbouring region. If there are multiple neighbouring regions with known BAF we use the highest confident region (i.e. highest BAF count) to infer.

At this stage we have determined a copy number and minor allele ploidy for every base in the genome.

### *10. Validation of purity, ploidy and copy number output*

We performed three validations to evaluate the purity and ploidy estimates and copy number profile obtained from PURPLE.

**1. Validation of purity estimates through cell line in-silico dilutions**
The purity estimates of PURPLE were validated using the tumor cell line COLO829. We created diluted in-silico mixture models of the tumor and blood cell lines from COLO829 with simulated purities of 20%, 30%, 40%, 60%, 80% and 100%, and ran PURPLE on the simulated BAM files against the reference sample.
The PURPLE estimates were found to match the simulation very closely as shown in the table below:

| Simulated Purity | PURPLE estimated purity | Difference |
|---|---|---|
| 20% | 20% | 0% |
| 30% | 30% | 0% |
| 40% | 40% | 0% |
| 50% | 50% | 0% |
| 60% | 60% | 0% |
| 80% | 81% | 1% |
| 100% | 100% | 0% |

**2. Validation of absolute copy number predictions by FISH**

We also validated the absolute copy number results for PURPLE by comparing the WGS analysis results of the COLO-829 tumor vs normal cell line pair with DNA Fluorescence In Situ Hybridization (FISH) results for the centromeric region of chromosome 9, 13, 16 and 18 (CEP9, CEP13, CEP16, CEP18) and for the 2p23 ALK locus and the 9p24 JAK2 locus. In total, 100 COLO829 tumor cells were scored for each of the six FISH probes. For both assays the local copy-number as well as the percentage of DNA (PURPLE) or number of cells (FISH) is provided in the table below to indicate the intratumoral heterogeneity. The FISH and sequencing based results showed a very high concordance for the chromosomal copy numbers and the intratumoral heterogeneity (COLO-829 cell line heterogeneity has been described previously[19]).

| Genomic region | PURPLE ploidy and purity | FISH copy number |
|---|---|---|
| Centromere Chr 9 | 3.7-4.0 : 53-57% | *2n : 33%*<br>3n : 9%<br>4n : 58% |

| | | |
|---|---|---|
| Centromere Chr 13 | 3.2 : 55% | *2n* : *41%*<br>3n : 59% |
| Centromere Chr 16 | *2.0 : 100%* | *2n* : *100%* |
| Centromere Chr 18 | 2.8-2.9 : 67-71% | 2n : 38%<br>3n : 62% |
| ALK (2p23) | 3.1 : 67% | 2n : 21%<br>3n : 79% |
| JAK2 (9p24) | *2.0 : 100%* | *2n* : *100%* |

**3. Comparison of PURPLE purity and ploidy estimates on patient samples with ASCAT**

To validate PURPLE on real patient data, we compared the purity and ploidy outputs from PURPLE to the widely used copy number tool ASCAT[20] for 65 randomly selected samples from our cohort. ASCAT was run on GC corrected data using default parameters except for gamma which was set to 1 which is recommended for massively parallel sequencing data.

The following charts show a comparison of ASCAT vs PURPLE purity and ploidy results with 55 of 65 samples (85%) in agreement to within 10% absolute purity and relative sample ploidy.

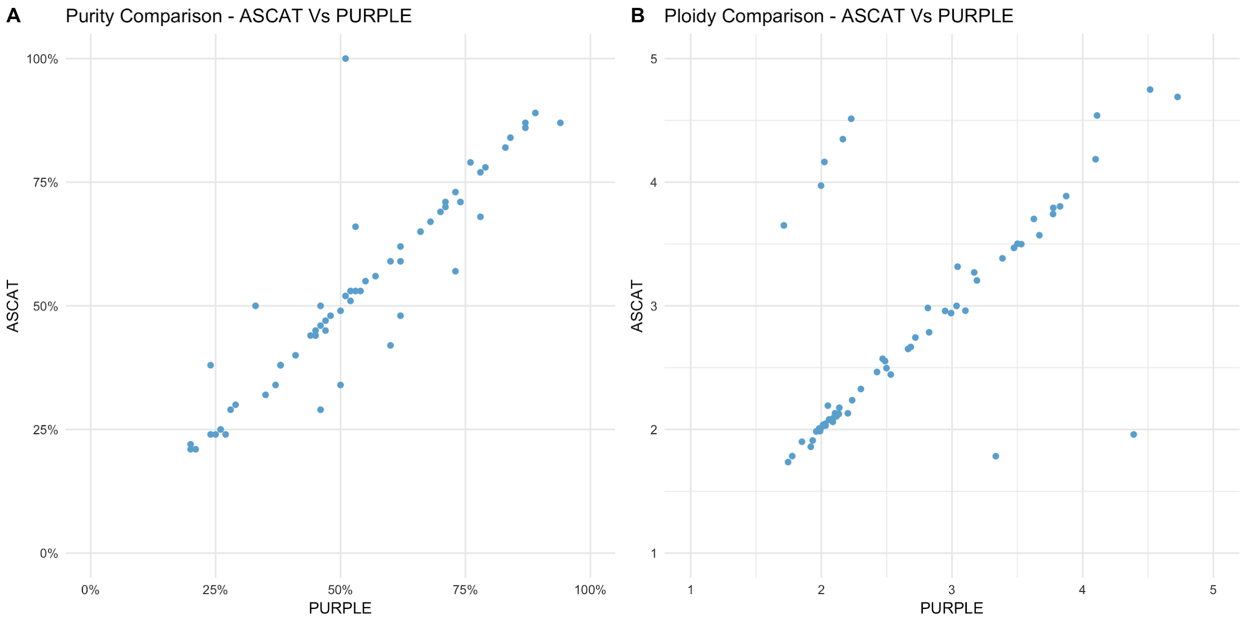


There are 2 types of differences observed in the remaining 10 samples:

- Purity differences for highly diploid samples - this is unsurprising as PURPLE has additional functionality which is not dependent on copy number alterations in the tumor for highly diploid samples to fit the somatic ploidies whereas ASCAT does not.
- Whole genome duplication (WGD) vs no whole genome duplication - In 5 of the samples ASCAT calls a WGD event whereas PURPLE does not and in 2 samples the opposite occurs. This reflects the tradeoff in the purity and ploidy determination between penalising higher ploidy solutions which are more degenerate vs lower ploidy solutions with more subclonality. Manual inspection of purity-corrected fitted minor allele ploidy plots reveals in all of the 5 cases where

ASCAT calls a WGD that whilst there is subclonality in each of these cases in the PURPLE solution there is no subclonal peak at 0.5 copy number, nor is there a 0.5 somatic ploidy peak, suggesting that the the WGD solution is less likely. Conversely, in the 2 cases where PURPLE only calls a WGD, manual inspection shows that the ASCAT solution would be prefered in one case and the PURPLE solution in the other.

In summary, overall concordance is very high between PURPLE and ASCAT. There appears to be little systematic bias to either calling lower or higher ploidy solutions between methods, and where PURPLE differs from ASCAT it more often than not appears to be the more plausible solution.

### *11. Sample filtering based on copy number output*

Following our copy number calling, samples were QC filtered from the analysis based on 4 criteria:

- **NO_TUMOR** - If PURPLE fails to find any aneuploidy AND the number of somatic SNVs found is less than 1,000 then the sample is marked as NO_TUMOR.
- **MIN_PURITY** - We exclude samples with a fitted purity of <20%
- **FAIL_SEGMENT** - We remove samples with more than 120 copy number segments unsupported at either end by SV breakpoints. This step was added to remove samples with extreme GC bias, with differences in depth of up to or in excess of 10x between high and low GC regions. GC normalisation is unreliable when the corrections are so extreme so we filter.
- **FAIL_DELETED_GENES -** We removed any samples with more than 280 deleted genes. This QC step was added after observing that in a handful of samples with high MB scale positive GC bias we sometimes systematically underestimate the copy number in high GC regions. This can lead us to incorrectly infer homozygous loss of entire chromosomes, particularly on chromosome 19.

Where multiple biopsies exist for a single patient, we always choose the highest purity sample for our analysis of mutational load and drivers.

### *12. Impact of sequencing depth coverage and analysis pipeline on somatic variant calling sensitivity*

#### 12.1 Downsampling

To assess the impact of our sequencing depth on variant calling sensitivity, we selected 10 samples at random, downsampled the BAMs by 50% to about 50x average read coverage (similar to the mean depth of the PCAWG cohort[21]). We then reran the identical somatic variant calling pipeline.

Comparing the output to the original runs, we found near identical purities and ploidies for the down sampled runs (Extended Data Fig. 2). We observed an average decrease in sensitivity of 10% for SNV, 15% for MNV, 19% for SV, and 2% for INDEL.

The relatively small drop in indel calling sensitivity upon downsampling is caused by hard-coded setting in STRELKA. Strelka has a hard cutoff at 10% VAF for INDELs of less than 5 bases length (which is 99% of INDELs in our dataset) for both 50x and 100x depth whereas for SNVs the cutoff is fixed at ~5 supporting reads independent of read depth. This likely results in underestimation of subclonal INDELs in our dataset but does not affect specificity.

#### 12.2 Assessment of impact of pipeline differences on variant calling

Using the EGA data portal, we downloaded tumor and matching normal bam files from the following ICGC-PCAWG donor IDs: DO1011, DO1012, DO1020, DO217826, DO218065, DO218174, DO218583, DO218656, DO218669, DO218709, DO218796, DO220823, DO46350, DO46366, DO46370, DO46372, DO46386, DO46412, DO46448, DO46551, DO46591, DO52543, DO52554, DO52559 (The ICGC-PCAWG controlled data access was granted under the Data Access Compliance Office (DACO) Application Number: DACO-1050905). Subsequently, these bam files were reformatted to FASTQ files (SamToFastq PICARD v2.1.0 https://github.com/broadinstitute/picard) which were realigned to reference genome GRCH37 lacking the GL0000XX contigs (BWA-mem v0.7.5a2). These FASTQ files were used as input for the standard HMF analysis workflow as described above. The obtained somatic events were compared with the consensus somatic calls that were generated by the PCAWG SNV Calling Working Groups[22,23].

We compared the mutational load for each variant type on each of the 24 samples (Extended Data Fig.5). In the comparison we excluded trinucleotide MNVs (not called in the PCAWG pipeline and accounting for 11% of HMF MNVs). We also excluded translocations found in the HMF pipeline with a somatic ploidy of less than 0.2. These appear to be a pervasive artefact found by the HMF SV pipeline in many of the 24 PCAWG samples making up 17% of all variants in the PCAWG samples (compared to only 2% in the HMF cohort) and 91% of all SV with less than 0.2 somatic ploidy (compared to 33% in the HMF cohort). We also observed that the PCAWG pipeline makes no SV calls with less than 500 bases in length and few less than 1000 bases in length, whereas this is a common variant type in the HMF cohort making up 17% of all SV.

The comparison of HMF and PCAWG pipelines after excluding these variants on a per sample basis are presented in Extended Data Fig 5. The overall difference in variants called by each pipeline is summarised in the table below

| **Total # of variants found in 24 PCAWG validation samples** | | | |
|---|---|---|---|
| **Variant Type** | **HMF Pipeline** | **PCAWG Pipeline** | **PCAWG relative sensitivity** |
| SNV | 228435 | 235944 | +3% |
| INDEL | 59189 | 33736 | -43% |
| MNV* | 1521 | 1591 | +5% |
| SV** | 5451 | 5405 | -1% |

* Only dinucleotide MNV are compared for this analysis (3+ nucleotide not present in PCAWG cohort)
** SV with 1000 bases excluded from the analysis (not present in PCAWG cohort)

A more detailed analysis at a per variant level shows that for each variant type between 83% and 88% of the PCAWG called variants are also called by HMF, rising to 88% to 96% after excluding variants found with <=5 supporting reads in PCAWG, but only 34% to 52% for variants with <= 5 reads as presented in the table below. Somatic variant calling at low read support is a trade-off between precision and sensitivity. Since the HMF pipeline has been optimised for ~100x depth vs the ~50x depth in PCAWG, it is not surprising that the cut-offs are more stringent in the HMF variant calling.

| **% of PCAWG PASS variants found in HMF pipeline** | | | |
|---|---|---|---|
| **Variant Type** | **ALL Variants** | **>5 supporting reads in PCAWG** | **<=5 supporting reads in PCAWG** |
| SNV | 88% | 96% | 34% |
| INDEL | 87% | 88% | 48% |
| MNV | 82% | 93% | 50% |

| | | | |
|---|---|---|---|
| SV | 86% | N/A | N/A |

Conversely, HMF called variants are found by PCAWG 91% of the time for SNV, 86% of the time for MNV but only 50% for INDELs.

| % of HMF PASS variants found in PCAWG pipeline | |
|---|---|
| Variant Type | All Variants |
| SNV | 91% |
| INDEL | 50% |
| MNV* | 86% |
| SV** | 85% |

* Only dinucleotide MNV are compared for this analysis (3+ nucleotide not present in PCAWG cohort)
** SV with 1000 bases excluded from the analysis (not present in PCAWG cohort)

The significant difference in INDEL sensitivity is largely explained by INDELs in long repeats and microsatelites, with only 3% of HMF INDELs in a repeat sequence of more than 10 consecutive repeats found by PCAWG. A significant proportion of INDELs were also filtered by the panel of normals in PCAWG. Since 85% of INDELs fall in microsatellites or repeats, and only a limited number of such sites exist in the human genome, many genuine INDEL variants will be found in a panel of normals, so this result is also not unexpected.

Taken together with the results of the downsampling experiments, and allowing for the variant types not called in the PCAWG cohort, we expect approximately the following overall differences in reported TMB:

| Expected change in TMB in HMF cohort from downsampling to PCAWG depth and using PCAWG variant calling pipeline | | | | |
|---|---|---|---|---|
| Variant Type | Pipeline sensitivity difference | Depth sensitivity difference | Variant types not called in PCAWG pipeline | Total expected TMB difference |
| SNV | +3% | -10% | na | -7% |
| INDEL | -43% | -2% | na | -45% |
| MNV | +5% | -15% | -11% | -21% |
| SV | -1% | -19% | -17% | -37% |

### *13. Germline predisposition variant calling*

We searched for germline variants in a broad list of 152 germline predisposition genes curated by Huang et al[24]. For SNV and INDEL, using the germline variant calling outputs from the GATK HaplotypeCaller[4], we filtered for variants affecting the canonical transcript of these 152 genes which have the following coding or splice effects:

- All SNV Nonsense, INDEL Frameshift or SNV Splice Acceptor/Donor, excluding variants marked in ClinVar[25] as 'Benign/Likely_benign', 'Benign', 'Likely_benign'.
- Missense and synonymous variants, only if marked in clinvar as ‘Pathogenic’ or ‘Likely Pathogenic’, excluding pathogenic disease indications which are clearly unrelated to cancer.

Variants which were found with a median germline VAF across all samples of less than 0.2 or greater than 0.8 were filtered as likely mapping artefacts. We further excluded frameshift variants which are found to be exactly offset by other frameshift variants (thereby creating an in-frame protein product), which actually involved more than 50% of samples in which such events occur.

This yielded 550 potential germline predisposition point mutations across the 2,399 samples in our cohort. For each variant, we determined the genotype in the germline (HET or HOM) and also assessed in the tumor sample whether there is a 2nd somatic hit, and whether the wild type or the variant itself has been lost (see chapter 13: biallelic status evaluation methods). We also searched in the 152 genes for copy number deletions that were heterozygous in the germline with subsequent homozygous loss in the tumor and found an additional 16 of such germline copy number events, giving a total of 566 variants altogether.

We observed that for the variants in many of the 152 predisposition genes that a loss of wild type in the tumor via LOH was lower than the average rate of LOH across the cohort and that fewer than 5% of observed variants had a 2nd somatic hit in the same gene. Moreover, in many of these genes the ALT variant was lost via LOH as frequently as the wild type, suggesting that a significant portion of the 566 variants may be passengers. For our downstream analysis and driver catalog, we therefore restricted our analysis to a more conservative 'High Confidence' list including only the 25 cancer related genes in the ACMG secondary findings reporting guidelines (v2.0)[26], together with 4 curated genes (CDKN2A, CHEK2, BAP1 & ATM), selected because these are the only additional genes from the larger list of 152 genes with a significantly elevated proportion of called germline variants with loss of wild type in the tumor sample.

The following table summarises the statistics for the high confidence and low confidence genes:

| Genes | Total germline predisposition SNV & INDEL | % with loss of wild type OR somatic hit in tumor | % with loss of germline ALT variant in tumor |
|---|---|---|---|
| High Confidence: ACMG + 4 curated genes | 211 | 53.1% | 10.4% |
| Low Confidence: Rest of 152 panel | 355 | 16.3% | 13.1% |

Outside the 29 high confidence genes, the germline variant itself is lost almost as frequently via LOH as the remaining wild type in the tumor, whereas for the high confidence ACMG + curated genes, there is an observed loss of wild type allele in over half of all variants.

For the additional 4 curated genes, the numbers are as follows:

| Gene | Count germline predisposition SNV & INDEL | % with loss of wild type in tumor sample | % with loss of germline variant in tumor sample |
|---|---|---|---|
| ATM | 17 | 52.9% | 11.8% |
| BAP1 | 5 | 66.7% | 0% |
| CHEK2 | 72 | 36.1% | 13.9% |
| CDKN2A | 3 | 66.7% | 33.3% |

Germline variants with loss of ALT variants in the tumor were also excluded from the final list used in our analyses, leading to a final inclusion of 189 variants from the high confidence panel.

Supplementary Table 6 contains the full catalog of high and low confidence germline variants.

### *14. Clonality and biallelic status of point mutations*

For each point mutation we determined the clonality and biallelic status by comparing the estimated ploidy of the variant to the local copy number at the exact base of the variant. The ploidy of each variant is calculated by adjusting the observed VAF by the purity and then multiplying by the local copy number to work out the absolute number of chromatids that contain the variant.

We mark a mutation as biallelic (i.e. no wild type remaining) if Variant Ploidy > Local Copy Number - 0.5. The 0.5 tolerance is used to allow for the binomial distribution of VAF measurements for each variant. For example, if the local copy number is 2 than any somatic variant with measured ploidy > 1.5 is marked as biallelic.

For each variant we also determine a probability that it is subclonal. This is achieved via a two-step process

**1. Fit the somatic ploidies for each sample into a set of clonal and subclonal peaks**
We apply an iterative algorithm to find peaks in the ploidy distribution:

- Determine the peak by finding the highest density of variants within +/- 0.1 of every 0.01 ploidy bucket.
- Sample the variants within a 0.05 ploidy range around the peak.
- For each sampled variant, use a binomial distribution to estimate the likelihood that the variant would appear in all other 0.05 ploidy buckets.
- Sum the expected variants from the peak across all ploidy buckets and subtract from the distribution.
- Repeat the process with the next peak

This process yields a set of ploidy peaks, each with a ploidy and a total density (i.e. count of variants). To avoid overfitting small amounts of noise in the distribution, we filter out any peaks that account for less than 40% of the variants in the ploidy bucket at the peak itself. After this filtering we scale the fitted peaks by a constant so that the sum of fitted peaks = the total variant count of the sample.

Finally we mark a peak as subclonal if the peak ploidy < 0.85.

**2. Calculate the probability that each individual variant belongs to each peak**
Once we have fitted the somatic ploidy peaks and determined their clonality, we can calculate the subclonal likelihood for any individual variant as the proportion of subclonal variants at that same ploidy.

The following diagram illustrates this process for a typical sample. Figure A shows the histogram of somatic ploidy for all SNV and INDEL in blue. Superimposed are four peaks in different colours fitted from the sample as described above. The red filled peak is below the 0.85 threshold and is thus considered subclonal. The black line shows the overall fitted ploidy distribution. Figure B shows the likelihood of a variant being subclonal at any given ploidy.

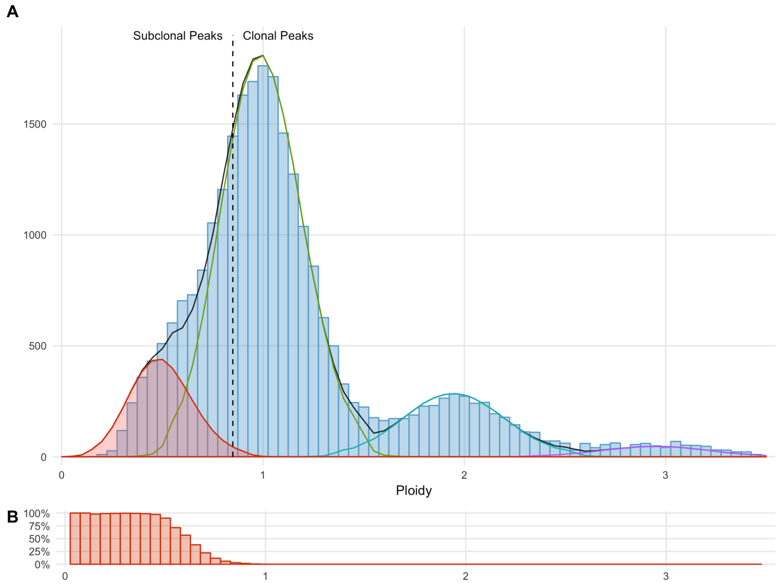


Subclonal counts in this paper are calculated as the total density of the subclonal peaks for each sample. Subclonal driver counts are calculated as the sum across the driver catalog of subclonal probability * driver likelihood (driver likelihood is explained in detail in chapter 20).

### *15. WGD status determination*

We implement a simple heuristic that determines if Whole Genome Duplication has occurred:

**Major allele Ploidy >1.5 on at least 50% of at least 11 autosomes**

The principle behind this heuristic is that if sufficient independent chromosomes are predominantly duplicated, the most parsimonious explanation is that the duplication occurred in a single genome-wide event.

The number of duplicated autosomes per sample (ie. the number of autosomes which satisfy the above rule) follows a bimodal distribution with 95% of samples have either <= 6 or > =15 autosomes duplicated. Hence, the classification of a genome as WGD is not particularly sensitive to the choice of cut-off as is evident the following chart:

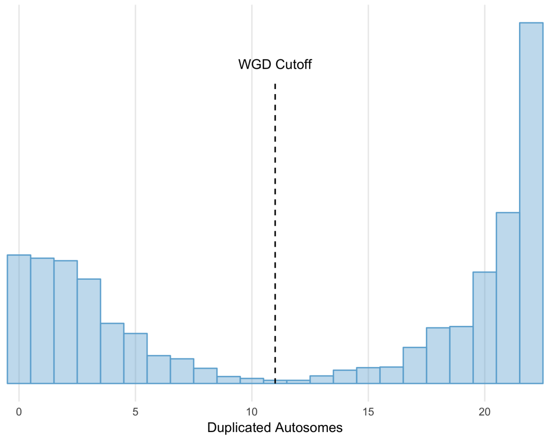

### 16. MSI status determination

To determine the MSI status of all samples we used the method described by the MSISeq tool[27]. In brief, we count the number of INDELS per million bases occuring in homopolymers of 5 or more bases or dinucleotide, trinucleotide and tetranucleotide sequences of repeat count 4 or more. MSIseq scores ranged from 0.004 up to 98.63, with a long tail towards lower MSI scores as shown in the following chart:

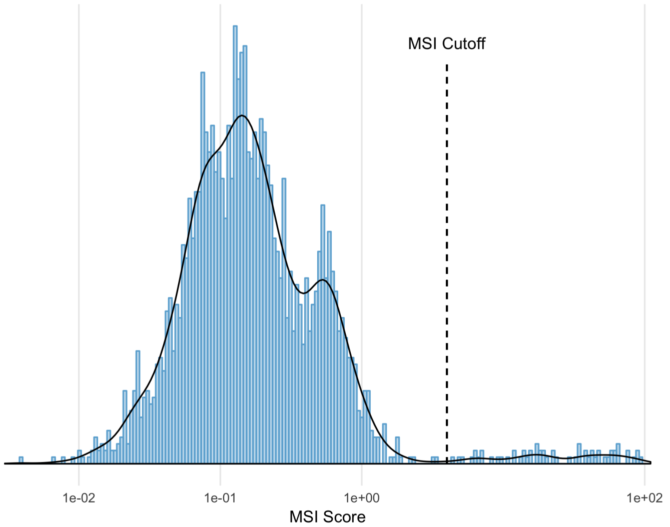


To be able to accurately set and validate the MSIseq cutoff for classification of MSI we compared the WGS results with the standard, routinely used MSI assessment using a 5-marker PCR panel (BAT25, BAT26, NR21, NR24 and MONO27 markers). For a batch of 48 pre-selected samples, the MSI PCR assay was blindly performed by an independent ISO-accredited pathology laboratory. Both the binary MSI and MSS classifications were determined, but also the number of positive markers.

A sample was considered as MSI if two or more of the five markers were score as positive (instable). PCR-based analysis identified 16 MSI samples, all of which were also identified by MSIseq with scores >4. MSIseq identified one sample that was missed by PCR-based analysis, although this sample showed microsatellite instability for one out the five markers. The MSIseq scores thus highly correlate with the number of positive MSI PCR markers and all, except one, samples with an elevated score are classified as MSI by pathology. Based on this data we determined the best cutoff for MSIseq classification to be at a **score of 4**.

Results of the PCR-based and WGS based MSI classification are summarized in the table below. The sensitivity of WGS-based MSI classification on this set was 100% (95%CI 82.6 – 100%) with a specificity of 97% (95%CI 88.2-96.9%). The calculated Cohen's kappa score was 0.954 (95%CI 0-696-0.954), indicative of a very high agreement.

| | PCR-MSS | PCR-MSI | Total |
|---|---|---|---|
| **MSISeq -MSS** | 31 | 0 | 31 |
| **MSIseq - MSI** | 1 | 16 | 17 |
| **Total** | 32 | 16 | 48 |

### 17. Holistic gene panel for driver discovery

We used Ensembl[15] release 89 as a basis for our gene definitions and have taken the union of Entrez identifiable genes and protein coding genes as our base panel.

Certain genes have multiple definitions. NPIPA7 for example has two definitions which are equally valid, ENSG00000214967 and ENSG00000183889. To solve this we select a single gene definition based on the following steps:

1) Exclude non protein coding genes.
2) Favour genes that are present in both Havana and Ensembl.
3) Select gene with longest transcript.

This returns our final gene panel tally to 25,963 genes of which 20,083 genes are protein coding. For each gene we chose the canonical transcript or the longest if no canonical transcript exists.

For CDKN2A, we included both the p16 and p14arf transcripts in the analysis given the known importance of both transcripts to tumorigenesis[28] and the fact that the two transcripts use alternate reading frames in the same exon.

### 18. Significantly mutated driver genes discovery

Using all SNV and INDEL variants from the holistic gene panel, we ran dNdScv[29] to find significantly mutated genes (SMGs) and also to estimate the proportion of missense, nonsense, essential splice site and INDEL variants which are drivers in each individual gene in the panel.

Pan cancer and at an individual cancer level we tested the normalised dNdS rates against a null hypothesis that dNdS = 1 for each variant subtype. To identify SMGs in our cohort we used a strict significance cutoff of $q<0.01$.

Two of the newly discovered SMG candidates were subsequently removed via manual curation as they were deemed to be likely artefacts of our methods:

- POM121L12 - found only to be significant due to an extreme covariate value in dNdScv
- TRIM49B - found to have poor mappability on nearly all its variants and a known close paralog

### 19. Significantly amplified & deleted driver gene discovery

To search for significantly amplified and deleted genes we first calculated the minimum exonic copy number per gene across our holistic gene panel. For amplifications, we searched for all the genes with high level amplifications only (defined as minimum Exonic Copy number > 3 * sample ploidy). For deletions, we searched for all the genes in each sample with either full or partial gene homozygous deletions (defined as minimum exonic copy number < 0.5). The Y chromosome was excluded from the deletion analysis since the Y chromosome is deleted altogether in 35% of all male cancer samples in our cohort and hence is difficult to distinguish at the gene level.

We then searched separately for amplifications and deletions, on a per chromosome basis, for the most significant focal peaks, using an iterative GISTIC-like peel off method[30], specifically:

- Find the highest scoring gene.
    - For deletions the score is simply the count of samples with homozygous deletions in the gene.
    - For amplifications, we need to consider both the count and strength of the amplification so we use:

    - score = sum(log2(copy number/ sample ploidy)).
- Record gene as a peak, and mark all consecutive genes with a score within 15% and 25% of the highest score for deletions and amplifications respectively as part of the candidate peak.
- 'Peel' off all samples which contributed to the peak across the entire chromosome.
- Repeat the process.

A filter was applied where we removed deletions from a handful of noisy copy number regions in the genome where we found more than 50% of the observed deletions were not supported on either breakend by a structural variant.

Most of the deletion peaks resolve clearly to a single target gene reflecting the fact that homozygous deletions are highly focal, but for amplifications this is not the case and the majority of our peaks have 10 or more candidates. We therefore annotated the peaks, to choose a single putative target gene using an objective set of automated curation rules in order of precedence:

- If more than 50% of the copy number events in the peeled samples include the telomere or centromere than mark as <CHR>_<ARM>_<TELOMERE/CENTROMERE>
- Else choose highest scoring candidate gene which matches a list of actionable amplifications from OncoKB, CGI and CIViC clinical annotation DBs
- Else choose highest scoring candidate gene found in our panel of significantly mutated genes.
- Else choose highest scoring candidate gene found in cosmic census
- Else choose highest scoring protein coding candidate gene
- Else choose longest non-coding candidate gene

Finally, we filter the peaks to only highly significant deletions and amplifications using the following rules

- Deletions => Keep any peak with > 5 homozygous deletions
- Amplifications => Keep any peak with score > 29

These cut-offs were chosen using a binomial model which assumes the probability of any given gene being observed to be randomly deleted or highly amplified is equal to the average number of genes amplified or deleted in each event divided by the total number of genes considered. The cut-offs were chosen to be the lowest score with a q-value below 0.25. Since amplifications are generally much broader (averaged genes affected per event of 41.6 compared to just 5.4 for deletions) a much higher number of genes is required to reach significance.

The calculation details for the cut-offs are presented in the table below.

| | Cohort data | | | | | | Statistical Calculations | | | | |
|---|---|---|---|---|---|---|---|---|---|---|---|
| | Count of events | Sum Scores | Count of genes affected | Avg genes affected per event | Avg score / event | Total genes tested | Probability event overlaps a given gene | Score cutoff | P value of cutoff | Significant findings | Q Value |
| Dels | 4,915 | 4,915 | 26,676 | 5.4 | 1.0 | 25,965 | 0.00021 | 5 | 0.00068 | 117 | 0.15 |
| Amps | 3,925 | 6,959 | 163,393 | 41.6 | 1.8 | 25,965 | 0.00160 | 29 | 0.00030 | 33 | 0.23 |

This model is likely to be highly conservative as it assumes that all the events are passengers, whereas in fact a high proportion contain driver genes.

### 20. Fragile site annotation

Homozygous deletions were also annotated as common fragile site (CFS) based on their genomic characteristics. This annotation is not definitive, but is useful as CFS are known to be regions of high genomic instability. Hence despite being significantly deleted, their status as a genuine cancer driver remains unclear.

There is no absolute agreement on which regions should be classified as CFS, but two well-known features are a strong enrichment in long genes and a high rate of observed deletions of up to 1 megabase[31]. Hence for this analysis we classified a gene as a fragile site if it met all the following criteria:

- Total length of gene > 500,000 bases
- More than 30% of all SVs with breakpoints that disrupt the gene are deletions with length greater than 20,000 bases and less than 1 megabase.
- The gene is not found to be significantly mutated (by dNdScv) in our cohort or in Martincorena et al.[29].

Using these criteria we annotated the following list of 16 Genes as fragile:

| Gene | Chr | Start position | Length (bases) | Total Disruptive SV Count | % of SV that are DELs (>20kb & <1MB) |
|---|---|---|---|---|---|
| LRP1B | 2 | 140,988,992 | 1,900,278 | 1,272 | 0.469 |
| FHIT | 3 | 59,735,036 | 1,502,097 | 2,128 | 0.596 |
| LSAMP | 3 | 115,521,235 | 2,194,860 | 1,306 | 0.364 |
| NAALADL2 | 3 | 174,156,363 | 1,367,065 | 1,198 | 0.456 |
| CCSER1 | 4 | 91,048,686 | 1,474,378 | 1,398 | 0.441 |
| PDE4D | 5 | 58,264,865 | 1,553,082 | 1,166 | 0.458 |
| GMDS | 6 | 1,624,041 | 621,885 | 399 | 0.441 |
| PARK2 | 6 | 161,768,452 | 1,380,351 | 1,296 | 0.555 |
| IMMP2L | 7 | 110,303,110 | 899,463 | 1,028 | 0.444 |
| PTPRD | 9 | 8,314,246 | 2,298,477 | 1,264 | 0.309 |
| PRKG1 | 10 | 52,750,945 | 1,307,165 | 781 | 0.318 |
| GPHN | 14 | 66,974,125 | 674,395 | 291 | 0.306 |
| WWOX | 16 | 78,133,310 | 1,113,254 | 1,319 | 0.541 |
| MACROD2 | 20 | 13,976,015 | 2,057,827 | 3,039 | 0.605 |

| | | | | | |
|---|---|---|---|---|---|
| DMD | X | 31,115,794 | 2,241,764 | 789 | 0.328 |
| DIAPH2 | X | 95,939,662 | 920,334 | 331 | 0.381 |

We also noted that 4 other significantly deleted genes (STS,HDHD1,LRRN3 and LINC00290), though not fulfilling the length criteria above have a particularly high proportion of deletion SVs between 20kb and 1 megabase (over 60%) and hence were also marked as fragile:

| Gene | Chr | Start position | Length (bases) | Total Disruptive SV Count | % of SV that are DELs (>20kb & <1MB) |
|---|---|---|---|---|---|
| LINC00290 | 4 | 181,985,242 | 95,060 | 64 | 0.641 |
| LRRN3 | 7 | 110,731,062 | 34,448 | 70 | 0.686 |
| STS | X | 7,137,497 | 135,354 | 168 | 0.649 |
| HDHD1 | X | 6,966,961 | 99,270 | 126 | 0.659 |

Two of these genes (STS and HDHD1) fall in a previously identified CFS region (FRAXB) and a third, LRNN3, falls in another knowns CFS region (FRAX7). The final one, LINC00290 is a long non-coding RNA with an unknown status as cancer driver.

### *21. Somatic driver catalog construction*

We created a catalog of each and every driver in our cohort across all variant types on a per patient basis. This was done in a similar incremental manner to Sabarinathan et al[32] (N. Lopez, personal communication) whereby we first calculated the number of drivers in a broad panel of known and significantly mutated genes across the full cohort, and then assigned the drivers for each gene to individual patients by ranking and prioritising each of the observed variants. Key points of difference in this study were both the prioritisation mechanism used and our choice to ascribe each mutation a probability of being a driver rather than a binary cutoff based on absolute ranking.

The four detailed steps to create the catalog are described below:

**1. Create a panel of driver genes for point mutations using significantly mutated genes and known drivers**

We created a gene panel using the union of

- Martincorena significantly mutated genes[29] (filtered to significance of q<0.01)
- HMF significantly mutated genes (q<0.01) at global level or at cancer type level
- Cosmic Curated Genes[12] (v83)

**2. Determine TSG or Oncogene status of each significantly mutated gene**

We used a logistic regression model to classify the genes in our pane as either tumor suppressor gene (TSG) or oncogene. We trained the model using unambiguous classifications from the Comic curated genes, i.e. a gene was considered either a Oncogene or TSG but not both. We determined that the dNdS missense and nonsense ratios (w_missense and w_nonsense) are both significant predictors of the classification. The coefficients are given in the table below.

| | Estimate | Std. Error | z value | Pr(>\|z\|) |
|---|---|---|---|---|

| | | | | |
|---|---|---|---|---|
| intercept | 0.1830 | 0.3926 | 0.466 | 0.64106 |
| w_missense | -0.6869 | 0.2643 | -2.599 | 0.00936 |
| w_nonsense | 0.5237 | 0.1116 | 4.691 | 2.72e-06 |

We applied the model to all significantly mutated genes in Matincorena and HMF as well as any ambiguous Cosmic curated genes.

The following figure shows all genes that have classified using the logistic regression model. Figures A and C show the likelihood of a gene being classified as a TSG under a single variate logistic model of w_missense and w_nonsense respectively. Figure B shows the classification after the multivariate regression using both predictors.

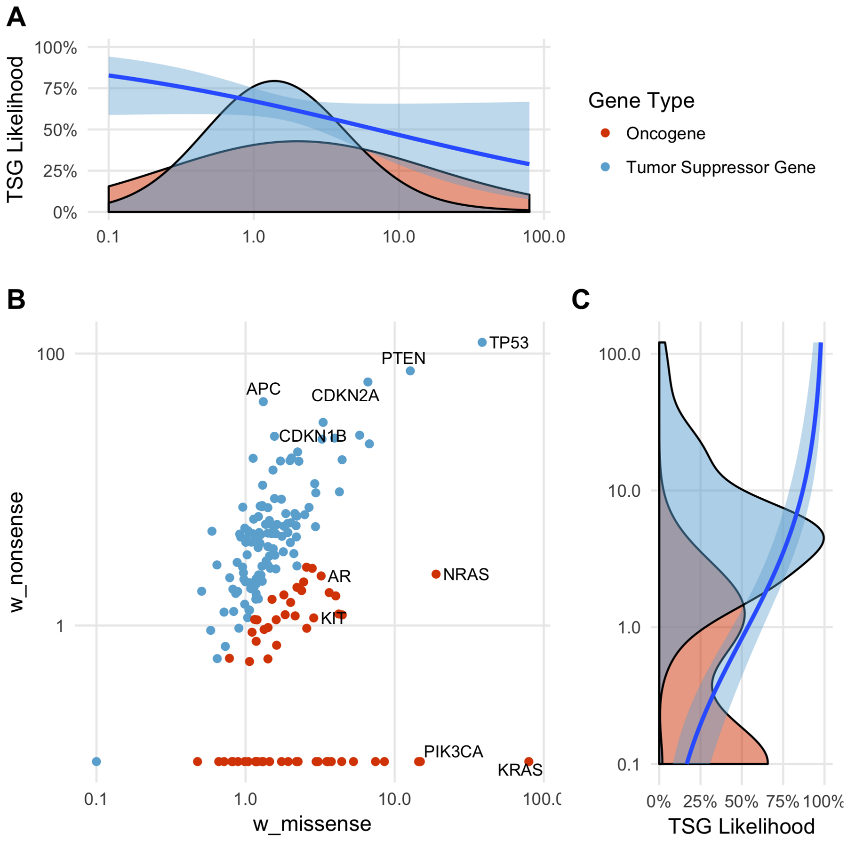


**3. Add drivers from all variant classes to the catalog**
Variants were added to the driver catalog which met any of the following criteria

- All missense and inframe indels for panel oncogenes
- All non synonymous and essential splice point mutations for tumor suppressor genes
- All high level amplifications (min exonic copy number > 3 * sample ploidy) for both significantly amplified target genes and panel oncogenes
- All homozygous deletions for significantly deleted target genes and panel TSG (except for the Y chromosome as described before)
- All known or promiscuous inframe gene fusions as described above
- Recurrent TERT promoter mutations

**4. Calculate a per sample driver likelihood for each gene in the catalog**
A driver likelihood estimate between 0 and 1 was calculated for each variant in the gene panel to ensure that only excess mutations are used for determining the number of drivers in cancer cohort groups or at the individual sample level. High level amplifications, Deletions, Fusions, and TERT promoter mutations are all rare so were assumed to have a likelihood of 1 when found affecting a driver gene, but for coding mutations we need to account for the large number of passenger point mutations that are present throughout the genome and thus also in driver genes.

For coding mutations we also marked coding mutations that are highly likely to be drivers and/or highly unlikely to have occurred as passengers as driver likelihood of 1, specifically:

- Known hotspot variants
- Variants within 5 bases of a known pathogenic hotspot in oncogenes
- Inframe indels in oncogenes with repeat count < 8 repeats. Longer repeat count contexts are excluded as these are often mutated by chance in MSI samples
- Biallelic variants in tumor suppressor genes

For the remaining variants (non-hotspot missense variants in oncogenes and non-biallelic variants in TSG) these were only assigned a > 0 driver likelihood where there was a remaining excess of unallocated drivers based on the calculated dNdS rates in that gene across the cohort after applying the above rules. Any remaining point mutations were assigned a driver likelihood between 0 and 1 using a bayesian statistic to calculate a sample specific likelihood of each gene based on the type of variant observed (missense, nonsense, splice or INDEL) and taking into account the mutational load of the sample. The principle behind the method is that the likelihood of a passenger variant occuring in a particular sample should be approximately proportional to the tumor mutational burden and hence variants in samples with lower mutational burden are more likely to be drivers.

The sample specific likelihood of a residual excess variant being a driver is estimated for each gene using the following formula:

P(Driver|Variant) = P(Driver) / (P(Driver) + P(Variant|Non-Driver) * (1-P(Driver)))

where P(Driver) in a given gene is assumed to be equal across all samples in the cohort, ie:

P(Driver) = (residual unallocated drivers in gene) / # of samples in cohort

And P(Variant|Non-Driver), the probability of observing n or more passenger variants of a particular variant type in a sample in a given gene, is assumed to vary according to tumor mutational burden, and is modelled as a poisson process:

P(Variant|Non-Driver) = 1 - poisson(λ = TMB(Sample) / TMB(Cohort) * (# of passenger variants in cohort),k=n-1)

All counts reported in the paper at a per cancer type or sample level refer to the sum of driver likelihoods for that cancer type or sample.

### *22. Driver co-occurrence analysis*

To examine the co-occurence of drivers, the driver-gene catalog was filtered to exclude fusions and any driver with a driver likelihood of < 0.5. Separately for each cancer type, every pair of driver genes was tested to see whether they co-occur more or less frequently than expected if they were independent using Fisher's Exact Test. The results were adjusted to a FDR using the number of gene-pair comparison being tested in each cancer type cohort. Gene pairs with a positive correlation which were on the same chromosome were excluded from the analysis as they are frequently co-amplified or deleted by chance.

### *23. Actionability analysis*

To determine clinical actionability of the variants observed in each sample, we mapped all variants to 3 external clinical annotation databases

- OncoKB[9] (download = 01-mar-2018)
- CGI[8] (update: 17-jan-2018)
- CIViC[7] (download = 01-mar-2018)

In order to be able to aggregate and compare this data, we have mapped each of the databases to a common data model using the following rules:

**1. Level of evidence mapping**

The 3 databases we used in this study define different level for evidence items, depending on evidence strength. In order to be able to aggregate and compare this data, we have mapped the CGI and OncoKB evidence levels on the CIViC evidence levels defined at: https://civicdb.org/help/evidence/evidence-levels.

| HMF | CIViC | CGI | OncoKB |
|---|---|---|---|
| A | A | FDA guidelines,<br>NCCN guidelines, NCCN/CAP guidelines, CPIC guidelines, European Leukemia Net guideline | 1<br>2<br>R1 |
| B | B | Clinical trials,<br>Late trials,<br>Late trials,Pre-clinical | 3<br>R2 |
| C | C | Early trials,<br>Case report | |
| D | D | Pre-clinical | 4,R3 |

In this study we considered only A and B level variants. This classification roughly corresponds to the recently proposed ESMO Scale for Clinical Actionability of molecular Targets (ESCAT)[33] as follows:
HMF A: ESCAT I-A+B (for on label) and I-C (for off-label)
HMF B: ESCAT II-A+B (for on label) and III-A (for off-label)

**2. Response type Mapping**

We also mapped response type to a common data model. First we filtered out evidence items from the annotation databases that do not lead to clinical actionability (for example prognostic biomarkers). The remaining evidence items were mapped as either responsive or resistant based on the following rules:

| HMF | CIViC | CGI | OncoKB |
|---|---|---|---|
| Responsive | Sensitivity | Responsive | 1<br>2 |

| | | | |
|---|---|---|---|
| | | | 3<br>4 |
| Resistant | Resistant or Non-Response | Resistant | R1<br>R2<br>R3 |

**3. Mutation/Event type mapping**

Each evidence item was mapped to HMF data as one of 4 event types according to the following criteria:

| HMF Event type | Matching Criteria |
|---|---|
| Somatic Point Mutation | HGVS / genomic coordinates converted to chromosome, position, ref and alt and mapped to exact variants in our database |
| Somatic Range Event | Matched to missense / inframe variants in Oncogenes and any non-synonymous variant in TSG contained within a defined range, either exon level, transcript level or specific coordinates. Where a transcript was not specified, the canonical transcript was always used to map coordinates |
| Somatic CNA | 'Deletion' mapped to homozygous deletions and 'Amplification' mapped to high level amplification (>3x sample ploidy) |
| Fusion | Exact matching to an inframe fusion in our database. For OncoKB 'loss-of-function' fusions were excluded |

A small number of items from CIViC level B evidence level were deemed either not specific enough or insufficiently supportive of actionability for this study and were filtered:

- Evidence items supporting TP53, KRAS & PTEN as actionable
- Evidence items supporting actionability with 'chemotherapy' (ie. chemotherapy in general rather than a specific treatment), 'aspirin' or 'steroids'

Finally, a number of suspicious fusions from each of the databases were curated by either changing the 5' and 3' partners or filtered out altogether based on referring to the original evidence sources, specifically:

| HMF Curation | CIViC | CGI | OncoKB |
|---|---|---|---|
| Filtered Fusions | BRAF - CUL1 | RET - TPCN1 | |
| 5' and 3' partners exchanged | | ABL1 - BCR<br>PDGFRA - FIP1L1<br>PDGFB - COL1A1 | ROS1 - CD74<br>EP300 - MLL<br>EP300 - MOZ<br>RET - CCDC6 |

Some of the more complex event types from the 3 databases have not been fully interpreted and have been excluded from this analysis.

#### 4. Cancer type mapping

Each evidence event mapped was also determined to be either on-label (ie. evidence supports treatment in that specific cancer type) or off-label (evidence exists in another cancer type) for each specific sample. To do this, we have annotated both the patient cancer types and the database cancer types with relevant DOIDs, using the disease ontology database available at: http://disease-ontology.org.

Patient cancer types from the HMF database were annotated according to the following table:

| HMF tumor type | DOID |
|---|---|
| Biliary | 4607 |
| Bone/Soft tissue | 201;9253 |
| Breast | 1612 |
| CNS | 3620;3070 |
| Colon/Rectum | 9256;219 |
| CUP | - |
| Esophagus | 5041;4944 |
| Head and neck | 11934;8618 |
| Kidney | 263;8411 |
| Liver | 3571 |
| Lung | 1324 |
| Mesothelioma | 1790 |
| NET | - |
| Other | - |
| Ovary | 2394 |
| Pancreas | 1793 |
| Prostate | 10283 |
| Skin | 4159 |
| Stomach | 10534 |
| Urinary tract | 3996 |
| Uterus | 363 |

Database cancer types were mapped to a DOID by automatically querying the ontology on the disease names. Some CIViC evidence items are already annotated with a DOID in the database, this was used if present. We also manually annotated with DOIDs some of the database cancer types that failed the automatic query:

| cancerType | DOID | Ontology term |
|---|---|---|
| All Tumors | 162 | cancer |
| Any cancer type | 162 | cancer |
| B cell lymphoma | 707 | B-cell lymphoma |
| Billiary tract | 4607 | biliary tract cancer |
| Bladder | 11054 | urinary bladder cancer |

| | | |
|---|---|---|
| Cervix | 4362 | cervical cancer |
| CNS Cancer | 3620 | central nervous system cancer |
| Dedifferentiated Liposarcoma | 3382 | liposarcoma |
| Endometrium | 1380 | endometrial cancer |
| Esophagogastric Cancer | 5041 | esophageal cancer |
| Gastrointestinal stromal | 9253 | gastrointestinal stromal tumor |
| Giant cell astrocytoma | 3069 | astrocytoma |
| Hairy-Cell leukemia | 285 | hairy cell leukemia |
| Head and neck | 11934 | head and neck cancer |
| Head and neck squamous | 5520 | head and neck squamous cell carcinoma |
| Hepatic carcinoma | 686 | liver carcinoma |
| Hepatocellular Mixed Fibrolamellar Carcinoma | 0080182 | mixed fibrolamellar hepatocellular carcinoma |
| Inflammatory myofibroblastic | 0050905 | inflammatory myofibroblastic tumor |
| Lung | 1324 | lung cancer |
| Lung squamous cell | 3907 | lung squamous cell carcinoma |
| Melanoma | 8923 | Skin melanoma |
| Mesothelioma | 1790 | malignant mesothelioma |
| Neuroendocrine | 169 | neuroendocrine tumor |
| Non-small cell lung | 3908 | non-small cell lung carcinoma |
| Ovary | 2394 | ovarian cancer |
| Pancreas | 1793 | pancreatic cancer |
| Renal | 263 | kidney cancer |
| Salivary glands | 8850 | salivary gland cancer |
| Stomach | 10534 | stomach cancer |
| Thymic | 3277 | thymus cancer |
| Thyroid | 1781 | thyroid cancer |
| Well-Differentiated Liposarcoma | 3382 | liposarcoma |

In case a matching DOID was found for the disease, we annotated the disease with a DOID set consisting of: the disease DOID, all the children DOIDs and all the parent disease DOIDs.

A treatment is defined as on-label if any of the DOIDs of the patient cancer is present in the DOID set of the disease.

**5. MSI actionability**

Samples classified as MSI in our driver catalog were also mapped as actionable at level A evidence based on clinical annotation in the OncoKb database

**6. Aggregation of evidence**

For each actionable mutation in each sample, we aggregated all the mapped evidence that was available supporting both on-label and off-label treatments at an A or B evidence level. Treatments that also had evidence supporting resistance based on other biomarkers in the sample at the same or higher level were excluded as non-actionable.

For each sample we reported the highest level of actionability, ranked first by evidence level and then by on-label vs off-label.

### 24. Data availability

All data described in this study is freely available for academic use from the Hartwig Medical Foundation through standardized procedures and request forms which can be found at https://www.hartwigmedicalfoundation.nl/en. Briefly, a data request can be initiated by filling out the standard form in which intended use of the requested data is motivated. First, an advice on scientific feasibility and validity is obtained from experts in the field which is used as input by an independent Data Access Board who also evaluates if the intended use of the data is compatible with the consent given by the patients and if there would be any applicable legal or ethical constraints. Upon formal approval by the Data Access Board, a standard license agreement which does not have any restrictions regarding Intellectual Property resulting from the data analysis needs to be signed by an official organisation representative before access to the data is granted. Raw data files will be made available through a dedicated download portal with two-factor authentication.
