## Supplementary Image 1 for "Pan-cancer whole genome analyses of metastatic solid tumors"

Biliary

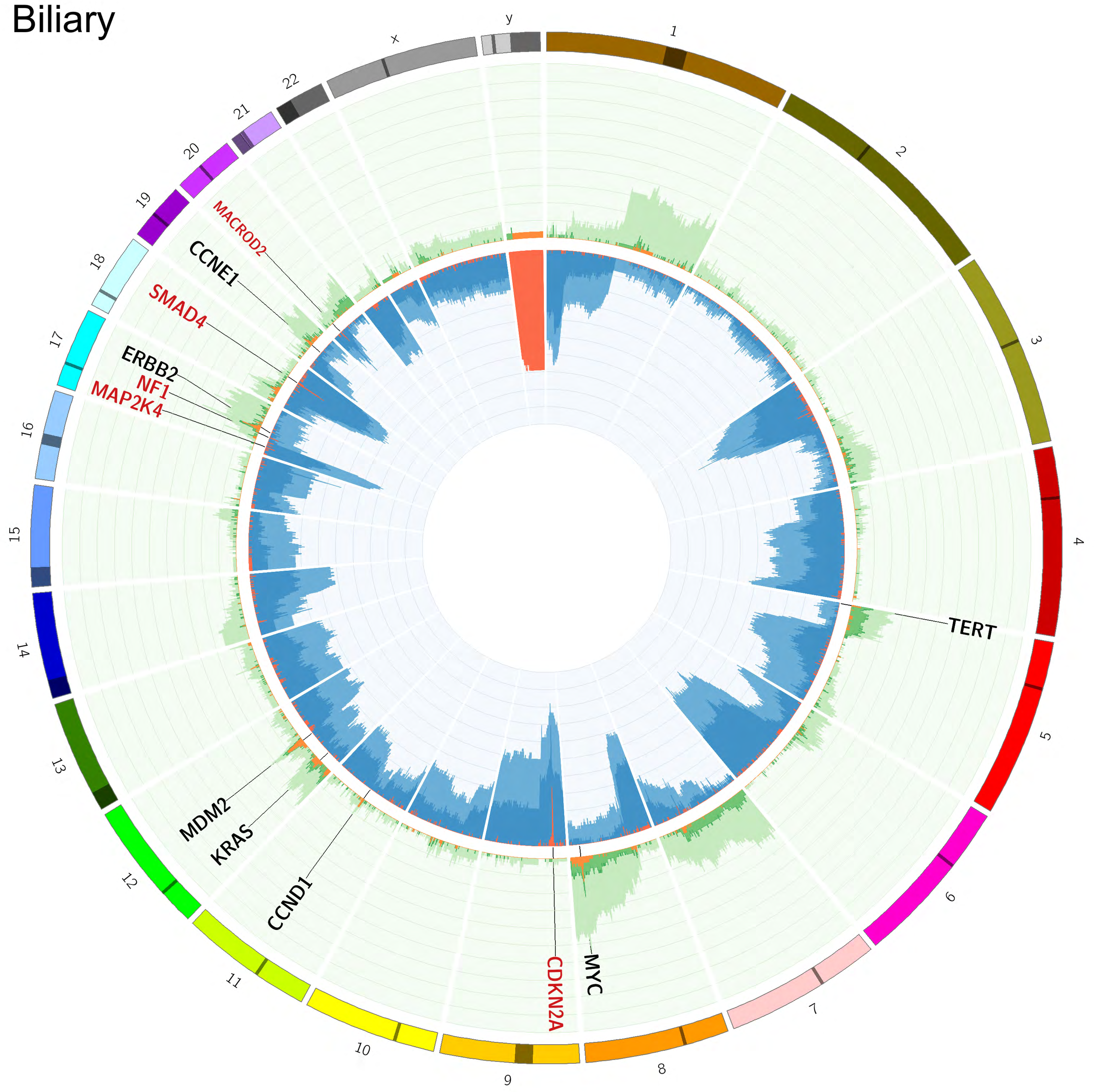

Bone/Soft tissue

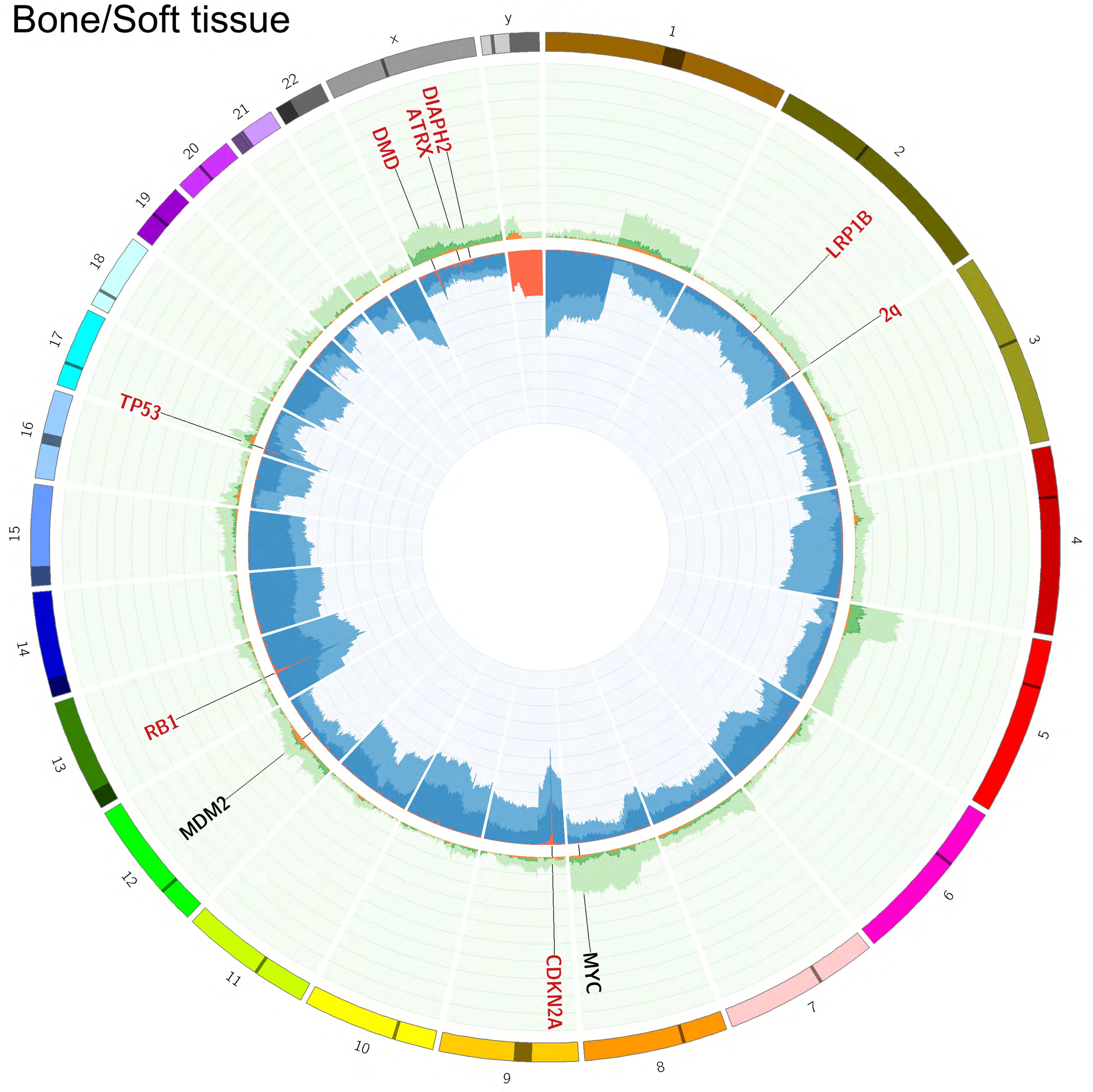

### Breast

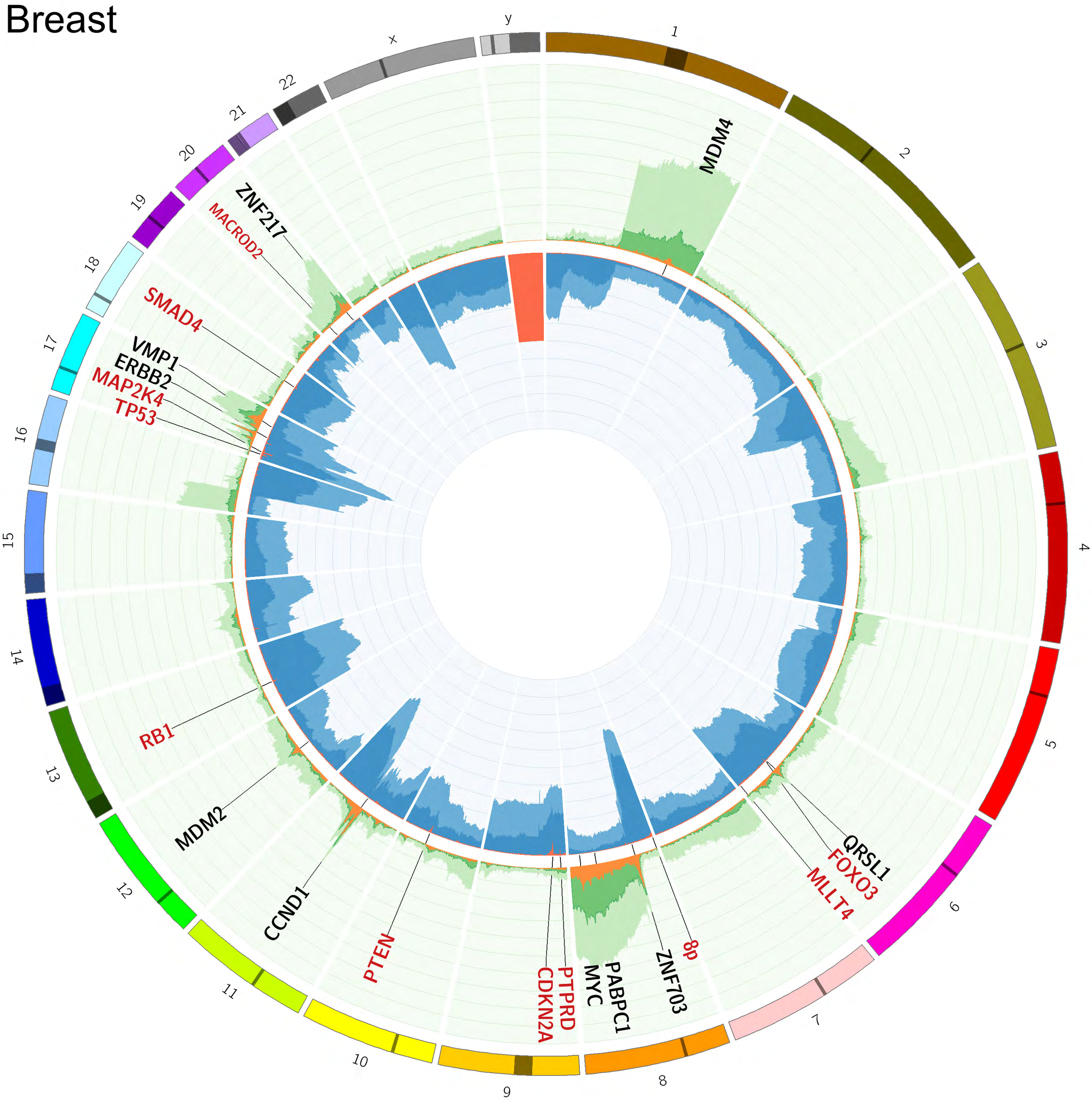

y
1
x
22
21
20
19
18
17
16
15
14
13
12
11
10
9
8
7
6
5
4
3
2
MDM4
ZNF217
MACROD2
SMAD4
VMP1
ERBB2
MAP2K4
TP53
RB1
MDM2
CCND1
PTEN
CDKN2A
PTPRD
MYC
PABPC1
ZNF703
8p
MLLT4
FOXO3
QRSL1

### CNS

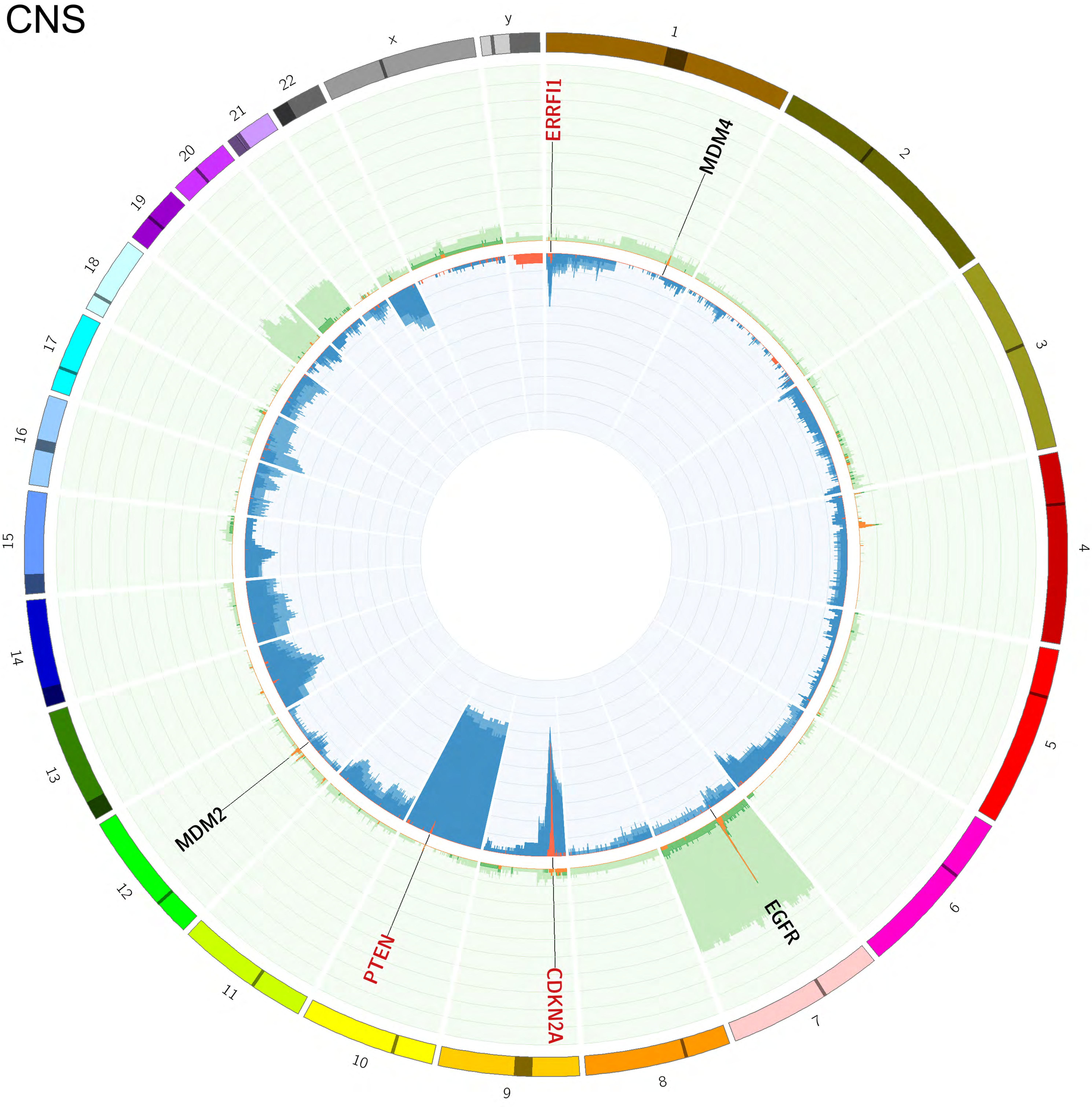

y
1
x
22
21
20
19
18
17
16
15
14
13
12
11
10
9
8
7
6
5
4
3
2
ERRFI1
MDM4
MDM2
PTEN
CDKN2A
EGFR

### Colon/Rectum

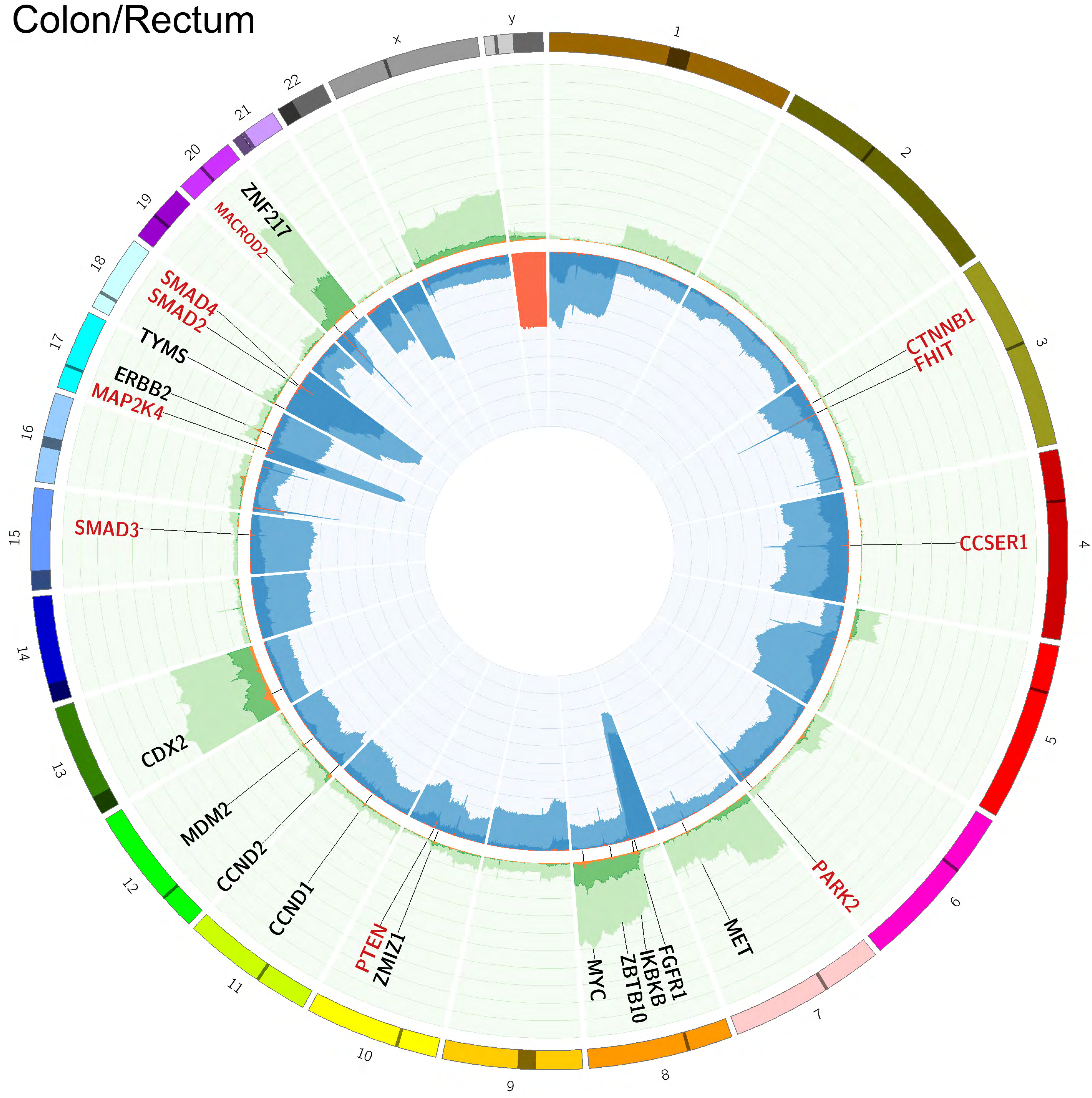

### CUP

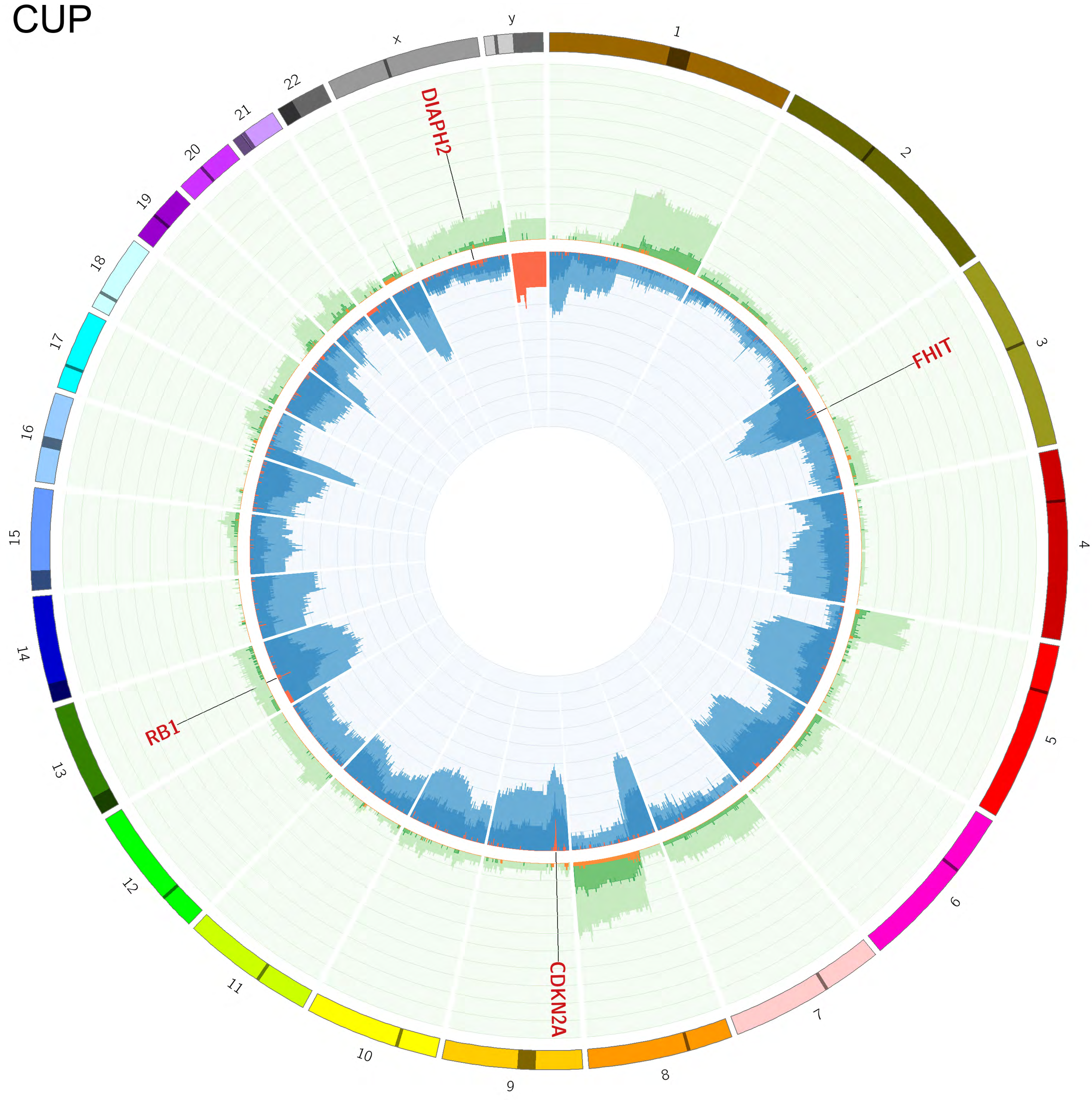

### Esophagus

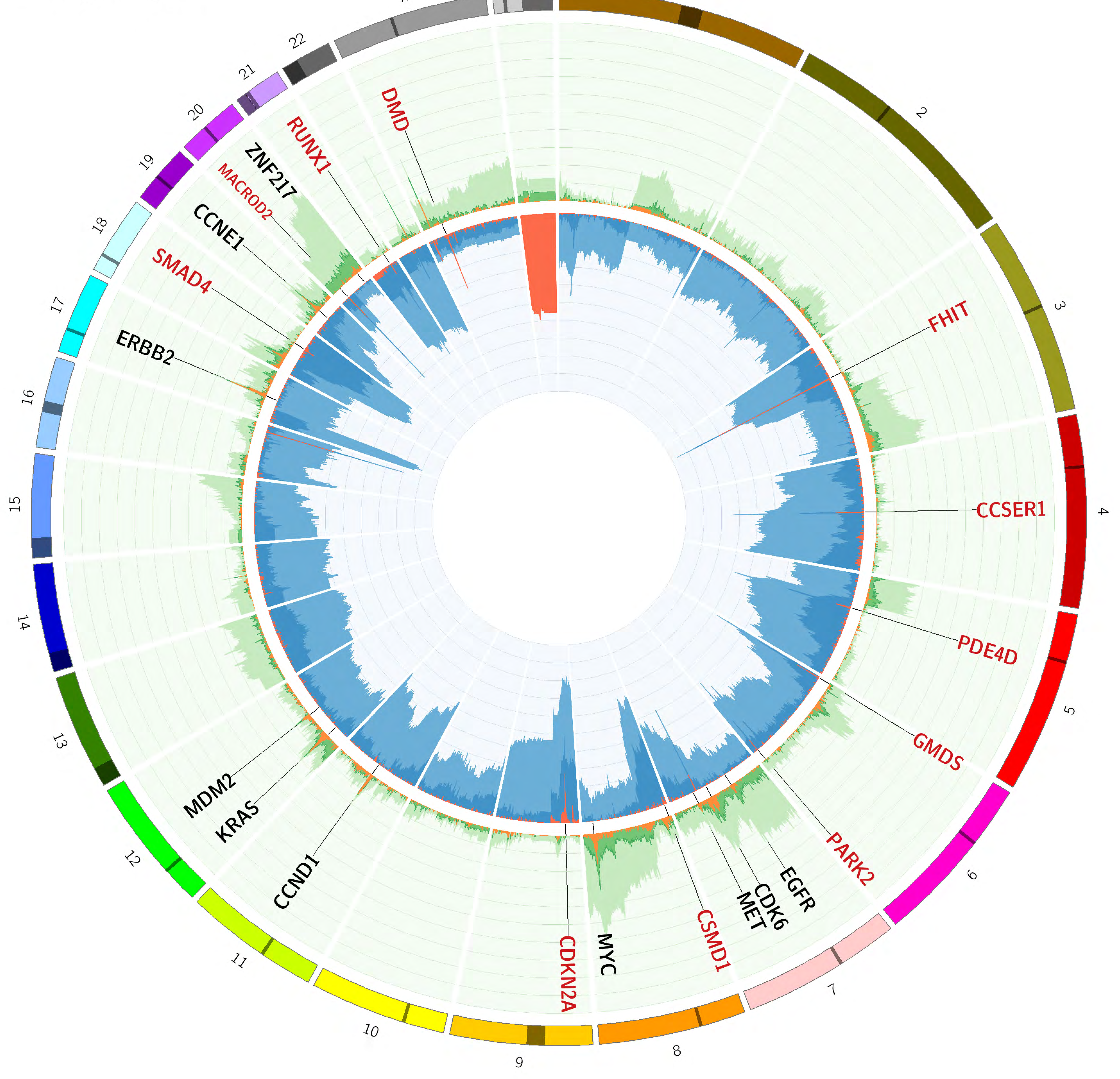

### Head and neck

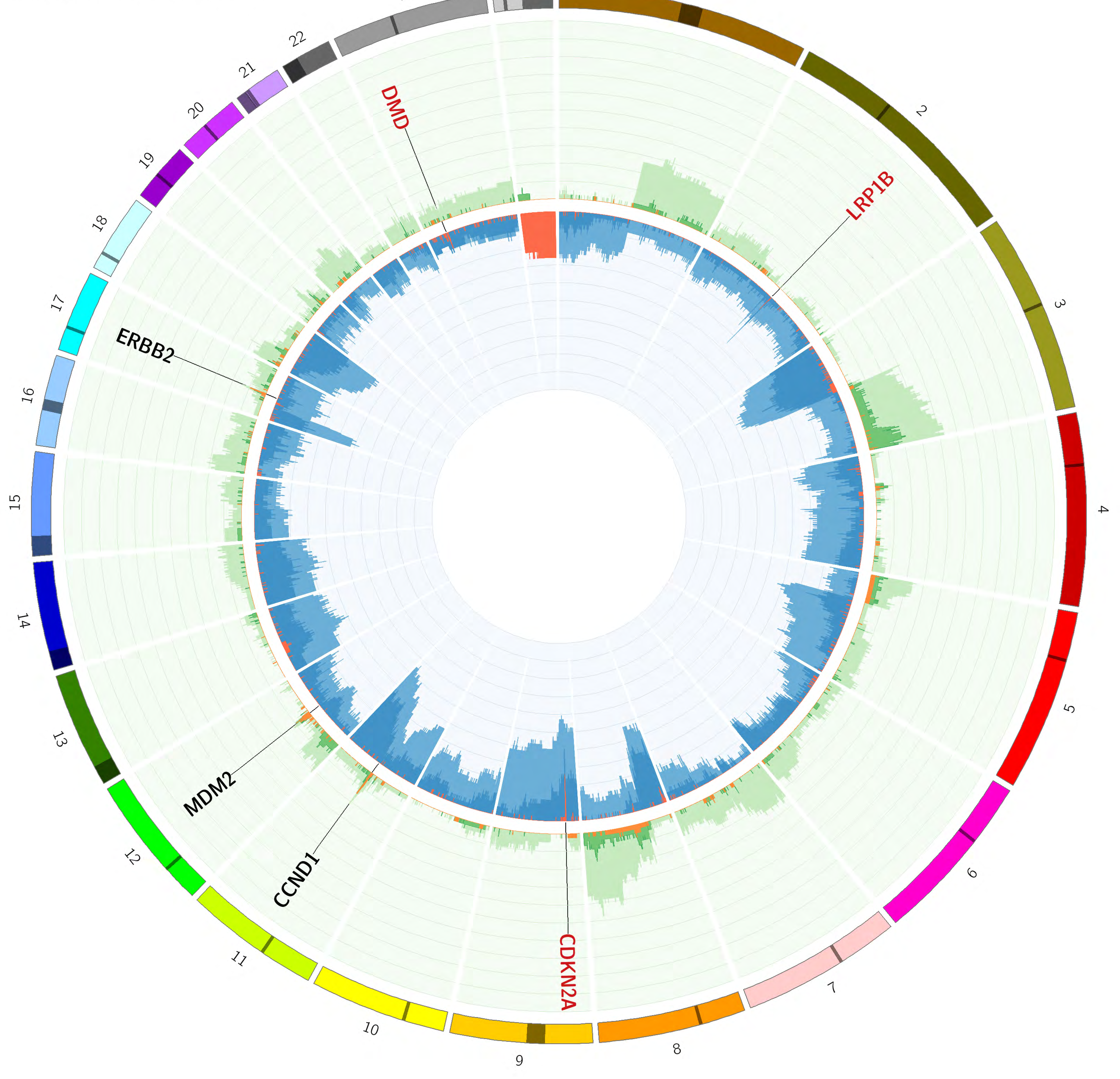

Kidney

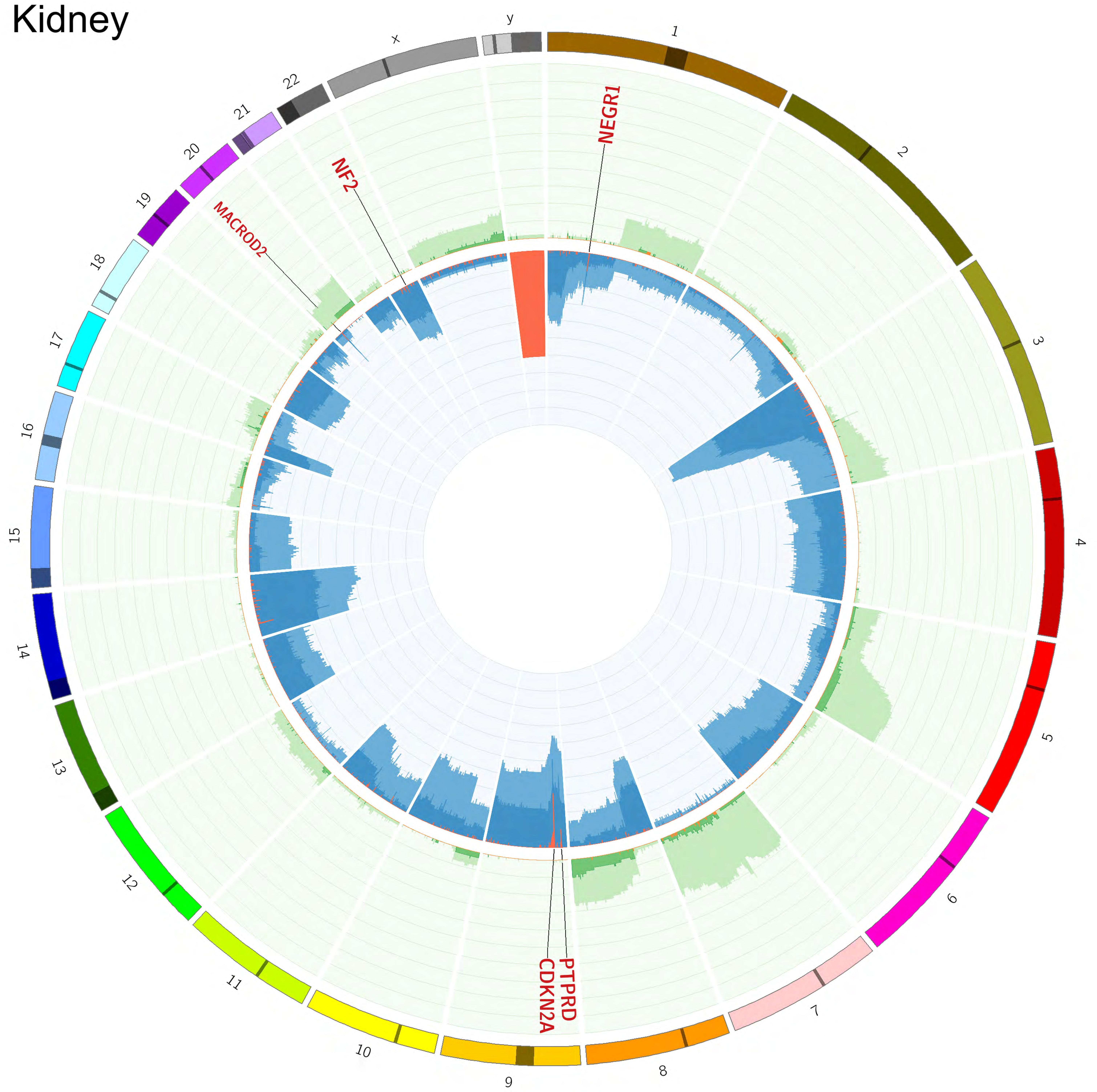

Liver

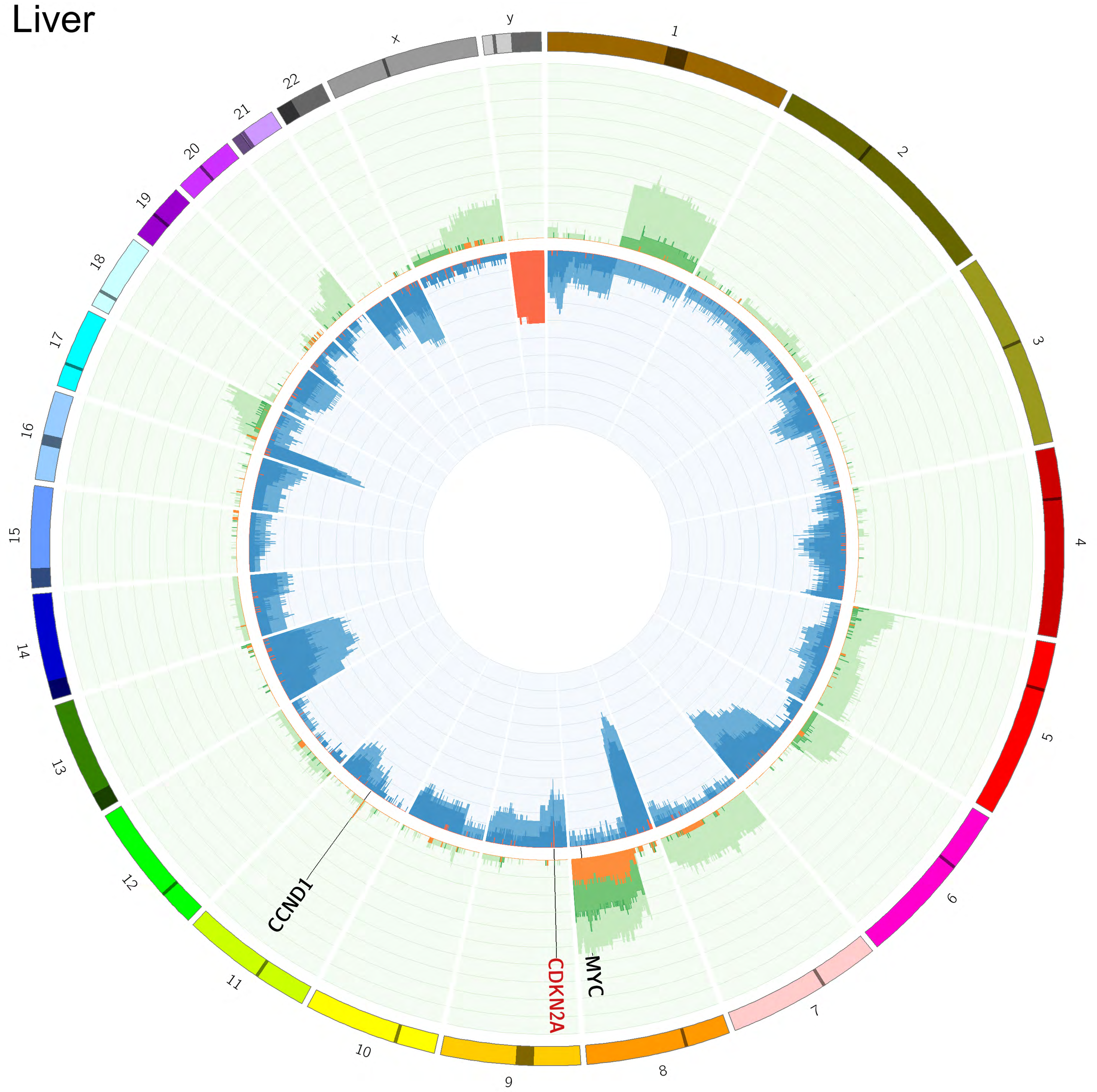

Lung

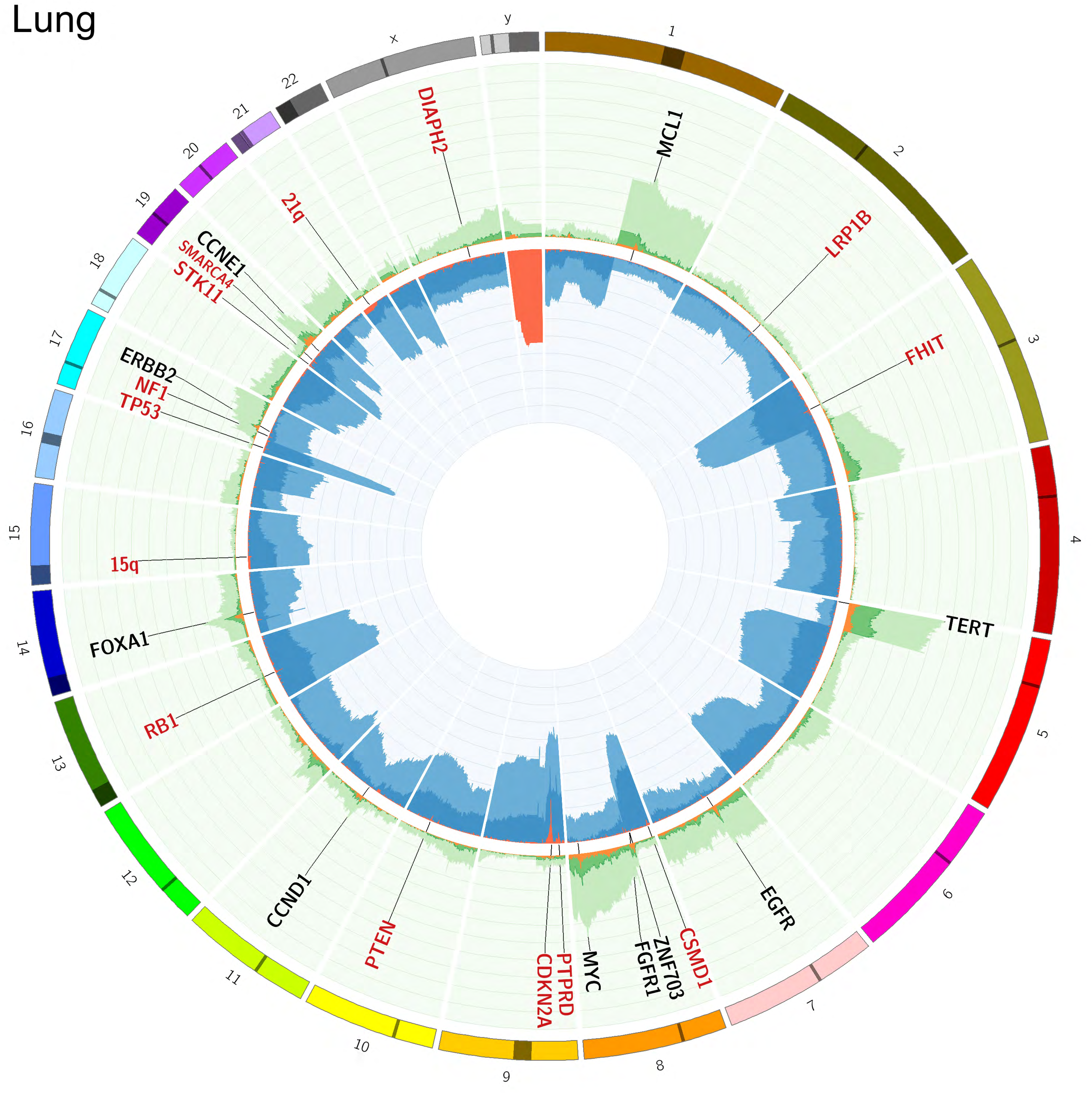

Mesothelioma

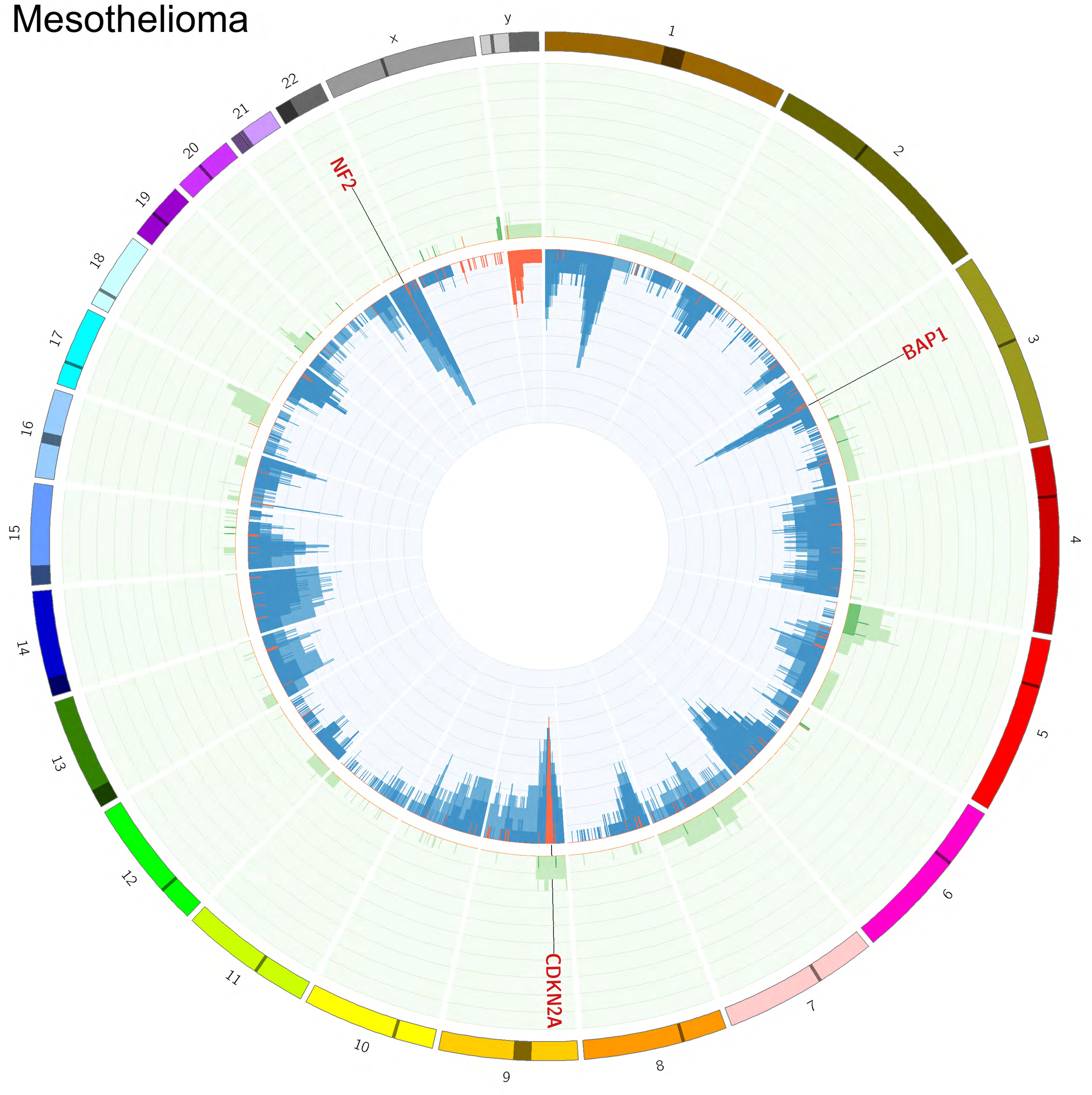

NET

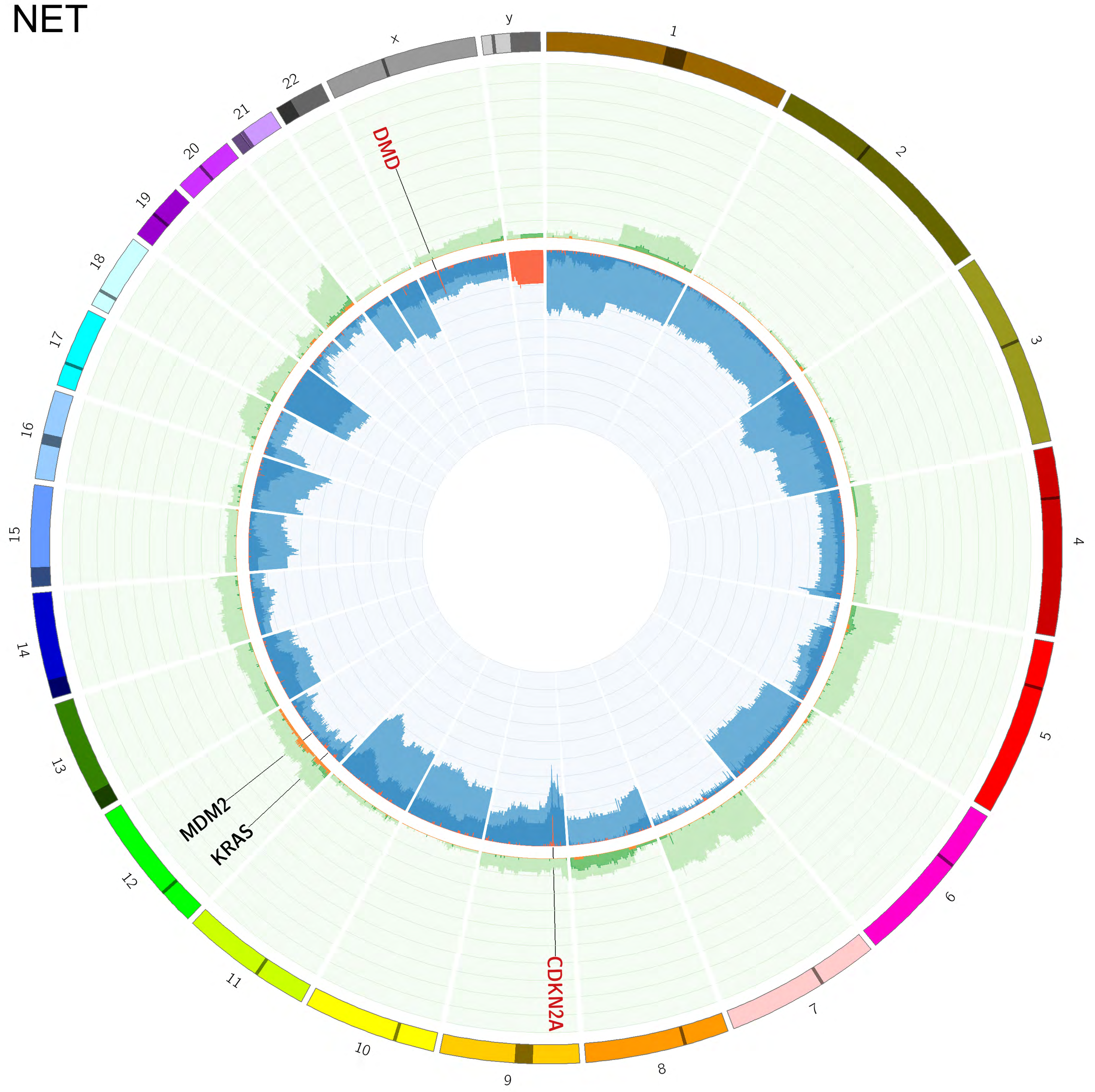

x
y
DMD
MDM2
KRAS
CDKN2A

Ovary

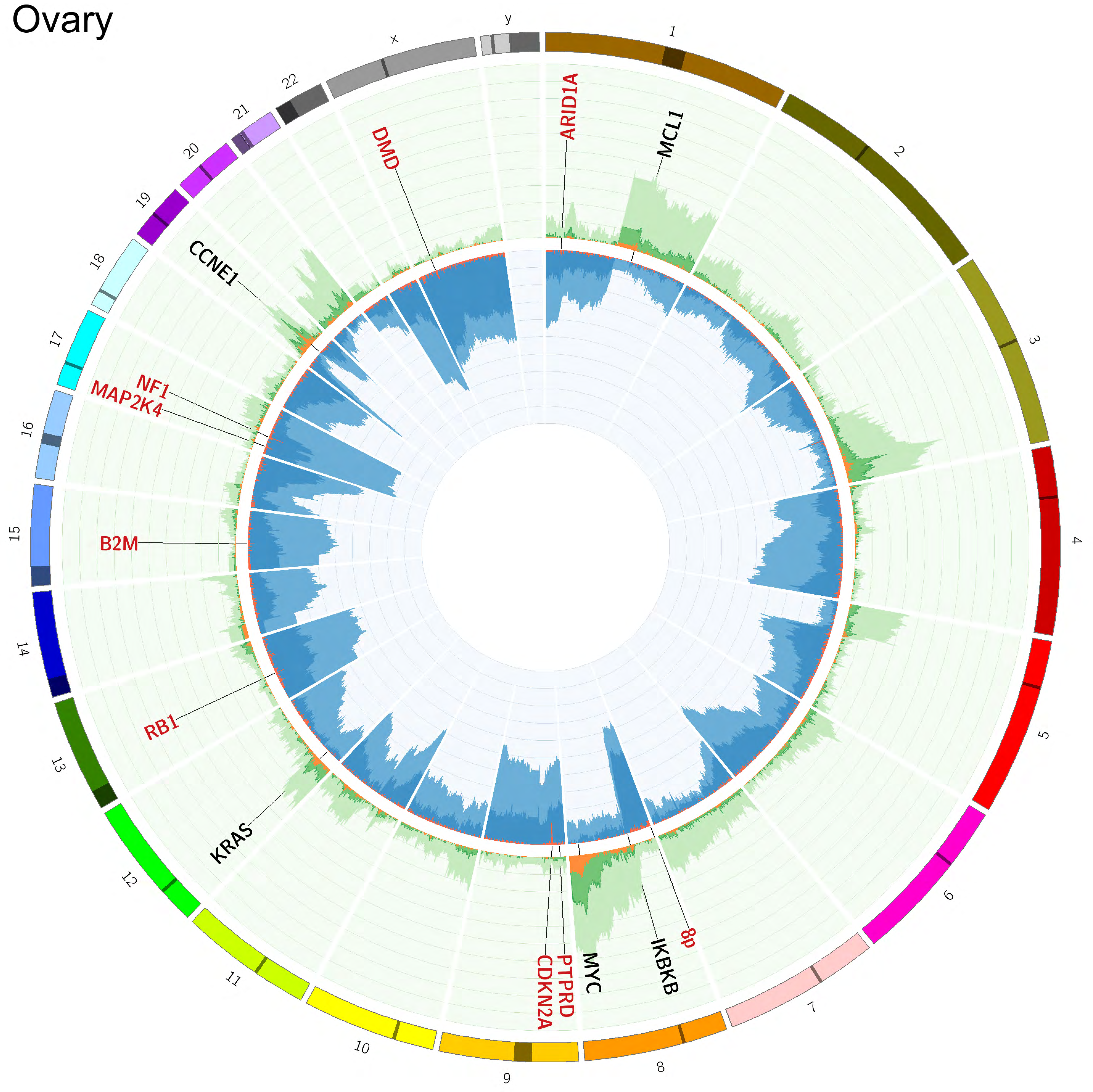

x
y
ARID1A
MCL1
DMD
CCNE1
NF1
MAP2K4
B2M
RB1
KRAS
CDKN2A
PTPRD
MYC
IKBKB
8p

Pancreas

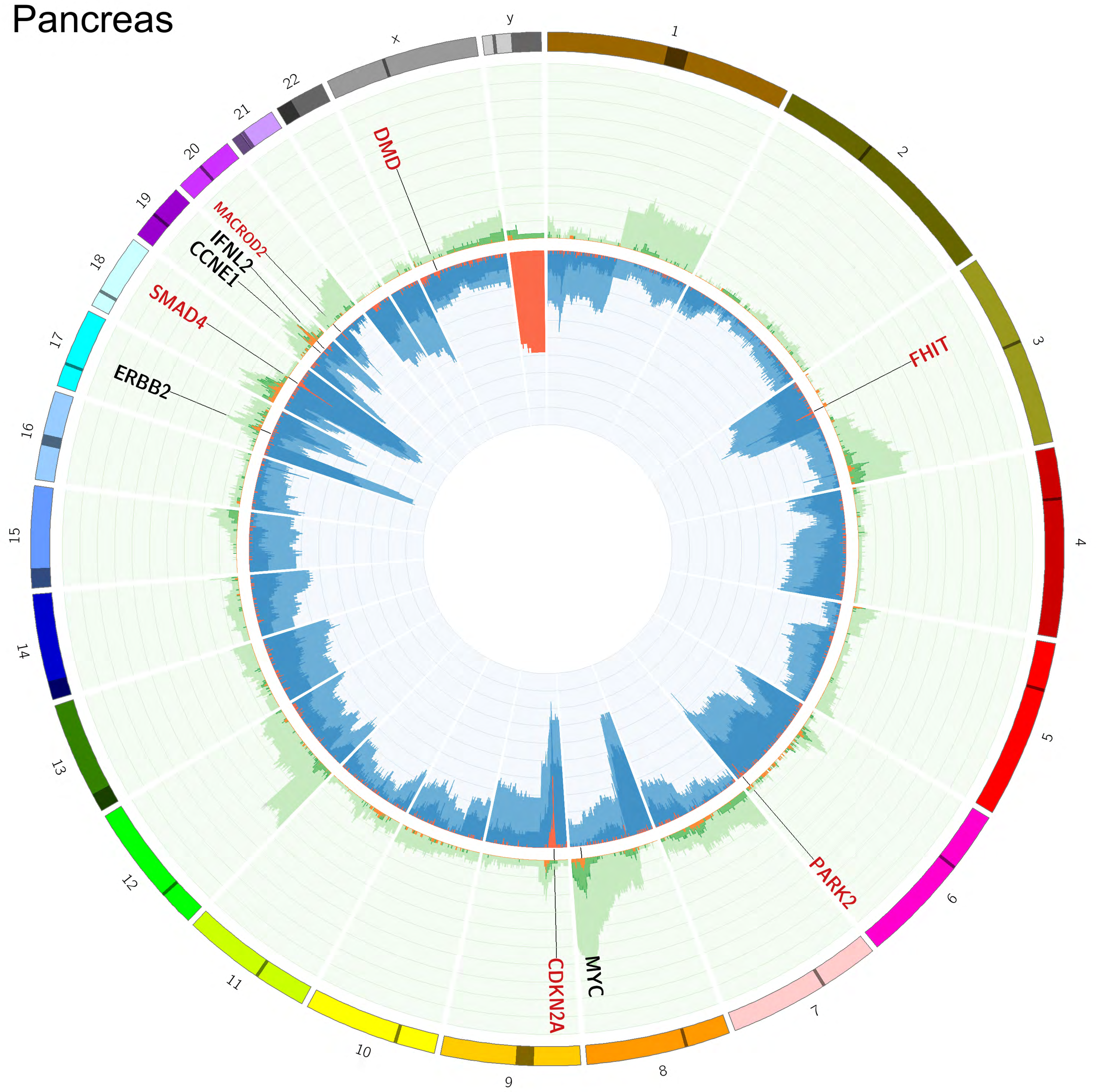

Prostate

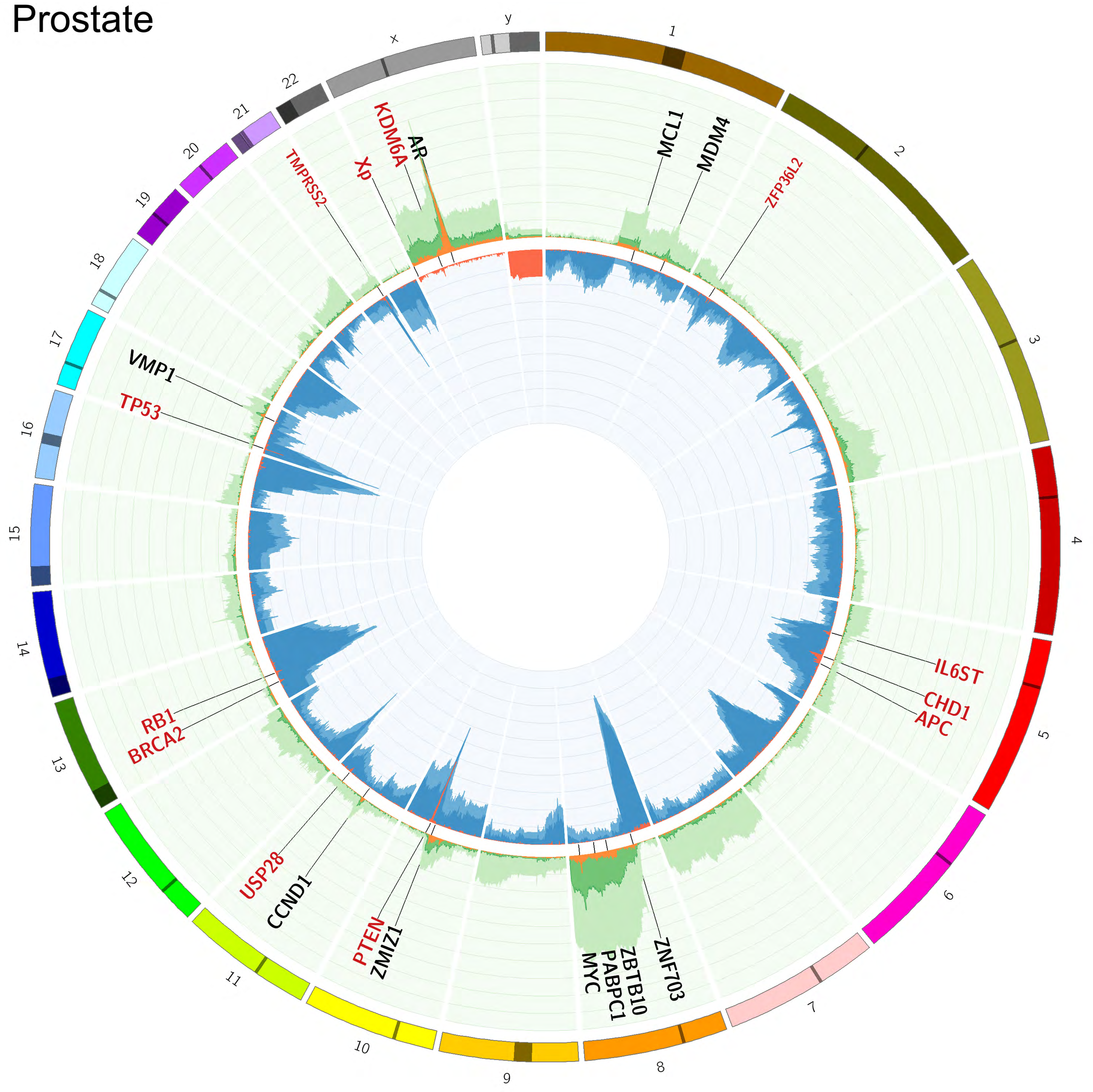

### Skin

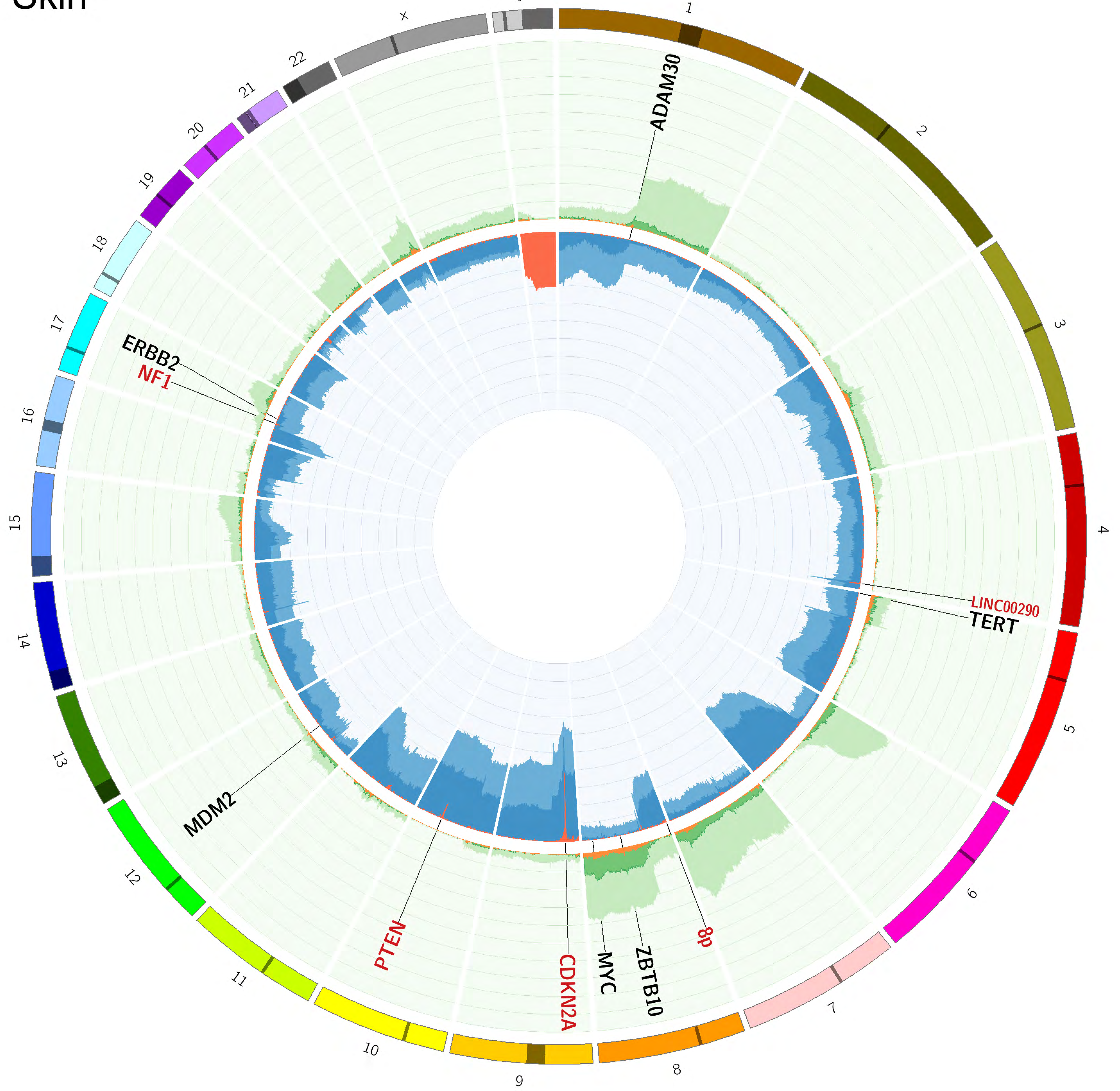

### Stomach

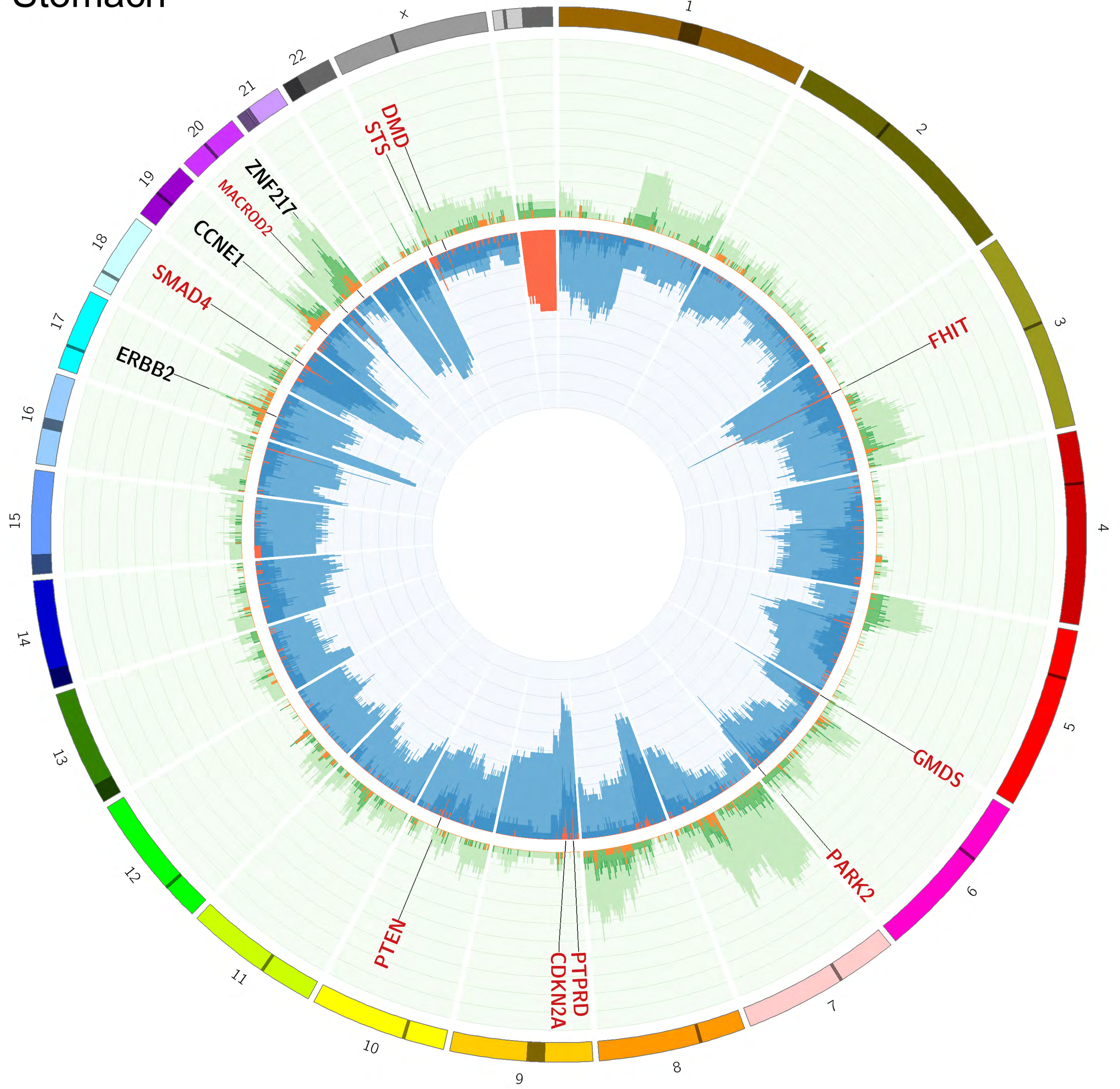

### Urinary tract

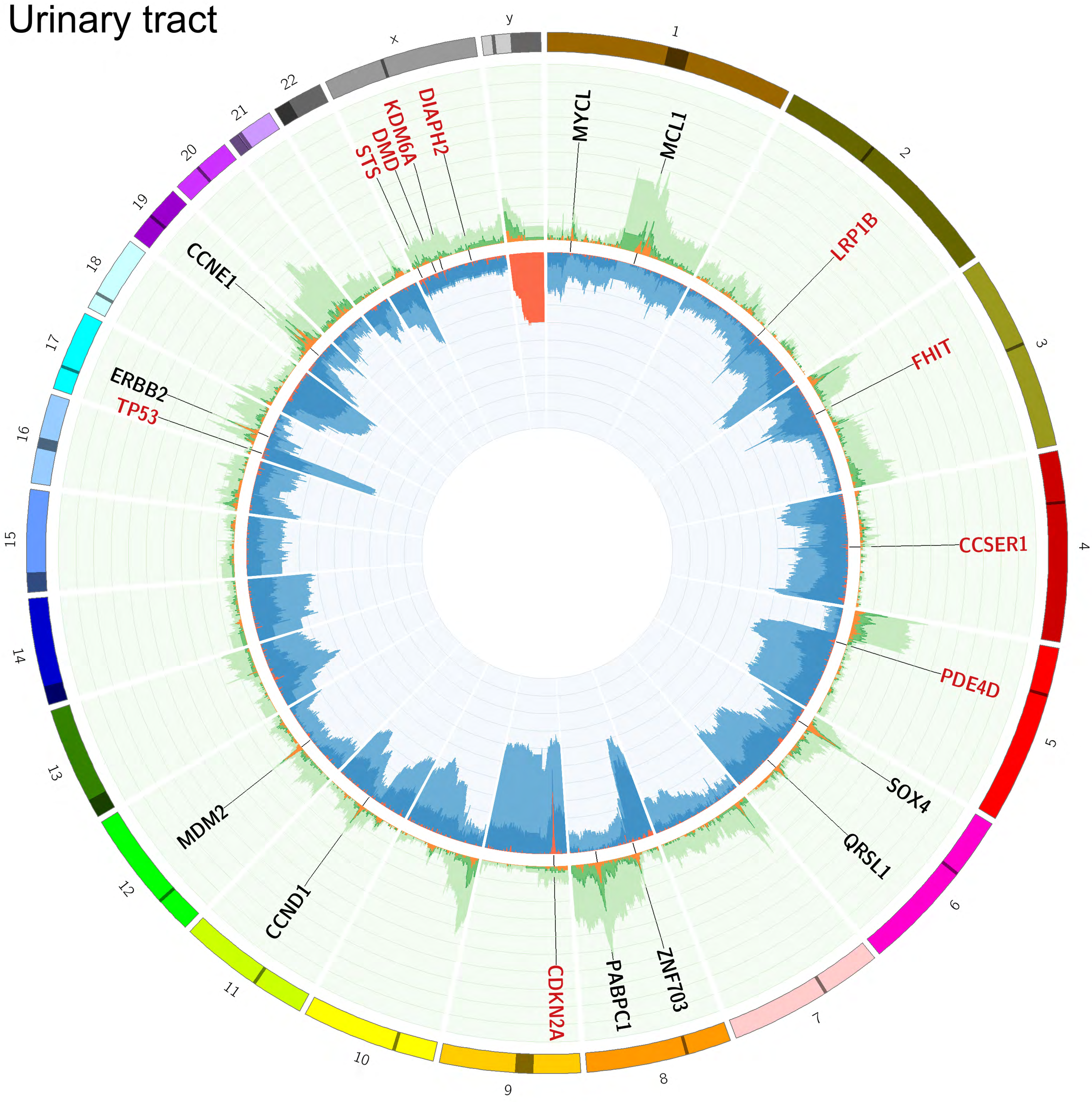

### Uterus

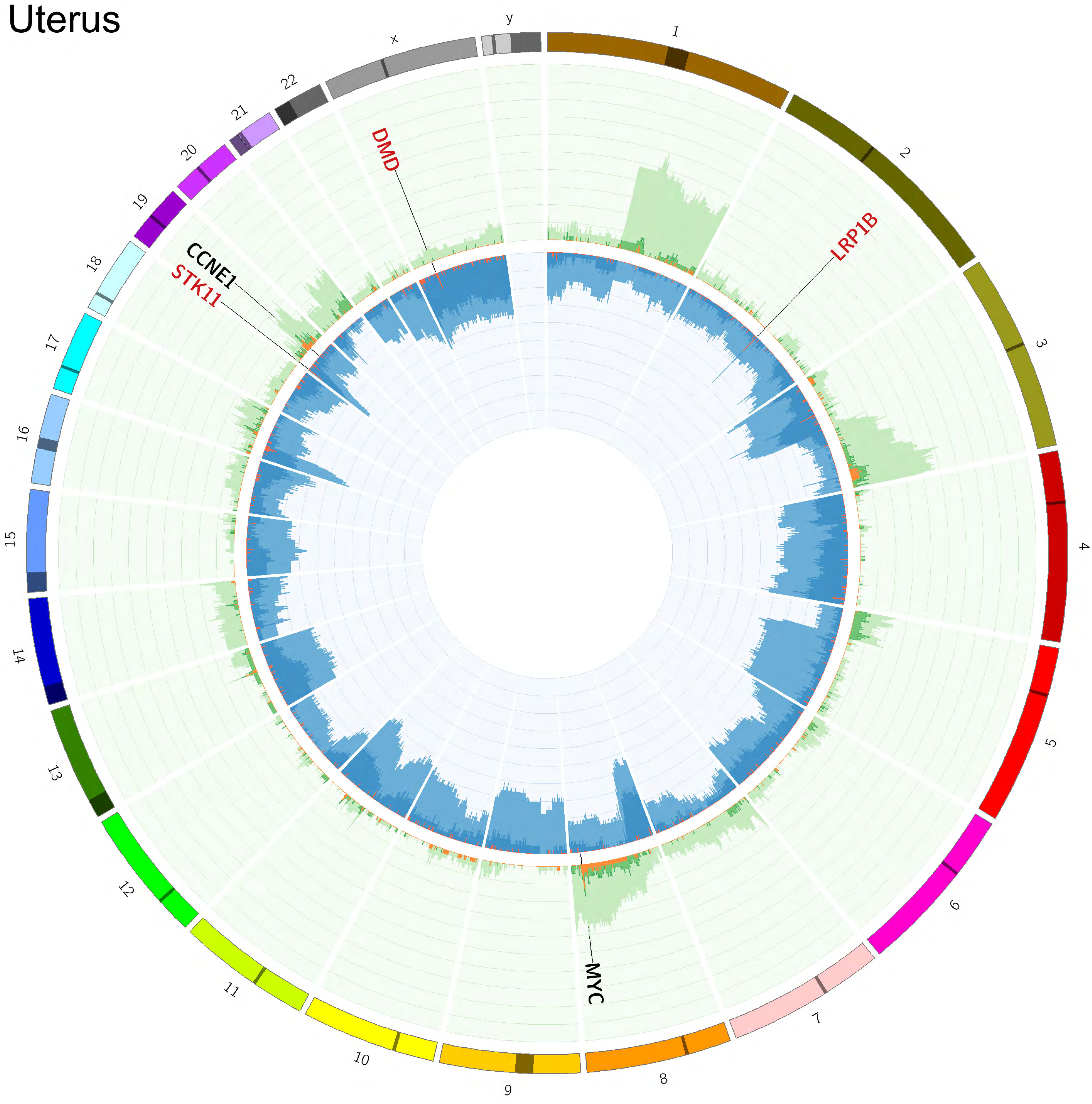
