## Supplementary Image 2 for "Pan-cancer whole genome analyses of metastatic solid tumors"

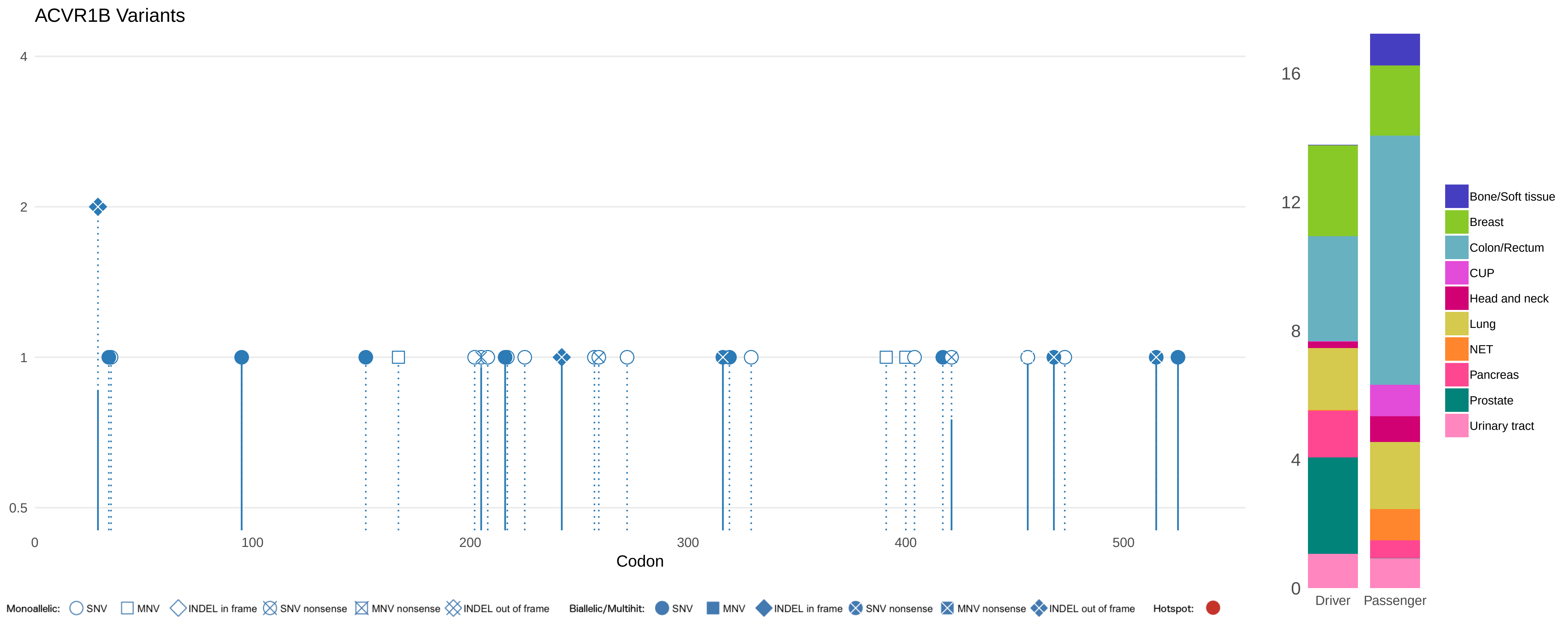
ACVR1B Variants
4
2
1
0.5
0
100
200
300
400
500
Codon
Monoallelic: SNV MNV INDEL in frame SNV nonsense MNV nonsense INDEL out of frame
Biallelic/Multihit: SNV MNV INDEL in frame SNV nonsense MNV nonsense INDEL out of frame
Hotspot:
16
12
8
4
0
Driver
Passenger
Bone/Soft tissue
Breast
Colon/Rectum
CUP
Head and neck
Lung
NET
Pancreas
Prostate
Urinary tract

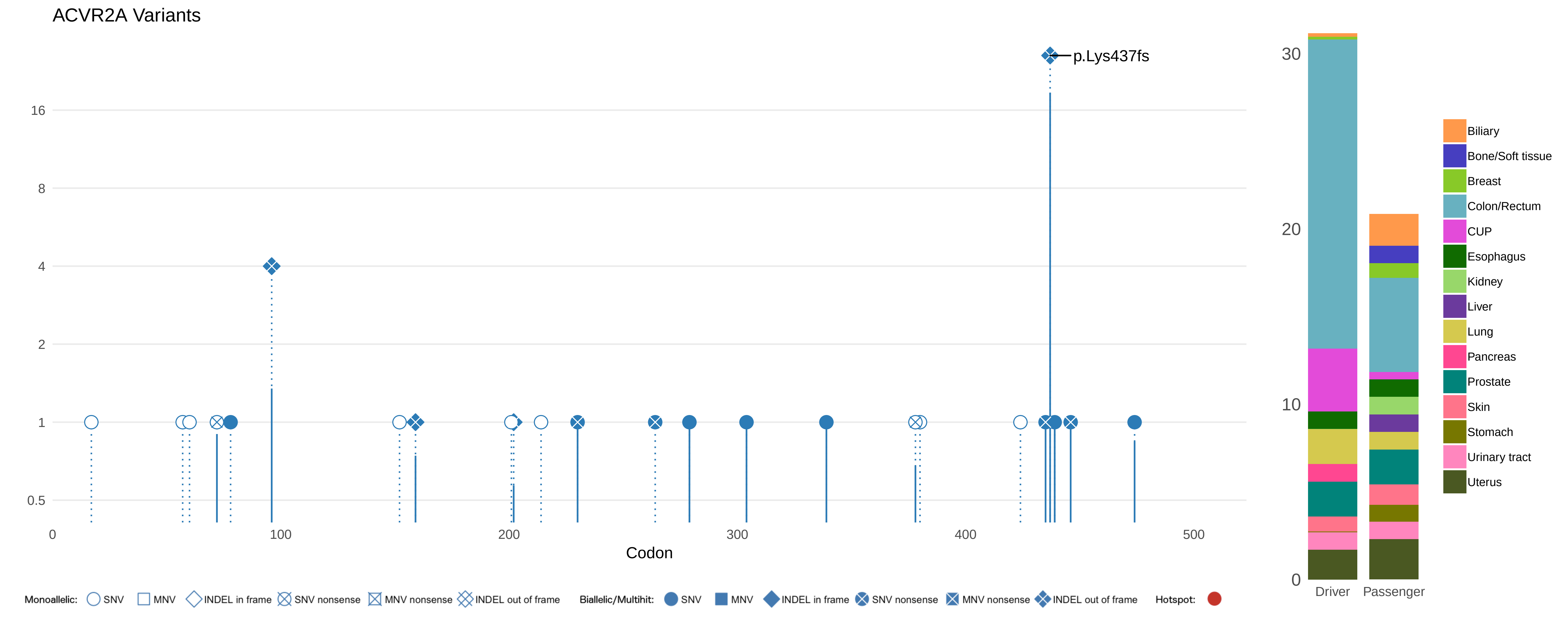
ACVR2A Variants
p.Lys437fs
16
8
4
2
1
0.5
0
100
200
300
400
500
Codon
30
20
10
0
Driver
Passenger
Biliary
Bone/Soft tissue
Breast
Colon/Rectum
CUP
Esophagus
Kidney
Liver
Lung
Pancreas
Prostate
Skin
Stomach
Urinary tract
Uterus
Monoallelic: SNV MNV INDEL in frame SNV nonsense MNV nonsense INDEL out of frame
Biallelic/Multihit: SNV MNV INDEL in frame SNV nonsense MNV nonsense INDEL out of frame
Hotspot:

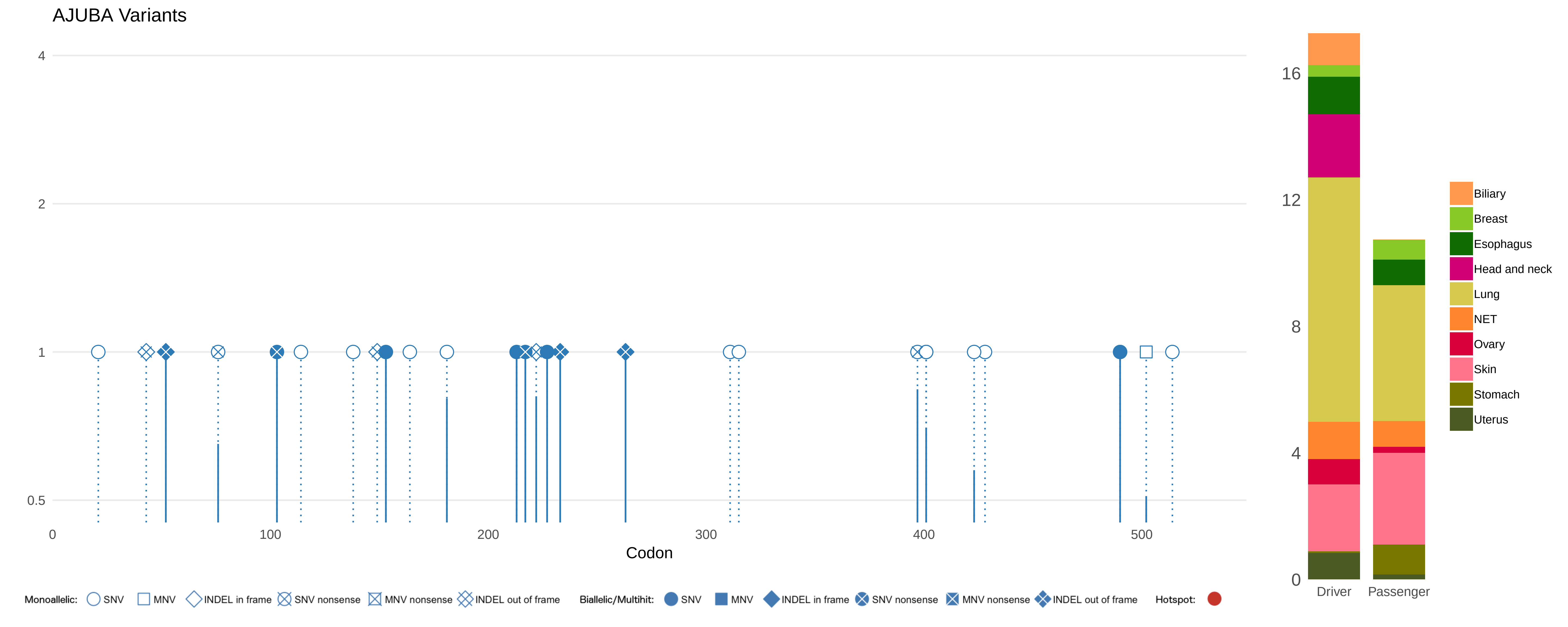
AJUBA Variants
4
2
1
0.5
0
100
200
300
400
500
Codon
16
12
8
4
0
Driver
Passenger
Biliary
Breast
Esophagus
Head and neck
Lung
NET
Ovary
Skin
Stomach
Uterus
Monoallelic: SNV MNV INDEL in frame SNV nonsense MNV nonsense INDEL out of frame
Biallelic/Multihit: SNV MNV INDEL in frame SNV nonsense MNV nonsense INDEL out of frame
Hotspot:

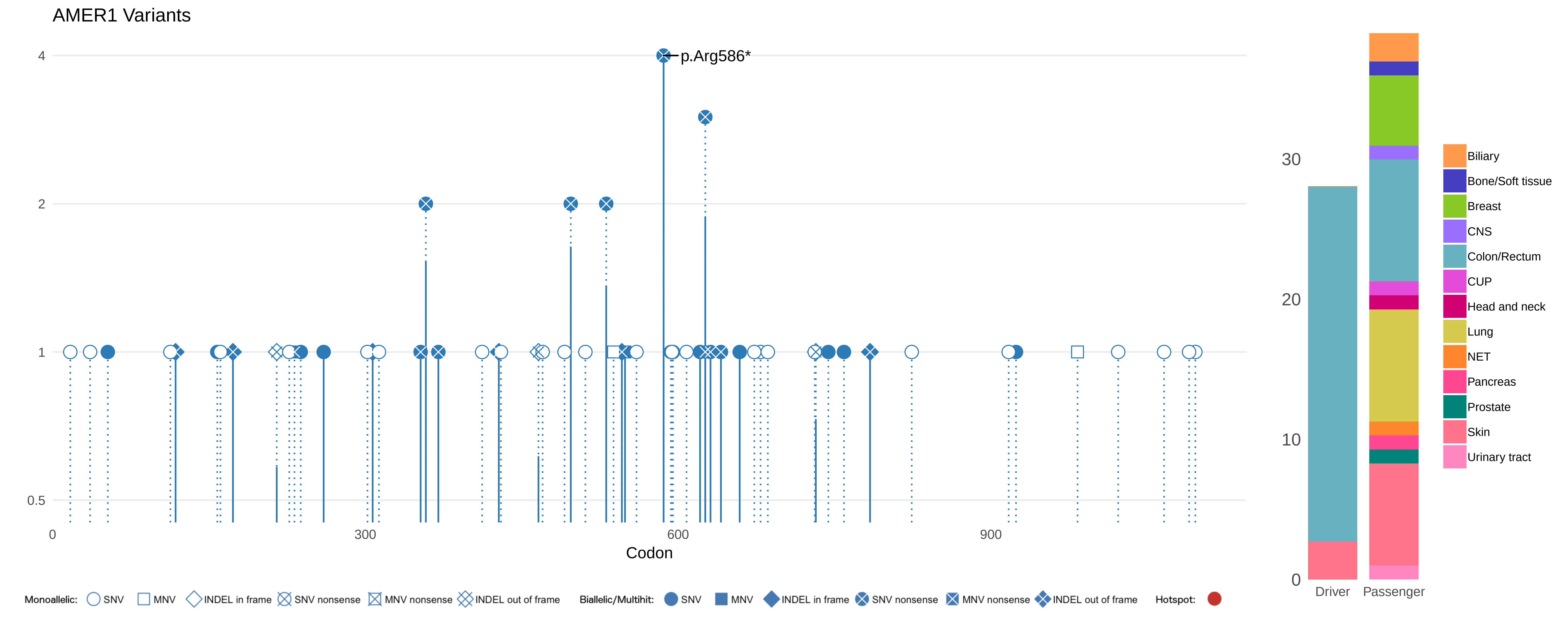
AMER1 Variants
p.Arg586*
4
2
1
0.5
0
300
600
900
Codon
30
20
10
0
Driver
Passenger
Biliary
Bone/Soft tissue
Breast
CNS
Colon/Rectum
CUP
Head and neck
Lung
NET
Pancreas
Prostate
Skin
Urinary tract
Monoallelic: SNV MNV INDEL in frame SNV nonsense MNV nonsense INDEL out of frame
Biallelic/Multihit: SNV MNV INDEL in frame SNV nonsense MNV nonsense INDEL out of frame
Hotspot:

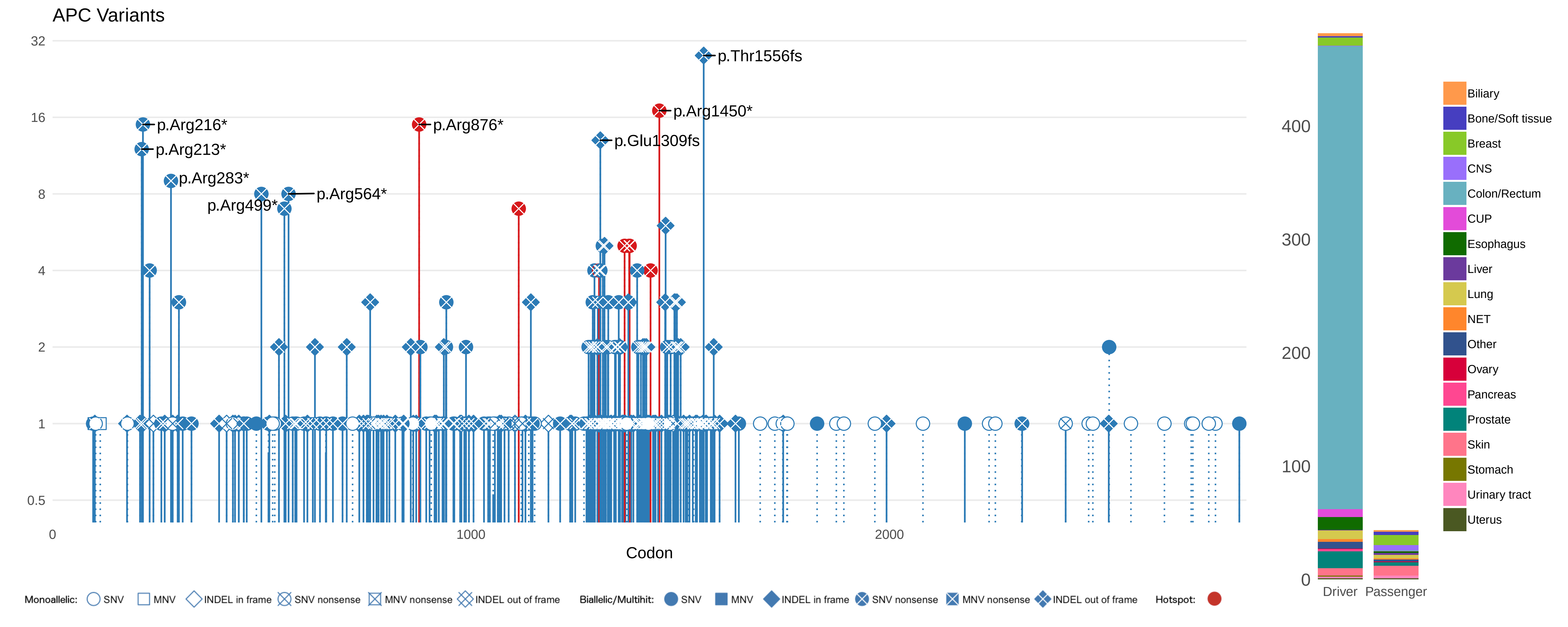

APC Variants
32
16
8
4
2
1
0.5
0
1000
2000
Codon
p.Thr1556fs
p.Arg1450*
p.Arg216*
p.Arg876*
p.Glu1309fs
p.Arg213*
p.Arg283*
p.Arg564*
p.Arg499*
400
300
200
100
0
Driver
Passenger
Biliary
Bone/Soft tissue
Breast
CNS
Colon/Rectum
CUP
Esophagus
Liver
Lung
NET
Other
Ovary
Pancreas
Prostate
Skin
Stomach
Urinary tract
Uterus
Monoallelic: SNV MNV INDEL in frame SNV nonsense MNV nonsense INDEL out of frame
Biallelic/Multihit: SNV MNV INDEL in frame SNV nonsense MNV nonsense INDEL out of frame
Hotspot:

ARHGAP35 Variants

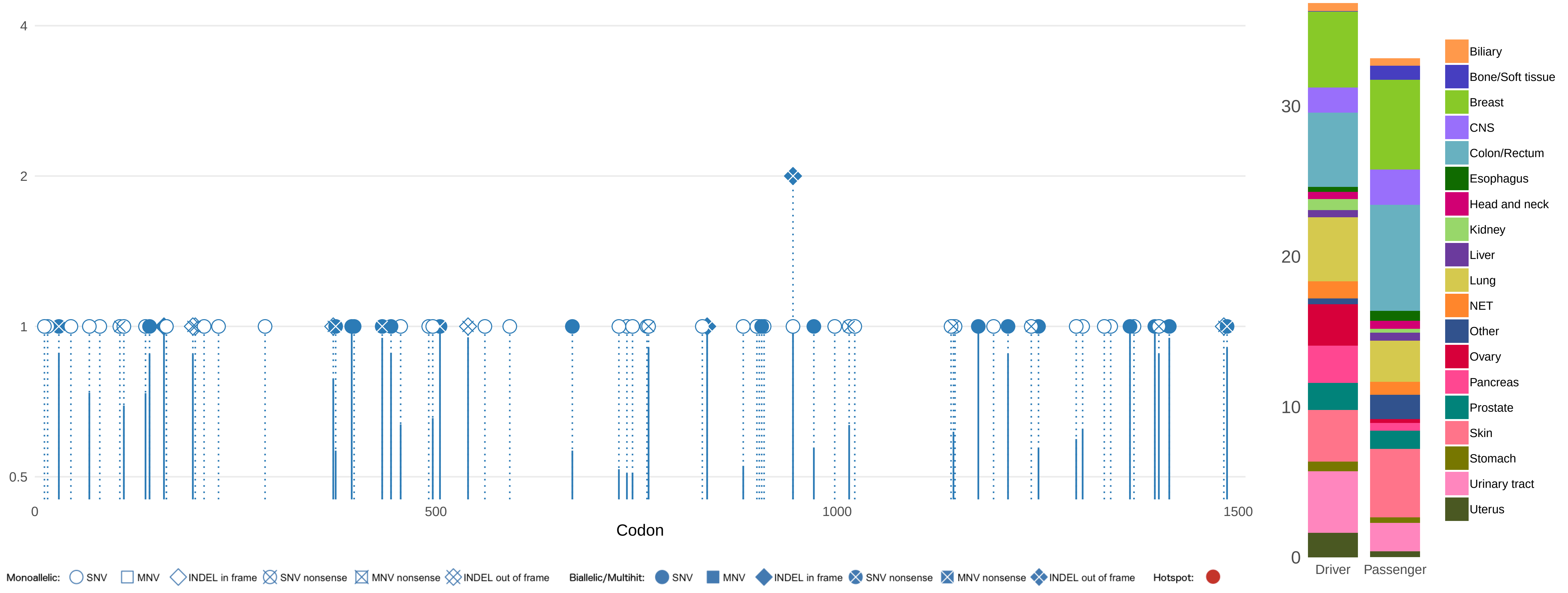

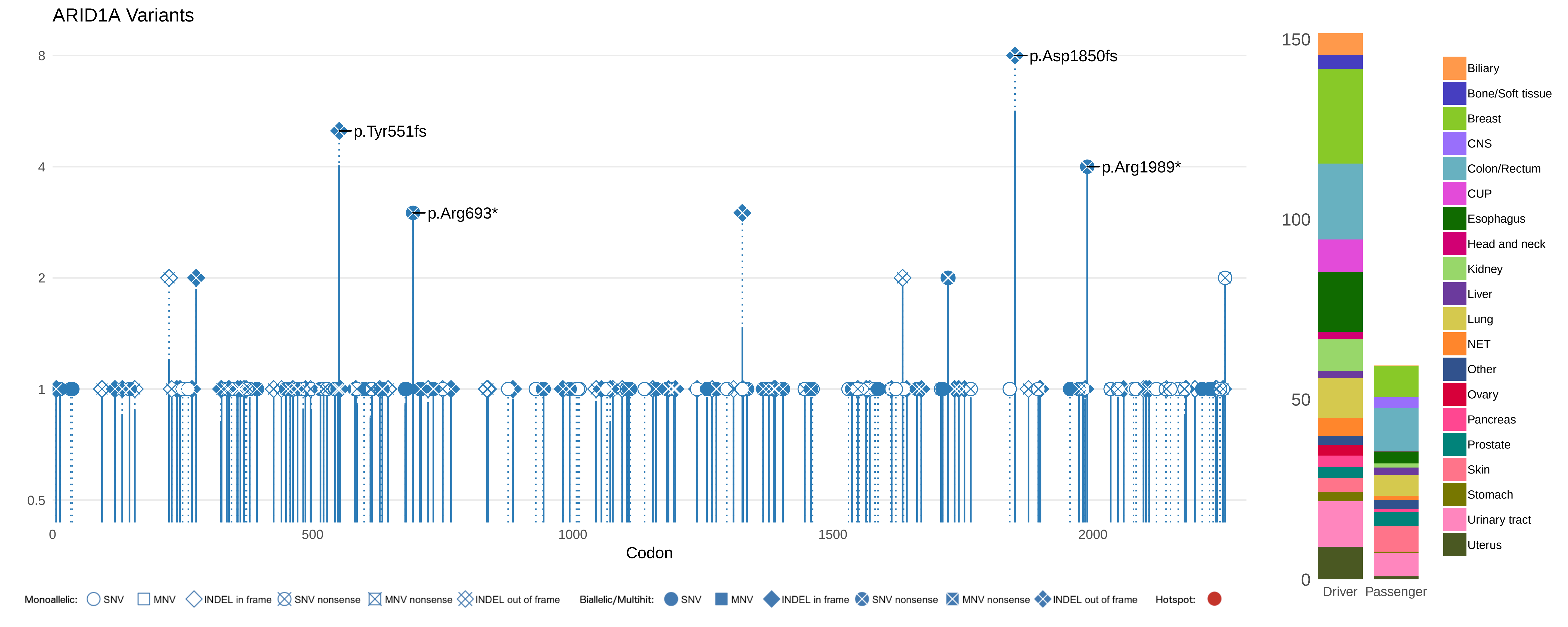

ARID1A Variants
8
4
2
1
0.5
p.Asp1850fs
p.Tyr551fs
p.Arg693*
p.Arg1989*
0
500
1000
1500
2000
Codon
Monoallelic: SNV MNV INDEL in frame SNV nonsense MNV nonsense INDEL out of frame
Biallelic/Multihit: SNV MNV INDEL in frame SNV nonsense MNV nonsense INDEL out of frame
Hotspot:
150
100
50
0
Driver
Passenger
Biliary
Bone/Soft tissue
Breast
CNS
Colon/Rectum
CUP
Esophagus
Head and neck
Kidney
Liver
Lung
NET
Other
Ovary
Pancreas
Prostate
Skin
Stomach
Urinary tract
Uterus

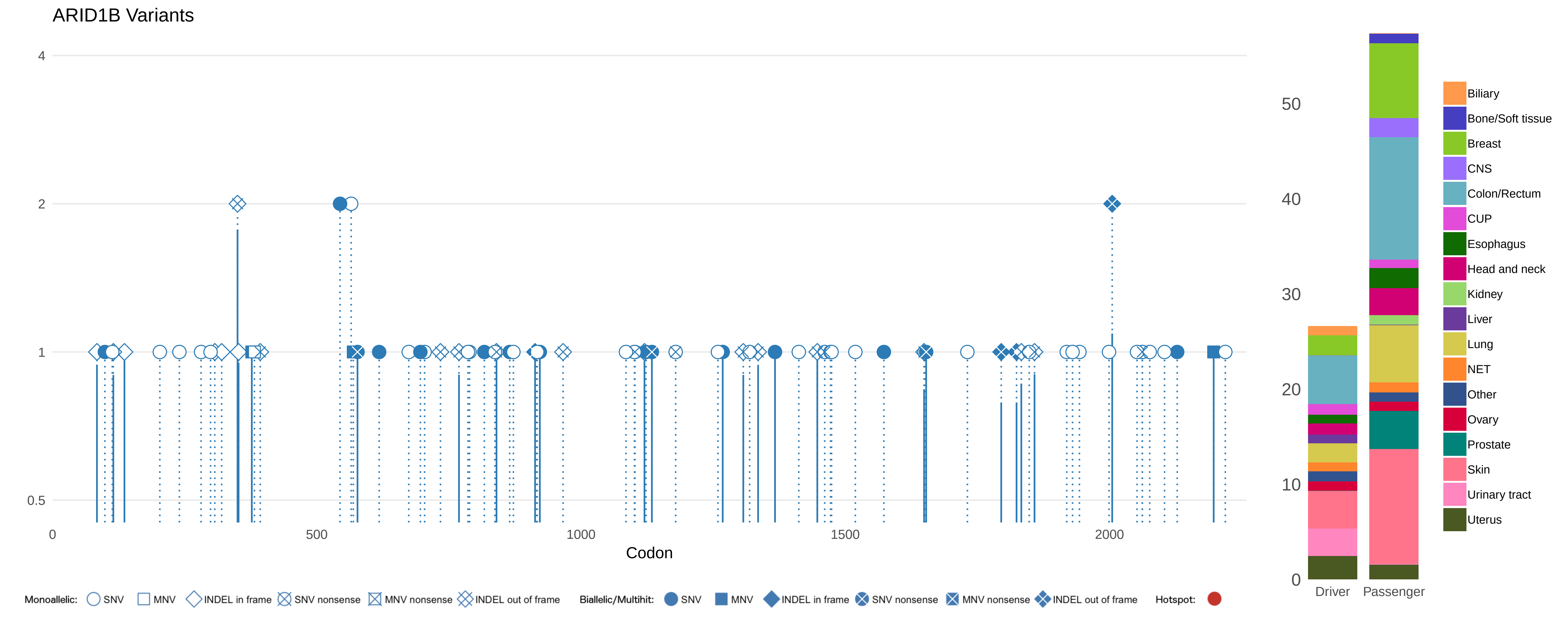

ARID1B Variants
4
2
1
0.5
0
500
1000
1500
2000
Codon
50
40
30
20
10
0
Driver
Passenger
Biliary
Bone/Soft tissue
Breast
CNS
Colon/Rectum
CUP
Esophagus
Head and neck
Kidney
Liver
Lung
NET
Other
Ovary
Prostate
Skin
Urinary tract
Uterus
Monoallelic: SNV MNV INDEL in frame SNV nonsense MNV nonsense INDEL out of frame
Biallelic/Multihit: SNV MNV INDEL in frame SNV nonsense MNV nonsense INDEL out of frame
Hotspot:

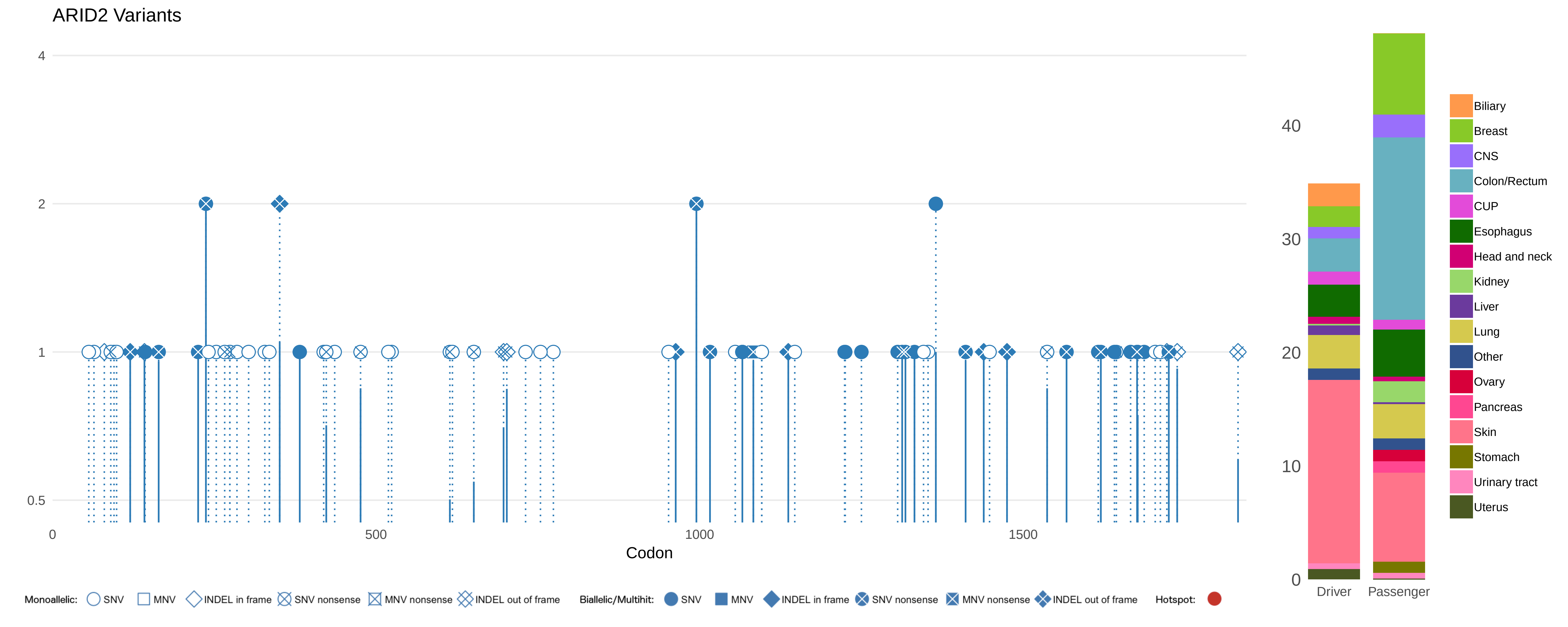

ARID2 Variants
4
2
1
0.5
0
500
1000
1500
Codon
40
30
20
10
0
Driver
Passenger
Biliary
Breast
CNS
Colon/Rectum
CUP
Esophagus
Head and neck
Kidney
Liver
Lung
Other
Ovary
Pancreas
Skin
Stomach
Urinary tract
Uterus
Monoallelic: SNV MNV INDEL in frame SNV nonsense MNV nonsense INDEL out of frame
Biallelic/Multihit: SNV MNV INDEL in frame SNV nonsense MNV nonsense INDEL out of frame
Hotspot:

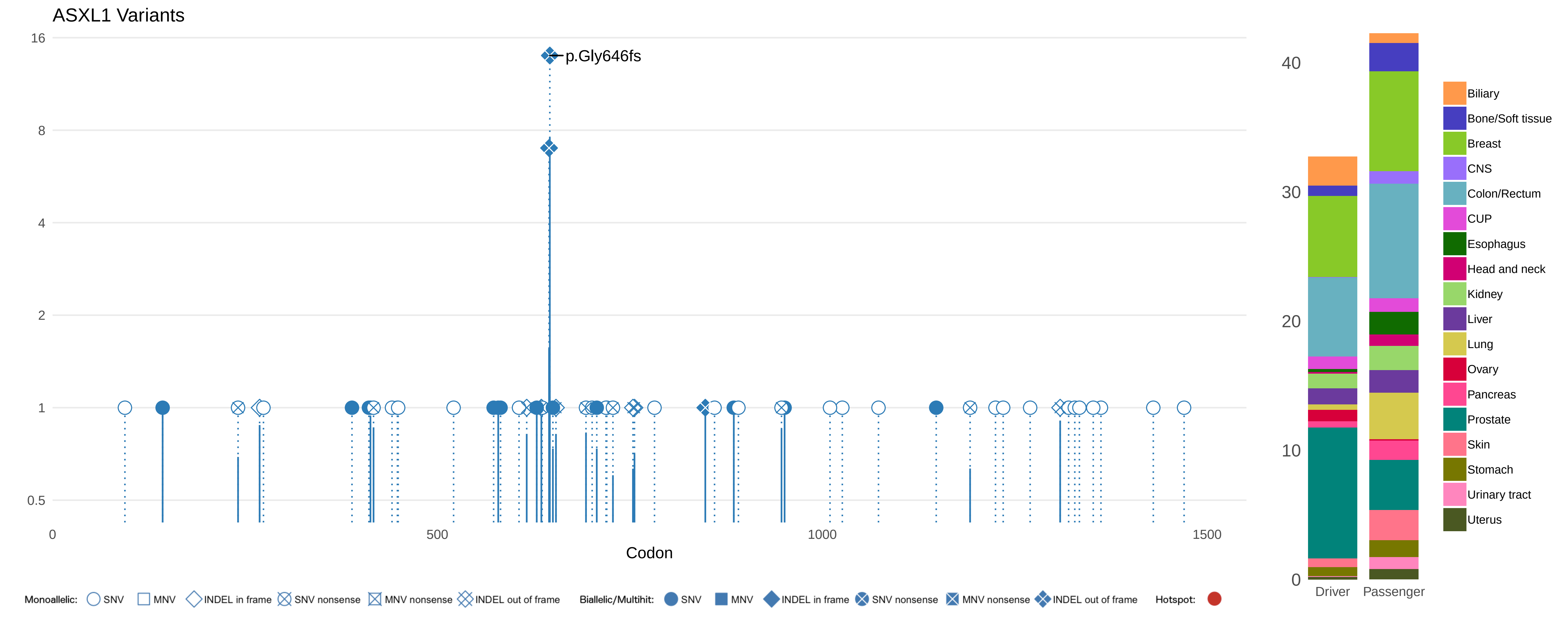

ASXL1 Variants
16
8
4
2
1
0.5
p.Gly646fs
0
500
1000
1500
Codon
40
30
20
10
0
Driver
Passenger
Biliary
Bone/Soft tissue
Breast
CNS
Colon/Rectum
CUP
Esophagus
Head and neck
Kidney
Liver
Lung
Ovary
Pancreas
Prostate
Skin
Stomach
Urinary tract
Uterus
Monoallelic: SNV MNV INDEL in frame SNV nonsense MNV nonsense INDEL out of frame
Biallelic/Multihit: SNV MNV INDEL in frame SNV nonsense MNV nonsense INDEL out of frame
Hotspot:

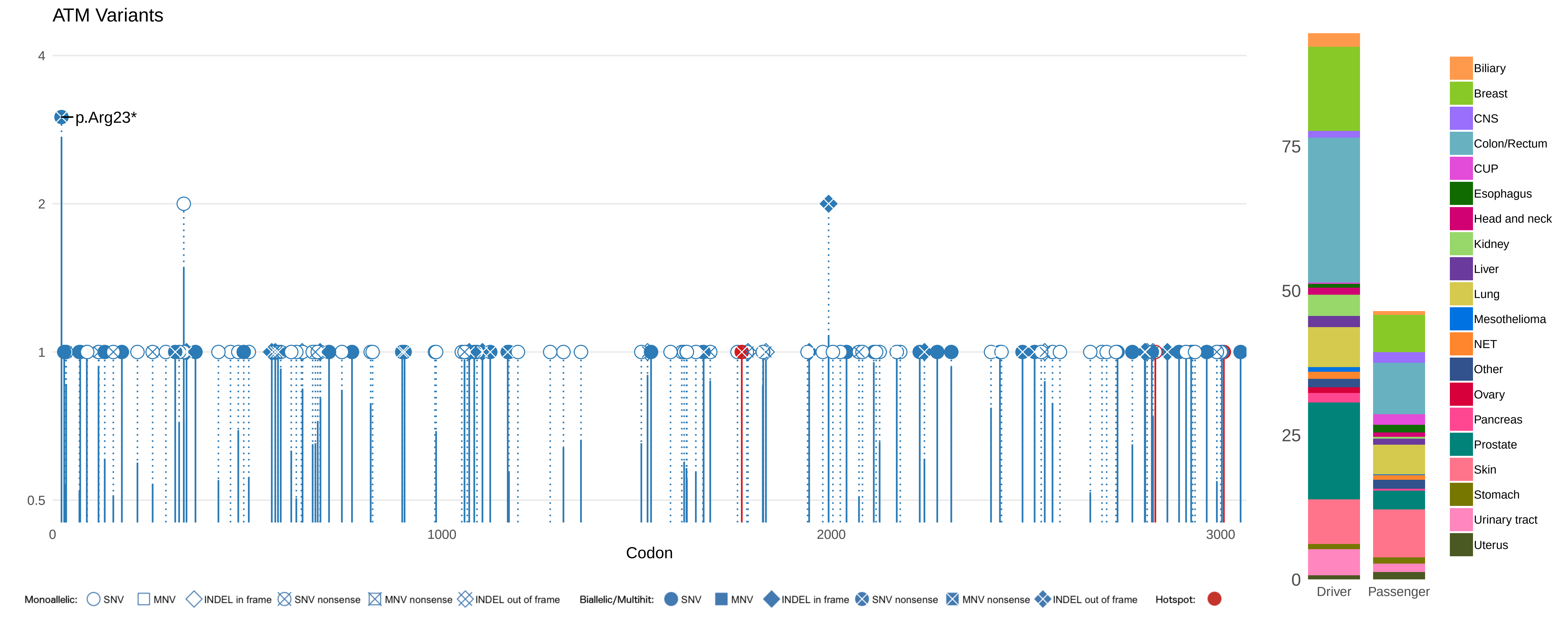

ATM Variants
p.Arg23*
4
2
1
0.5
0
1000
2000
3000
Codon
Monoallelic: SNV MNV INDEL in frame SNV nonsense MNV nonsense INDEL out of frame
Biallelic/Multihit: SNV MNV INDEL in frame SNV nonsense MNV nonsense INDEL out of frame
Hotspot:
75
50
25
0
Driver
Passenger
Biliary
Breast
CNS
Colon/Rectum
CUP
Esophagus
Head and neck
Kidney
Liver
Lung
Mesothelioma
NET
Other
Ovary
Pancreas
Prostate
Skin
Stomach
Urinary tract
Uterus

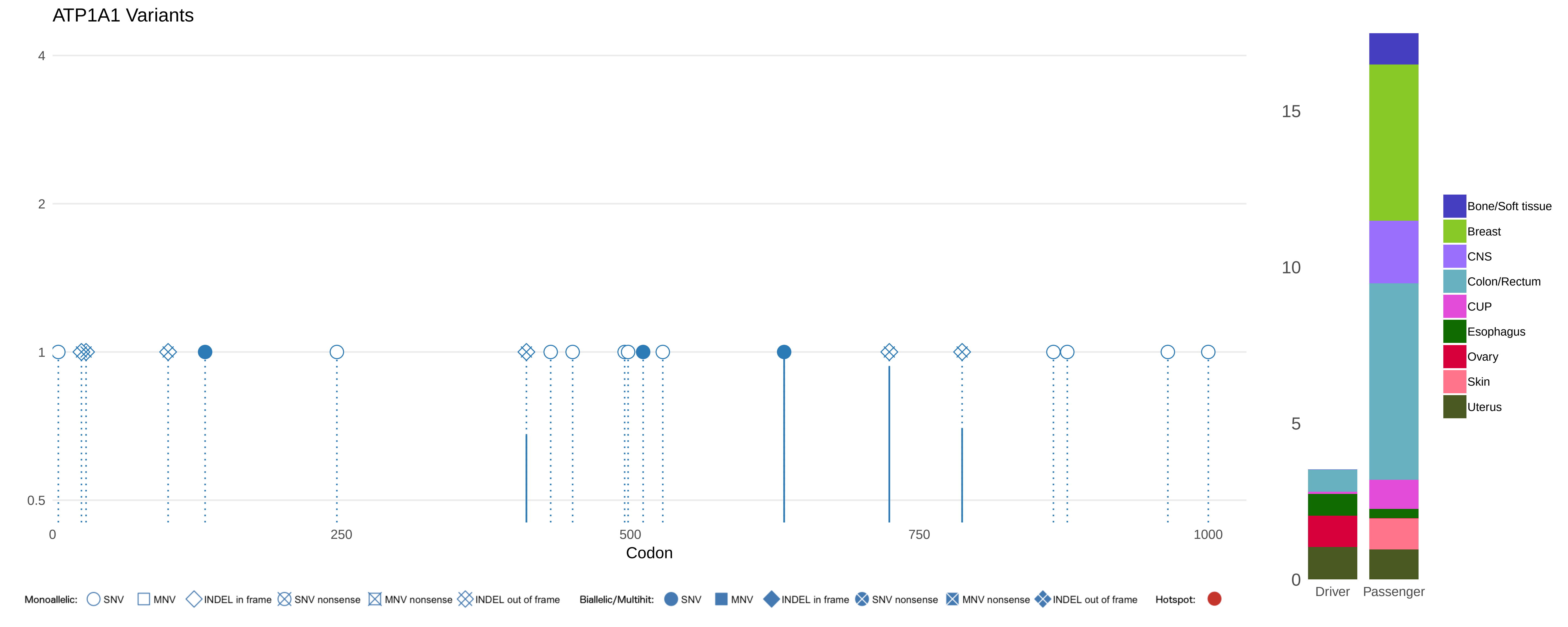

ATP1A1 Variants
4
2
1
0.5
0
250
500
750
1000
Codon
15
10
5
0
Driver
Passenger
Bone/Soft tissue
Breast
CNS
Colon/Rectum
CUP
Esophagus
Ovary
Skin
Uterus
Monoallelic: SNV MNV INDEL in frame SNV nonsense MNV nonsense INDEL out of frame
Biallelic/Multihit: SNV MNV INDEL in frame SNV nonsense MNV nonsense INDEL out of frame
Hotspot:

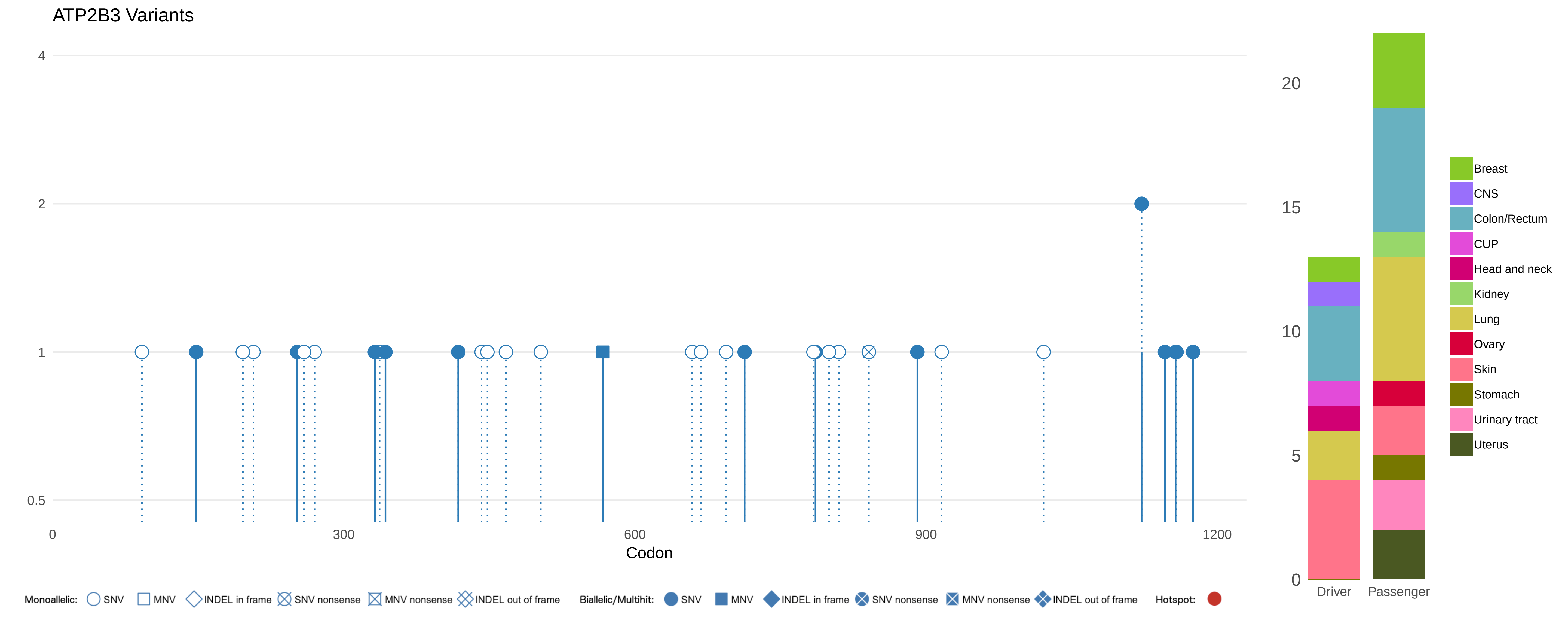
ATP2B3 Variants
4
2
1
0.5
0
300
600
900
1200
Codon
20
15
10
5
0
Driver
Passenger
Breast
CNS
Colon/Rectum
CUP
Head and neck
Kidney
Lung
Ovary
Skin
Stomach
Urinary tract
Uterus
Monoallelic: SNV MNV INDEL in frame SNV nonsense MNV nonsense INDEL out of frame
Biallelic/Multihit: SNV MNV INDEL in frame SNV nonsense MNV nonsense INDEL out of frame
Hotspot:

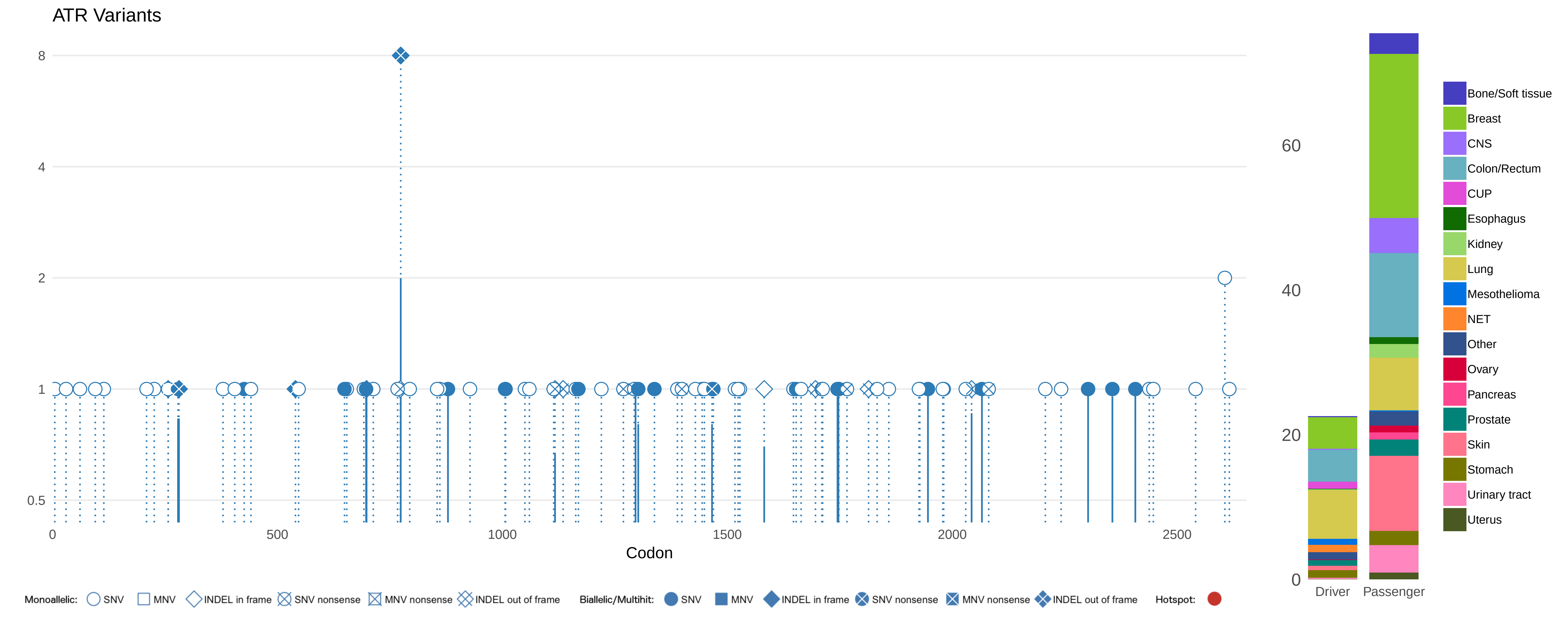

ATR Variants
8
4
2
1
0.5
0
500
1000
1500
2000
2500
Codon
60
40
20
0
Driver
Passenger
Bone/Soft tissue
Breast
CNS
Colon/Rectum
CUP
Esophagus
Kidney
Lung
Mesothelioma
NET
Other
Ovary
Pancreas
Prostate
Skin
Stomach
Urinary tract
Uterus
Monoallelic: SNV MNV INDEL in frame SNV nonsense MNV nonsense INDEL out of frame
Biallelic/Multihit: SNV MNV INDEL in frame SNV nonsense MNV nonsense INDEL out of frame
Hotspot:

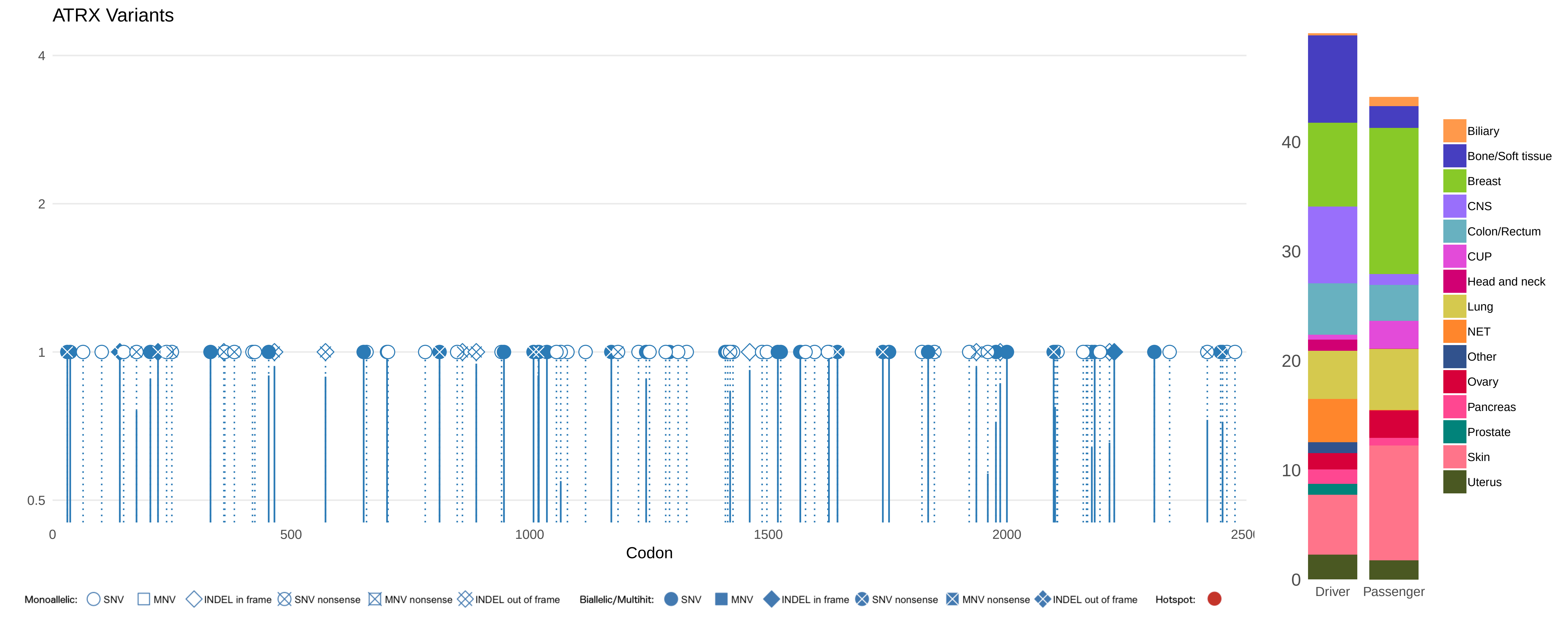

ATRX Variants
4
2
1
0.5
0
500
1000
1500
2000
2500
Codon
Monoallelic: SNV MNV INDEL in frame SNV nonsense MNV nonsense INDEL out of frame
Biallelic/Multihit: SNV MNV INDEL in frame SNV nonsense MNV nonsense INDEL out of frame
Hotspot:
40
30
20
10
0
Driver
Passenger
Biliary
Bone/Soft tissue
Breast
CNS
Colon/Rectum
CUP
Head and neck
Lung
NET
Other
Ovary
Pancreas
Prostate
Skin
Uterus

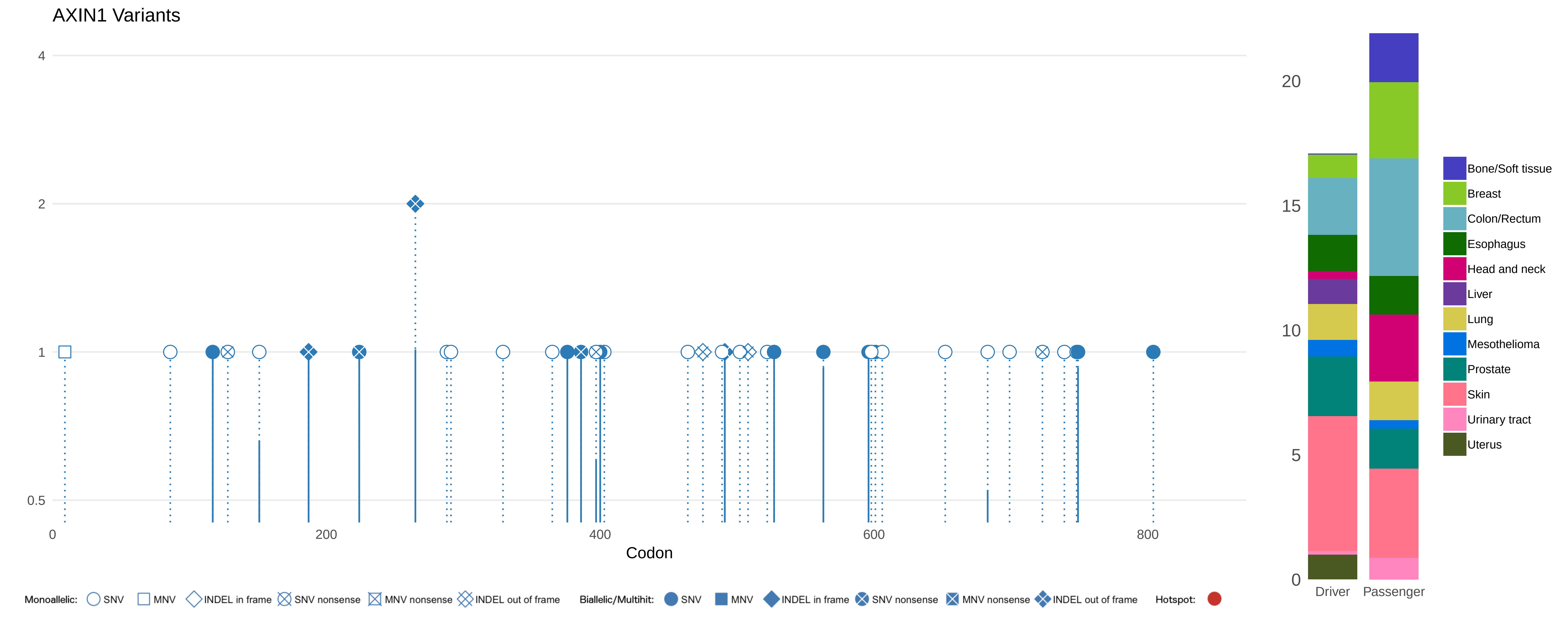
AXIN1 Variants
4
2
1
0.5
0
200
400
600
800
Codon
20
15
10
5
0
Driver
Passenger
Bone/Soft tissue
Breast
Colon/Rectum
Esophagus
Head and neck
Liver
Lung
Mesothelioma
Prostate
Skin
Urinary tract
Uterus
Monoallelic: SNV MNV INDEL in frame SNV nonsense MNV nonsense INDEL out of frame
Biallelic/Multihit: SNV MNV INDEL in frame SNV nonsense MNV nonsense INDEL out of frame
Hotspot:

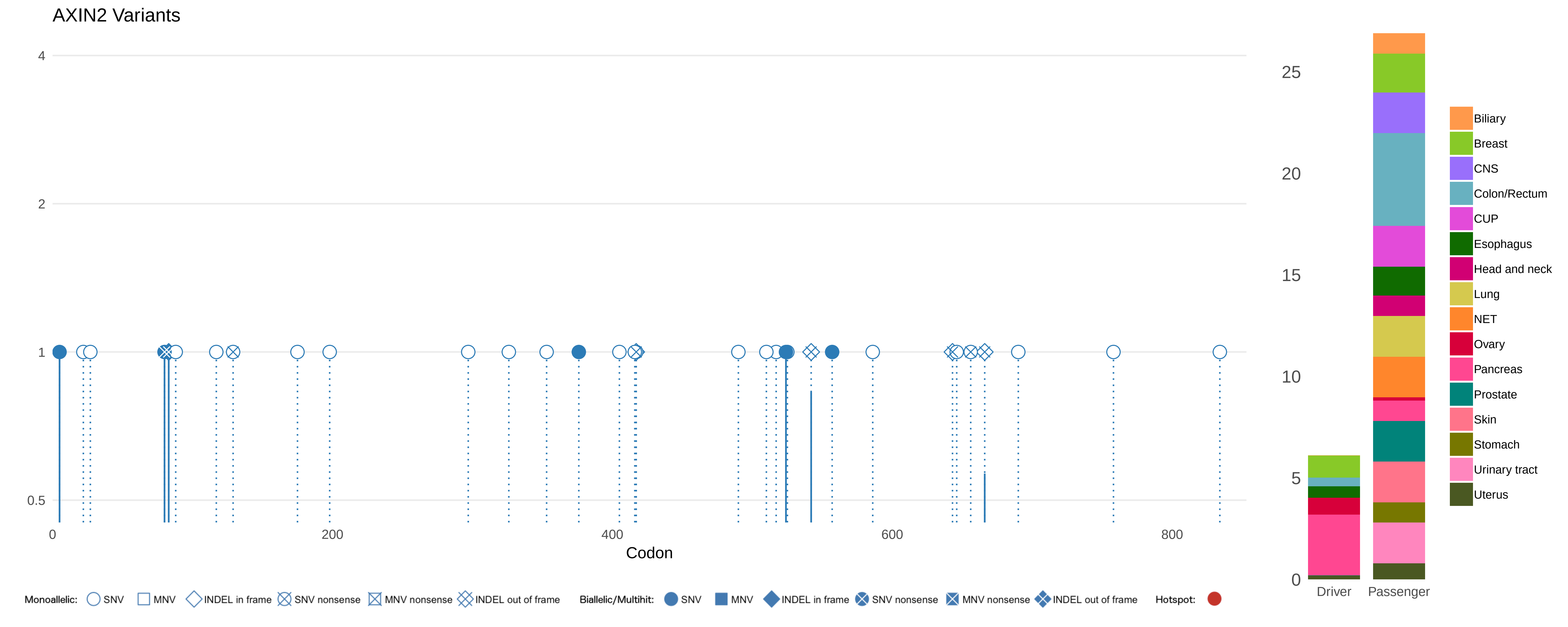
AXIN2 Variants
4
2
1
0.5
0
200
400
600
800
Codon
Monoallelic: SNV MNV INDEL in frame SNV nonsense MNV nonsense INDEL out of frame
Biallelic/Multihit: SNV MNV INDEL in frame SNV nonsense MNV nonsense INDEL out of frame
Hotspot:
25
20
15
10
5
0
Driver
Passenger
Biliary
Breast
CNS
Colon/Rectum
CUP
Esophagus
Head and neck
Lung
NET
Ovary
Pancreas
Prostate
Skin
Stomach
Urinary tract
Uterus

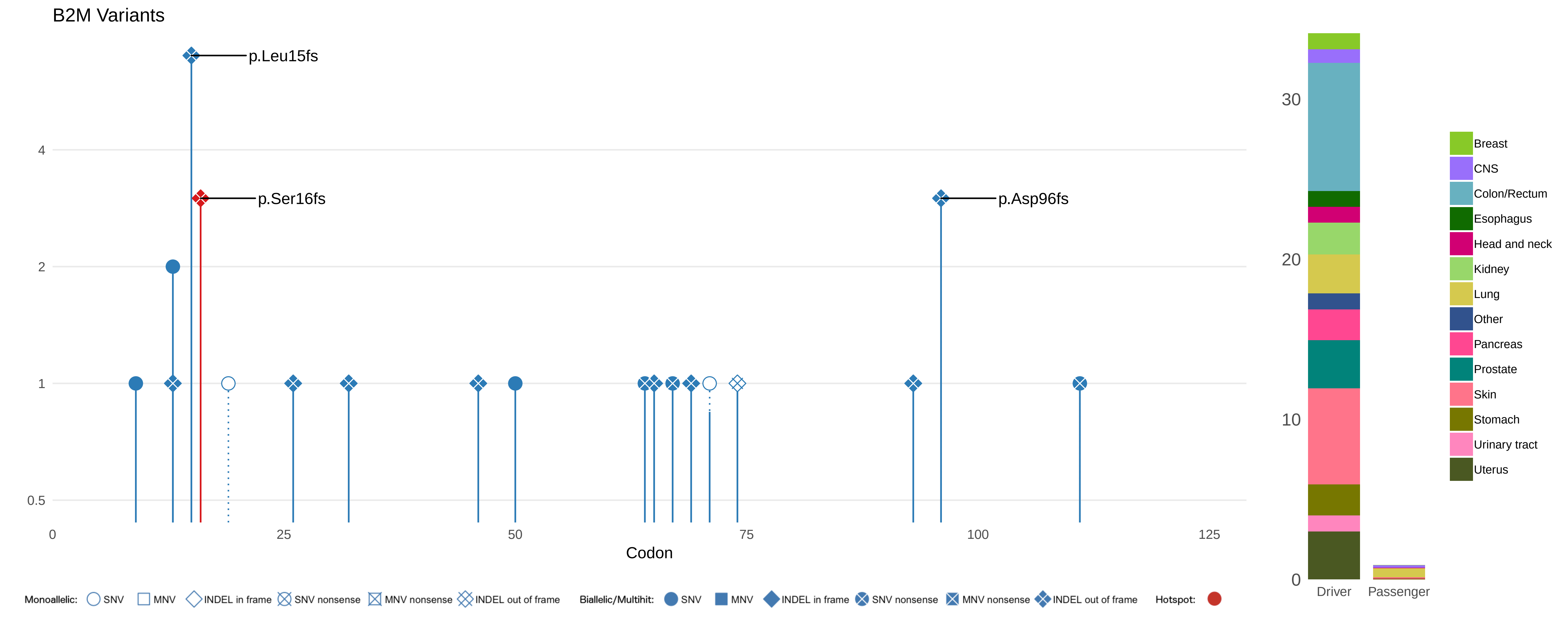

B2M Variants
p.Leu15fs
p.Ser16fs
p.Asp96fs
4
2
1
0.5
0
25
50
75
100
125
Codon
30
20
10
0
Driver
Passenger
Breast
CNS
Colon/Rectum
Esophagus
Head and neck
Kidney
Lung
Other
Pancreas
Prostate
Skin
Stomach
Urinary tract
Uterus
Monoallelic: SNV MNV INDEL in frame SNV nonsense MNV nonsense INDEL out of frame
Biallelic/Multihit: SNV MNV INDEL in frame SNV nonsense MNV nonsense INDEL out of frame
Hotspot:

### BAP1 Variants

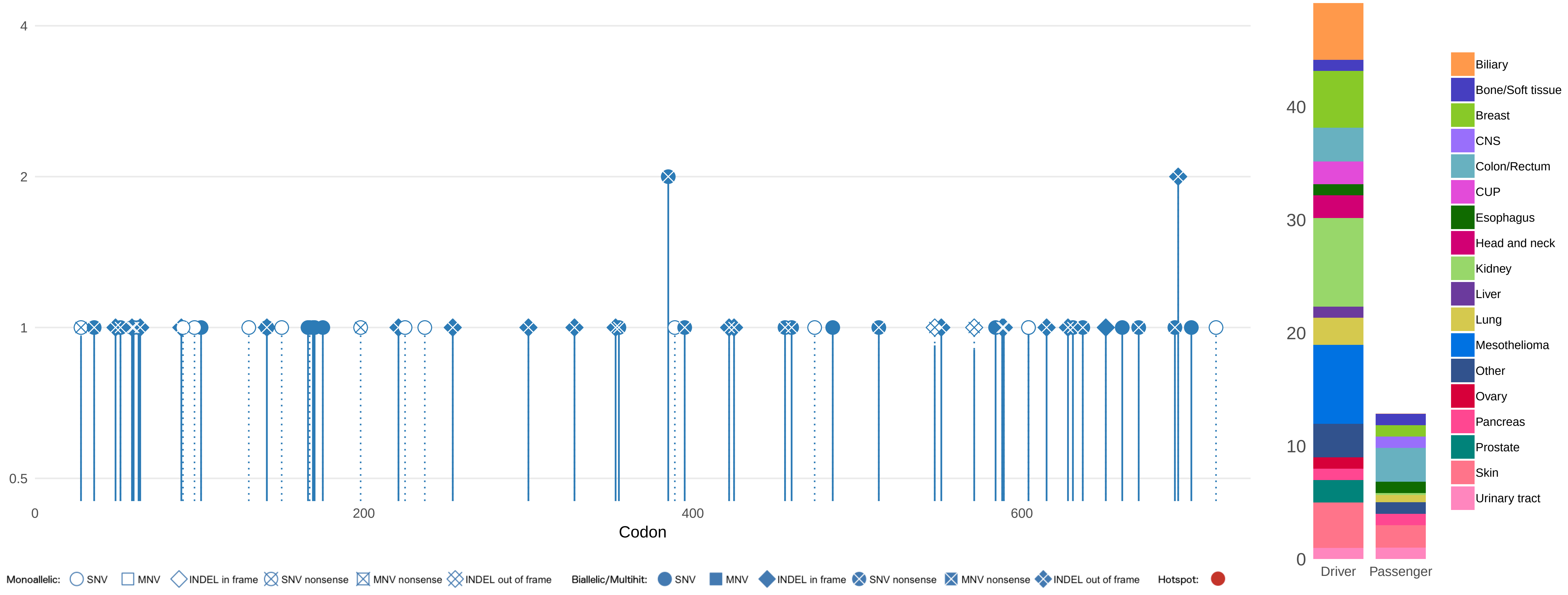

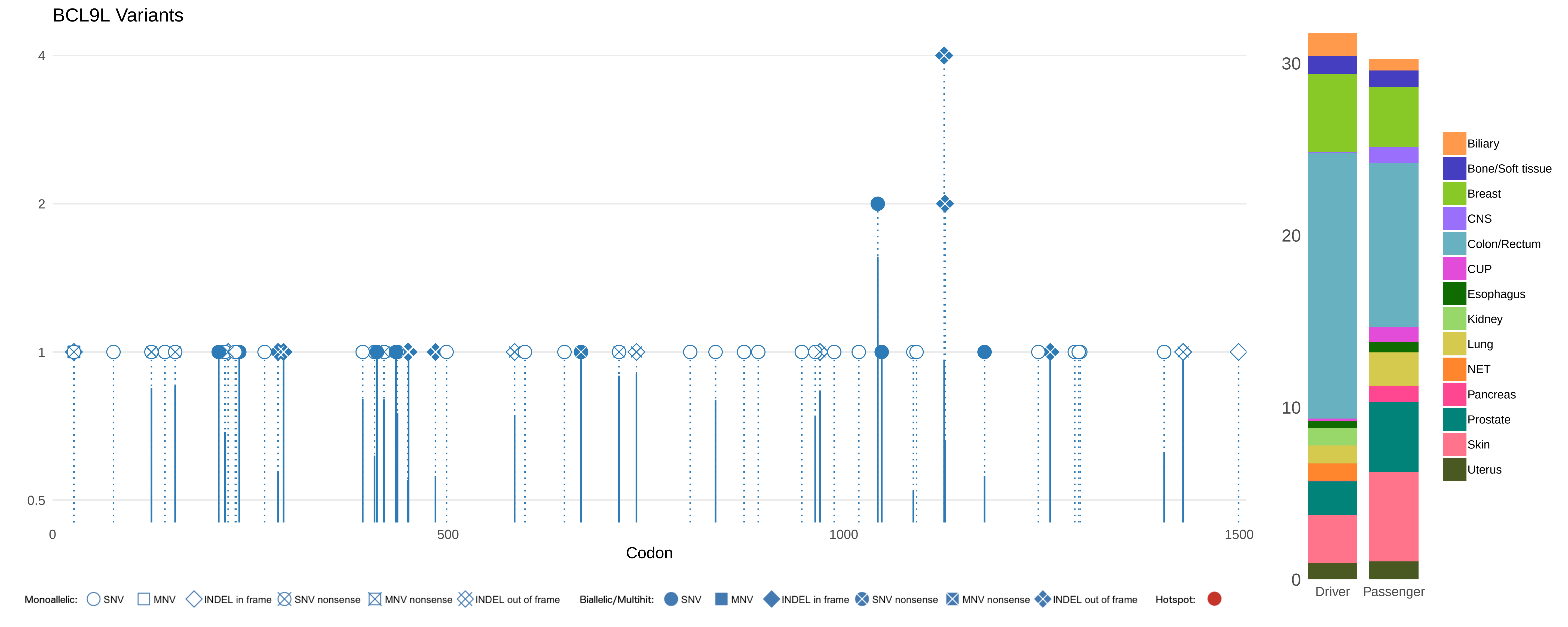

BCL9L Variants
4
2
1
0.5
0
500
1000
1500
Codon
30
20
10
0
Driver
Passenger
Biliary
Bone/Soft tissue
Breast
CNS
Colon/Rectum
CUP
Esophagus
Kidney
Lung
NET
Pancreas
Prostate
Skin
Uterus
Monoallelic: SNV MNV INDEL in frame SNV nonsense MNV nonsense INDEL out of frame
Biallelic/Multihit: SNV MNV INDEL in frame SNV nonsense MNV nonsense INDEL out of frame
Hotspot:

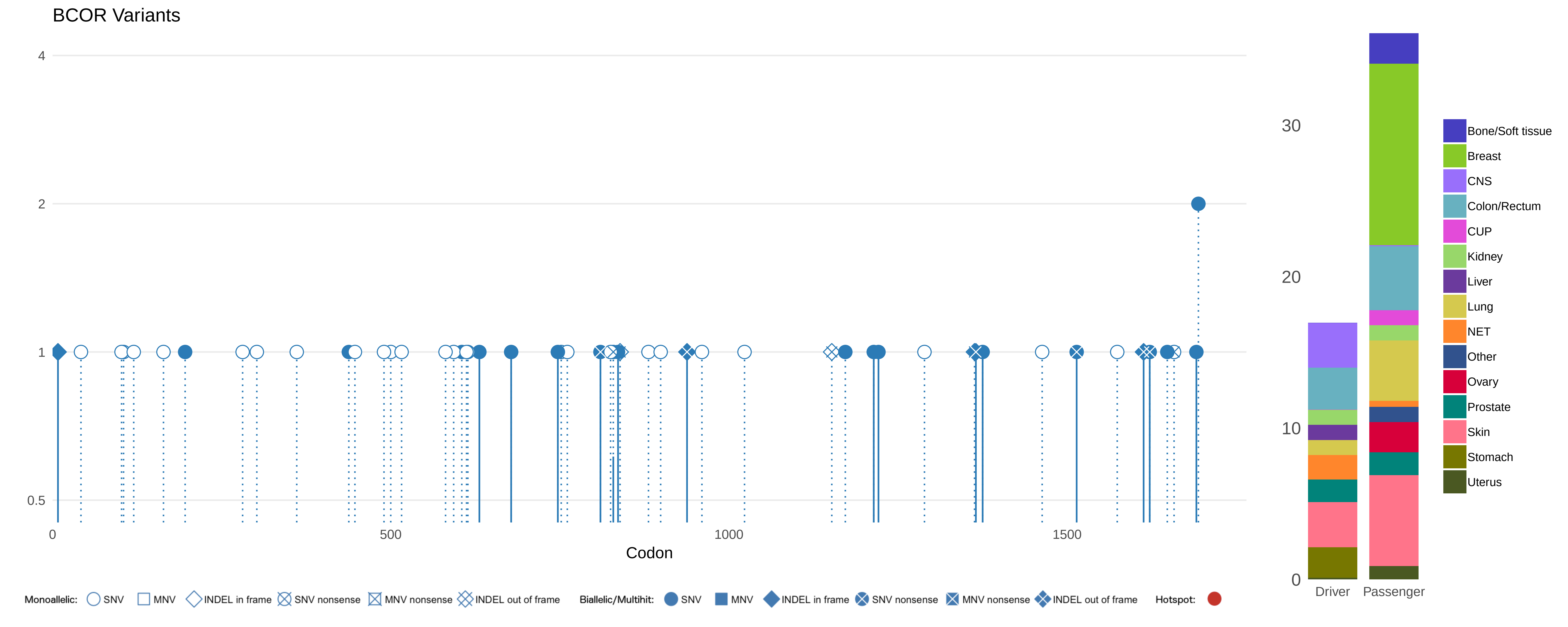
BCOR Variants
4
2
1
0.5
0
500
1000
1500
Codon
30
20
10
0
Driver
Passenger
Bone/Soft tissue
Breast
CNS
Colon/Rectum
CUP
Kidney
Liver
Lung
NET
Other
Ovary
Prostate
Skin
Stomach
Uterus
Monoallelic: SNV MNV INDEL in frame SNV nonsense MNV nonsense INDEL out of frame
Biallelic/Multihit: SNV MNV INDEL in frame SNV nonsense MNV nonsense INDEL out of frame
Hotspot:

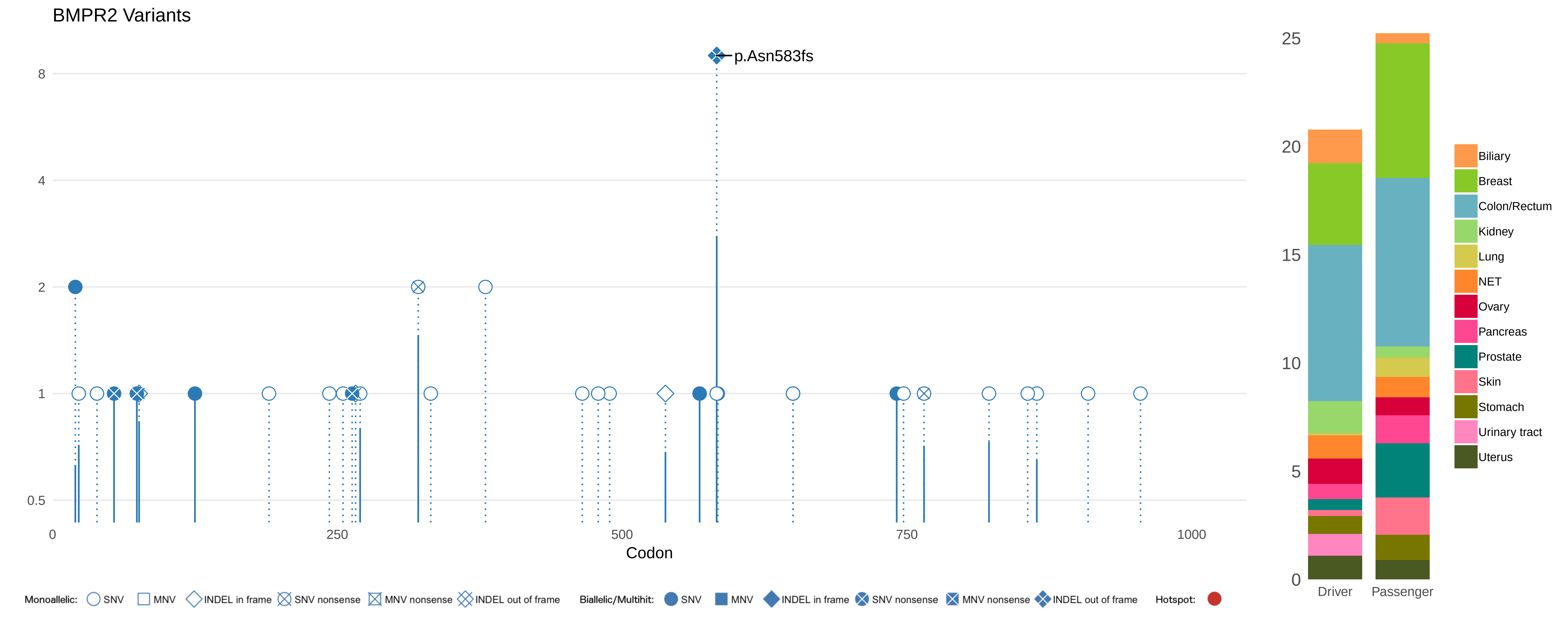
BMPR2 Variants
p.Asn583fs
8
4
2
1
0.5
0
250
500
750
1000
Codon
25
20
15
10
5
0
Driver
Passenger
Biliary
Breast
Colon/Rectum
Kidney
Lung
NET
Ovary
Pancreas
Prostate
Skin
Stomach
Urinary tract
Uterus
Monoallelic: SNV MNV INDEL in frame SNV nonsense MNV nonsense INDEL out of frame
Biallelic/Multihit: SNV MNV INDEL in frame SNV nonsense MNV nonsense INDEL out of frame
Hotspot:

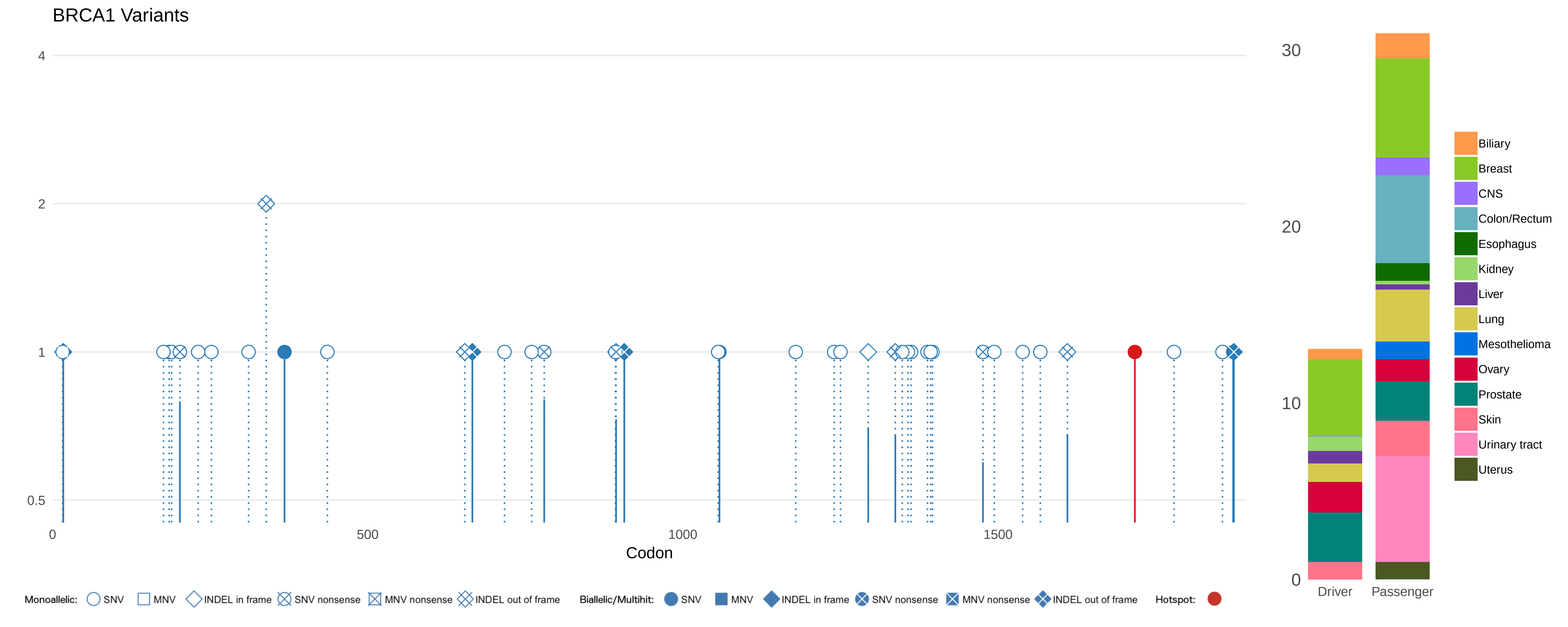
BRCA1 Variants
4
2
1
0.5
0
500
1000
1500
Codon
30
20
10
0
Driver
Passenger
Biliary
Breast
CNS
Colon/Rectum
Esophagus
Kidney
Liver
Lung
Mesothelioma
Ovary
Prostate
Skin
Urinary tract
Uterus
Monoallelic: SNV MNV INDEL in frame SNV nonsense MNV nonsense INDEL out of frame
Biallelic/Multihit: SNV MNV INDEL in frame SNV nonsense MNV nonsense INDEL out of frame
Hotspot:

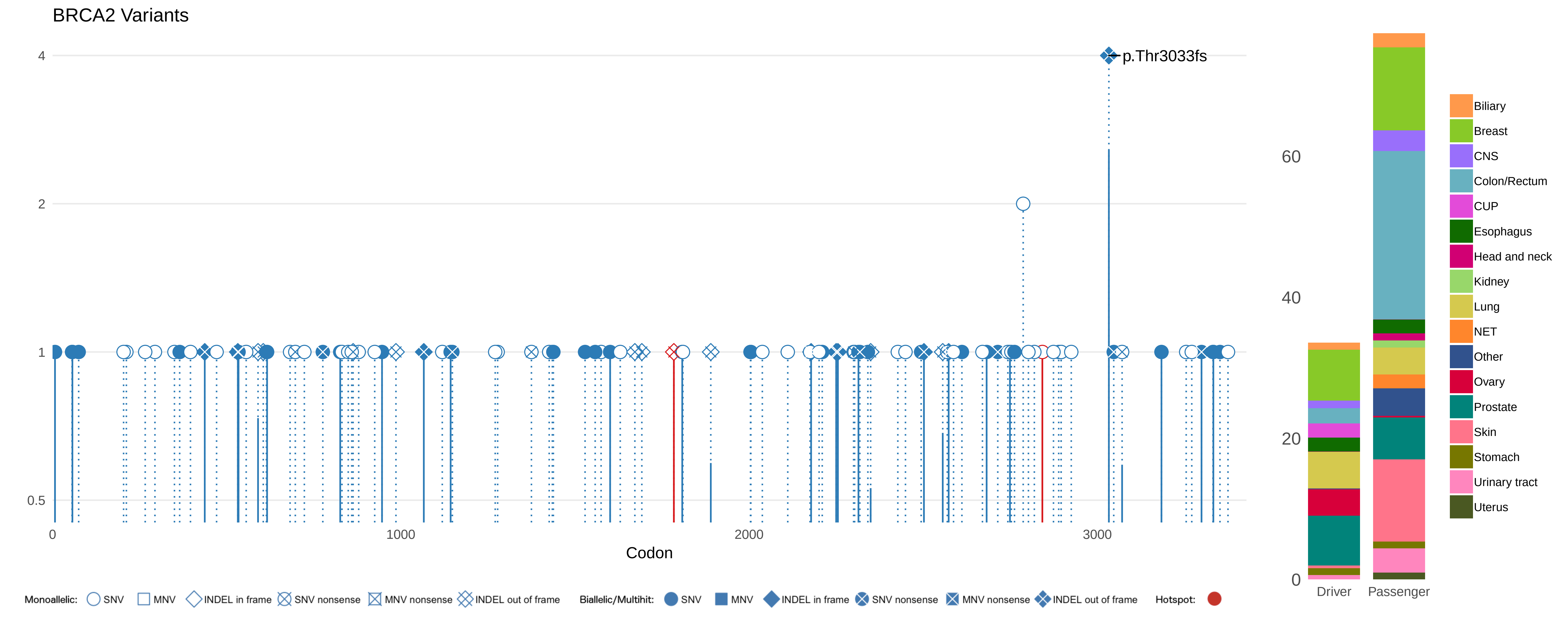

BRCA2 Variants
p.Thr3033fs
4
2
1
0.5
0
1000
2000
3000
Codon
60
40
20
0
Driver
Passenger
Biliary
Breast
CNS
Colon/Rectum
CUP
Esophagus
Head and neck
Kidney
Lung
NET
Other
Ovary
Prostate
Skin
Stomach
Urinary tract
Uterus
Monoallelic: SNV MNV INDEL in frame SNV nonsense MNV nonsense INDEL out of frame
Biallelic/Multihit: SNV MNV INDEL in frame SNV nonsense MNV nonsense INDEL out of frame
Hotspot:

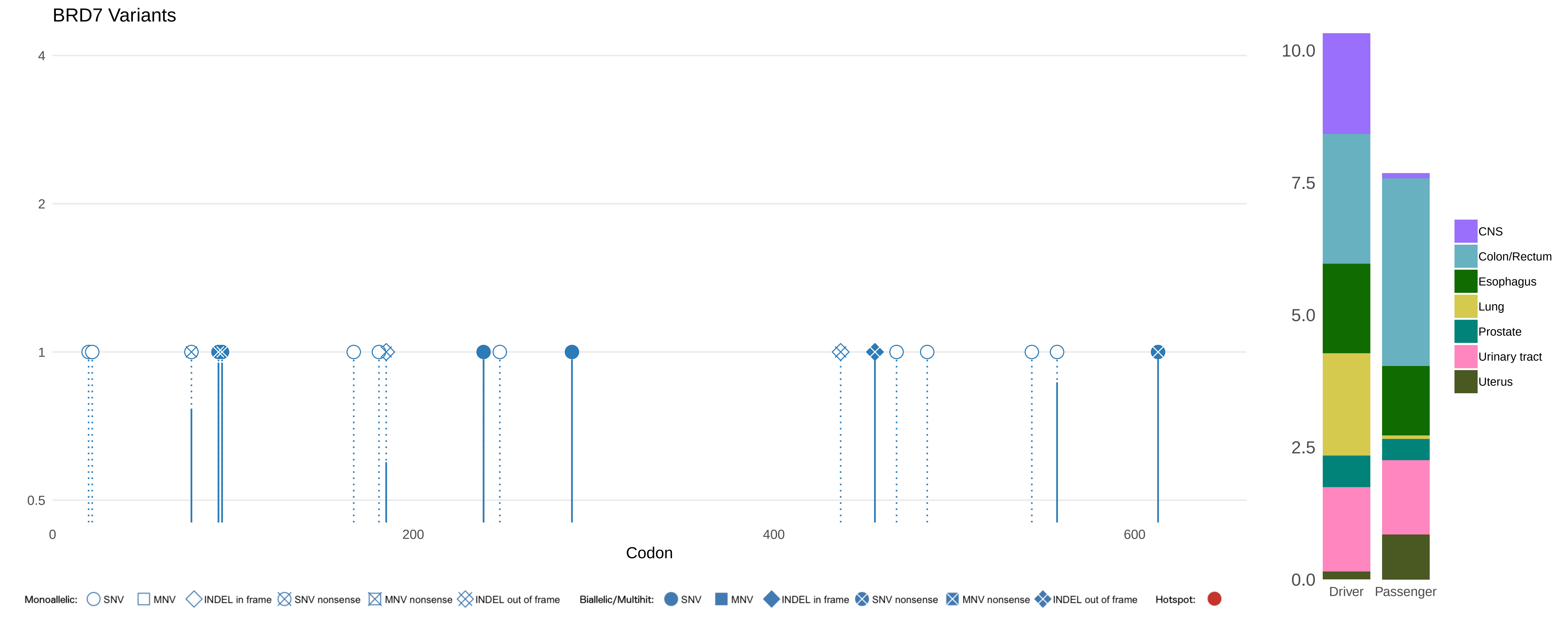

BRD7 Variants
4
2
1
0.5
0
200
400
600
Codon
10.0
7.5
5.0
2.5
0.0
Driver
Passenger
CNS
Colon/Rectum
Esophagus
Lung
Prostate
Urinary tract
Uterus
Monoallelic: SNV MNV INDEL in frame SNV nonsense MNV nonsense INDEL out of frame
Biallelic/Multihit: SNV MNV INDEL in frame SNV nonsense MNV nonsense INDEL out of frame
Hotspot:

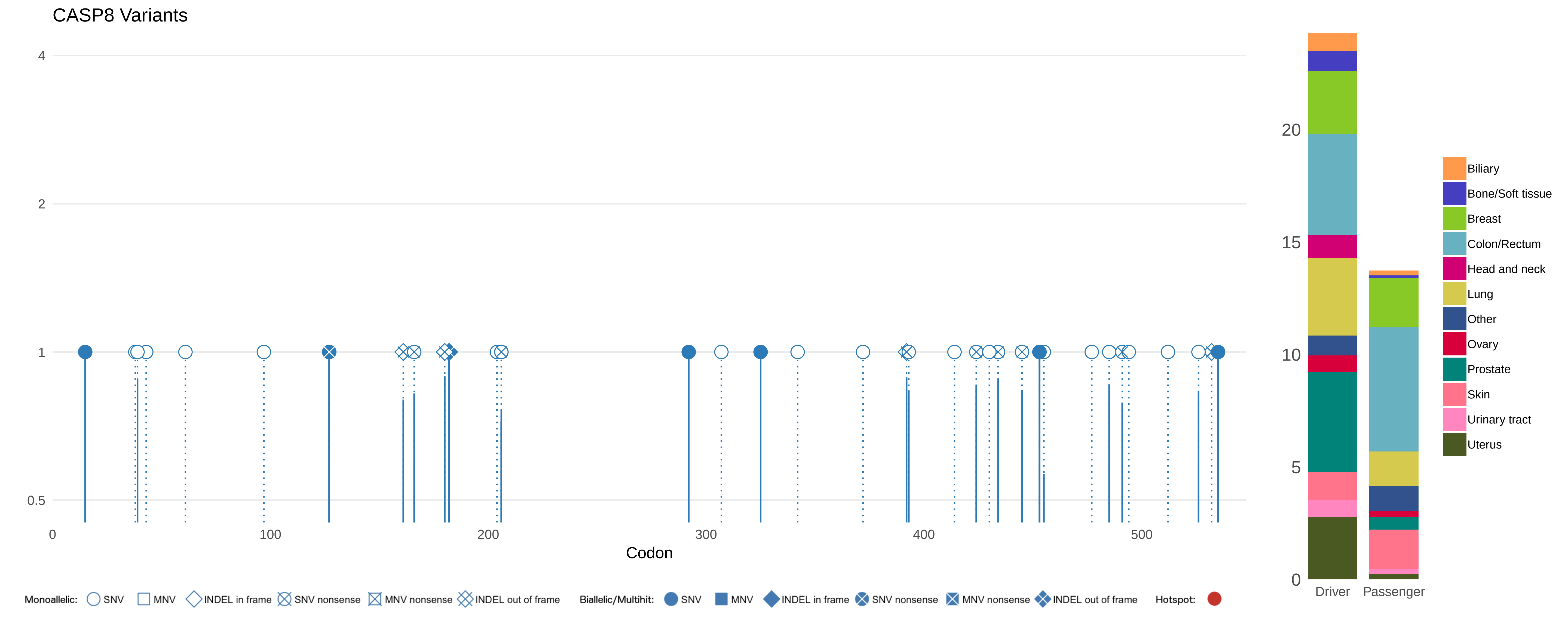
CASP8 Variants
4
2
1
0.5
0
100
200
300
400
500
Codon
20
15
10
5
0
Driver
Passenger
Biliary
Bone/Soft tissue
Breast
Colon/Rectum
Head and neck
Lung
Other
Ovary
Prostate
Skin
Urinary tract
Uterus
Monoallelic: SNV MNV INDEL in frame SNV nonsense MNV nonsense INDEL out of frame
Biallelic/Multihit: SNV MNV INDEL in frame SNV nonsense MNV nonsense INDEL out of frame
Hotspot:

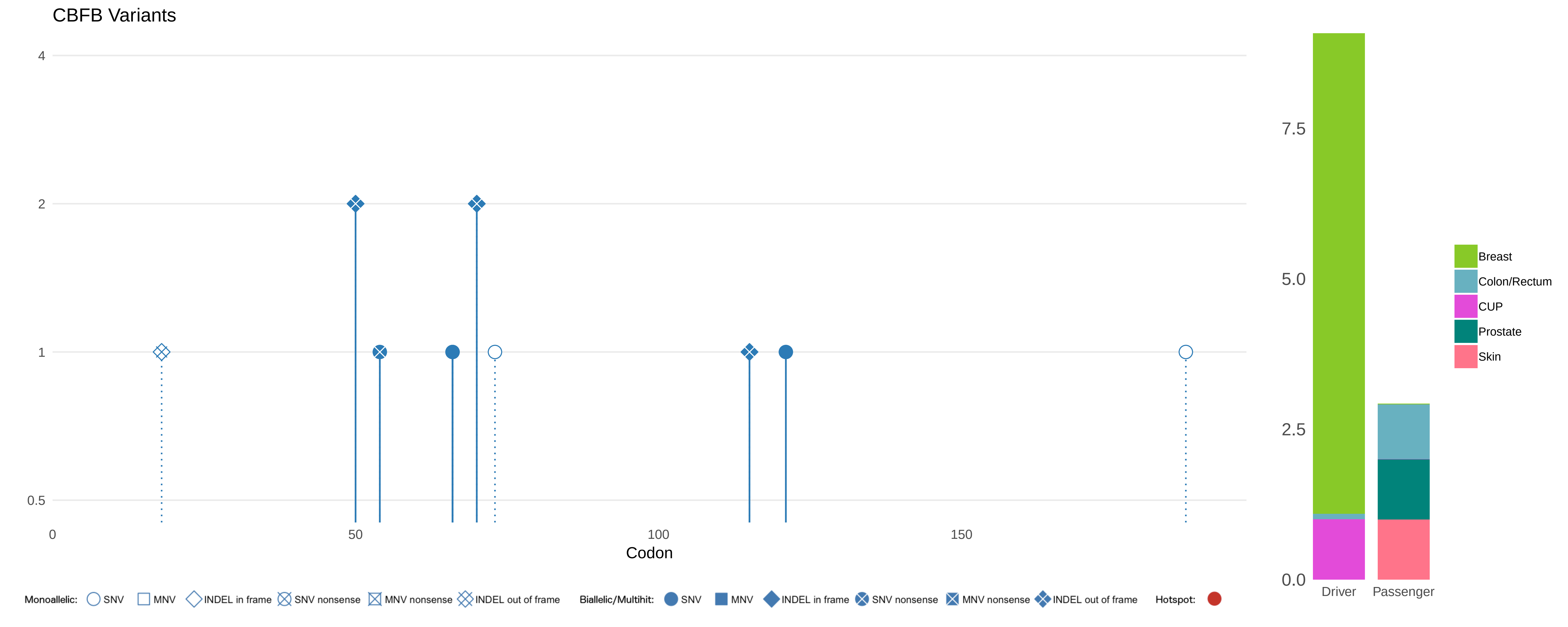
CBFB Variants
4
2
1
0.5
0
50
100
150
Codon
7.5
5.0
2.5
0.0
Driver
Passenger
Breast
Colon/Rectum
CUP
Prostate
Skin
Monoallelic: SNV MNV INDEL in frame SNV nonsense MNV nonsense INDEL out of frame
Biallelic/Multihit: SNV MNV INDEL in frame SNV nonsense MNV nonsense INDEL out of frame
Hotspot:

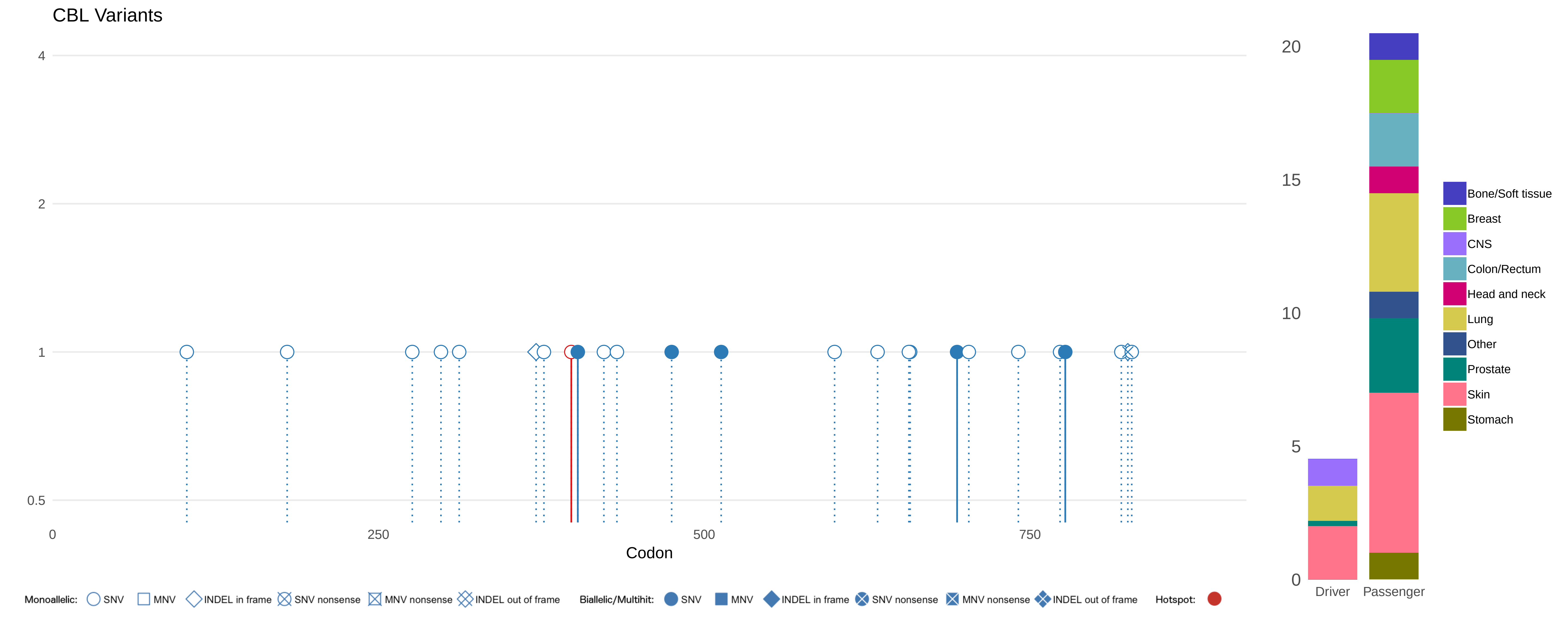

CBL Variants
4
2
1
0.5
0
250
500
750
Codon
20
15
10
5
0
Driver
Passenger
Bone/Soft tissue
Breast
CNS
Colon/Rectum
Head and neck
Lung
Other
Prostate
Skin
Stomach
Monoallelic: SNV MNV INDEL in frame SNV nonsense MNV nonsense INDEL out of frame
Biallelic/Multihit: SNV MNV INDEL in frame SNV nonsense MNV nonsense INDEL out of frame
Hotspot:

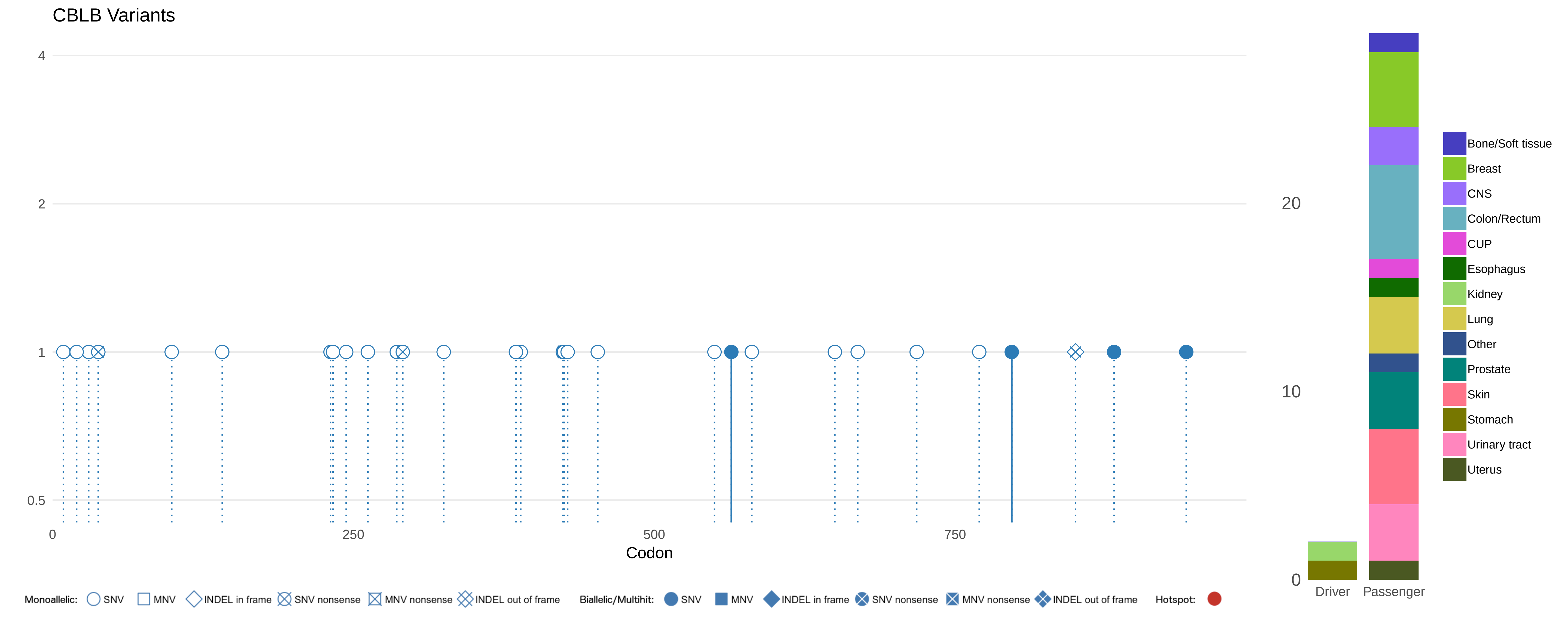
CBLB Variants
4
2
1
0.5
0
250
500
750
Codon
20
10
0
Driver
Passenger
Bone/Soft tissue
Breast
CNS
Colon/Rectum
CUP
Esophagus
Kidney
Lung
Other
Prostate
Skin
Stomach
Urinary tract
Uterus
Monoallelic: SNV MNV INDEL in frame SNV nonsense MNV nonsense INDEL out of frame
Biallelic/Multihit: SNV MNV INDEL in frame SNV nonsense MNV nonsense INDEL out of frame
Hotspot:

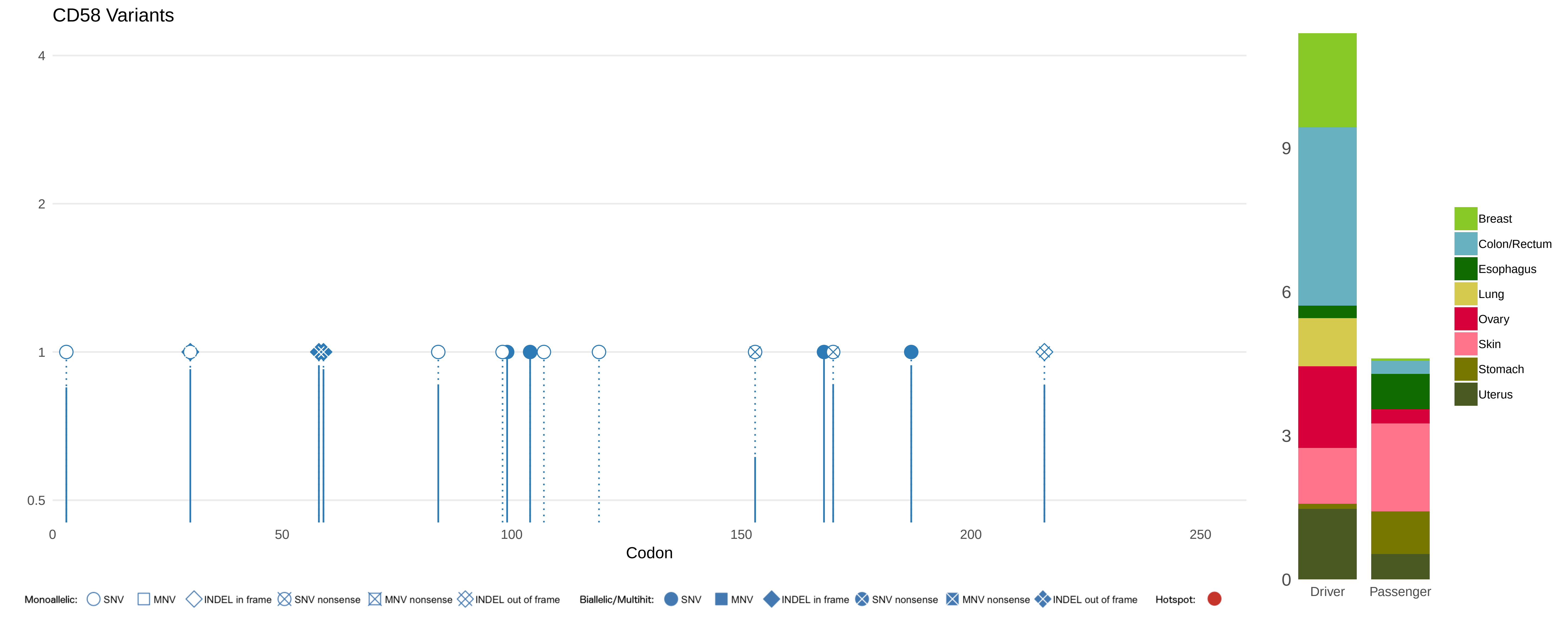

CD58 Variants
4
2
1
0.5
0
50
100
150
200
250
Codon
Monoallelic: SNV MNV INDEL in frame SNV nonsense MNV nonsense INDEL out of frame
Biallelic/Multihit: SNV MNV INDEL in frame SNV nonsense MNV nonsense INDEL out of frame
Hotspot:
9
6
3
0
Driver
Passenger
Breast
Colon/Rectum
Esophagus
Lung
Ovary
Skin
Stomach
Uterus

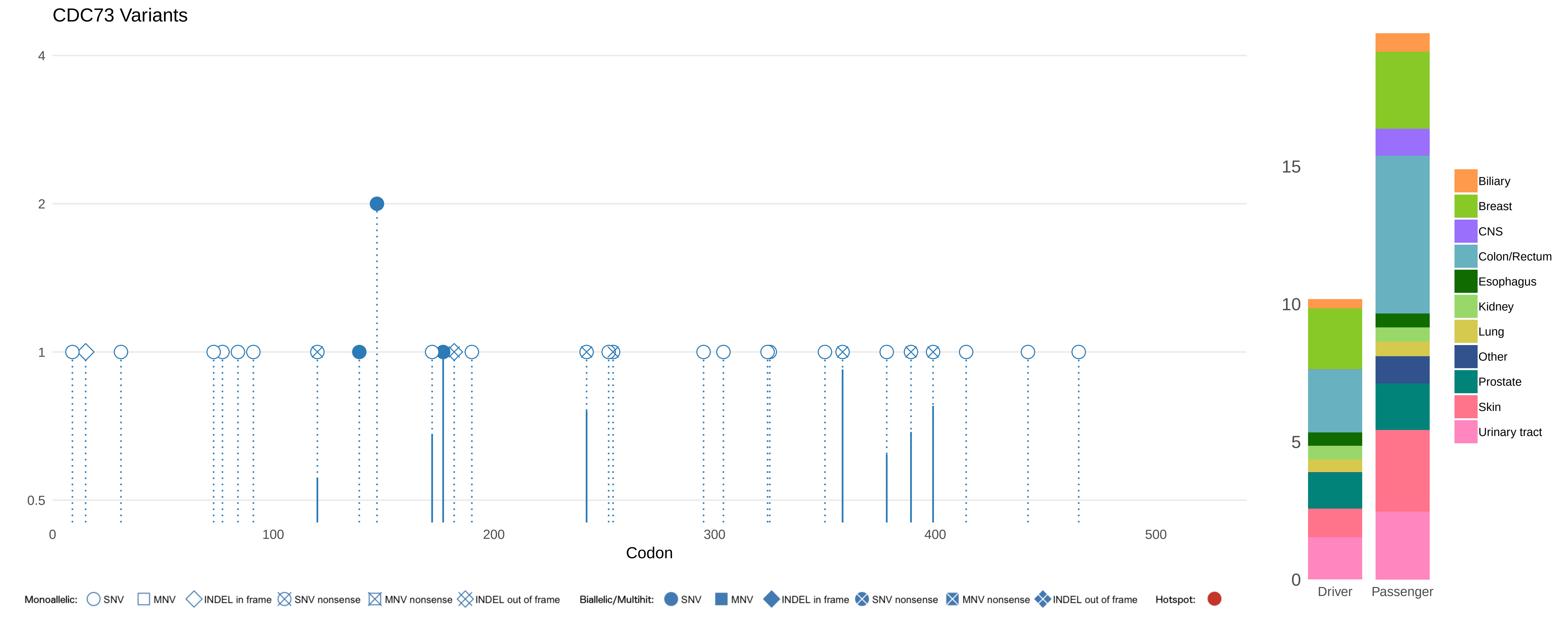

CDC73 Variants
4
2
1
0.5
0
100
200
300
400
500
Codon
Monoallelic: SNV MNV INDEL in frame SNV nonsense MNV nonsense INDEL out of frame
Biallelic/Multihit: SNV MNV INDEL in frame SNV nonsense MNV nonsense INDEL out of frame
Hotspot:
15
10
5
0
Driver
Passenger
Biliary
Breast
CNS
Colon/Rectum
Esophagus
Kidney
Lung
Other
Prostate
Skin
Urinary tract

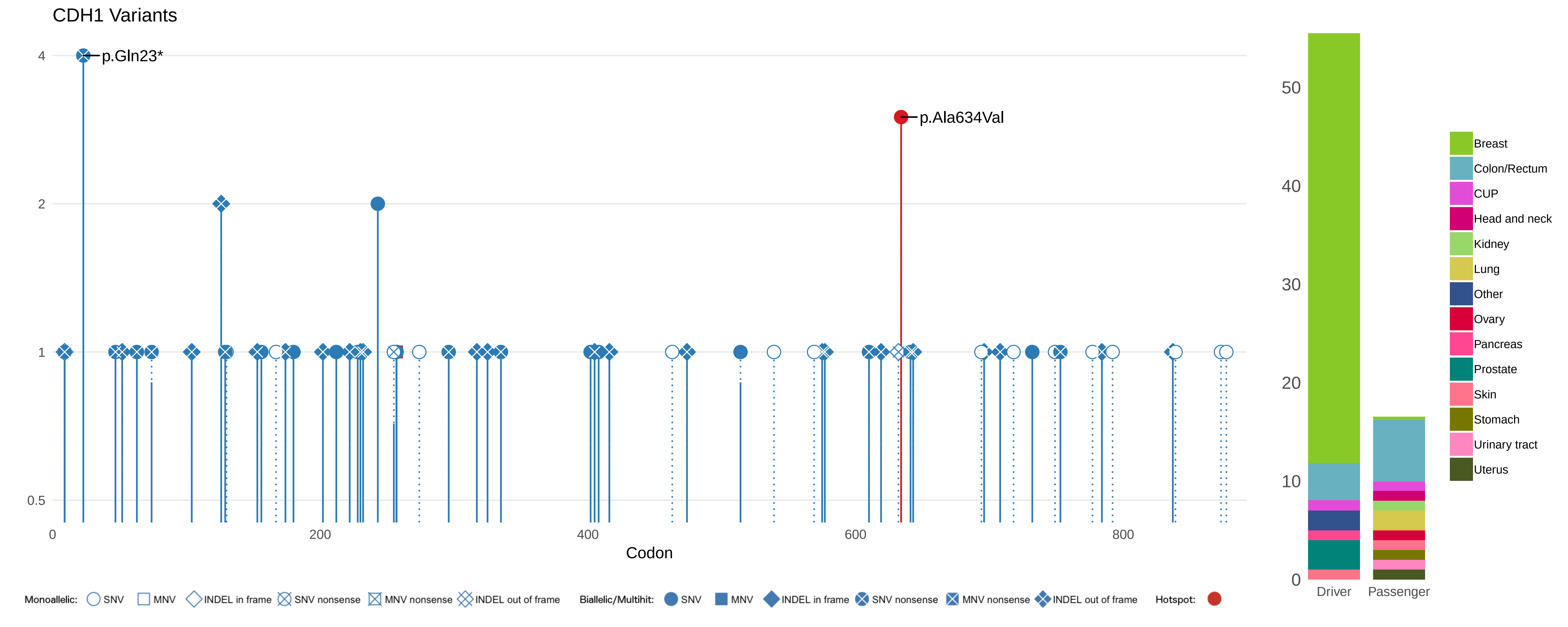

CDH1 Variants
p.Gln23*
p.Ala634Val
4
2
1
0.5
0
200
400
600
800
Codon
Monoallelic: SNV MNV INDEL in frame SNV nonsense MNV nonsense INDEL out of frame
Biallelic/Multihit: SNV MNV INDEL in frame SNV nonsense MNV nonsense INDEL out of frame
Hotspot:
50
40
30
20
10
0
Driver
Passenger
Breast
Colon/Rectum
CUP
Head and neck
Kidney
Lung
Other
Ovary
Pancreas
Prostate
Skin
Stomach
Urinary tract
Uterus

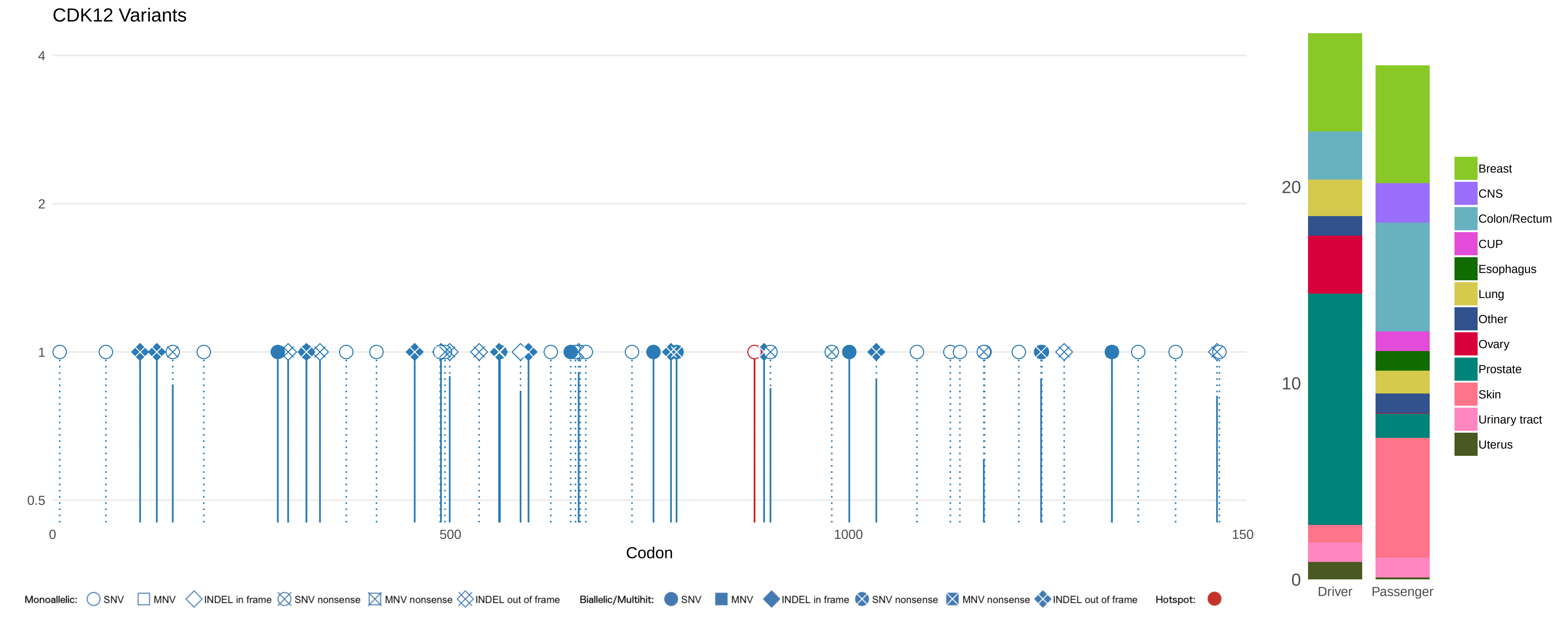
CDK12 Variants
4
2
1
0.5
0
500
1000
150
Codon
20
10
0
Driver
Passenger
Breast
CNS
Colon/Rectum
CUP
Esophagus
Lung
Other
Ovary
Prostate
Skin
Urinary tract
Uterus
Monoallelic: SNV MNV INDEL in frame SNV nonsense MNV nonsense INDEL out of frame
Biallelic/Multihit: SNV MNV INDEL in frame SNV nonsense MNV nonsense INDEL out of frame
Hotspot:

### CDKN1A Variants

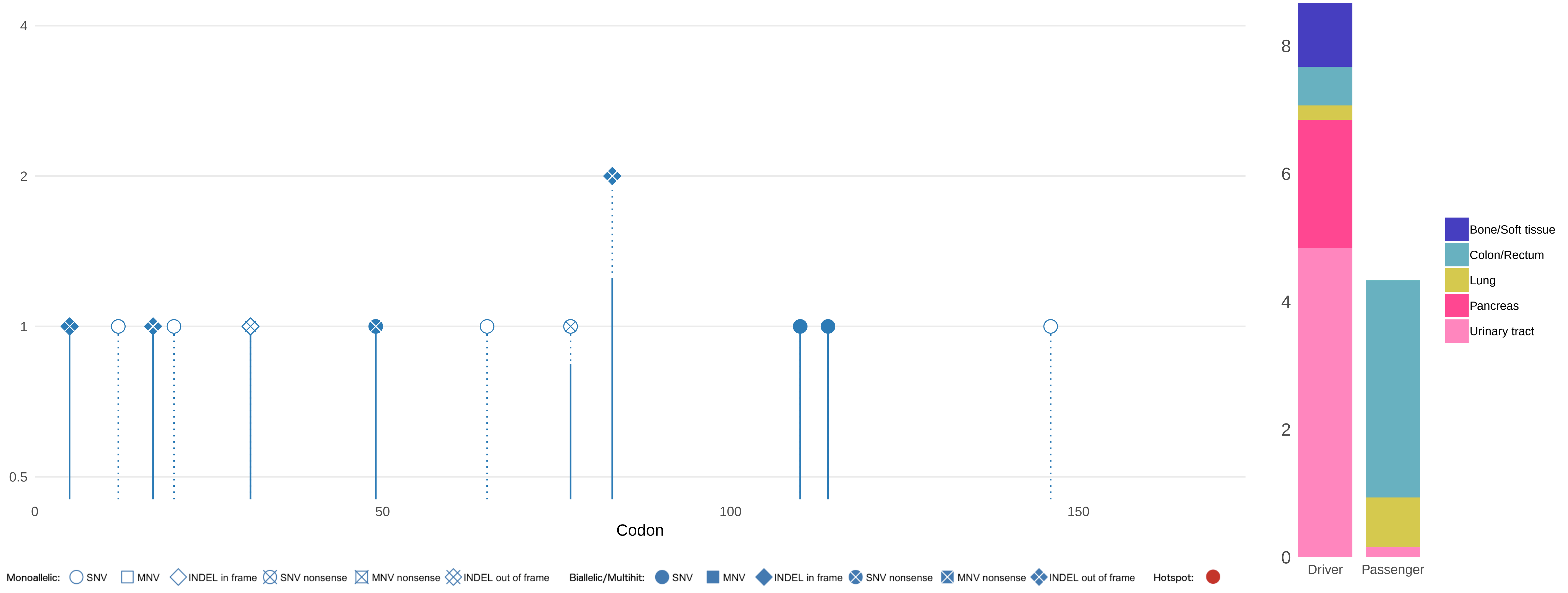

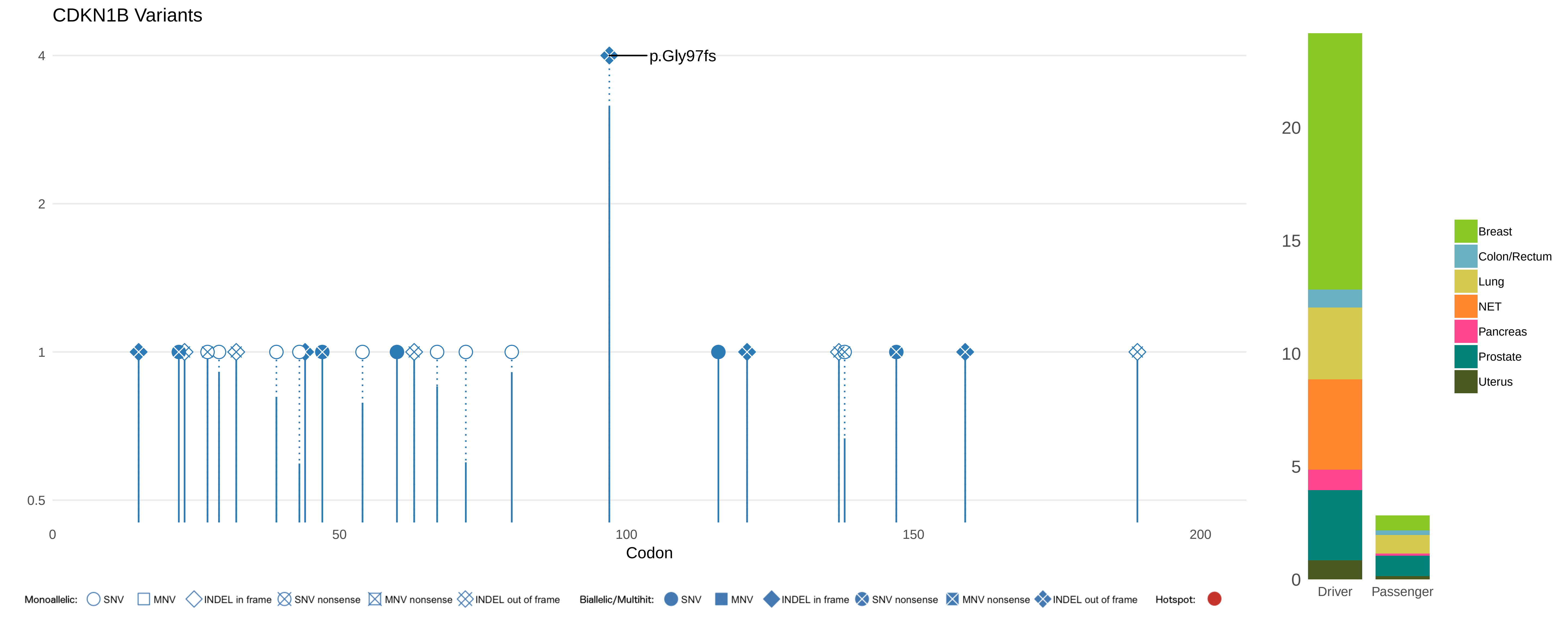

CDKN1B Variants
p.Gly97fs
4
2
1
0.5
0
50
100
150
200
Codon
20
15
10
5
0
Driver
Passenger
Breast
Colon/Rectum
Lung
NET
Pancreas
Prostate
Uterus
Monoallelic:
SNV
MNV
INDEL in frame
SNV nonsense
MNV nonsense
INDEL out of frame
Biallelic/Multihit:
SNV
MNV
INDEL in frame
SNV nonsense
MNV nonsense
INDEL out of frame
Hotspot:

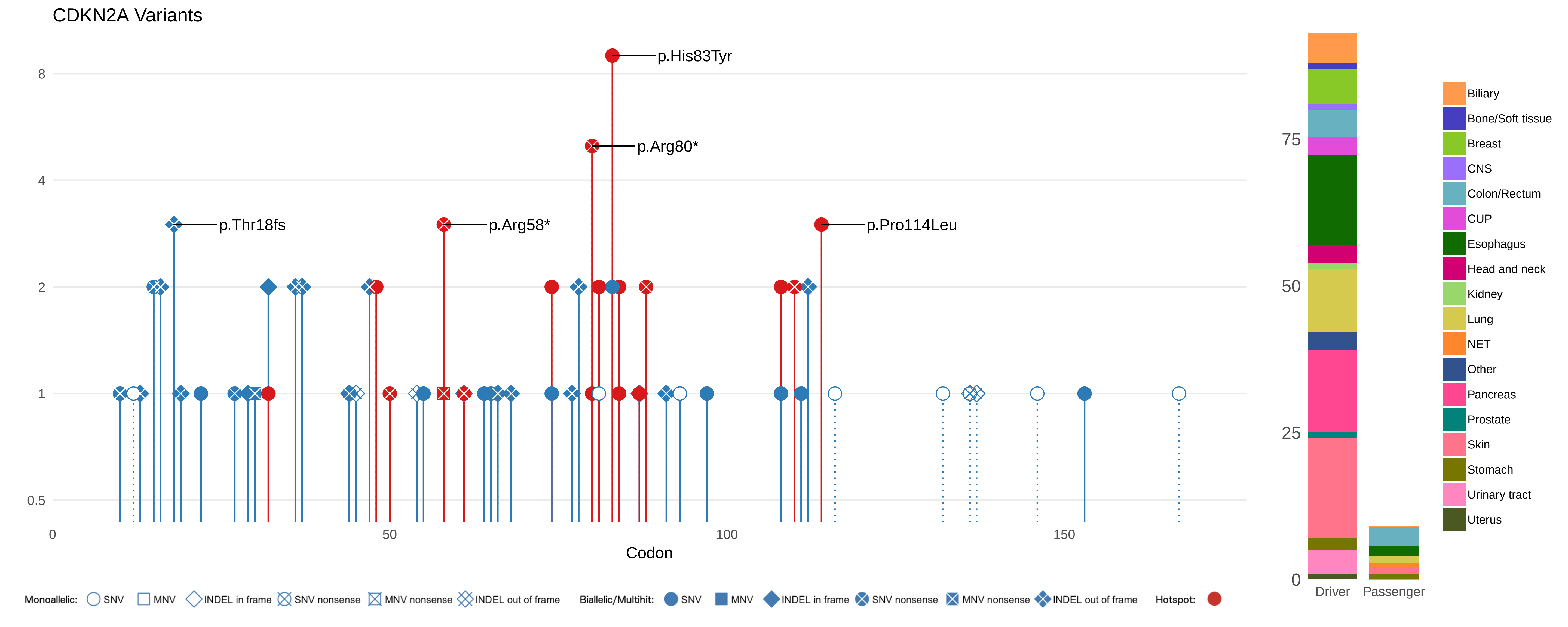

CDKN2A Variants
p.His83Tyr
p.Arg80*
p.Thr18fs
p.Arg58*
p.Pro114Leu
8
4
2
1
0.5
0
50
100
150
Codon
75
50
25
0
Driver
Passenger
Biliary
Bone/Soft tissue
Breast
CNS
Colon/Rectum
CUP
Esophagus
Head and neck
Kidney
Lung
NET
Other
Pancreas
Prostate
Skin
Stomach
Urinary tract
Uterus
Monoallelic: SNV MNV INDEL in frame SNV nonsense MNV nonsense INDEL out of frame
Biallelic/Multihit: SNV MNV INDEL in frame SNV nonsense MNV nonsense INDEL out of frame
Hotspot:

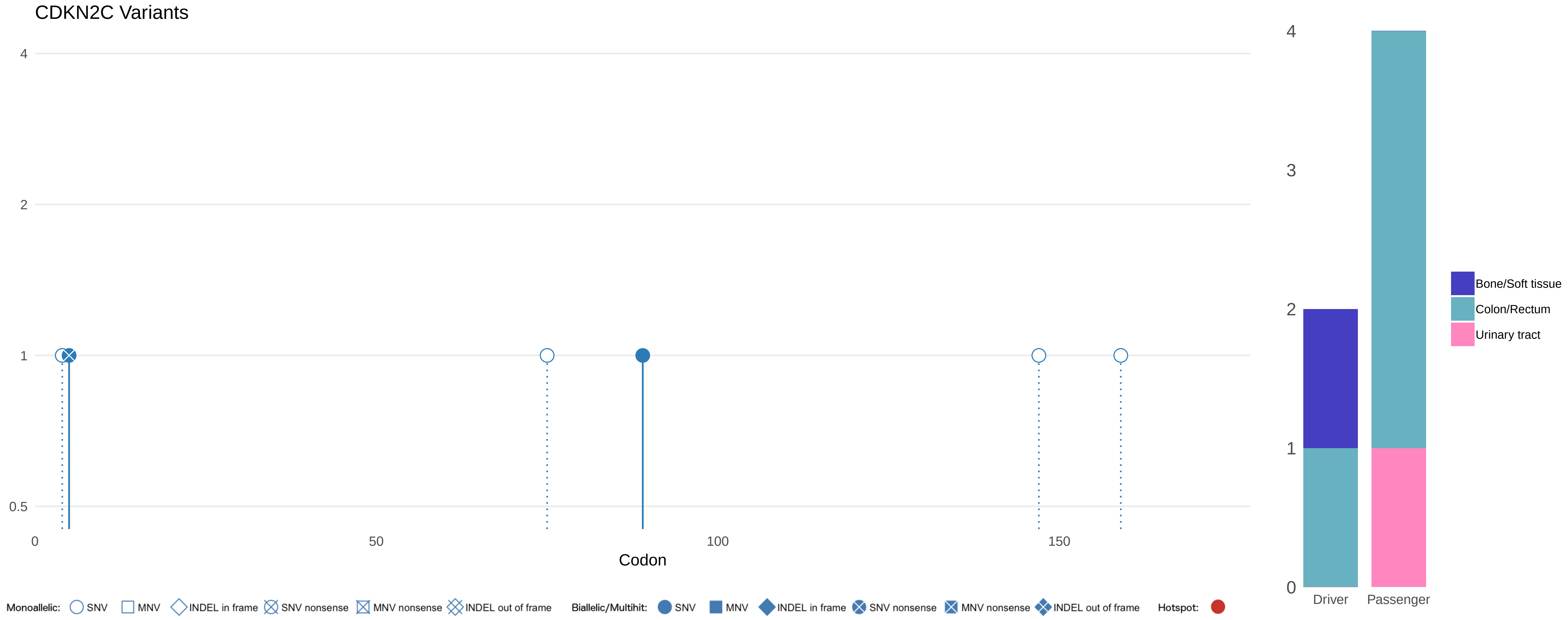

CDKN2C Variants
4
2
1
0.5
0
50
100
150
Codon
4
3
2
1
0
Driver
Passenger
Bone/Soft tissue
Colon/Rectum
Urinary tract
Monoallelic: SNV MNV INDEL in frame SNV nonsense MNV nonsense INDEL out of frame
Biallelic/Multihit: SNV MNV INDEL in frame SNV nonsense MNV nonsense INDEL out of frame
Hotspot:

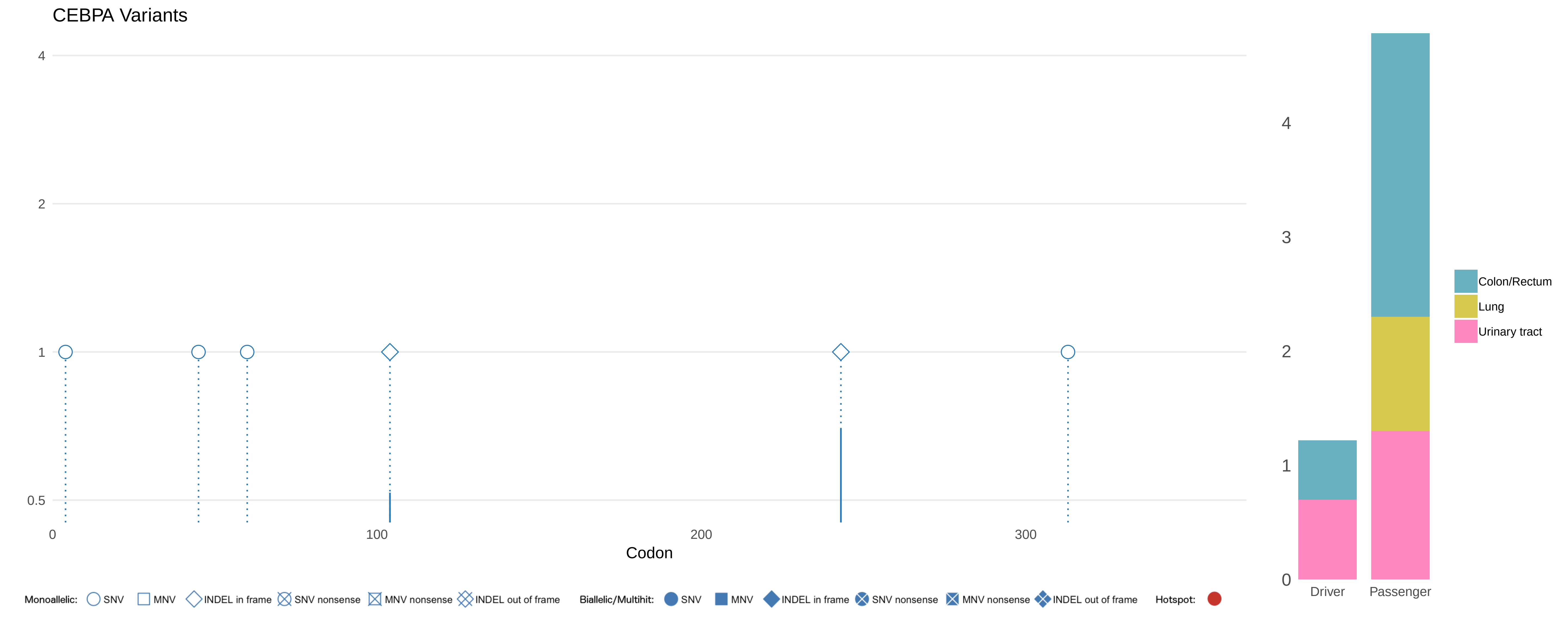
CEBPA Variants
4
2
1
0.5
0
100
200
300
Codon
Monoallelic: SNV MNV INDEL in frame SNV nonsense MNV nonsense INDEL out of frame
Biallelic/Multihit: SNV MNV INDEL in frame SNV nonsense MNV nonsense INDEL out of frame
Hotspot:
4
3
2
1
0
Driver
Passenger
Colon/Rectum
Lung
Urinary tract

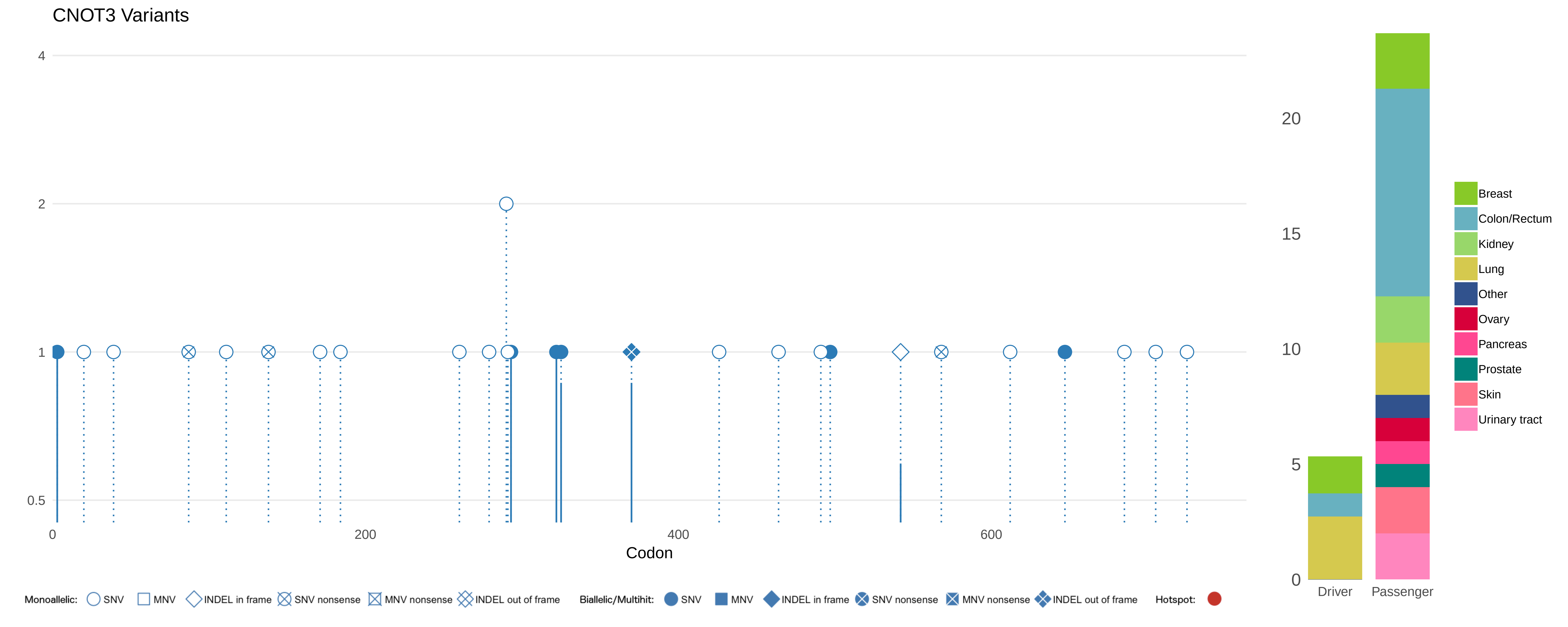
CNOT3 Variants
4
2
1
0.5
0
200
400
600
Codon
20
15
10
5
0
Driver
Passenger
Breast
Colon/Rectum
Kidney
Lung
Other
Ovary
Pancreas
Prostate
Skin
Urinary tract
Monoallelic: SNV MNV INDEL in frame SNV nonsense MNV nonsense INDEL out of frame
Biallelic/Multihit: SNV MNV INDEL in frame SNV nonsense MNV nonsense INDEL out of frame
Hotspot:

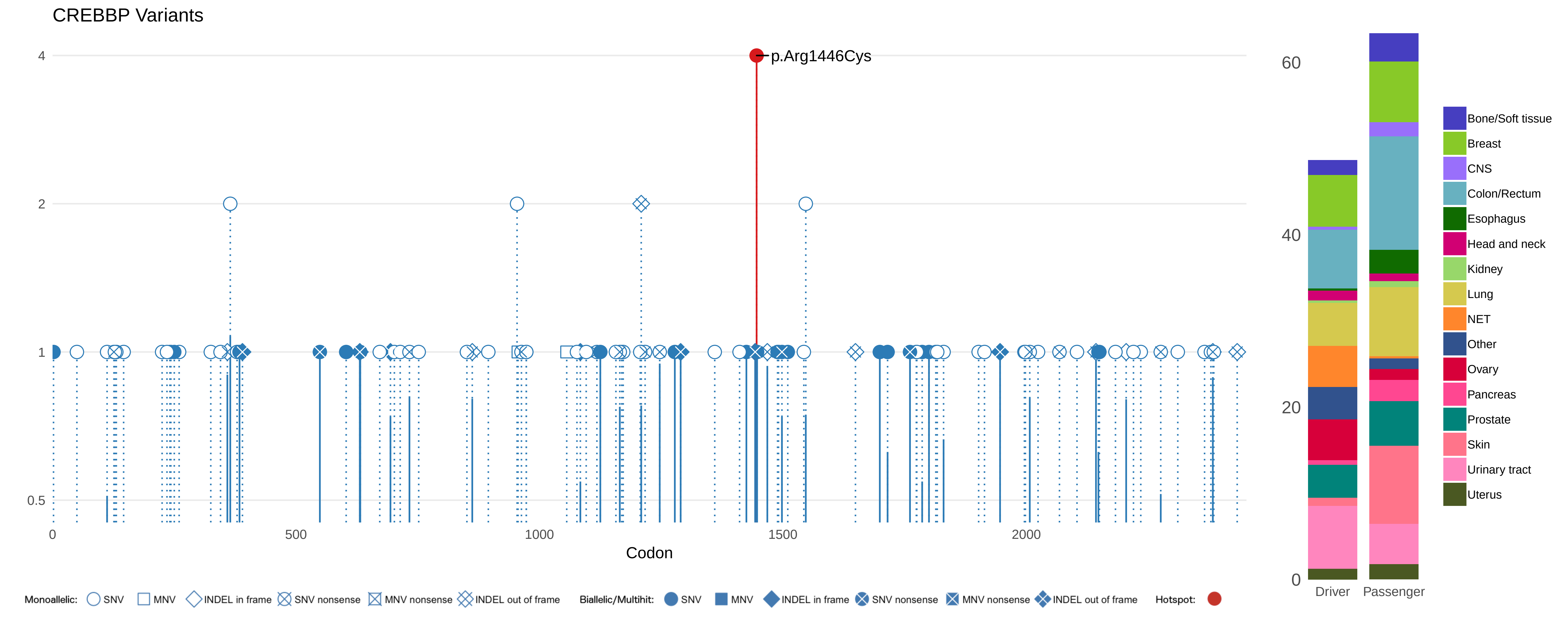

CREBBP Variants
p.Arg1446Cys
4
2
1
0.5
0
500
1000
1500
2000
Codon
60
40
20
0
Driver
Passenger
Bone/Soft tissue
Breast
CNS
Colon/Rectum
Esophagus
Head and neck
Kidney
Lung
NET
Other
Ovary
Pancreas
Prostate
Skin
Urinary tract
Uterus
Monoallelic: SNV MNV INDEL in frame SNV nonsense MNV nonsense INDEL out of frame
Biallelic/Multihit: SNV MNV INDEL in frame SNV nonsense MNV nonsense INDEL out of frame
Hotspot:

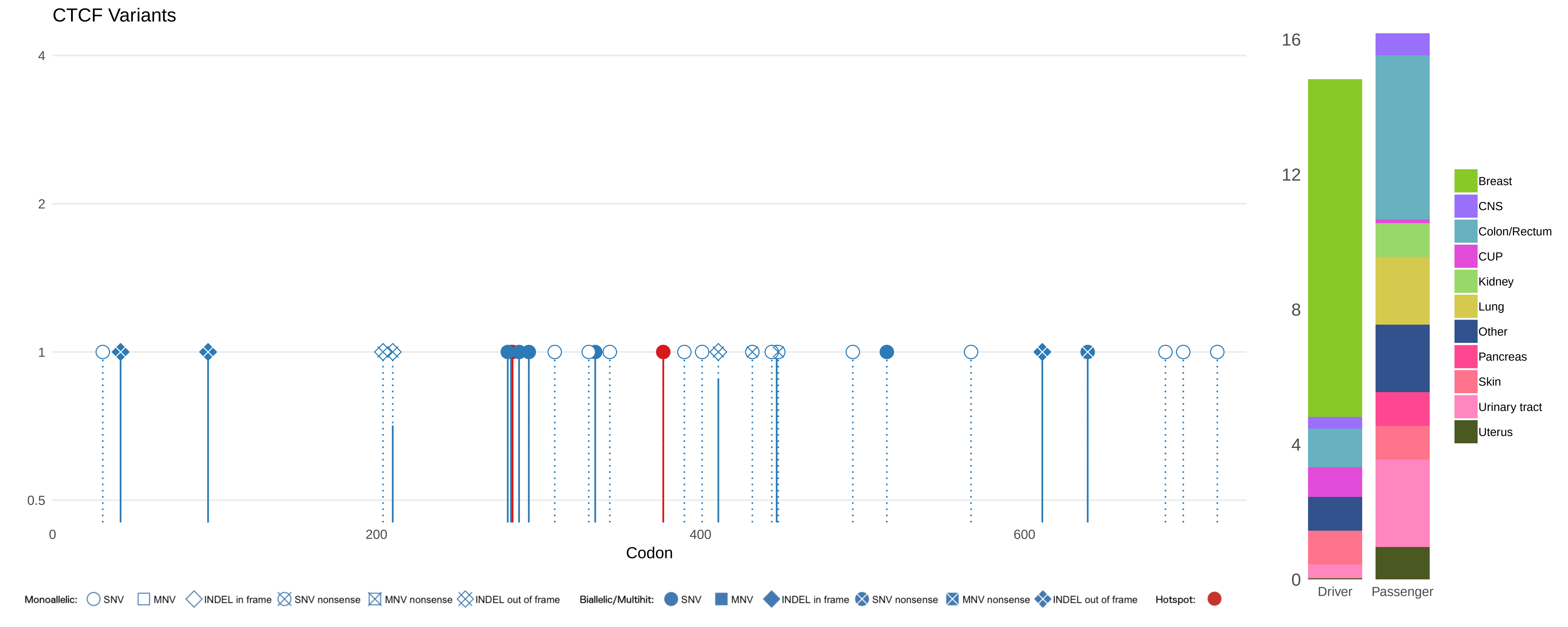

CTCF Variants
4
2
1
0.5
0
200
400
600
Codon
16
12
8
4
0
Driver
Passenger
Breast
CNS
Colon/Rectum
CUP
Kidney
Lung
Other
Pancreas
Skin
Urinary tract
Uterus
Monoallelic: SNV MNV INDEL in frame SNV nonsense MNV nonsense INDEL out of frame
Biallelic/Multihit: SNV MNV INDEL in frame SNV nonsense MNV nonsense INDEL out of frame
Hotspot:

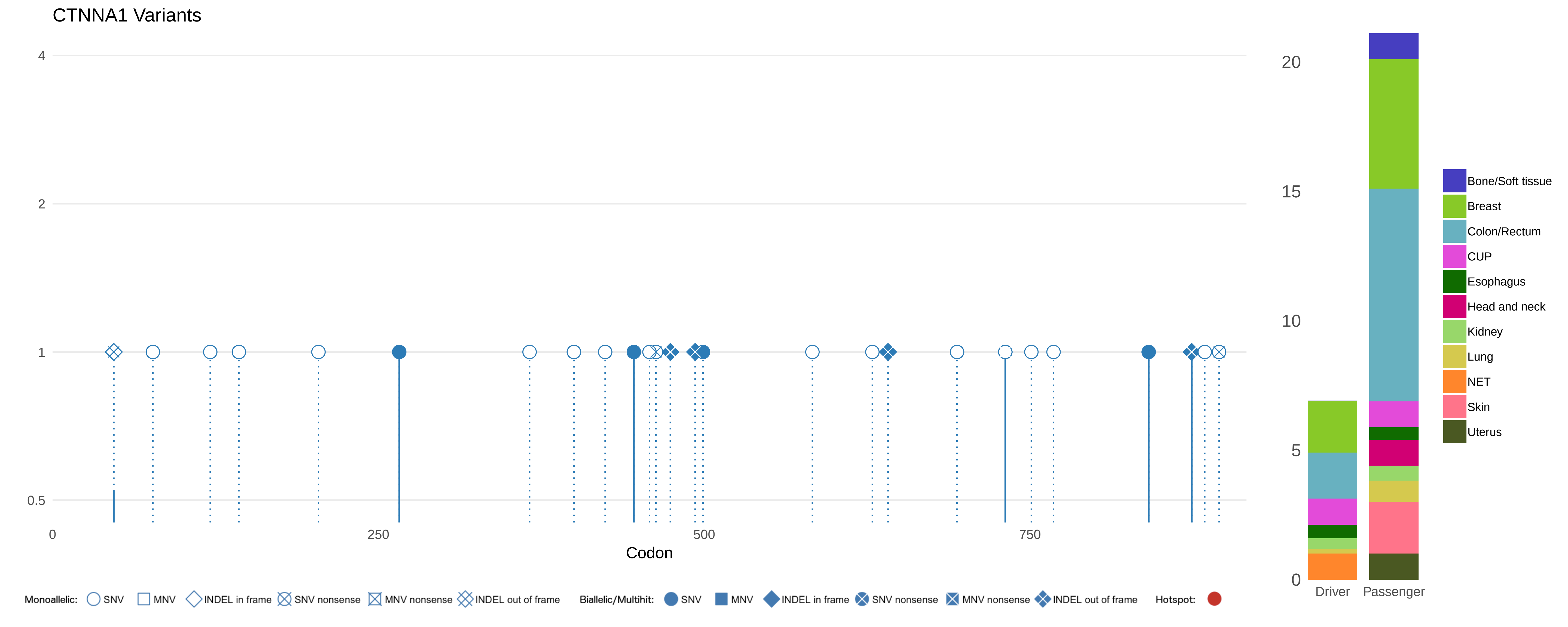
CTNNA1 Variants
4
2
1
0.5
0
250
500
750
Codon
20
15
10
5
0
Driver
Passenger
Bone/Soft tissue
Breast
Colon/Rectum
CUP
Esophagus
Head and neck
Kidney
Lung
NET
Skin
Uterus
Monoallelic: SNV MNV INDEL in frame SNV nonsense MNV nonsense INDEL out of frame
Biallelic/Multihit: SNV MNV INDEL in frame SNV nonsense MNV nonsense INDEL out of frame
Hotspot:

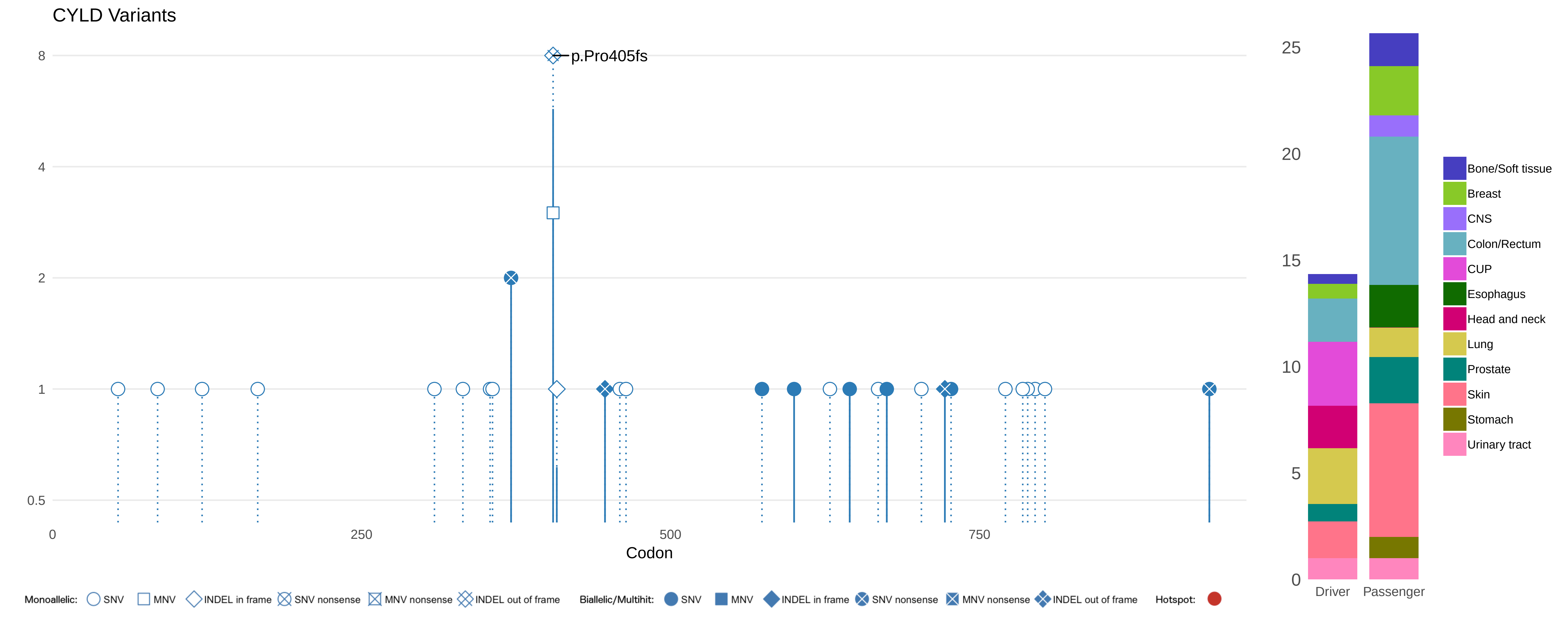

CYLD Variants
p.Pro405fs
8
4
2
1
0.5
0
250
500
750
Codon
25
20
15
10
5
0
Driver
Passenger
Bone/Soft tissue
Breast
CNS
Colon/Rectum
CUP
Esophagus
Head and neck
Lung
Prostate
Skin
Stomach
Urinary tract
Monoallelic: SNV MNV INDEL in frame SNV nonsense MNV nonsense INDEL out of frame
Biallelic/Multihit: SNV MNV INDEL in frame SNV nonsense MNV nonsense INDEL out of frame
Hotspot:

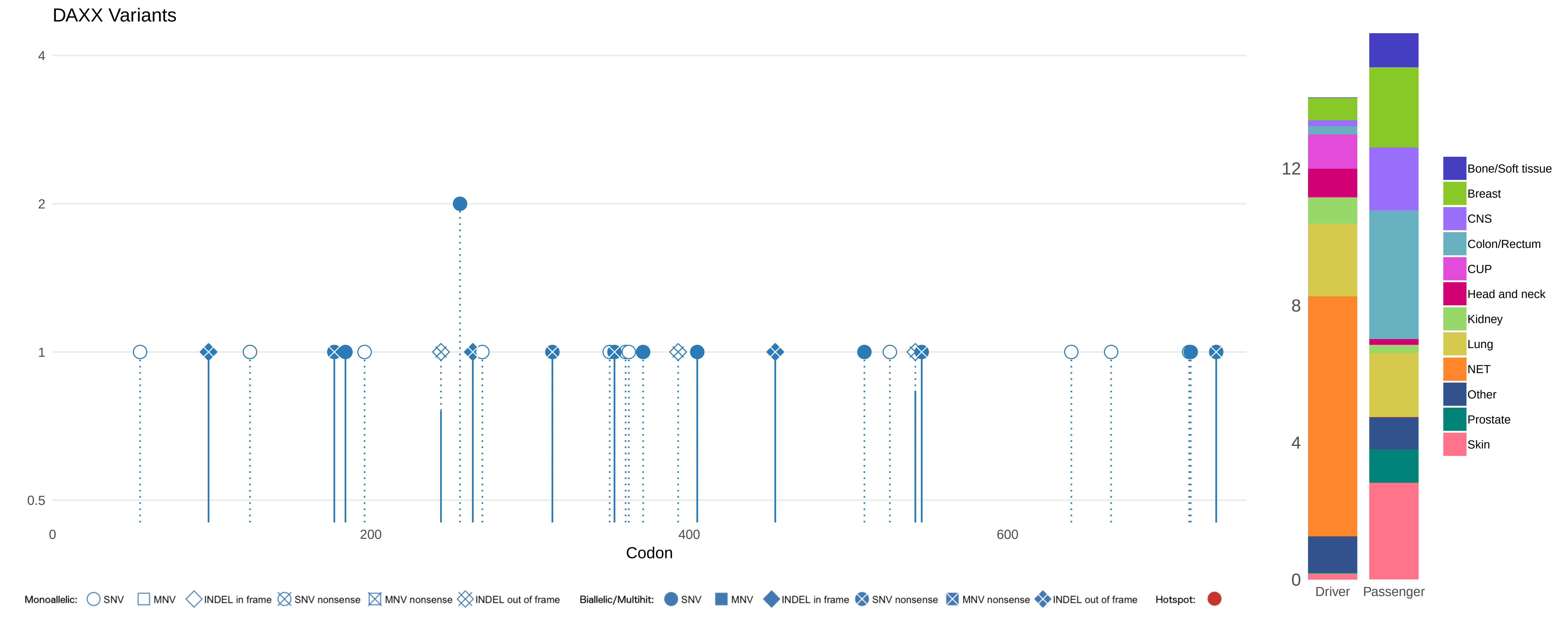

DAXX Variants
4
2
1
0.5
0
200
400
600
Codon
Monoallelic: SNV MNV INDEL in frame SNV nonsense MNV nonsense INDEL out of frame
Biallelic/Multihit: SNV MNV INDEL in frame SNV nonsense MNV nonsense INDEL out of frame
Hotspot:
12
8
4
0
Driver
Passenger
Bone/Soft tissue
Breast
CNS
Colon/Rectum
CUP
Head and neck
Kidney
Lung
NET
Other
Prostate
Skin

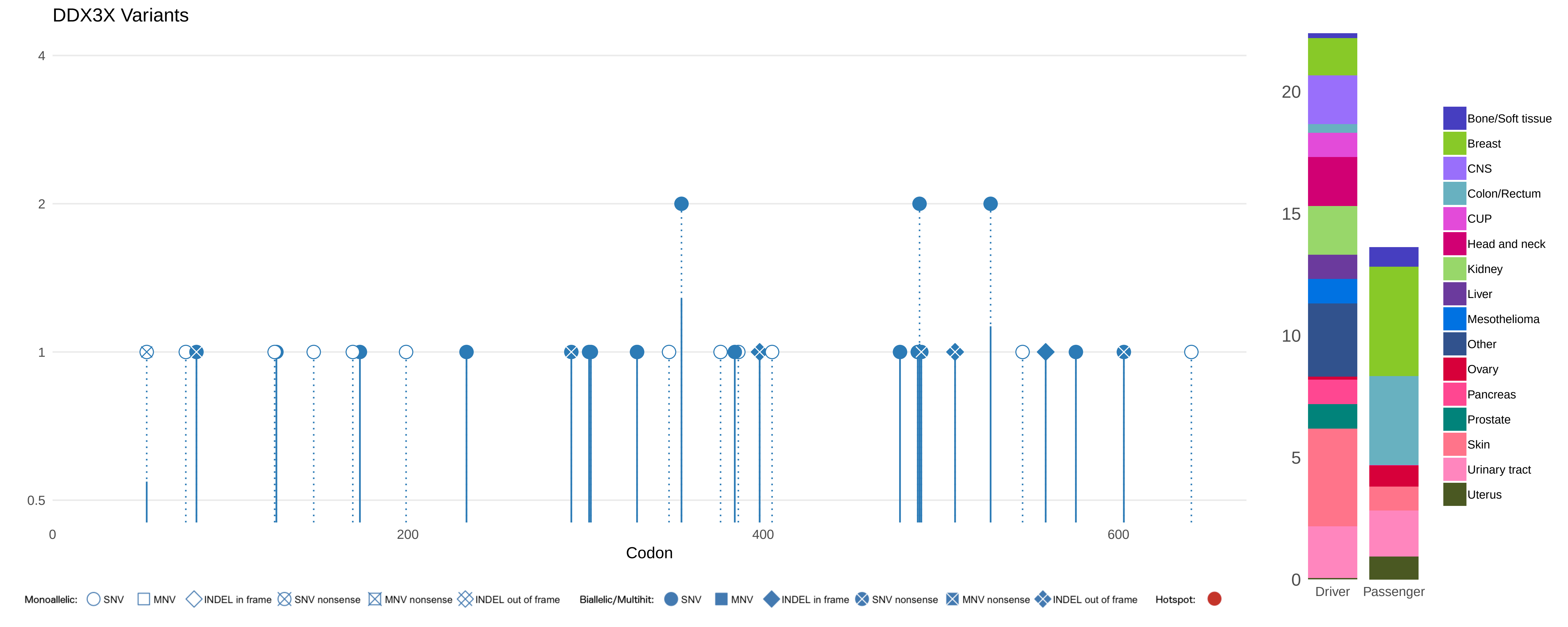

DDX3X Variants
4
2
1
0.5
0
200
400
600
Codon
Monoallelic: SNV MNV INDEL in frame SNV nonsense MNV nonsense INDEL out of frame
Biallelic/Multihit: SNV MNV INDEL in frame SNV nonsense MNV nonsense INDEL out of frame
Hotspot:
20
15
10
5
0
Driver
Passenger
Bone/Soft tissue
Breast
CNS
Colon/Rectum
CUP
Head and neck
Kidney
Liver
Mesothelioma
Other
Ovary
Pancreas
Prostate
Skin
Urinary tract
Uterus

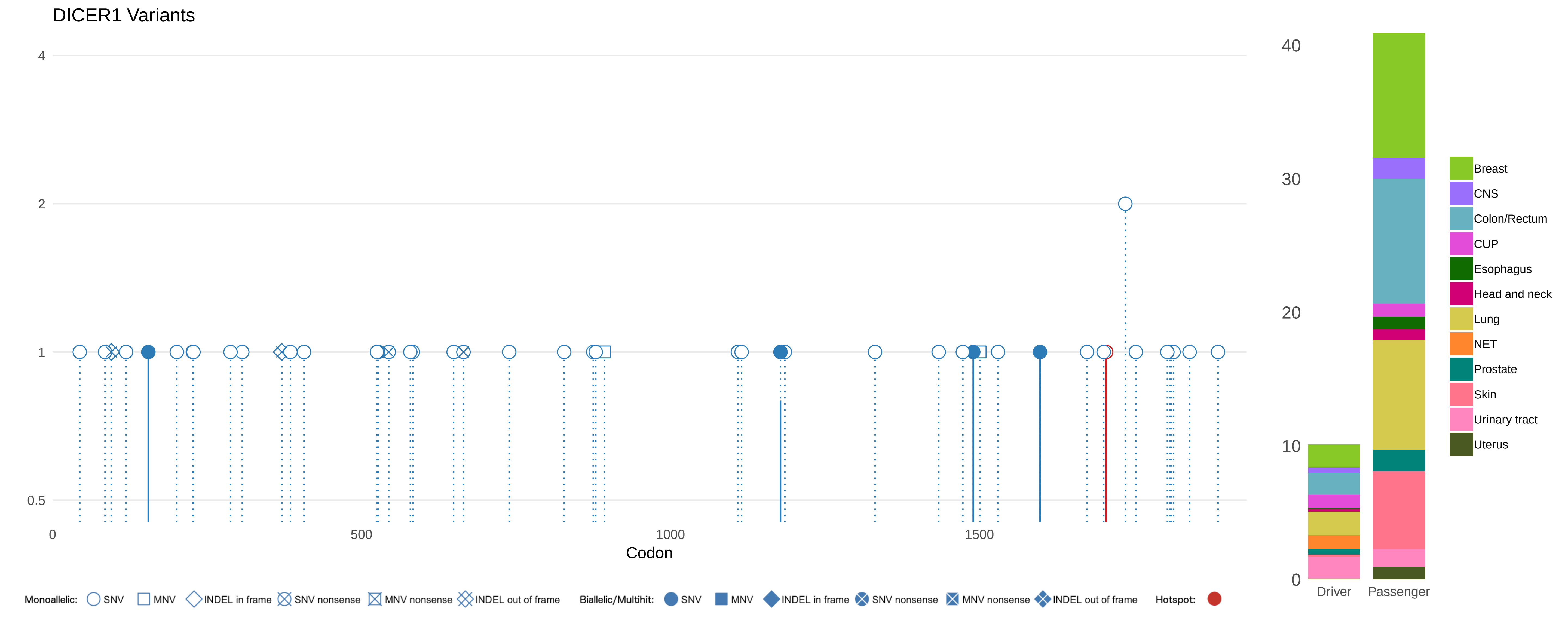

DICER1 Variants
4
2
1
0.5
0
500
1000
1500
Codon
Monoallelic: SNV MNV INDEL in frame SNV nonsense MNV nonsense INDEL out of frame
Biallelic/Multihit: SNV MNV INDEL in frame SNV nonsense MNV nonsense INDEL out of frame
Hotspot:
40
30
20
10
0
Driver
Passenger
Breast
CNS
Colon/Rectum
CUP
Esophagus
Head and neck
Lung
NET
Prostate
Skin
Urinary tract
Uterus

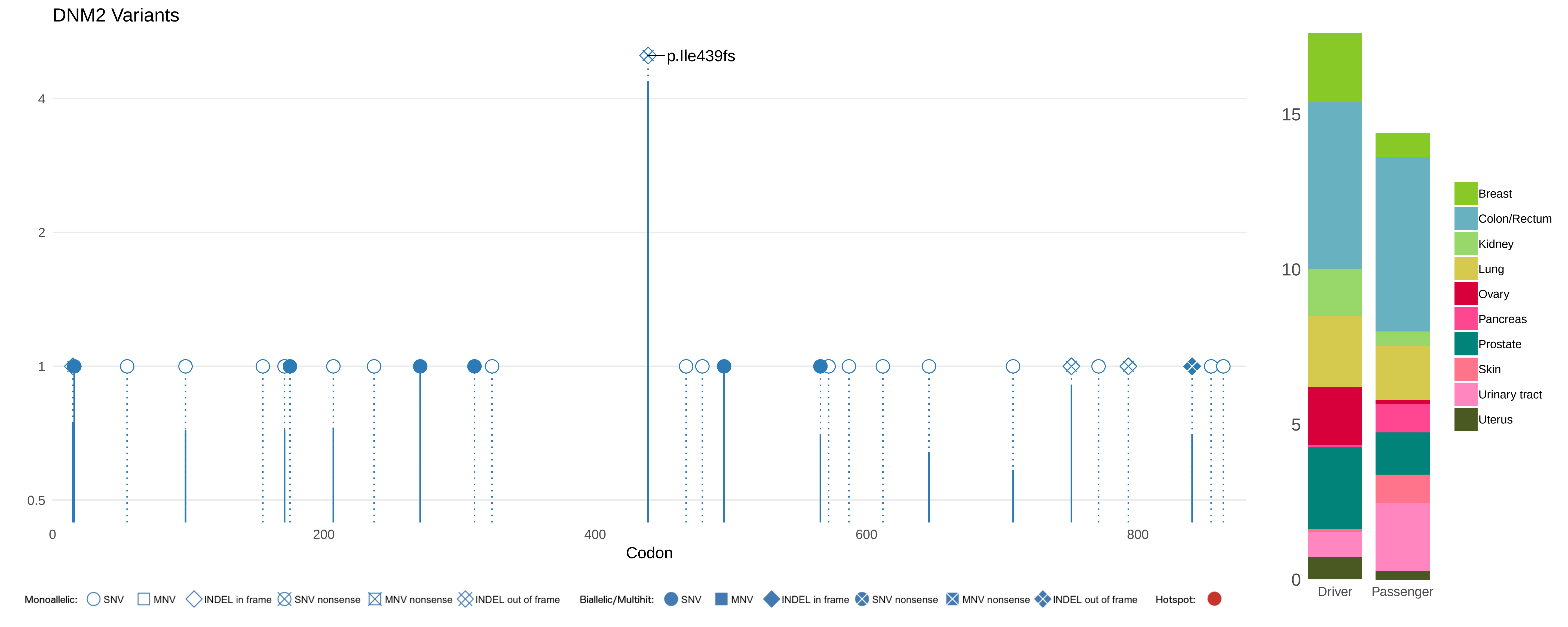
DNM2 Variants
p.Ile439fs
4
2
1
0.5
0
200
400
600
800
Codon
15
10
5
0
Driver
Passenger
Breast
Colon/Rectum
Kidney
Lung
Ovary
Pancreas
Prostate
Skin
Urinary tract
Uterus
Monoallelic: SNV MNV INDEL in frame SNV nonsense MNV nonsense INDEL out of frame
Biallelic/Multihit: SNV MNV INDEL in frame SNV nonsense MNV nonsense INDEL out of frame
Hotspot:

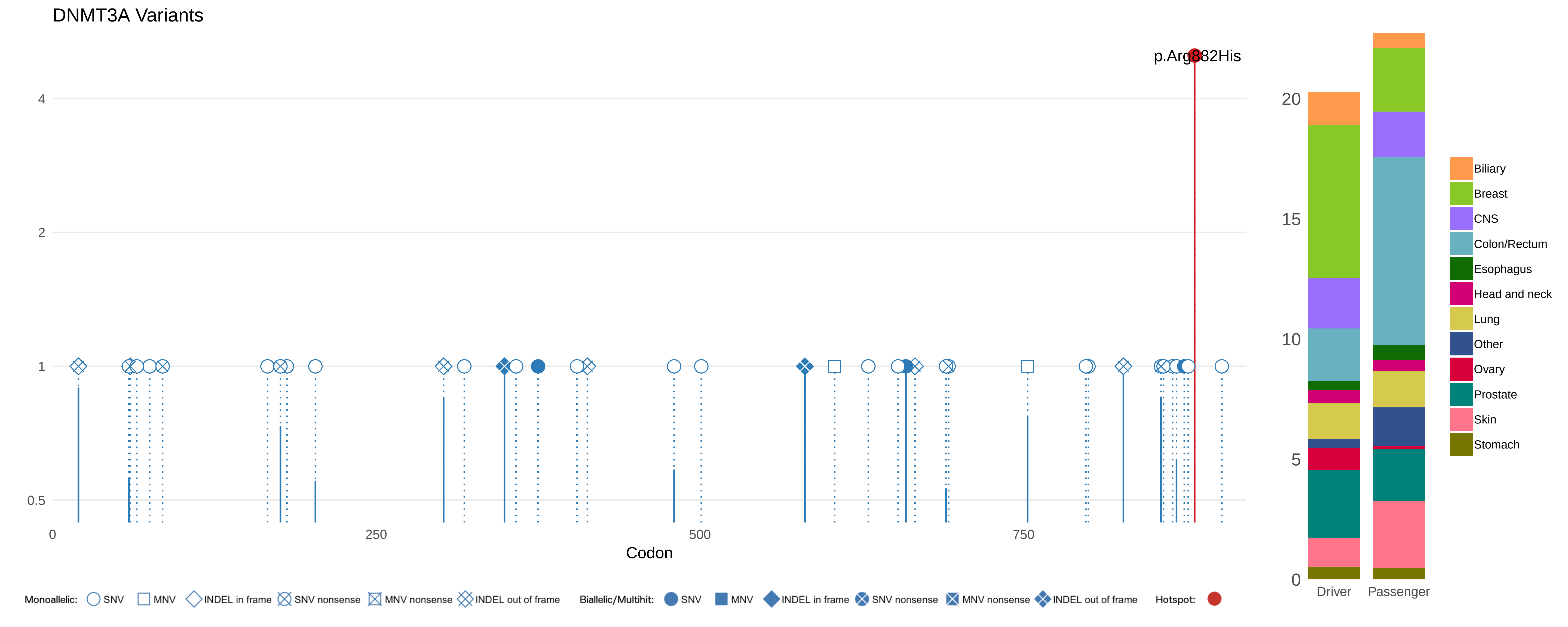
DNMT3A Variants
p.Arg882His
4
2
1
0.5
0
250
500
750
Codon
20
15
10
5
0
Driver
Passenger
Biliary
Breast
CNS
Colon/Rectum
Esophagus
Head and neck
Lung
Other
Ovary
Prostate
Skin
Stomach
Monoallelic: SNV MNV INDEL in frame SNV nonsense MNV nonsense INDEL out of frame
Biallelic/Multihit: SNV MNV INDEL in frame SNV nonsense MNV nonsense INDEL out of frame
Hotspot:

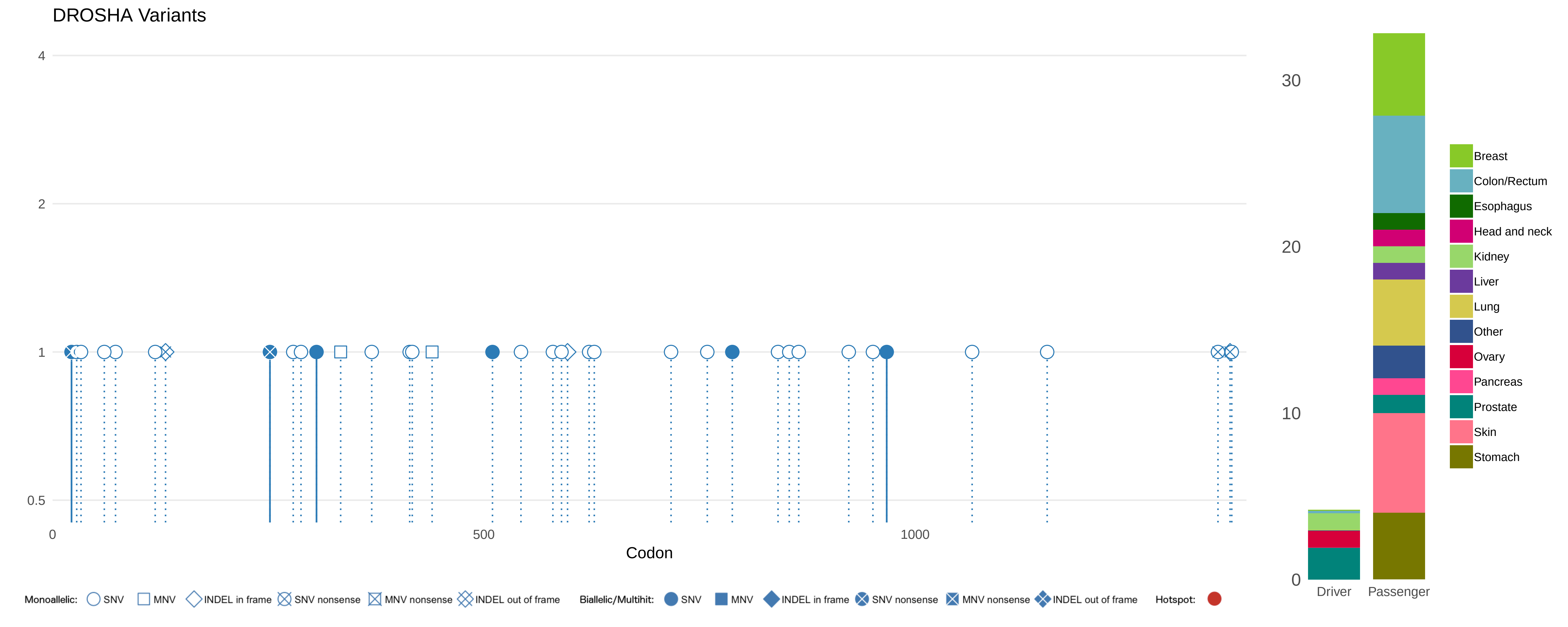
DROSHA Variants
4
2
1
0.5
0
500
1000
Codon
30
20
10
0
Driver
Passenger
Breast
Colon/Rectum
Esophagus
Head and neck
Kidney
Liver
Lung
Other
Ovary
Pancreas
Prostate
Skin
Stomach
Monoallelic: SNV MNV INDEL in frame SNV nonsense MNV nonsense INDEL out of frame
Biallelic/Multihit: SNV MNV INDEL in frame SNV nonsense MNV nonsense INDEL out of frame
Hotspot:

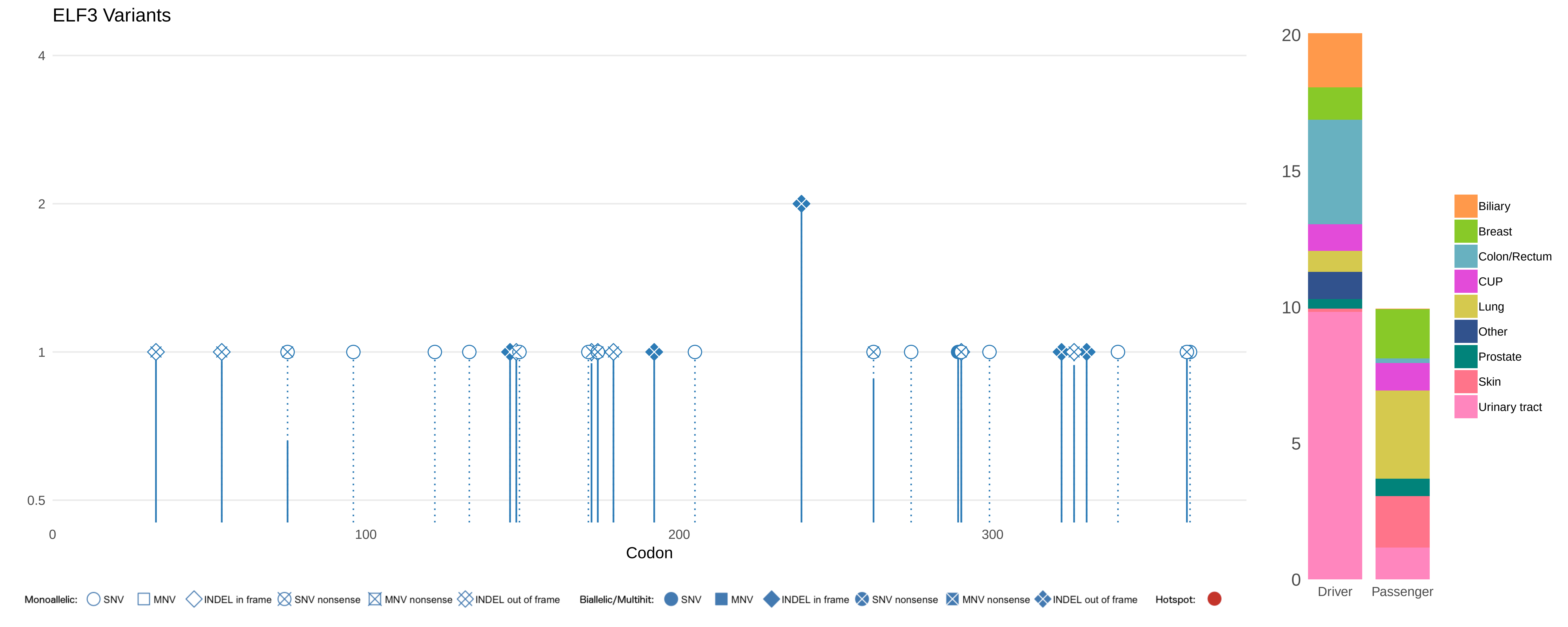

ELF3 Variants
4
2
1
0.5
0
100
200
300
Codon
20
15
10
5
0
Driver
Passenger
Biliary
Breast
Colon/Rectum
CUP
Lung
Other
Prostate
Skin
Urinary tract
Monoallelic: SNV MNV INDEL in frame SNV nonsense MNV nonsense INDEL out of frame
Biallelic/Multihit: SNV MNV INDEL in frame SNV nonsense MNV nonsense INDEL out of frame
Hotspot:

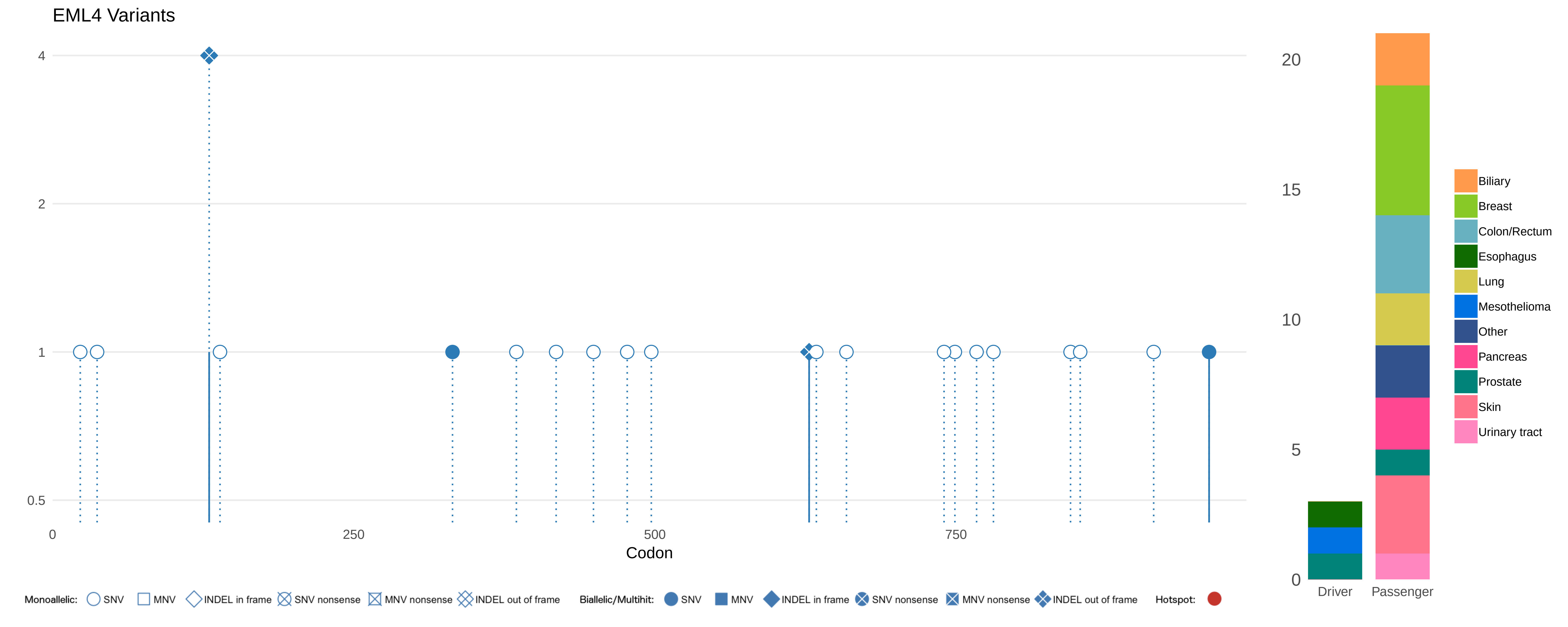

EML4 Variants
4
2
1
0.5
0
250
500
750
Codon
20
15
10
5
0
Driver
Passenger
Biliary
Breast
Colon/Rectum
Esophagus
Lung
Mesothelioma
Other
Pancreas
Prostate
Skin
Urinary tract
Monoallelic: SNV MNV INDEL in frame SNV nonsense MNV nonsense INDEL out of frame
Biallelic/Multihit: SNV MNV INDEL in frame SNV nonsense MNV nonsense INDEL out of frame
Hotspot:

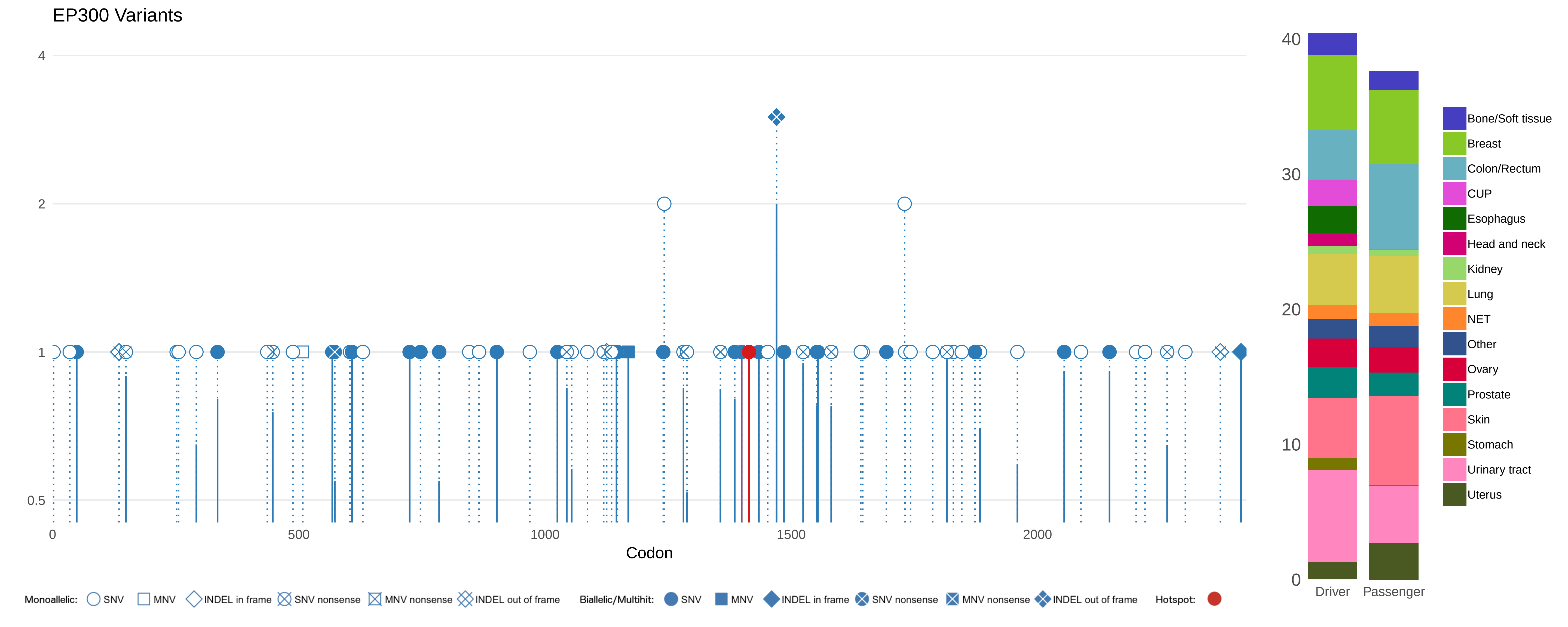

EP300 Variants
4
2
1
0.5
0
500
1000
1500
2000
Codon
Monoallelic: SNV MNV INDEL in frame SNV nonsense MNV nonsense INDEL out of frame
Biallelic/Multihit: SNV MNV INDEL in frame SNV nonsense MNV nonsense INDEL out of frame
Hotspot:
40
30
20
10
0
Driver
Passenger
Bone/Soft tissue
Breast
Colon/Rectum
CUP
Esophagus
Head and neck
Kidney
Lung
NET
Other
Ovary
Prostate
Skin
Stomach
Urinary tract
Uterus

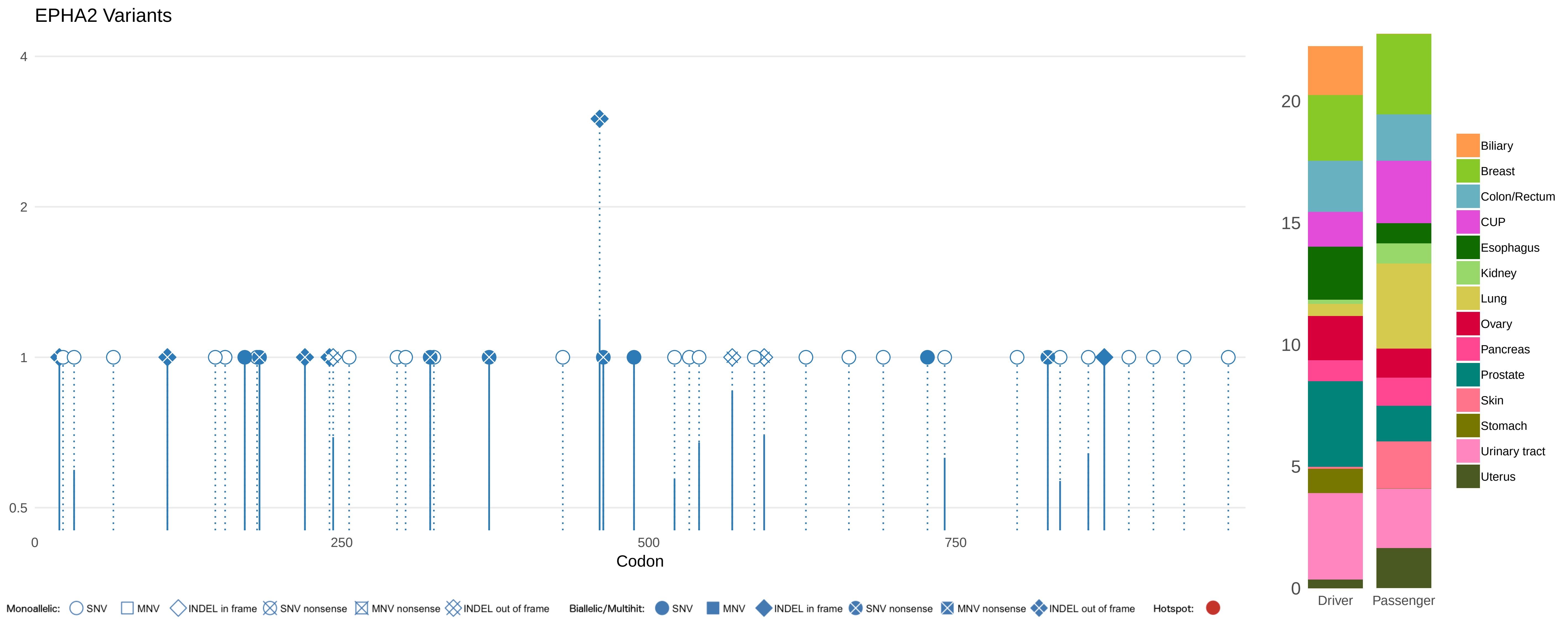

EPHA2 Variants
4
2
1
0.5
0
250
500
750
Codon
20
15
10
5
0
Driver
Passenger
Biliary
Breast
Colon/Rectum
CUP
Esophagus
Kidney
Lung
Ovary
Pancreas
Prostate
Skin
Stomach
Urinary tract
Uterus
Monoallelic: SNV MNV INDEL in frame SNV nonsense MNV nonsense INDEL out of frame
Biallelic/Multihit: SNV MNV INDEL in frame SNV nonsense MNV nonsense INDEL out of frame
Hotspot:

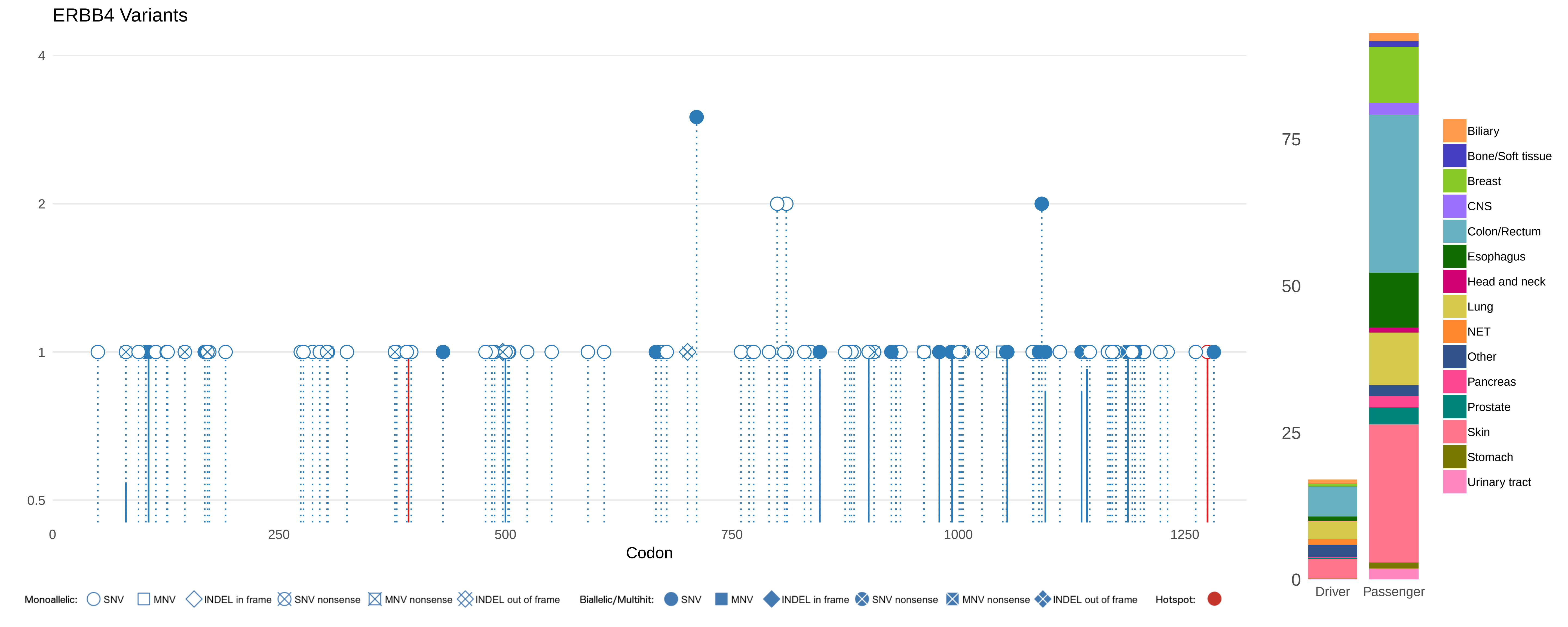
ERBB4 Variants
4
2
1
0.5
0
250
500
750
1000
1250
Codon
Monoallelic: SNV MNV INDEL in frame SNV nonsense MNV nonsense INDEL out of frame
Biallelic/Multihit: SNV MNV INDEL in frame SNV nonsense MNV nonsense INDEL out of frame
Hotspot:
75
50
25
0
Driver
Passenger
Biliary
Bone/Soft tissue
Breast
CNS
Colon/Rectum
Esophagus
Head and neck
Lung
NET
Other
Pancreas
Prostate
Skin
Stomach
Urinary tract

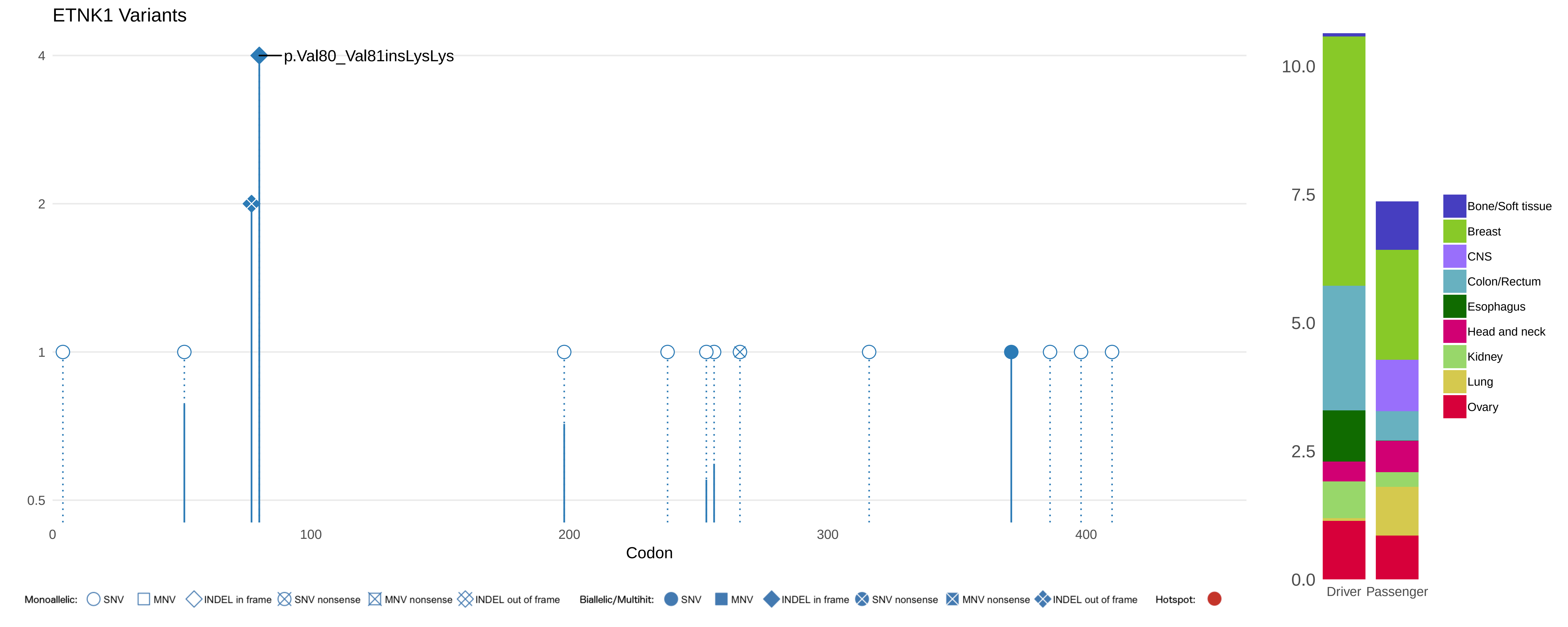

ETNK1 Variants
p.Val80_Val81insLysLys
4
2
1
0.5
0
100
200
300
400
Codon
10.0
7.5
5.0
2.5
0.0
Driver
Passenger
Bone/Soft tissue
Breast
CNS
Colon/Rectum
Esophagus
Head and neck
Kidney
Lung
Ovary
Monoallelic: SNV MNV INDEL in frame SNV nonsense MNV nonsense INDEL out of frame
Biallelic/Multihit: SNV MNV INDEL in frame SNV nonsense MNV nonsense INDEL out of frame
Hotspot:

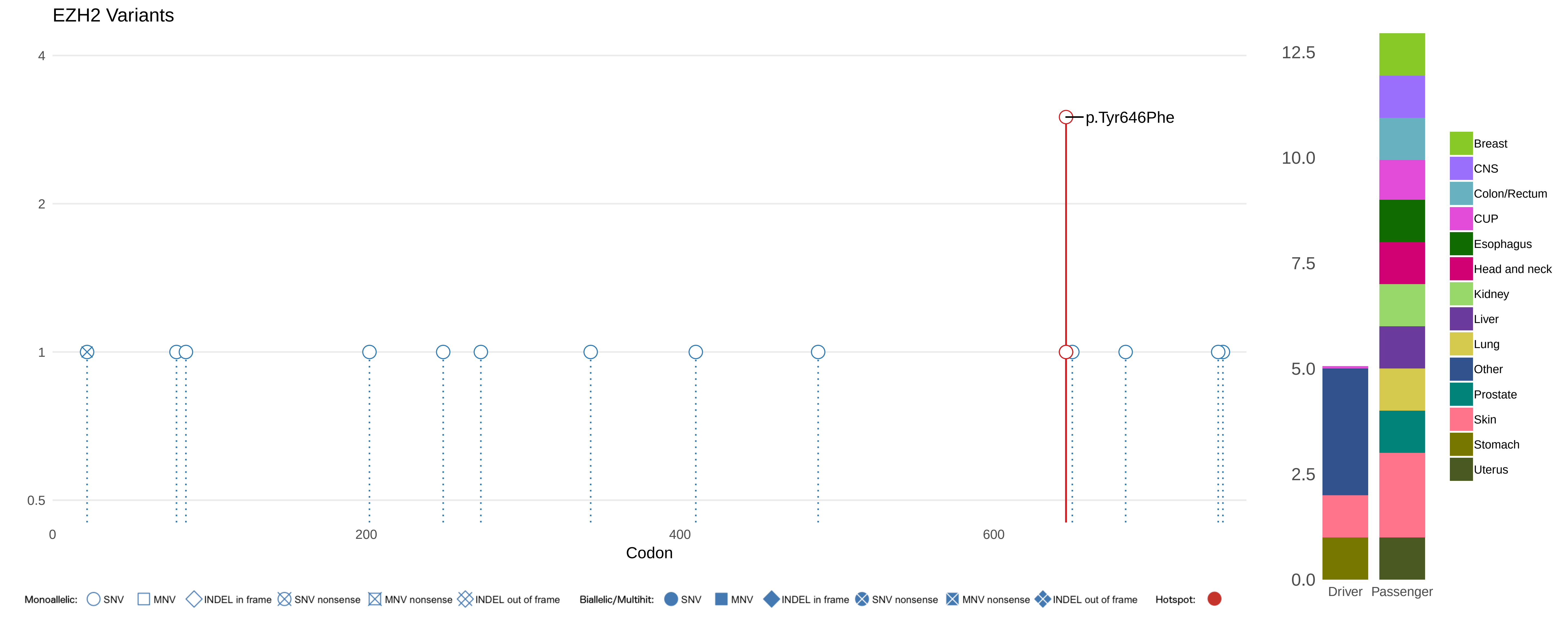

EZH2 Variants
4
2
1
0.5
0
200
400
600
Codon
p.Tyr646Phe
12.5
10.0
7.5
5.0
2.5
0.0
Driver
Passenger
Breast
CNS
Colon/Rectum
CUP
Esophagus
Head and neck
Kidney
Liver
Lung
Other
Prostate
Skin
Stomach
Uterus
Monoallelic: SNV MNV INDEL in frame SNV nonsense MNV nonsense INDEL out of frame
Biallelic/Multihit: SNV MNV INDEL in frame SNV nonsense MNV nonsense INDEL out of frame
Hotspot:

#### FAT1 Variants

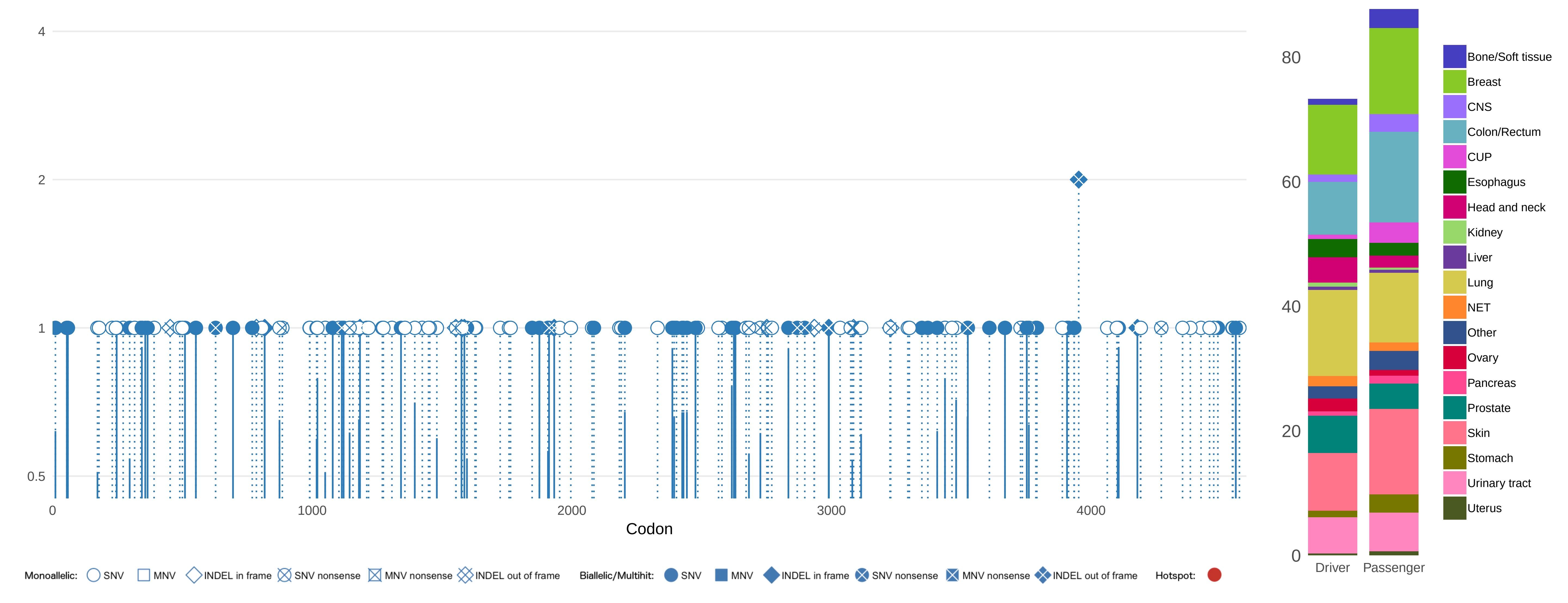

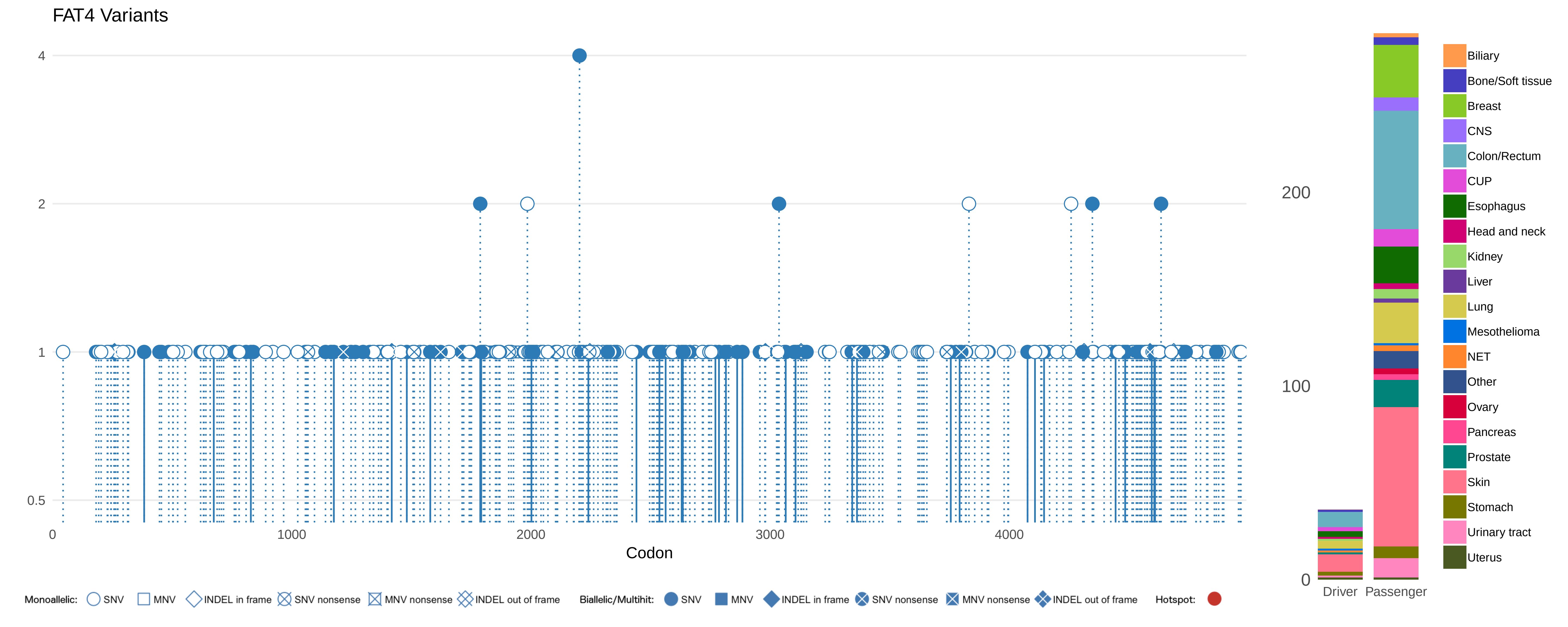
FAT4 Variants
4
2
1
0.5
0
1000
2000
3000
4000
Codon
200
100
0
Driver
Passenger
Biliary
Bone/Soft tissue
Breast
CNS
Colon/Rectum
CUP
Esophagus
Head and neck
Kidney
Liver
Lung
Mesothelioma
NET
Other
Ovary
Pancreas
Prostate
Skin
Stomach
Urinary tract
Uterus
Monoallelic: SNV MNV INDEL in frame SNV nonsense MNV nonsense INDEL out of frame
Biallelic/Multihit: SNV MNV INDEL in frame SNV nonsense MNV nonsense INDEL out of frame
Hotspot:

### FBXO11 Variants

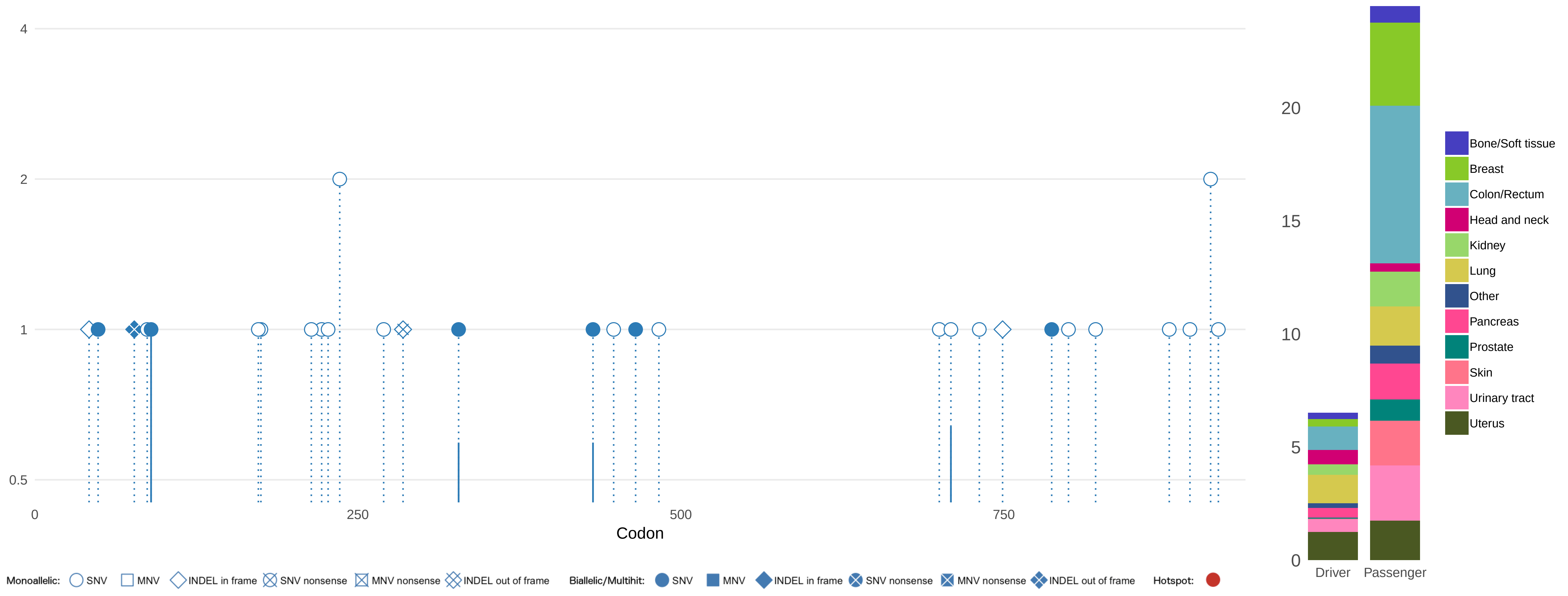

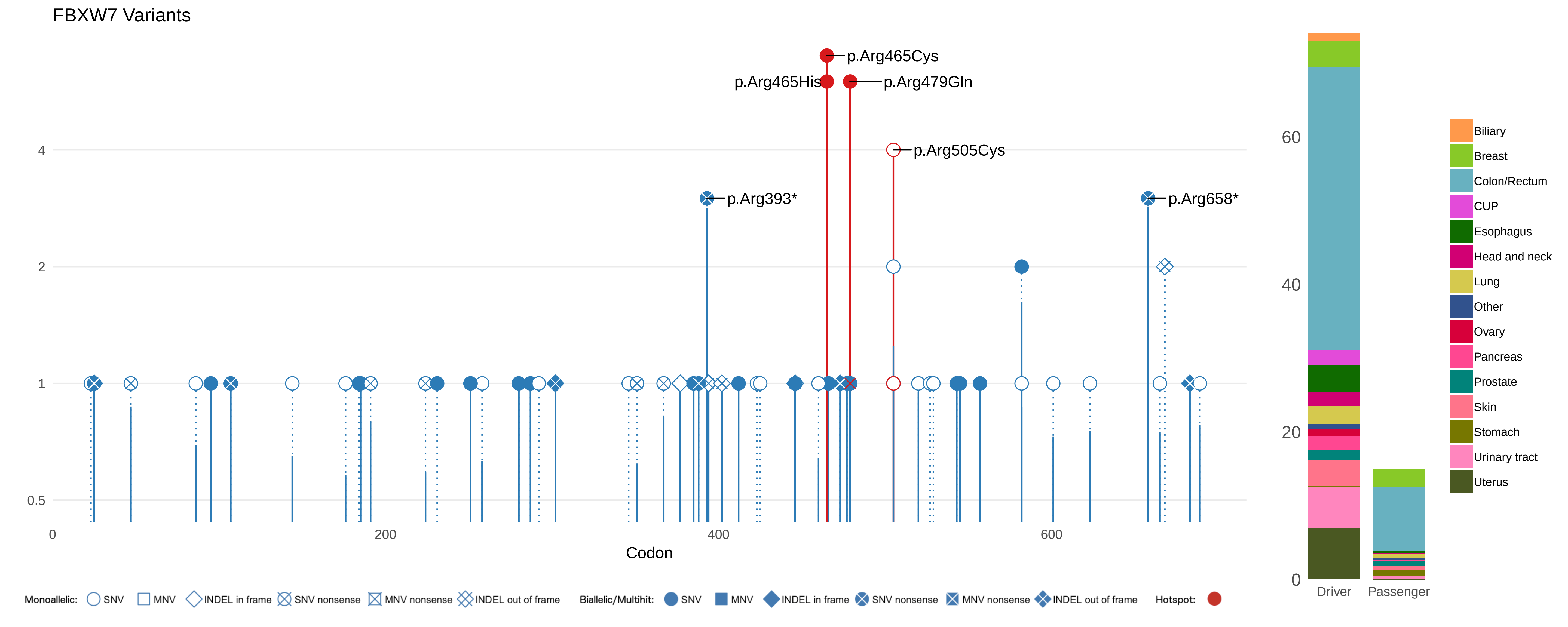
FBXW7 Variants
p.Arg465Cys
p.Arg465His
p.Arg479Gln
p.Arg505Cys
p.Arg393*
p.Arg658*
4
2
1
0.5
0
200
400
600
Codon
Monoallelic: SNV MNV INDEL in frame SNV nonsense MNV nonsense INDEL out of frame
Biallelic/Multihit: SNV MNV INDEL in frame SNV nonsense MNV nonsense INDEL out of frame
Hotspot:
60
40
20
0
Driver
Passenger
Biliary
Breast
Colon/Rectum
CUP
Esophagus
Head and neck
Lung
Other
Ovary
Pancreas
Prostate
Skin
Stomach
Urinary tract
Uterus

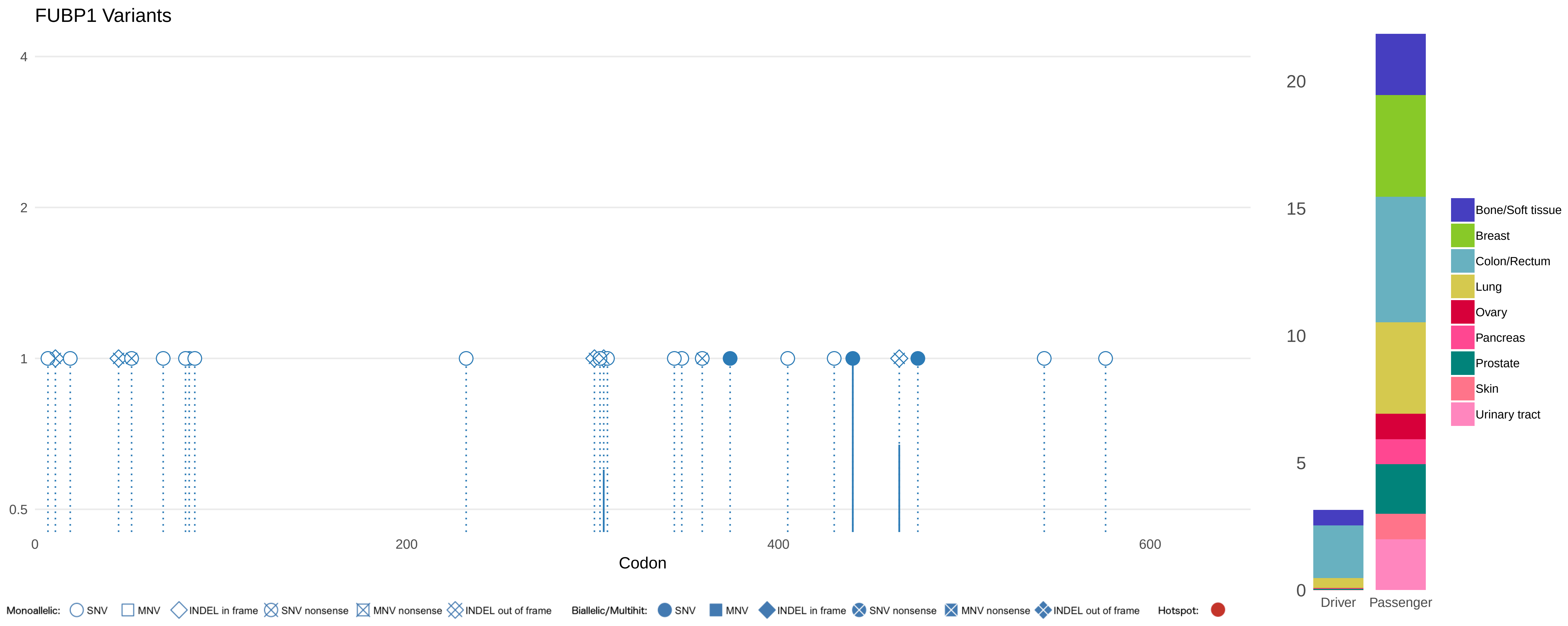
FUBP1 Variants
4
2
1
0.5
0
200
400
600
Codon
Monoallelic: SNV MNV INDEL in frame SNV nonsense MNV nonsense INDEL out of frame
Biallelic/Multihit: SNV MNV INDEL in frame SNV nonsense MNV nonsense INDEL out of frame
Hotspot:
20
15
10
5
0
Driver
Passenger
Bone/Soft tissue
Breast
Colon/Rectum
Lung
Ovary
Pancreas
Prostate
Skin
Urinary tract

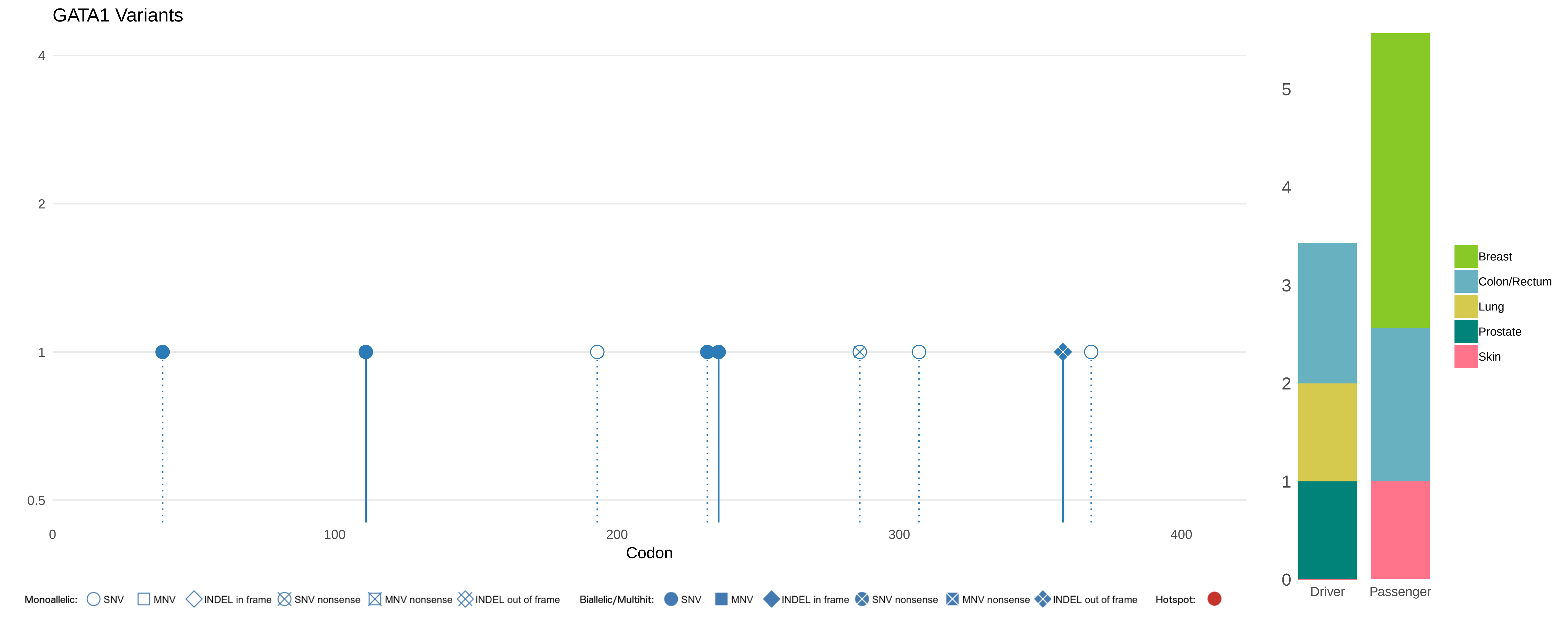

GATA1 Variants
4
2
1
0.5
0
100
200
300
400
Codon
5
4
3
2
1
0
Driver
Passenger
Breast
Colon/Rectum
Lung
Prostate
Skin
Monoallelic: SNV MNV INDEL in frame SNV nonsense MNV nonsense INDEL out of frame
Biallelic/Multihit: SNV MNV INDEL in frame SNV nonsense MNV nonsense INDEL out of frame
Hotspot:

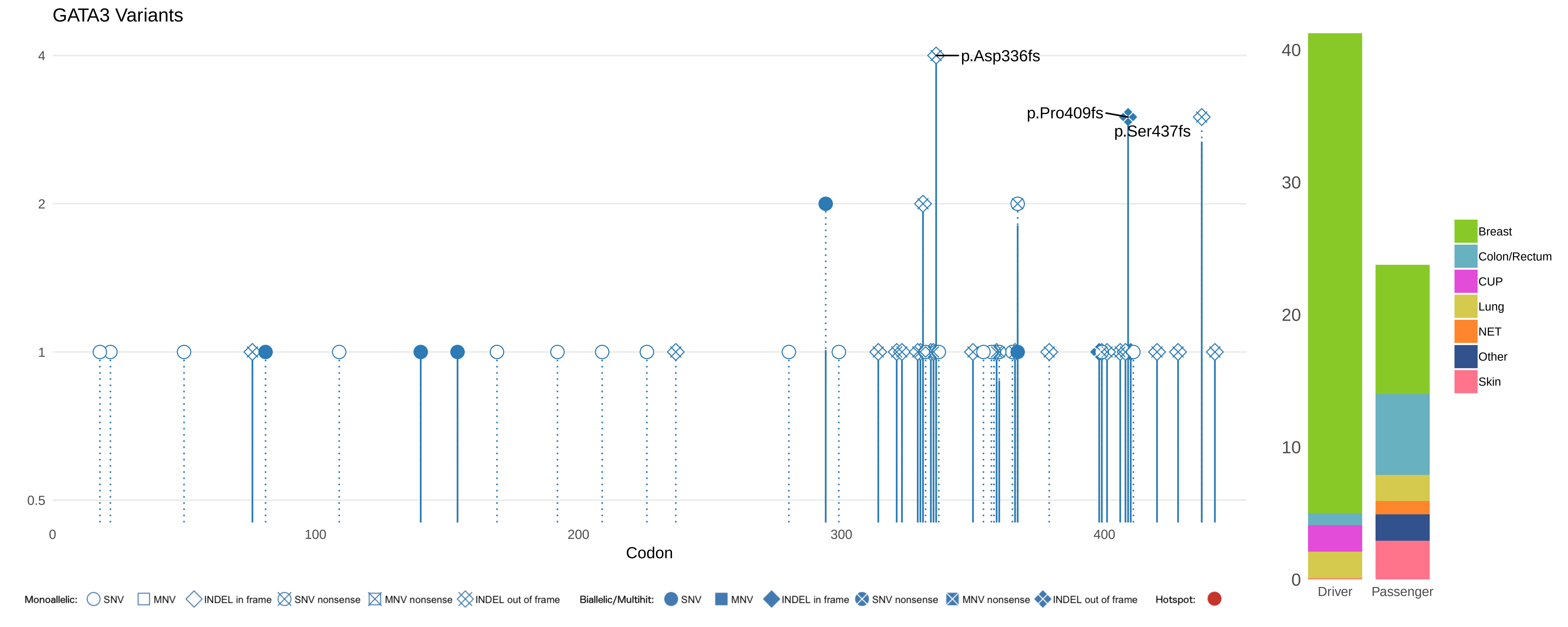

GATA3 Variants
p.Asp336fs
p.Pro409fs
p.Ser437fs
4
2
1
0.5
0
100
200
300
400
Codon
40
30
20
10
0
Driver
Passenger
Breast
Colon/Rectum
CUP
Lung
NET
Other
Skin
Monoallelic: SNV MNV INDEL in frame SNV nonsense MNV nonsense INDEL out of frame
Biallelic/Multihit: SNV MNV INDEL in frame SNV nonsense MNV nonsense INDEL out of frame
Hotspot:

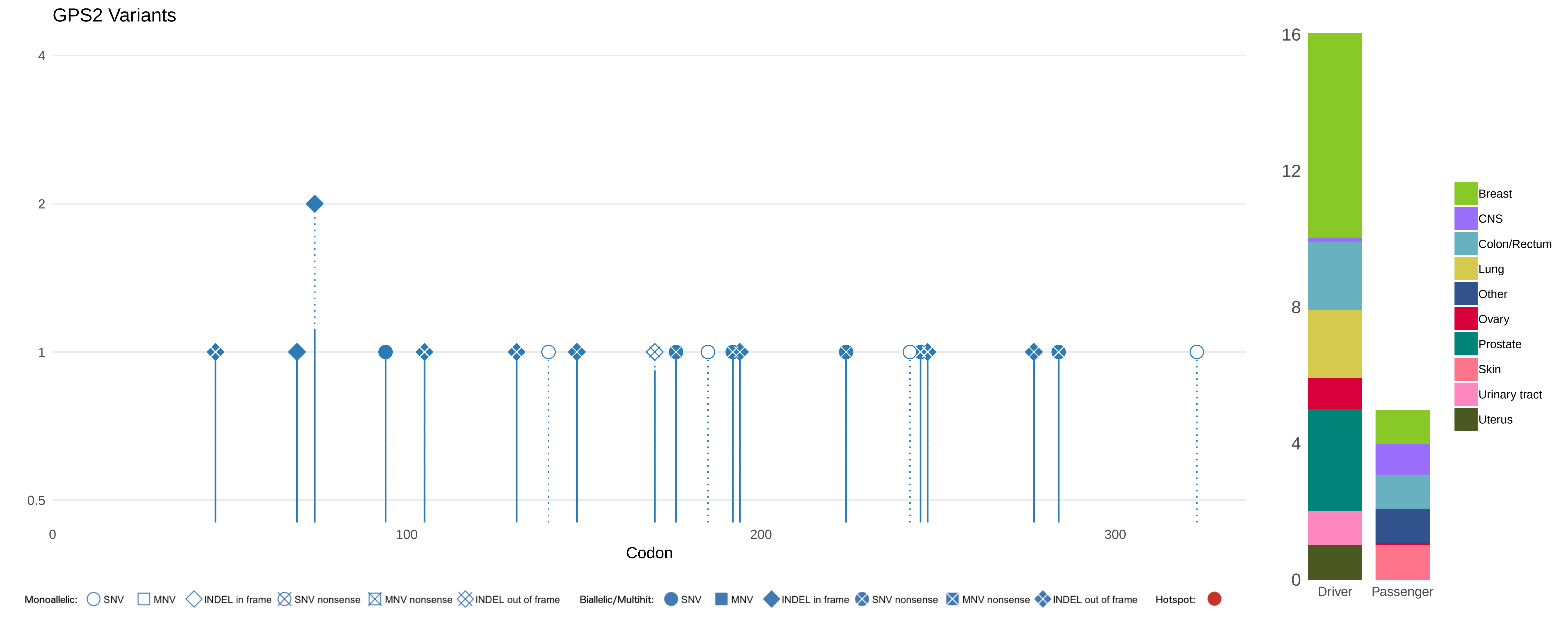
GPS2 Variants
4
2
1
0.5
0
100
200
300
Codon
16
12
8
4
0
Driver
Passenger
Breast
CNS
Colon/Rectum
Lung
Other
Ovary
Prostate
Skin
Urinary tract
Uterus
Monoallelic: SNV MNV INDEL in frame SNV nonsense MNV nonsense INDEL out of frame
Biallelic/Multihit: SNV MNV INDEL in frame SNV nonsense MNV nonsense INDEL out of frame
Hotspot:

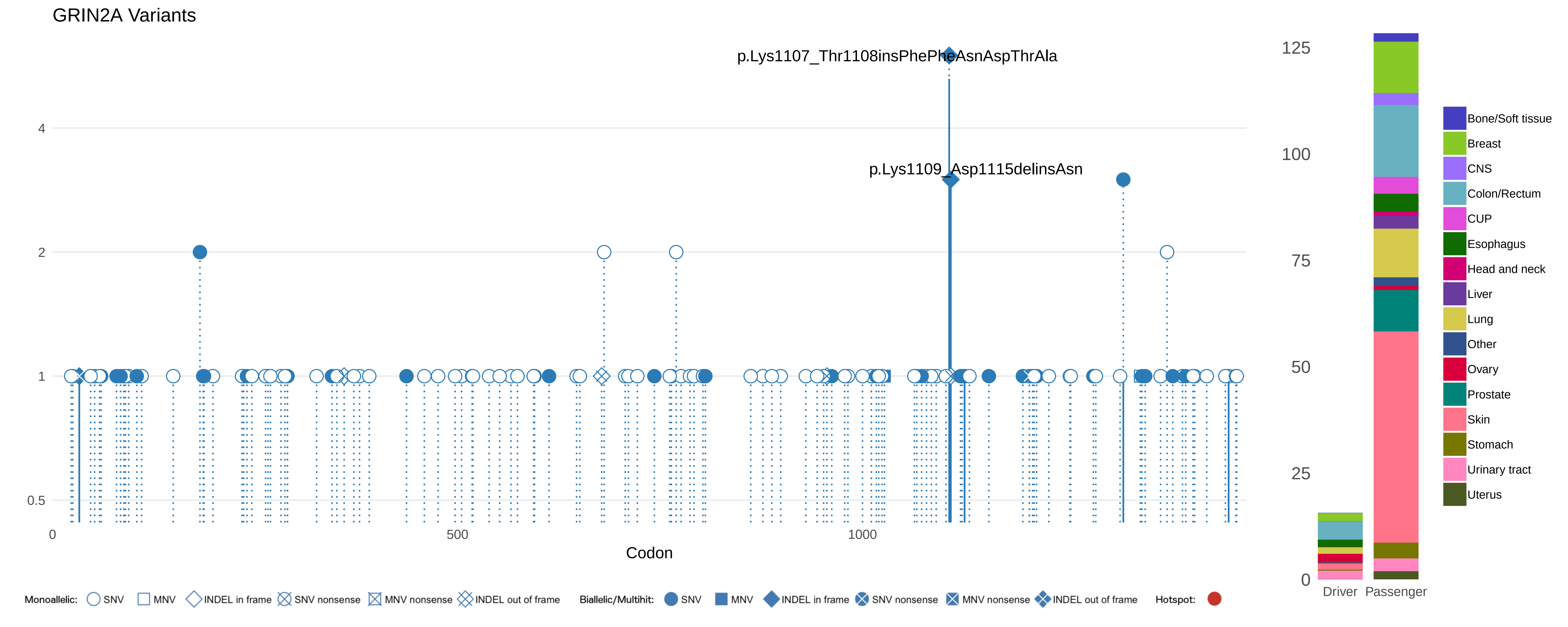
GRIN2A Variants
p.Lys1107_Thr1108insPhePheAsnAspThrAla
p.Lys1109_Asp1115delinsAsn
4
2
1
0.5
0
500
1000
Codon
125
100
75
50
25
0
Driver
Passenger
Bone/Soft tissue
Breast
CNS
Colon/Rectum
CUP
Esophagus
Head and neck
Liver
Lung
Other
Ovary
Prostate
Skin
Stomach
Urinary tract
Uterus
Monoallelic: SNV MNV INDEL in frame SNV nonsense MNV nonsense INDEL out of frame
Biallelic/Multihit: SNV MNV INDEL in frame SNV nonsense MNV nonsense INDEL out of frame
Hotspot:

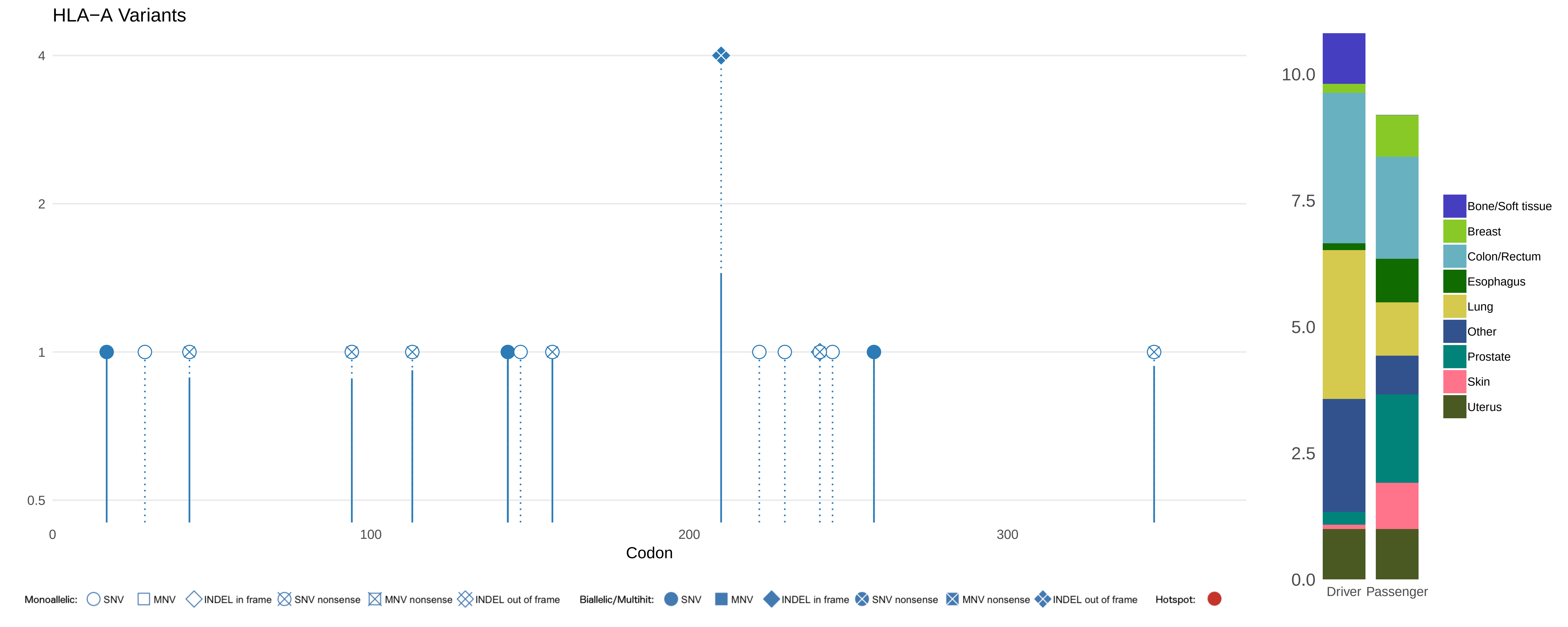

HLA−A Variants
4
2
1
0.5
0
100
200
300
Codon
10.0
7.5
5.0
2.5
0.0
Driver
Passenger
Bone/Soft tissue
Breast
Colon/Rectum
Esophagus
Lung
Other
Prostate
Skin
Uterus
Monoallelic: SNV MNV INDEL in frame SNV nonsense MNV nonsense INDEL out of frame
Biallelic/Multihit: SNV MNV INDEL in frame SNV nonsense MNV nonsense INDEL out of frame
Hotspot:

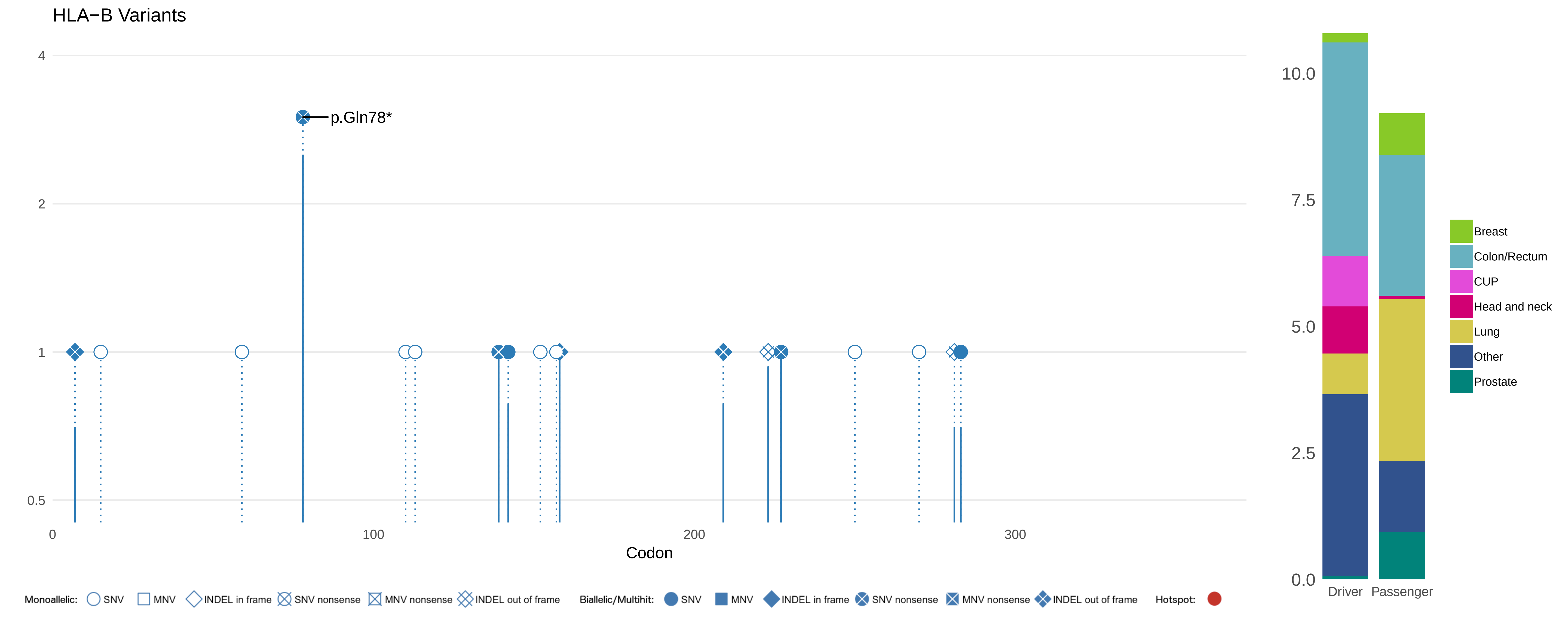

HLA−B Variants
p.Gln78*
4
2
1
0.5
0
100
200
300
Codon
10.0
7.5
5.0
2.5
0.0
Driver
Passenger
Breast
Colon/Rectum
CUP
Head and neck
Lung
Other
Prostate
Monoallelic: SNV MNV INDEL in frame SNV nonsense MNV nonsense INDEL out of frame
Biallelic/Multihit: SNV MNV INDEL in frame SNV nonsense MNV nonsense INDEL out of frame
Hotspot:

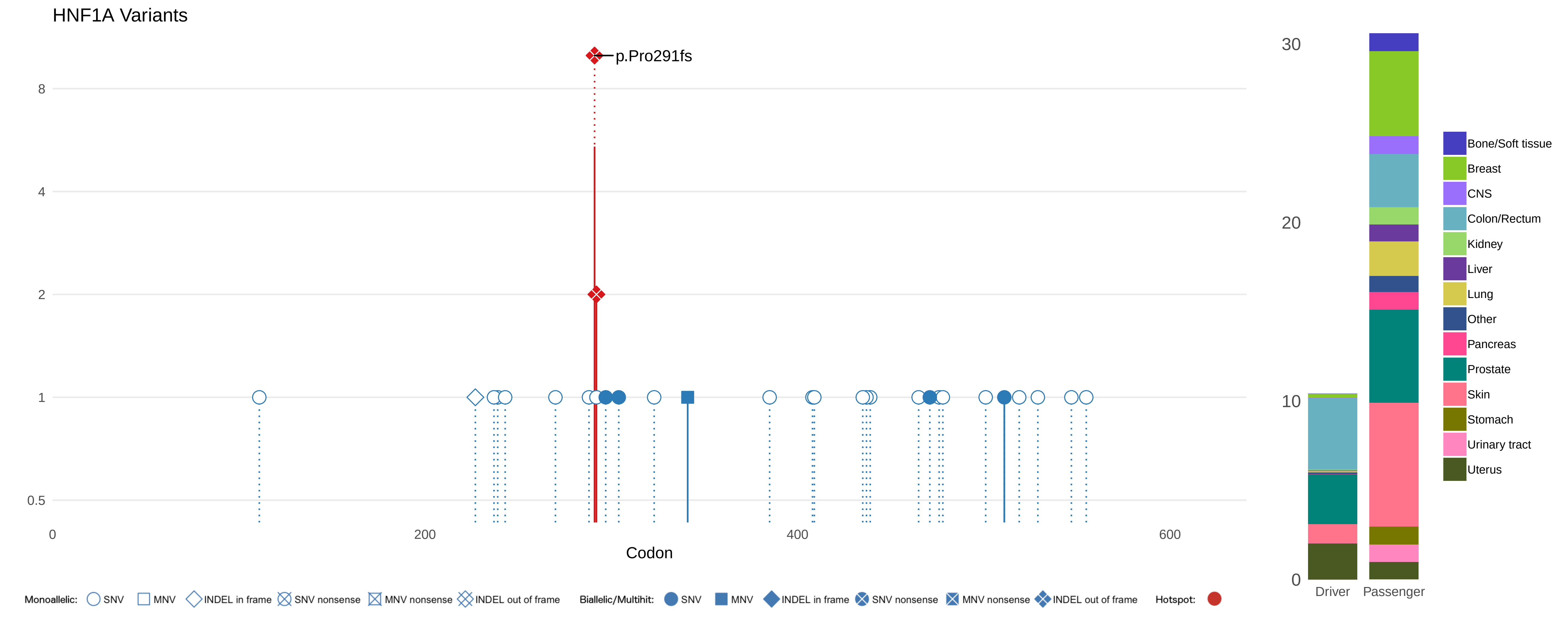
HNF1A Variants
p.Pro291fs
8
4
2
1
0.5
0
200
400
600
Codon
30
20
10
0
Driver
Passenger
Bone/Soft tissue
Breast
CNS
Colon/Rectum
Kidney
Liver
Lung
Other
Pancreas
Prostate
Skin
Stomach
Urinary tract
Uterus
Monoallelic: SNV MNV INDEL in frame SNV nonsense MNV nonsense INDEL out of frame
Biallelic/Multihit: SNV MNV INDEL in frame SNV nonsense MNV nonsense INDEL out of frame
Hotspot:

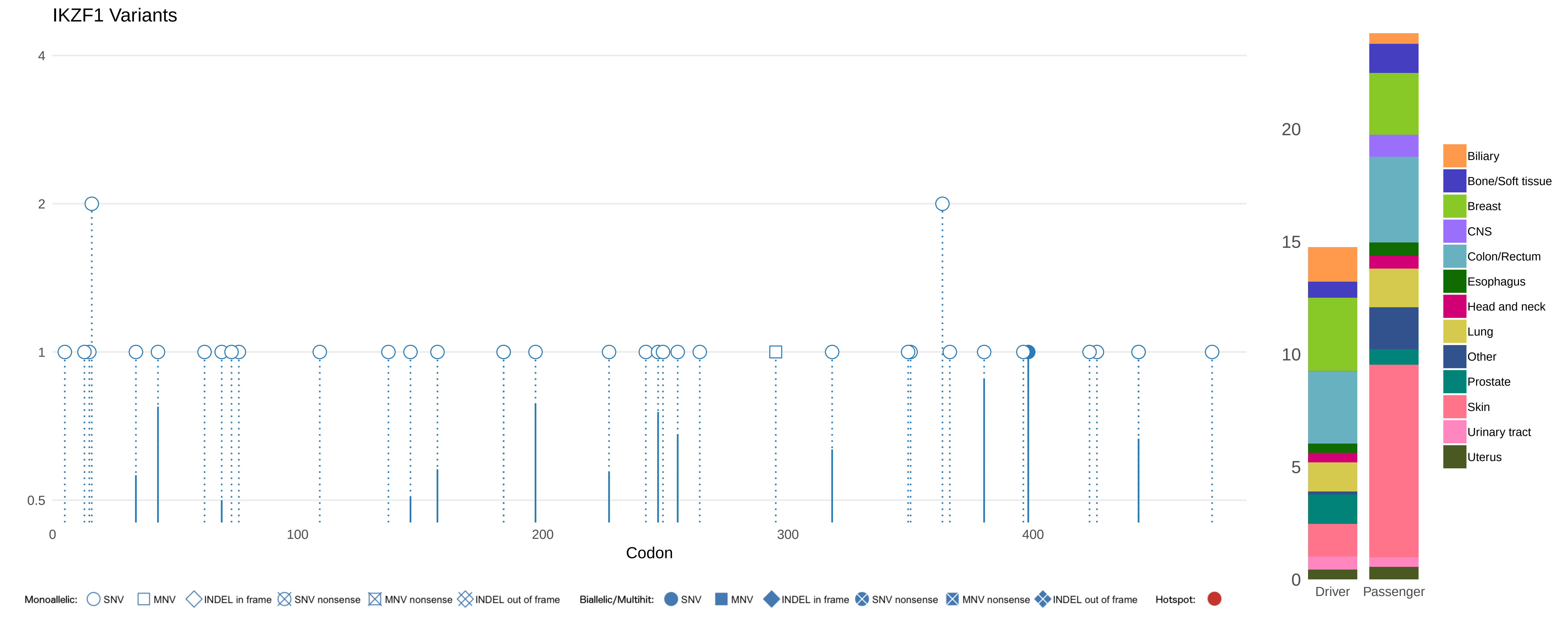

IKZF1 Variants
4
2
1
0.5
0
100
200
300
400
Codon
Monoallelic: SNV MNV INDEL in frame SNV nonsense MNV nonsense INDEL out of frame
Biallelic/Multihit: SNV MNV INDEL in frame SNV nonsense MNV nonsense INDEL out of frame
Hotspot:
20
15
10
5
0
Driver
Passenger
Biliary
Bone/Soft tissue
Breast
CNS
Colon/Rectum
Esophagus
Head and neck
Lung
Other
Prostate
Skin
Urinary tract
Uterus

### JAK1 Variants

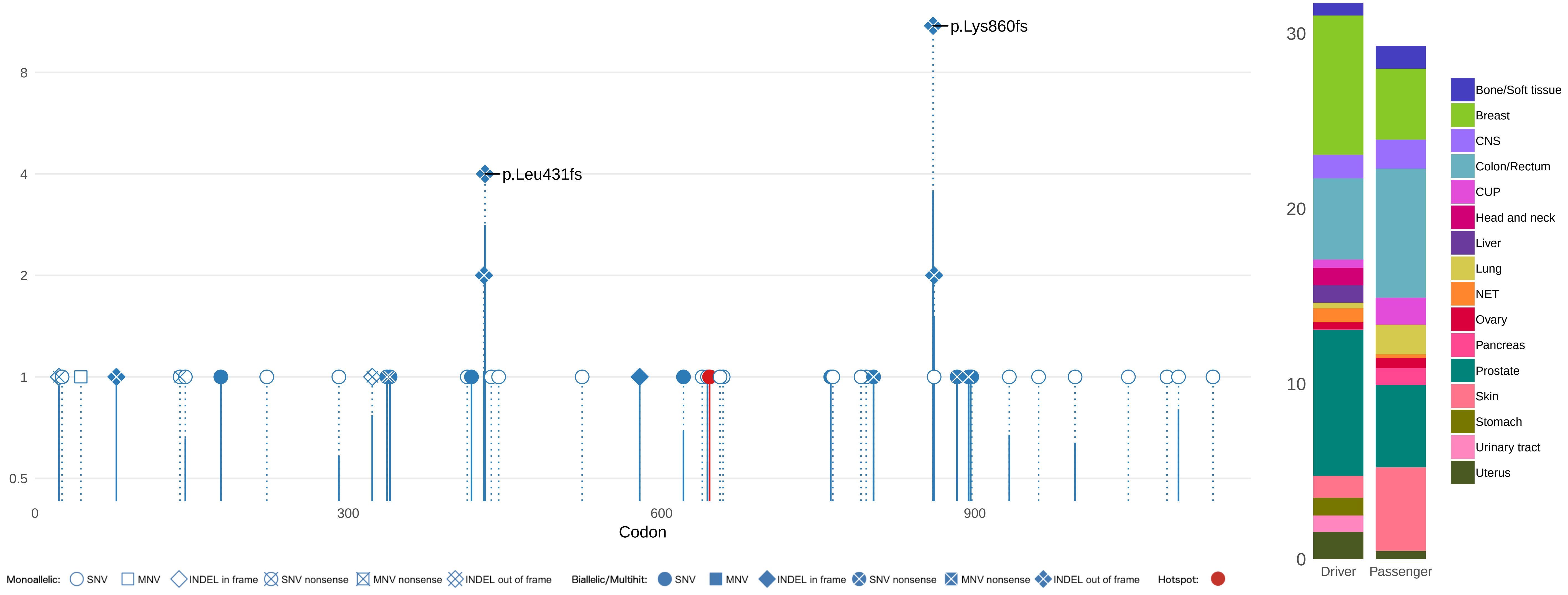

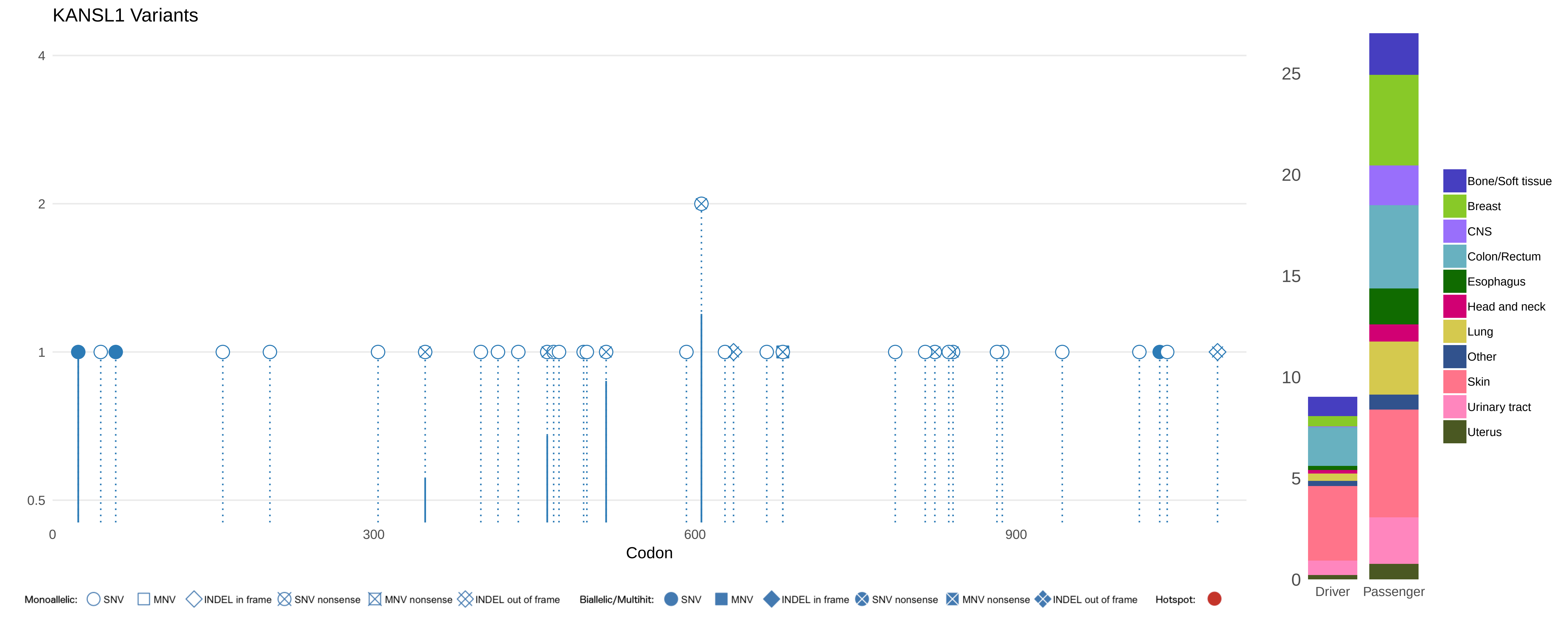
KANSL1 Variants
4
2
1
0.5
0
300
600
900
Codon
25
20
15
10
5
0
Driver
Passenger
Bone/Soft tissue
Breast
CNS
Colon/Rectum
Esophagus
Head and neck
Lung
Other
Skin
Urinary tract
Uterus
Monoallelic: SNV MNV INDEL in frame SNV nonsense MNV nonsense INDEL out of frame
Biallelic/Multihit: SNV MNV INDEL in frame SNV nonsense MNV nonsense INDEL out of frame
Hotspot:

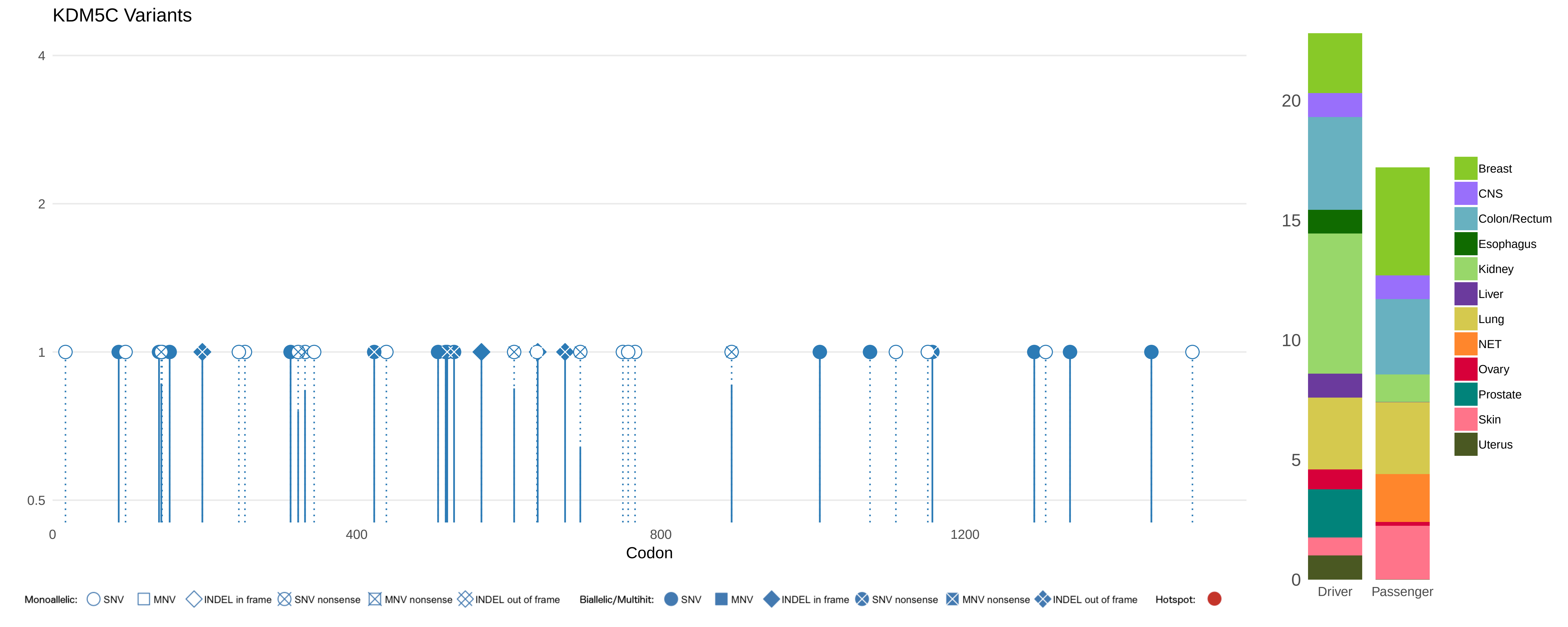
KDM5C Variants
4
2
1
0.5
0
400
800
1200
Codon
20
15
10
5
0
Driver
Passenger
Breast
CNS
Colon/Rectum
Esophagus
Kidney
Liver
Lung
NET
Ovary
Prostate
Skin
Uterus
Monoallelic:
SNV
MNV
INDEL in frame
SNV nonsense
MNV nonsense
INDEL out of frame
Biallelic/Multihit:
SNV
MNV
INDEL in frame
SNV nonsense
MNV nonsense
INDEL out of frame
Hotspot:

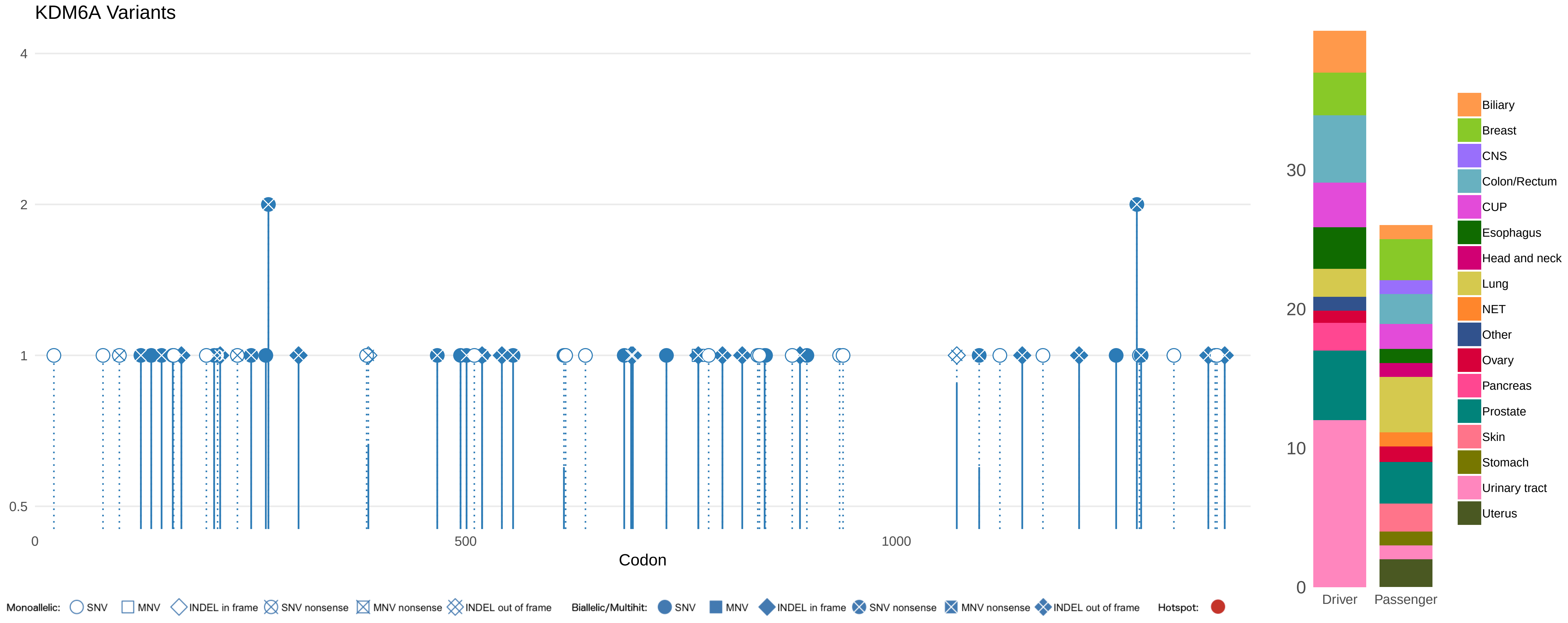

KDM6A Variants
4
2
1
0.5
0
500
1000
Codon
30
20
10
0
Driver
Passenger
Biliary
Breast
CNS
Colon/Rectum
CUP
Esophagus
Head and neck
Lung
NET
Other
Ovary
Pancreas
Prostate
Skin
Stomach
Urinary tract
Uterus
Monoallelic: SNV MNV INDEL in frame SNV nonsense MNV nonsense INDEL out of frame
Biallelic/Multihit: SNV MNV INDEL in frame SNV nonsense MNV nonsense INDEL out of frame
Hotspot:

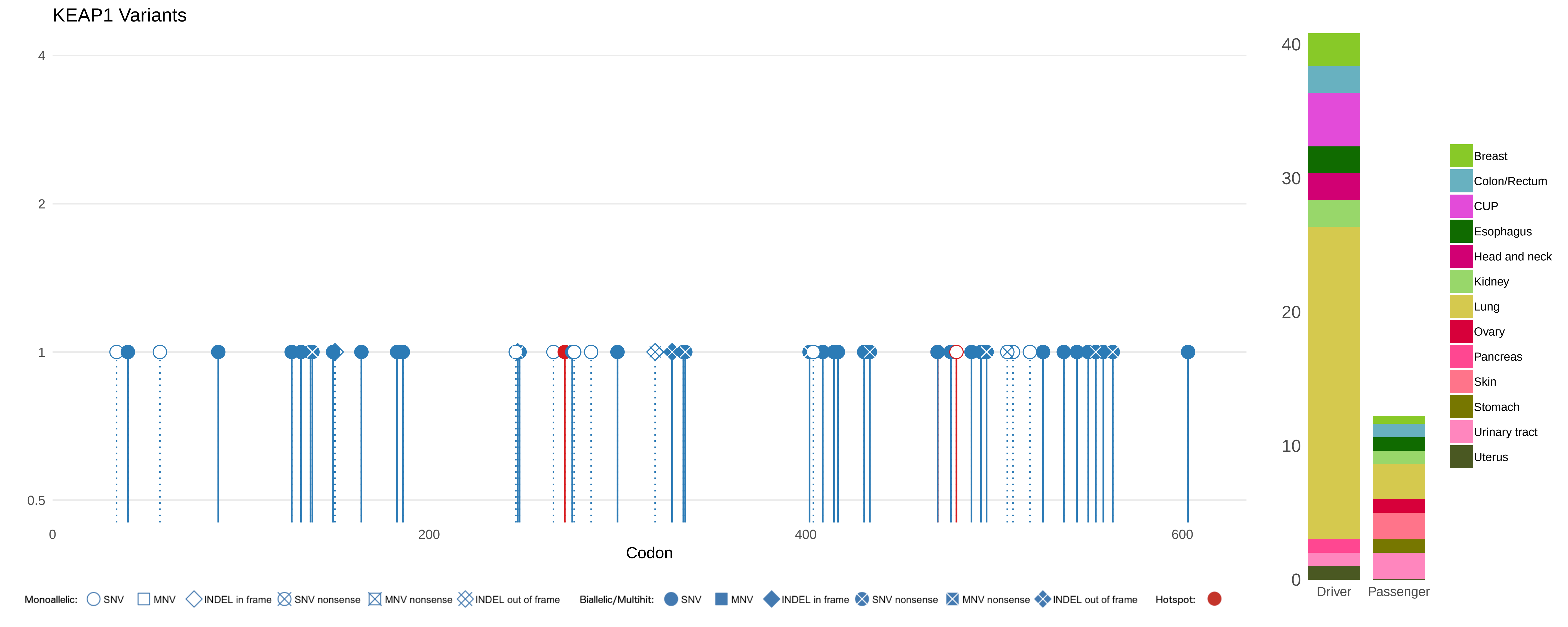

KEAP1 Variants
4
2
1
0.5
0
200
400
600
Codon
40
30
20
10
0
Driver
Passenger
Breast
Colon/Rectum
CUP
Esophagus
Head and neck
Kidney
Lung
Ovary
Pancreas
Skin
Stomach
Urinary tract
Uterus
Monoallelic: SNV MNV INDEL in frame SNV nonsense MNV nonsense INDEL out of frame
Biallelic/Multihit: SNV MNV INDEL in frame SNV nonsense MNV nonsense INDEL out of frame
Hotspot:

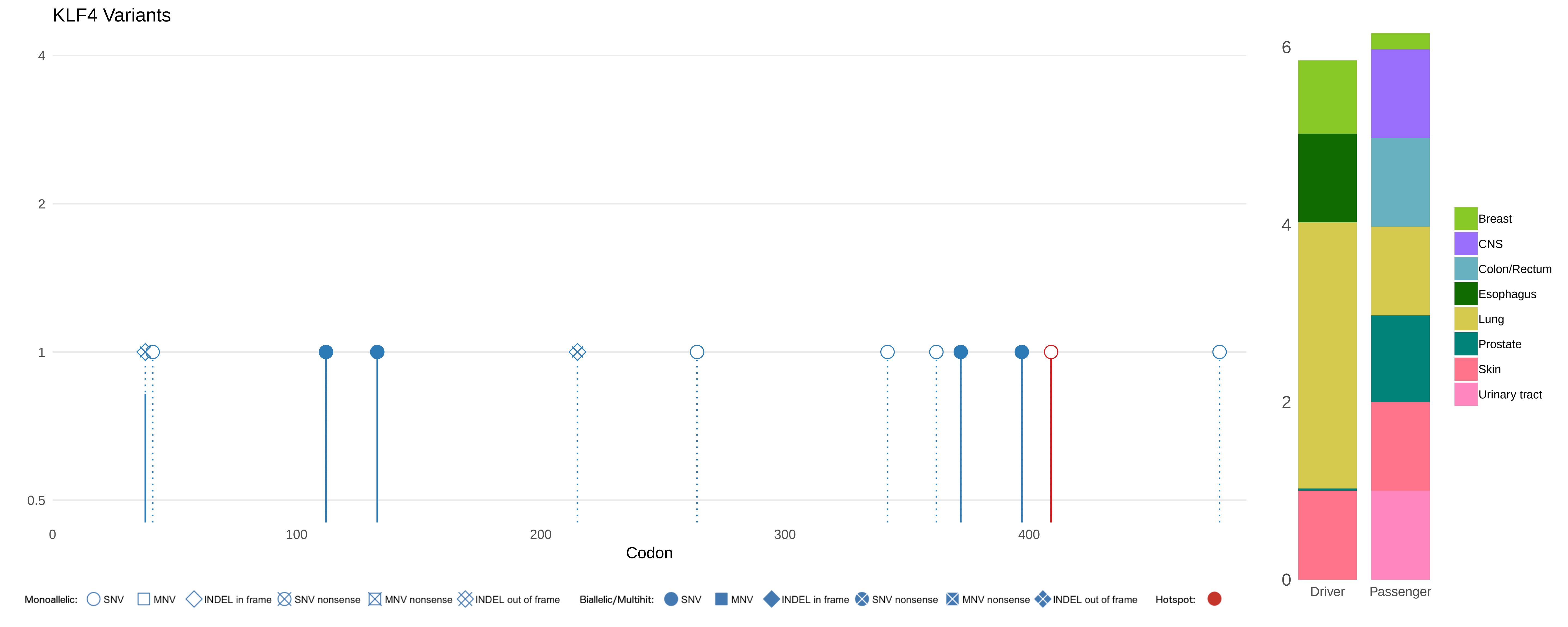
KLF4 Variants
4
2
1
0.5
0
100
200
300
400
Codon
6
4
2
0
Driver
Passenger
Breast
CNS
Colon/Rectum
Esophagus
Lung
Prostate
Skin
Urinary tract
Monoallelic: SNV MNV INDEL in frame SNV nonsense MNV nonsense INDEL out of frame
Biallelic/Multihit: SNV MNV INDEL in frame SNV nonsense MNV nonsense INDEL out of frame
Hotspot:

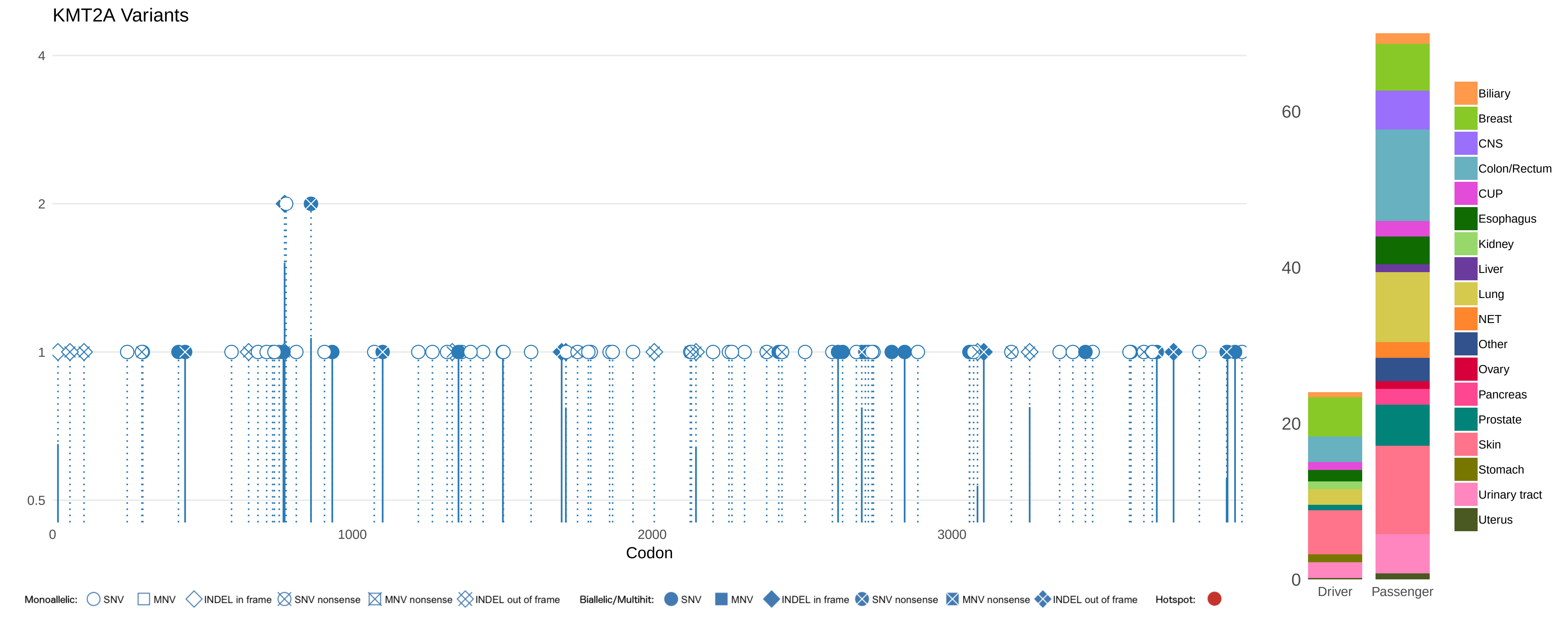
KMT2A Variants
4
2
1
0.5
0
1000
2000
3000
Codon
60
40
20
0
Driver
Passenger
Biliary
Breast
CNS
Colon/Rectum
CUP
Esophagus
Kidney
Liver
Lung
NET
Other
Ovary
Pancreas
Prostate
Skin
Stomach
Urinary tract
Uterus
Monoallelic: SNV MNV INDEL in frame SNV nonsense MNV nonsense INDEL out of frame
Biallelic/Multihit: SNV MNV INDEL in frame SNV nonsense MNV nonsense INDEL out of frame
Hotspot:

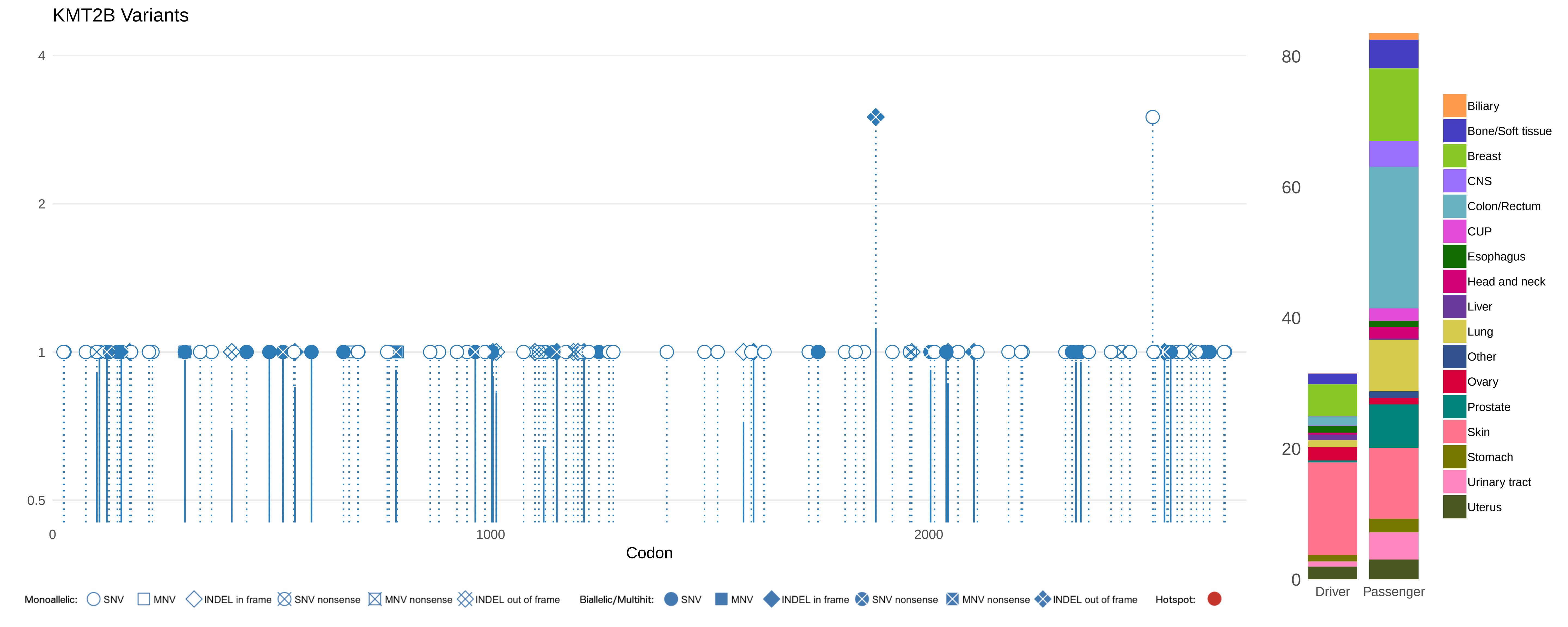
KMT2B Variants
4
2
1
0.5
0
1000
2000
Codon
80
60
40
20
0
Driver
Passenger
Biliary
Bone/Soft tissue
Breast
CNS
Colon/Rectum
CUP
Esophagus
Head and neck
Liver
Lung
Other
Ovary
Prostate
Skin
Stomach
Urinary tract
Uterus
Monoallelic: SNV MNV INDEL in frame SNV nonsense MNV nonsense INDEL out of frame
Biallelic/Multihit: SNV MNV INDEL in frame SNV nonsense MNV nonsense INDEL out of frame
Hotspot:

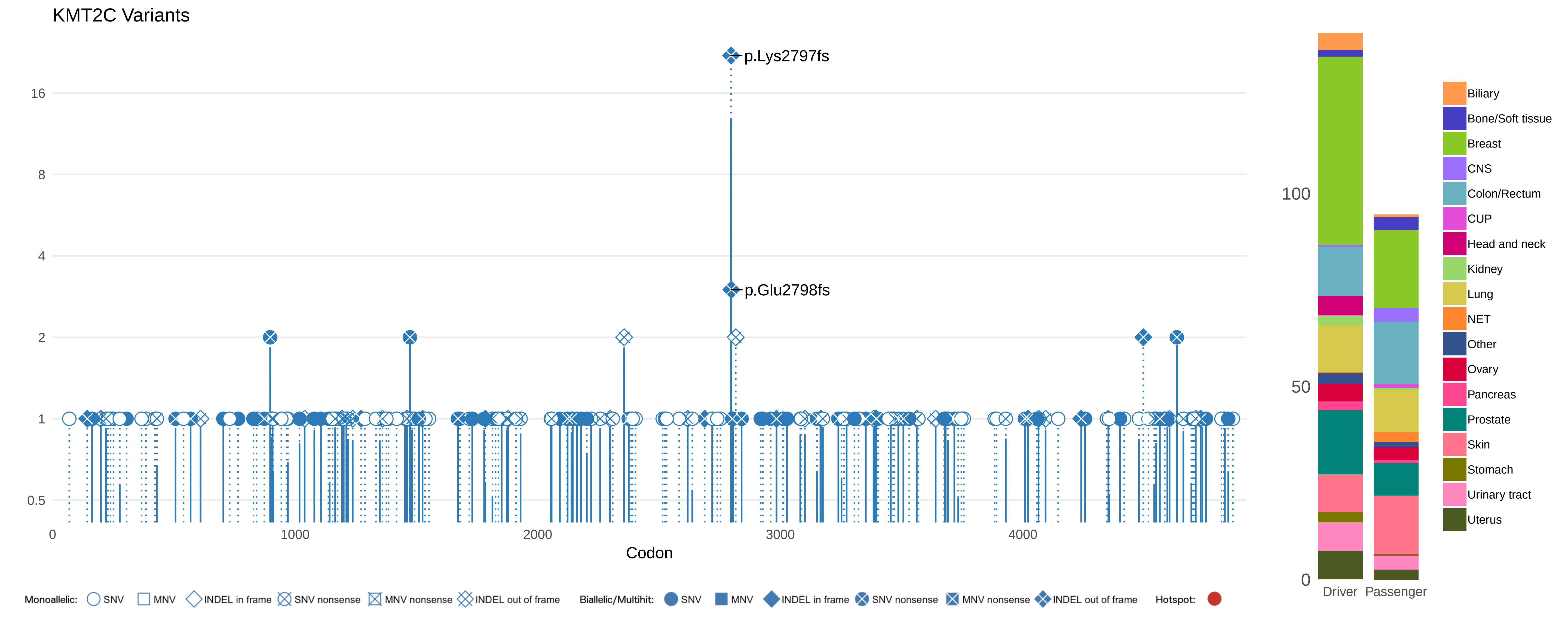

KMT2C Variants
p.Lys2797fs
p.Glu2798fs
16
8
4
2
1
0.5
0
1000
2000
3000
4000
Codon
100
50
0
Driver
Passenger
Biliary
Bone/Soft tissue
Breast
CNS
Colon/Rectum
CUP
Head and neck
Kidney
Lung
NET
Other
Ovary
Pancreas
Prostate
Skin
Stomach
Urinary tract
Uterus
Monoallelic: SNV MNV INDEL in frame SNV nonsense MNV nonsense INDEL out of frame
Biallelic/Multihit: SNV MNV INDEL in frame SNV nonsense MNV nonsense INDEL out of frame
Hotspot:

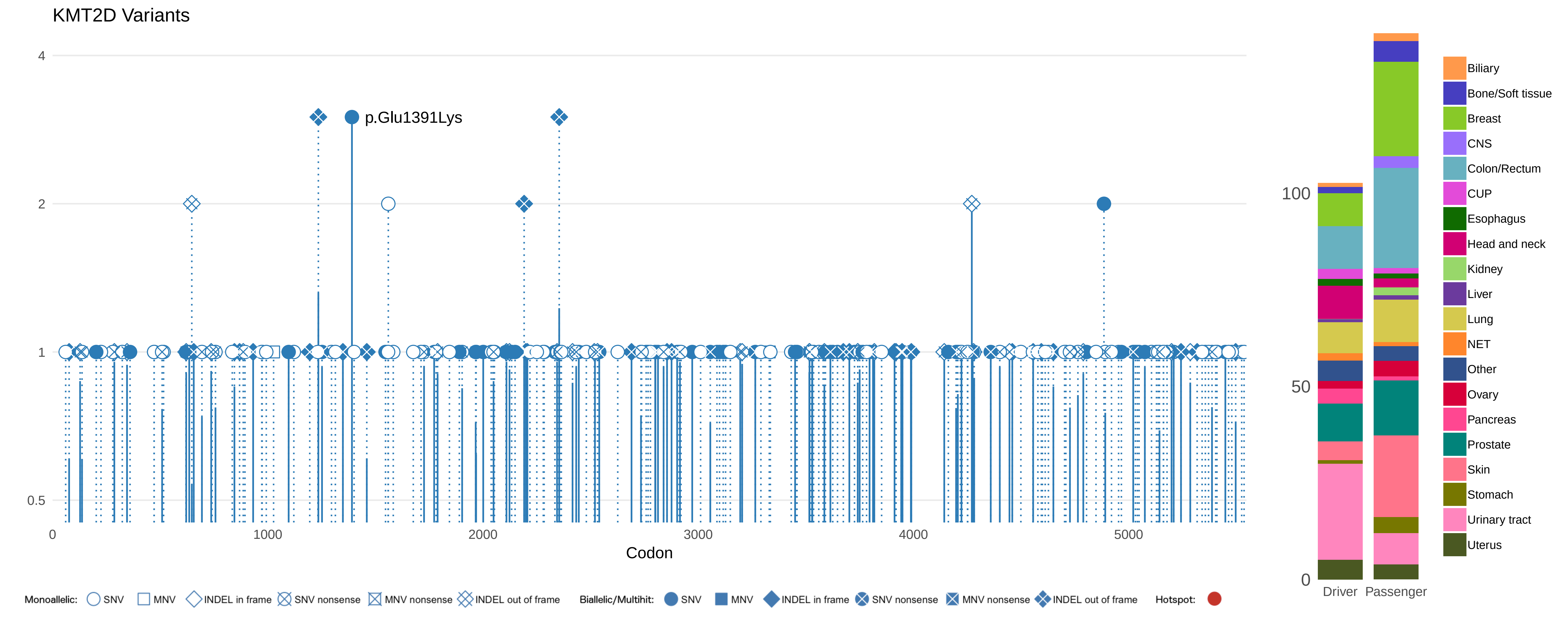

KMT2D Variants
p.Glu1391Lys
4
2
1
0.5
0
1000
2000
3000
4000
5000
Codon
Monoallelic: SNV MNV INDEL in frame SNV nonsense MNV nonsense INDEL out of frame
Biallelic/Multihit: SNV MNV INDEL in frame SNV nonsense MNV nonsense INDEL out of frame
Hotspot:
100
50
0
Driver
Passenger
Biliary
Bone/Soft tissue
Breast
CNS
Colon/Rectum
CUP
Esophagus
Head and neck
Kidney
Liver
Lung
NET
Other
Ovary
Pancreas
Prostate
Skin
Stomach
Urinary tract
Uterus

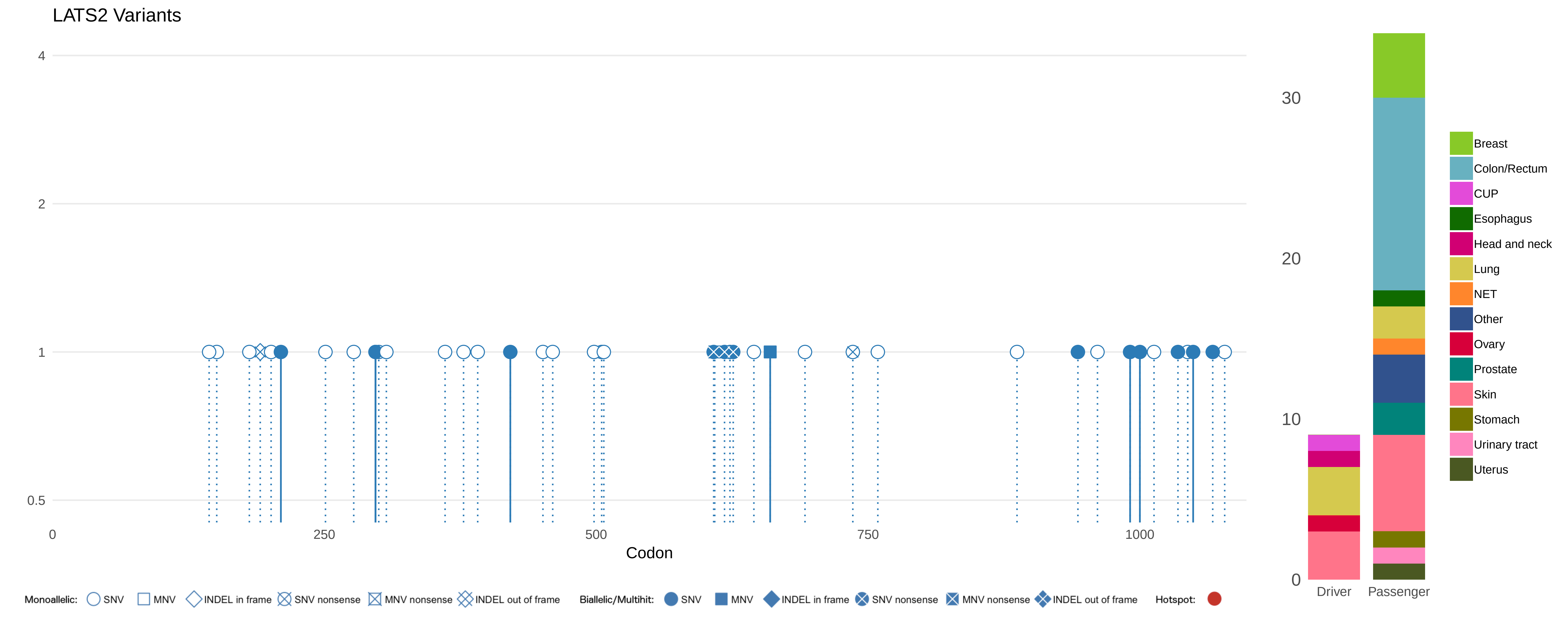
LATS2 Variants
4
2
1
0.5
0
250
500
750
1000
Codon
30
20
10
0
Driver
Passenger
Breast
Colon/Rectum
CUP
Esophagus
Head and neck
Lung
NET
Other
Ovary
Prostate
Skin
Stomach
Urinary tract
Uterus
Monoallelic: SNV MNV INDEL in frame SNV nonsense MNV nonsense INDEL out of frame
Biallelic/Multihit: SNV MNV INDEL in frame SNV nonsense MNV nonsense INDEL out of frame
Hotspot:

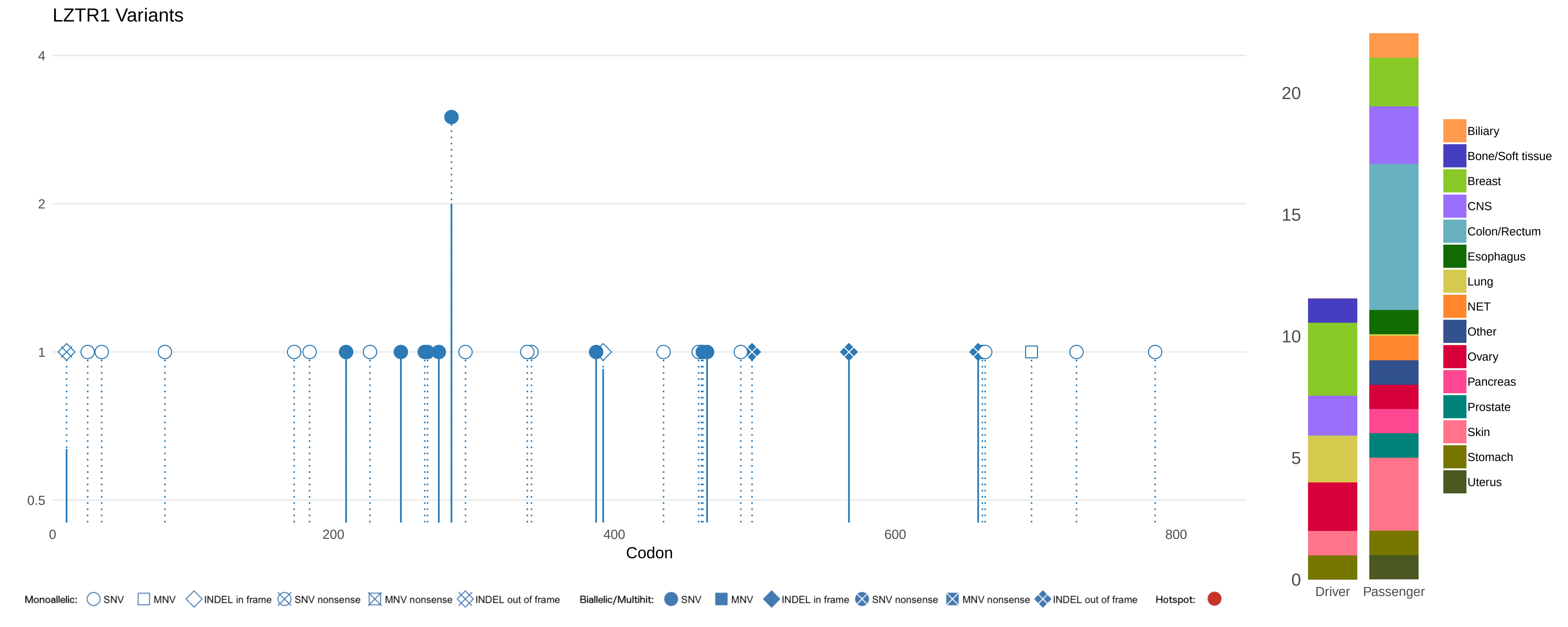
LZTR1 Variants
4
2
1
0.5
0
200
400
600
800
Codon
20
15
10
5
0
Driver
Passenger
Biliary
Bone/Soft tissue
Breast
CNS
Colon/Rectum
Esophagus
Lung
NET
Other
Ovary
Pancreas
Prostate
Skin
Stomach
Uterus
Monoallelic: SNV MNV INDEL in frame SNV nonsense MNV nonsense INDEL out of frame
Biallelic/Multihit: SNV MNV INDEL in frame SNV nonsense MNV nonsense INDEL out of frame
Hotspot:

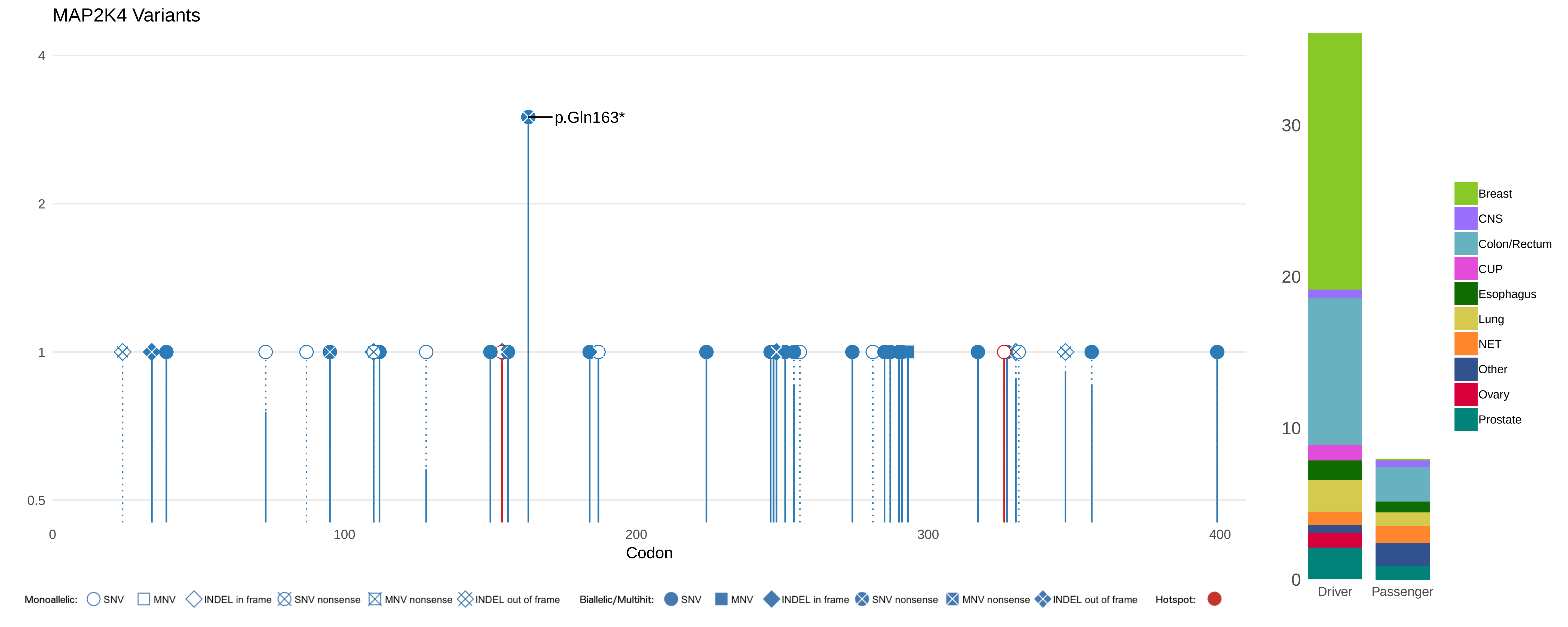

MAP2K4 Variants
p.Gln163*
4
2
1
0.5
0
100
200
300
400
Codon
30
20
10
0
Driver
Passenger
Breast
CNS
Colon/Rectum
CUP
Esophagus
Lung
NET
Other
Ovary
Prostate
Monoallelic: SNV MNV INDEL in frame SNV nonsense MNV nonsense INDEL out of frame
Biallelic/Multihit: SNV MNV INDEL in frame SNV nonsense MNV nonsense INDEL out of frame
Hotspot:

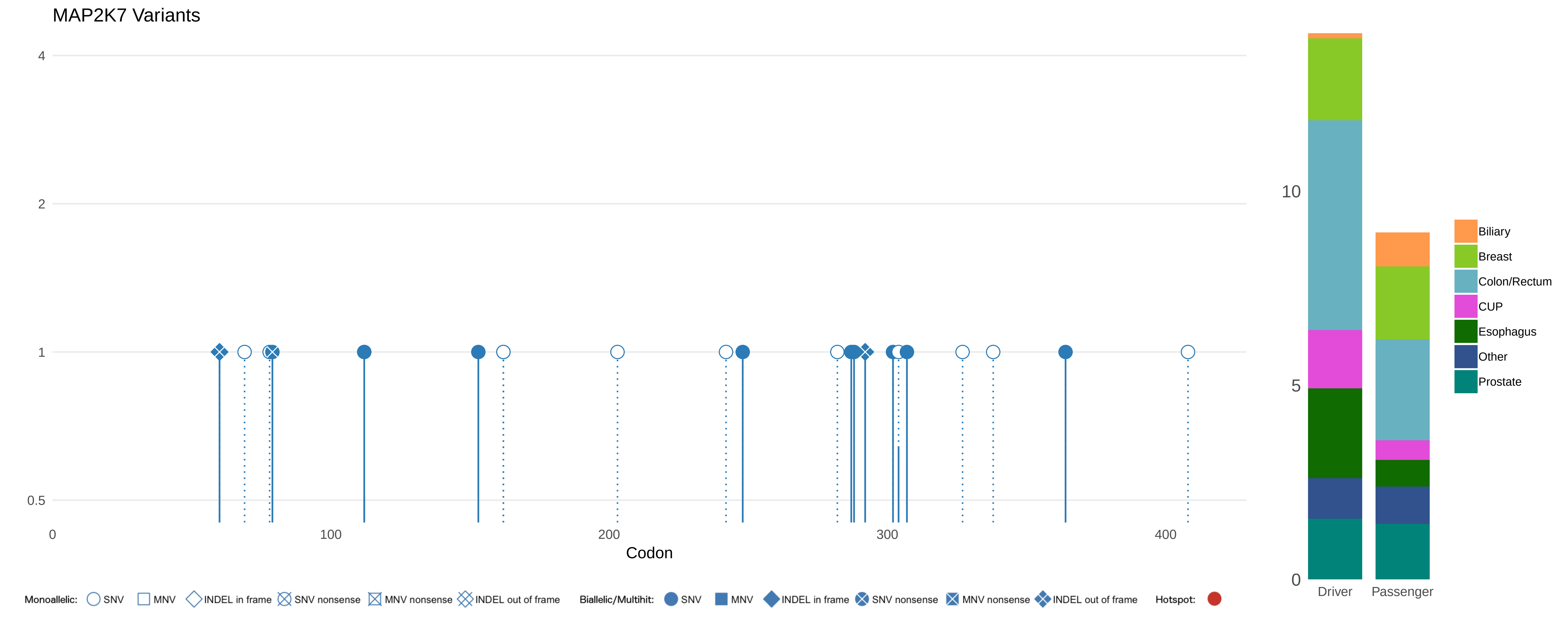
MAP2K7 Variants
4
2
1
0.5
0
100
200
300
400
Codon
Monoallelic: SNV MNV INDEL in frame SNV nonsense MNV nonsense INDEL out of frame
Biallelic/Multihit: SNV MNV INDEL in frame SNV nonsense MNV nonsense INDEL out of frame
Hotspot:
10
5
0
Driver
Passenger
Biliary
Breast
Colon/Rectum
CUP
Esophagus
Other
Prostate

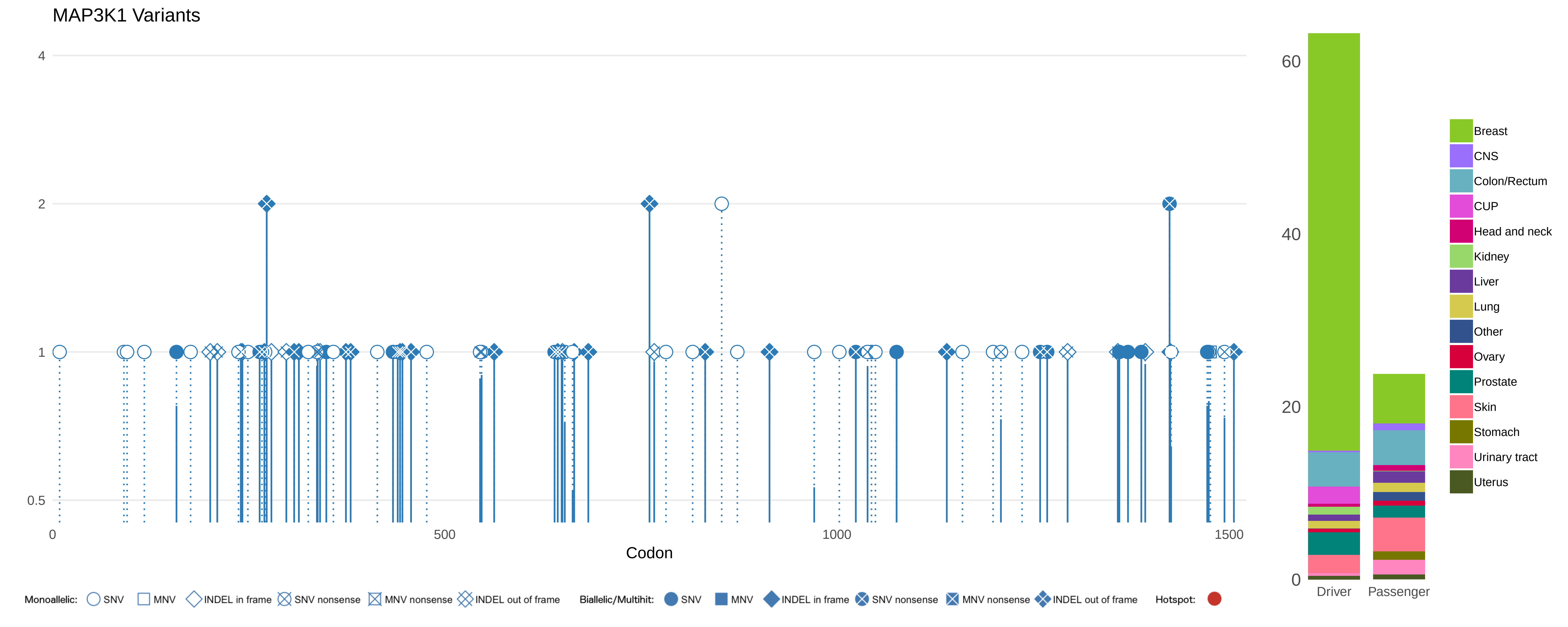

MAP3K1 Variants
4
2
1
0.5
0
500
1000
1500
Codon
60
40
20
0
Driver
Passenger
Breast
CNS
Colon/Rectum
CUP
Head and neck
Kidney
Liver
Lung
Other
Ovary
Prostate
Skin
Stomach
Urinary tract
Uterus
Monoallelic: SNV MNV INDEL in frame SNV nonsense MNV nonsense INDEL out of frame
Biallelic/Multihit: SNV MNV INDEL in frame SNV nonsense MNV nonsense INDEL out of frame
Hotspot:

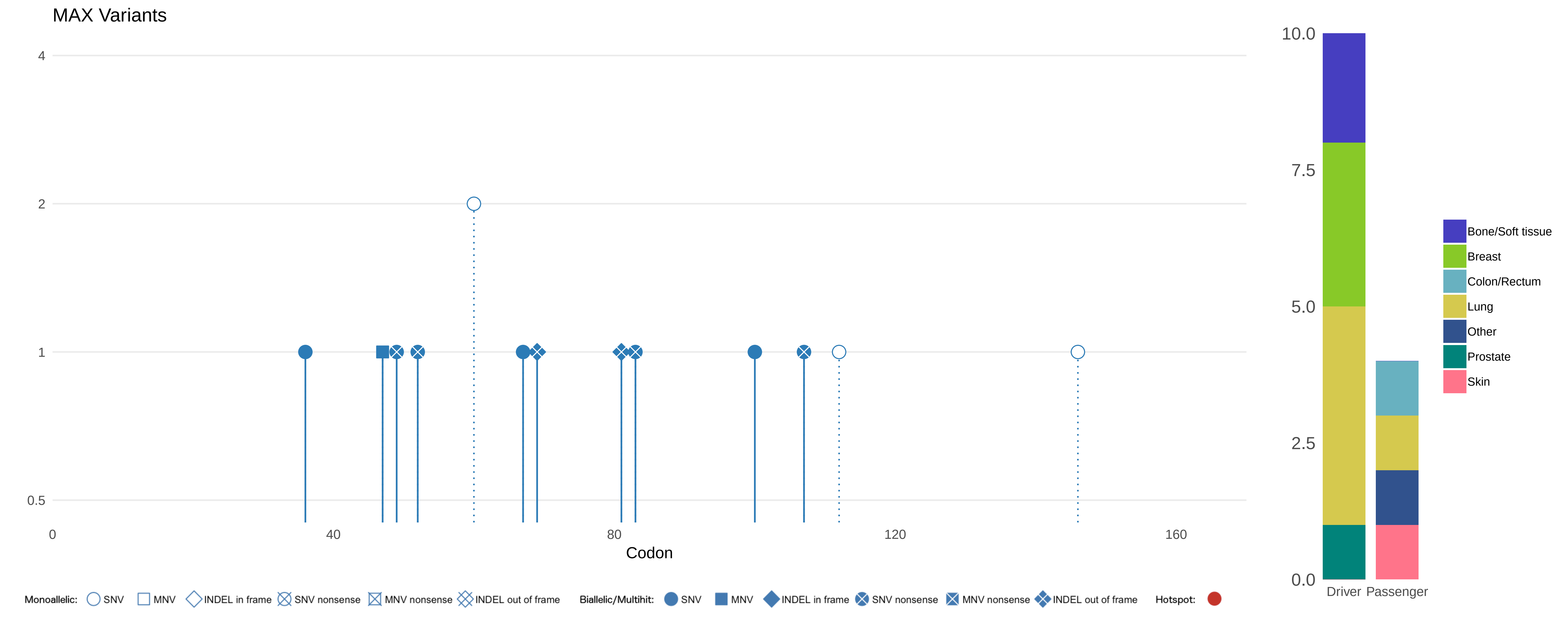
MAX Variants
4
2
1
0.5
0
40
80
120
160
Codon
10.0
7.5
5.0
2.5
0.0
Driver
Passenger
Bone/Soft tissue
Breast
Colon/Rectum
Lung
Other
Prostate
Skin
Monoallelic:
SNV
MNV
INDEL in frame
SNV nonsense
MNV nonsense
INDEL out of frame
Biallelic/Multihit:
SNV
MNV
INDEL in frame
SNV nonsense
MNV nonsense
INDEL out of frame
Hotspot:

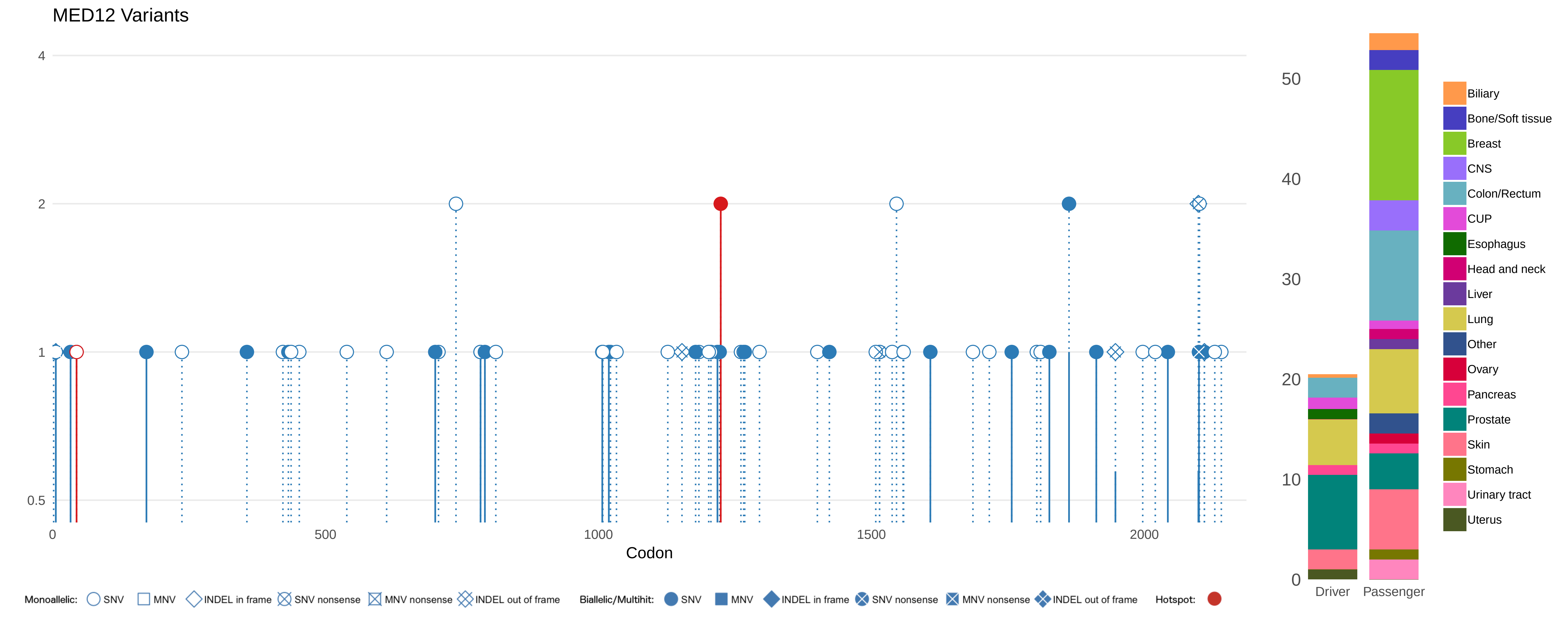

MED12 Variants
4
2
1
0.5
0
500
1000
1500
2000
Codon
Monoallelic: SNV MNV INDEL in frame SNV nonsense MNV nonsense INDEL out of frame
Biallelic/Multihit: SNV MNV INDEL in frame SNV nonsense MNV nonsense INDEL out of frame
Hotspot:
50
40
30
20
10
0
Driver
Passenger
Biliary
Bone/Soft tissue
Breast
CNS
Colon/Rectum
CUP
Esophagus
Head and neck
Liver
Lung
Other
Ovary
Pancreas
Prostate
Skin
Stomach
Urinary tract
Uterus

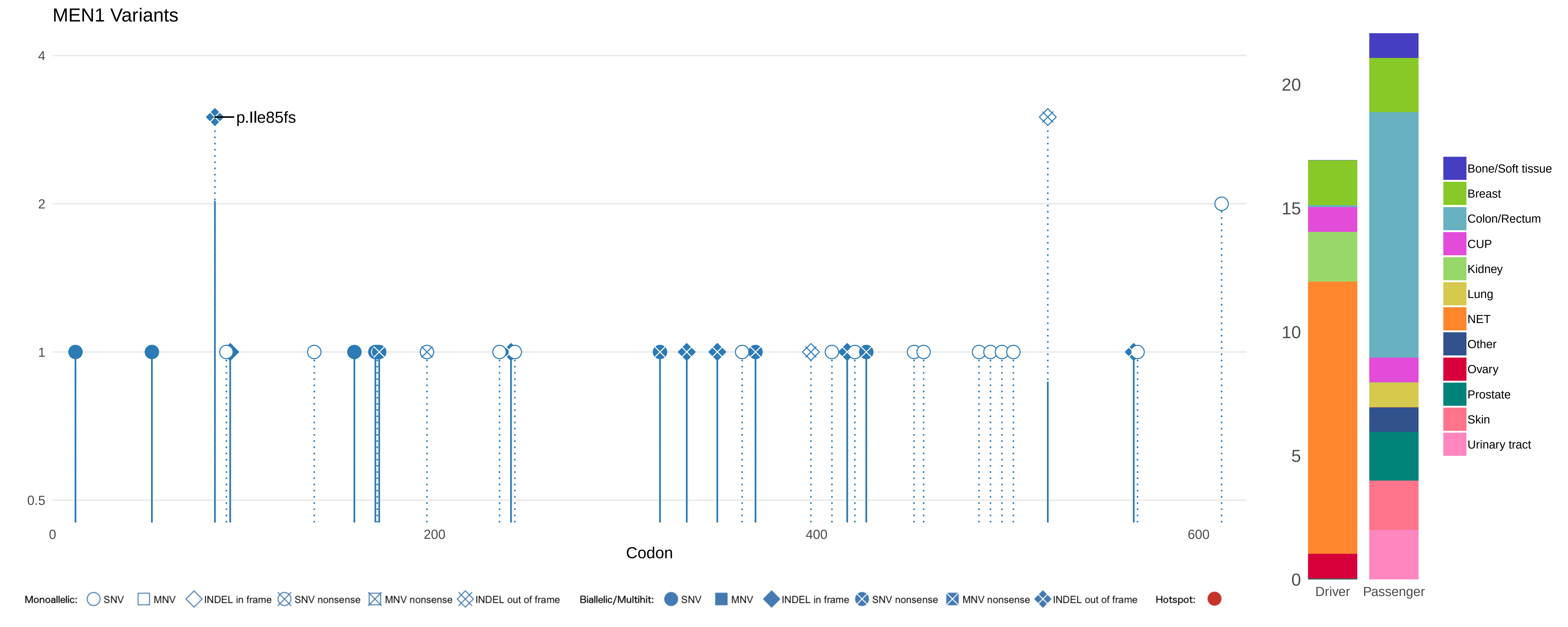
MEN1 Variants
p.Ile85fs
4
2
1
0.5
0
200
400
600
Codon
20
15
10
5
0
Driver
Passenger
Bone/Soft tissue
Breast
Colon/Rectum
CUP
Kidney
Lung
NET
Other
Ovary
Prostate
Skin
Urinary tract
Monoallelic:
SNV
MNV
INDEL in frame
SNV nonsense
MNV nonsense
INDEL out of frame
Biallelic/Multihit:
SNV
MNV
INDEL in frame
SNV nonsense
MNV nonsense
INDEL out of frame
Hotspot:

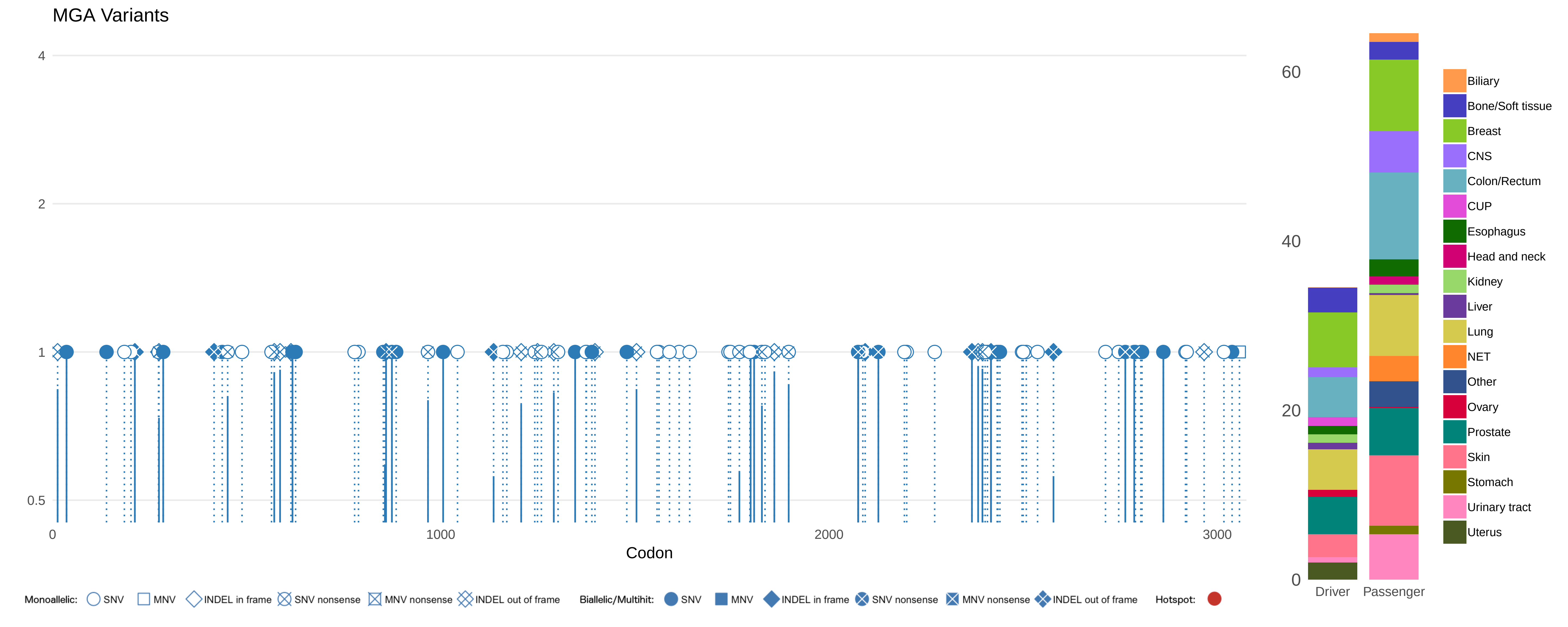

MGA Variants
4
2
1
0.5
0
1000
2000
3000
Codon
Monoallelic: SNV MNV INDEL in frame SNV nonsense MNV nonsense INDEL out of frame
Biallelic/Multihit: SNV MNV INDEL in frame SNV nonsense MNV nonsense INDEL out of frame
Hotspot:
60
40
20
0
Driver
Passenger
Biliary
Bone/Soft tissue
Breast
CNS
Colon/Rectum
CUP
Esophagus
Head and neck
Kidney
Liver
Lung
NET
Other
Ovary
Prostate
Skin
Stomach
Urinary tract
Uterus

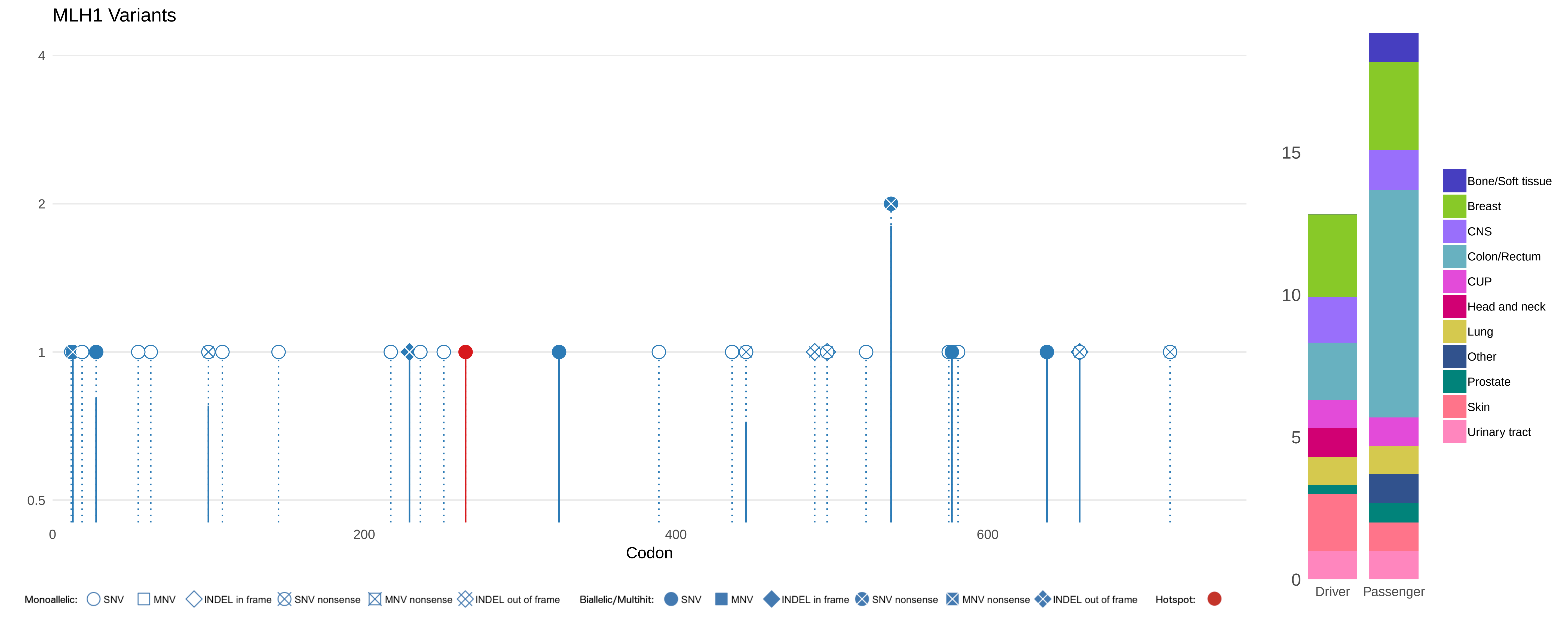
MLH1 Variants
4
2
1
0.5
0
200
400
600
Codon
15
10
5
0
Driver
Passenger
Bone/Soft tissue
Breast
CNS
Colon/Rectum
CUP
Head and neck
Lung
Other
Prostate
Skin
Urinary tract
Monoallelic: SNV MNV INDEL in frame SNV nonsense MNV nonsense INDEL out of frame
Biallelic/Multihit: SNV MNV INDEL in frame SNV nonsense MNV nonsense INDEL out of frame
Hotspot:

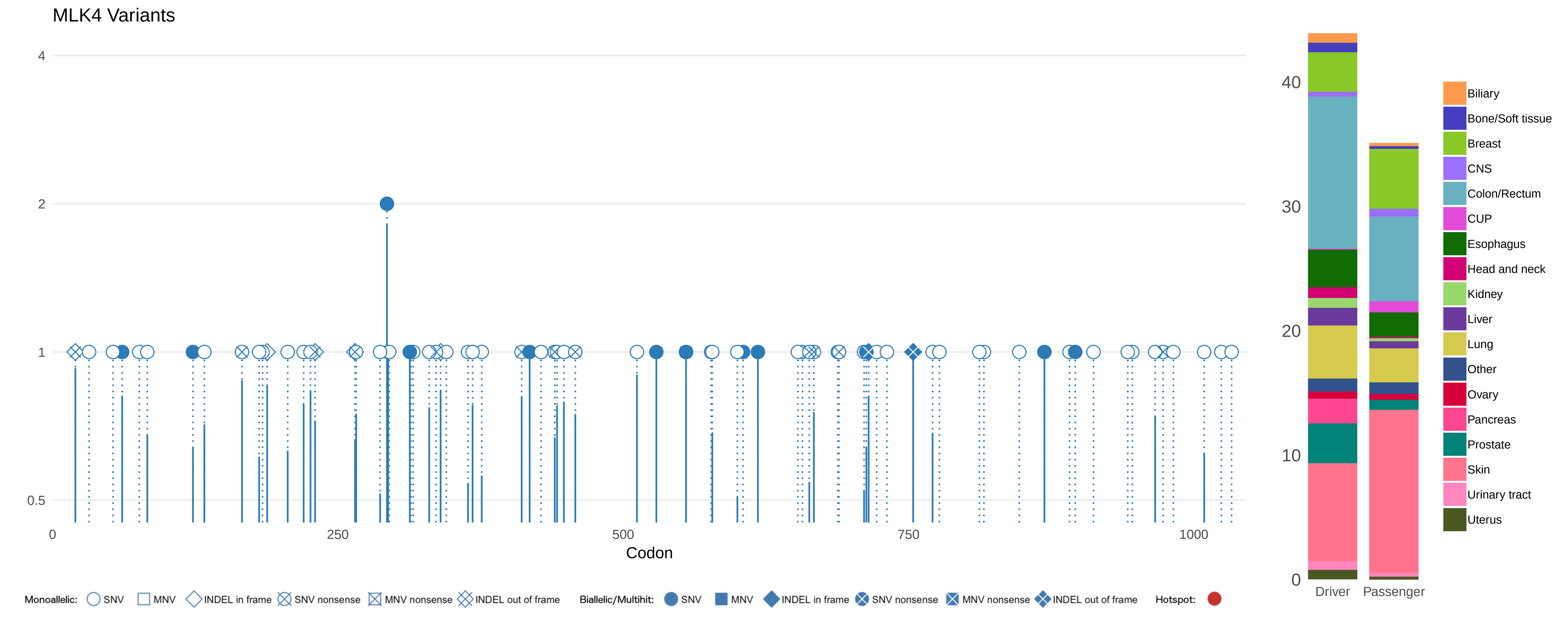

MLK4 Variants
4
2
1
0.5
0
250
500
750
1000
Codon
40
30
20
10
0
Driver
Passenger
Biliary
Bone/Soft tissue
Breast
CNS
Colon/Rectum
CUP
Esophagus
Head and neck
Kidney
Liver
Lung
Other
Ovary
Pancreas
Prostate
Skin
Urinary tract
Uterus
Monoallelic: SNV MNV INDEL in frame SNV nonsense MNV nonsense INDEL out of frame
Biallelic/Multihit: SNV MNV INDEL in frame SNV nonsense MNV nonsense INDEL out of frame
Hotspot:

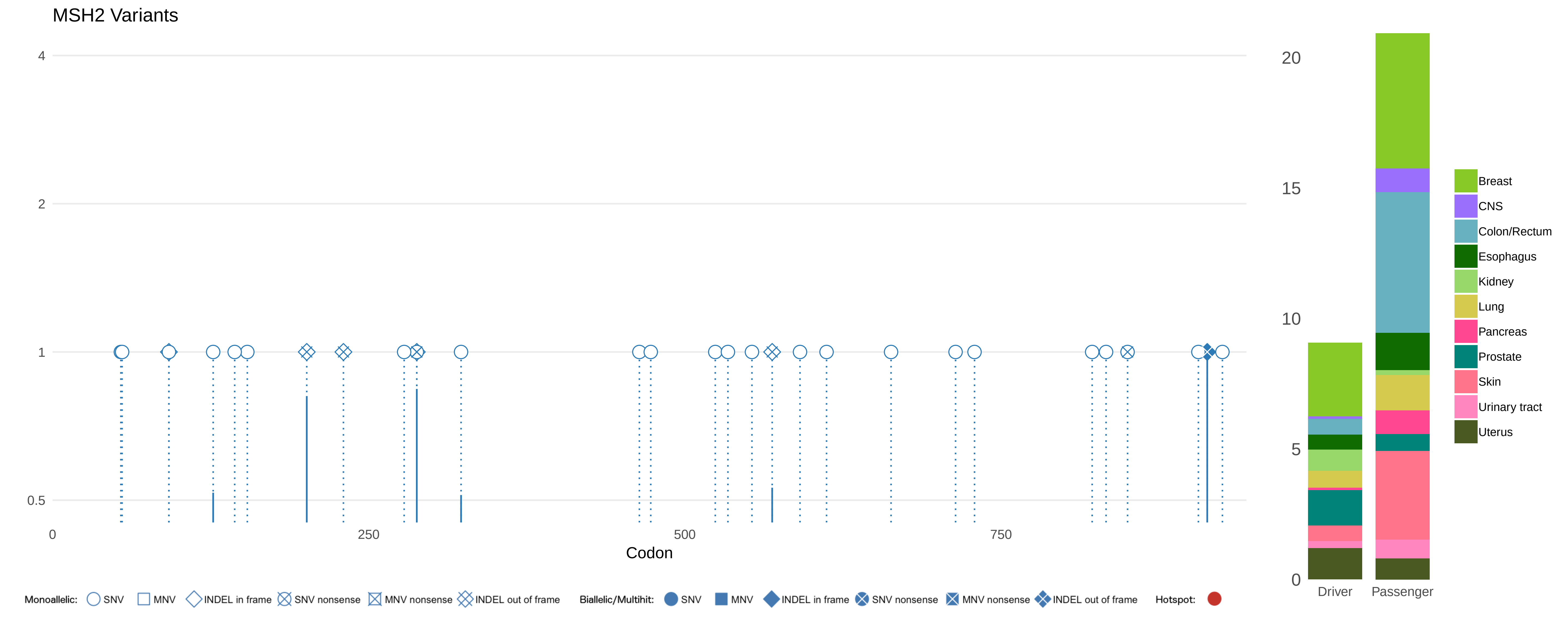
MSH2 Variants
4
2
1
0.5
0
250
500
750
Codon
20
15
10
5
0
Driver
Passenger
Breast
CNS
Colon/Rectum
Esophagus
Kidney
Lung
Pancreas
Prostate
Skin
Urinary tract
Uterus
Monoallelic: SNV MNV INDEL in frame SNV nonsense MNV nonsense INDEL out of frame
Biallelic/Multihit: SNV MNV INDEL in frame SNV nonsense MNV nonsense INDEL out of frame
Hotspot:

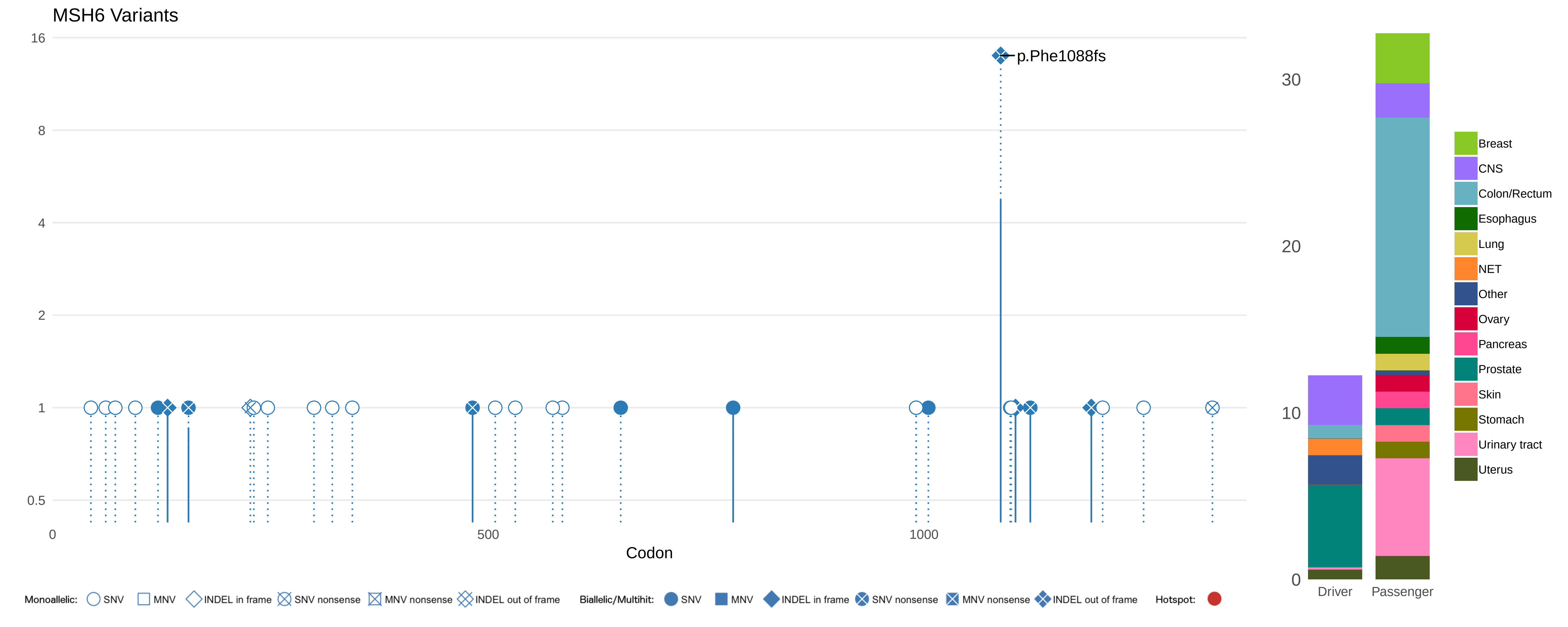

MSH6 Variants
16
8
4
2
1
0.5
0
500
1000
Codon
p.Phe1088fs
30
20
10
0
Driver
Passenger
Breast
CNS
Colon/Rectum
Esophagus
Lung
NET
Other
Ovary
Pancreas
Prostate
Skin
Stomach
Urinary tract
Uterus
Monoallelic: SNV MNV INDEL in frame SNV nonsense MNV nonsense INDEL out of frame
Biallelic/Multihit: SNV MNV INDEL in frame SNV nonsense MNV nonsense INDEL out of frame
Hotspot:

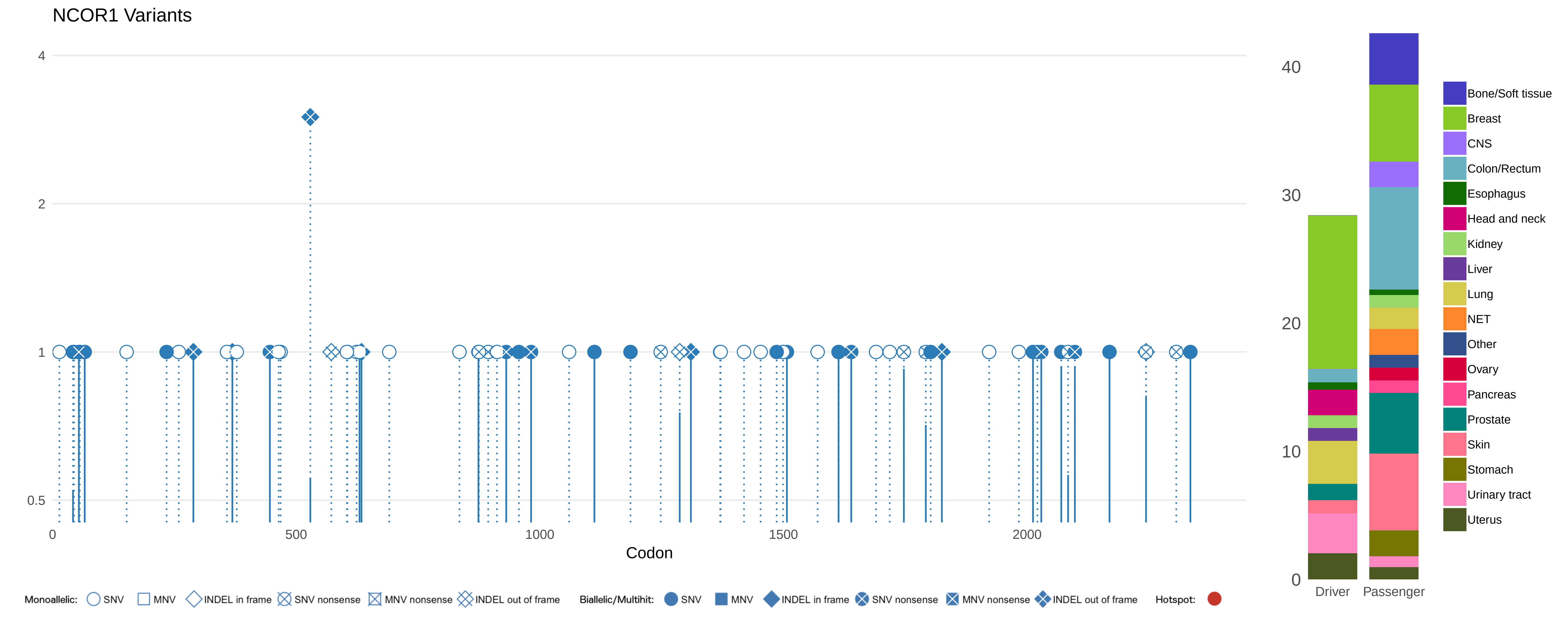

NCOR1 Variants
4
2
1
0.5
0
500
1000
1500
2000
Codon
Monoallelic: SNV MNV INDEL in frame SNV nonsense MNV nonsense INDEL out of frame
Biallelic/Multihit: SNV MNV INDEL in frame SNV nonsense MNV nonsense INDEL out of frame
Hotspot:
40
30
20
10
0
Driver
Passenger
Bone/Soft tissue
Breast
CNS
Colon/Rectum
Esophagus
Head and neck
Kidney
Liver
Lung
NET
Other
Ovary
Pancreas
Prostate
Skin
Stomach
Urinary tract
Uterus

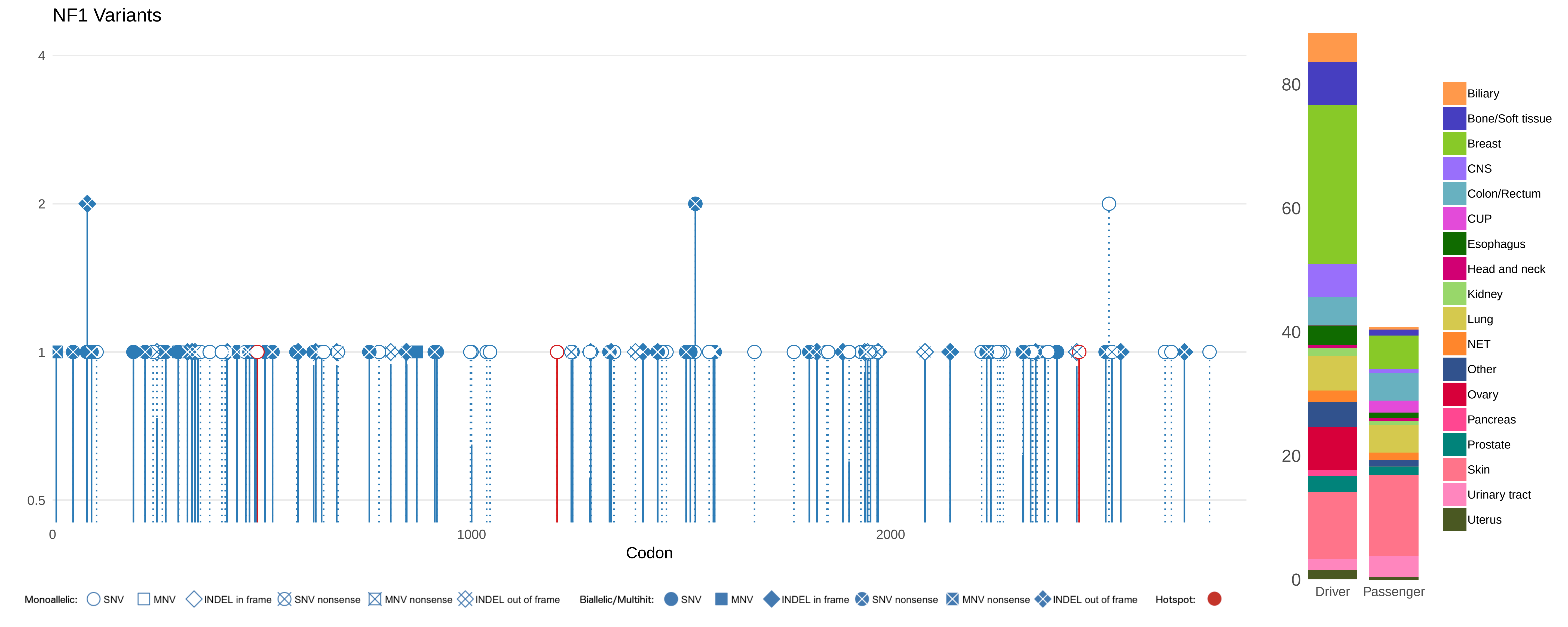

NF1 Variants
4
2
1
0.5
0
1000
2000
Codon
Monoallelic: SNV MNV INDEL in frame SNV nonsense MNV nonsense INDEL out of frame
Biallelic/Multihit: SNV MNV INDEL in frame SNV nonsense MNV nonsense INDEL out of frame
Hotspot:
0
20
40
60
80
Driver
Passenger
Biliary
Bone/Soft tissue
Breast
CNS
Colon/Rectum
CUP
Esophagus
Head and neck
Kidney
Lung
NET
Other
Ovary
Pancreas
Prostate
Skin
Urinary tract
Uterus

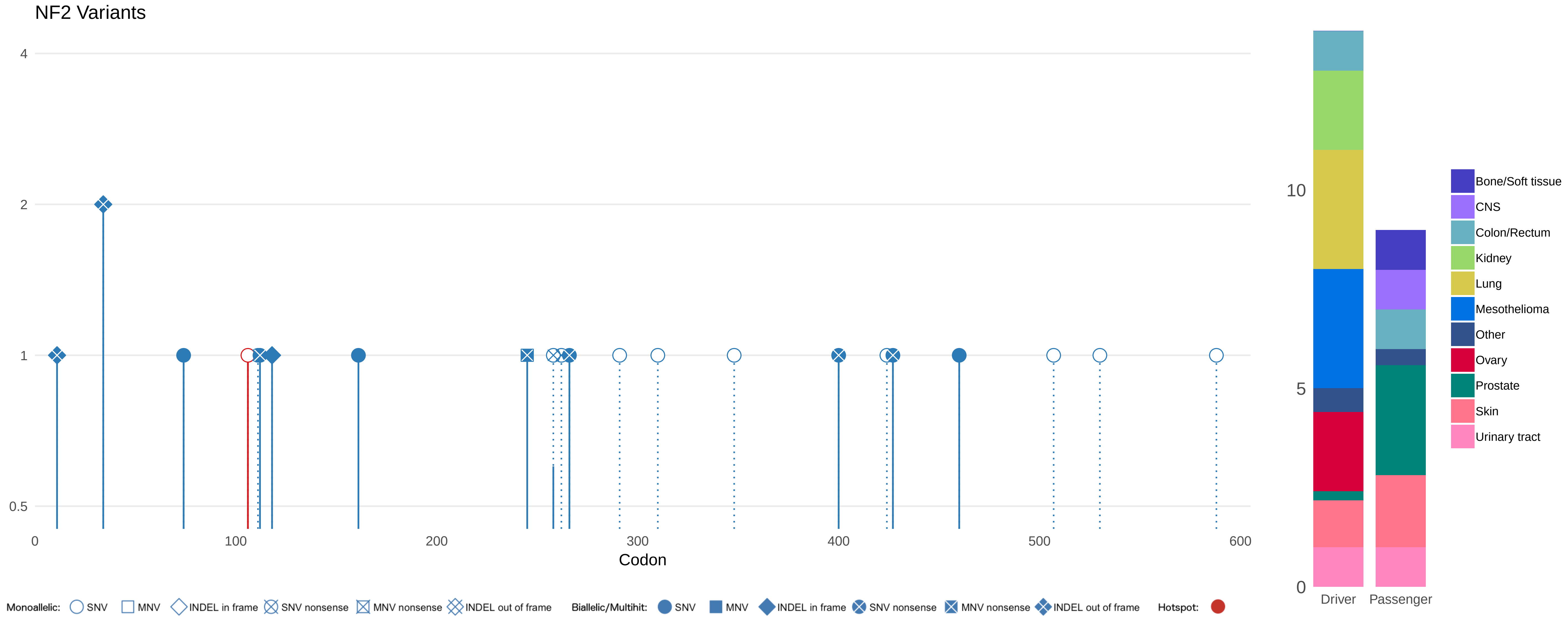

NF2 Variants
4
2
1
0.5
0
100
200
300
400
500
600
Codon
10
5
0
Driver
Passenger
Bone/Soft tissue
CNS
Colon/Rectum
Kidney
Lung
Mesothelioma
Other
Ovary
Prostate
Skin
Urinary tract
Monoallelic: SNV MNV INDEL in frame SNV nonsense MNV nonsense INDEL out of frame
Biallelic/Multihit: SNV MNV INDEL in frame SNV nonsense MNV nonsense INDEL out of frame
Hotspot:

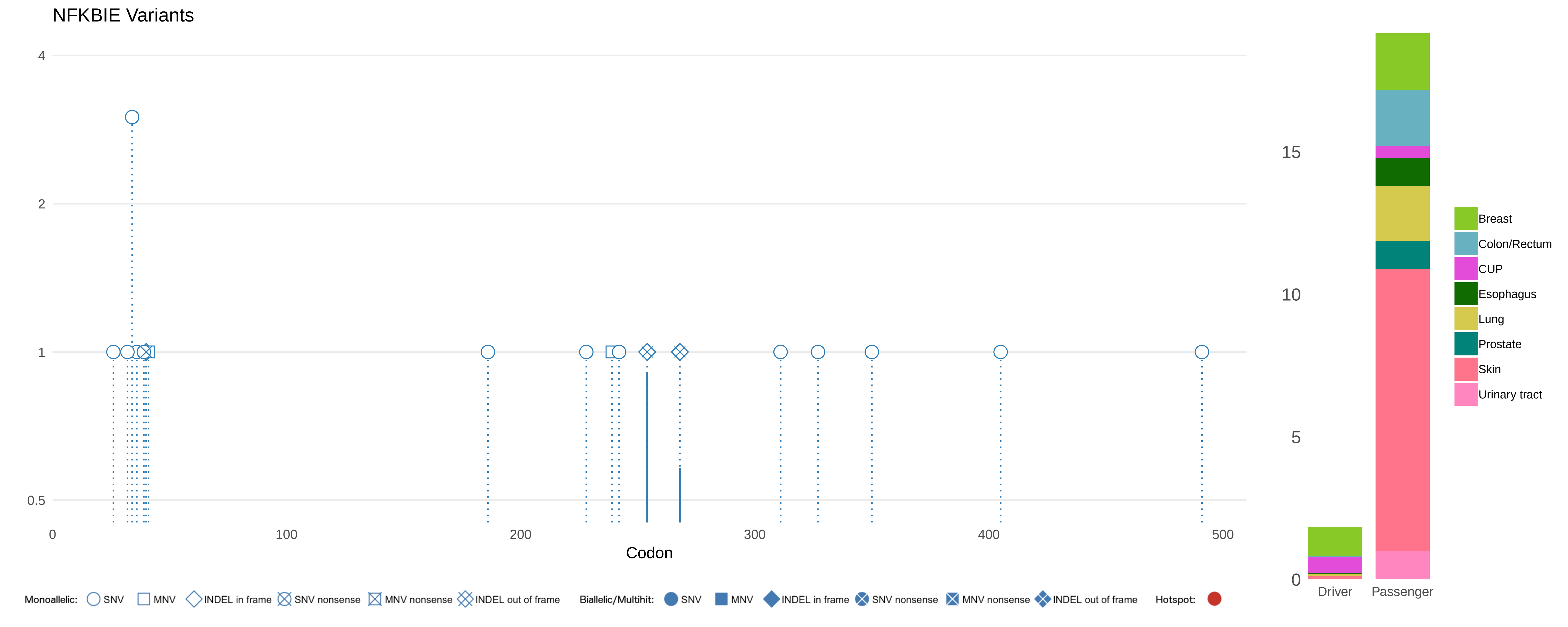

NFKBIE Variants
4
2
1
0.5
0
100
200
300
400
500
Codon
15
10
5
0
Driver
Passenger
Breast
Colon/Rectum
CUP
Esophagus
Lung
Prostate
Skin
Urinary tract
Monoallelic: SNV MNV INDEL in frame SNV nonsense MNV nonsense INDEL out of frame
Biallelic/Multihit: SNV MNV INDEL in frame SNV nonsense MNV nonsense INDEL out of frame
Hotspot:

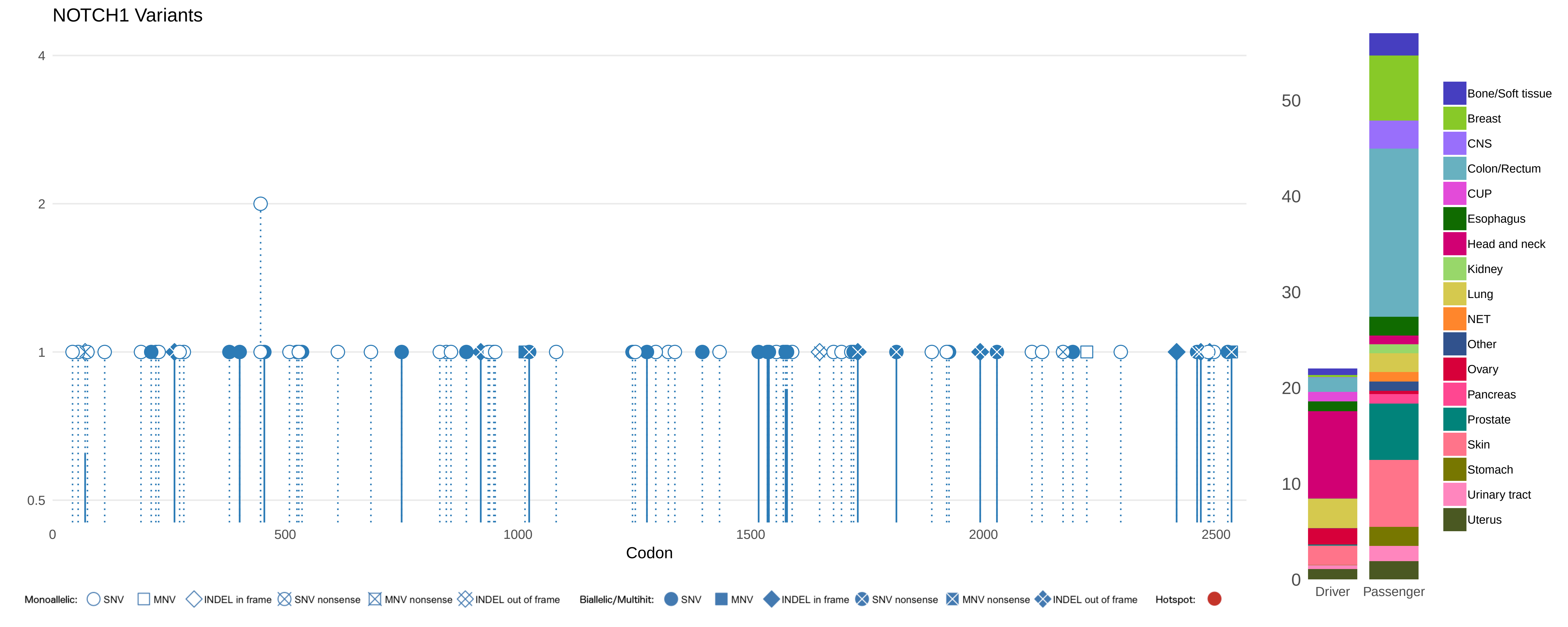

NOTCH1 Variants
4
2
1
0.5
0
500
1000
1500
2000
2500
Codon
50
40
30
20
10
0
Driver
Passenger
Bone/Soft tissue
Breast
CNS
Colon/Rectum
CUP
Esophagus
Head and neck
Kidney
Lung
NET
Other
Ovary
Pancreas
Prostate
Skin
Stomach
Urinary tract
Uterus
Monoallelic: SNV MNV INDEL in frame SNV nonsense MNV nonsense INDEL out of frame
Biallelic/Multihit: SNV MNV INDEL in frame SNV nonsense MNV nonsense INDEL out of frame
Hotspot:

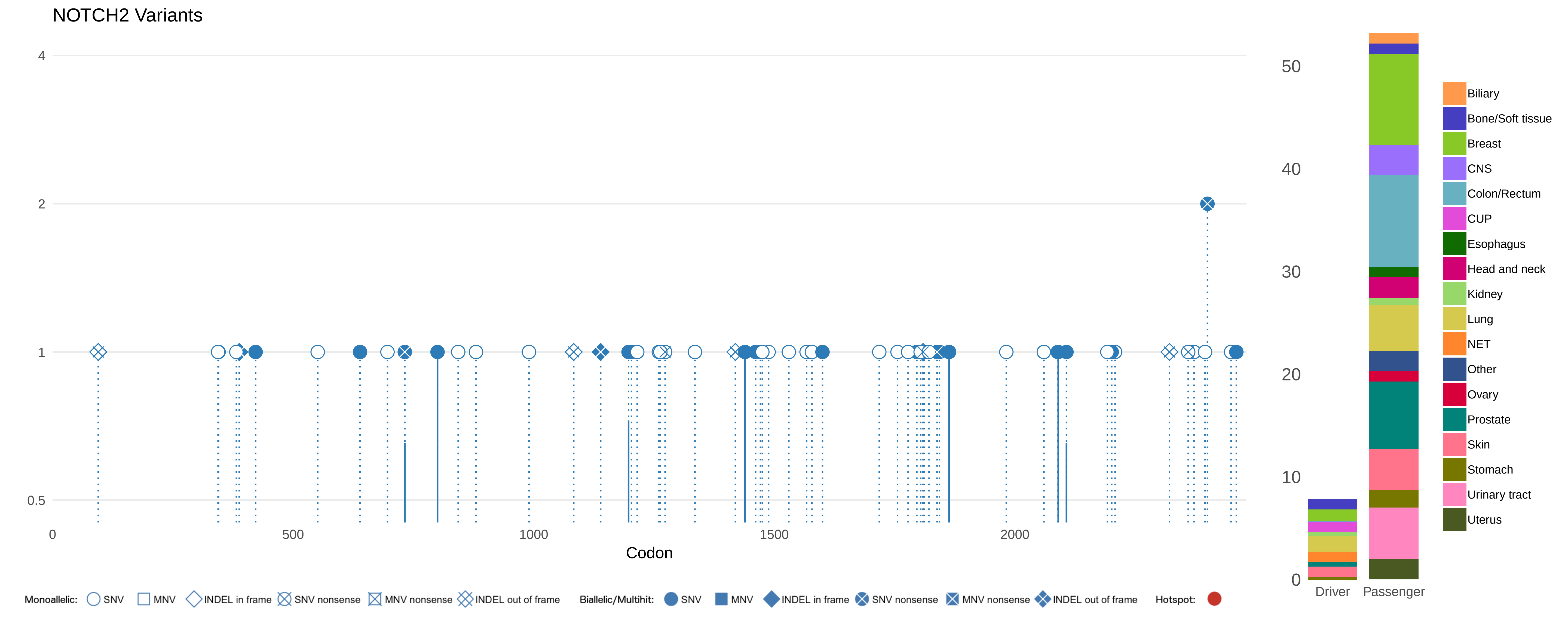

NOTCH2 Variants
4
2
1
0.5
0
500
1000
1500
2000
Codon
50
40
30
20
10
0
Driver
Passenger
Biliary
Bone/Soft tissue
Breast
CNS
Colon/Rectum
CUP
Esophagus
Head and neck
Kidney
Lung
NET
Other
Ovary
Prostate
Skin
Stomach
Urinary tract
Uterus
Monoallelic: SNV MNV INDEL in frame SNV nonsense MNV nonsense INDEL out of frame
Biallelic/Multihit: SNV MNV INDEL in frame SNV nonsense MNV nonsense INDEL out of frame
Hotspot:

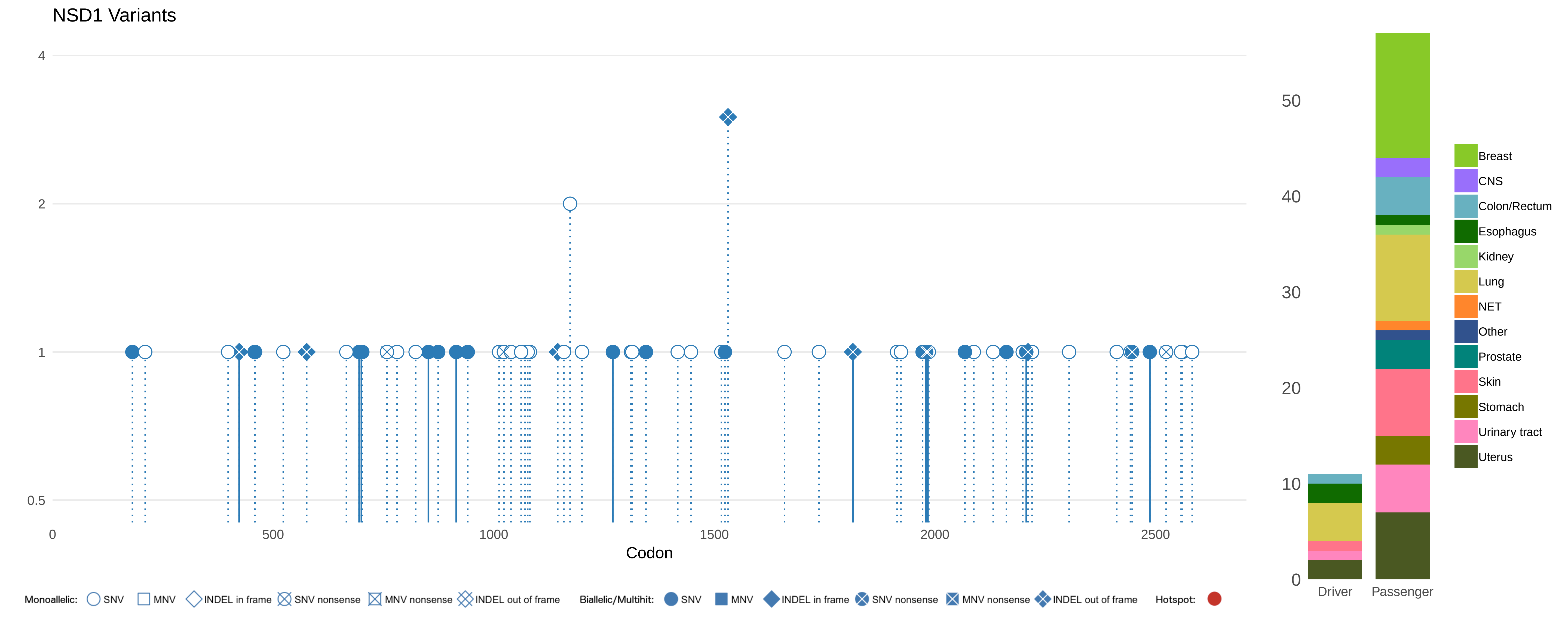
NSD1 Variants
4
2
1
0.5
0
500
1000
1500
2000
2500
Codon
Monoallelic: SNV MNV INDEL in frame SNV nonsense MNV nonsense INDEL out of frame
Biallelic/Multihit: SNV MNV INDEL in frame SNV nonsense MNV nonsense INDEL out of frame
Hotspot:
50
40
30
20
10
0
Driver
Passenger
Breast
CNS
Colon/Rectum
Esophagus
Kidney
Lung
NET
Other
Prostate
Skin
Stomach
Urinary tract
Uterus

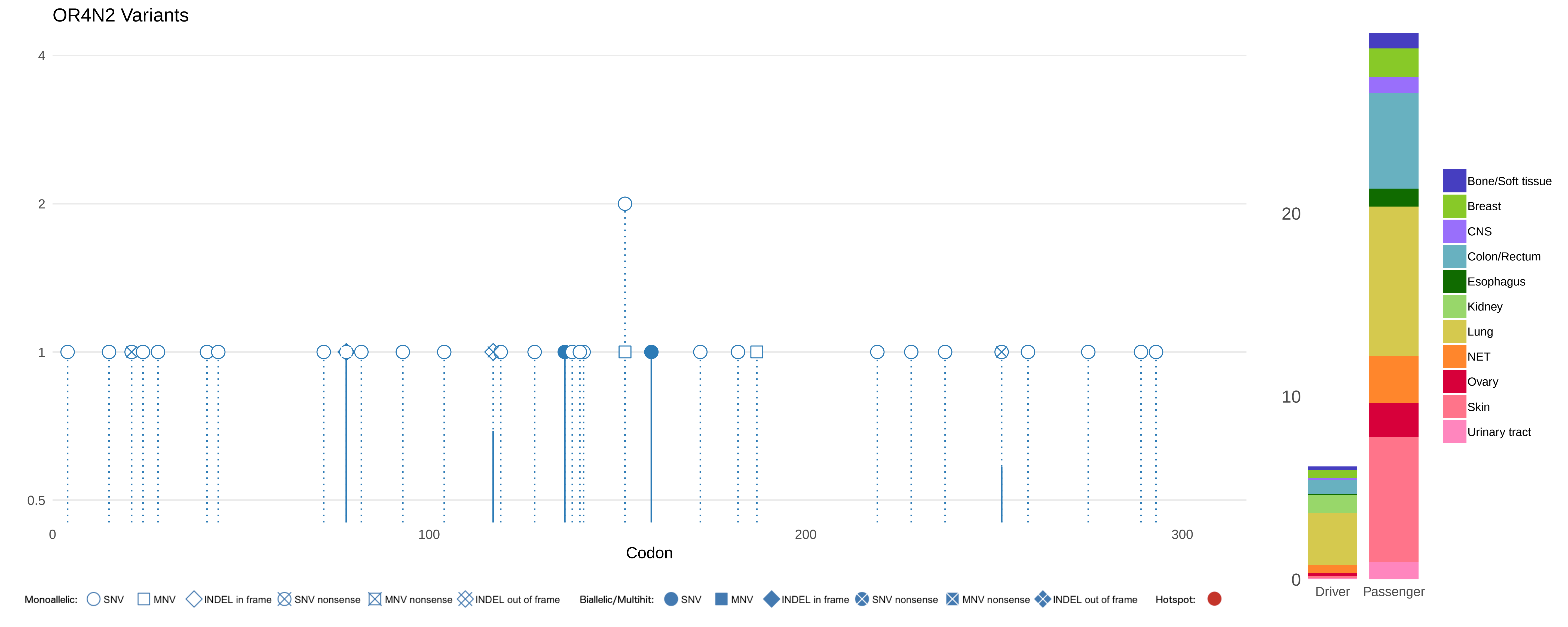
OR4N2 Variants
4
2
1
0.5
0
100
200
300
Codon
20
10
0
Driver
Passenger
Bone/Soft tissue
Breast
CNS
Colon/Rectum
Esophagus
Kidney
Lung
NET
Ovary
Skin
Urinary tract
Monoallelic: SNV MNV INDEL in frame SNV nonsense MNV nonsense INDEL out of frame
Biallelic/Multihit: SNV MNV INDEL in frame SNV nonsense MNV nonsense INDEL out of frame
Hotspot:

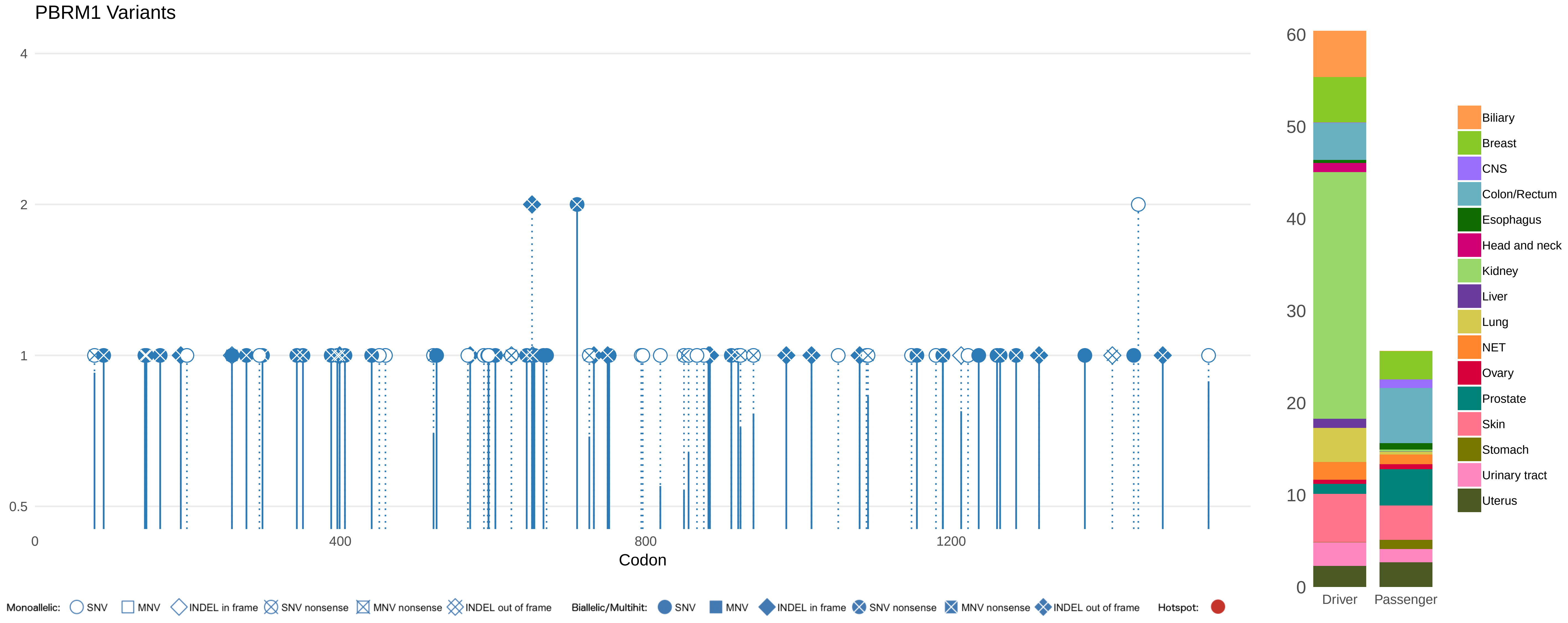

PBRM1 Variants
4
2
1
0.5
0
400
800
1200
Codon
60
50
40
30
20
10
0
Driver
Passenger
Biliary
Breast
CNS
Colon/Rectum
Esophagus
Head and neck
Kidney
Liver
Lung
NET
Ovary
Prostate
Skin
Stomach
Urinary tract
Uterus
Monoallelic: SNV MNV INDEL in frame SNV nonsense MNV nonsense INDEL out of frame
Biallelic/Multihit: SNV MNV INDEL in frame SNV nonsense MNV nonsense INDEL out of frame
Hotspot:

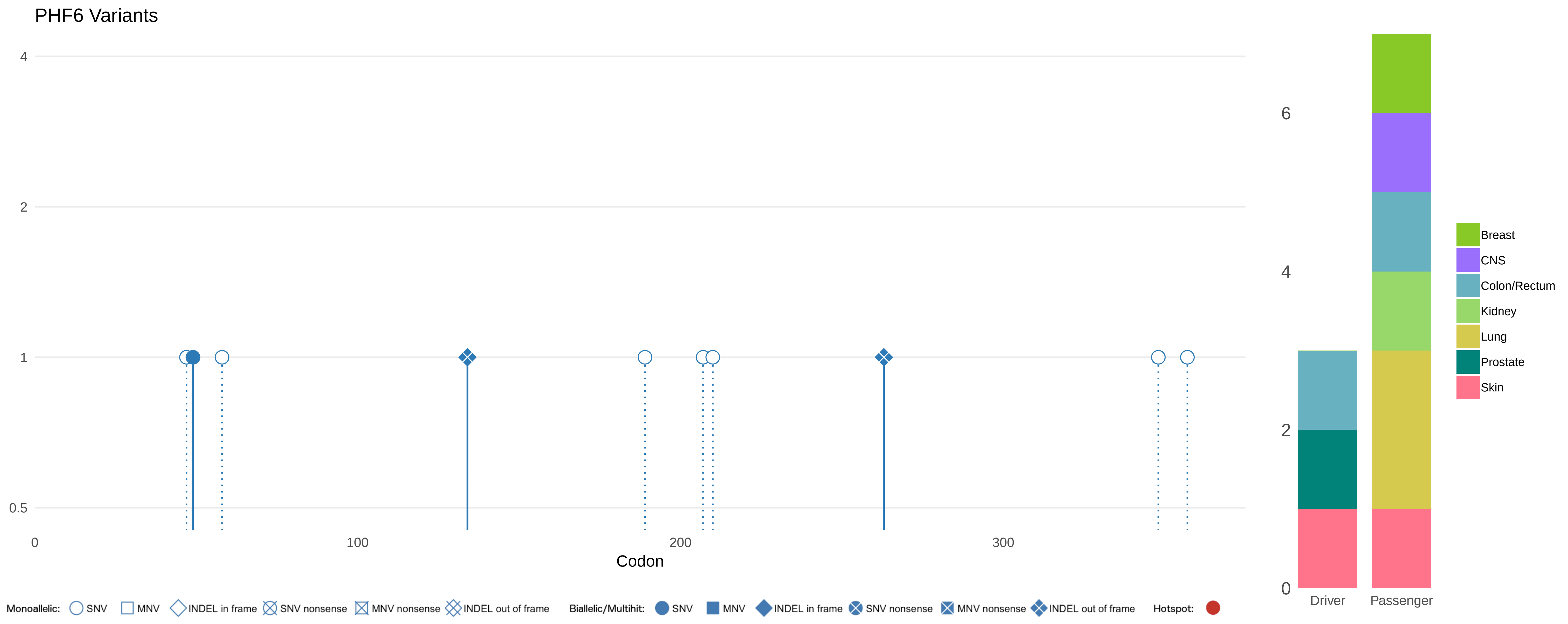
PHF6 Variants
4
2
1
0.5
0
100
200
300
Codon
Monoallelic: SNV MNV INDEL in frame SNV nonsense MNV nonsense INDEL out of frame
Biallelic/Multihit: SNV MNV INDEL in frame SNV nonsense MNV nonsense INDEL out of frame
Hotspot:
6
4
2
0
Driver
Passenger
Breast
CNS
Colon/Rectum
Kidney
Lung
Prostate
Skin

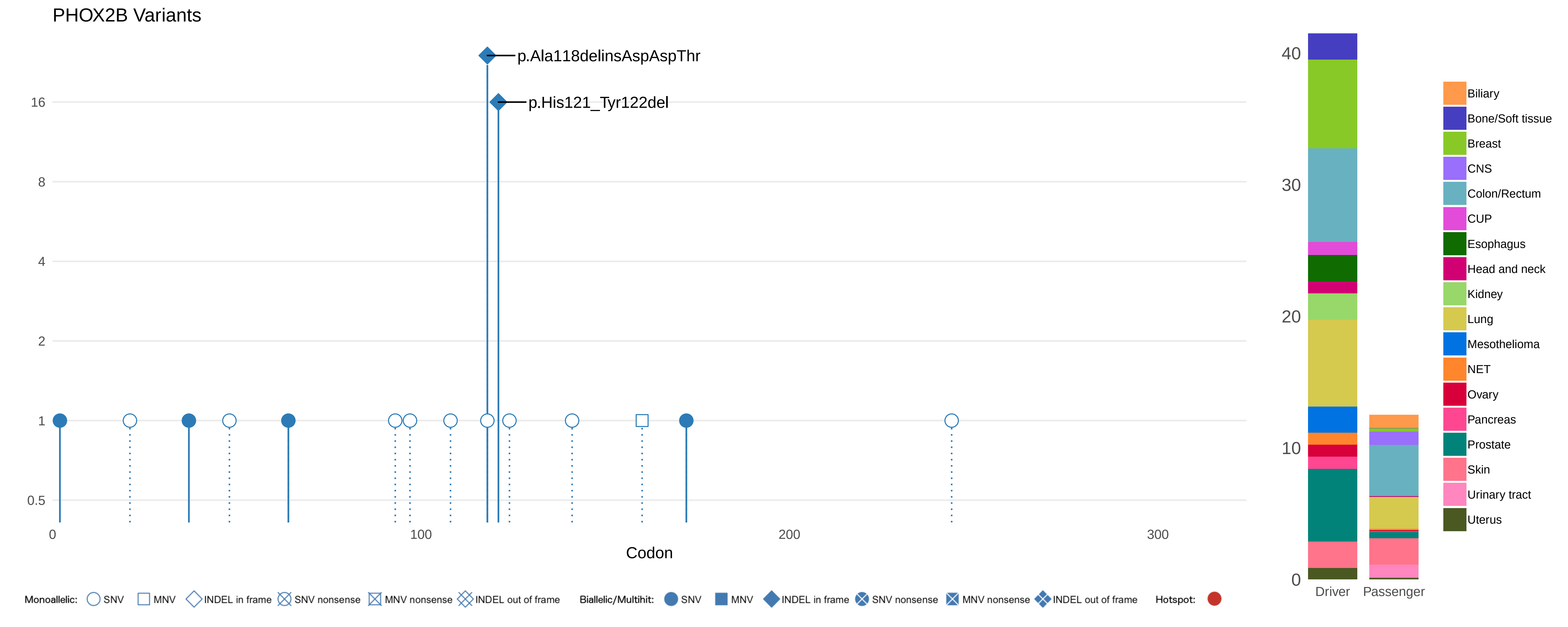

PHOX2B Variants
p.Ala118delinsAspAspThr
p.His121_Tyr122del
16
8
4
2
1
0.5
0
100
200
300
Codon
Monoallelic: SNV MNV INDEL in frame SNV nonsense MNV nonsense INDEL out of frame
Biallelic/Multihit: SNV MNV INDEL in frame SNV nonsense MNV nonsense INDEL out of frame
Hotspot:
40
30
20
10
0
Driver
Passenger
Biliary
Bone/Soft tissue
Breast
CNS
Colon/Rectum
CUP
Esophagus
Head and neck
Kidney
Lung
Mesothelioma
NET
Ovary
Pancreas
Prostate
Skin
Urinary tract
Uterus

### PIK3R1 Variants

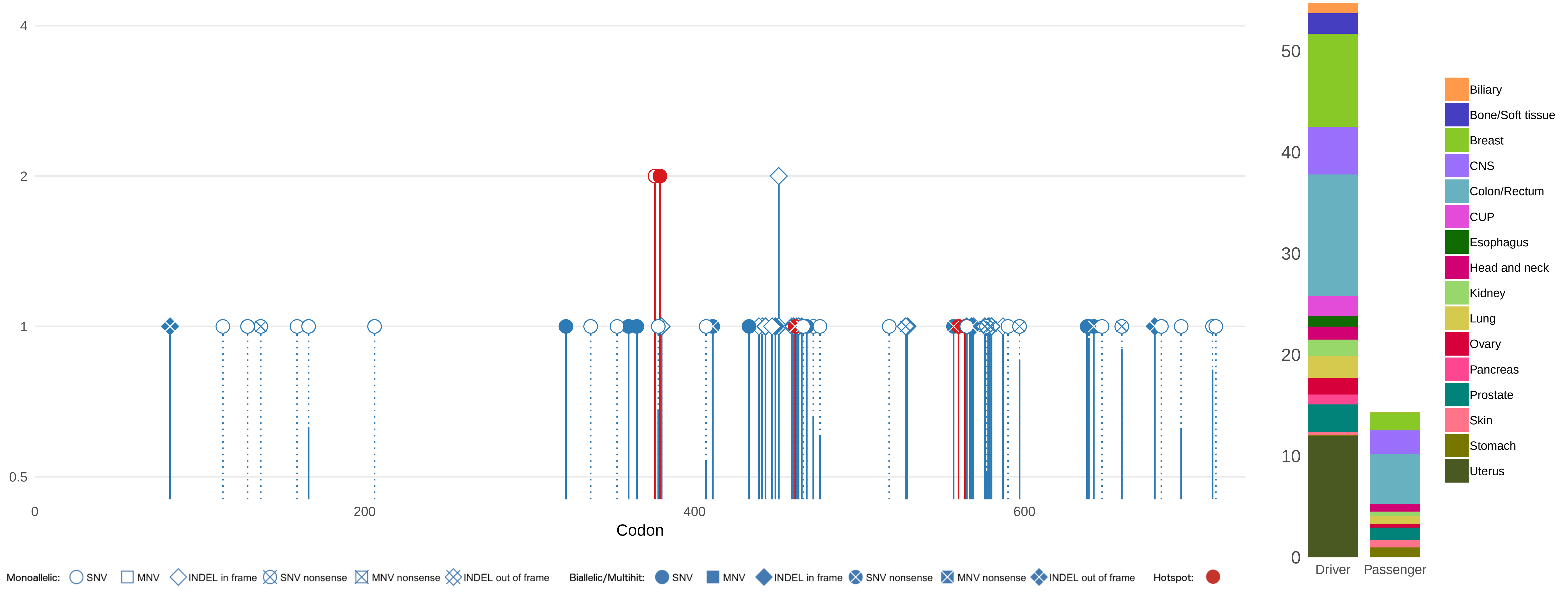

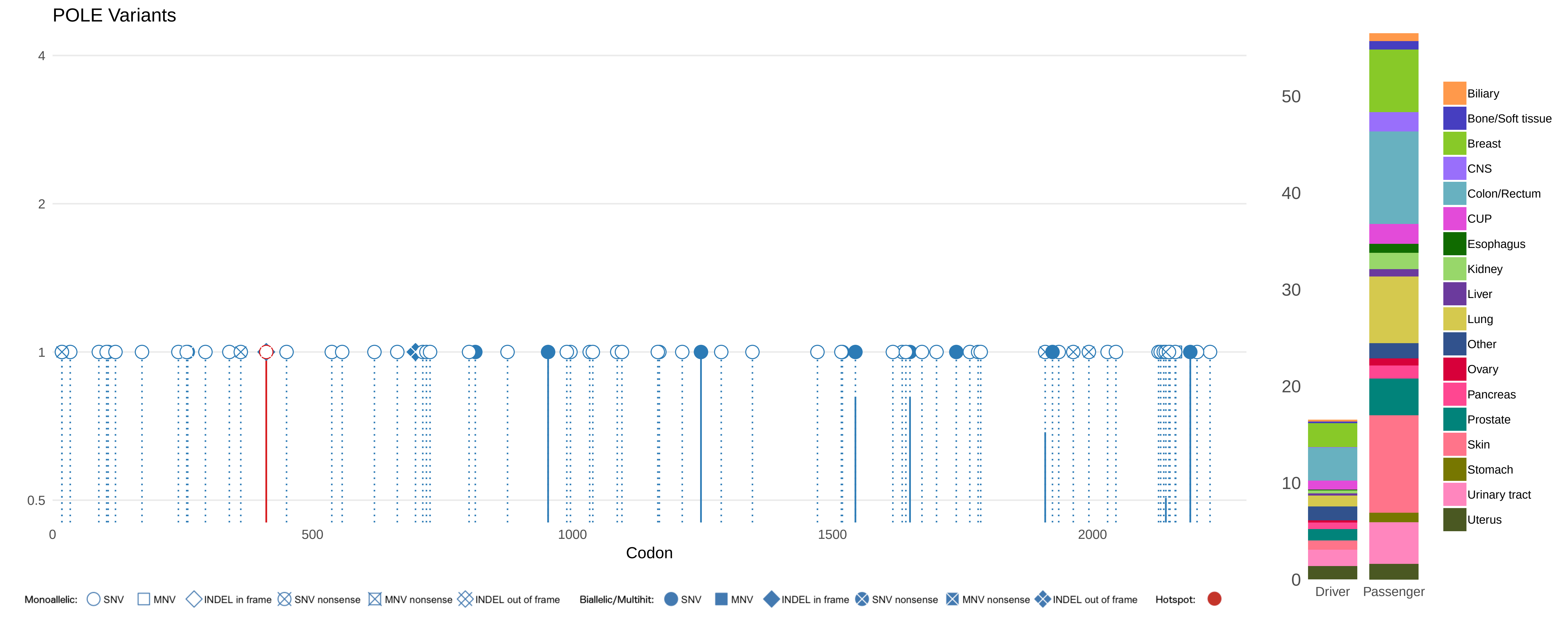

POLE Variants
4
2
1
0.5
0
500
1000
1500
2000
Codon
50
40
30
20
10
0
Driver
Passenger
Biliary
Bone/Soft tissue
Breast
CNS
Colon/Rectum
CUP
Esophagus
Kidney
Liver
Lung
Other
Ovary
Pancreas
Prostate
Skin
Stomach
Urinary tract
Uterus
Monoallelic: SNV MNV INDEL in frame SNV nonsense MNV nonsense INDEL out of frame
Biallelic/Multihit: SNV MNV INDEL in frame SNV nonsense MNV nonsense INDEL out of frame
Hotspot:

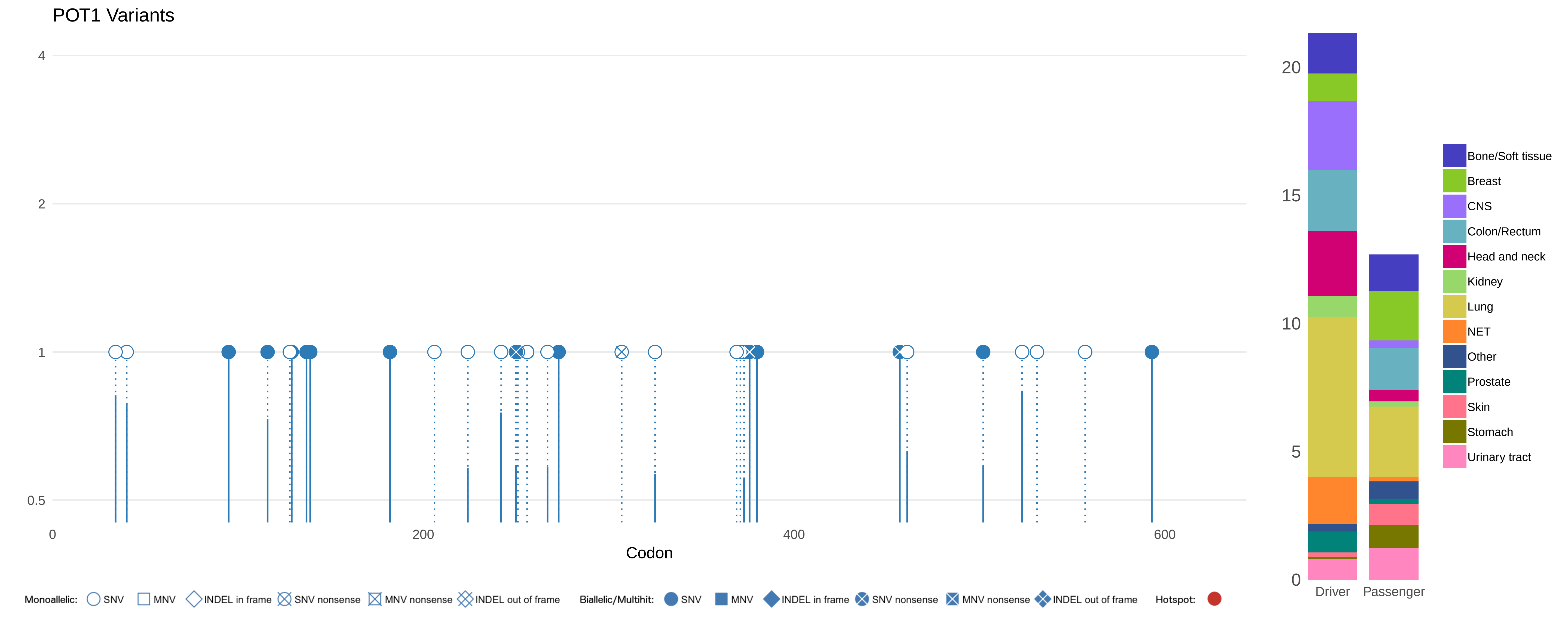

POT1 Variants
4
2
1
0.5
0
200
400
600
Codon
Monoallelic: SNV MNV INDEL in frame SNV nonsense MNV nonsense INDEL out of frame
Biallelic/Multihit: SNV MNV INDEL in frame SNV nonsense MNV nonsense INDEL out of frame
Hotspot:
20
15
10
5
0
Driver
Passenger
Bone/Soft tissue
Breast
CNS
Colon/Rectum
Head and neck
Kidney
Lung
NET
Other
Prostate
Skin
Stomach
Urinary tract

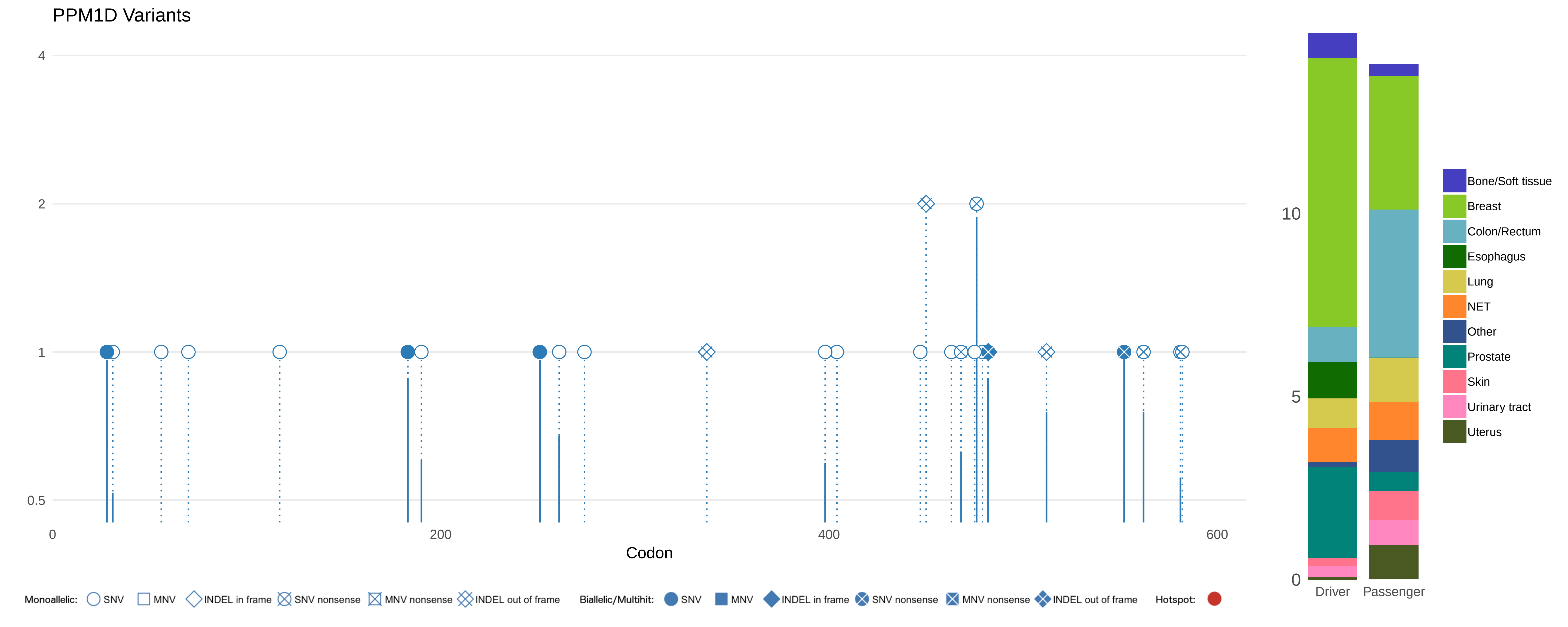

PPM1D Variants
4
2
1
0.5
0
200
400
600
Codon
Monoallelic: SNV MNV INDEL in frame SNV nonsense MNV nonsense INDEL out of frame
Biallelic/Multihit: SNV MNV INDEL in frame SNV nonsense MNV nonsense INDEL out of frame
Hotspot:
10
5
0
Driver
Passenger
Bone/Soft tissue
Breast
Colon/Rectum
Esophagus
Lung
NET
Other
Prostate
Skin
Urinary tract
Uterus

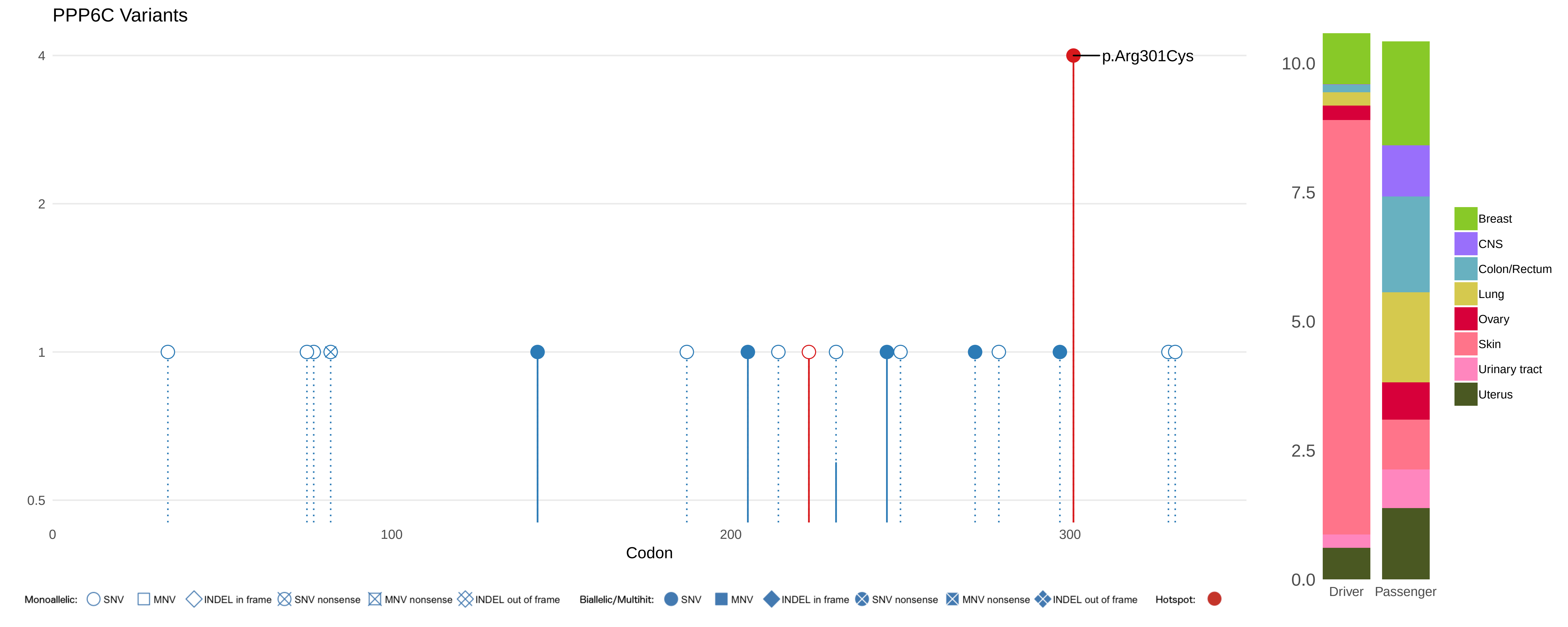
PPP6C Variants
4
2
1
0.5
p.Arg301Cys
0
100
200
300
Codon
Monoallelic: SNV MNV INDEL in frame SNV nonsense MNV nonsense INDEL out of frame
Biallelic/Multihit: SNV MNV INDEL in frame SNV nonsense MNV nonsense INDEL out of frame
Hotspot:
10.0
7.5
5.0
2.5
0.0
Driver
Passenger
Breast
CNS
Colon/Rectum
Lung
Ovary
Skin
Urinary tract
Uterus

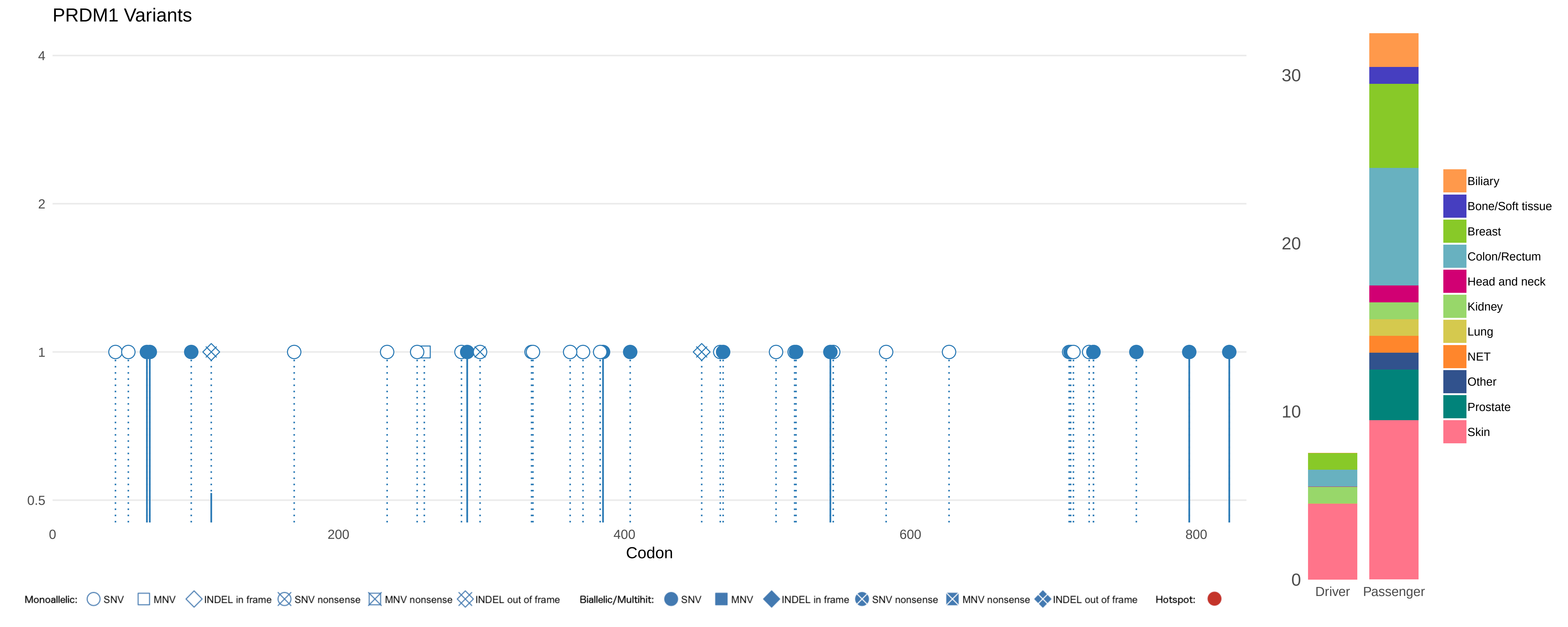
PRDM1 Variants
4
2
1
0.5
0
200
400
600
800
Codon
30
20
10
0
Driver
Passenger
Biliary
Bone/Soft tissue
Breast
Colon/Rectum
Head and neck
Kidney
Lung
NET
Other
Prostate
Skin
Monoallelic: SNV MNV INDEL in frame SNV nonsense MNV nonsense INDEL out of frame
Biallelic/Multihit: SNV MNV INDEL in frame SNV nonsense MNV nonsense INDEL out of frame
Hotspot:

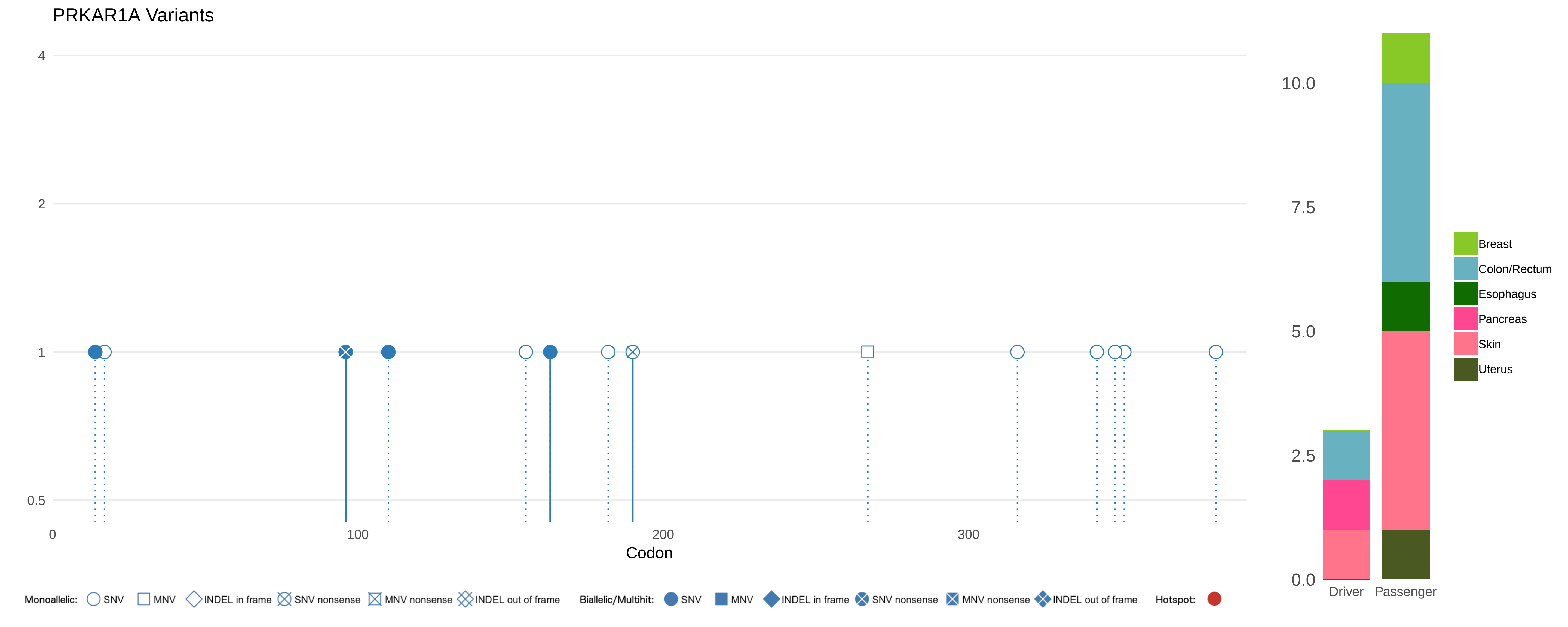
PRKAR1A Variants
4
2
1
0.5
0
100
200
300
Codon
10.0
7.5
5.0
2.5
0.0
Driver
Passenger
Breast
Colon/Rectum
Esophagus
Pancreas
Skin
Uterus
Monoallelic: SNV MNV INDEL in frame SNV nonsense MNV nonsense INDEL out of frame
Biallelic/Multihit: SNV MNV INDEL in frame SNV nonsense MNV nonsense INDEL out of frame
Hotspot:

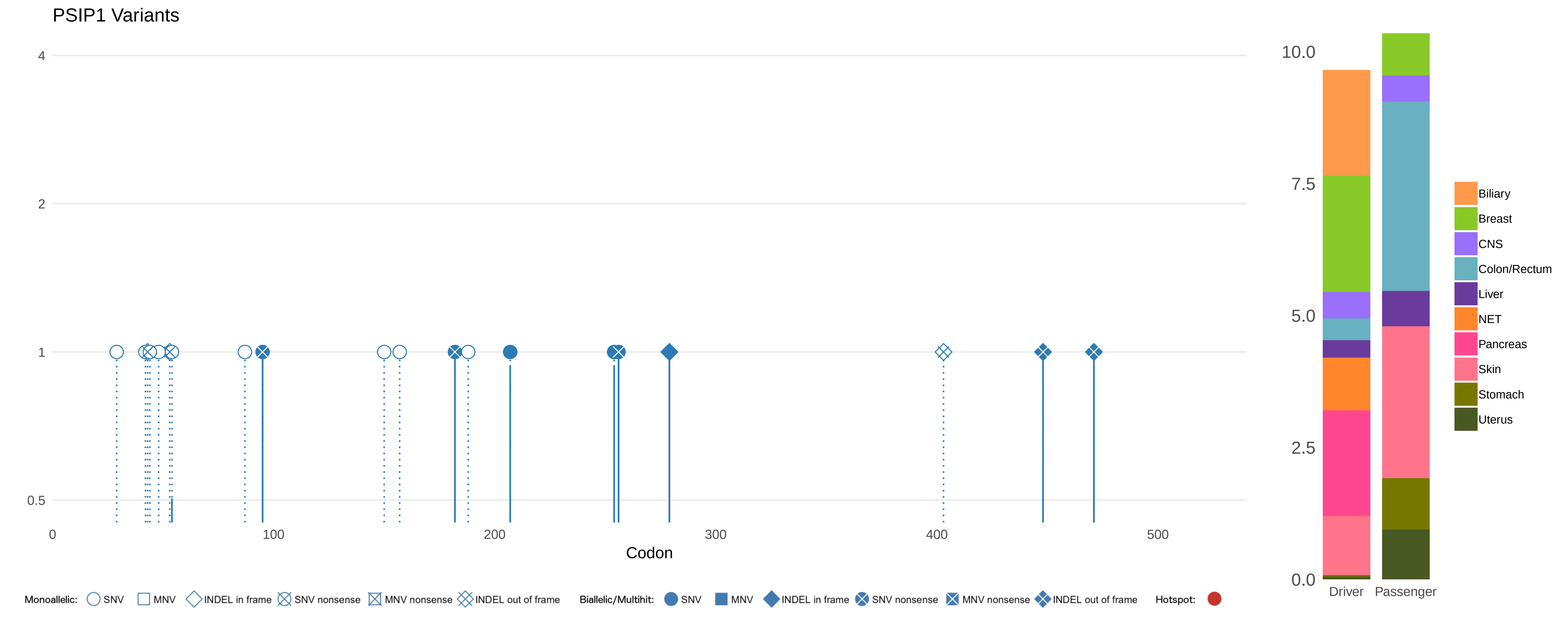
PSIP1 Variants
4
2
1
0.5
0
100
200
300
400
500
Codon
10.0
7.5
5.0
2.5
0.0
Driver
Passenger
Biliary
Breast
CNS
Colon/Rectum
Liver
NET
Pancreas
Skin
Stomach
Uterus
Monoallelic: SNV MNV INDEL in frame SNV nonsense MNV nonsense INDEL out of frame
Biallelic/Multihit: SNV MNV INDEL in frame SNV nonsense MNV nonsense INDEL out of frame
Hotspot:

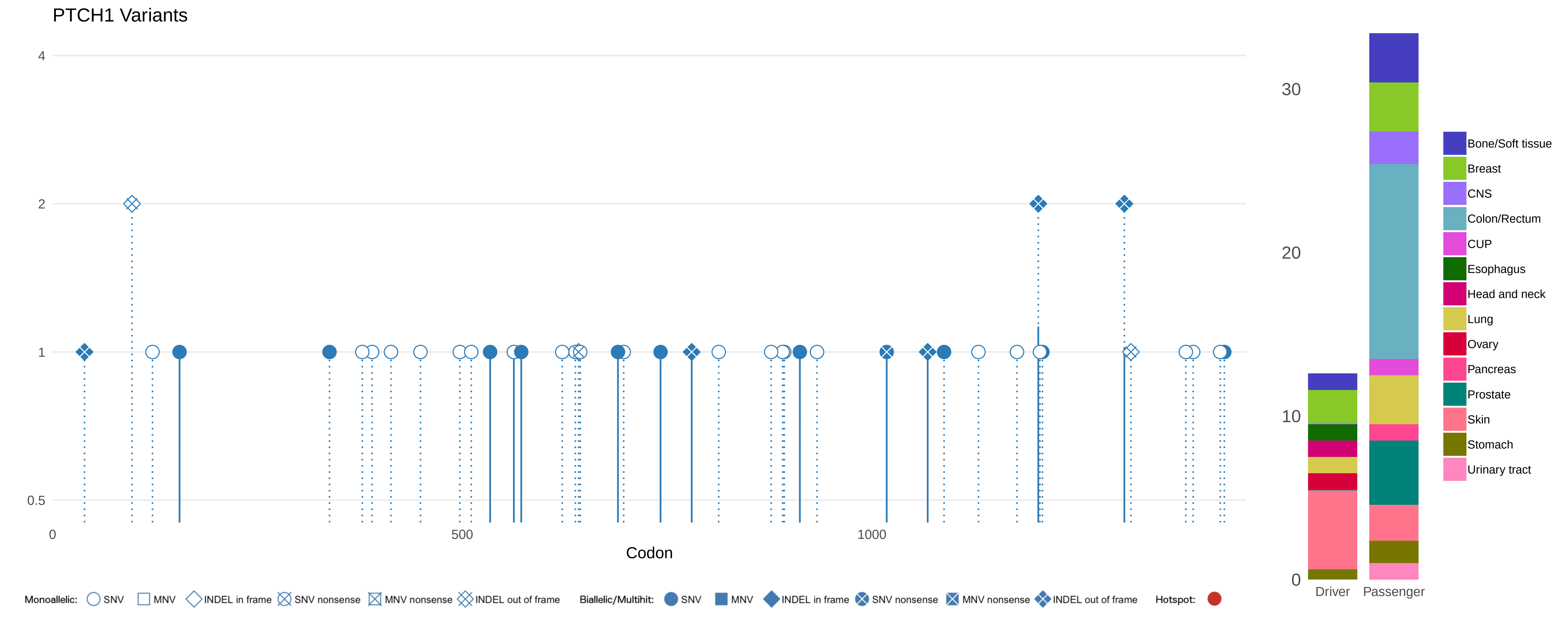

PTCH1 Variants
4
2
1
0.5
0
500
1000
Codon
30
20
10
0
Driver
Passenger
Bone/Soft tissue
Breast
CNS
Colon/Rectum
CUP
Esophagus
Head and neck
Lung
Ovary
Pancreas
Prostate
Skin
Stomach
Urinary tract
Monoallelic: SNV MNV INDEL in frame SNV nonsense MNV nonsense INDEL out of frame
Biallelic/Multihit: SNV MNV INDEL in frame SNV nonsense MNV nonsense INDEL out of frame
Hotspot:

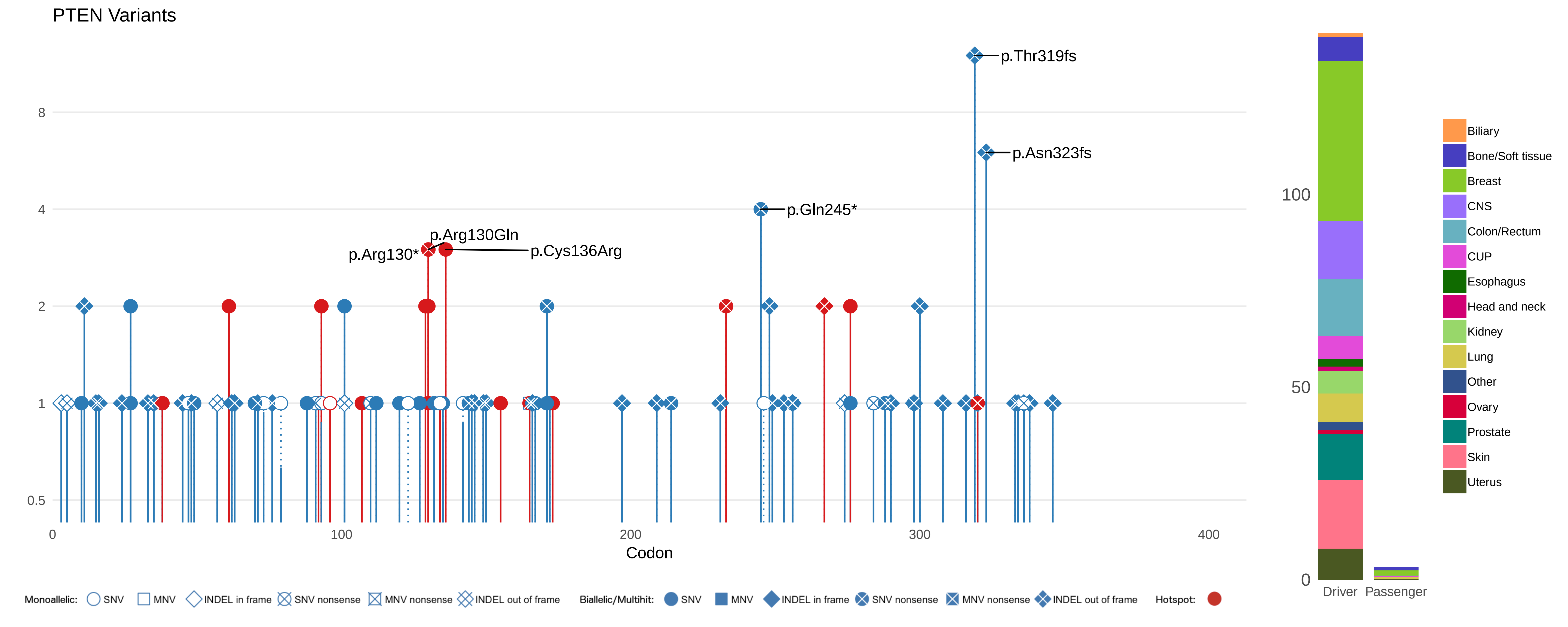

PTEN Variants
p.Thr319fs
p.Asn323fs
p.Gln245*
p.Arg130Gln
p.Arg130*
p.Cys136Arg
8
4
2
1
0.5
0
100
200
300
400
Codon
100
50
0
Driver
Passenger
Biliary
Bone/Soft tissue
Breast
CNS
Colon/Rectum
CUP
Esophagus
Head and neck
Kidney
Lung
Other
Ovary
Prostate
Skin
Uterus
Monoallelic: SNV MNV INDEL in frame SNV nonsense MNV nonsense INDEL out of frame
Biallelic/Multihit: SNV MNV INDEL in frame SNV nonsense MNV nonsense INDEL out of frame
Hotspot:

PTK6 Variants

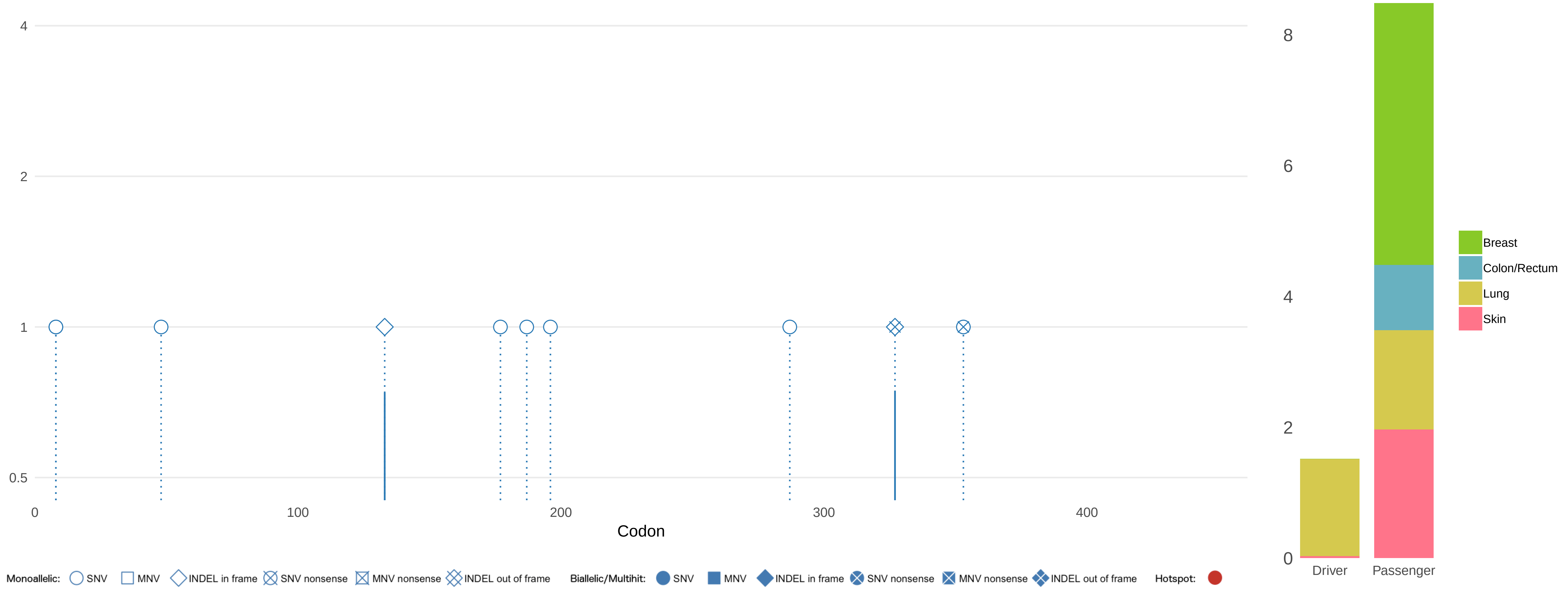

### PTPN13 Variants

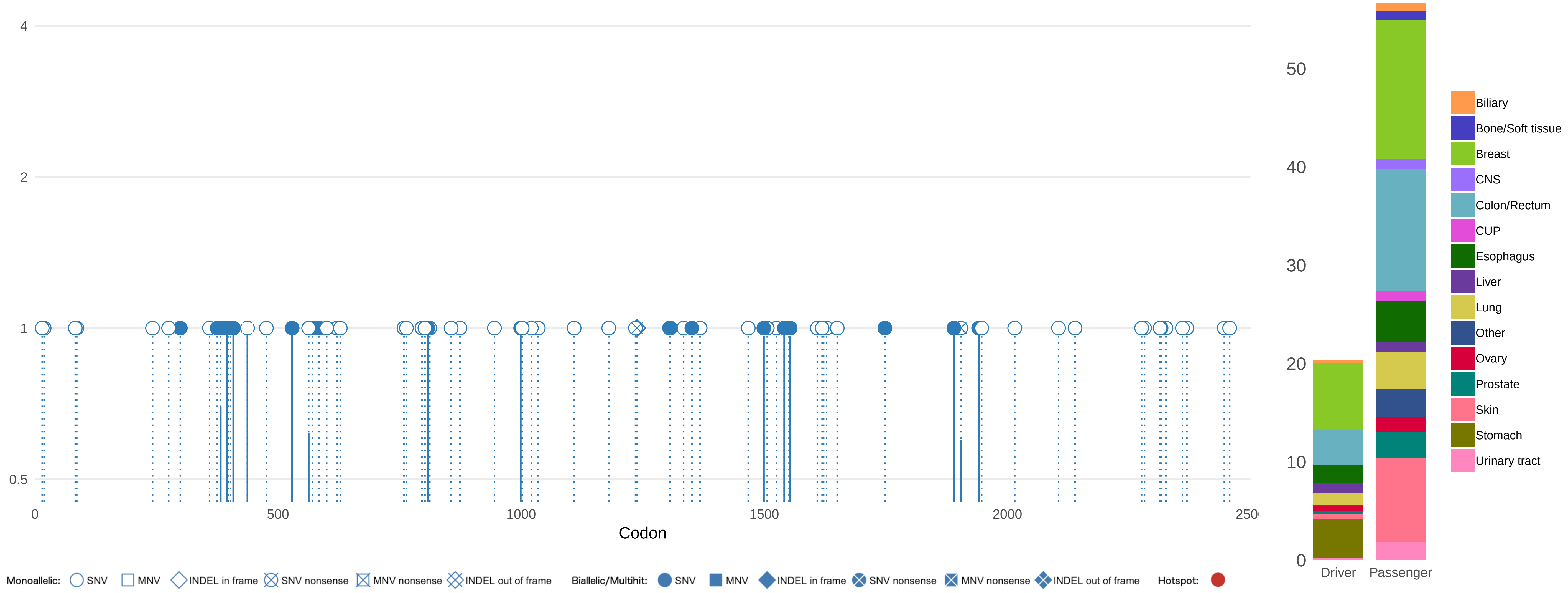

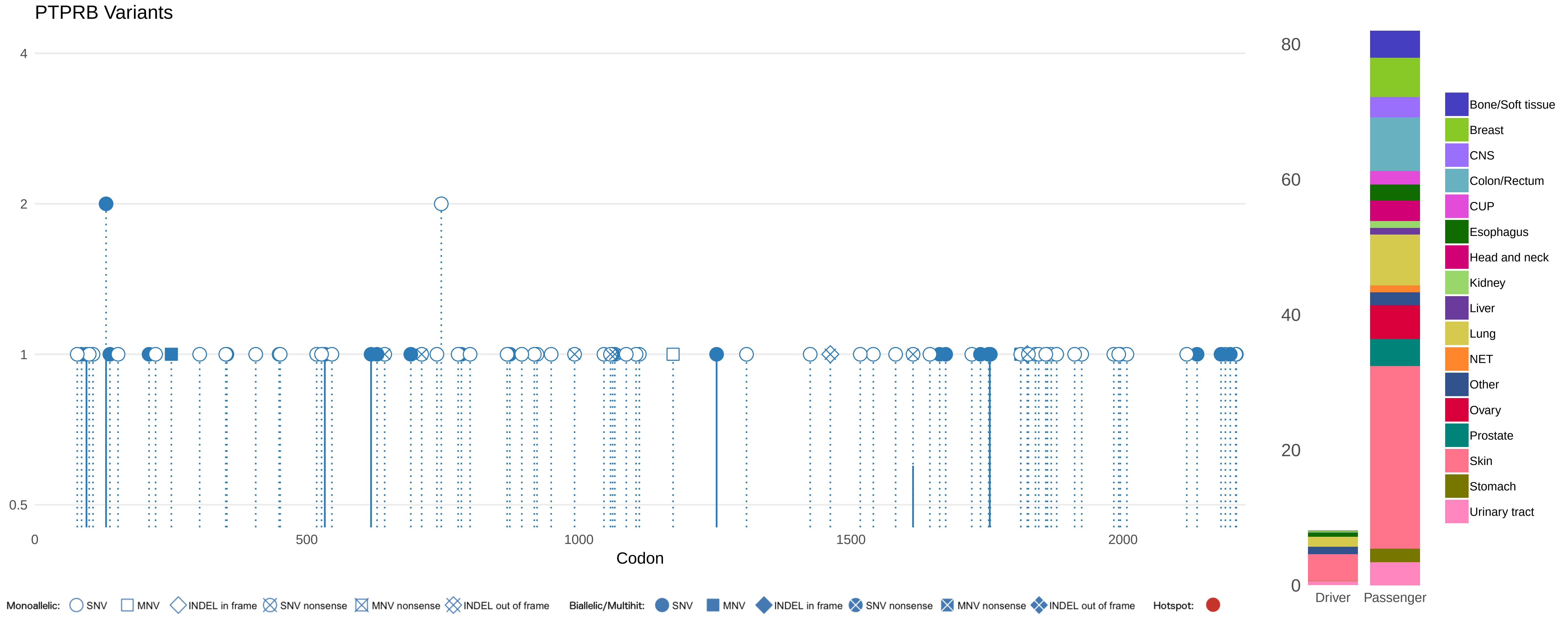

PTPRB Variants
4
2
1
0.5
0
500
1000
1500
2000
Codon
80
60
40
20
0
Driver
Passenger
Bone/Soft tissue
Breast
CNS
Colon/Rectum
CUP
Esophagus
Head and neck
Kidney
Liver
Lung
NET
Other
Ovary
Prostate
Skin
Stomach
Urinary tract
Monoallelic:
SNV
MNV
INDEL in frame
SNV nonsense
MNV nonsense
INDEL out of frame
Biallelic/Multihit:
SNV
MNV
INDEL in frame
SNV nonsense
MNV nonsense
INDEL out of frame
Hotspot:

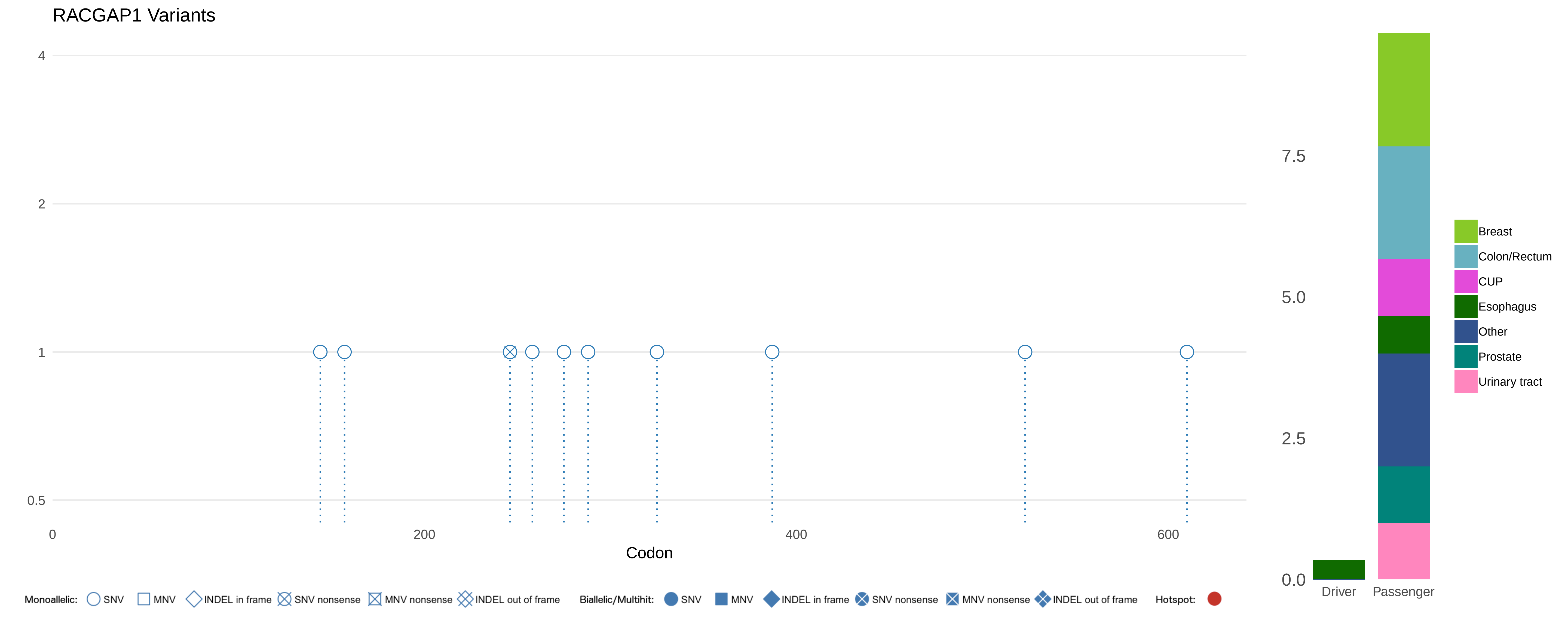

RACGAP1 Variants
4
2
1
0.5
0
200
400
600
Codon
Monoallelic: SNV MNV INDEL in frame SNV nonsense MNV nonsense INDEL out of frame
Biallelic/Multihit: SNV MNV INDEL in frame SNV nonsense MNV nonsense INDEL out of frame
Hotspot:
7.5
5.0
2.5
0.0
Driver
Passenger
Breast
Colon/Rectum
CUP
Esophagus
Other
Prostate
Urinary tract

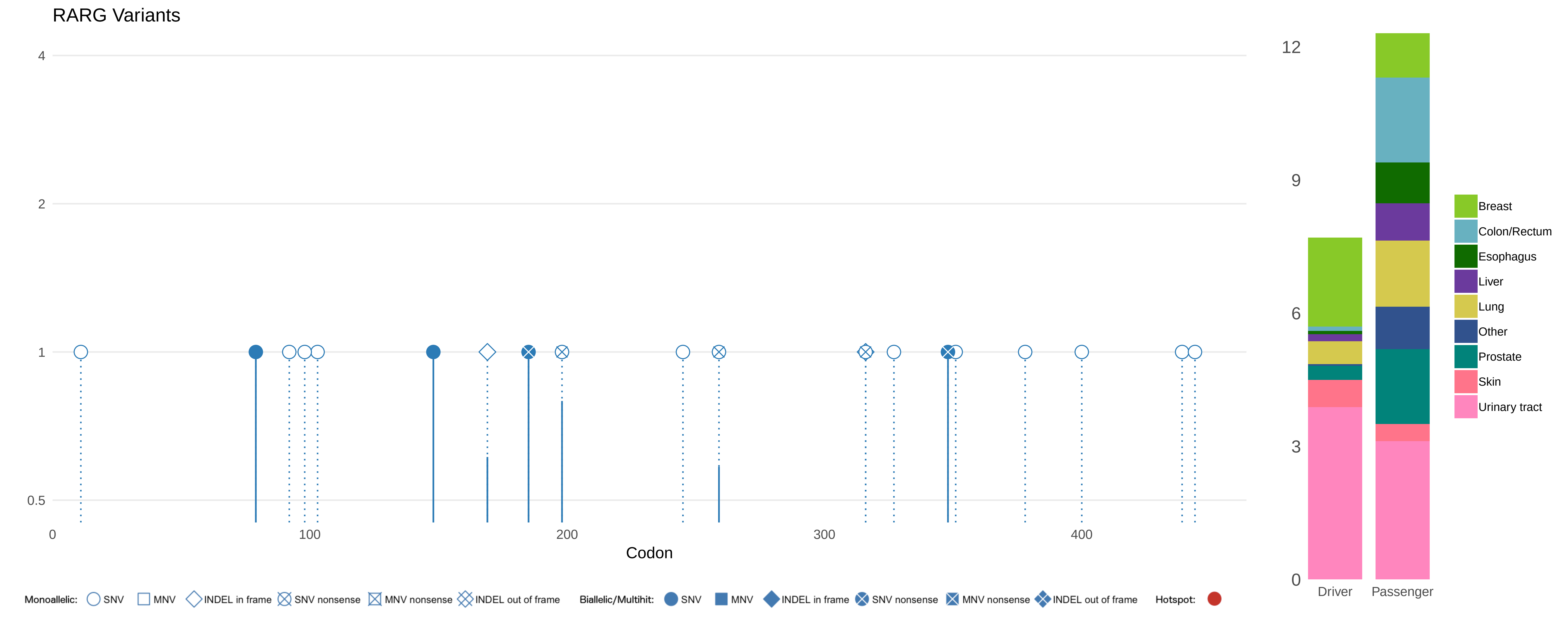

RARG Variants
4
2
1
0.5
0
100
200
300
400
Codon
12
9
6
3
0
Driver
Passenger
Breast
Colon/Rectum
Esophagus
Liver
Lung
Other
Prostate
Skin
Urinary tract
Monoallelic: SNV MNV INDEL in frame SNV nonsense MNV nonsense INDEL out of frame
Biallelic/Multihit: SNV MNV INDEL in frame SNV nonsense MNV nonsense INDEL out of frame
Hotspot:

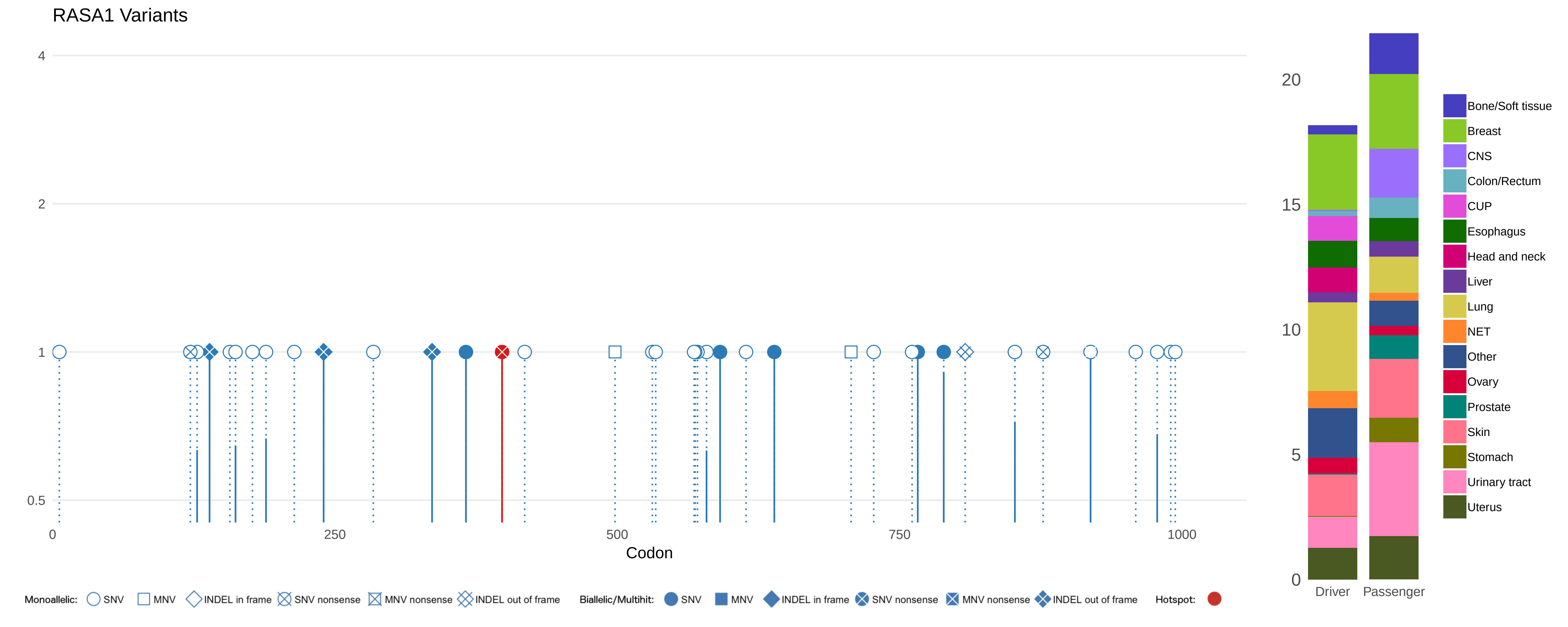
RASA1 Variants
4
2
1
0.5
0
250
500
750
1000
Codon
Monoallelic: SNV MNV INDEL in frame SNV nonsense MNV nonsense INDEL out of frame
Biallelic/Multihit: SNV MNV INDEL in frame SNV nonsense MNV nonsense INDEL out of frame
Hotspot:
20
15
10
5
0
Driver
Passenger
Bone/Soft tissue
Breast
CNS
Colon/Rectum
CUP
Esophagus
Head and neck
Liver
Lung
NET
Other
Ovary
Prostate
Skin
Stomach
Urinary tract
Uterus

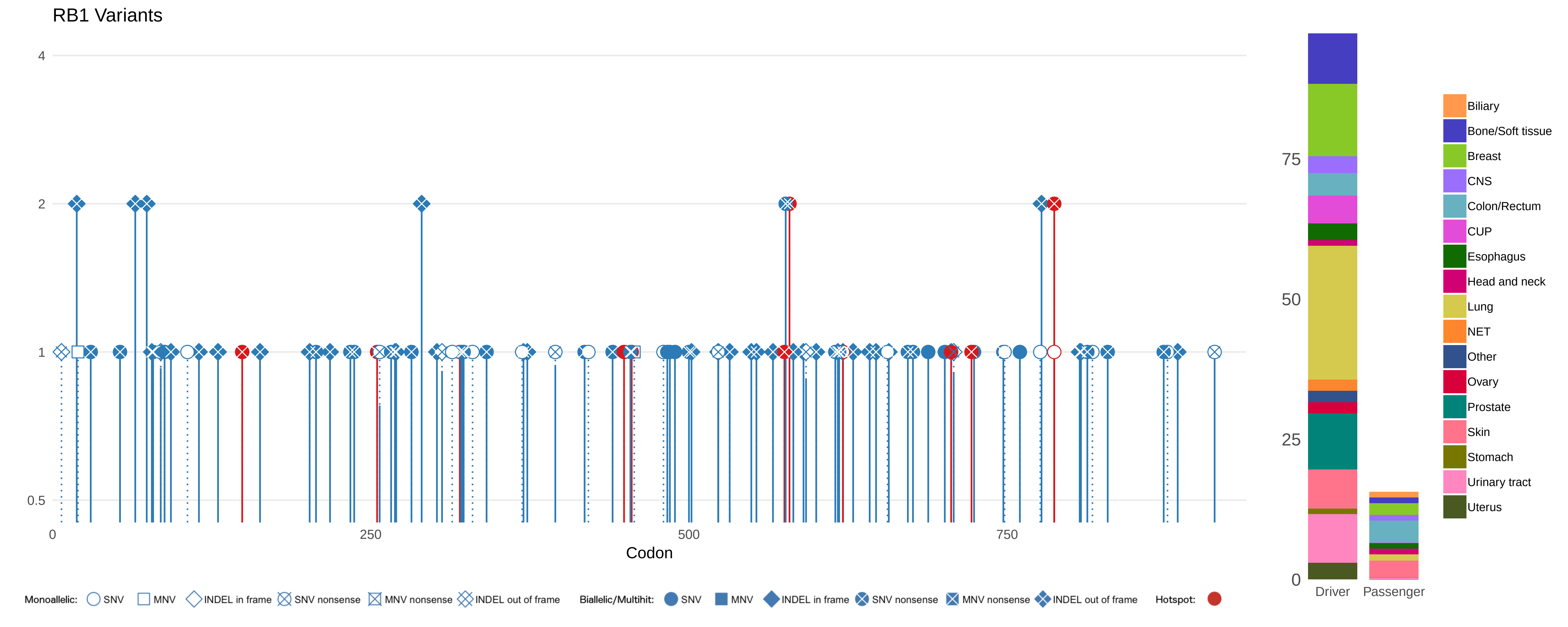
RB1 Variants
4
2
1
0.5
0
250
500
750
Codon
75
50
25
0
Driver
Passenger
Biliary
Bone/Soft tissue
Breast
CNS
Colon/Rectum
CUP
Esophagus
Head and neck
Lung
NET
Other
Ovary
Prostate
Skin
Stomach
Urinary tract
Uterus
Monoallelic: SNV MNV INDEL in frame SNV nonsense MNV nonsense INDEL out of frame
Biallelic/Multihit: SNV MNV INDEL in frame SNV nonsense MNV nonsense INDEL out of frame
Hotspot:

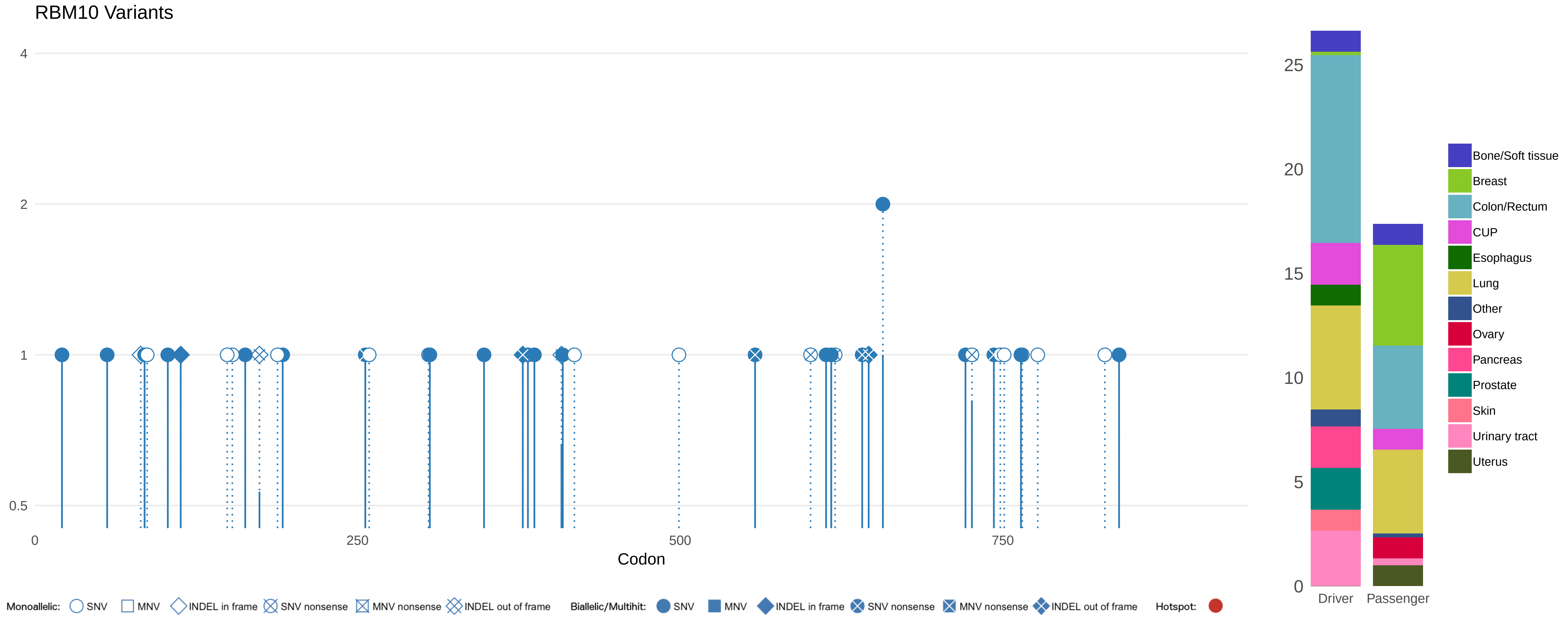

RBM10 Variants
4
2
1
0.5
0
250
500
750
Codon
25
20
15
10
5
0
Driver
Passenger
Bone/Soft tissue
Breast
Colon/Rectum
CUP
Esophagus
Lung
Other
Ovary
Pancreas
Prostate
Skin
Urinary tract
Uterus
Monoallelic: SNV MNV INDEL in frame SNV nonsense MNV nonsense INDEL out of frame
Biallelic/Multihit: SNV MNV INDEL in frame SNV nonsense MNV nonsense INDEL out of frame
Hotspot:

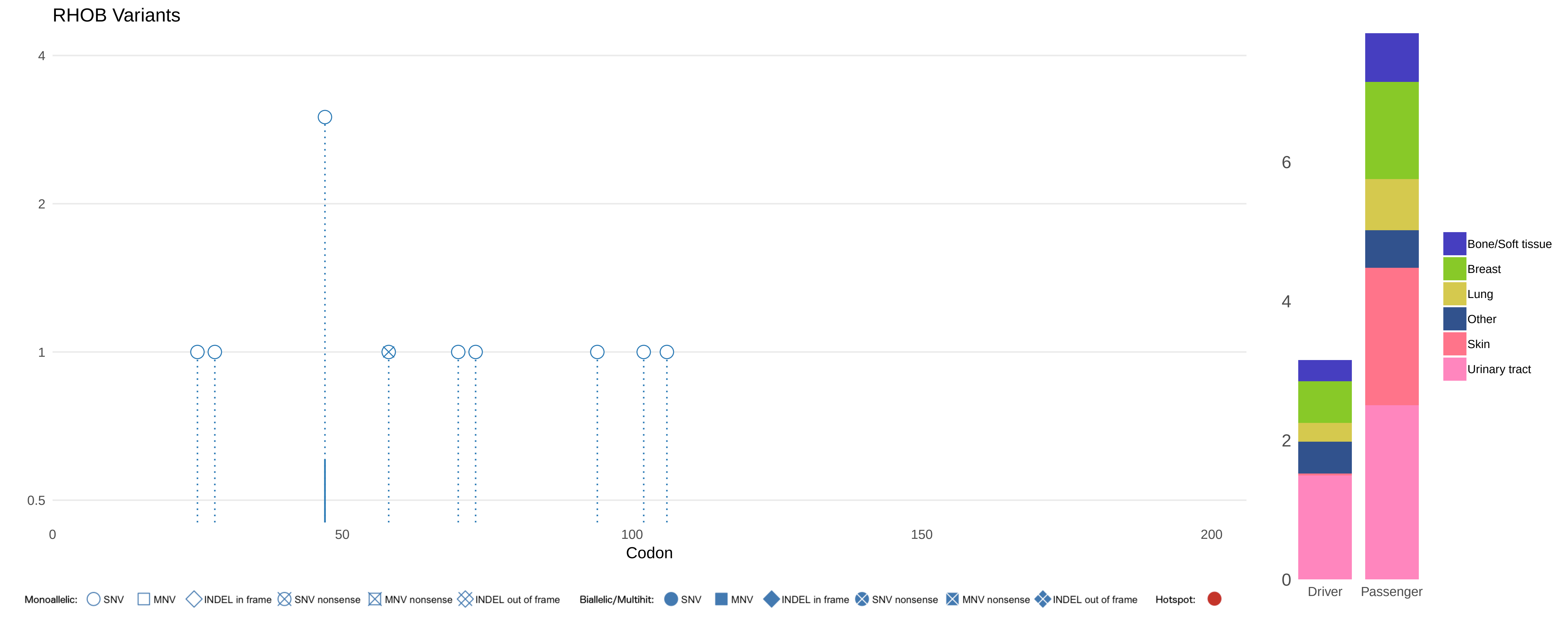
RHOB Variants
4
2
1
0.5
0
50
100
150
200
Codon
Monoallelic: SNV MNV INDEL in frame SNV nonsense MNV nonsense INDEL out of frame
Biallelic/Multihit: SNV MNV INDEL in frame SNV nonsense MNV nonsense INDEL out of frame
Hotspot:
6
4
2
0
Driver
Passenger
Bone/Soft tissue
Breast
Lung
Other
Skin
Urinary tract

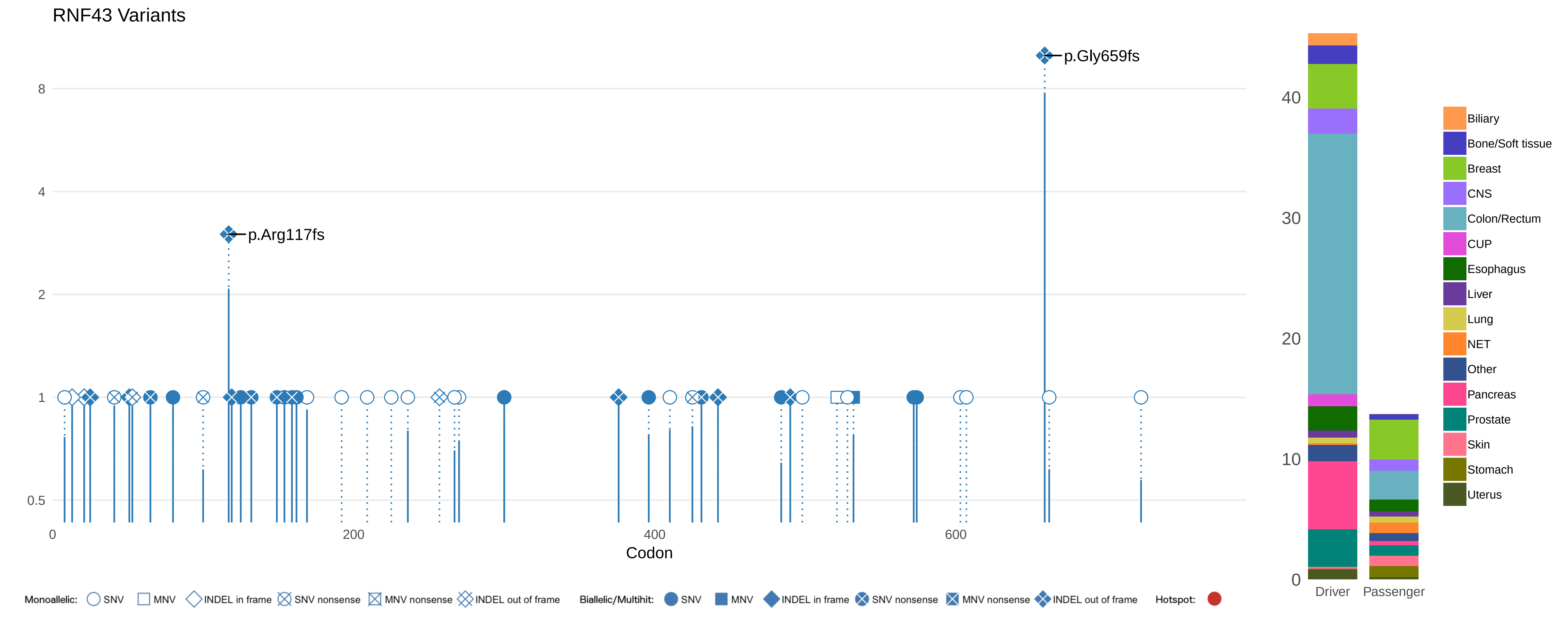

RNF43 Variants
p.Gly659fs
p.Arg117fs
8
4
2
1
0.5
0
200
400
600
Codon
Monoallelic: SNV MNV INDEL in frame SNV nonsense MNV nonsense INDEL out of frame
Biallelic/Multihit: SNV MNV INDEL in frame SNV nonsense MNV nonsense INDEL out of frame
Hotspot:
40
30
20
10
0
Driver
Passenger
Biliary
Bone/Soft tissue
Breast
CNS
Colon/Rectum
CUP
Esophagus
Liver
Lung
NET
Other
Pancreas
Prostate
Skin
Stomach
Uterus

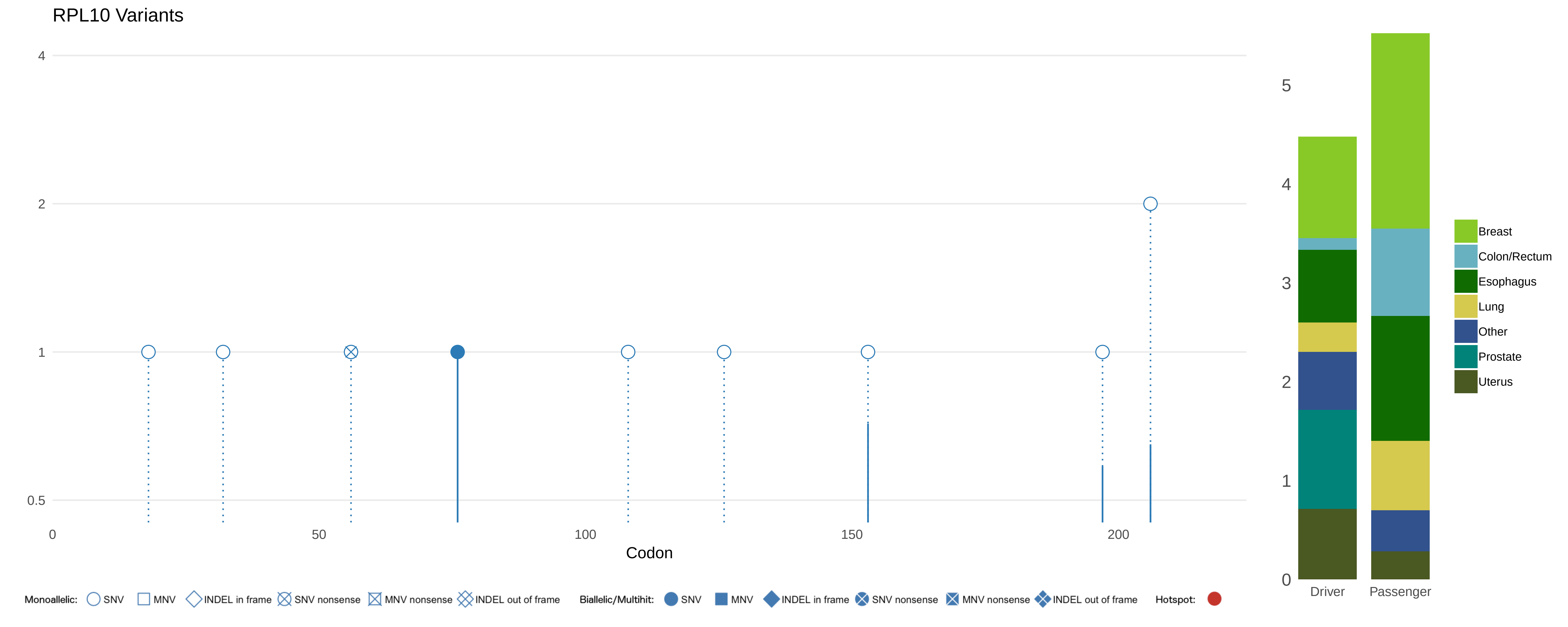

RPL10 Variants
4
2
1
0.5
0
50
100
150
200
Codon
5
4
3
2
1
0
Driver
Passenger
Breast
Colon/Rectum
Esophagus
Lung
Other
Prostate
Uterus
Monoallelic: SNV MNV INDEL in frame SNV nonsense MNV nonsense INDEL out of frame
Biallelic/Multihit: SNV MNV INDEL in frame SNV nonsense MNV nonsense INDEL out of frame
Hotspot:

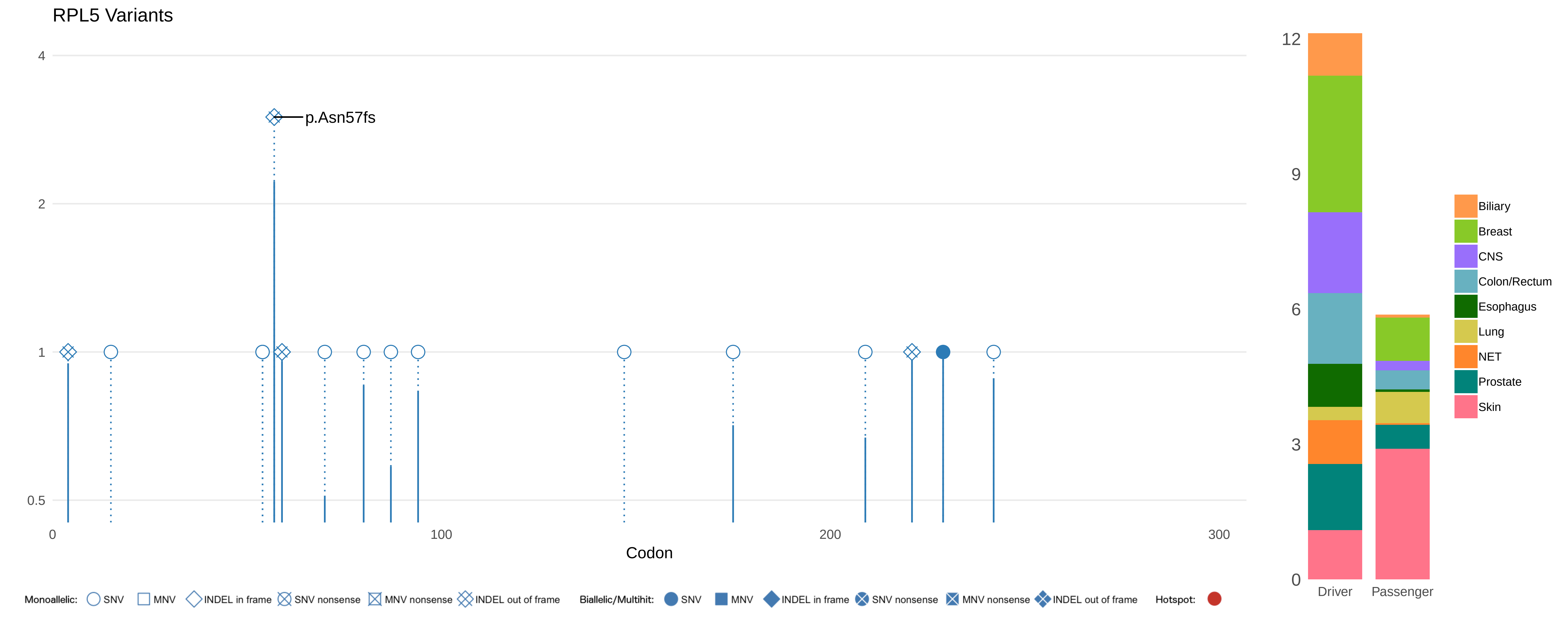

RPL5 Variants
4
2
1
0.5
p.Asn57fs
0
100
200
300
Codon
12
9
6
3
0
Driver
Passenger
Biliary
Breast
CNS
Colon/Rectum
Esophagus
Lung
NET
Prostate
Skin
Monoallelic: SNV MNV INDEL in frame SNV nonsense MNV nonsense INDEL out of frame
Biallelic/Multihit: SNV MNV INDEL in frame SNV nonsense MNV nonsense INDEL out of frame
Hotspot:

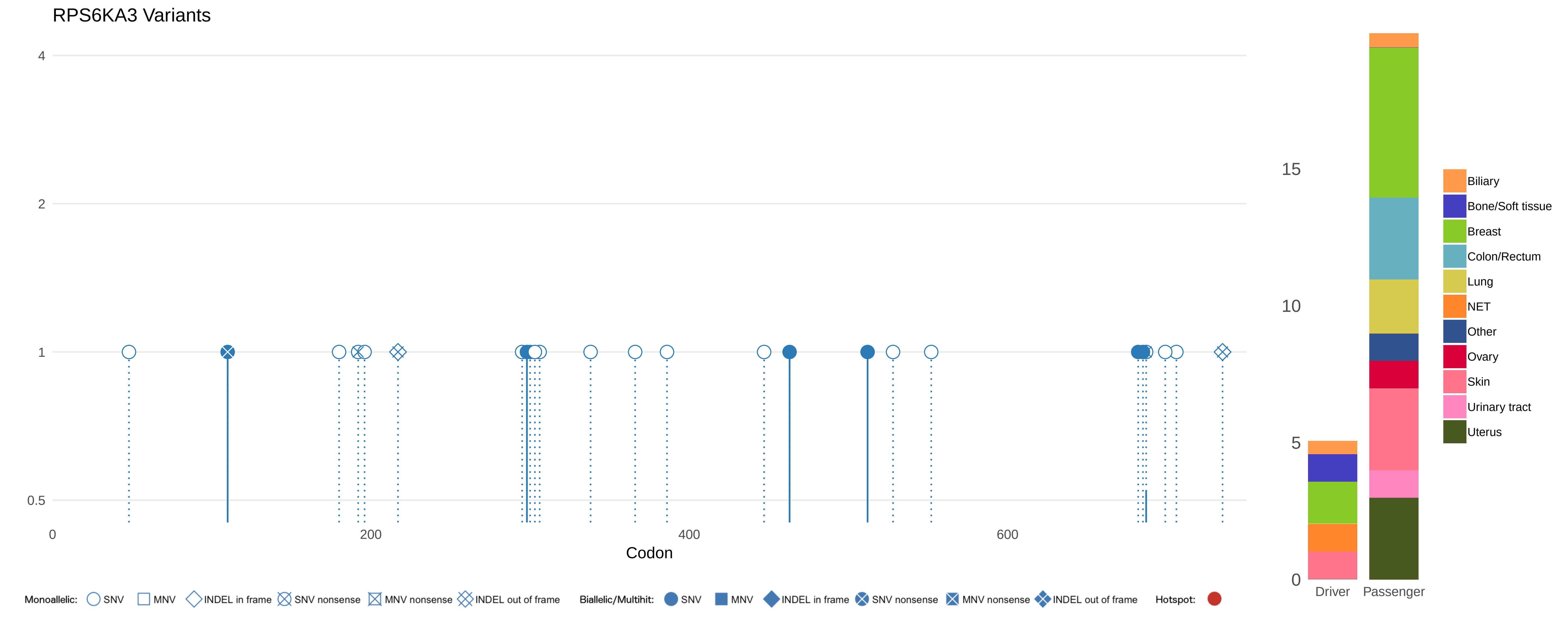
RPS6KA3 Variants
4
2
1
0.5
0
200
400
600
Codon
15
10
5
0
Driver
Passenger
Biliary
Bone/Soft tissue
Breast
Colon/Rectum
Lung
NET
Other
Ovary
Skin
Urinary tract
Uterus
Monoallelic: SNV MNV INDEL in frame SNV nonsense MNV nonsense INDEL out of frame
Biallelic/Multihit: SNV MNV INDEL in frame SNV nonsense MNV nonsense INDEL out of frame
Hotspot:

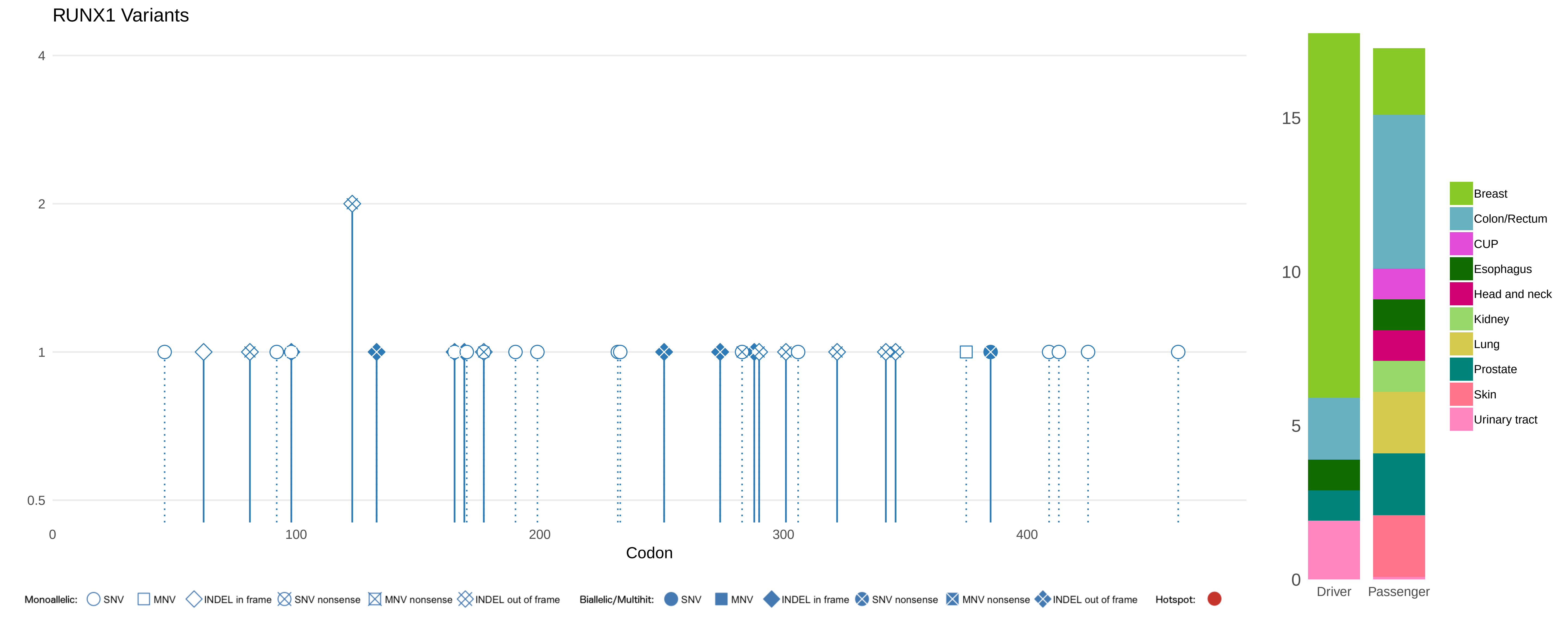
RUNX1 Variants
4
2
1
0.5
0
100
200
300
400
Codon
15
10
5
0
Driver
Passenger
Breast
Colon/Rectum
CUP
Esophagus
Head and neck
Kidney
Lung
Prostate
Skin
Urinary tract
Monoallelic:
SNV
MNV
INDEL in frame
SNV nonsense
MNV nonsense
INDEL out of frame
Biallelic/Multihit:
SNV
MNV
INDEL in frame
SNV nonsense
MNV nonsense
INDEL out of frame
Hotspot:

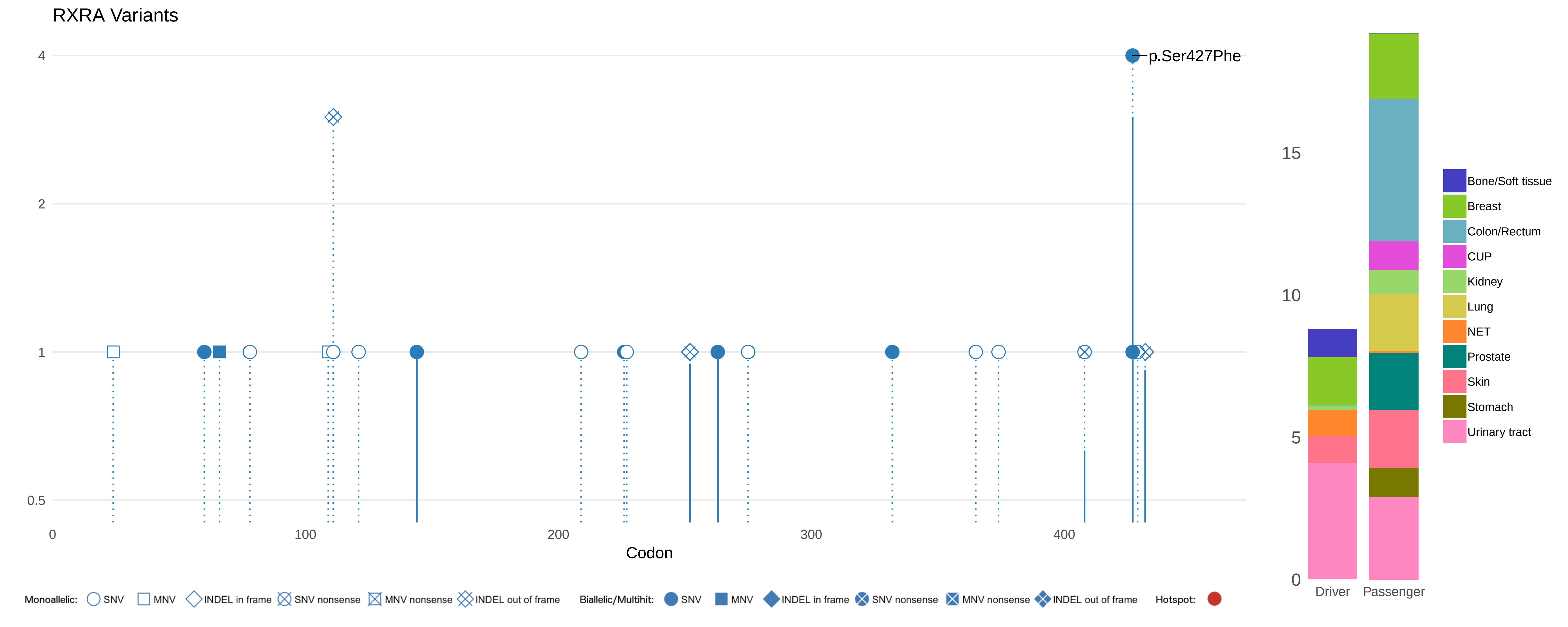

RXRA Variants
p.Ser427Phe
4
2
1
0.5
0
100
200
300
400
Codon
15
10
5
0
Driver
Passenger
Bone/Soft tissue
Breast
Colon/Rectum
CUP
Kidney
Lung
NET
Prostate
Skin
Stomach
Urinary tract
Monoallelic: SNV MNV INDEL in frame SNV nonsense MNV nonsense INDEL out of frame
Biallelic/Multihit: SNV MNV INDEL in frame SNV nonsense MNV nonsense INDEL out of frame
Hotspot:

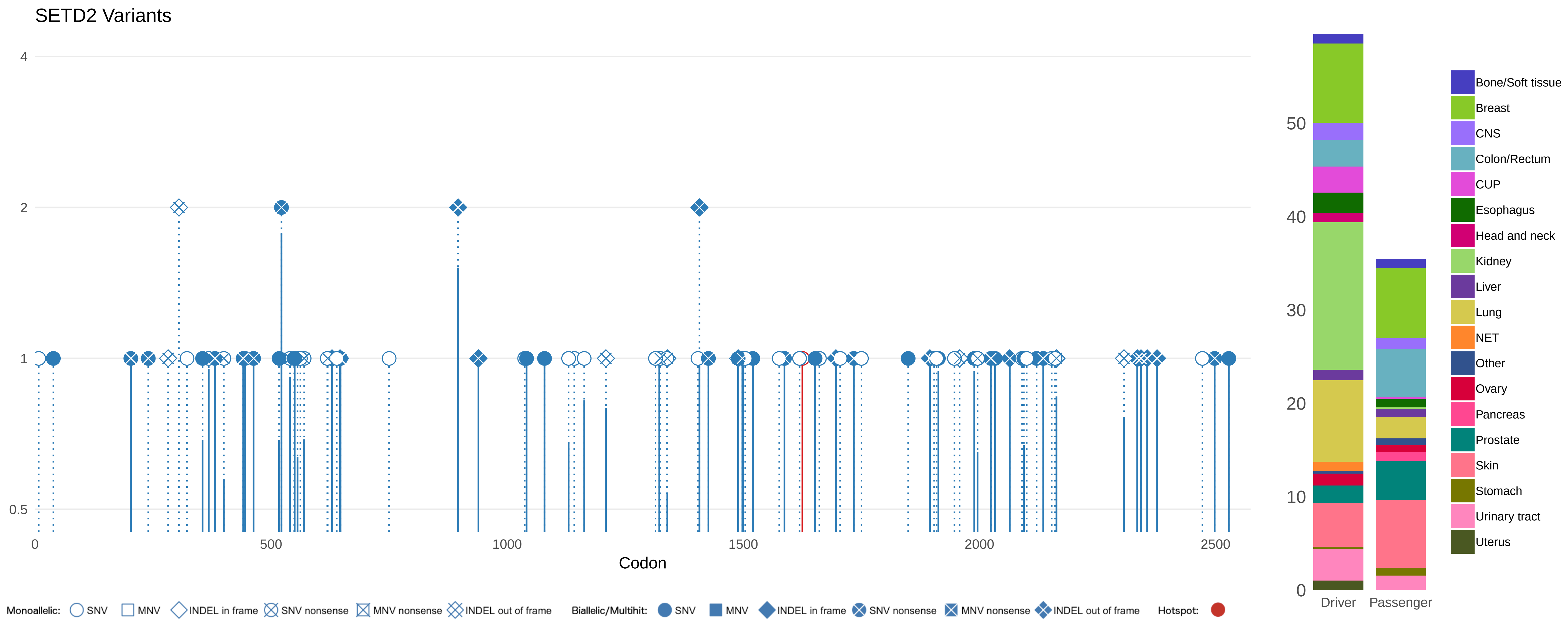
SETD2 Variants
4
2
1
0.5
0
500
1000
1500
2000
2500
Codon
Monoallelic: SNV MNV INDEL in frame SNV nonsense MNV nonsense INDEL out of frame
Biallelic/Multihit: SNV MNV INDEL in frame SNV nonsense MNV nonsense INDEL out of frame
Hotspot:
50
40
30
20
10
0
Driver
Passenger
Bone/Soft tissue
Breast
CNS
Colon/Rectum
CUP
Esophagus
Head and neck
Kidney
Liver
Lung
NET
Other
Ovary
Pancreas
Prostate
Skin
Stomach
Urinary tract
Uterus

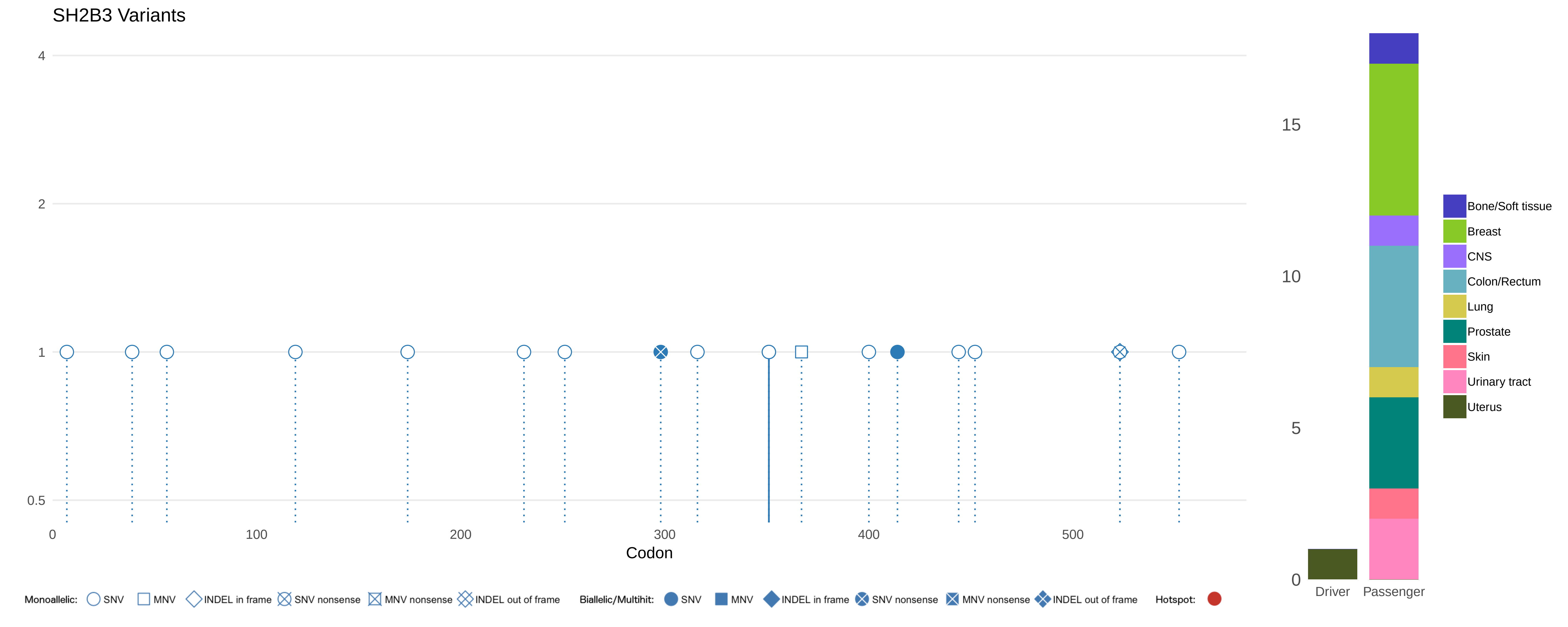

SH2B3 Variants
4
2
1
0.5
0
100
200
300
400
500
Codon
15
10
5
0
Driver
Passenger
Bone/Soft tissue
Breast
CNS
Colon/Rectum
Lung
Prostate
Skin
Urinary tract
Uterus
Monoallelic: SNV MNV INDEL in frame SNV nonsense MNV nonsense INDEL out of frame
Biallelic/Multihit: SNV MNV INDEL in frame SNV nonsense MNV nonsense INDEL out of frame
Hotspot:

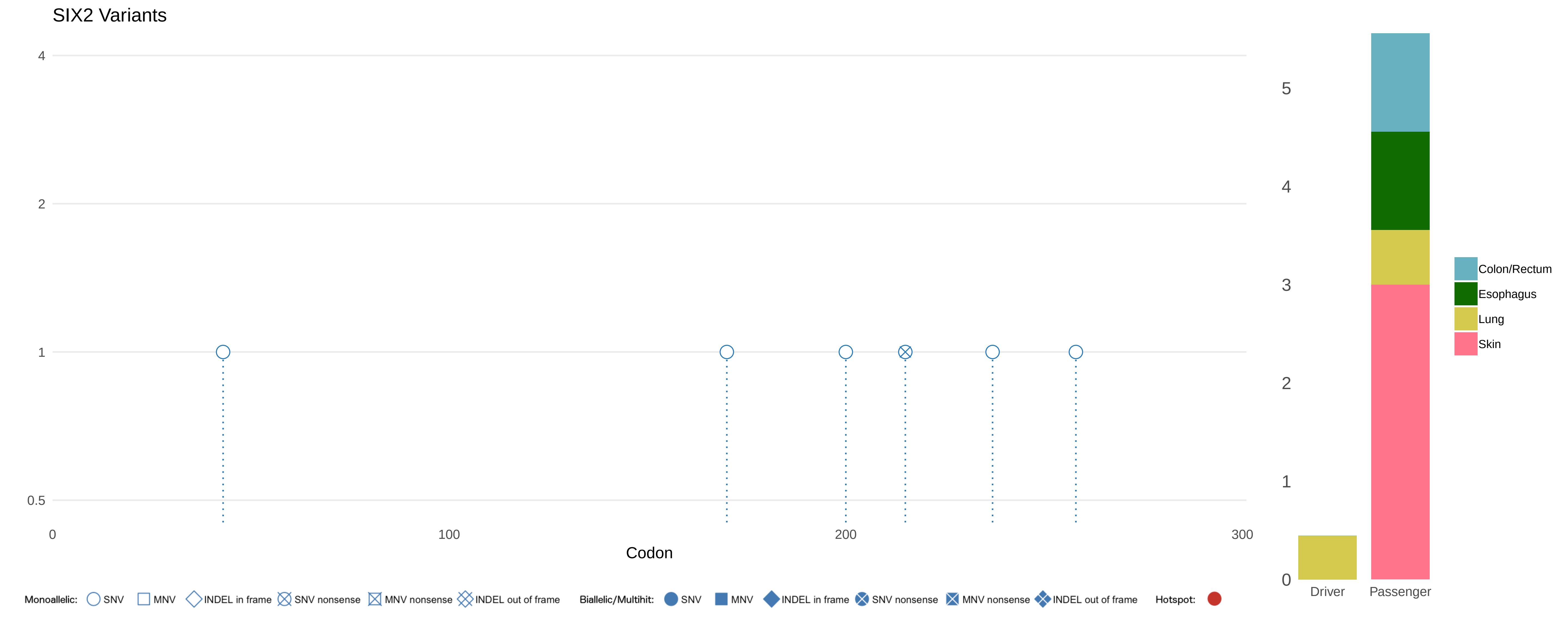
SIX2 Variants
4
2
1
0.5
0
100
200
300
Codon
5
4
3
2
1
0
Driver
Passenger
Colon/Rectum
Esophagus
Lung
Skin
Monoallelic: SNV MNV INDEL in frame SNV nonsense MNV nonsense INDEL out of frame
Biallelic/Multihit: SNV MNV INDEL in frame SNV nonsense MNV nonsense INDEL out of frame
Hotspot:

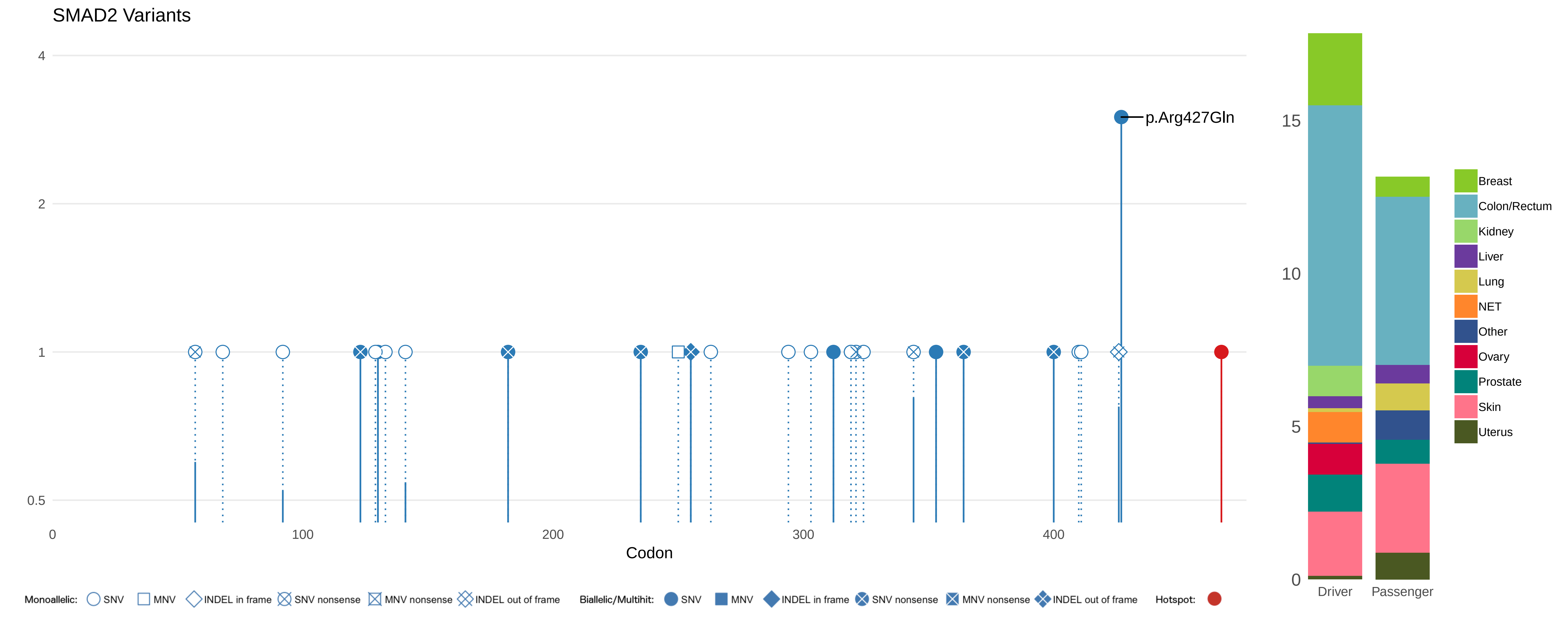
SMAD2 Variants
4
2
1
0.5
0
100
200
300
400
Codon
p.Arg427Gln
15
10
5
0
Driver
Passenger
Breast
Colon/Rectum
Kidney
Liver
Lung
NET
Other
Ovary
Prostate
Skin
Uterus
Monoallelic: SNV MNV INDEL in frame SNV nonsense MNV nonsense INDEL out of frame
Biallelic/Multihit: SNV MNV INDEL in frame SNV nonsense MNV nonsense INDEL out of frame
Hotspot:

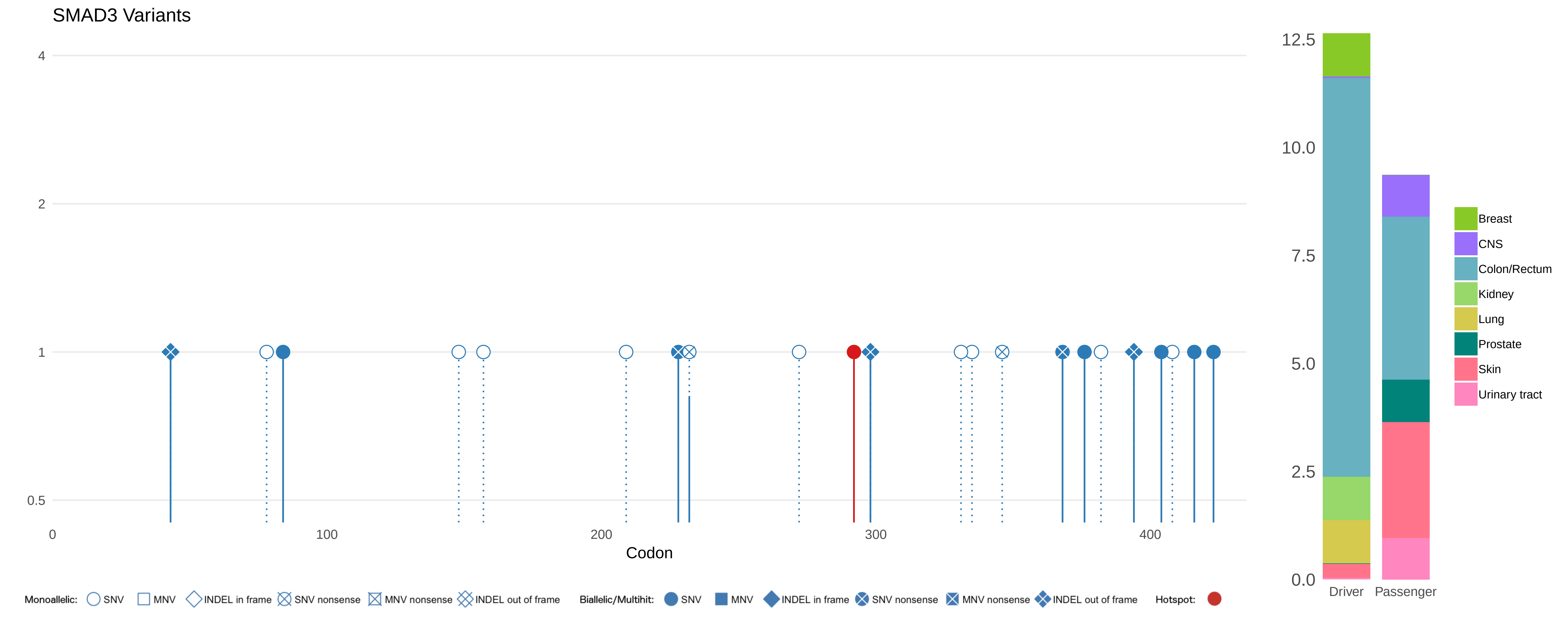
SMAD3 Variants
4
2
1
0.5
0
100
200
300
400
Codon
12.5
10.0
7.5
5.0
2.5
0.0
Driver
Passenger
Breast
CNS
Colon/Rectum
Kidney
Lung
Prostate
Skin
Urinary tract
Monoallelic: SNV MNV INDEL in frame SNV nonsense MNV nonsense INDEL out of frame
Biallelic/Multihit: SNV MNV INDEL in frame SNV nonsense MNV nonsense INDEL out of frame
Hotspot:

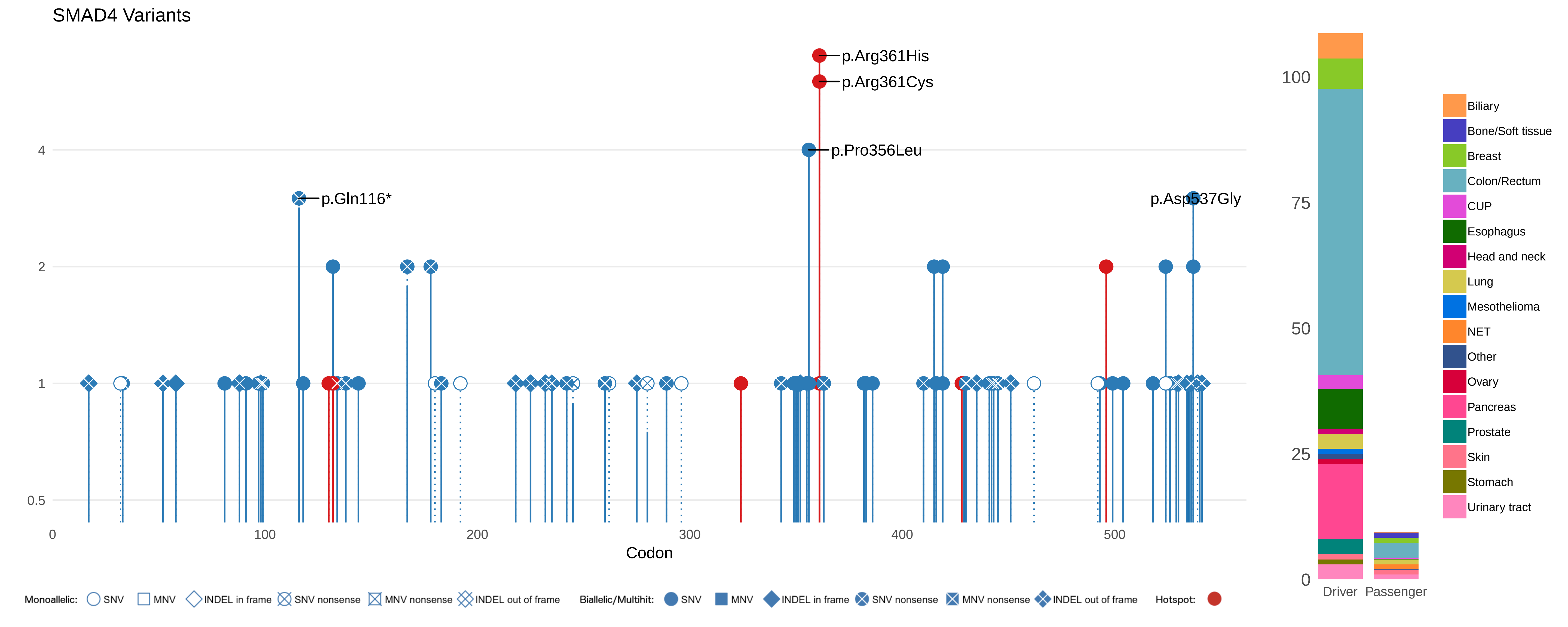

SMAD4 Variants
p.Arg361His
p.Arg361Cys
p.Pro356Leu
p.Gln116*
p.Asp537Gly
4
2
1
0.5
0
100
200
300
400
500
Codon
100
75
50
25
0
Driver
Passenger
Biliary
Bone/Soft tissue
Breast
Colon/Rectum
CUP
Esophagus
Head and neck
Lung
Mesothelioma
NET
Other
Ovary
Pancreas
Prostate
Skin
Stomach
Urinary tract
Monoallelic: SNV MNV INDEL in frame SNV nonsense MNV nonsense INDEL out of frame
Biallelic/Multihit: SNV MNV INDEL in frame SNV nonsense MNV nonsense INDEL out of frame
Hotspot:

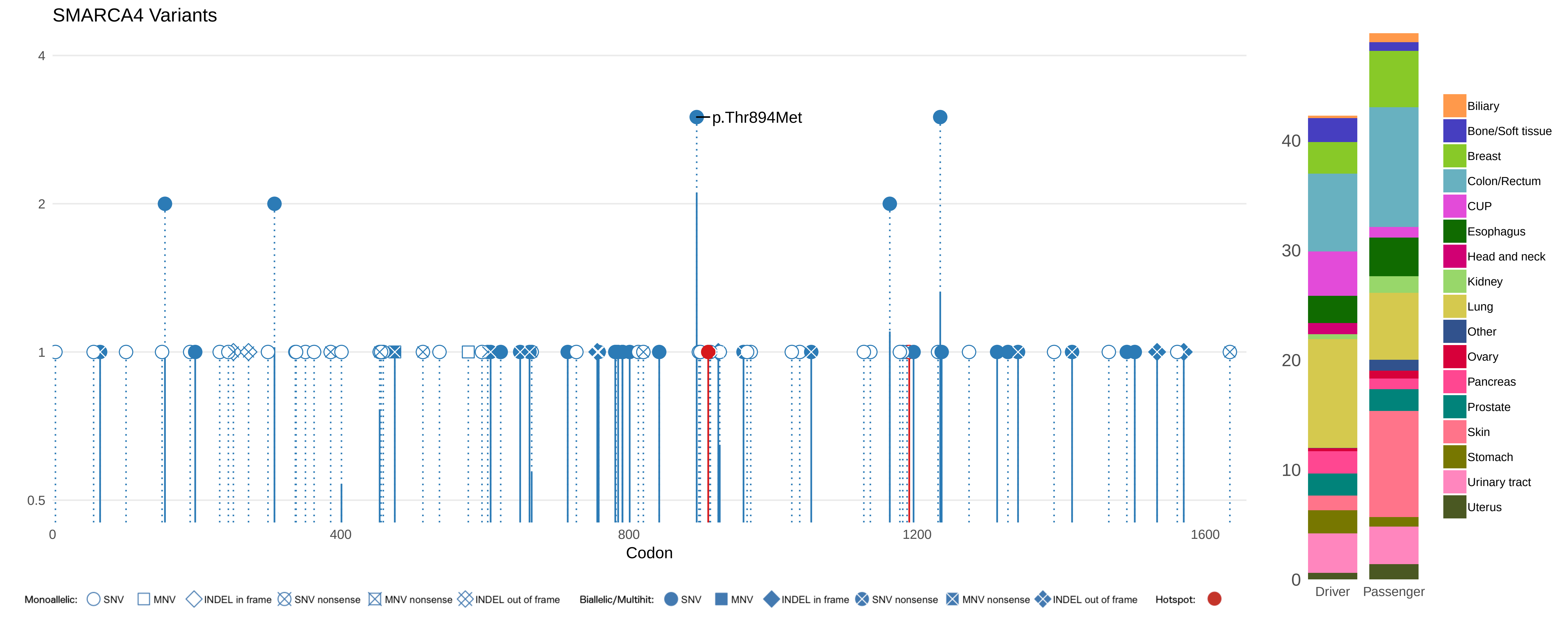

SMARCA4 Variants
4
2
1
0.5
p.Thr894Met
0
400
800
1200
1600
Codon
40
30
20
10
0
Driver
Passenger
Biliary
Bone/Soft tissue
Breast
Colon/Rectum
CUP
Esophagus
Head and neck
Kidney
Lung
Other
Ovary
Pancreas
Prostate
Skin
Stomach
Urinary tract
Uterus
Monoallelic: SNV MNV INDEL in frame SNV nonsense MNV nonsense INDEL out of frame
Biallelic/Multihit: SNV MNV INDEL in frame SNV nonsense MNV nonsense INDEL out of frame
Hotspot:

#### SMARCB1 Variants

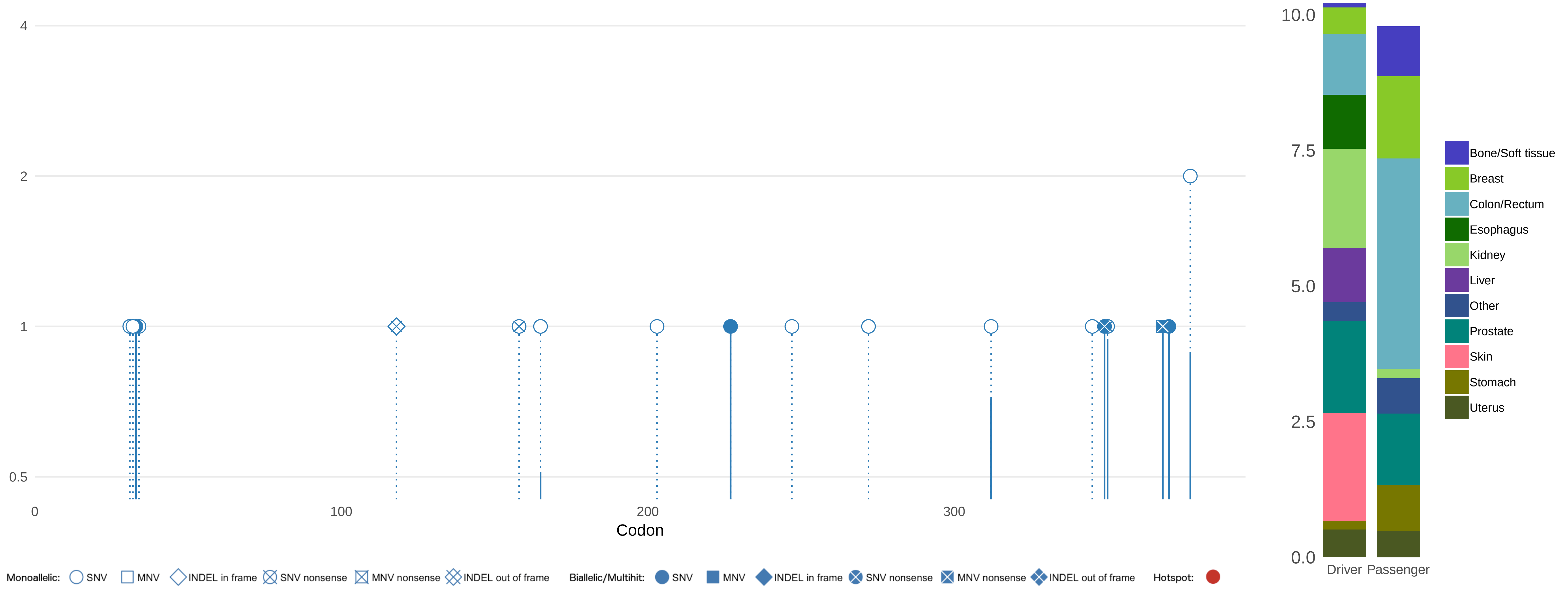

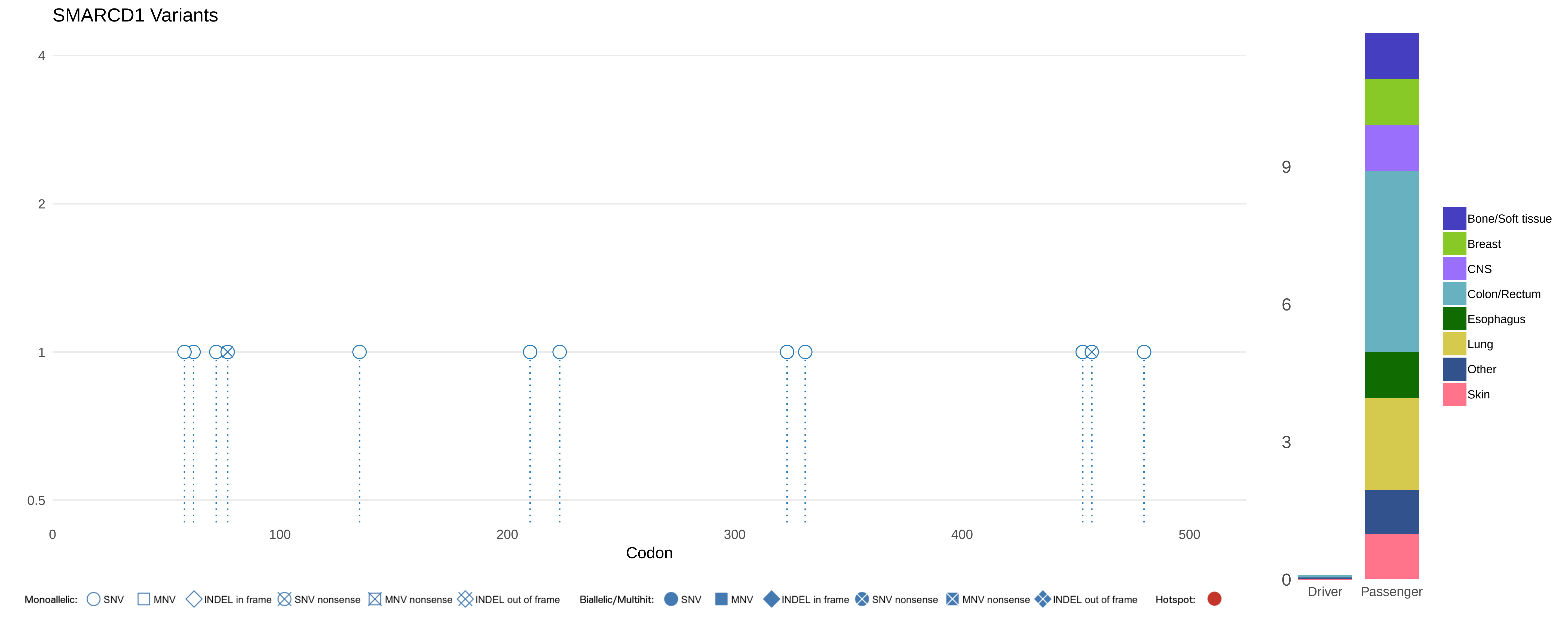
SMARCD1 Variants
4
2
1
0.5
0
100
200
300
400
500
Codon
9
6
3
0
Driver
Passenger
Bone/Soft tissue
Breast
CNS
Colon/Rectum
Esophagus
Lung
Other
Skin
Monoallelic: SNV MNV INDEL in frame SNV nonsense MNV nonsense INDEL out of frame
Biallelic/Multihit: SNV MNV INDEL in frame SNV nonsense MNV nonsense INDEL out of frame
Hotspot:

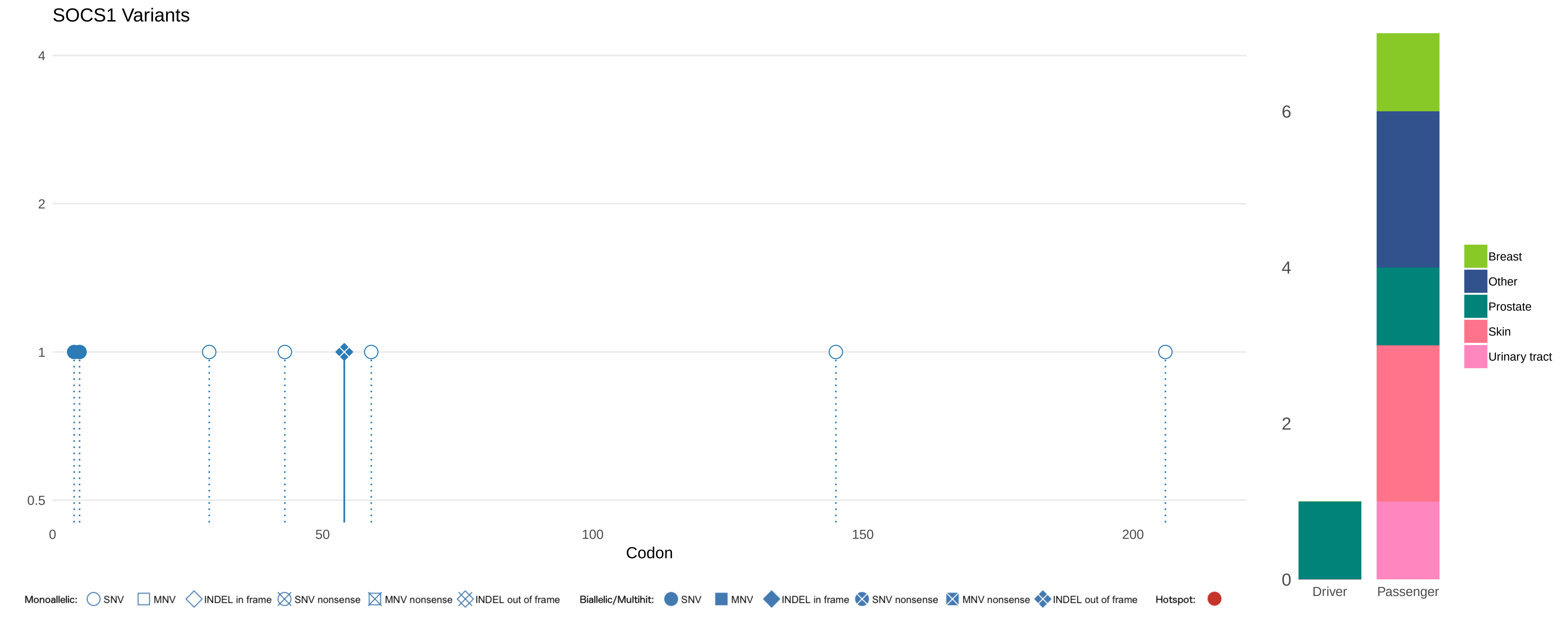
SOCS1 Variants
4
2
1
0.5
0
50
100
150
200
Codon
6
4
2
0
Driver
Passenger
Breast
Other
Prostate
Skin
Urinary tract
Monoallelic: SNV MNV INDEL in frame SNV nonsense MNV nonsense INDEL out of frame
Biallelic/Multihit: SNV MNV INDEL in frame SNV nonsense MNV nonsense INDEL out of frame
Hotspot:

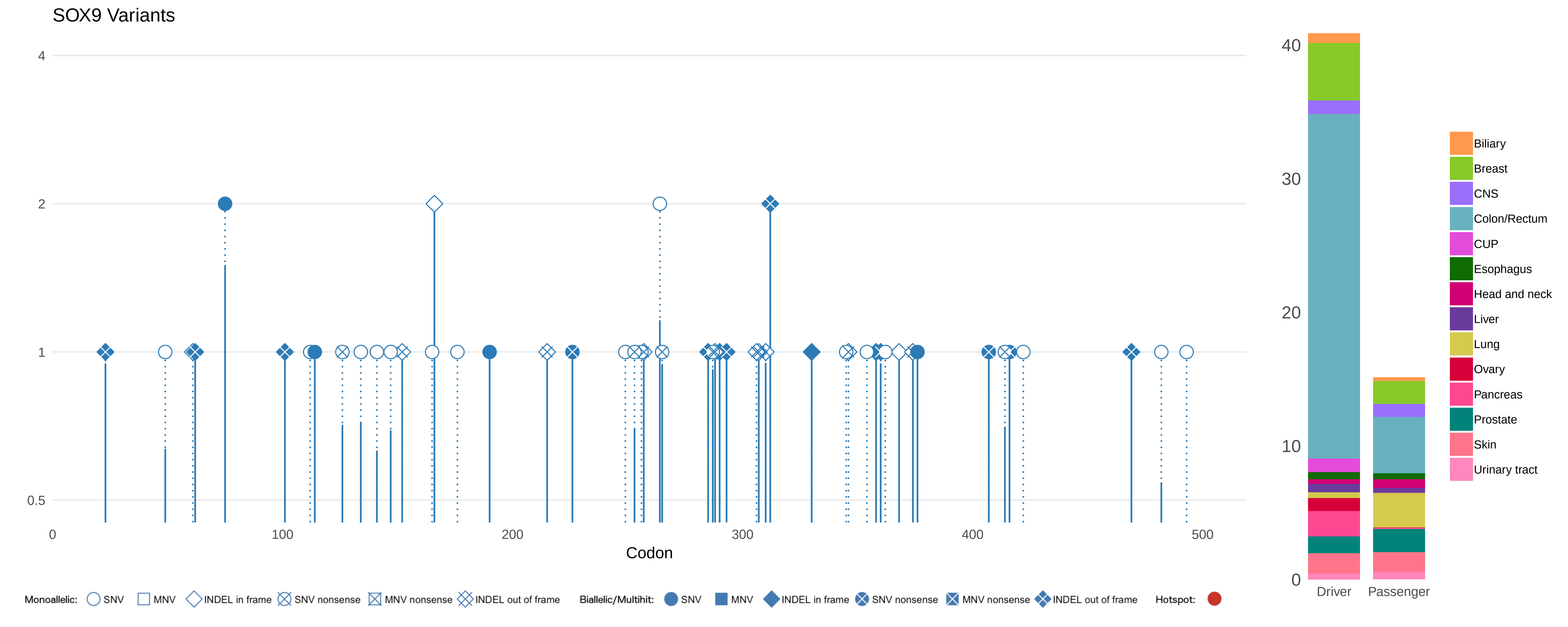

SOX9 Variants
4
2
1
0.5
0
100
200
300
400
500
Codon
40
30
20
10
0
Driver
Passenger
Biliary
Breast
CNS
Colon/Rectum
CUP
Esophagus
Head and neck
Liver
Lung
Ovary
Pancreas
Prostate
Skin
Urinary tract
Monoallelic: SNV MNV INDEL in frame SNV nonsense MNV nonsense INDEL out of frame
Biallelic/Multihit: SNV MNV INDEL in frame SNV nonsense MNV nonsense INDEL out of frame
Hotspot:

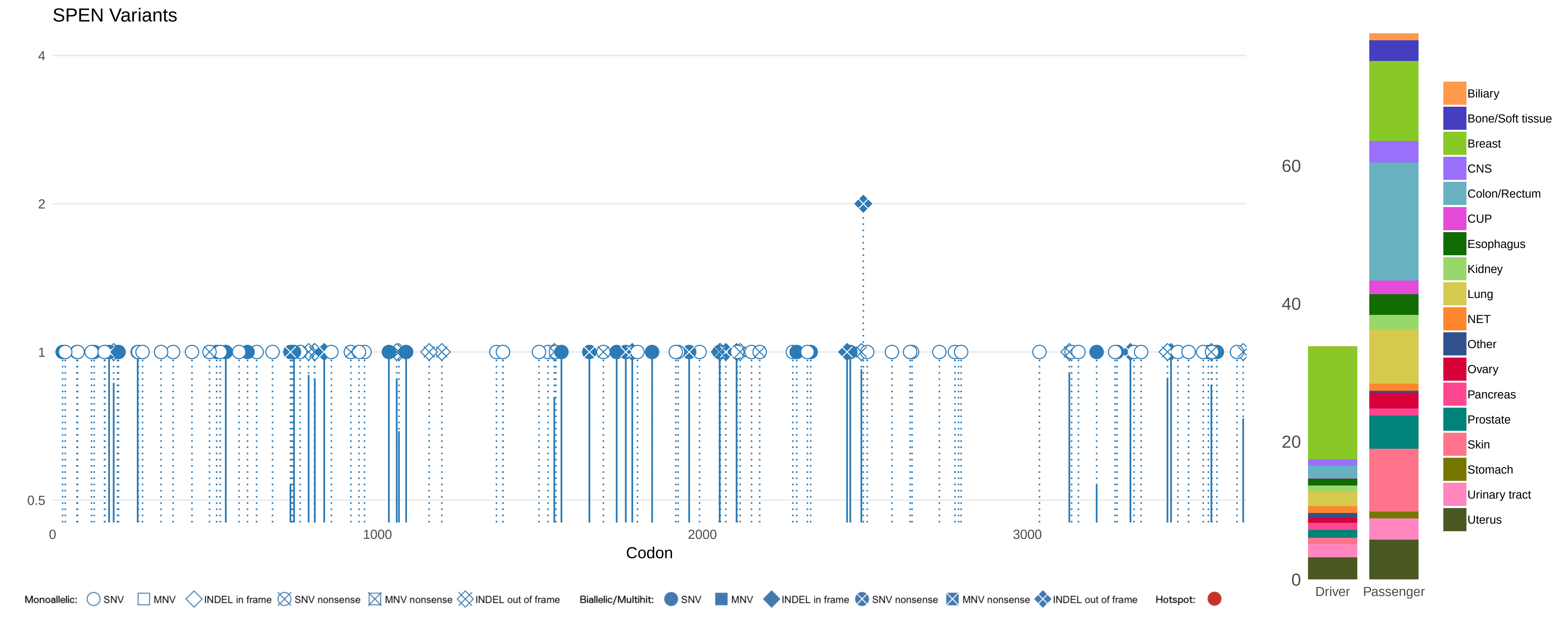
SPEN Variants
4
2
1
0.5
0
1000
2000
3000
Codon
60
40
20
0
Driver
Passenger
Biliary
Bone/Soft tissue
Breast
CNS
Colon/Rectum
CUP
Esophagus
Kidney
Lung
NET
Other
Ovary
Pancreas
Prostate
Skin
Stomach
Urinary tract
Uterus
Monoallelic: SNV MNV INDEL in frame SNV nonsense MNV nonsense INDEL out of frame
Biallelic/Multihit: SNV MNV INDEL in frame SNV nonsense MNV nonsense INDEL out of frame
Hotspot:

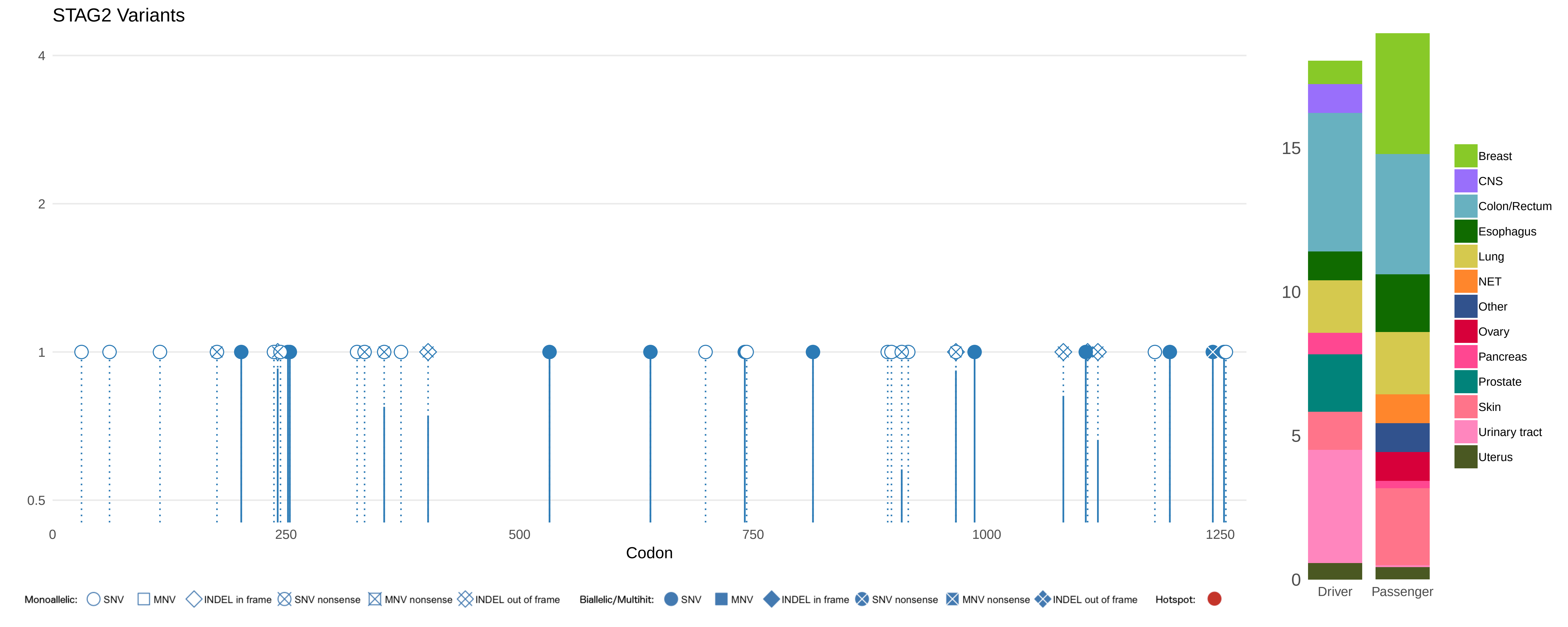
STAG2 Variants
4
2
1
0.5
0
250
500
750
1000
1250
Codon
15
10
5
0
Driver
Passenger
Breast
CNS
Colon/Rectum
Esophagus
Lung
NET
Other
Ovary
Pancreas
Prostate
Skin
Urinary tract
Uterus
Monoallelic: SNV MNV INDEL in frame SNV nonsense MNV nonsense INDEL out of frame
Biallelic/Multihit: SNV MNV INDEL in frame SNV nonsense MNV nonsense INDEL out of frame
Hotspot:

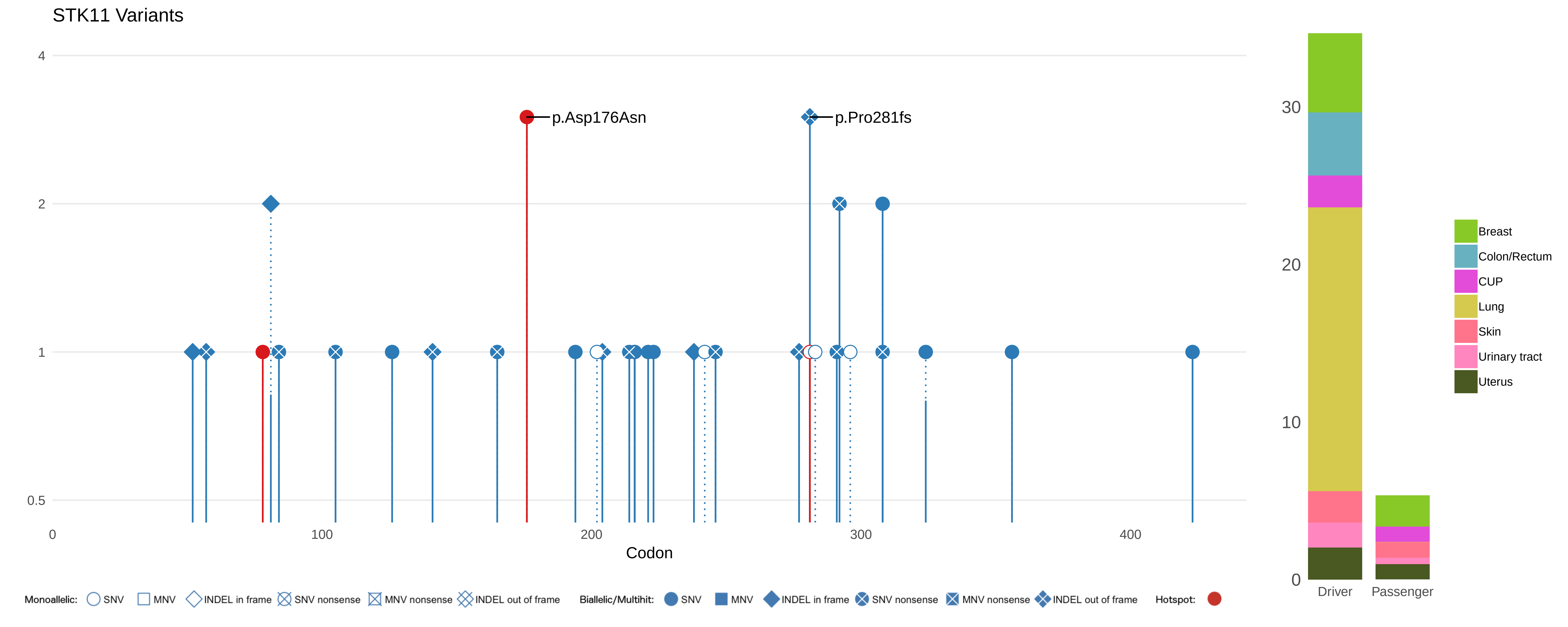

STK11 Variants
4
2
1
0.5
0
100
200
300
400
Codon
p.Asp176Asn
p.Pro281fs
30
20
10
0
Driver
Passenger
Breast
Colon/Rectum
CUP
Lung
Skin
Urinary tract
Uterus
Monoallelic:
SNV
MNV
INDEL in frame
SNV nonsense
MNV nonsense
INDEL out of frame
Biallelic/Multihit:
SNV
MNV
INDEL in frame
SNV nonsense
MNV nonsense
INDEL out of frame
Hotspot:

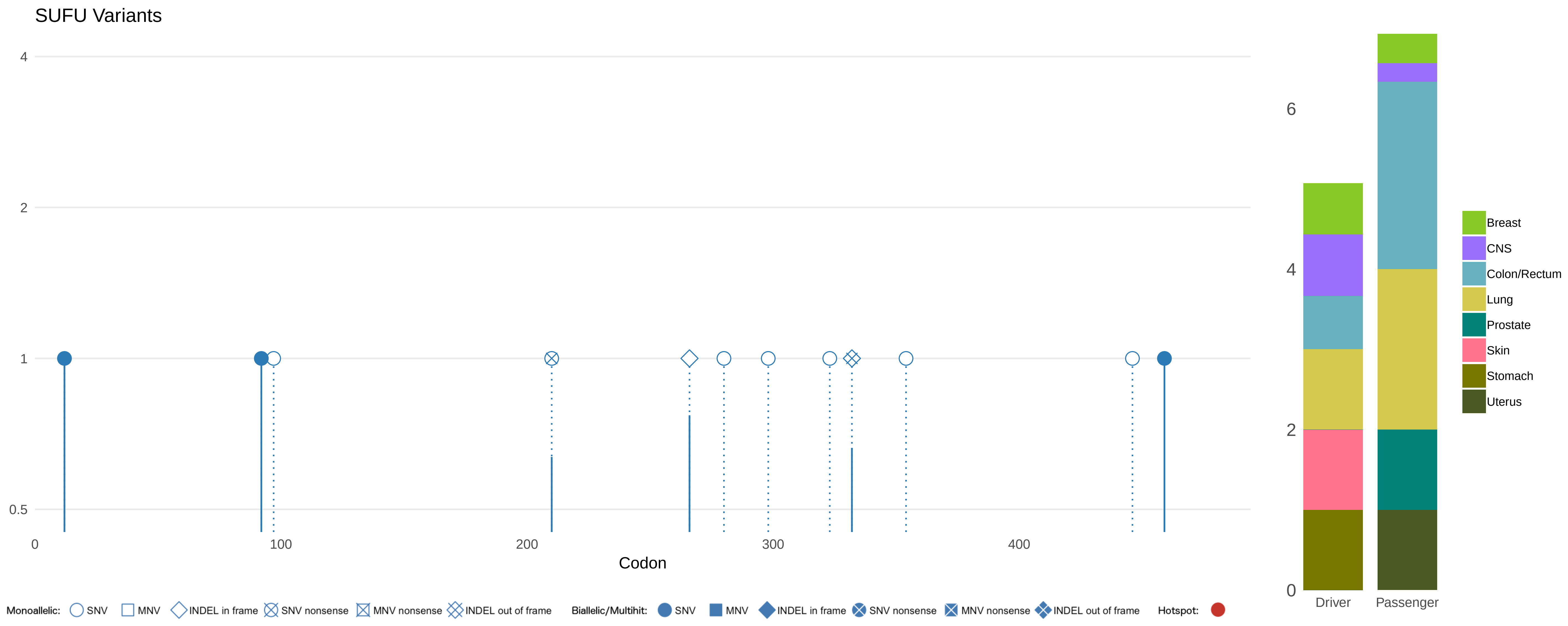
SUFU Variants
4
2
1
0.5
0
100
200
300
400
Codon
Monoallelic: SNV MNV INDEL in frame SNV nonsense MNV nonsense INDEL out of frame
Biallelic/Multihit: SNV MNV INDEL in frame SNV nonsense MNV nonsense INDEL out of frame
Hotspot:
6
4
2
0
Driver
Passenger
Breast
CNS
Colon/Rectum
Lung
Prostate
Skin
Stomach
Uterus

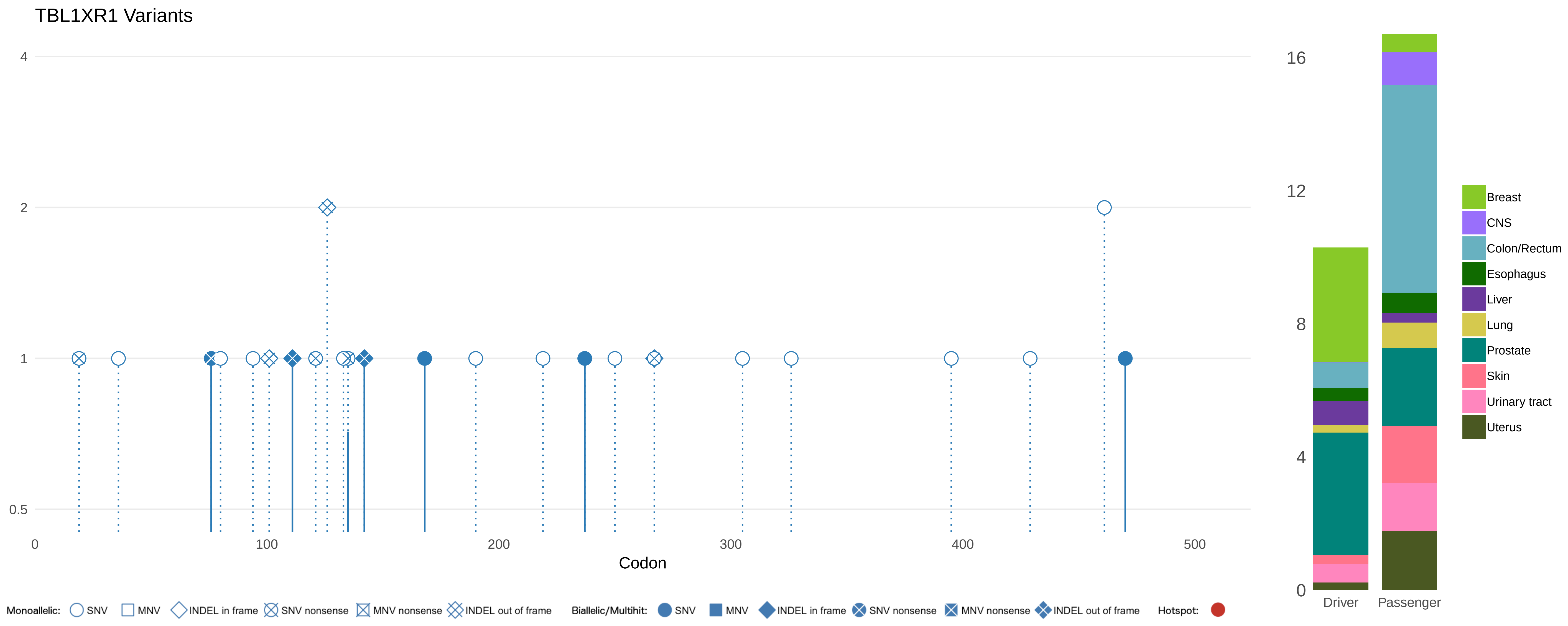

TBL1XR1 Variants
4
2
1
0.5
0
100
200
300
400
500
Codon
Monoallelic: SNV MNV INDEL in frame SNV nonsense MNV nonsense INDEL out of frame
Biallelic/Multihit: SNV MNV INDEL in frame SNV nonsense MNV nonsense INDEL out of frame
Hotspot:
16
12
8
4
0
Driver
Passenger
Breast
CNS
Colon/Rectum
Esophagus
Liver
Lung
Prostate
Skin
Urinary tract
Uterus

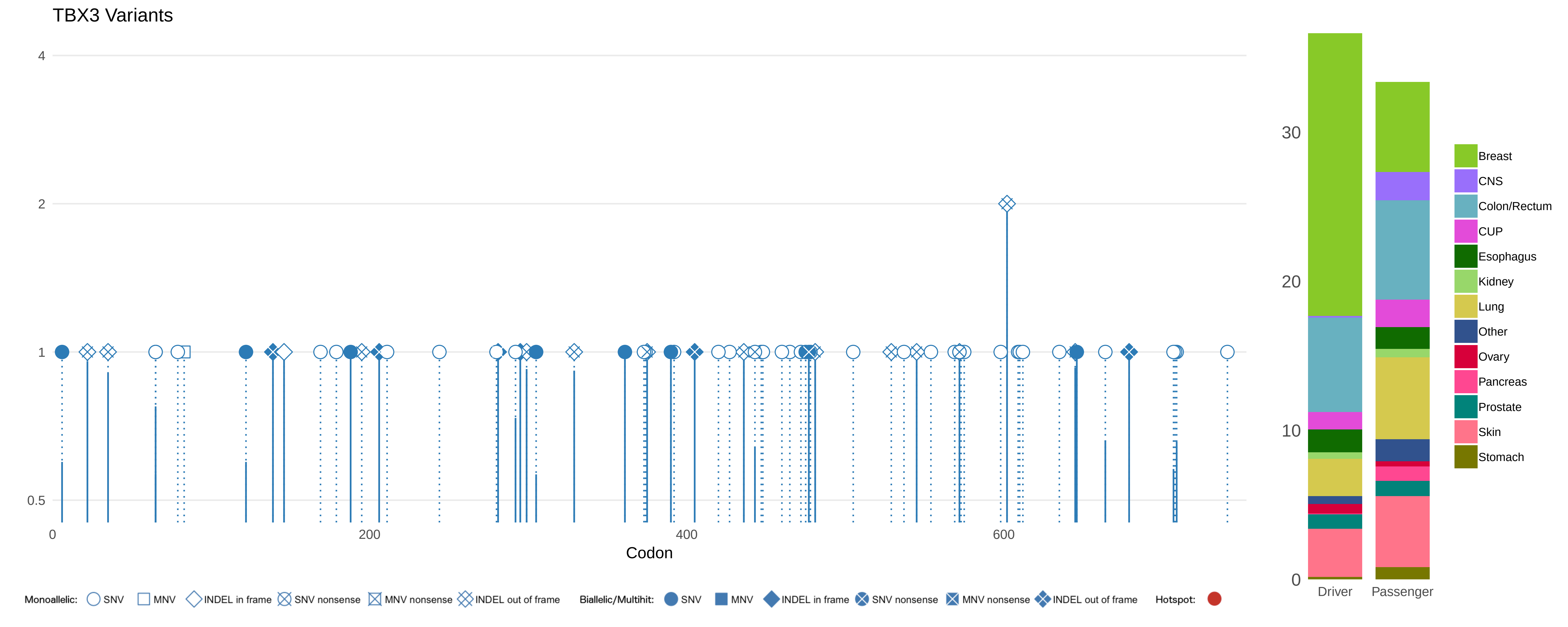
TBX3 Variants
4
2
1
0.5
0
200
400
600
Codon
Monoallelic: SNV MNV INDEL in frame SNV nonsense MNV nonsense INDEL out of frame
Biallelic/Multihit: SNV MNV INDEL in frame SNV nonsense MNV nonsense INDEL out of frame
Hotspot:
30
20
10
0
Driver
Passenger
Breast
CNS
Colon/Rectum
CUP
Esophagus
Kidney
Lung
Other
Ovary
Pancreas
Prostate
Skin
Stomach

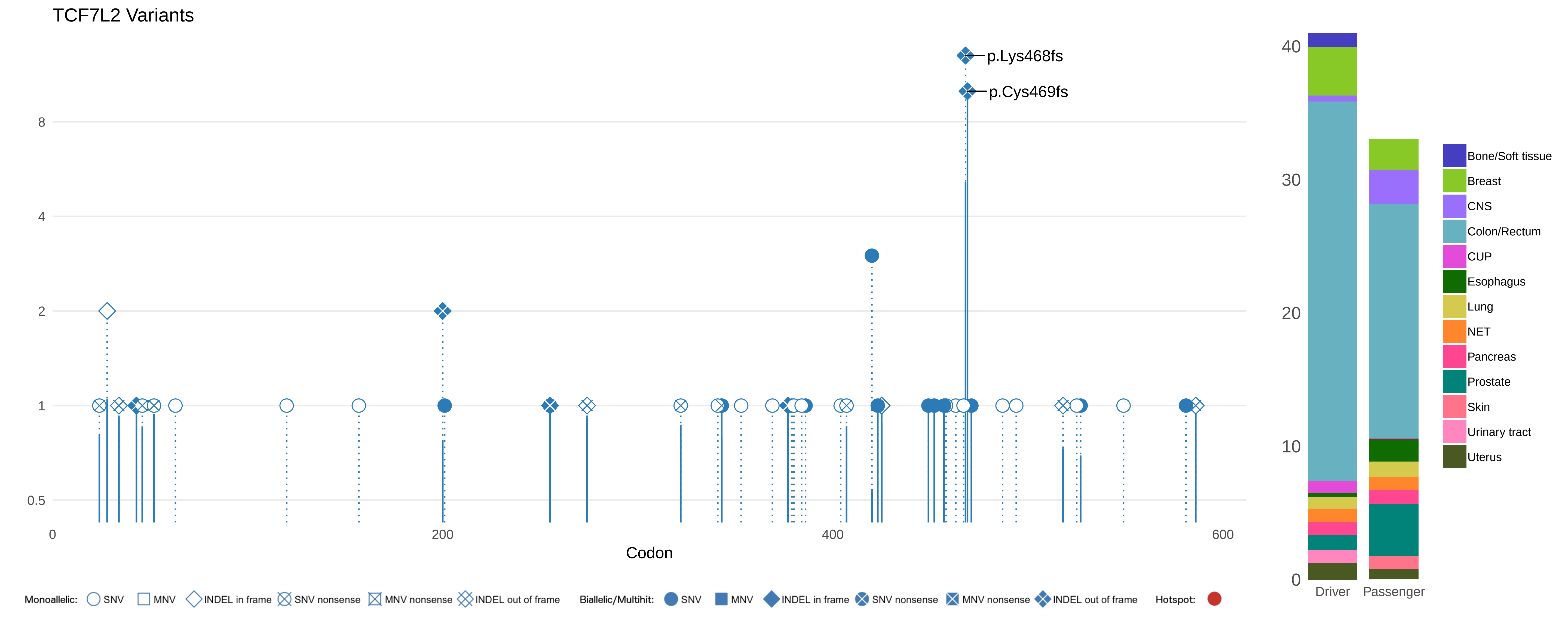
TCF7L2 Variants
p.Lys468fs
p.Cys469fs
8
4
2
1
0.5
0
200
400
600
Codon
40
30
20
10
0
Driver
Passenger
Bone/Soft tissue
Breast
CNS
Colon/Rectum
CUP
Esophagus
Lung
NET
Pancreas
Prostate
Skin
Urinary tract
Uterus
Monoallelic: SNV MNV INDEL in frame SNV nonsense MNV nonsense INDEL out of frame
Biallelic/Multihit: SNV MNV INDEL in frame SNV nonsense MNV nonsense INDEL out of frame
Hotspot:

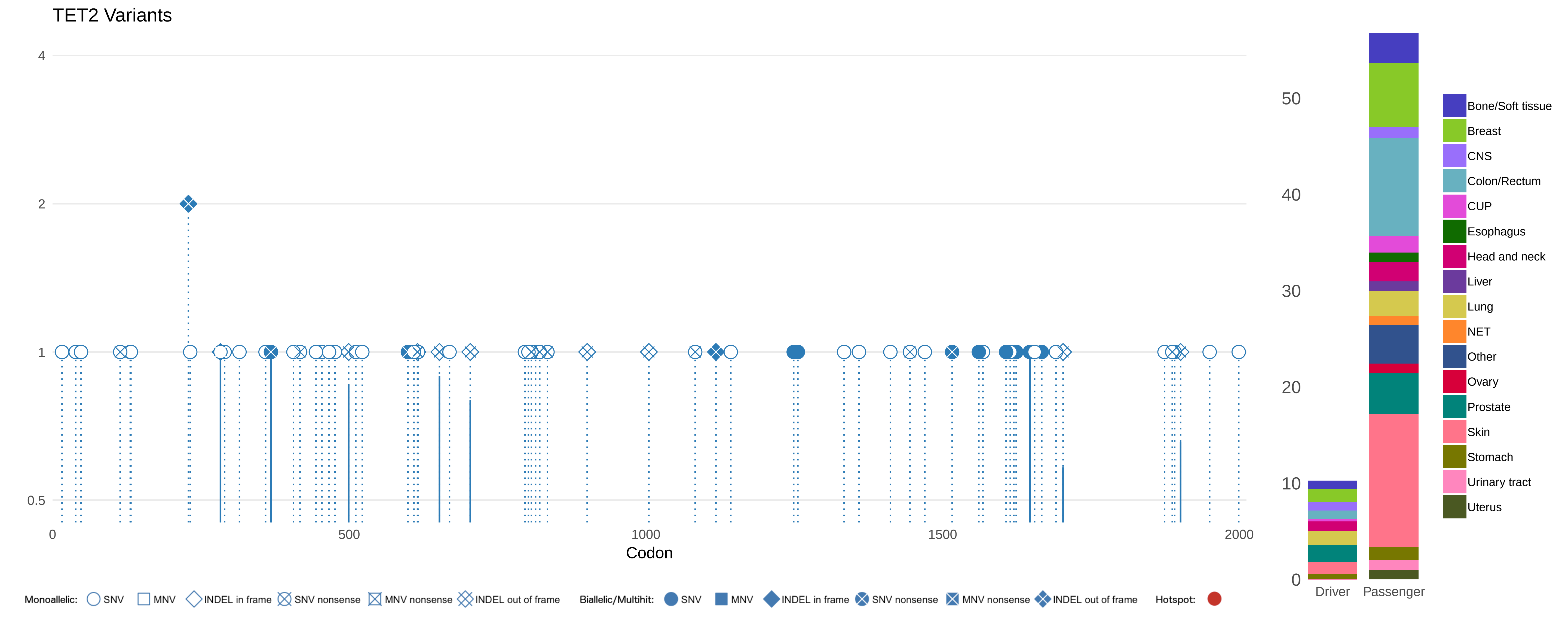

TET2 Variants
4
2
1
0.5
0
500
1000
1500
2000
Codon
50
40
30
20
10
0
Driver
Passenger
Bone/Soft tissue
Breast
CNS
Colon/Rectum
CUP
Esophagus
Head and neck
Liver
Lung
NET
Other
Ovary
Prostate
Skin
Stomach
Urinary tract
Uterus
Monoallelic: SNV MNV INDEL in frame SNV nonsense MNV nonsense INDEL out of frame
Biallelic/Multihit: SNV MNV INDEL in frame SNV nonsense MNV nonsense INDEL out of frame
Hotspot:

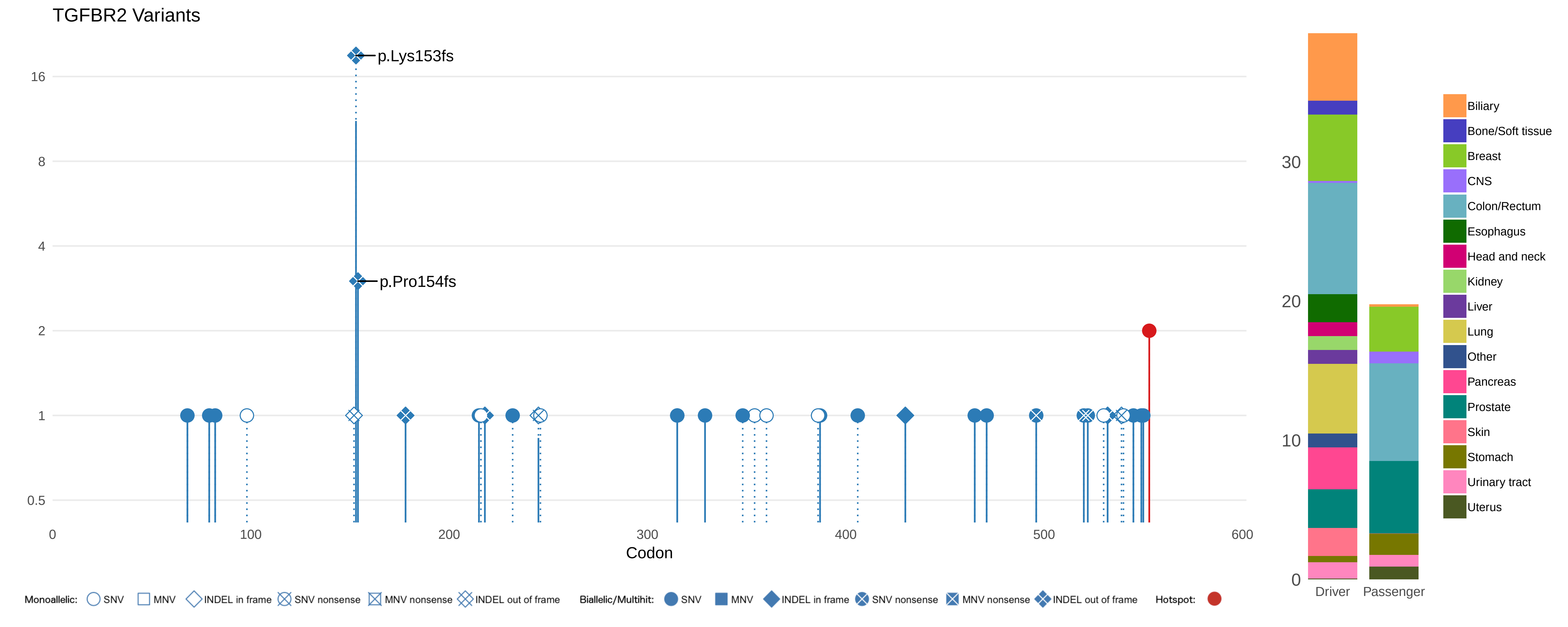
TGFBR2 Variants
p.Lys153fs
p.Pro154fs
16
8
4
2
1
0.5
0
100
200
300
400
500
600
Codon
30
20
10
0
Driver
Passenger
Biliary
Bone/Soft tissue
Breast
CNS
Colon/Rectum
Esophagus
Head and neck
Kidney
Liver
Lung
Other
Pancreas
Prostate
Skin
Stomach
Urinary tract
Uterus
Monoallelic: SNV MNV INDEL in frame SNV nonsense MNV nonsense INDEL out of frame
Biallelic/Multihit: SNV MNV INDEL in frame SNV nonsense MNV nonsense INDEL out of frame
Hotspot:

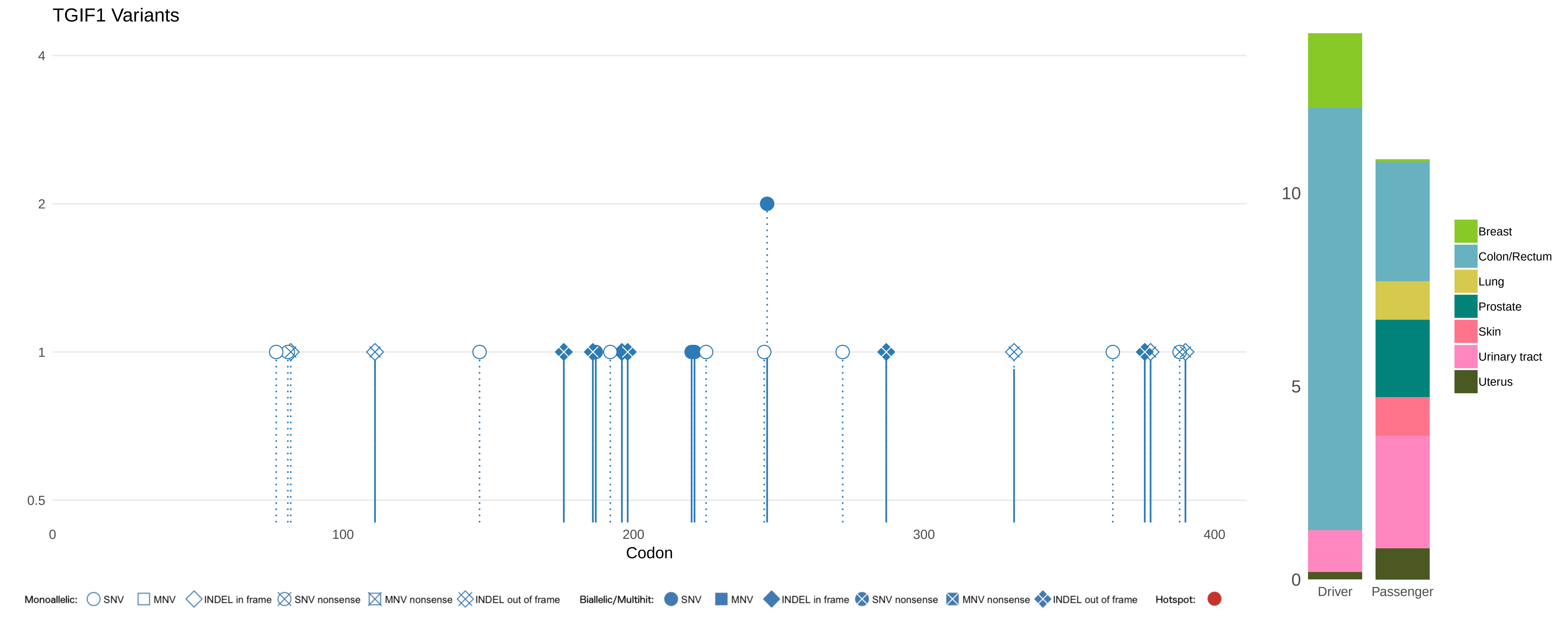
TGIF1 Variants
4
2
1
0.5
0
100
200
300
400
Codon
10
5
0
Driver
Passenger
Breast
Colon/Rectum
Lung
Prostate
Skin
Urinary tract
Uterus
Monoallelic:
SNV
MNV
INDEL in frame
SNV nonsense
MNV nonsense
INDEL out of frame
Biallelic/Multihit:
SNV
MNV
INDEL in frame
SNV nonsense
MNV nonsense
INDEL out of frame
Hotspot:

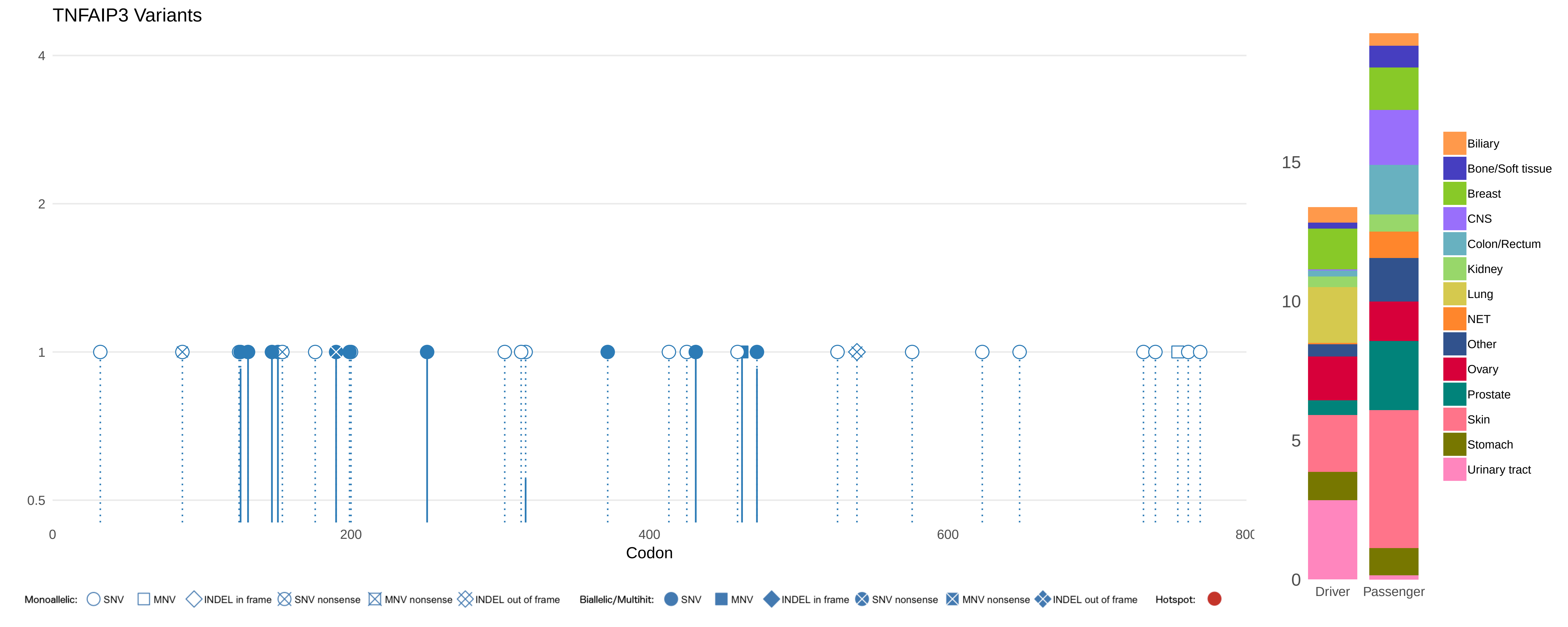
TNFAIP3 Variants
4
2
1
0.5
0
200
400
600
800
Codon
15
10
5
0
Driver
Passenger
Biliary
Bone/Soft tissue
Breast
CNS
Colon/Rectum
Kidney
Lung
NET
Other
Ovary
Prostate
Skin
Stomach
Urinary tract
Monoallelic: SNV MNV INDEL in frame SNV nonsense MNV nonsense INDEL out of frame
Biallelic/Multihit: SNV MNV INDEL in frame SNV nonsense MNV nonsense INDEL out of frame
Hotspot:

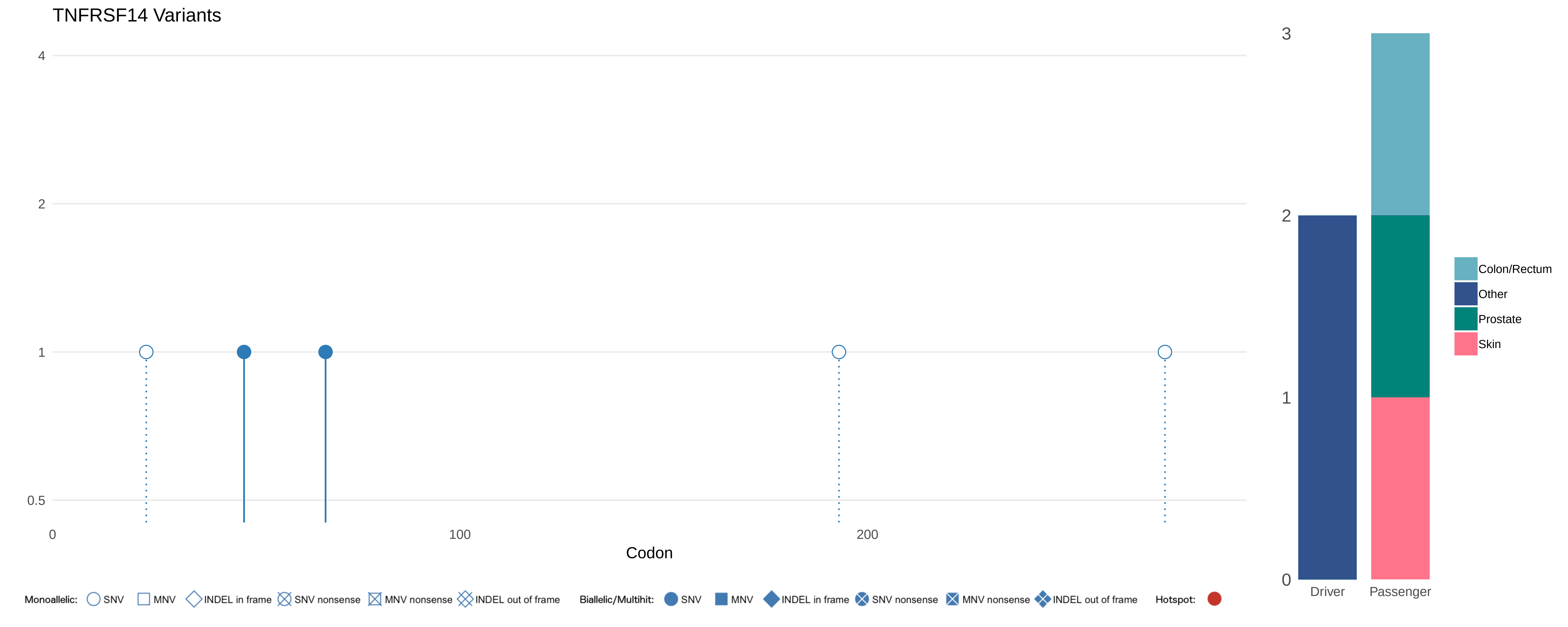
TNFRSF14 Variants
4
2
1
0.5
0
100
200
Codon
Monoallelic: SNV MNV INDEL in frame SNV nonsense MNV nonsense INDEL out of frame
Biallelic/Multihit: SNV MNV INDEL in frame SNV nonsense MNV nonsense INDEL out of frame
Hotspot:
3
2
1
0
Driver
Passenger
Colon/Rectum
Other
Prostate
Skin

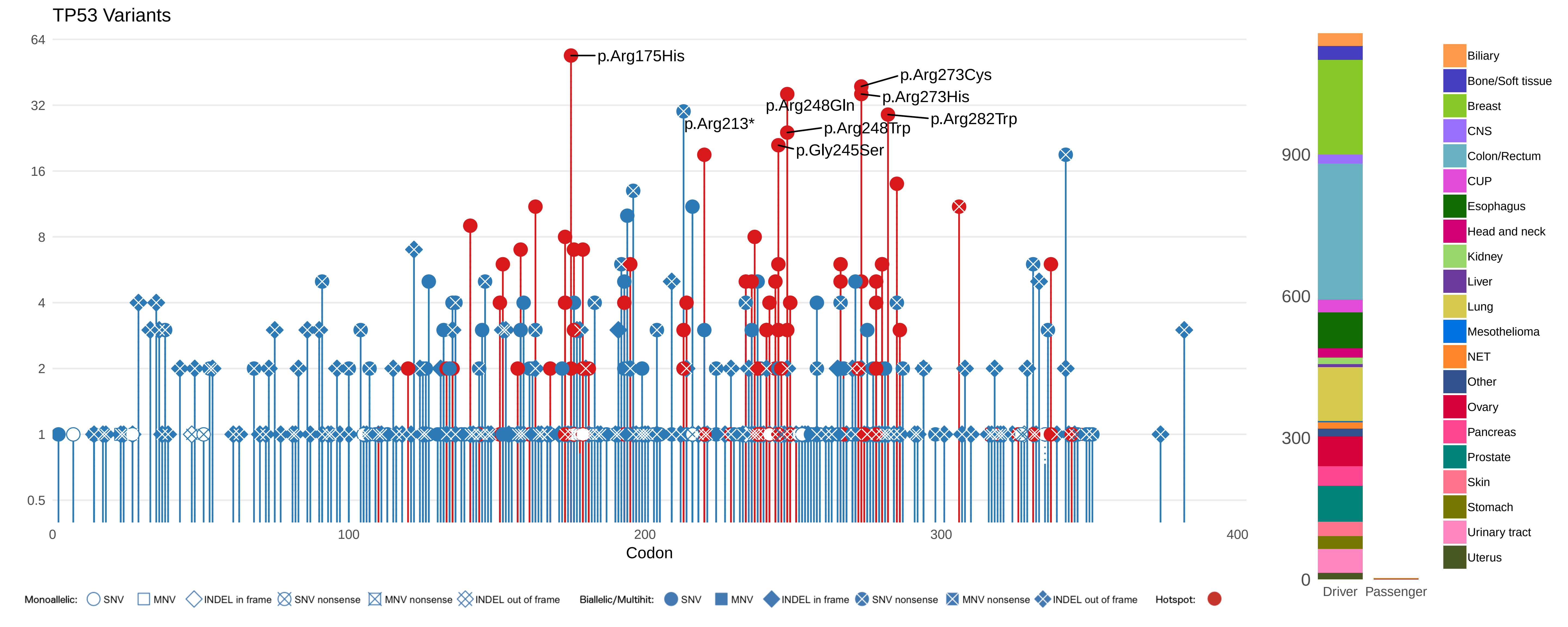
TP53 Variants
64
32
16
8
4
2
1
0.5
p.Arg175His
p.Arg273Cys
p.Arg273His
p.Arg248Gln
p.Arg282Trp
p.Arg213*
p.Arg248Trp
p.Gly245Ser
0
100
200
300
400
Codon
900
600
300
0
Driver
Passenger
Biliary
Bone/Soft tissue
Breast
CNS
Colon/Rectum
CUP
Esophagus
Head and neck
Kidney
Liver
Lung
Mesothelioma
NET
Other
Ovary
Pancreas
Prostate
Skin
Stomach
Urinary tract
Uterus
Monoallelic: SNV MNV INDEL in frame SNV nonsense MNV nonsense INDEL out of frame
Biallelic/Multihit: SNV MNV INDEL in frame SNV nonsense MNV nonsense INDEL out of frame
Hotspot:

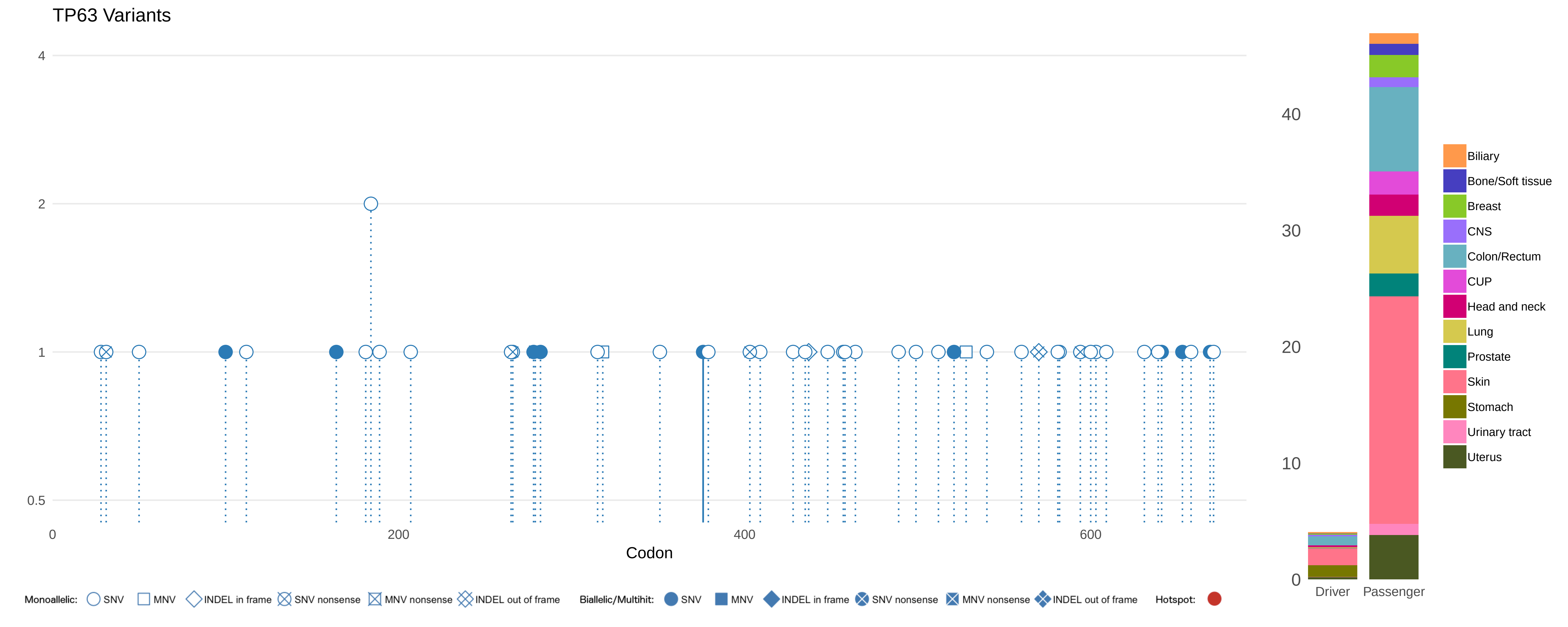
TP63 Variants
4
2
1
0.5
0
200
400
600
Codon
40
30
20
10
0
Driver
Passenger
Biliary
Bone/Soft tissue
Breast
CNS
Colon/Rectum
CUP
Head and neck
Lung
Prostate
Skin
Stomach
Urinary tract
Uterus
Monoallelic: SNV MNV INDEL in frame SNV nonsense MNV nonsense INDEL out of frame
Biallelic/Multihit: SNV MNV INDEL in frame SNV nonsense MNV nonsense INDEL out of frame
Hotspot:

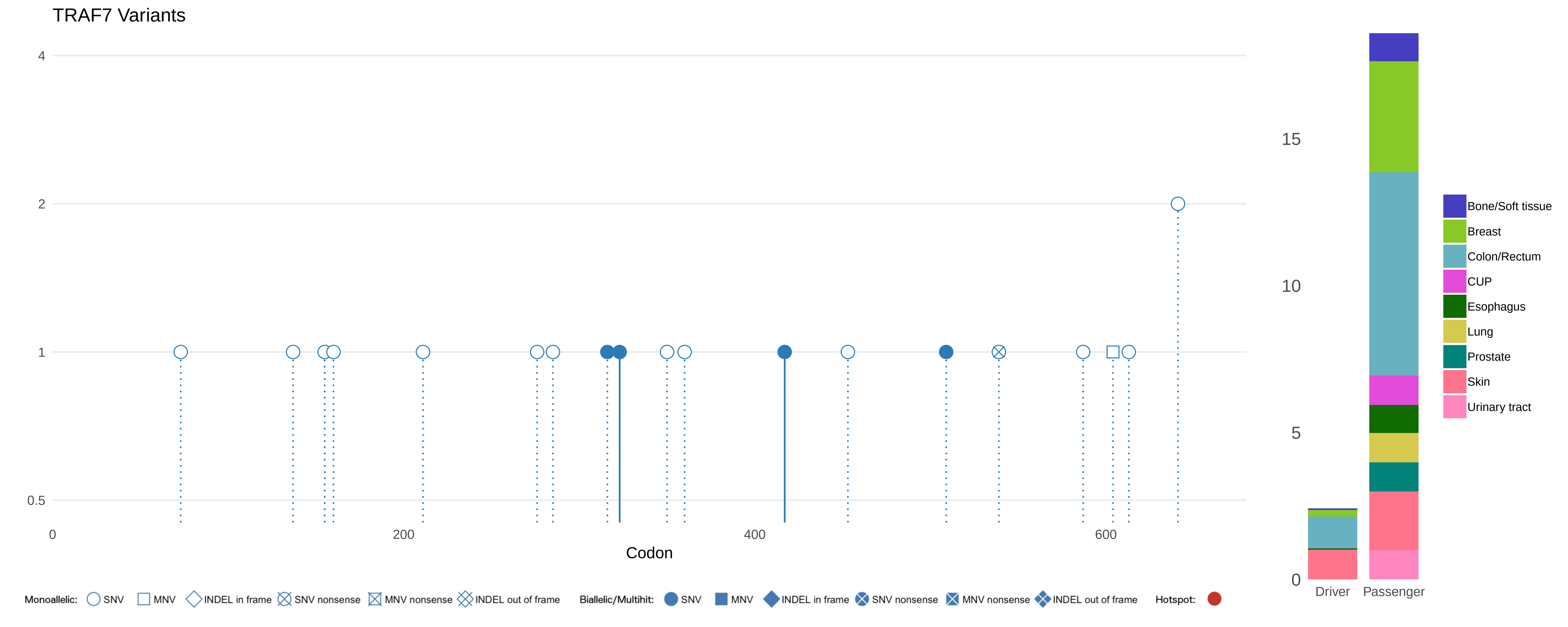
TRAF7 Variants
4
2
1
0.5
0
200
400
600
Codon
15
10
5
0
Driver
Passenger
Bone/Soft tissue
Breast
Colon/Rectum
CUP
Esophagus
Lung
Prostate
Skin
Urinary tract
Monoallelic: SNV MNV INDEL in frame SNV nonsense MNV nonsense INDEL out of frame
Biallelic/Multihit: SNV MNV INDEL in frame SNV nonsense MNV nonsense INDEL out of frame
Hotspot:

#### TSC1 Variants

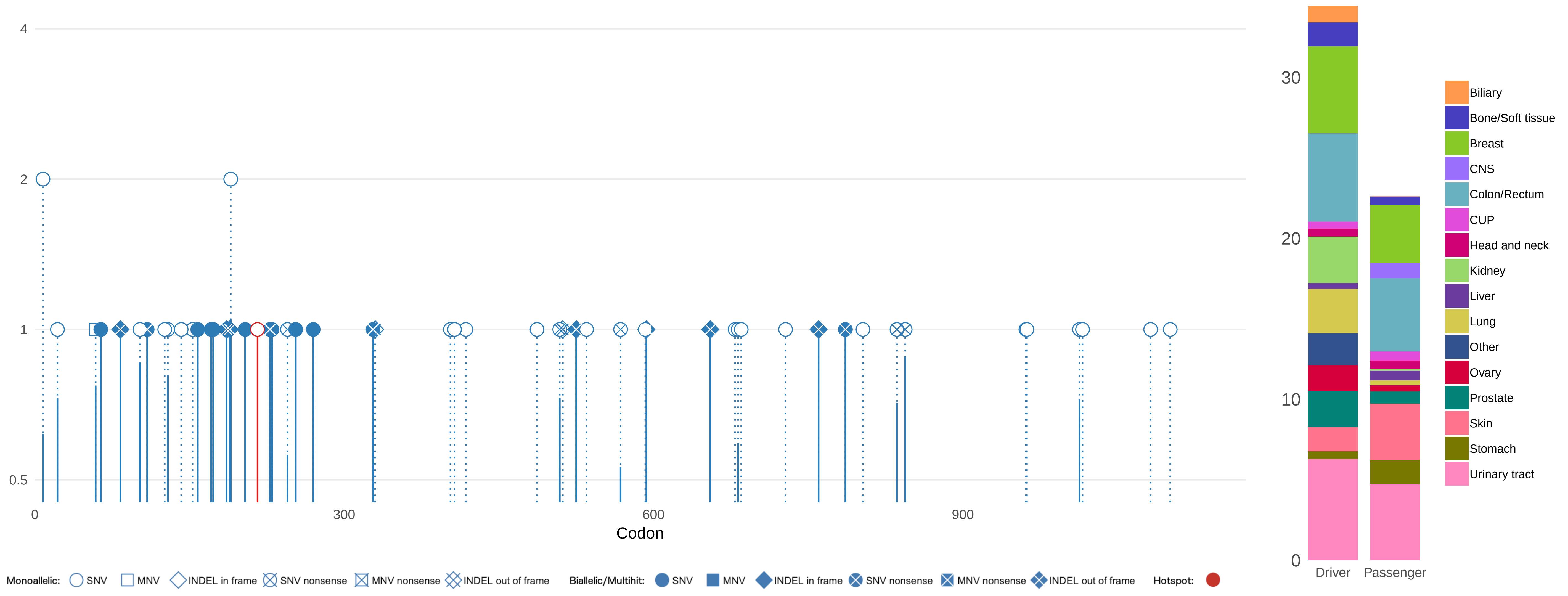

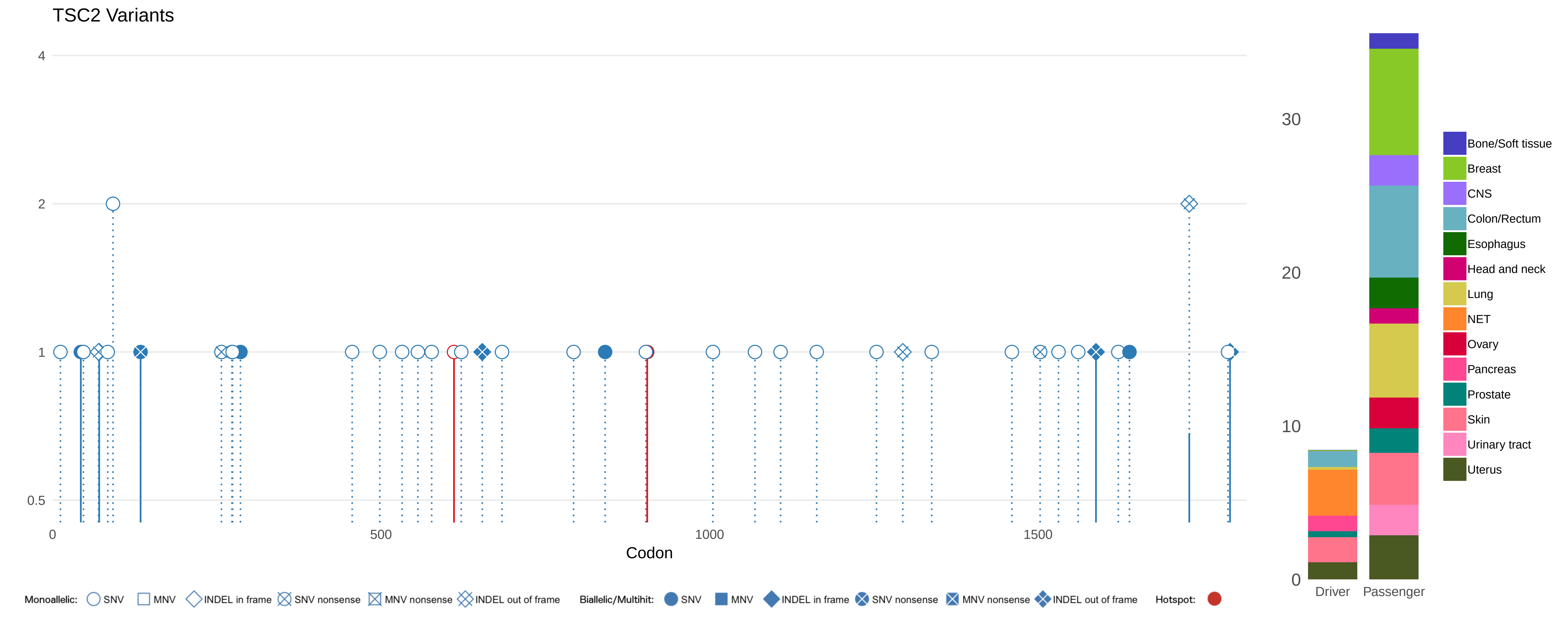
TSC2 Variants
4
2
1
0.5
0
500
1000
1500
Codon
30
20
10
0
Driver
Passenger
Bone/Soft tissue
Breast
CNS
Colon/Rectum
Esophagus
Head and neck
Lung
NET
Ovary
Pancreas
Prostate
Skin
Urinary tract
Uterus
Monoallelic: SNV MNV INDEL in frame SNV nonsense MNV nonsense INDEL out of frame
Biallelic/Multihit: SNV MNV INDEL in frame SNV nonsense MNV nonsense INDEL out of frame
Hotspot:

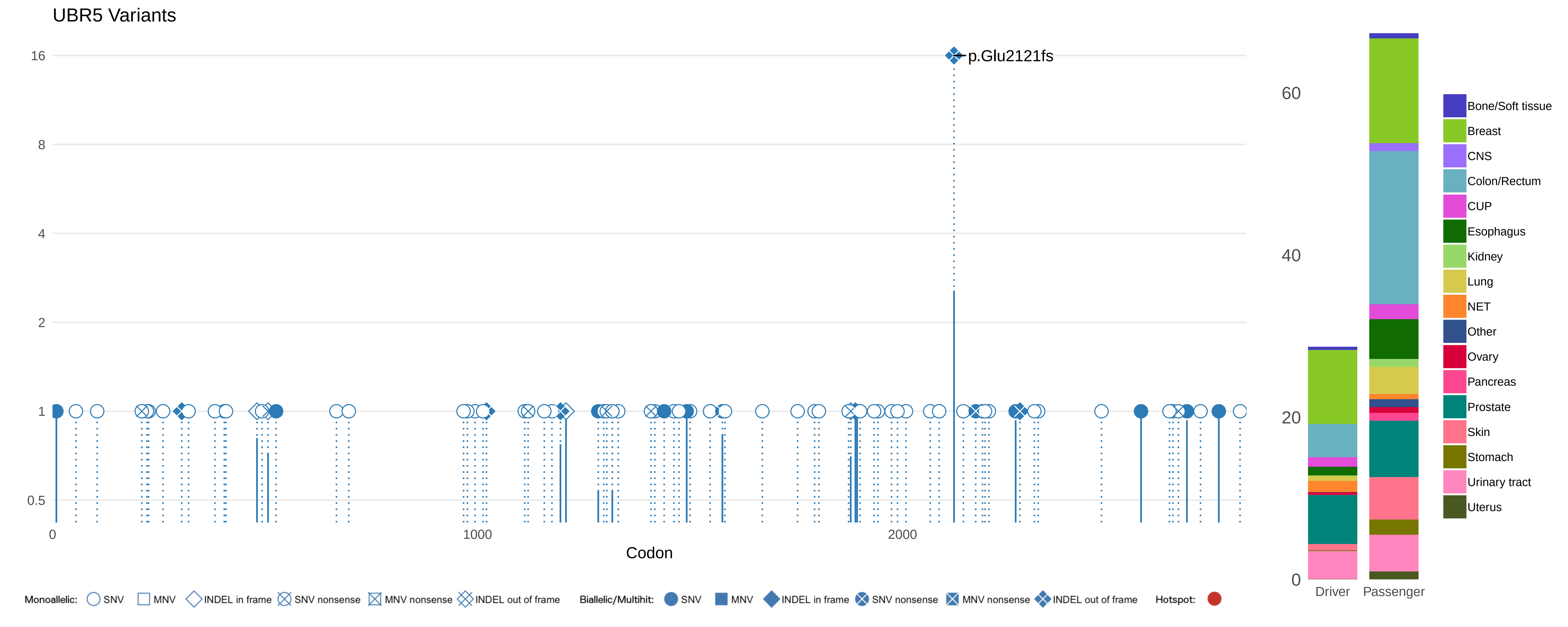

UBR5 Variants
16
8
4
2
1
0.5
p.Glu2121fs
0
1000
2000
Codon
60
40
20
0
Driver
Passenger
Bone/Soft tissue
Breast
CNS
Colon/Rectum
CUP
Esophagus
Kidney
Lung
NET
Other
Ovary
Pancreas
Prostate
Skin
Stomach
Urinary tract
Uterus
Monoallelic: SNV MNV INDEL in frame SNV nonsense MNV nonsense INDEL out of frame
Biallelic/Multihit: SNV MNV INDEL in frame SNV nonsense MNV nonsense INDEL out of frame
Hotspot:

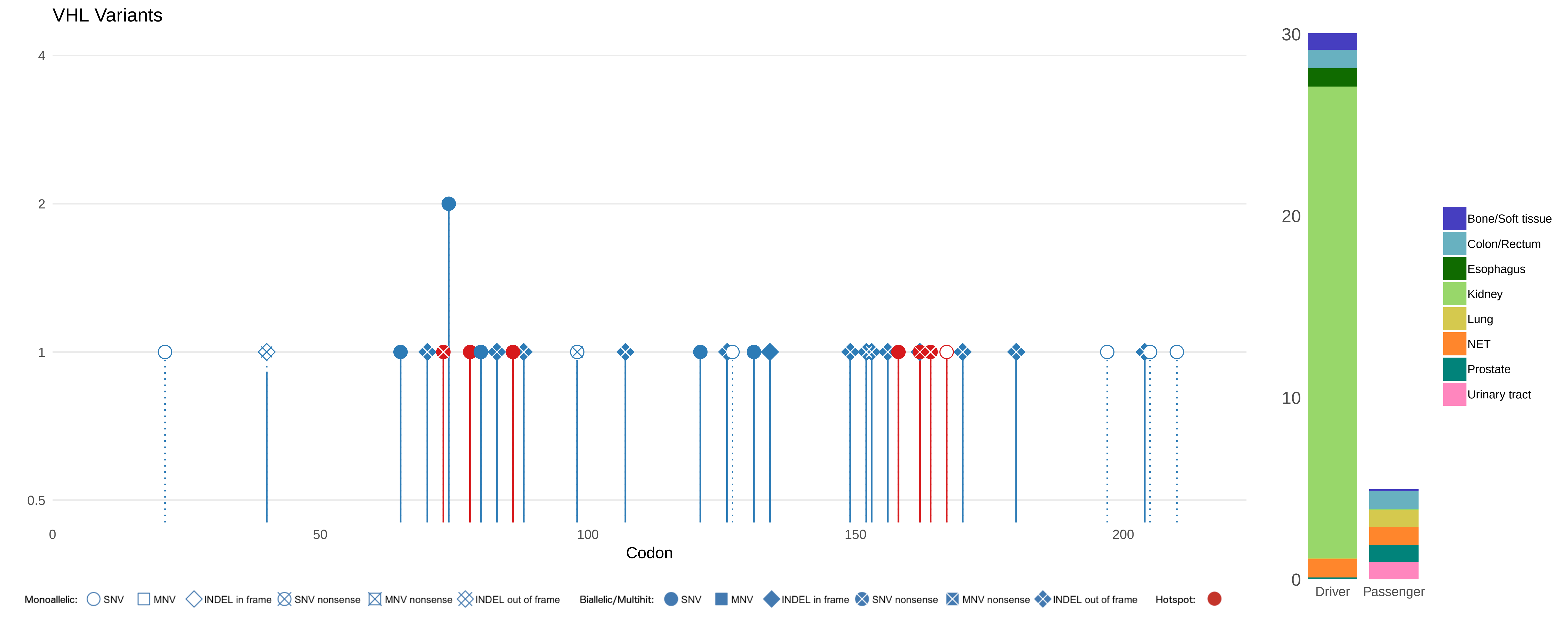
VHL Variants
4
2
1
0.5
0
50
100
150
200
Codon
Monoallelic: SNV MNV INDEL in frame SNV nonsense MNV nonsense INDEL out of frame
Biallelic/Multihit: SNV MNV INDEL in frame SNV nonsense MNV nonsense INDEL out of frame
Hotspot:
30
20
10
0
Driver
Passenger
Bone/Soft tissue
Colon/Rectum
Esophagus
Kidney
Lung
NET
Prostate
Urinary tract

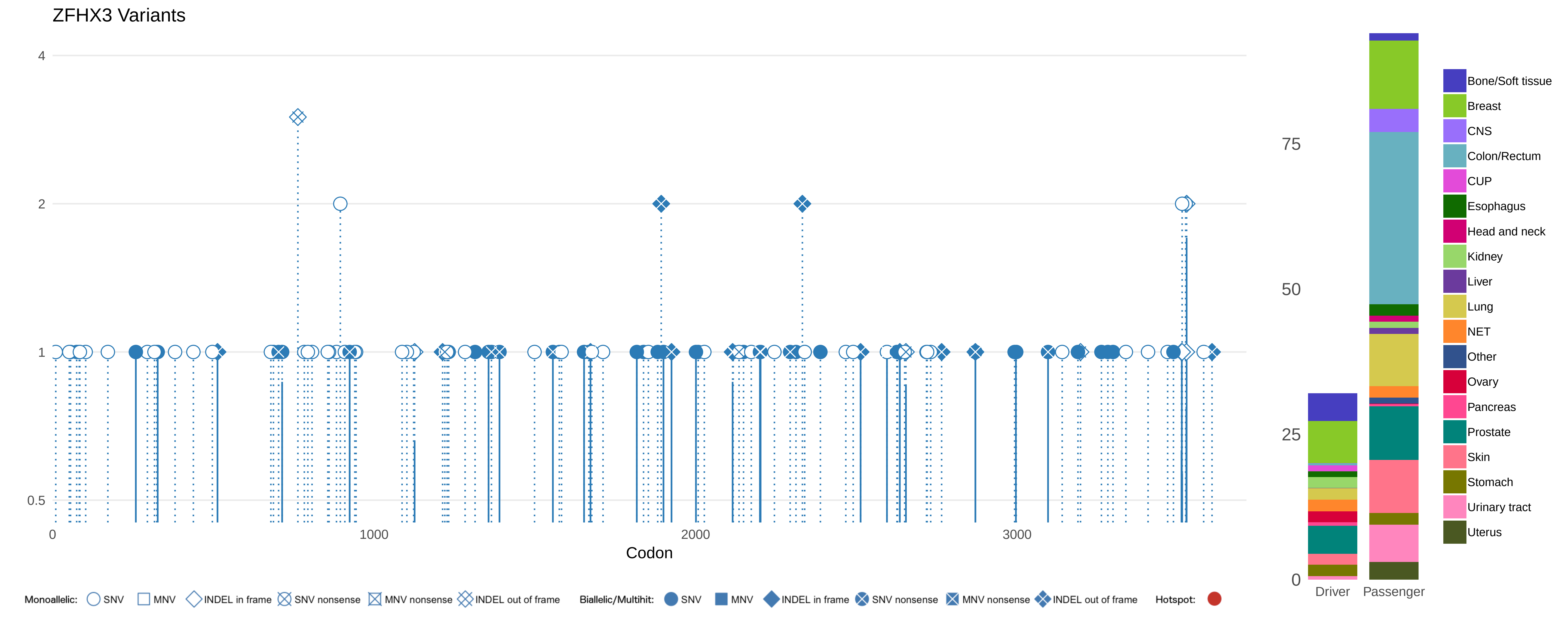

ZFHX3 Variants
4
2
1
0.5
0
1000
2000
3000
Codon
75
50
25
0
Driver
Passenger
Bone/Soft tissue
Breast
CNS
Colon/Rectum
CUP
Esophagus
Head and neck
Kidney
Liver
Lung
NET
Other
Ovary
Pancreas
Prostate
Skin
Stomach
Urinary tract
Uterus
Monoallelic: SNV MNV INDEL in frame SNV nonsense MNV nonsense INDEL out of frame
Biallelic/Multihit: SNV MNV INDEL in frame SNV nonsense MNV nonsense INDEL out of frame
Hotspot:

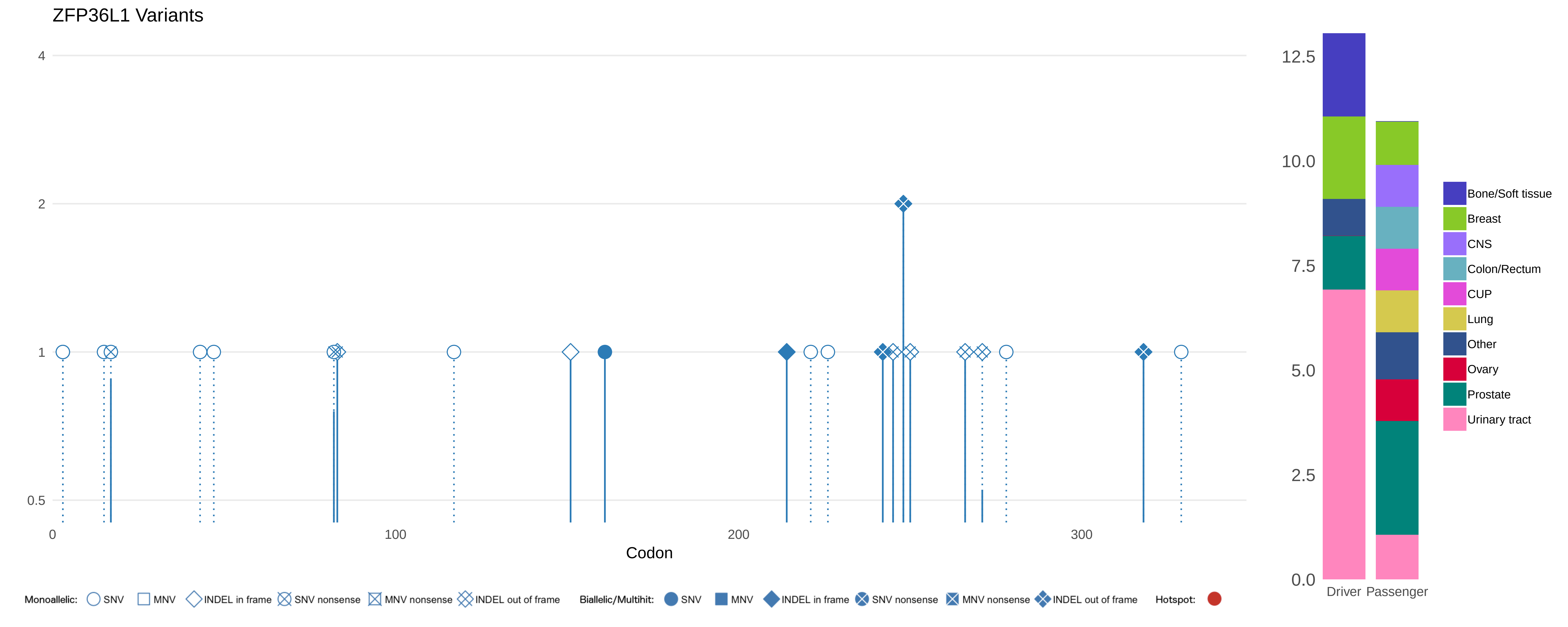

ZFP36L1 Variants
4
2
1
0.5
0
100
200
300
Codon
12.5
10.0
7.5
5.0
2.5
0.0
Driver
Passenger
Bone/Soft tissue
Breast
CNS
Colon/Rectum
CUP
Lung
Other
Ovary
Prostate
Urinary tract
Monoallelic: SNV MNV INDEL in frame SNV nonsense MNV nonsense INDEL out of frame
Biallelic/Multihit: SNV MNV INDEL in frame SNV nonsense MNV nonsense INDEL out of frame
Hotspot:

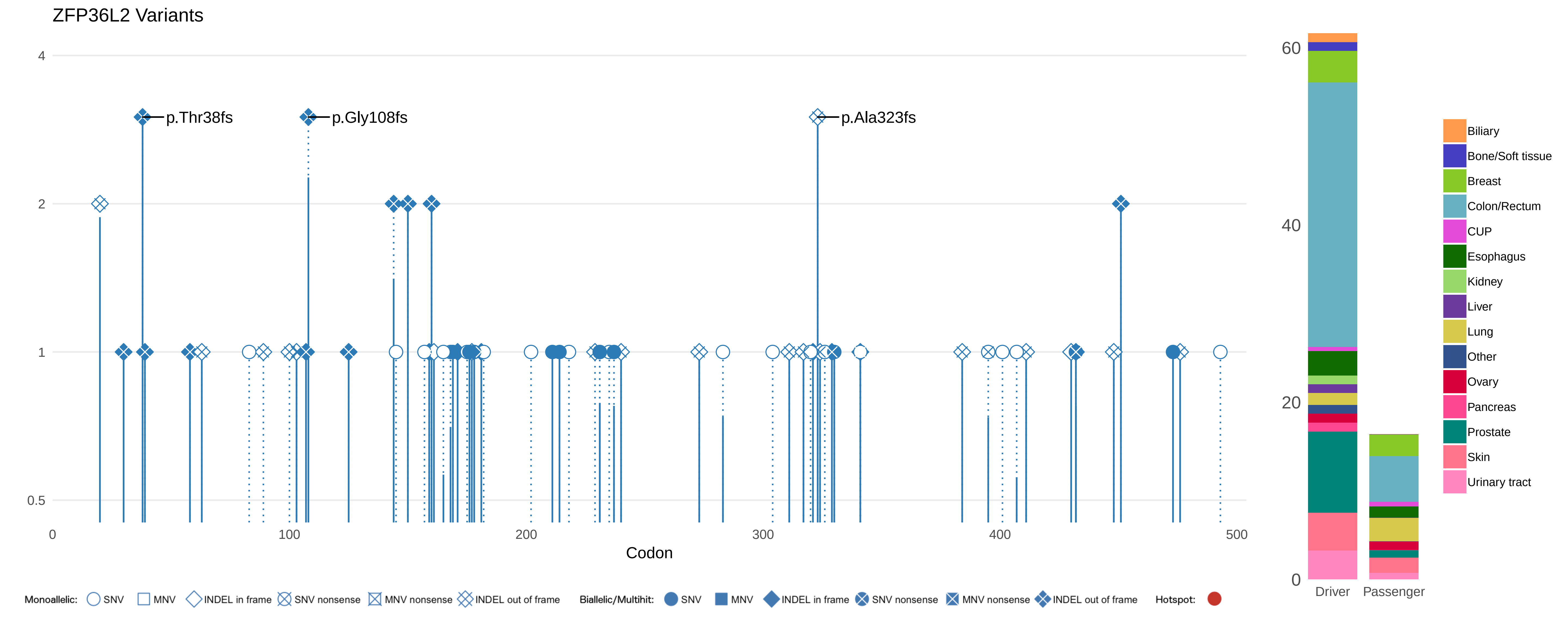

ZFP36L2 Variants
4
2
1
0.5
p.Thr38fs
p.Gly108fs
p.Ala323fs
0
100
200
300
400
500
Codon
60
40
20
0
Driver
Passenger
Biliary
Bone/Soft tissue
Breast
Colon/Rectum
CUP
Esophagus
Kidney
Liver
Lung
Other
Ovary
Pancreas
Prostate
Skin
Urinary tract
Monoallelic: SNV MNV INDEL in frame SNV nonsense MNV nonsense INDEL out of frame
Biallelic/Multihit: SNV MNV INDEL in frame SNV nonsense MNV nonsense INDEL out of frame
Hotspot:

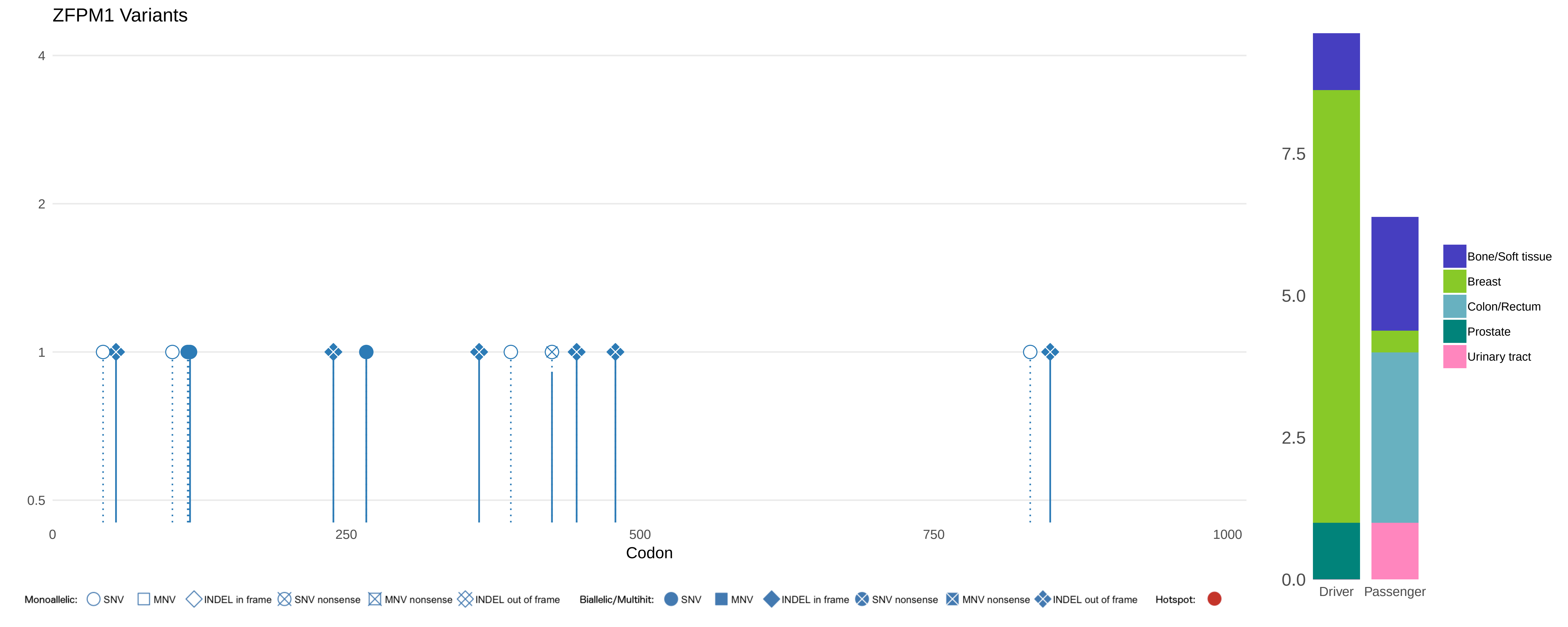

ZFPM1 Variants
4
2
1
0.5
0
250
500
750
1000
Codon
7.5
5.0
2.5
0.0
Driver
Passenger
Bone/Soft tissue
Breast
Colon/Rectum
Prostate
Urinary tract
Monoallelic: SNV MNV INDEL in frame SNV nonsense MNV nonsense INDEL out of frame
Biallelic/Multihit: SNV MNV INDEL in frame SNV nonsense MNV nonsense INDEL out of frame
Hotspot:

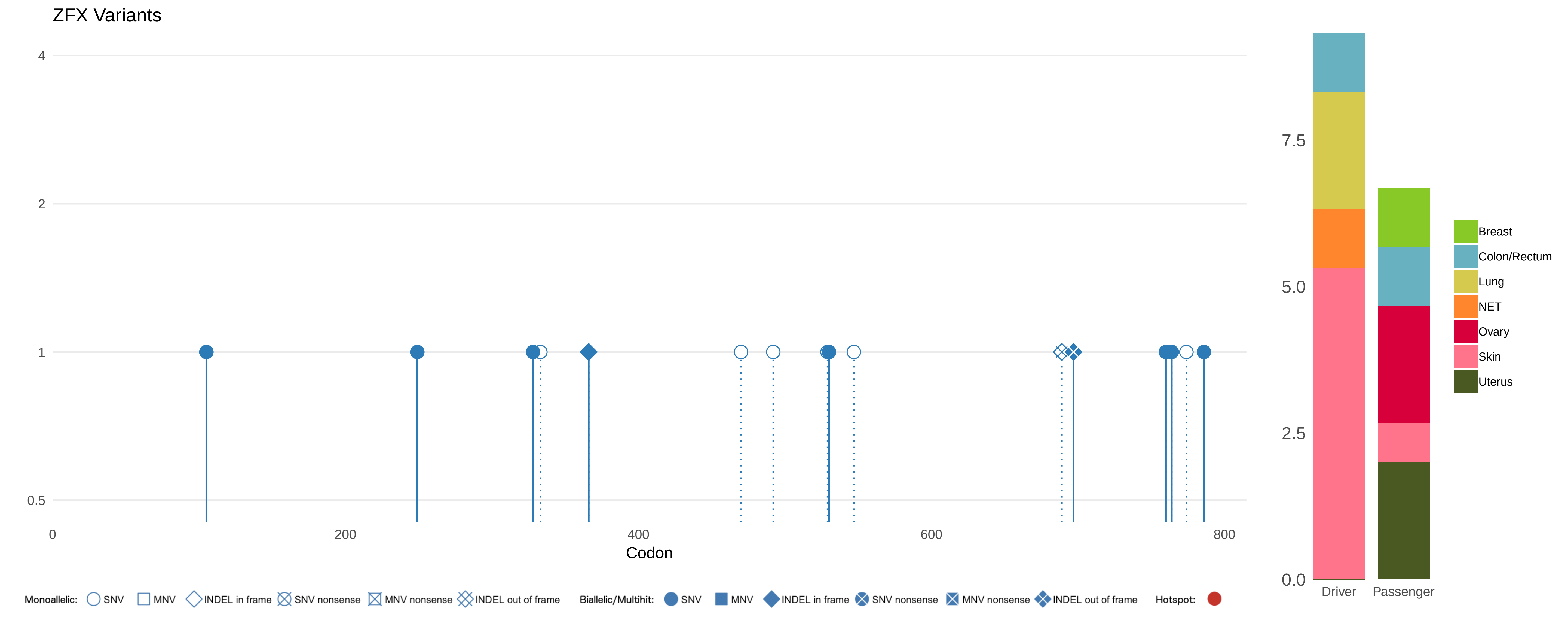
ZFX Variants
4
2
1
0.5
0
200
400
600
800
Codon
7.5
5.0
2.5
0.0
Driver
Passenger
Breast
Colon/Rectum
Lung
NET
Ovary
Skin
Uterus
Monoallelic: SNV MNV INDEL in frame SNV nonsense MNV nonsense INDEL out of frame
Biallelic/Multihit: SNV MNV INDEL in frame SNV nonsense MNV nonsense INDEL out of frame
Hotspot:

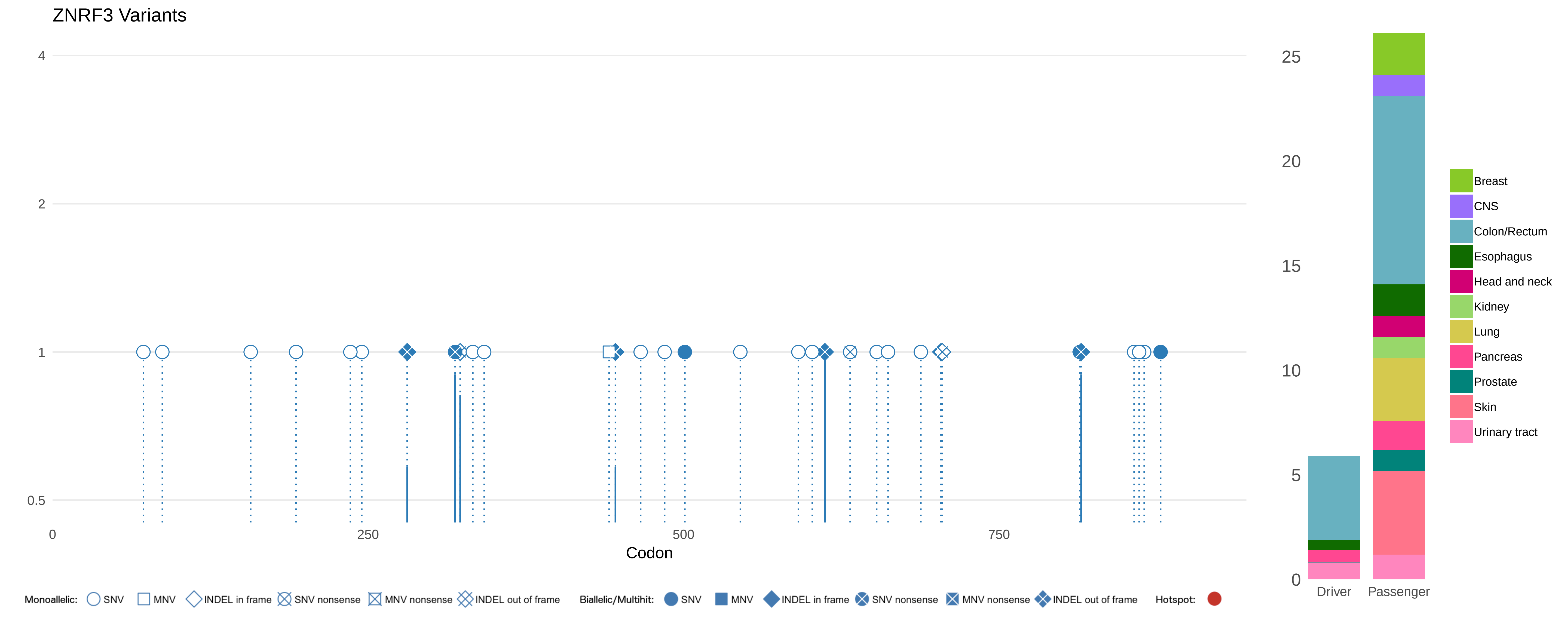

ZNRF3 Variants
4
2
1
0.5
0
250
500
750
Codon
25
20
15
10
5
0
Driver
Passenger
Breast
CNS
Colon/Rectum
Esophagus
Head and neck
Kidney
Lung
Pancreas
Prostate
Skin
Urinary tract
Monoallelic: SNV MNV INDEL in frame SNV nonsense MNV nonsense INDEL out of frame
Biallelic/Multihit: SNV MNV INDEL in frame SNV nonsense MNV nonsense INDEL out of frame
Hotspot:

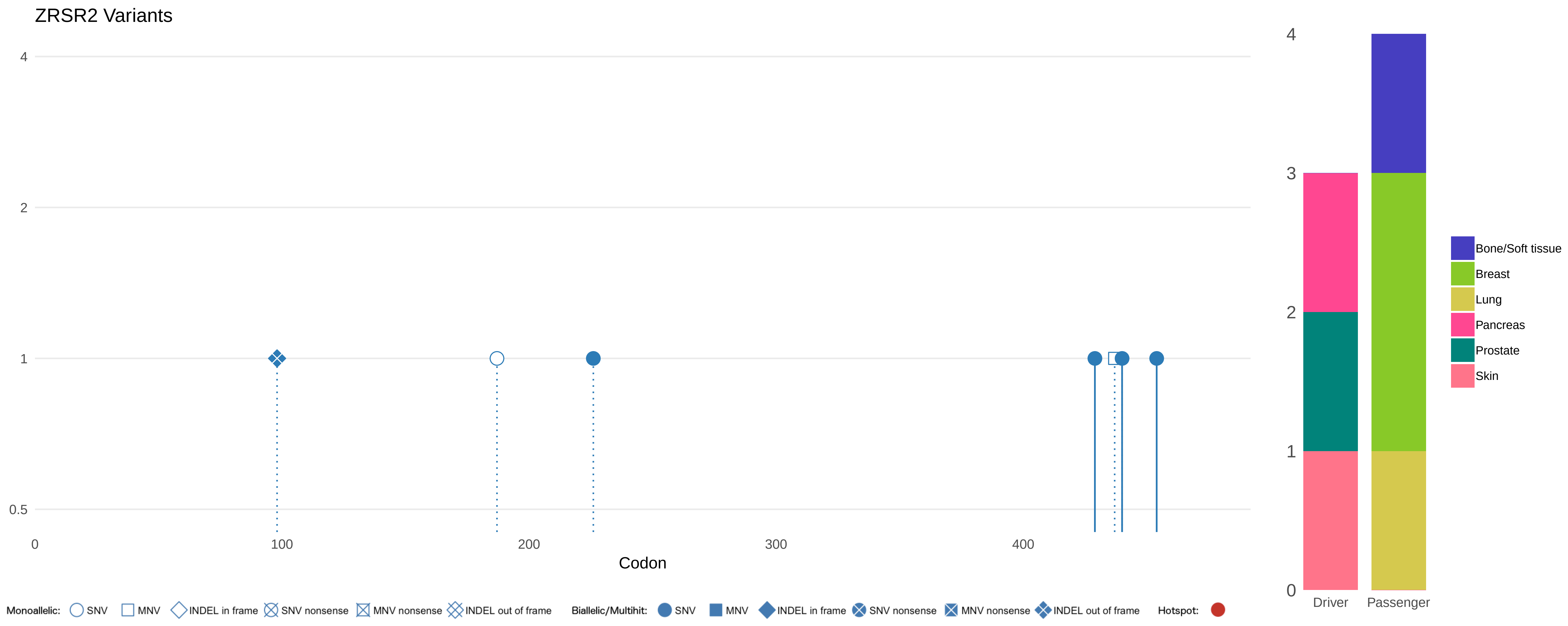

ZRSR2 Variants
4
2
1
0.5
0
100
200
300
400
Codon
Monoallelic: SNV MNV INDEL in frame SNV nonsense MNV nonsense INDEL out of frame
Biallelic/Multihit: SNV MNV INDEL in frame SNV nonsense MNV nonsense INDEL out of frame
Hotspot:
4
3
2
1
0
Driver
Passenger
Bone/Soft tissue
Breast
Lung
Pancreas
Prostate
Skin
