## Supplementary Image 3 for "Pan-cancer whole genome analyses of metastatic solid tumors"

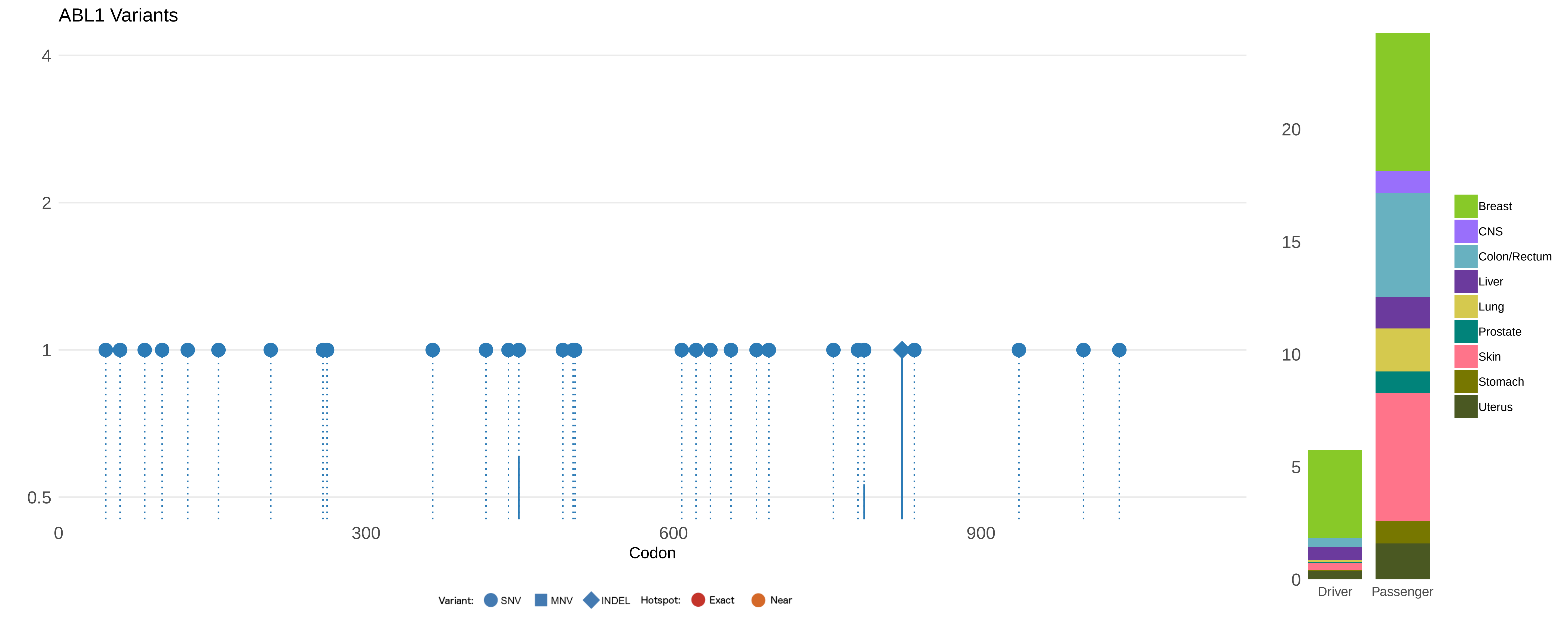

ABL1 Variants
4
2
1
0.5
0
300
600
900
Codon
20
15
10
5
0
Driver
Passenger
Breast
CNS
Colon/Rectum
Liver
Lung
Prostate
Skin
Stomach
Uterus
Variant:
SNV
MNV
INDEL
Hotspot:
Exact
Near

ACVR1 Variants

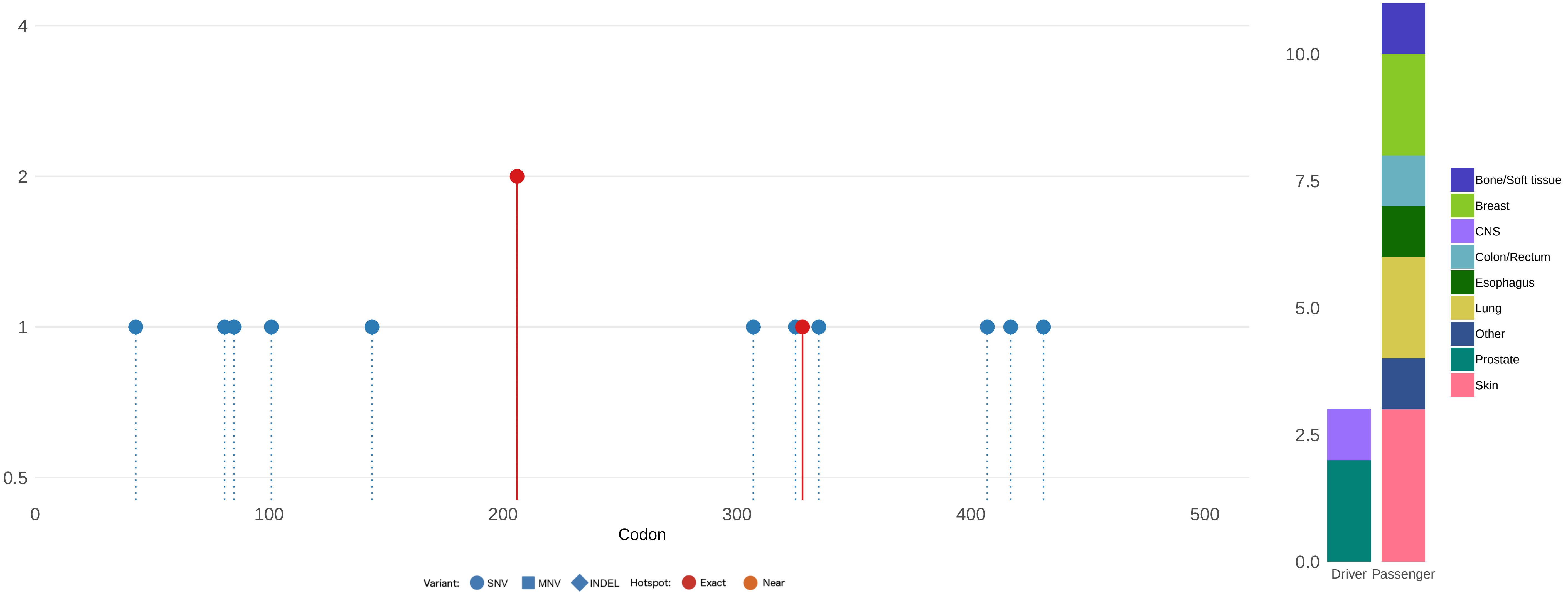

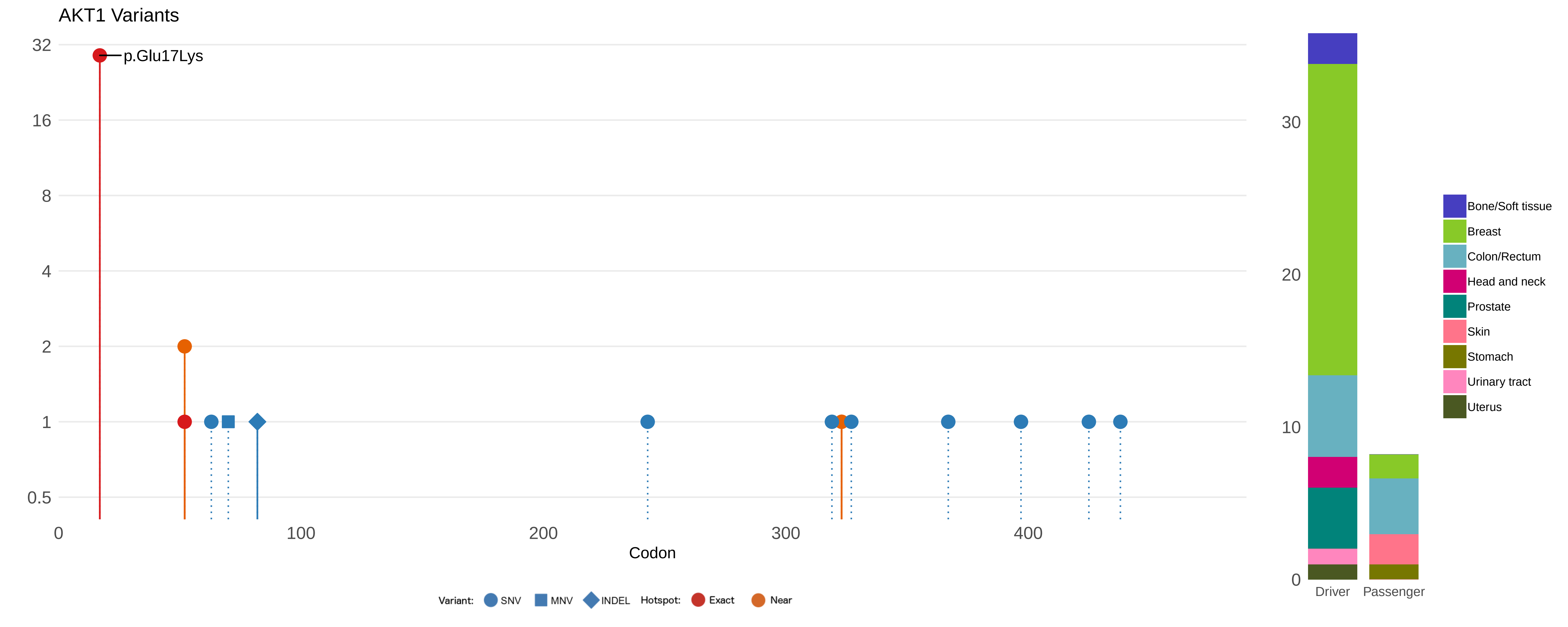

AKT1 Variants
p.Glu17Lys
32
16
8
4
2
1
0.5
0
100
200
300
400
Codon
Variant: SNV MNV INDEL Hotspot: Exact Near
30
20
10
0
Driver
Passenger
Bone/Soft tissue
Breast
Colon/Rectum
Head and neck
Prostate
Skin
Stomach
Urinary tract
Uterus

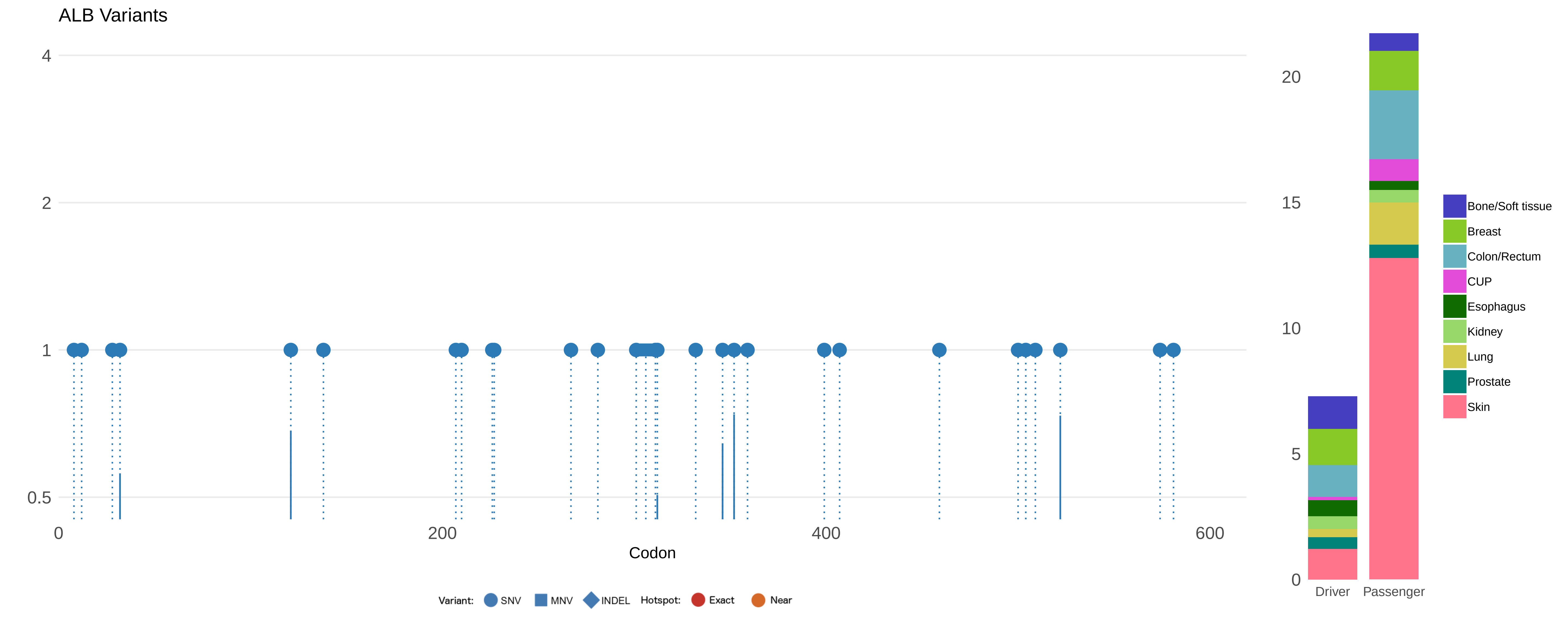

ALB Variants
4
2
1
0.5
0
200
400
600
Codon
Variant: SNV MNV INDEL Hotspot: Exact Near
20
15
10
5
0
Driver
Passenger
Bone/Soft tissue
Breast
Colon/Rectum
CUP
Esophagus
Kidney
Lung
Prostate
Skin

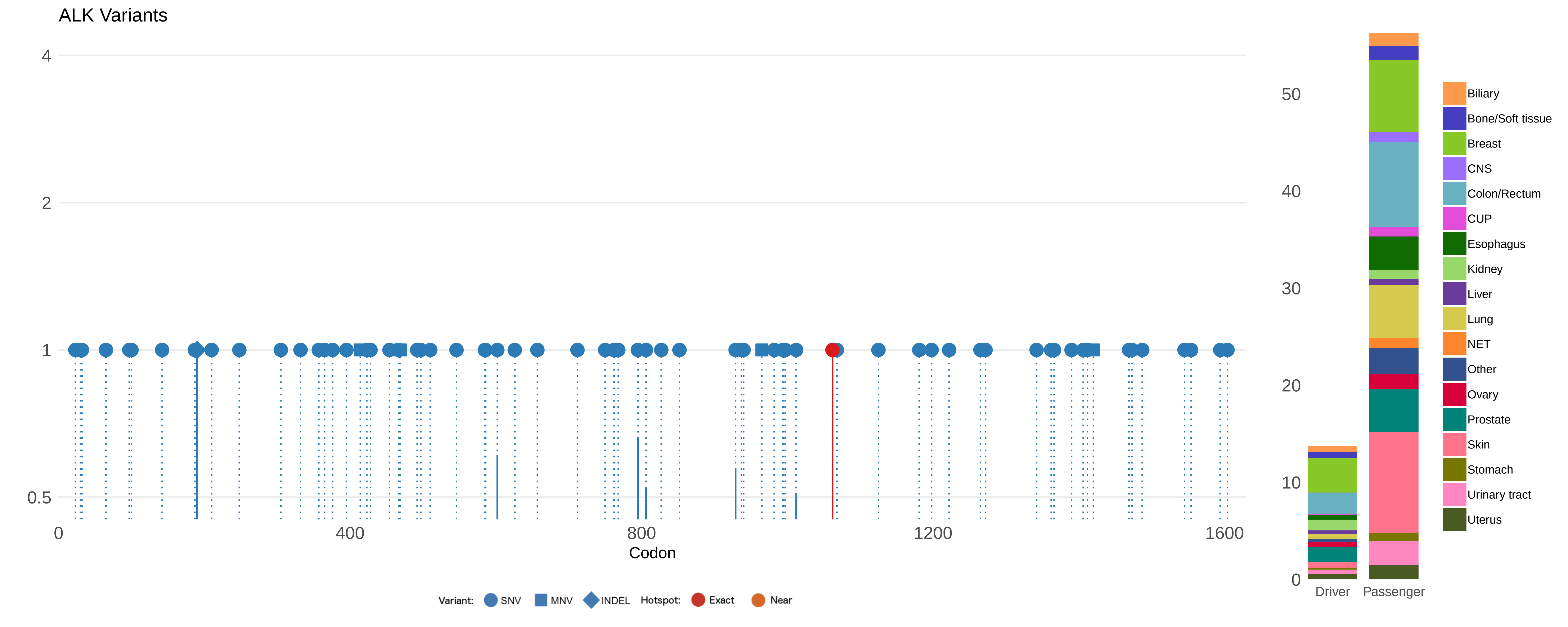

ALK Variants
4
2
1
0.5
0
400
800
1200
1600
Codon
Variant: SNV MNV INDEL Hotspot: Exact Near
50
40
30
20
10
0
Driver
Passenger
Biliary
Bone/Soft tissue
Breast
CNS
Colon/Rectum
CUP
Esophagus
Kidney
Liver
Lung
NET
Other
Ovary
Prostate
Skin
Stomach
Urinary tract
Uterus

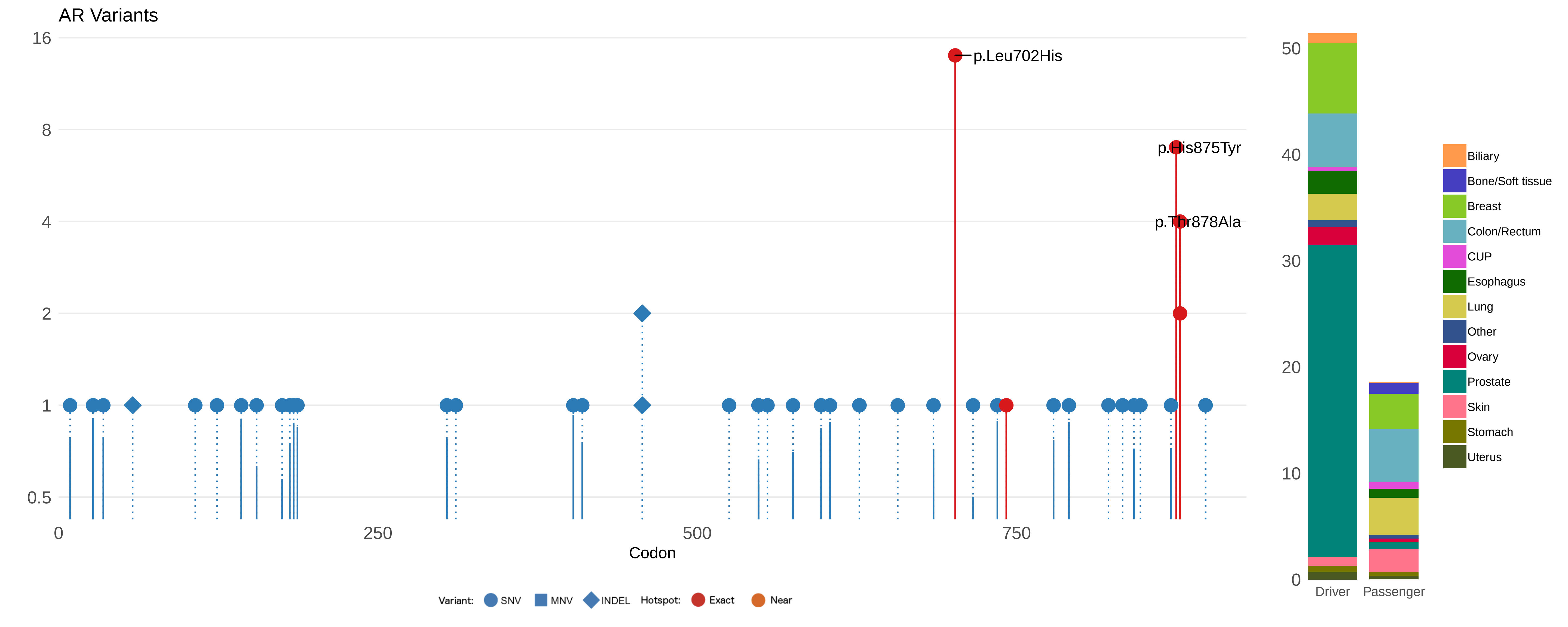
AR Variants
16
8
4
2
1
0.5
p.Leu702His
p.His875Tyr
p.Thr878Ala
0
250
500
750
Codon
Variant: SNV MNV INDEL Hotspot: Exact Near
50
40
30
20
10
0
Driver
Passenger
Biliary
Bone/Soft tissue
Breast
Colon/Rectum
CUP
Esophagus
Lung
Other
Ovary
Prostate
Skin
Stomach
Uterus

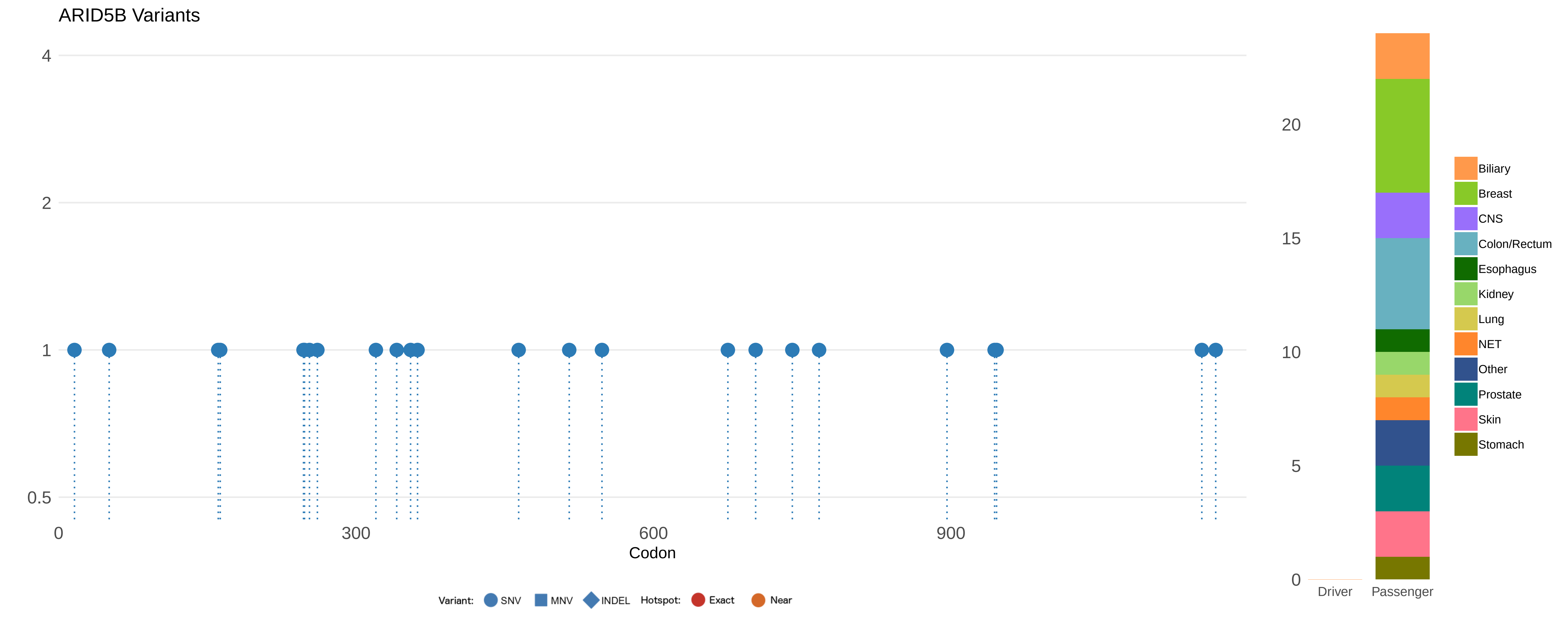
ARID5B Variants
4
2
1
0.5
0
300
600
900
Codon
20
15
10
5
0
Driver
Passenger
Biliary
Breast
CNS
Colon/Rectum
Esophagus
Kidney
Lung
NET
Other
Prostate
Skin
Stomach
Variant: SNV MNV INDEL Hotspot: Exact Near

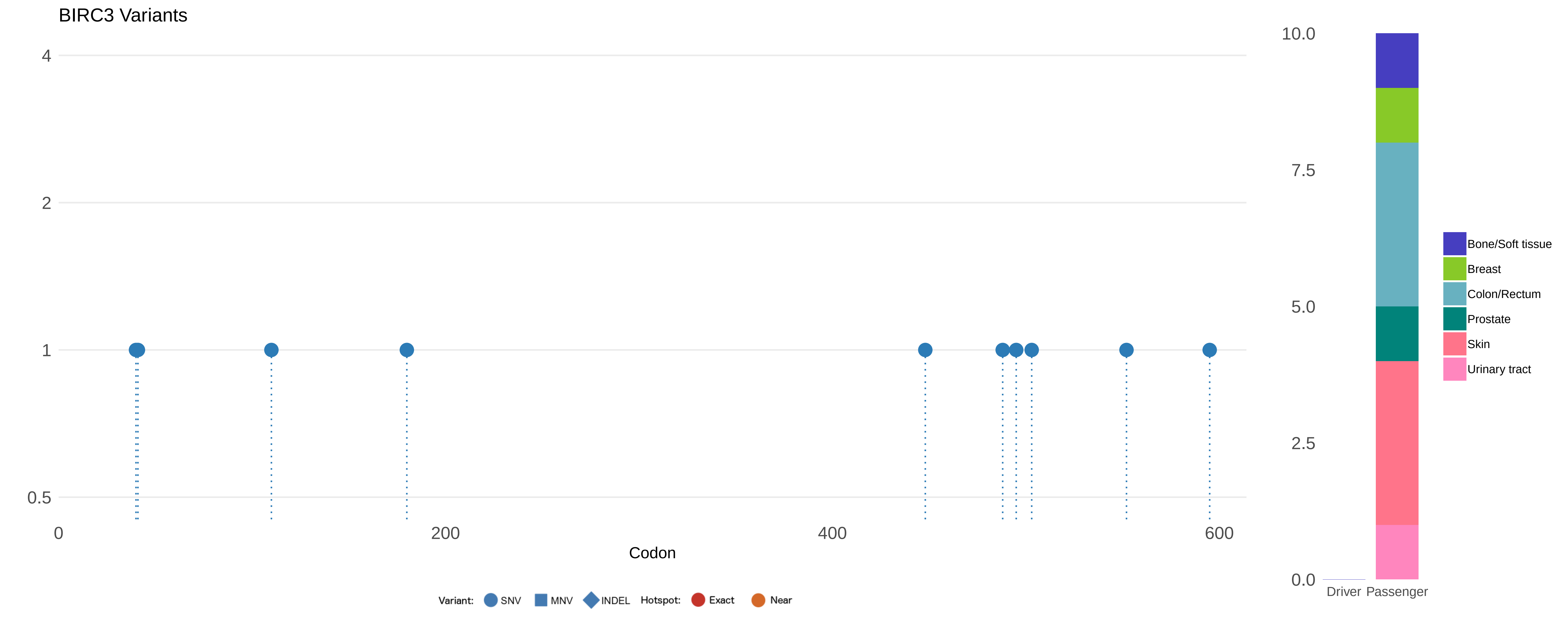

BIRC3 Variants
4
2
1
0.5
0
200
400
600
Codon
10.0
7.5
5.0
2.5
0.0
Driver
Passenger
Bone/Soft tissue
Breast
Colon/Rectum
Prostate
Skin
Urinary tract
Variant:
SNV
MNV
INDEL
Hotspot:
Exact
Near

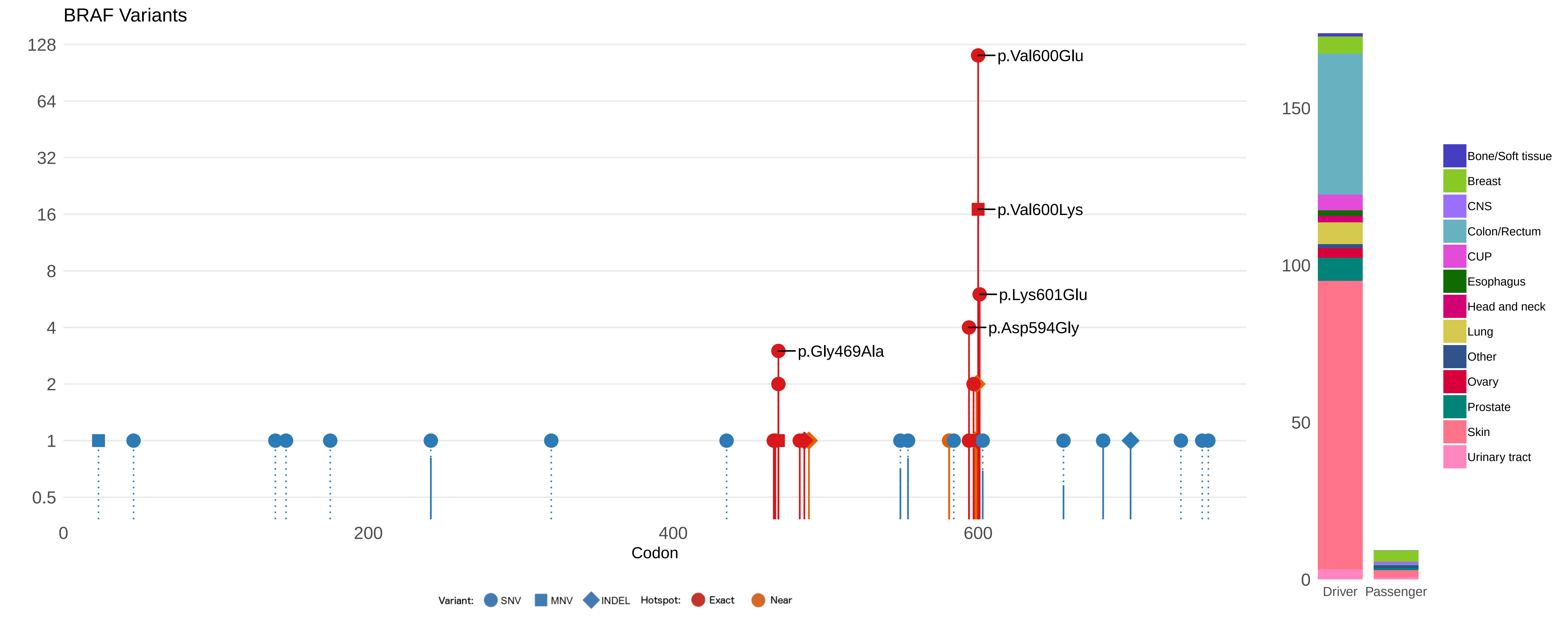

BRAF Variants
128
64
32
16
8
4
2
1
0.5
p.Val600Glu
p.Val600Lys
p.Lys601Glu
p.Asp594Gly
p.Gly469Ala
0
200
400
600
Codon
Variant: SNV MNV INDEL Hotspot: Exact Near
150
100
50
0
Driver
Passenger
Bone/Soft tissue
Breast
CNS
Colon/Rectum
CUP
Esophagus
Head and neck
Lung
Other
Ovary
Prostate
Skin
Urinary tract

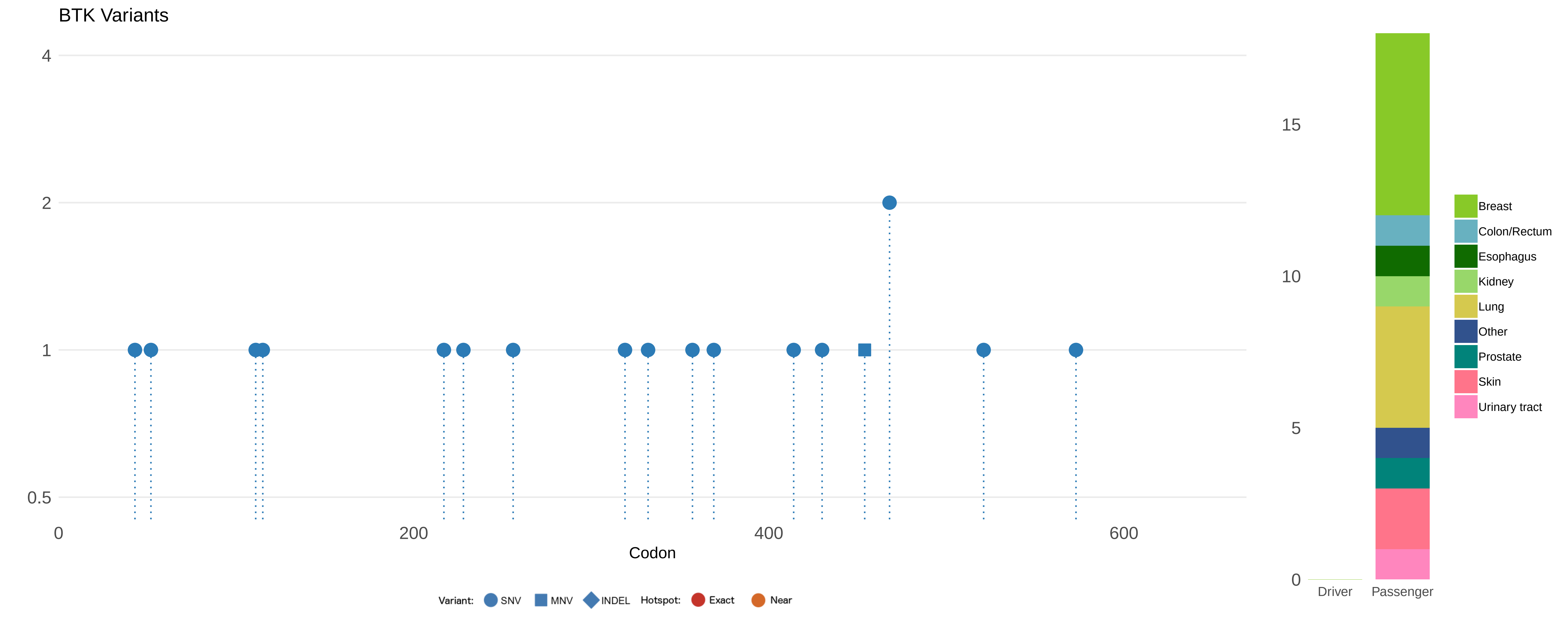
BTK Variants
4
2
1
0.5
0
200
400
600
Codon
Variant:
SNV
MNV
INDEL
Hotspot:
Exact
Near
15
10
5
0
Driver
Passenger
Breast
Colon/Rectum
Esophagus
Kidney
Lung
Other
Prostate
Skin
Urinary tract

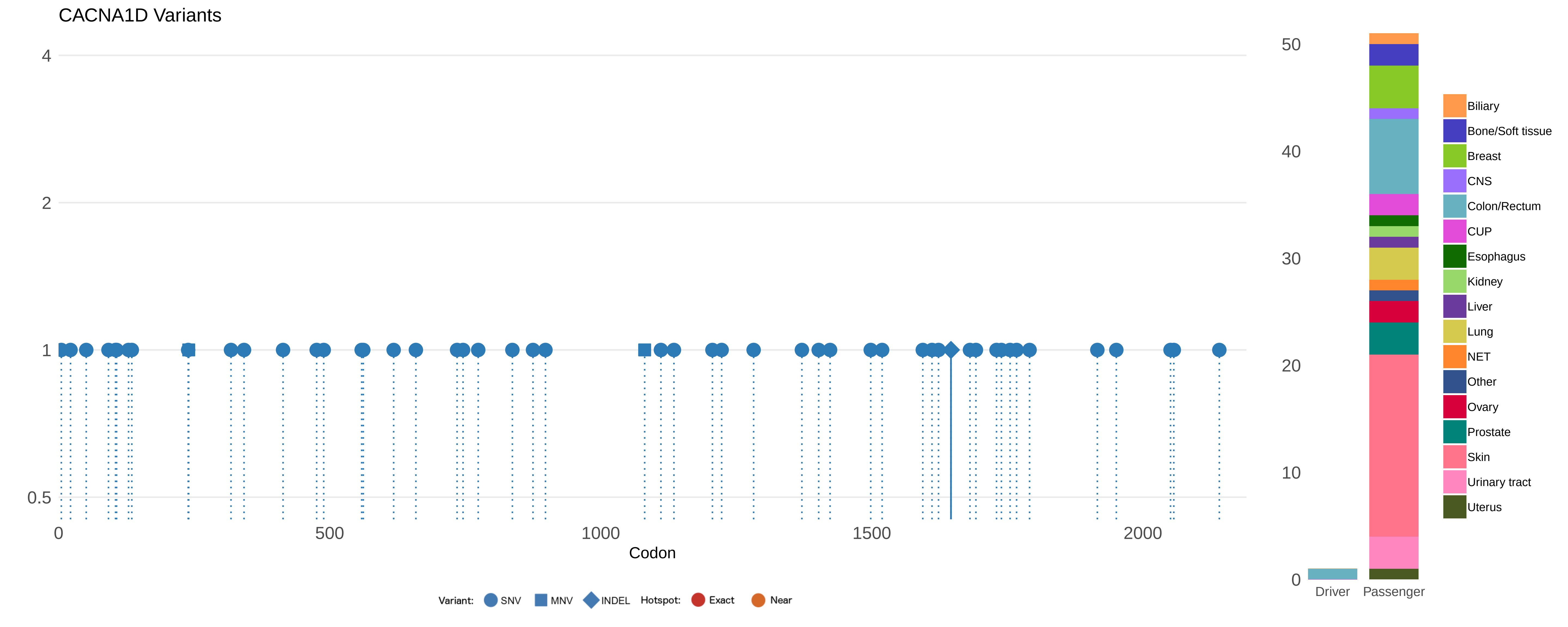

CACNA1D Variants
4
2
1
0.5
0
500
1000
1500
2000
Codon
Variant: SNV MNV INDEL Hotspot: Exact Near
50
40
30
20
10
0
Driver
Passenger
Biliary
Bone/Soft tissue
Breast
CNS
Colon/Rectum
CUP
Esophagus
Kidney
Liver
Lung
NET
Other
Ovary
Prostate
Skin
Urinary tract
Uterus

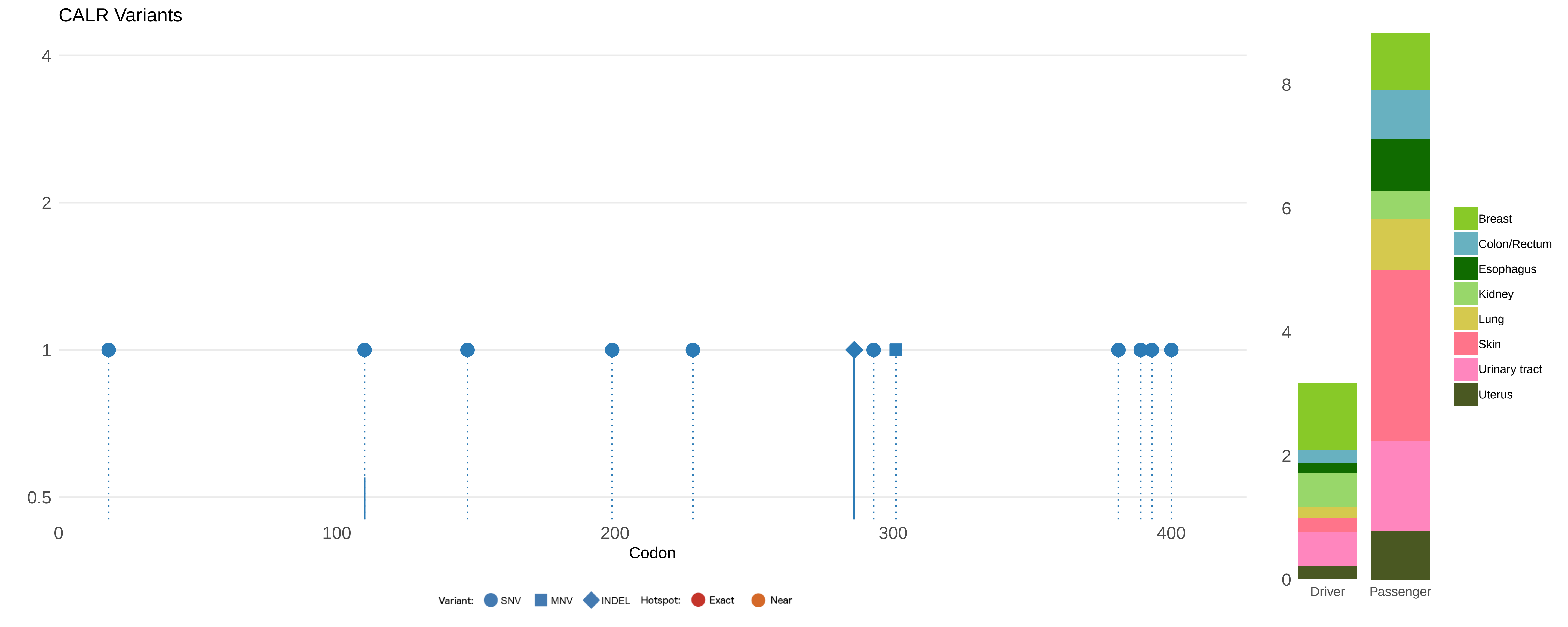

CALR Variants
4
2
1
0.5
0
100
200
300
400
Codon
Variant: SNV MNV INDEL Hotspot: Exact Near
8
6
4
2
0
Driver
Passenger
Breast
Colon/Rectum
Esophagus
Kidney
Lung
Skin
Urinary tract
Uterus

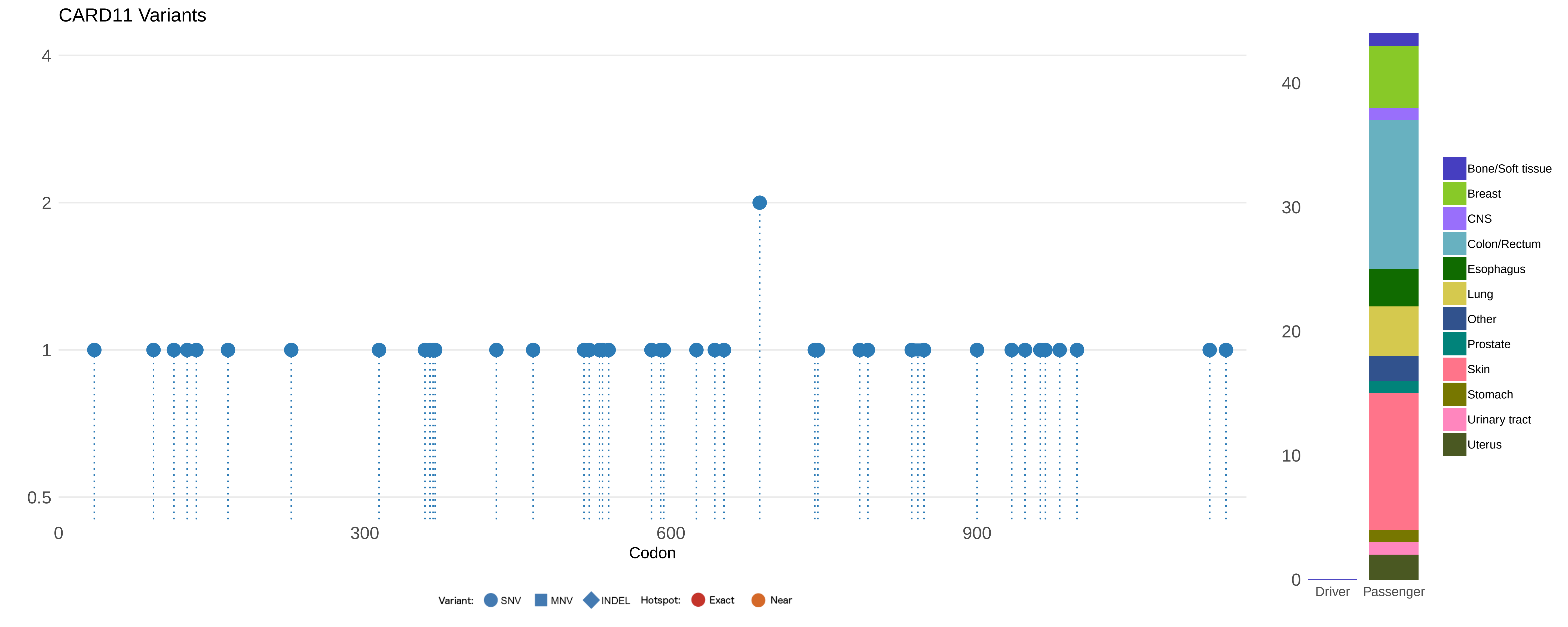

CARD11 Variants
4
2
1
0.5
0
300
600
900
Codon
40
30
20
10
0
Driver
Passenger
Bone/Soft tissue
Breast
CNS
Colon/Rectum
Esophagus
Lung
Other
Prostate
Skin
Stomach
Urinary tract
Uterus
Variant:
SNV
MNV
INDEL
Hotspot:
Exact
Near

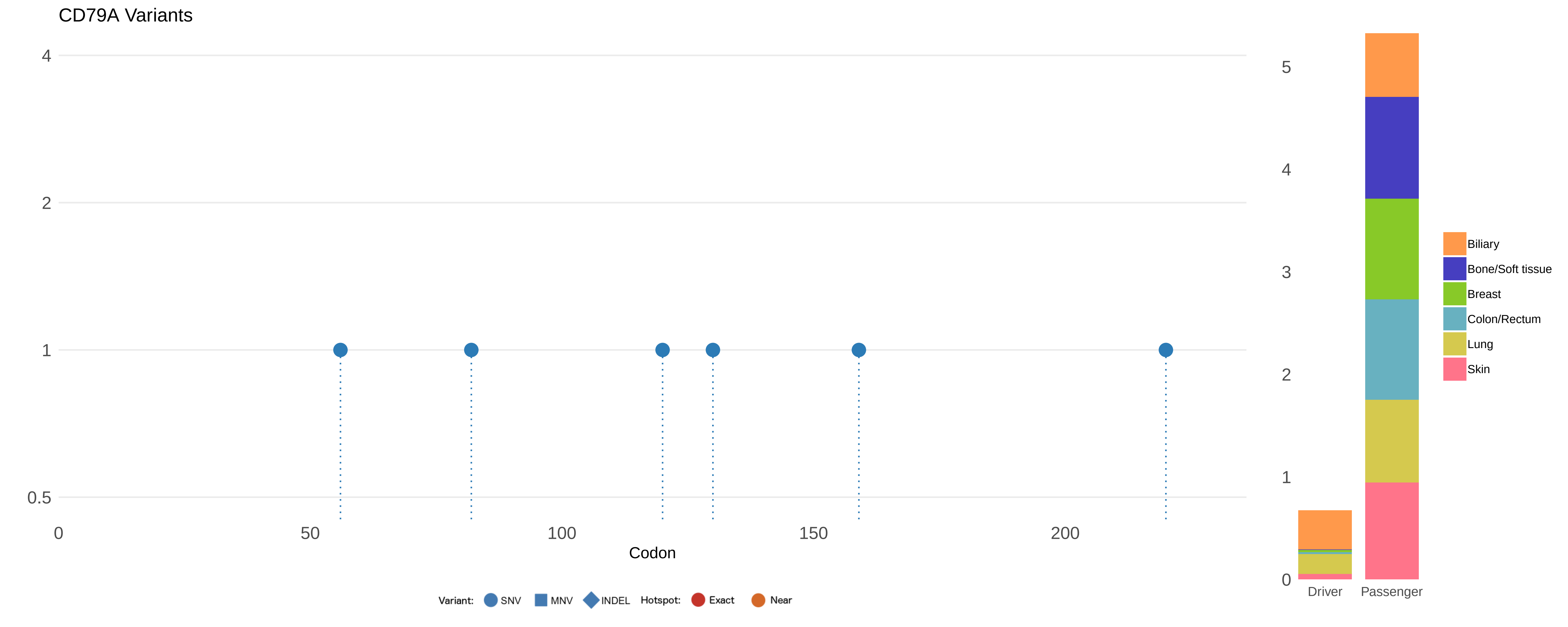
CD79A Variants
4
2
1
0.5
0
50
100
150
200
Codon
Variant: SNV MNV INDEL Hotspot: Exact Near
5
4
3
2
1
0
Driver
Passenger
Biliary
Bone/Soft tissue
Breast
Colon/Rectum
Lung
Skin

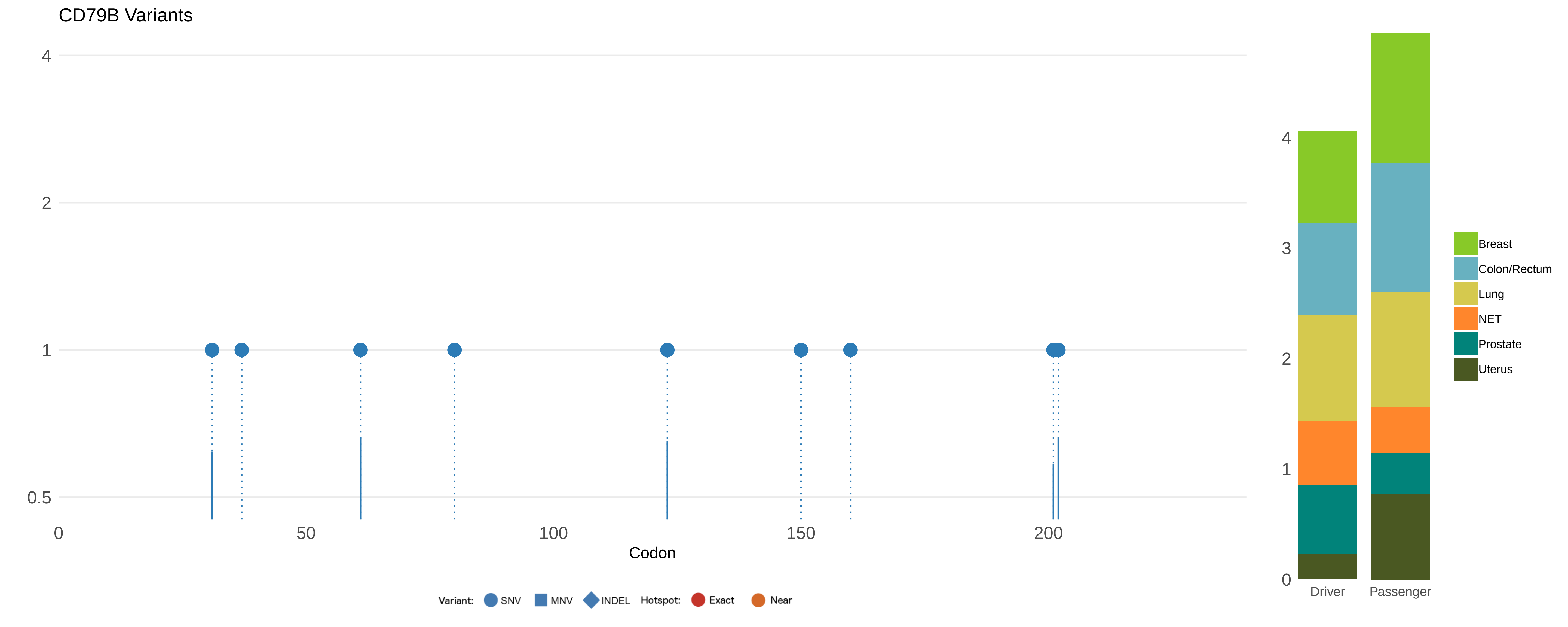

CD79B Variants
4
2
1
0.5
0
50
100
150
200
Codon
Variant: SNV MNV INDEL Hotspot: Exact Near
4
3
2
1
0
Driver
Passenger
Breast
Colon/Rectum
Lung
NET
Prostate
Uterus

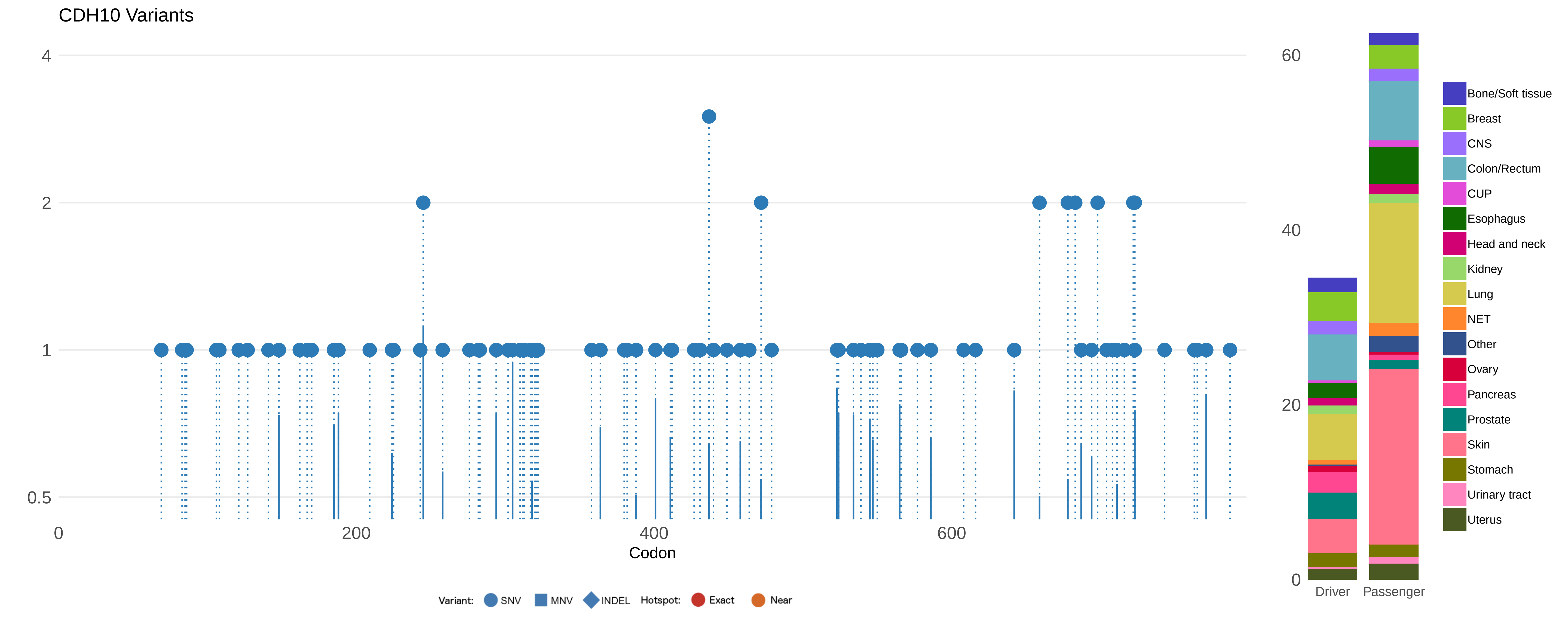

CDH10 Variants
4
2
1
0.5
0
200
400
600
Codon
Variant:
SNV
MNV
INDEL
Hotspot:
Exact
Near
60
40
20
0
Driver
Passenger
Bone/Soft tissue
Breast
CNS
Colon/Rectum
CUP
Esophagus
Head and neck
Kidney
Lung
NET
Other
Ovary
Pancreas
Prostate
Skin
Stomach
Urinary tract
Uterus

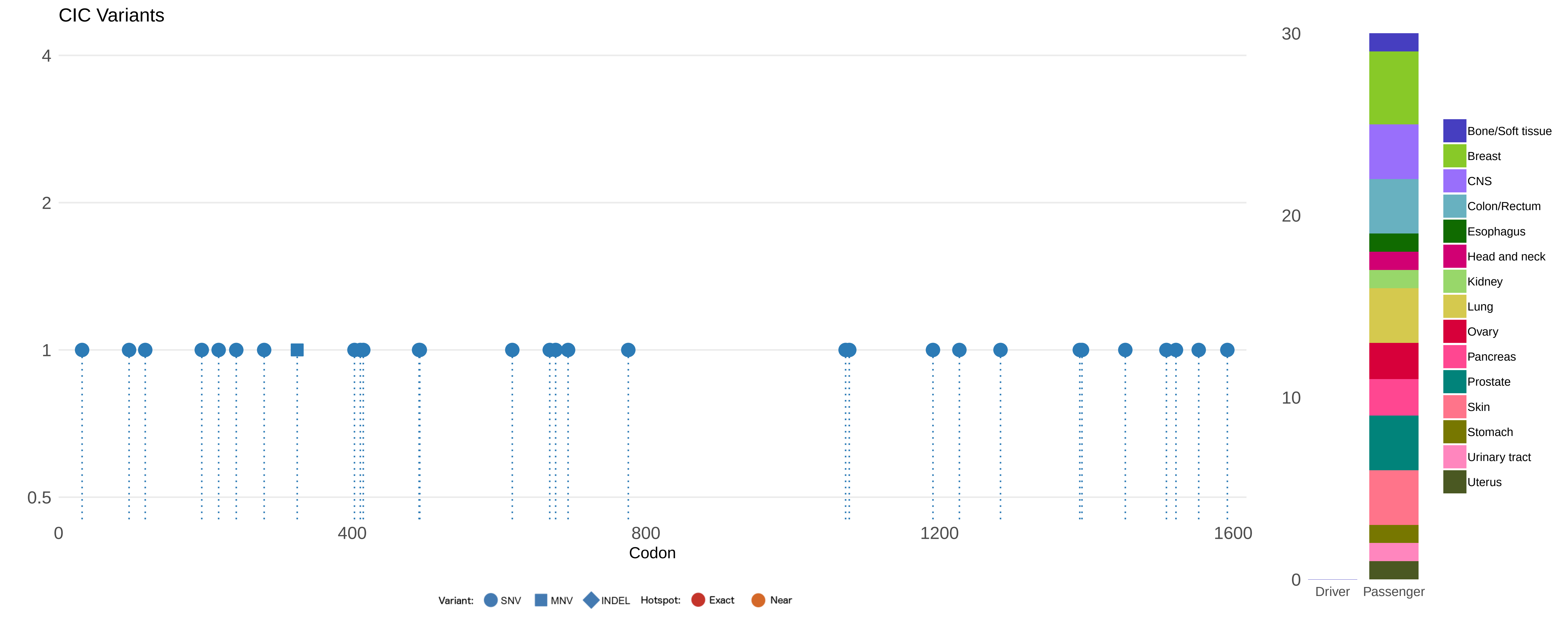
CIC Variants
4
2
1
0.5
0
400
800
1200
1600
Codon
Variant: SNV MNV INDEL Hotspot: Exact Near
30
20
10
0
Driver
Passenger
Bone/Soft tissue
Breast
CNS
Colon/Rectum
Esophagus
Head and neck
Kidney
Lung
Ovary
Pancreas
Prostate
Skin
Stomach
Urinary tract
Uterus

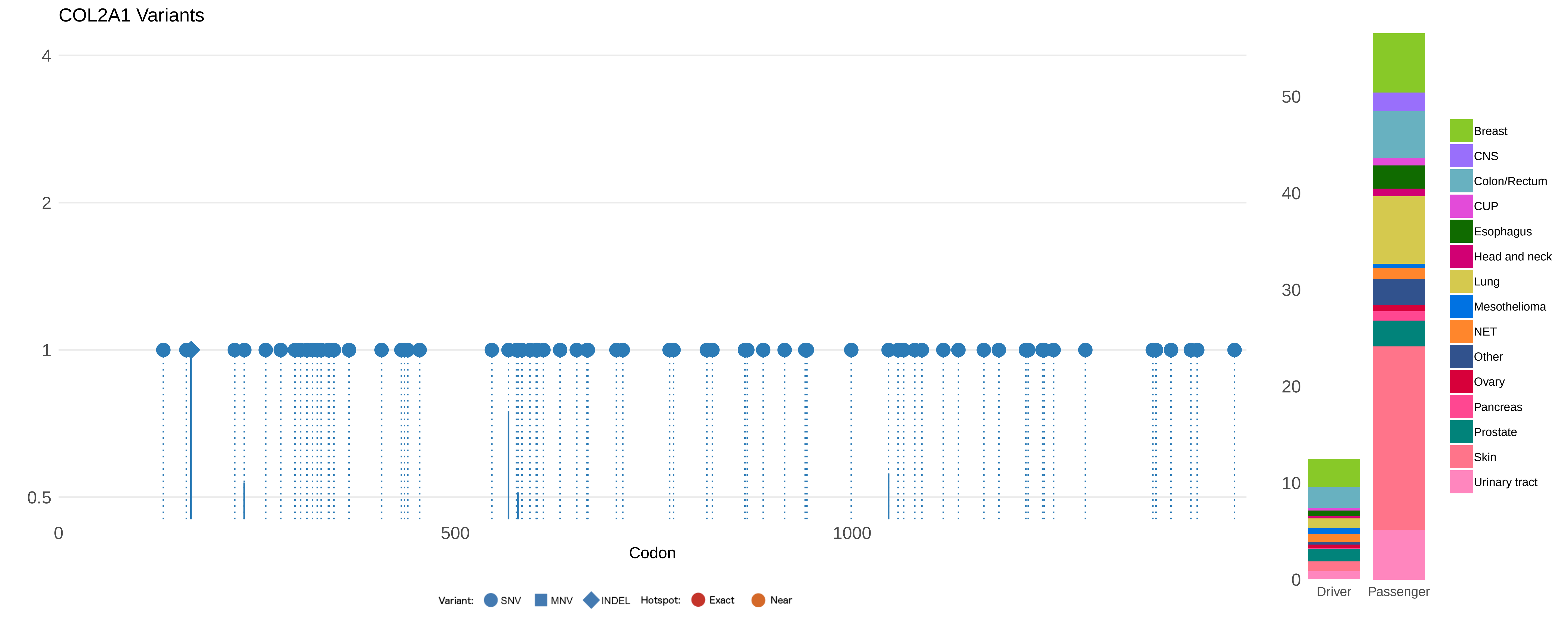

COL2A1 Variants
4
2
1
0.5
0
500
1000
Codon
Variant: SNV MNV INDEL Hotspot: Exact Near
50
40
30
20
10
0
Driver
Passenger
Breast
CNS
Colon/Rectum
CUP
Esophagus
Head and neck
Lung
Mesothelioma
NET
Other
Ovary
Pancreas
Prostate
Skin
Urinary tract

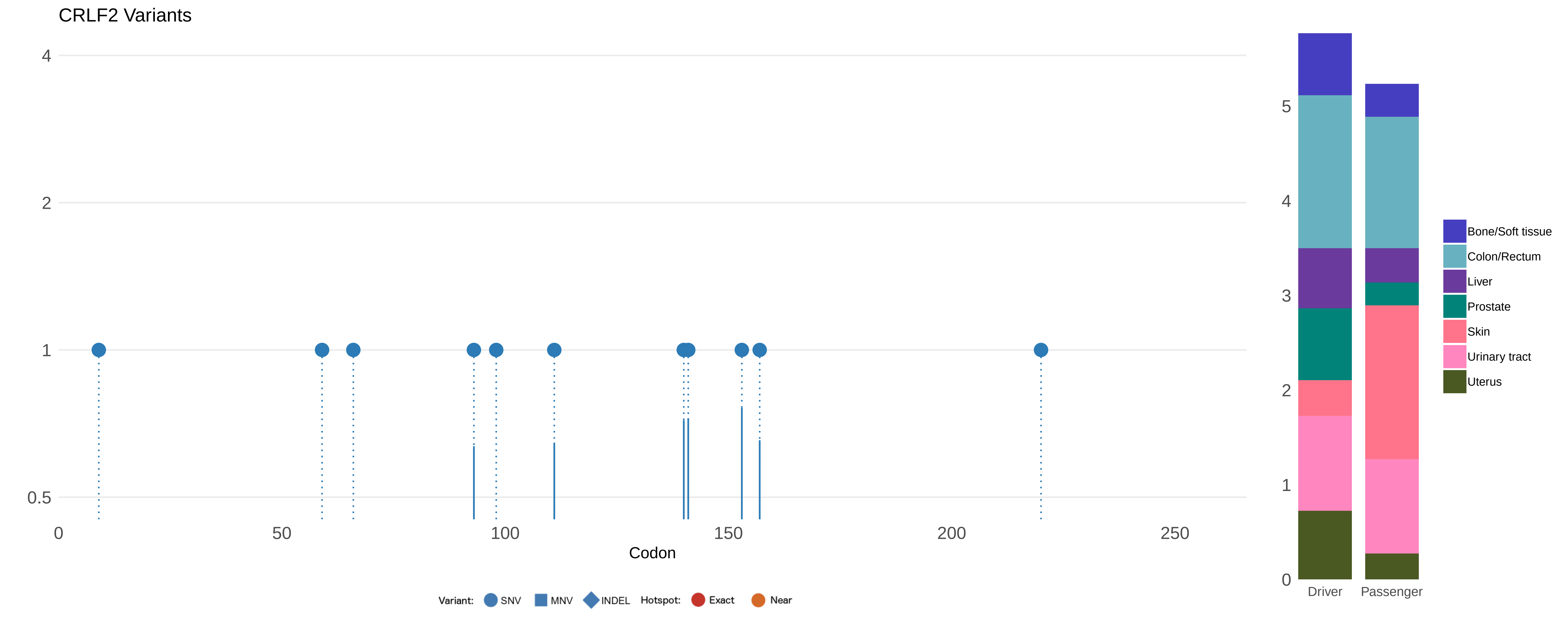

CRLF2 Variants
4
2
1
0.5
0
50
100
150
200
250
Codon
Variant: SNV MNV INDEL Hotspot: Exact Near
5
4
3
2
1
0
Driver
Passenger
Bone/Soft tissue
Colon/Rectum
Liver
Prostate
Skin
Urinary tract
Uterus

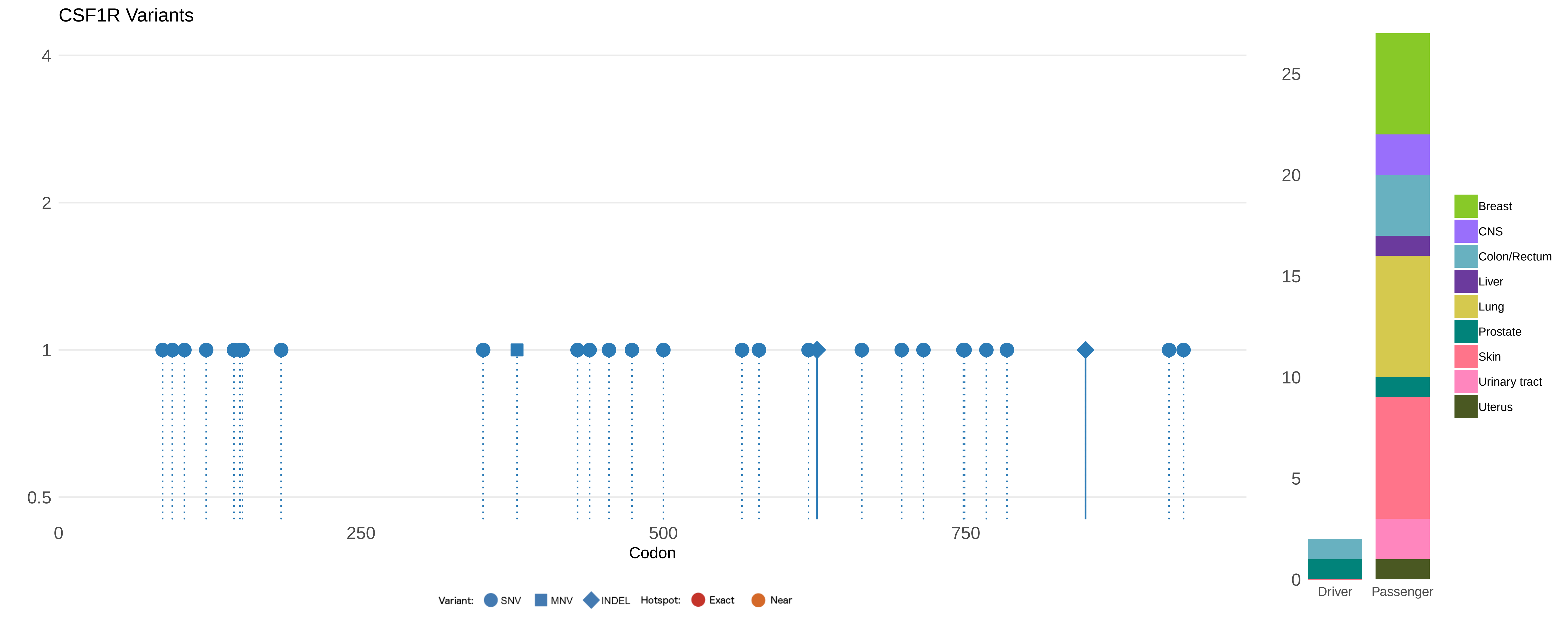
CSF1R Variants
4
2
1
0.5
0
250
500
750
Codon
Variant:
SNV
MNV
INDEL
Hotspot:
Exact
Near
25
20
15
10
5
0
Driver
Passenger
Breast
CNS
Colon/Rectum
Liver
Lung
Prostate
Skin
Urinary tract
Uterus

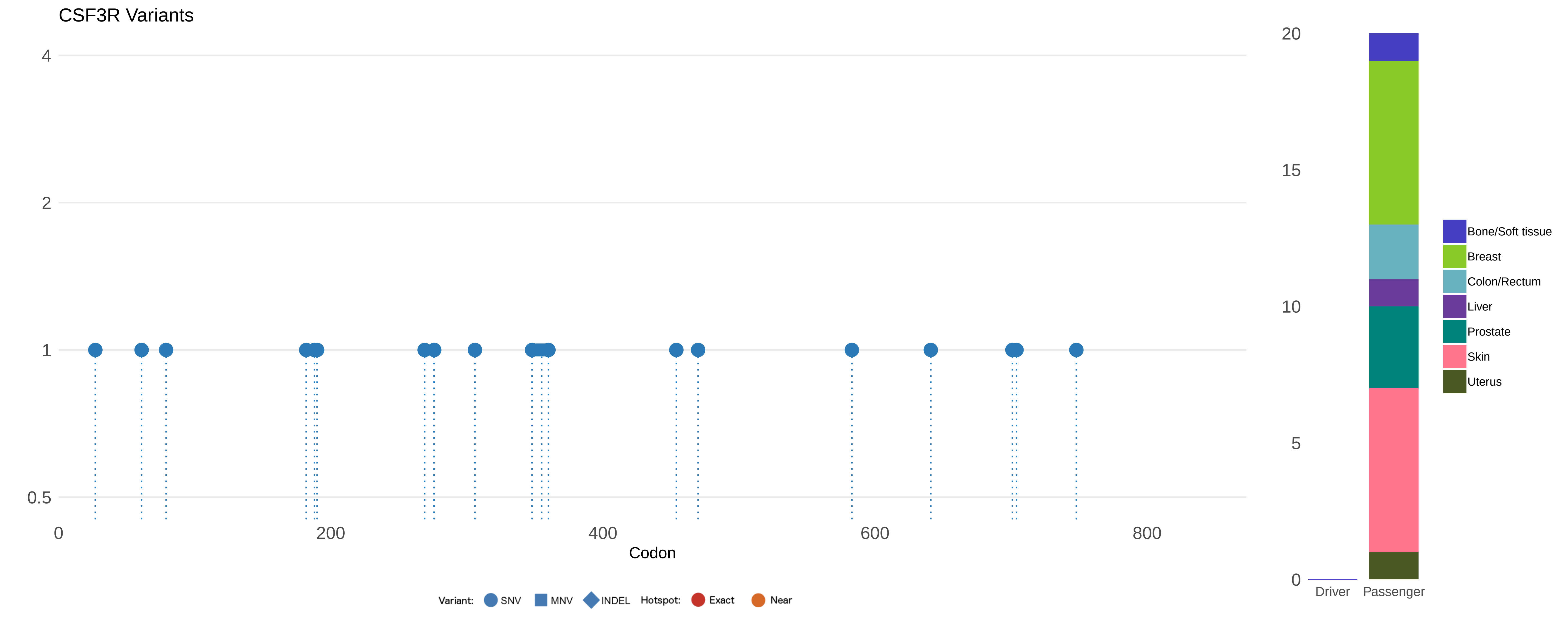
CSF3R Variants
4
2
1
0.5
0
200
400
600
800
Codon
Variant:
SNV
MNV
INDEL
Hotspot:
Exact
Near
20
15
10
5
0
Driver
Passenger
Bone/Soft tissue
Breast
Colon/Rectum
Liver
Prostate
Skin
Uterus

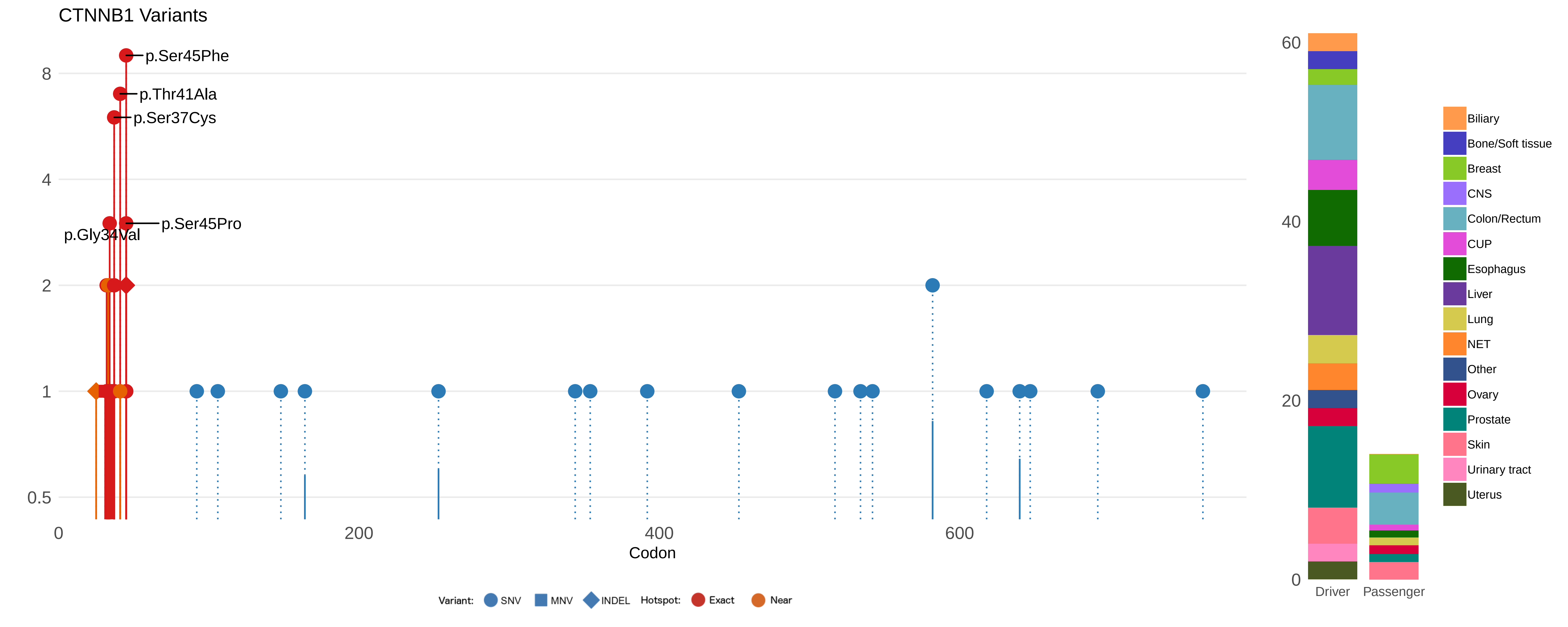
CTNNB1 Variants
p.Ser45Phe
p.Thr41Ala
p.Ser37Cys
p.Ser45Pro
p.Gly34Val
8
4
2
1
0.5
0
200
400
600
Codon
Variant: SNV MNV INDEL Hotspot: Exact Near
60
40
20
0
Driver
Passenger
Biliary
Bone/Soft tissue
Breast
CNS
Colon/Rectum
CUP
Esophagus
Liver
Lung
NET
Other
Ovary
Prostate
Skin
Urinary tract
Uterus

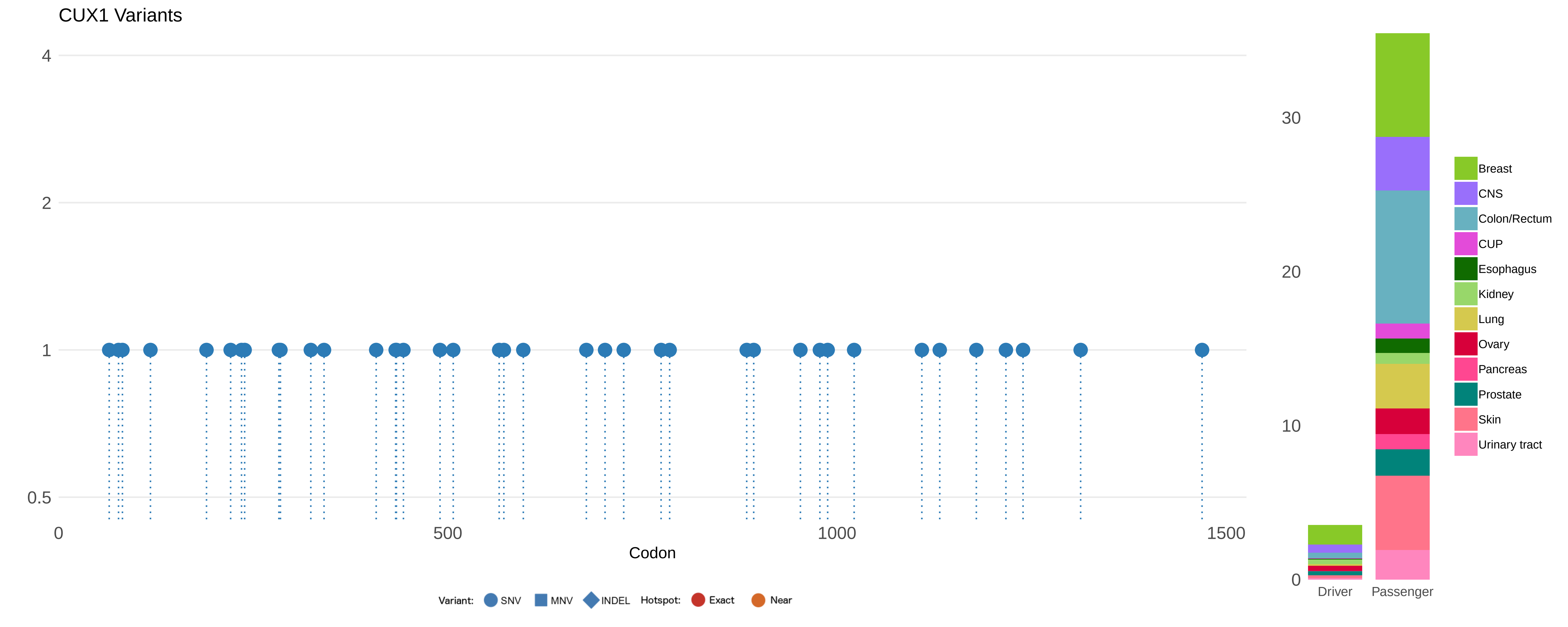
CUX1 Variants
4
2
1
0.5
0
500
1000
1500
Codon
Variant: SNV MNV INDEL Hotspot: Exact Near
30
20
10
0
Driver
Passenger
Breast
CNS
Colon/Rectum
CUP
Esophagus
Kidney
Lung
Ovary
Pancreas
Prostate
Skin
Urinary tract

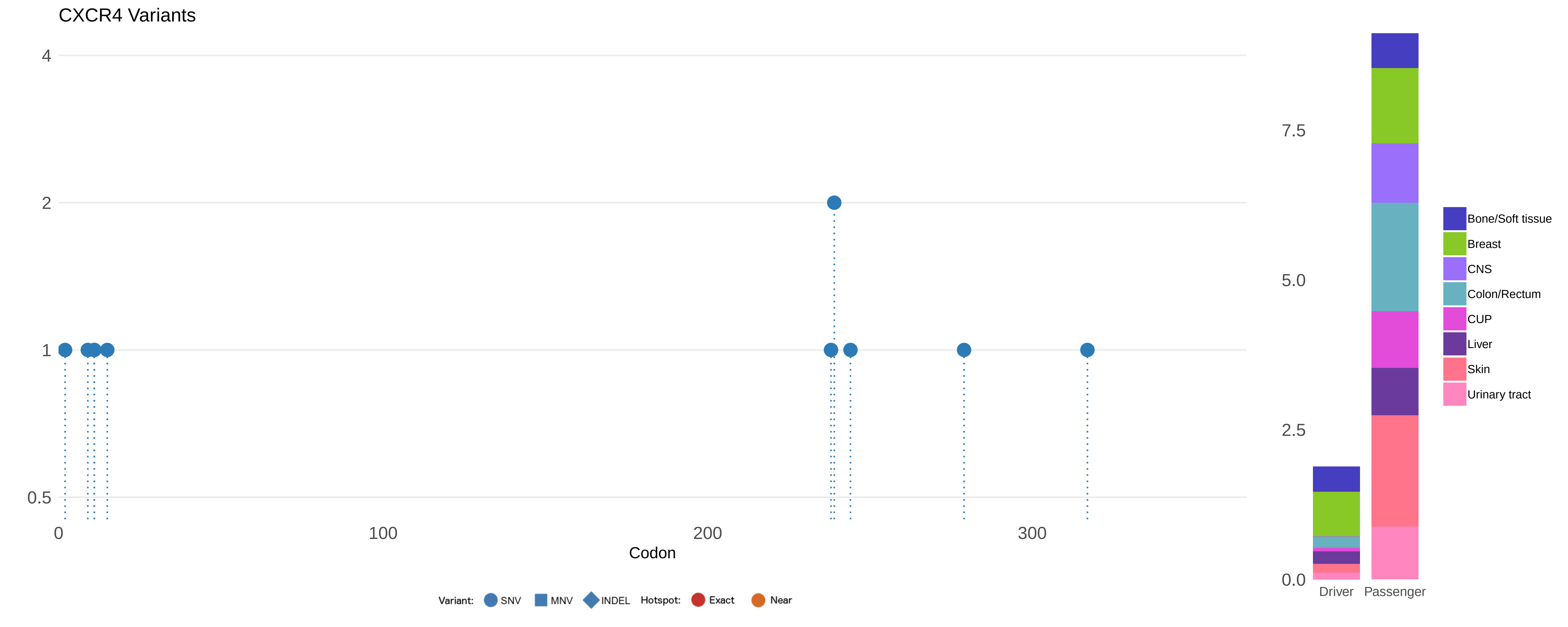

CXCR4 Variants
4
2
1
0.5
0
100
200
300
Codon
Variant: SNV MNV INDEL Hotspot: Exact Near
7.5
5.0
2.5
0.0
Driver
Passenger
Bone/Soft tissue
Breast
CNS
Colon/Rectum
CUP
Liver
Skin
Urinary tract

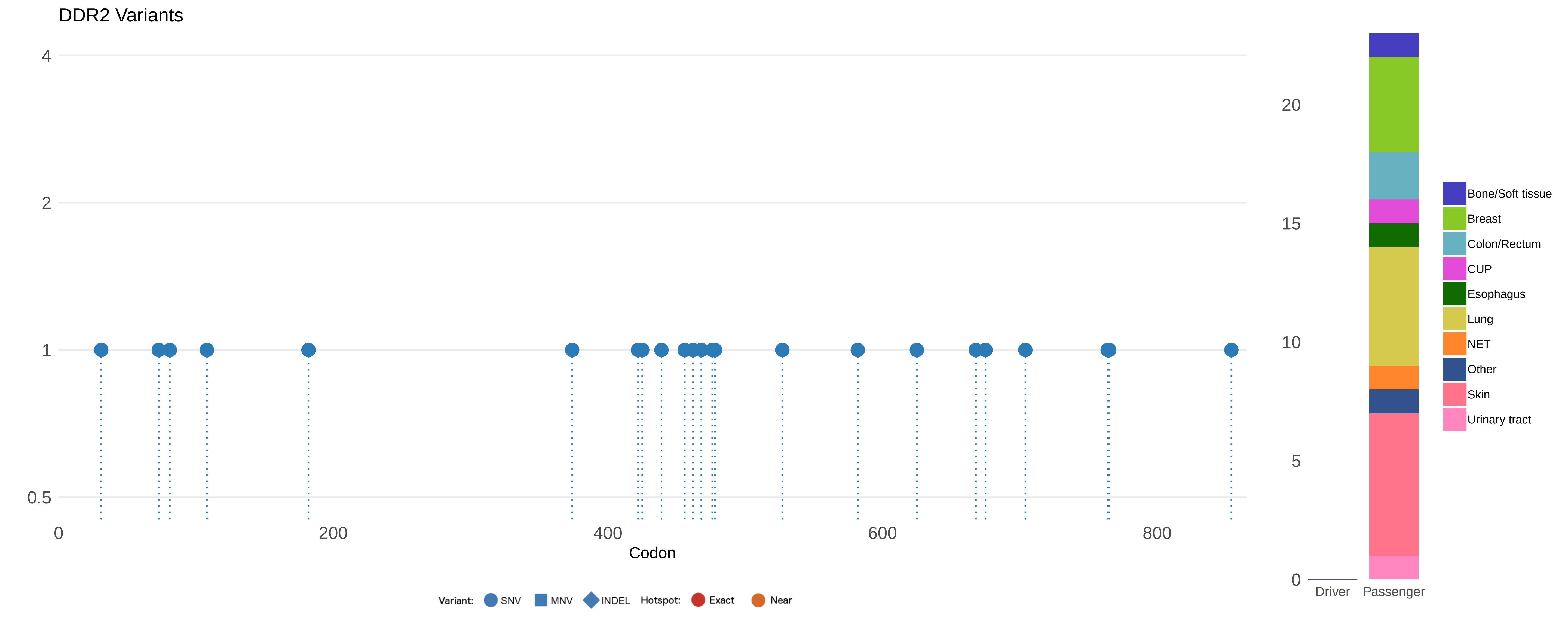
DDR2 Variants
4
2
1
0.5
0
200
400
600
800
Codon
Variant:
SNV
MNV
INDEL
Hotspot:
Exact
Near
20
15
10
5
0
Driver
Passenger
Bone/Soft tissue
Breast
Colon/Rectum
CUP
Esophagus
Lung
NET
Other
Skin
Urinary tract

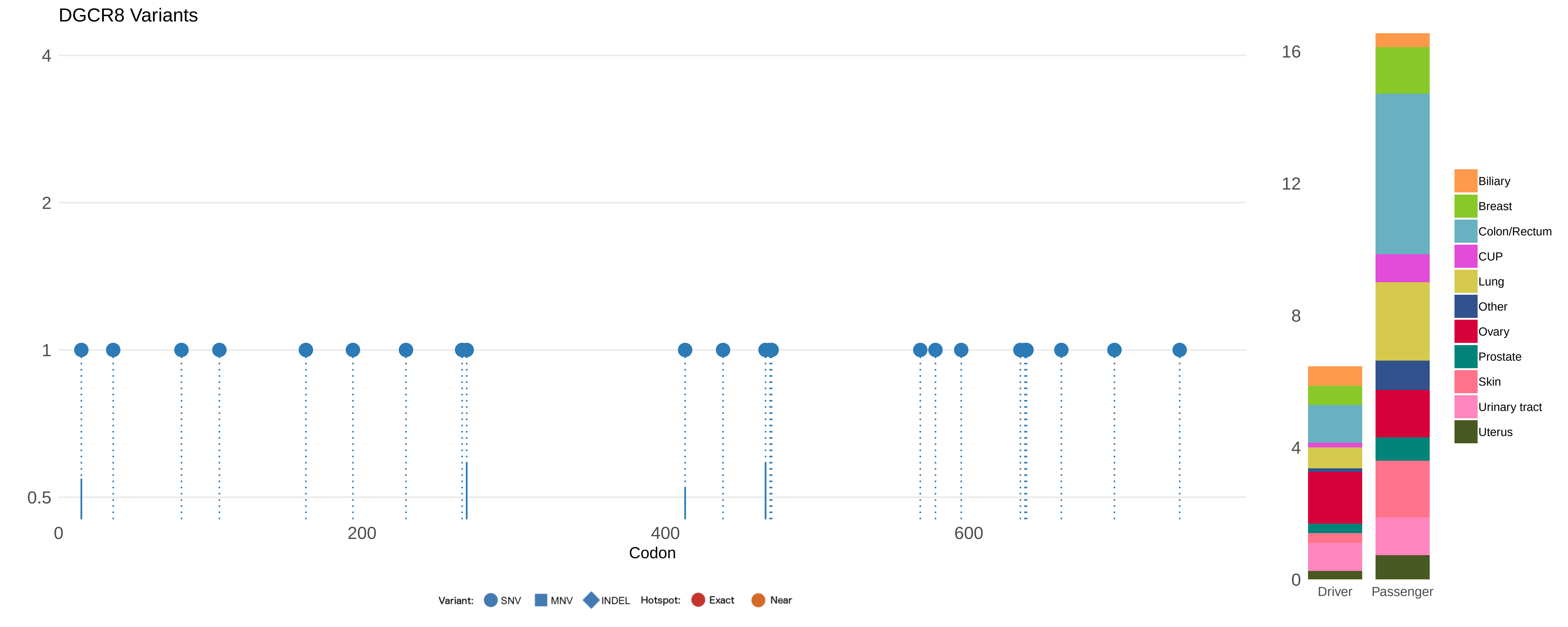

DGCR8 Variants
4
2
1
0.5
0
200
400
600
Codon
Variant:
SNV
MNV
INDEL
Hotspot:
Exact
Near
16
12
8
4
0
Driver
Passenger
Biliary
Breast
Colon/Rectum
CUP
Lung
Other
Ovary
Prostate
Skin
Urinary tract
Uterus

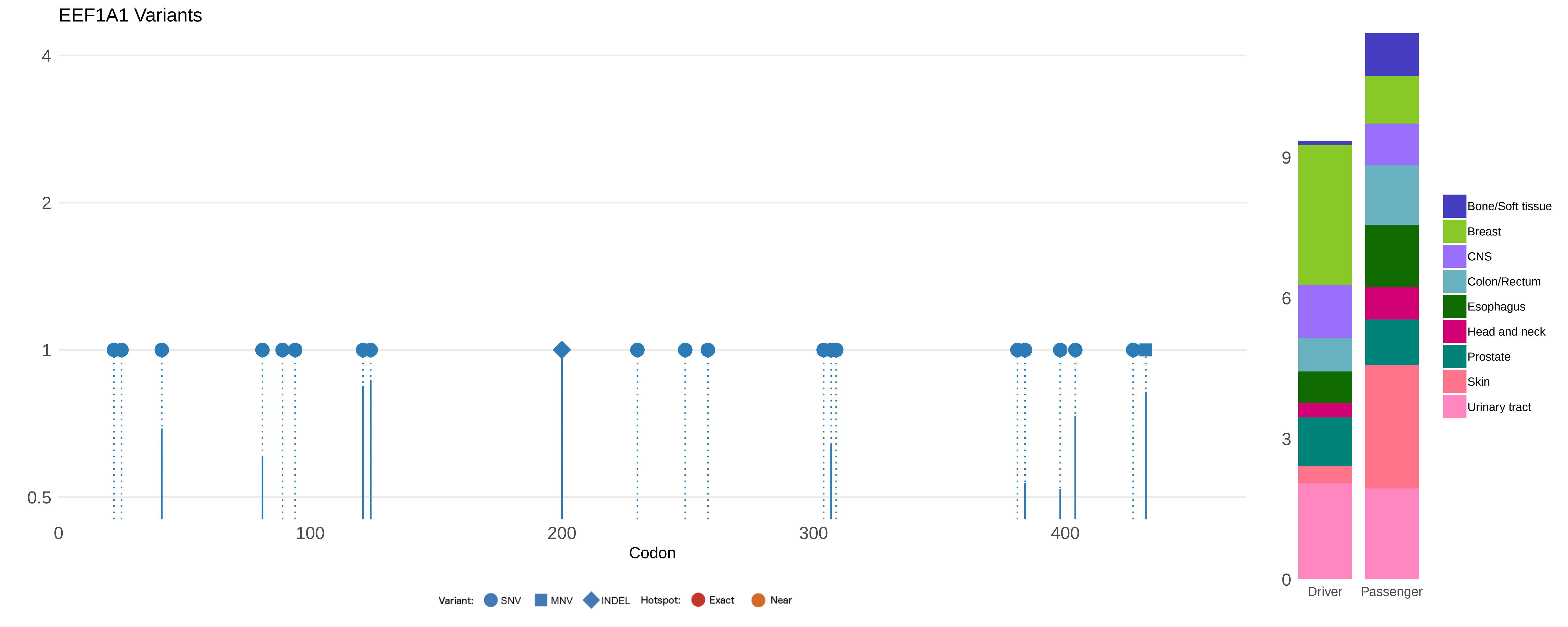
EEF1A1 Variants
4
2
1
0.5
0
100
200
300
400
Codon
Variant: SNV MNV INDEL Hotspot: Exact Near
9
6
3
0
Driver
Passenger
Bone/Soft tissue
Breast
CNS
Colon/Rectum
Esophagus
Head and neck
Prostate
Skin
Urinary tract

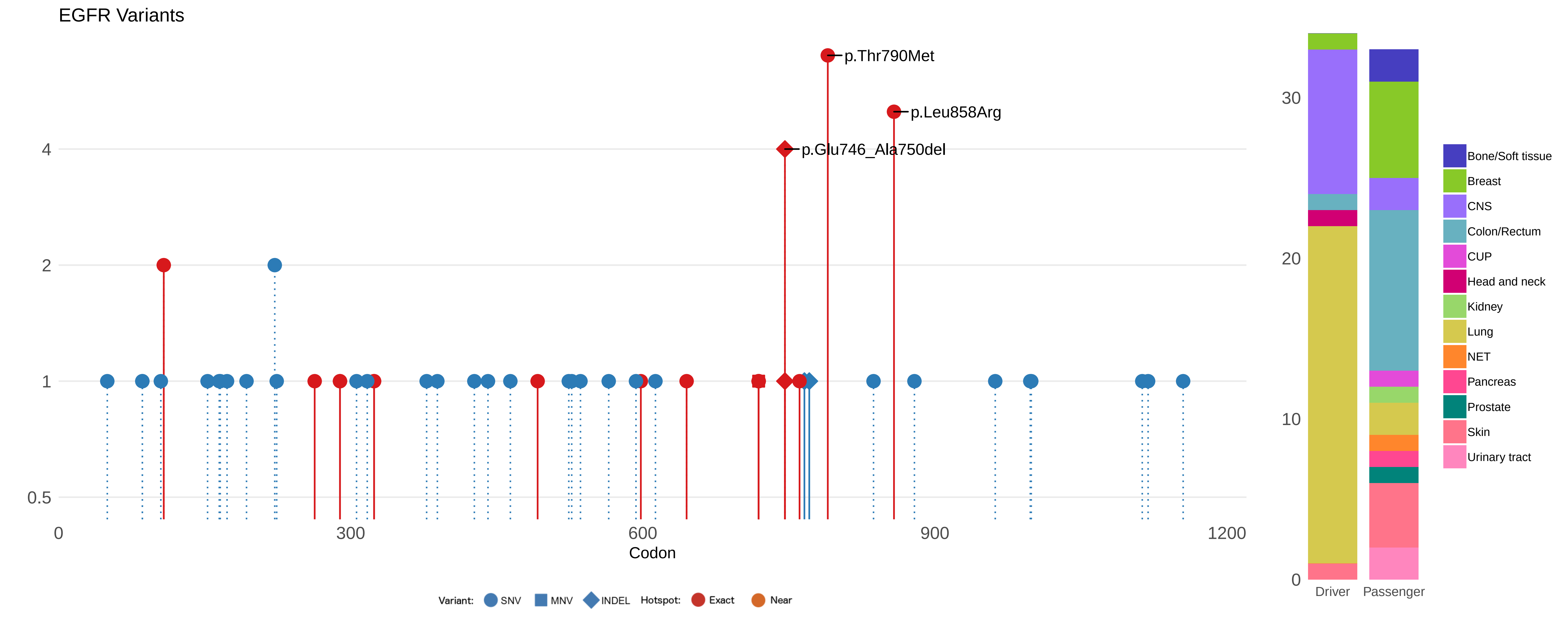
EGFR Variants
p.Thr790Met
p.Leu858Arg
p.Glu746_Ala750del
4
2
1
0.5
0
300
600
900
1200
Codon
Variant: SNV MNV INDEL Hotspot: Exact Near
30
20
10
0
Driver
Passenger
Bone/Soft tissue
Breast
CNS
Colon/Rectum
CUP
Head and neck
Kidney
Lung
NET
Pancreas
Prostate
Skin
Urinary tract

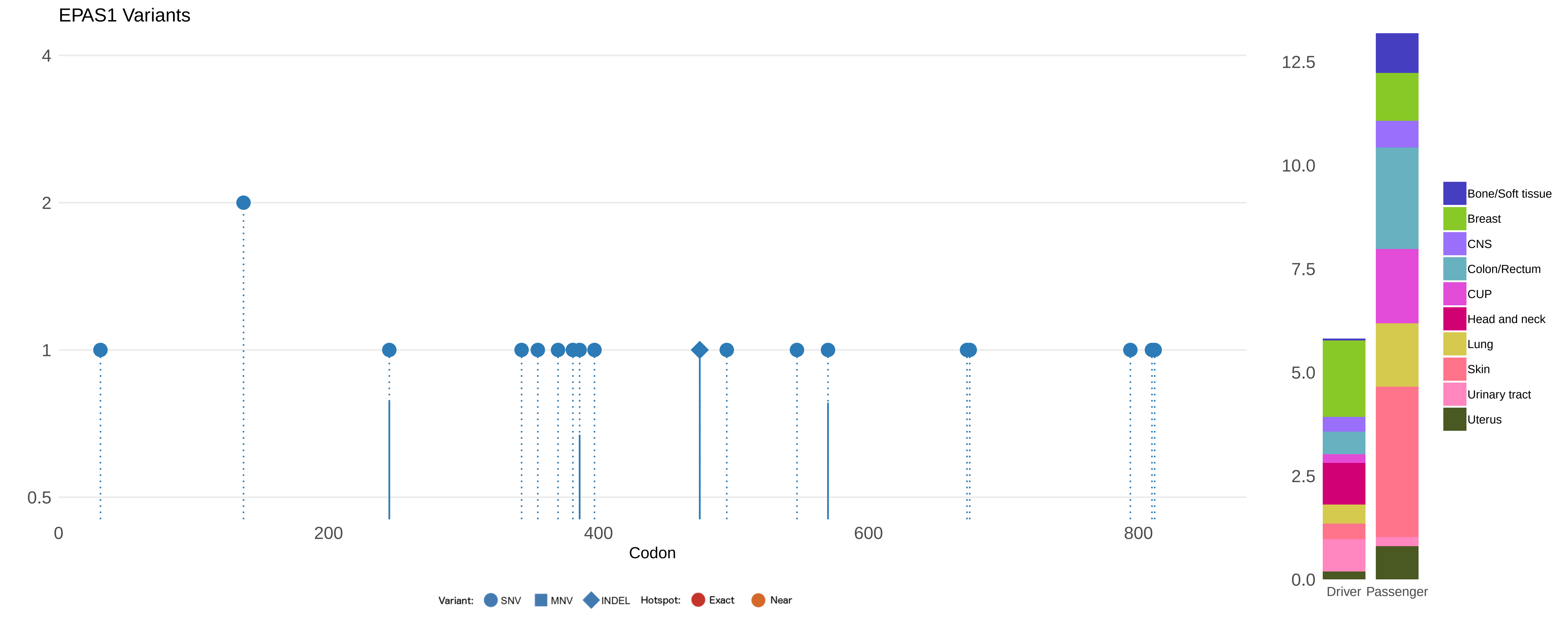

EPAS1 Variants
4
2
1
0.5
0
200
400
600
800
Codon
Variant: SNV MNV INDEL Hotspot: Exact Near
12.5
10.0
7.5
5.0
2.5
0.0
Driver
Passenger
Bone/Soft tissue
Breast
CNS
Colon/Rectum
CUP
Head and neck
Lung
Skin
Urinary tract
Uterus

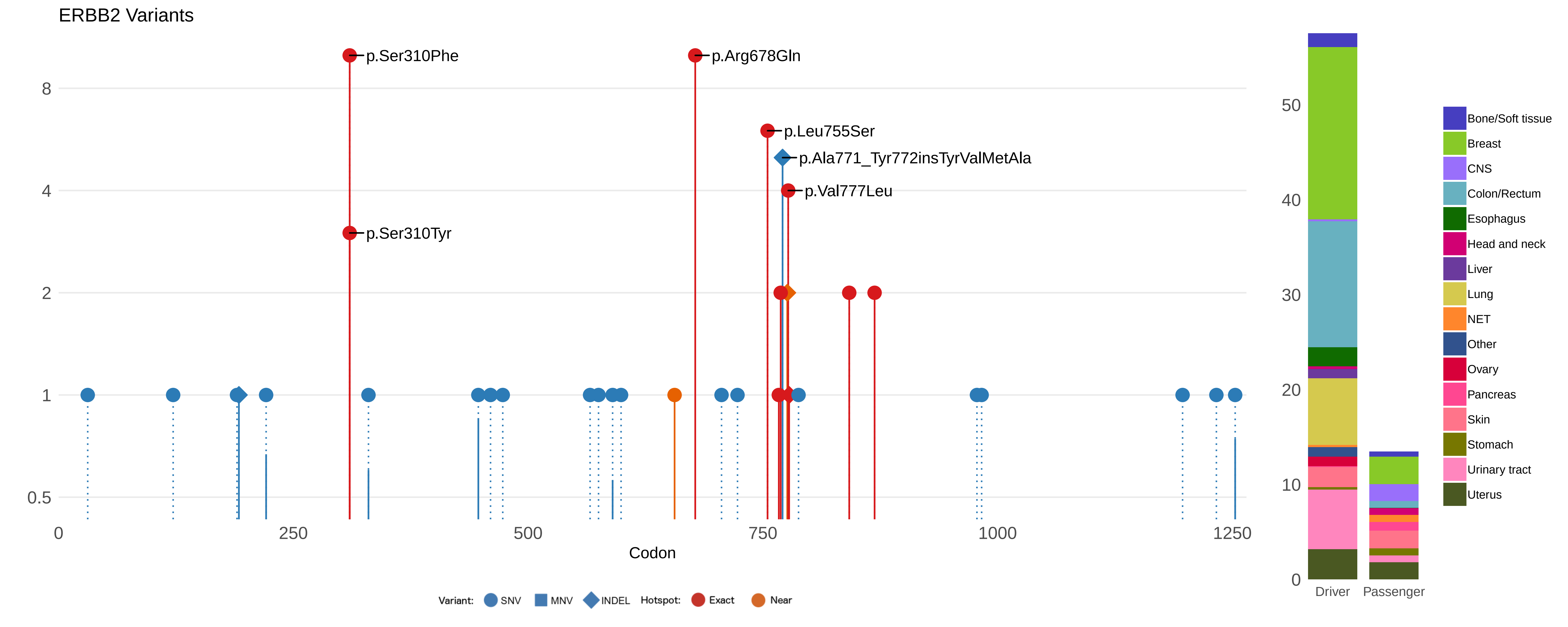

ERBB2 Variants
p.Ser310Phe
p.Arg678Gln
p.Leu755Ser
p.Ala771_Tyr772insTyrValMetAla
p.Val777Leu
p.Ser310Tyr
8
4
2
1
0.5
0
250
500
750
1000
1250
Codon
Variant:
SNV
MNV
INDEL
Hotspot:
Exact
Near
50
40
30
20
10
0
Driver
Passenger
Bone/Soft tissue
Breast
CNS
Colon/Rectum
Esophagus
Head and neck
Liver
Lung
NET
Other
Ovary
Pancreas
Skin
Stomach
Urinary tract
Uterus

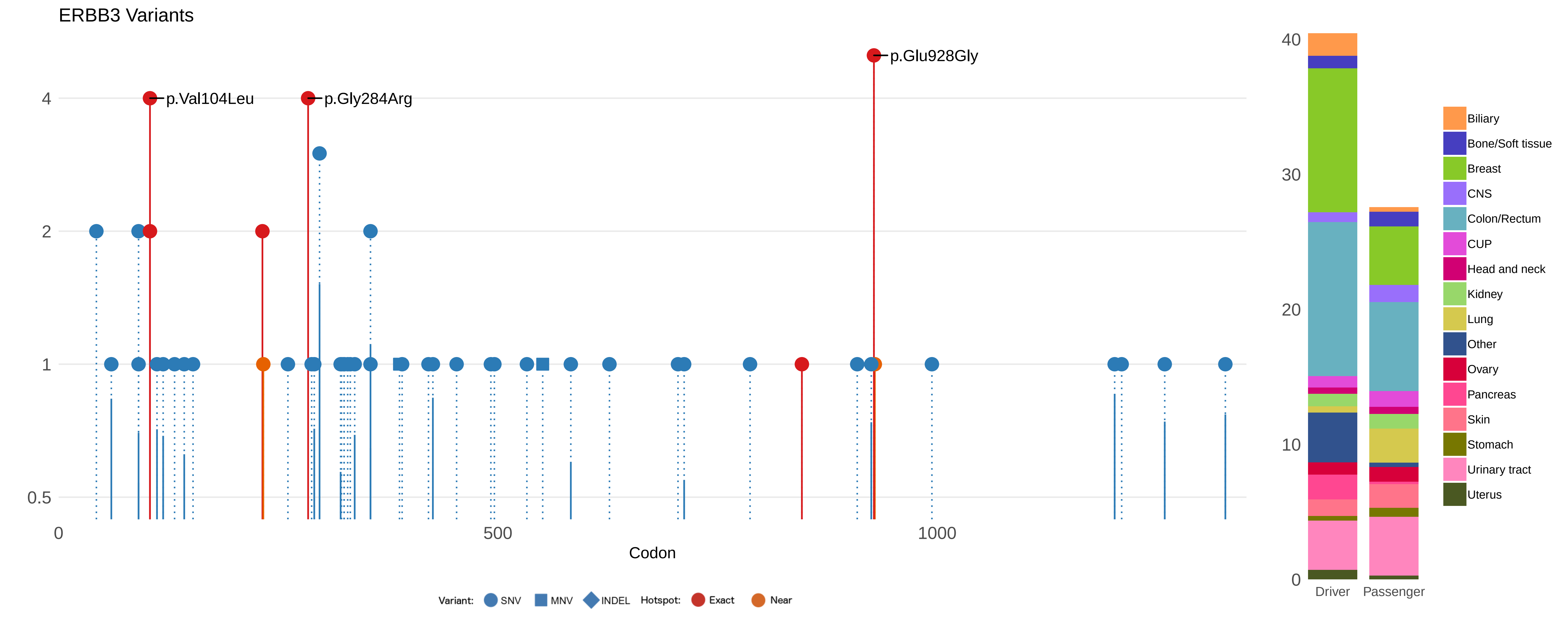

ERBB3 Variants
p.Val104Leu
p.Gly284Arg
p.Glu928Gly
4
2
1
0.5
0
500
1000
Codon
Variant: SNV MNV INDEL Hotspot: Exact Near
40
30
20
10
0
Driver
Passenger
Biliary
Bone/Soft tissue
Breast
CNS
Colon/Rectum
CUP
Head and neck
Kidney
Lung
Other
Ovary
Pancreas
Skin
Stomach
Urinary tract
Uterus

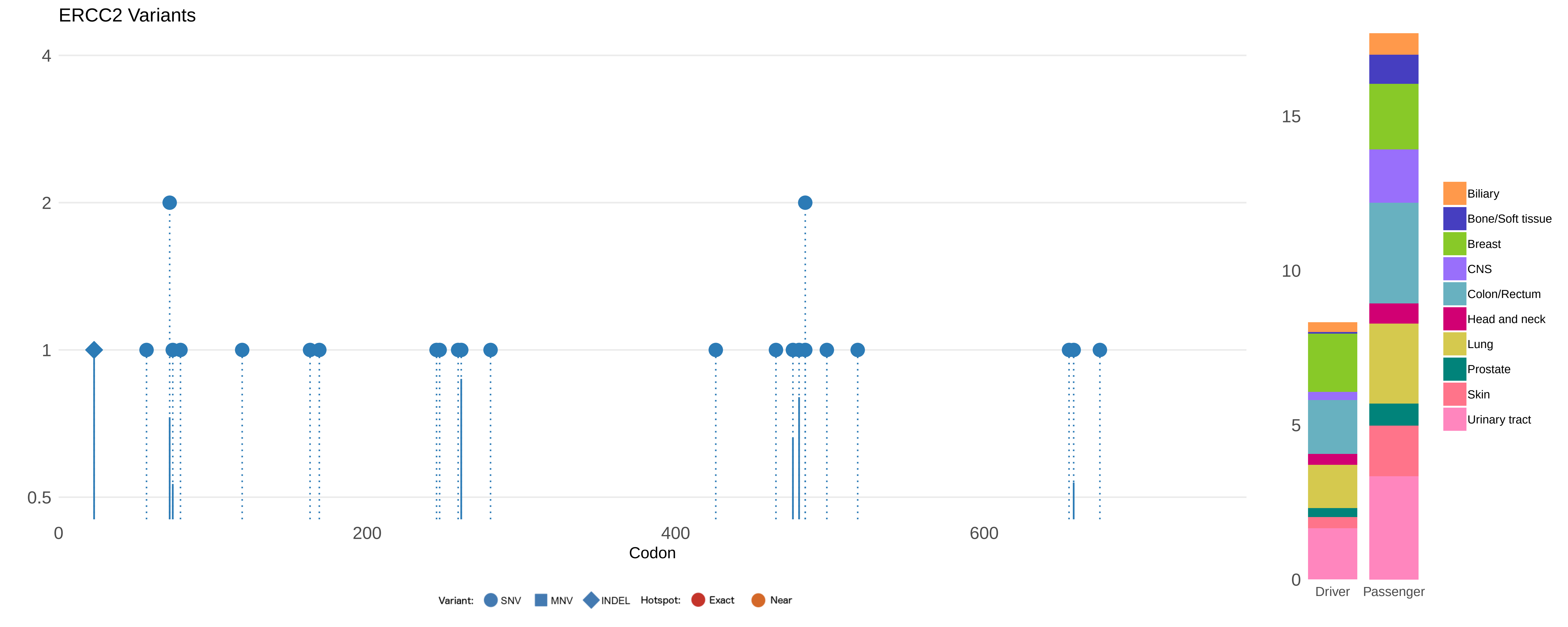

ERCC2 Variants
4
2
1
0.5
0
200
400
600
Codon
Variant:
SNV
MNV
INDEL
Hotspot:
Exact
Near
15
10
5
0
Driver
Passenger
Biliary
Bone/Soft tissue
Breast
CNS
Colon/Rectum
Head and neck
Lung
Prostate
Skin
Urinary tract

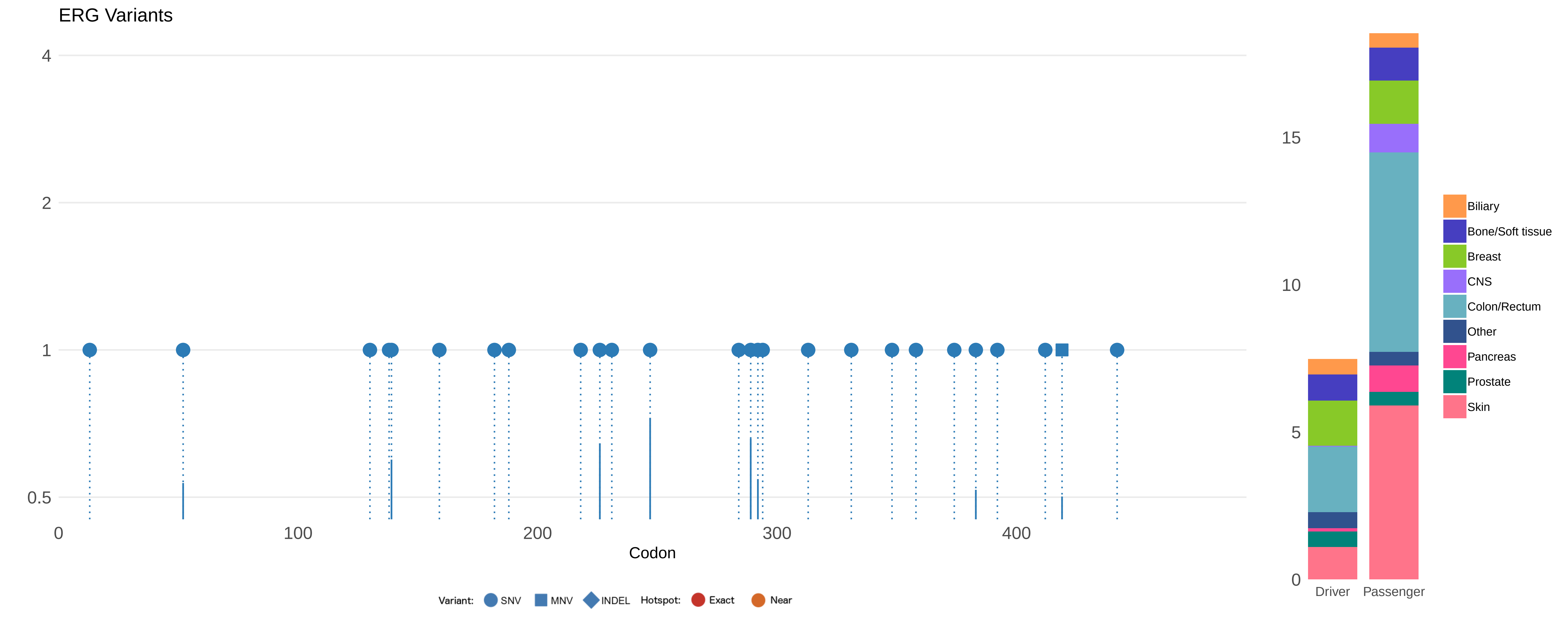
ERG Variants
4
2
1
0.5
0
100
200
300
400
Codon
Variant: SNV MNV INDEL Hotspot: Exact Near
15
10
5
0
Driver
Passenger
Biliary
Bone/Soft tissue
Breast
CNS
Colon/Rectum
Other
Pancreas
Prostate
Skin

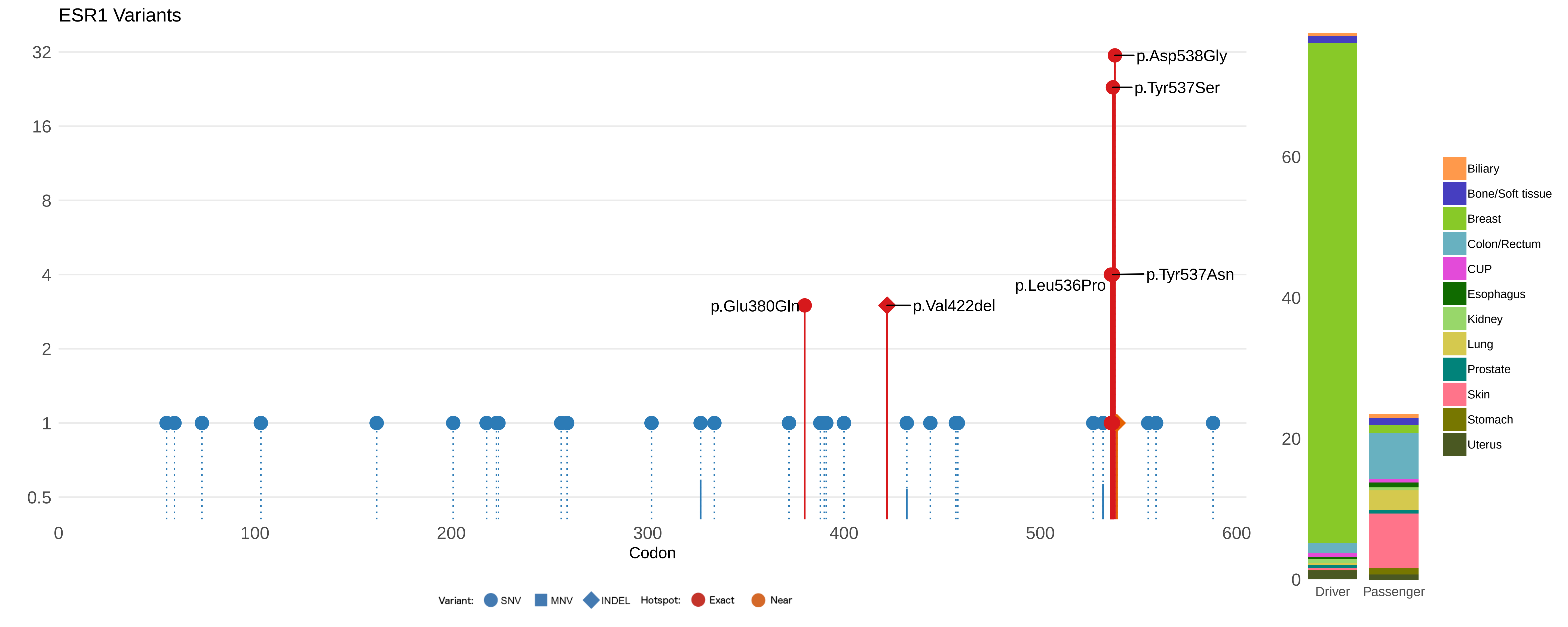

ESR1 Variants
32
16
8
4
2
1
0.5
p.Asp538Gly
p.Tyr537Ser
p.Tyr537Asn
p.Leu536Pro
p.Glu380Gln
p.Val422del
0
100
200
300
400
500
600
Codon
Variant: SNV MNV INDEL Hotspot: Exact Near
60
40
20
0
Driver
Passenger
Biliary
Bone/Soft tissue
Breast
Colon/Rectum
CUP
Esophagus
Kidney
Lung
Prostate
Skin
Stomach
Uterus

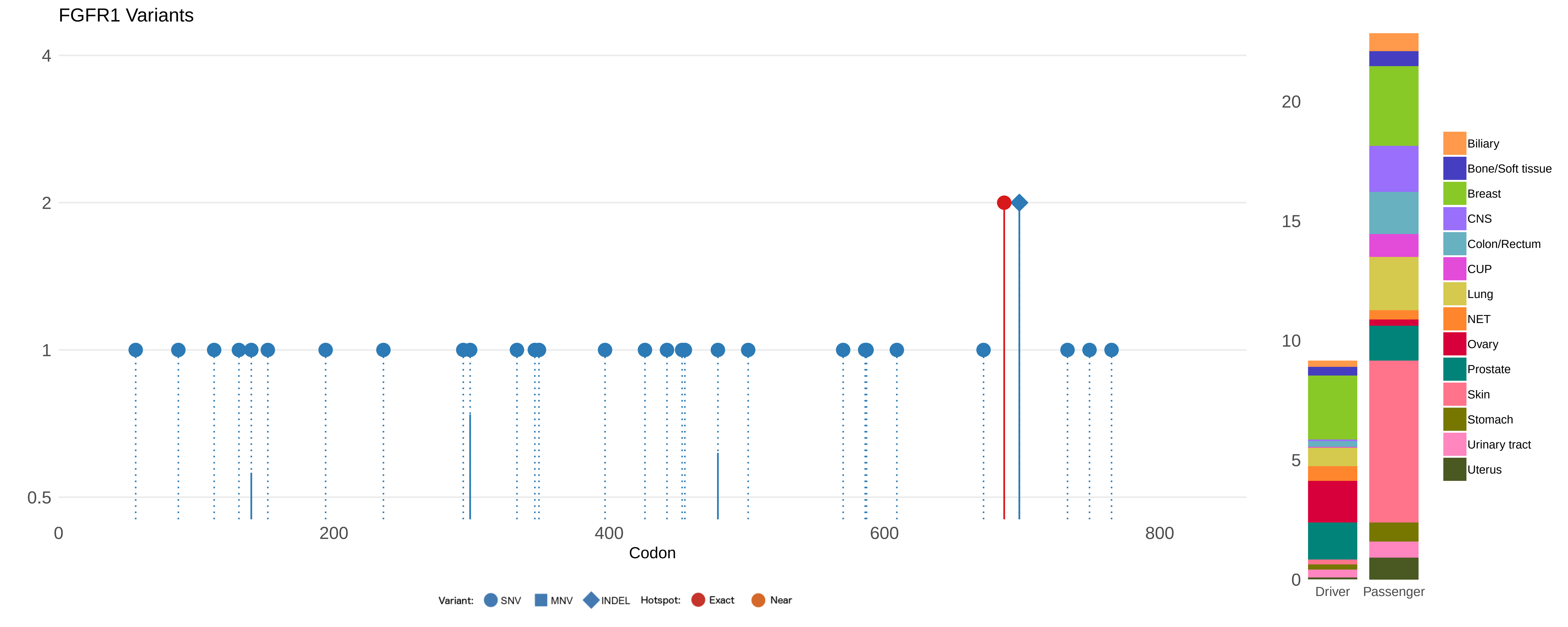
FGFR1 Variants
4
2
1
0.5
0
200
400
600
800
Codon
Variant:
SNV
MNV
INDEL
Hotspot:
Exact
Near
20
15
10
5
0
Driver
Passenger
Biliary
Bone/Soft tissue
Breast
CNS
Colon/Rectum
CUP
Lung
NET
Ovary
Prostate
Skin
Stomach
Urinary tract
Uterus

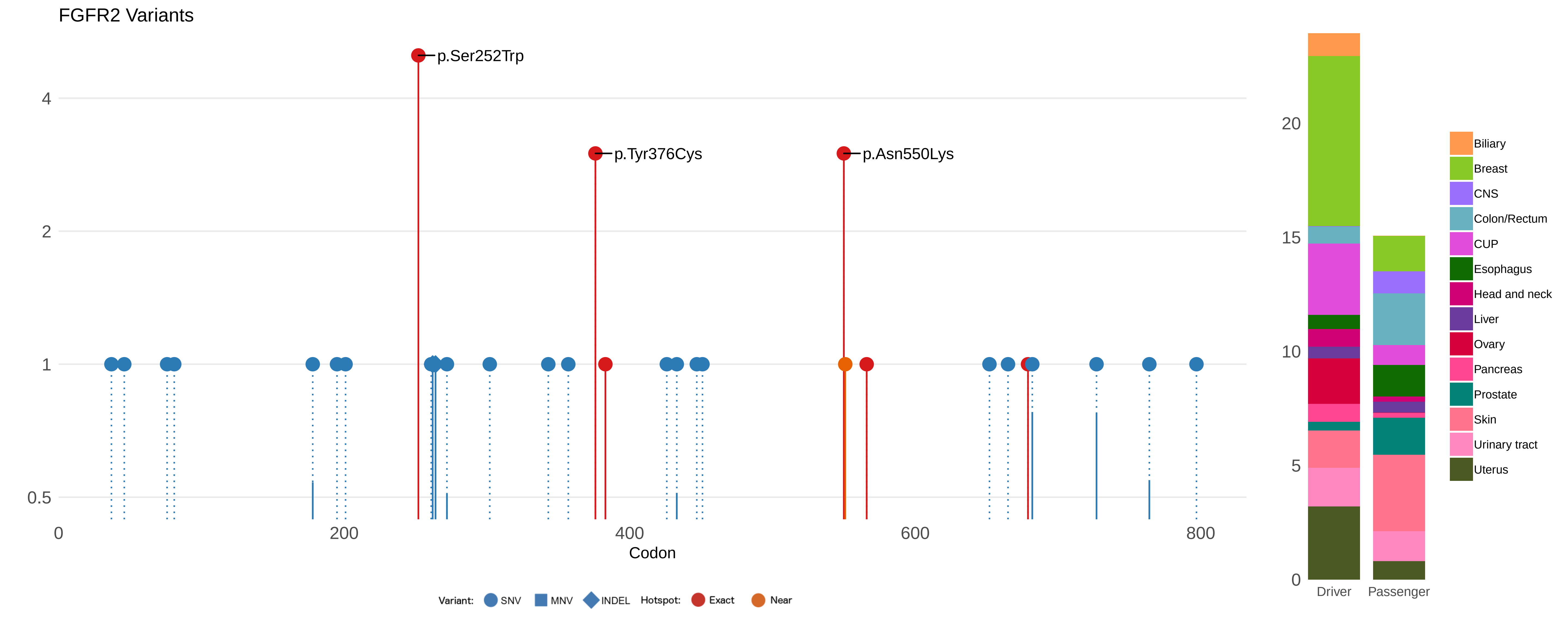

FGFR2 Variants
p.Ser252Trp
p.Tyr376Cys
p.Asn550Lys
4
2
1
0.5
0
200
400
600
800
Codon
Variant: SNV MNV INDEL Hotspot: Exact Near
20
15
10
5
0
Driver
Passenger
Biliary
Breast
CNS
Colon/Rectum
CUP
Esophagus
Head and neck
Liver
Ovary
Pancreas
Prostate
Skin
Urinary tract
Uterus

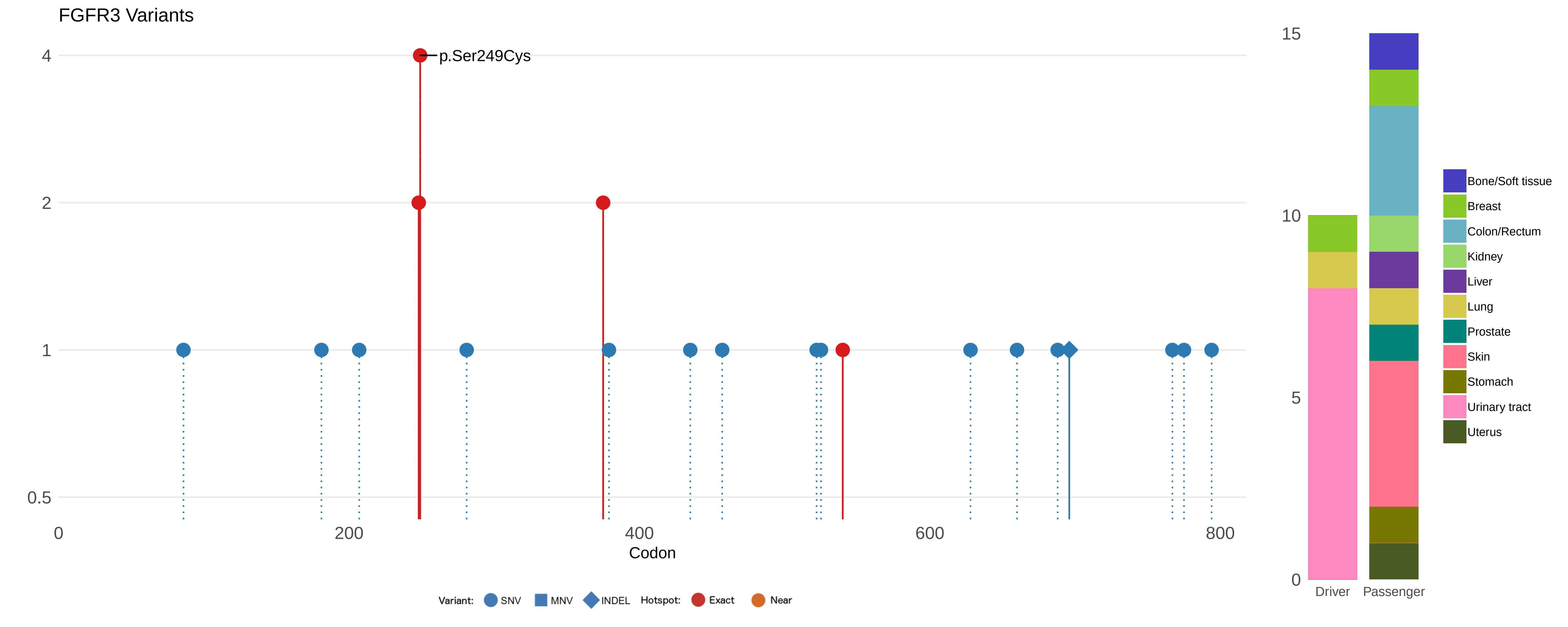

FGFR3 Variants
p.Ser249Cys
4
2
1
0.5
0
200
400
600
800
Codon
Variant:
SNV
MNV
INDEL
Hotspot:
Exact
Near
15
10
5
0
Driver
Passenger
Bone/Soft tissue
Breast
Colon/Rectum
Kidney
Liver
Lung
Prostate
Skin
Stomach
Urinary tract
Uterus

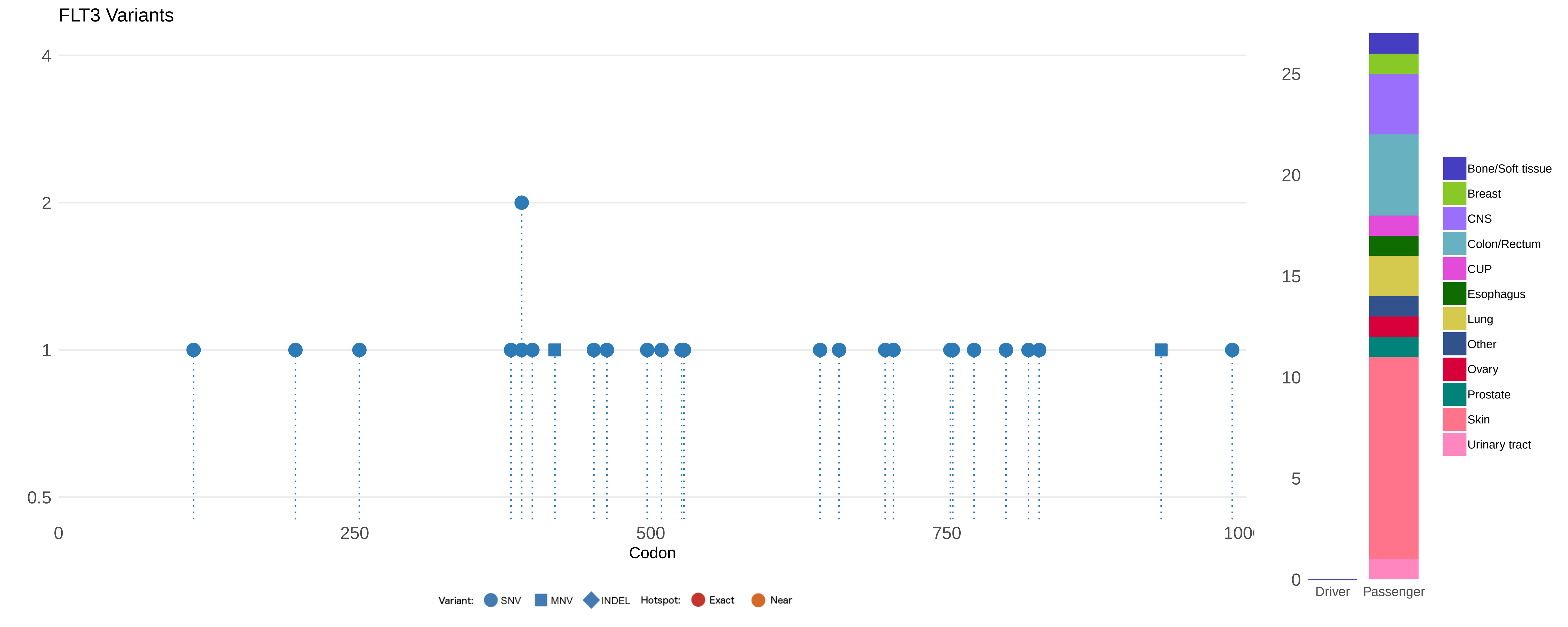

FLT3 Variants
4
2
1
0.5
0
250
500
750
100
Codon
Variant: SNV MNV INDEL Hotspot: Exact Near
25
20
15
10
5
0
Driver
Passenger
Bone/Soft tissue
Breast
CNS
Colon/Rectum
CUP
Esophagus
Lung
Other
Ovary
Prostate
Skin
Urinary tract

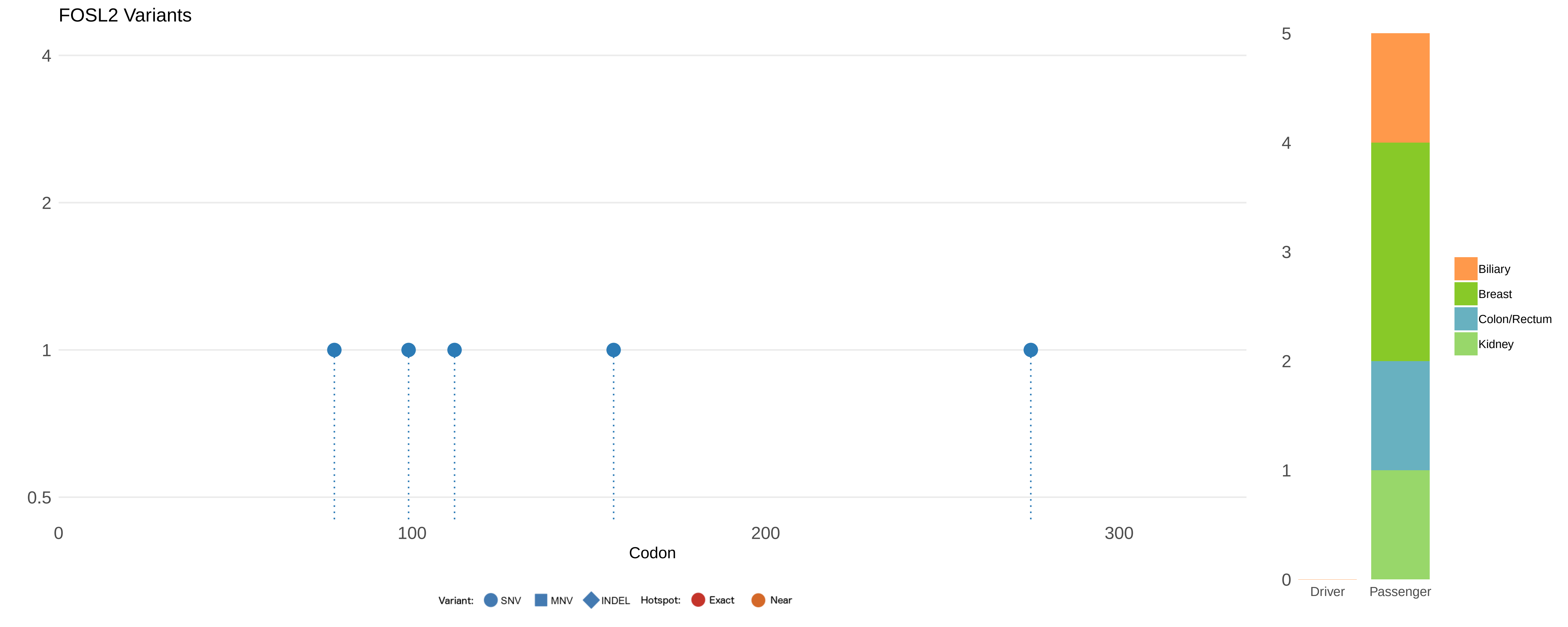
FOSL2 Variants
4
2
1
0.5
0
100
200
300
Codon
Variant:
SNV
MNV
INDEL
Hotspot:
Exact
Near
5
4
3
2
1
0
Driver
Passenger
Biliary
Breast
Colon/Rectum
Kidney

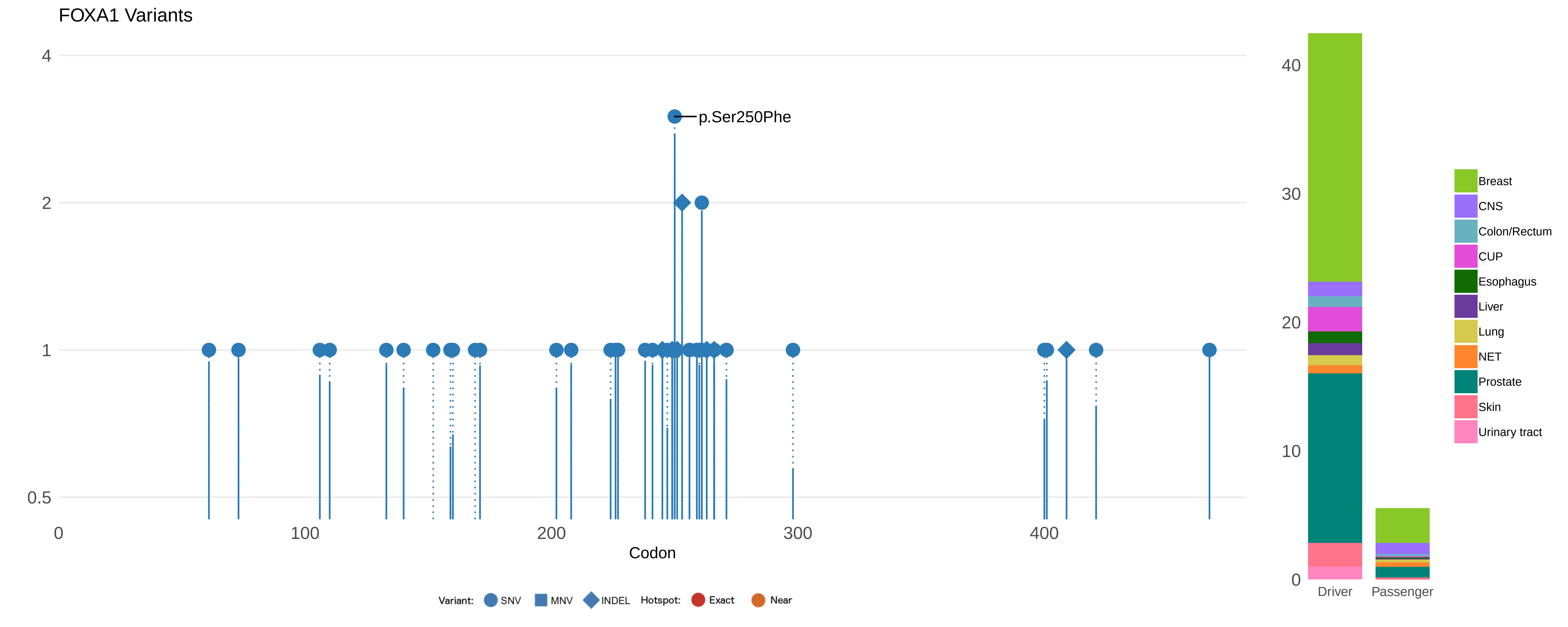

FOXA1 Variants
p.Ser250Phe
4
2
1
0.5
0
100
200
300
400
Codon
40
30
20
10
0
Driver
Passenger
Breast
CNS
Colon/Rectum
CUP
Esophagus
Liver
Lung
NET
Prostate
Skin
Urinary tract
Variant: SNV MNV INDEL Hotspot: Exact Near

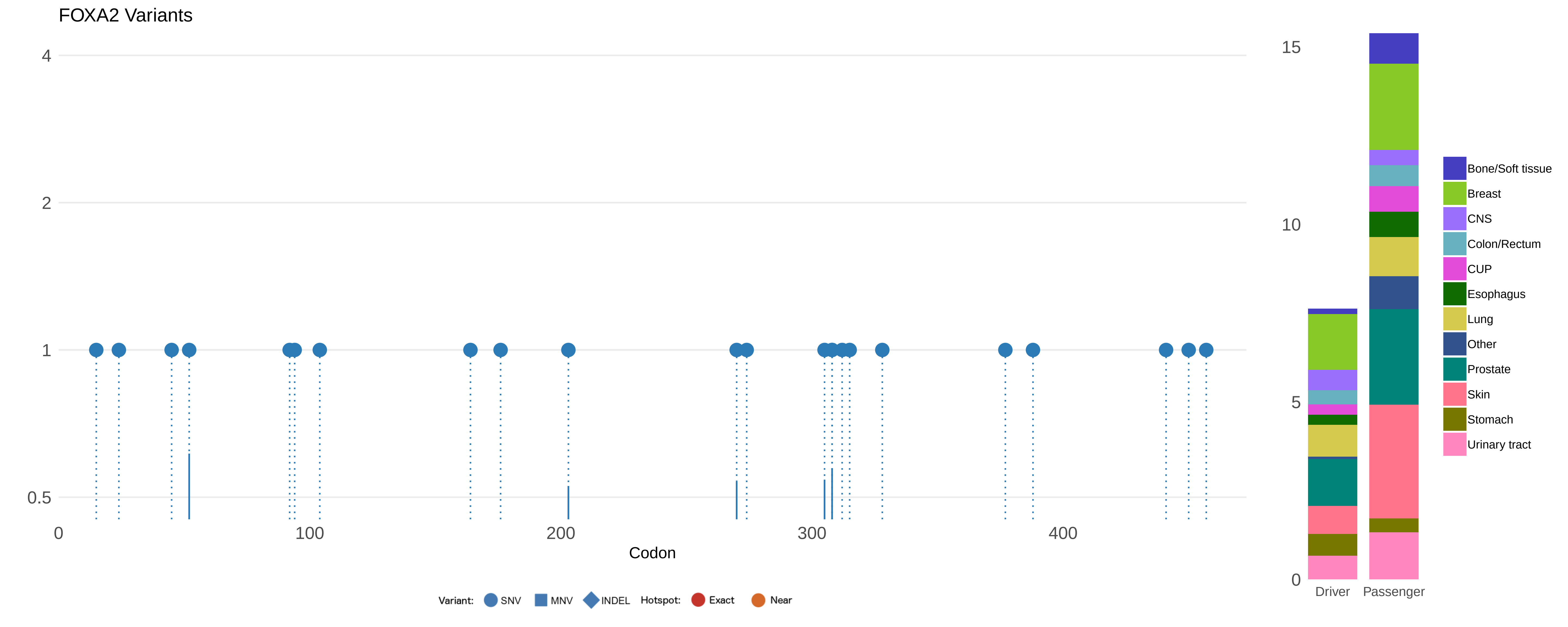

FOXA2 Variants
4
2
1
0.5
0
100
200
300
400
Codon
Variant: SNV MNV INDEL Hotspot: Exact Near
15
10
5
0
Driver
Passenger
Bone/Soft tissue
Breast
CNS
Colon/Rectum
CUP
Esophagus
Lung
Other
Prostate
Skin
Stomach
Urinary tract

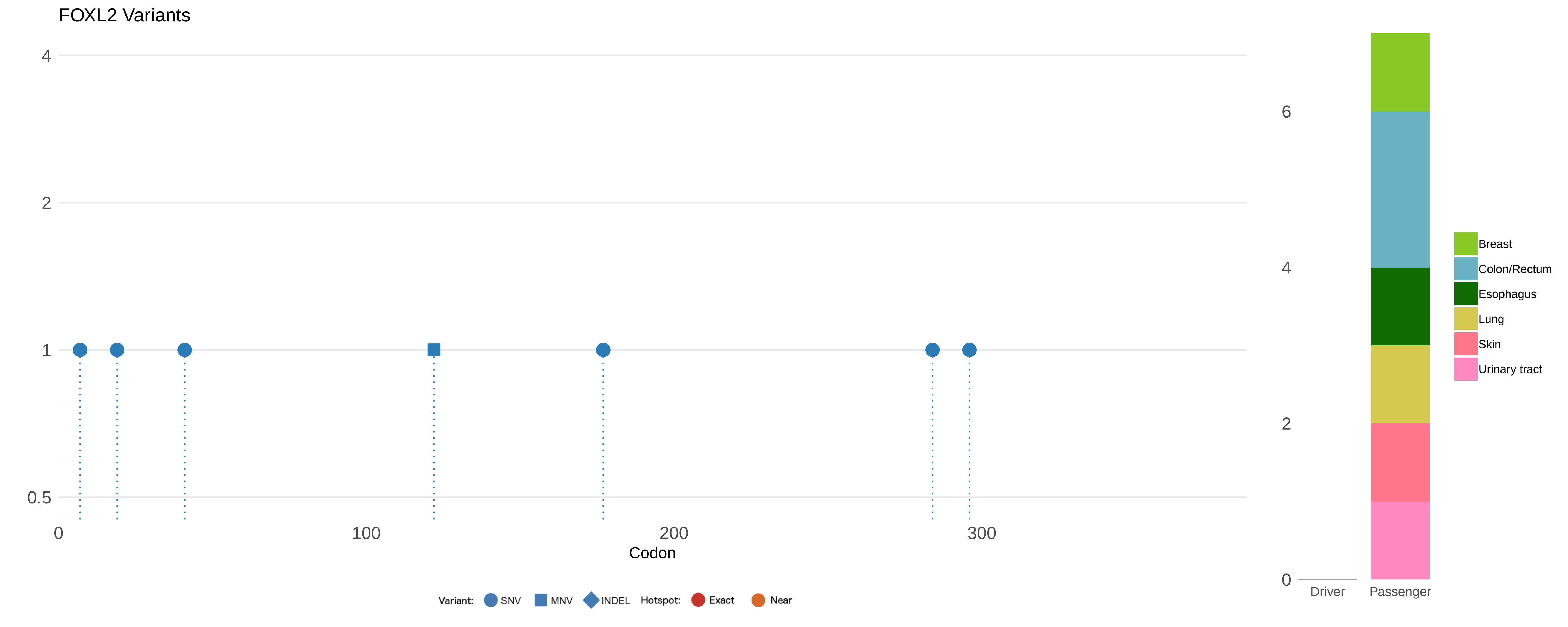
FOXL2 Variants
4
2
1
0.5
0
100
200
300
Codon
Variant:
SNV
MNV
INDEL
Hotspot:
Exact
Near
6
4
2
0
Driver
Passenger
Breast
Colon/Rectum
Esophagus
Lung
Skin
Urinary tract

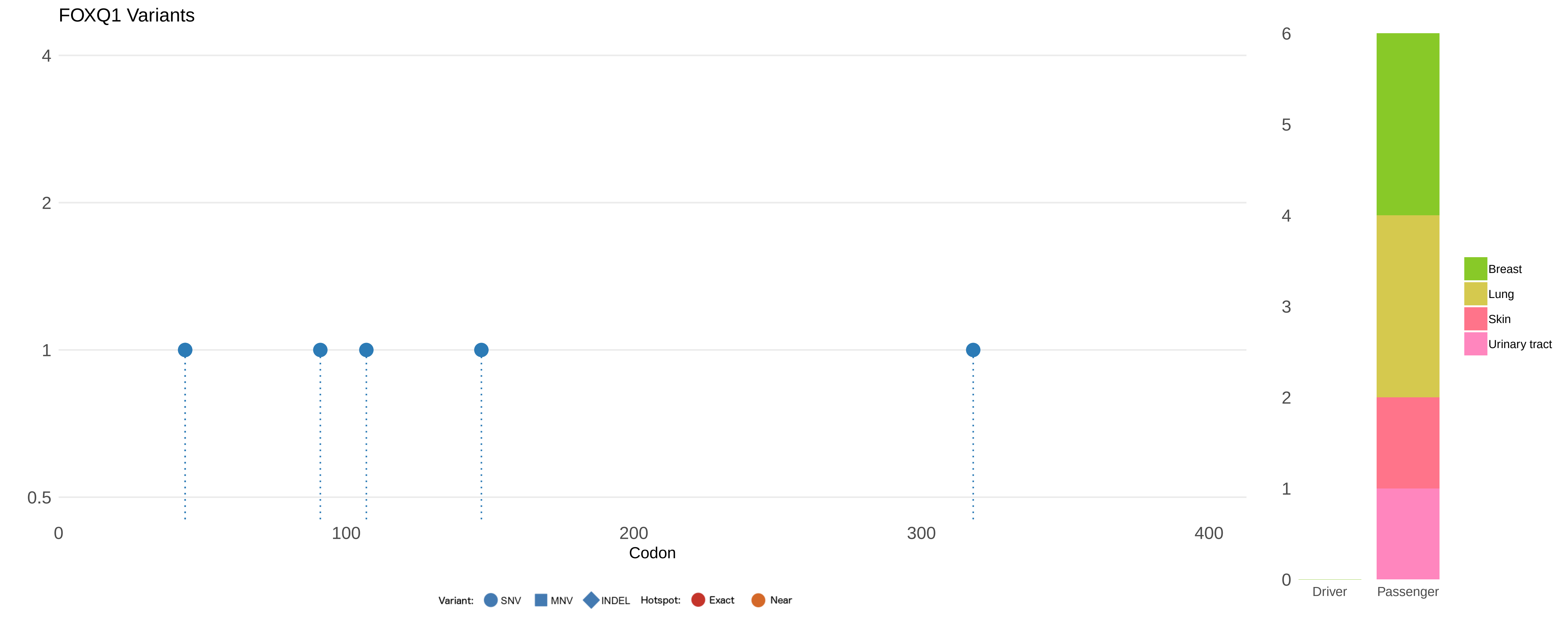
FOXQ1 Variants
4
2
1
0.5
0
100
200
300
400
Codon
Variant: SNV MNV INDEL Hotspot: Exact Near
6
5
4
3
2
1
0
Driver
Passenger
Breast
Lung
Skin
Urinary tract

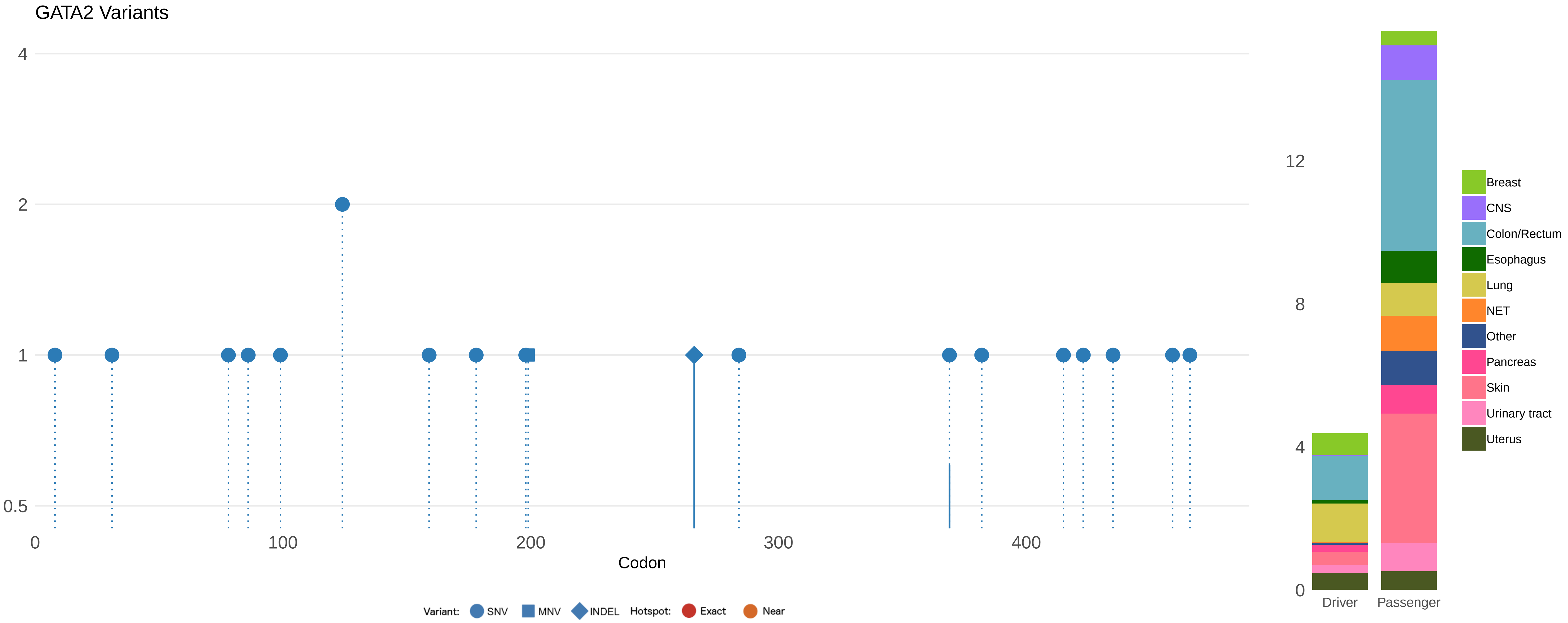

GATA2 Variants
4
2
1
0.5
0
100
200
300
400
Codon
Variant:
SNV
MNV
INDEL
Hotspot:
Exact
Near
12
8
4
0
Driver
Passenger
Breast
CNS
Colon/Rectum
Esophagus
Lung
NET
Other
Pancreas
Skin
Urinary tract
Uterus

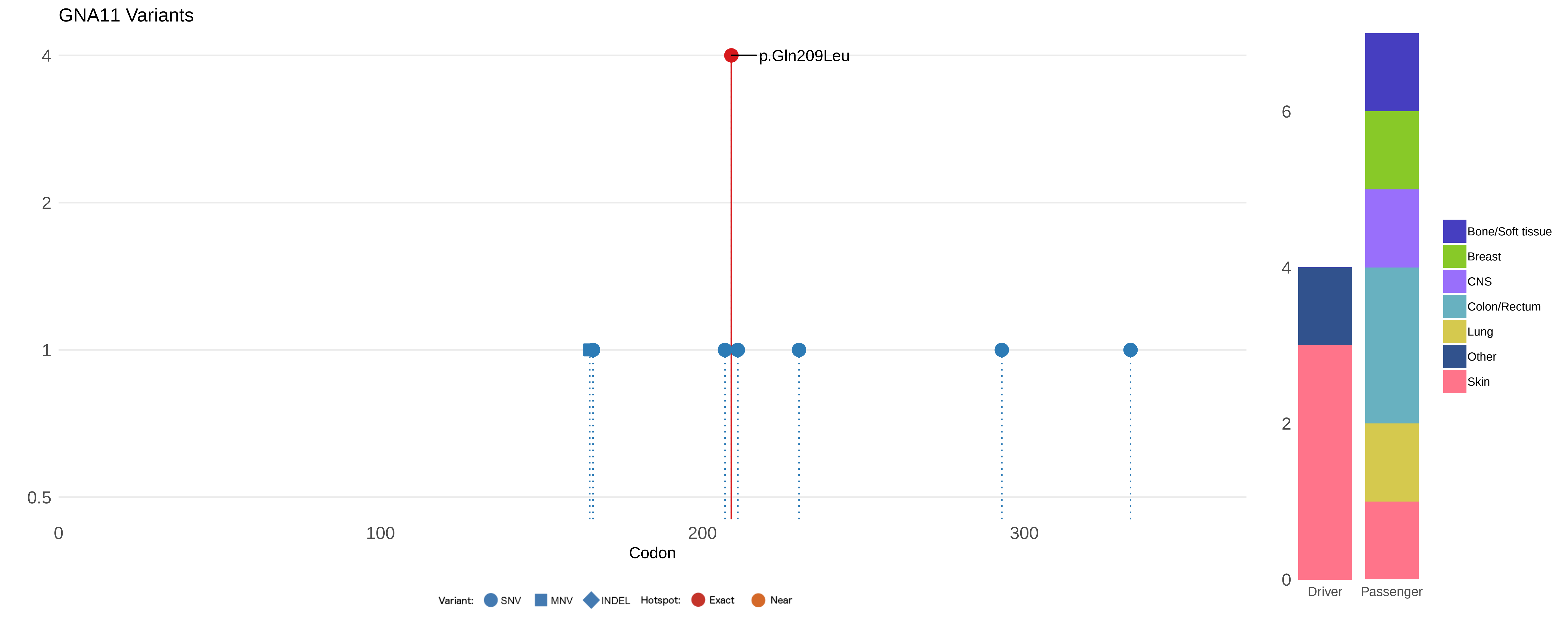

GNA11 Variants
p.Gln209Leu
4
2
1
0.5
0
100
200
300
Codon
Variant: SNV MNV INDEL Hotspot: Exact Near
6
4
2
0
Driver
Passenger
Bone/Soft tissue
Breast
CNS
Colon/Rectum
Lung
Other
Skin

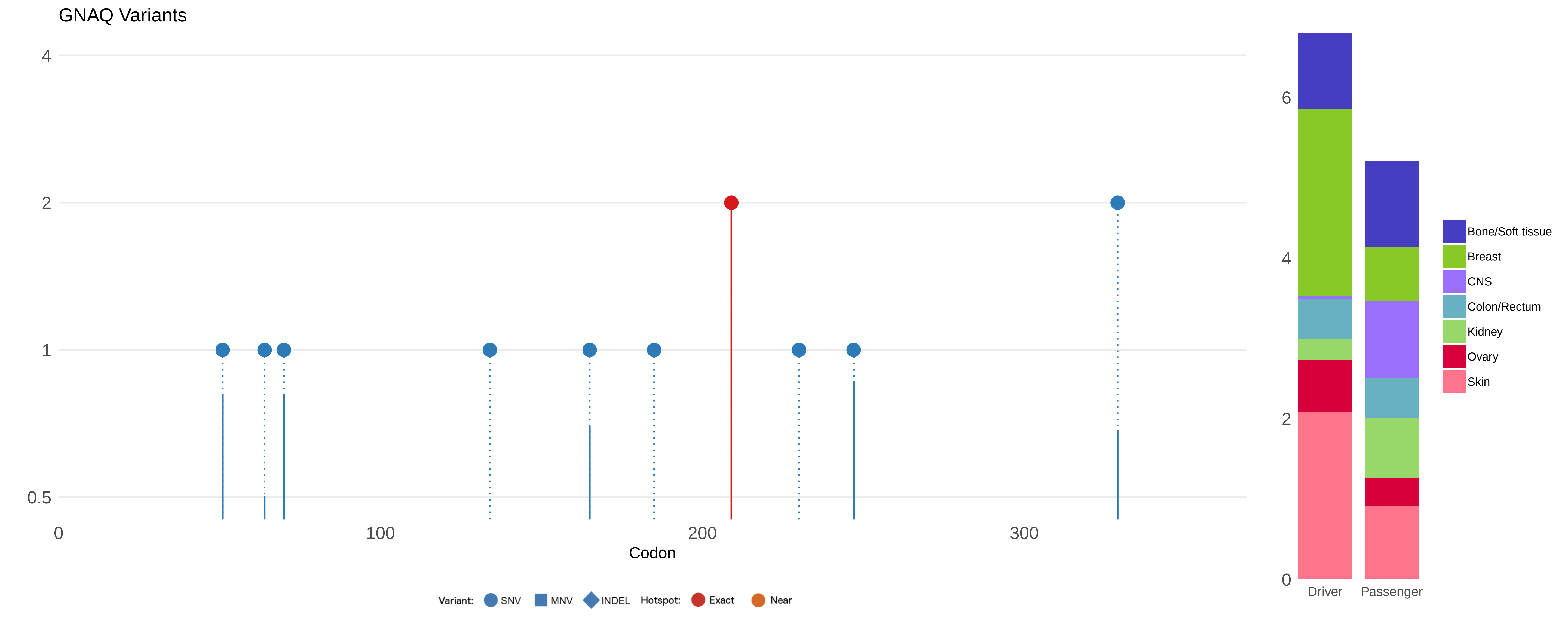

GNAQ Variants
4
2
1
0.5
0
100
200
300
Codon
Variant:
SNV
MNV
INDEL
Hotspot:
Exact
Near
6
4
2
0
Driver
Passenger
Bone/Soft tissue
Breast
CNS
Colon/Rectum
Kidney
Ovary
Skin

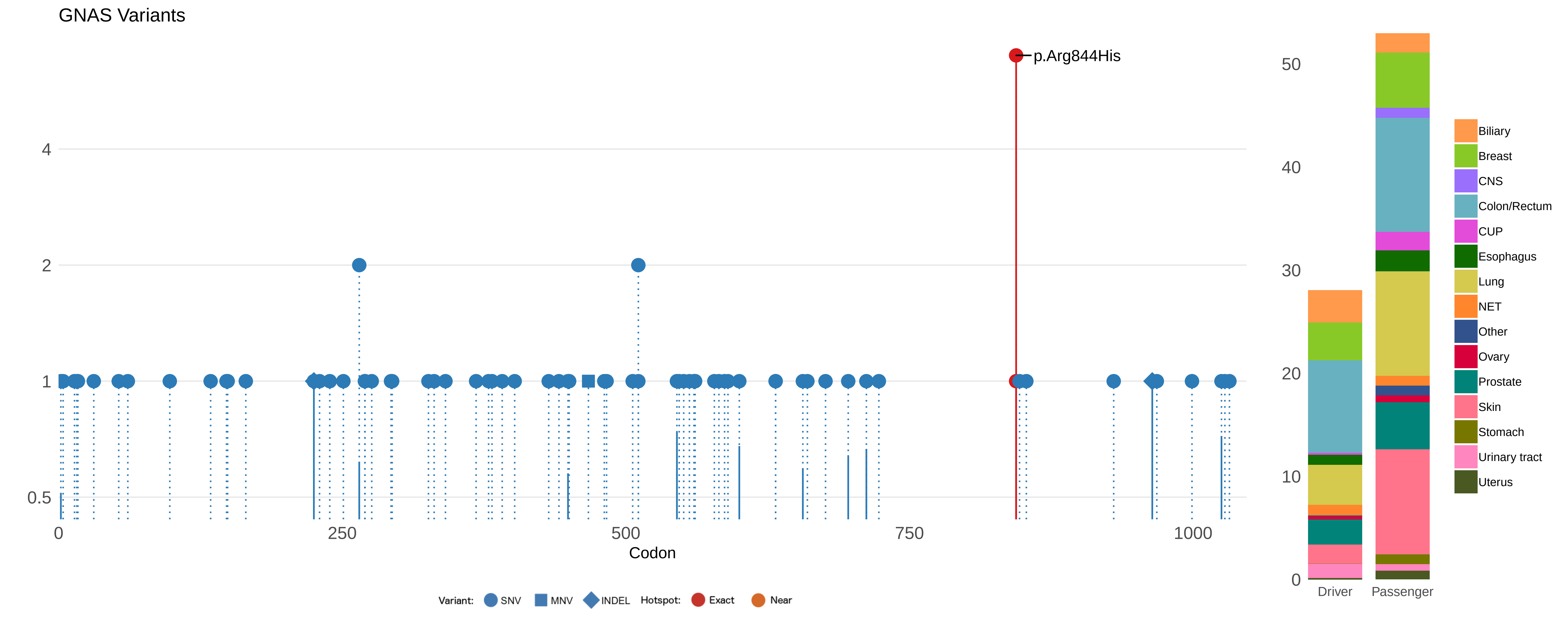
GNAS Variants
p.Arg844His
4
2
1
0.5
0
250
500
750
1000
Codon
Variant:
SNV
MNV
INDEL
Hotspot:
Exact
Near
50
40
30
20
10
0
Driver
Passenger
Biliary
Breast
CNS
Colon/Rectum
CUP
Esophagus
Lung
NET
Other
Ovary
Prostate
Skin
Stomach
Urinary tract
Uterus

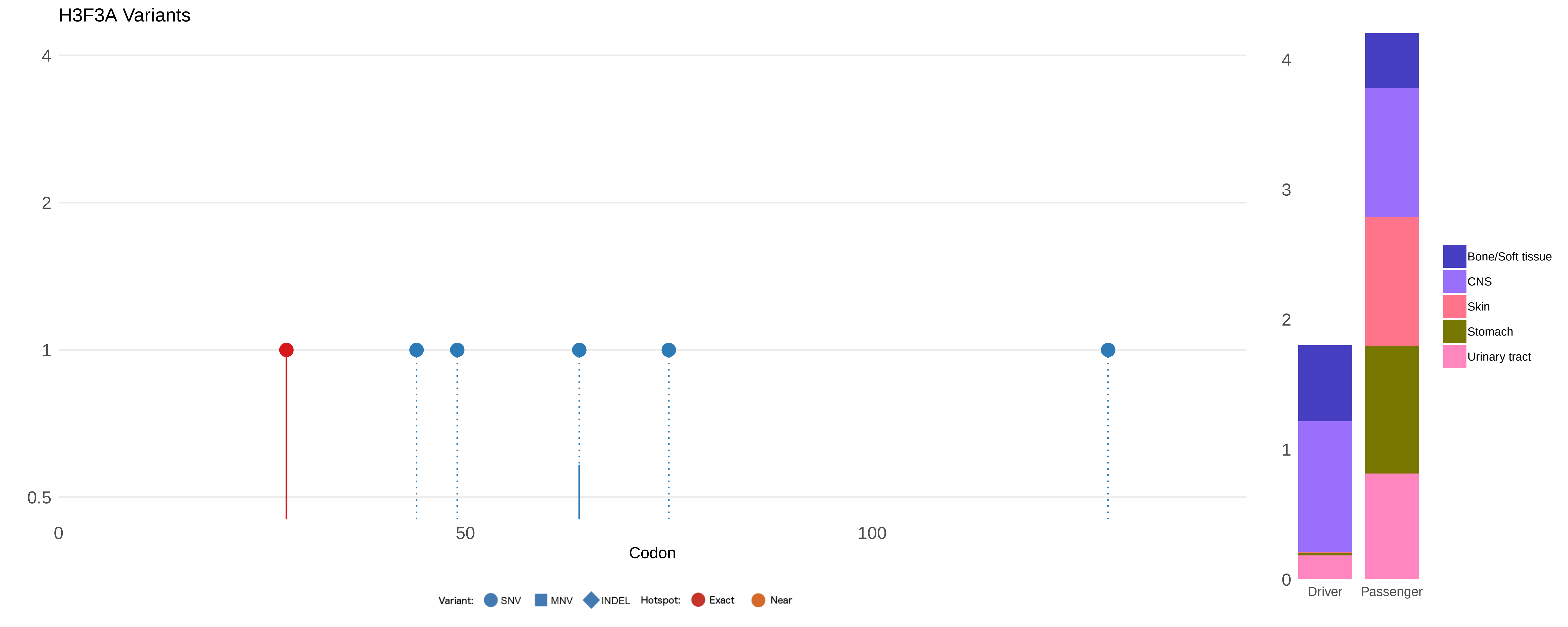

H3F3A Variants
4
2
1
0.5
0
50
100
Codon
Variant: SNV MNV INDEL Hotspot: Exact Near
4
3
2
1
0
Driver
Passenger
Bone/Soft tissue
CNS
Skin
Stomach
Urinary tract

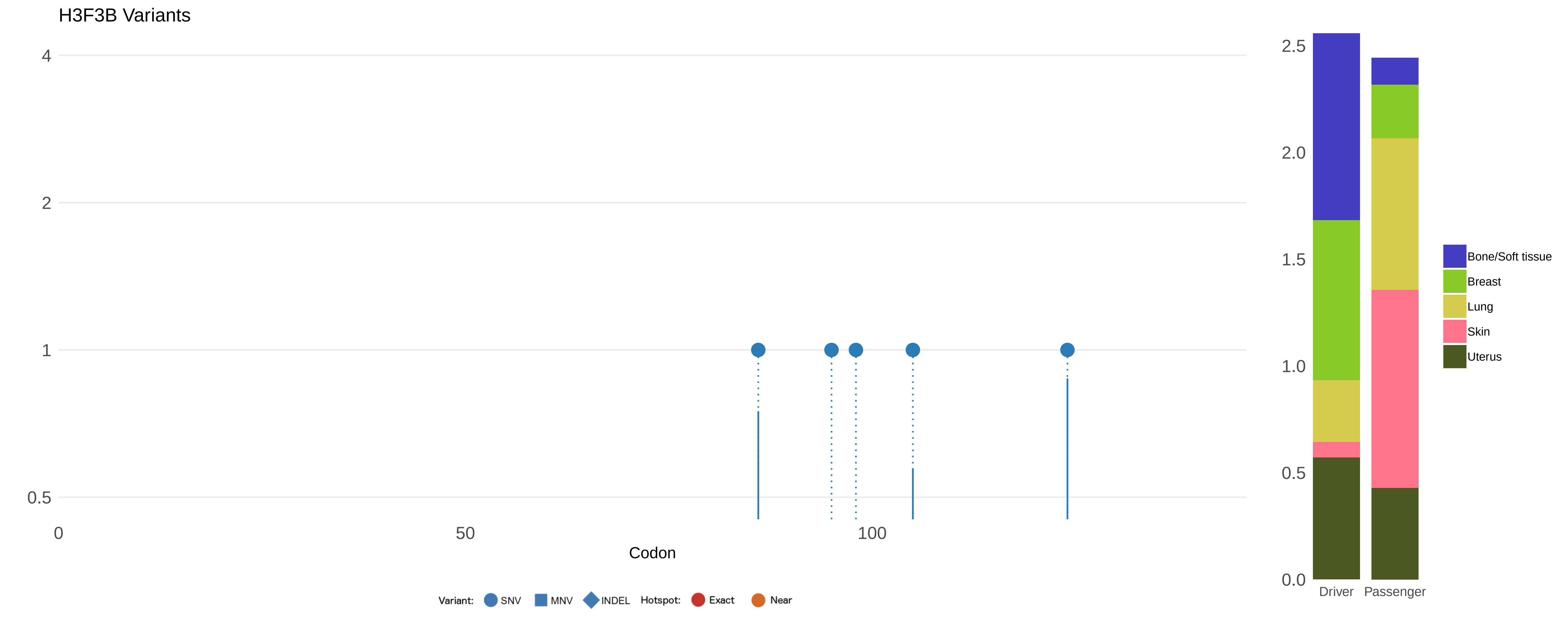

H3F3B Variants
4
2
1
0.5
0
50
100
Codon
Variant:
SNV
MNV
INDEL
Hotspot:
Exact
Near
2.5
2.0
1.5
1.0
0.5
0.0
Driver
Passenger
Bone/Soft tissue
Breast
Lung
Skin
Uterus

### HIF1A Variants

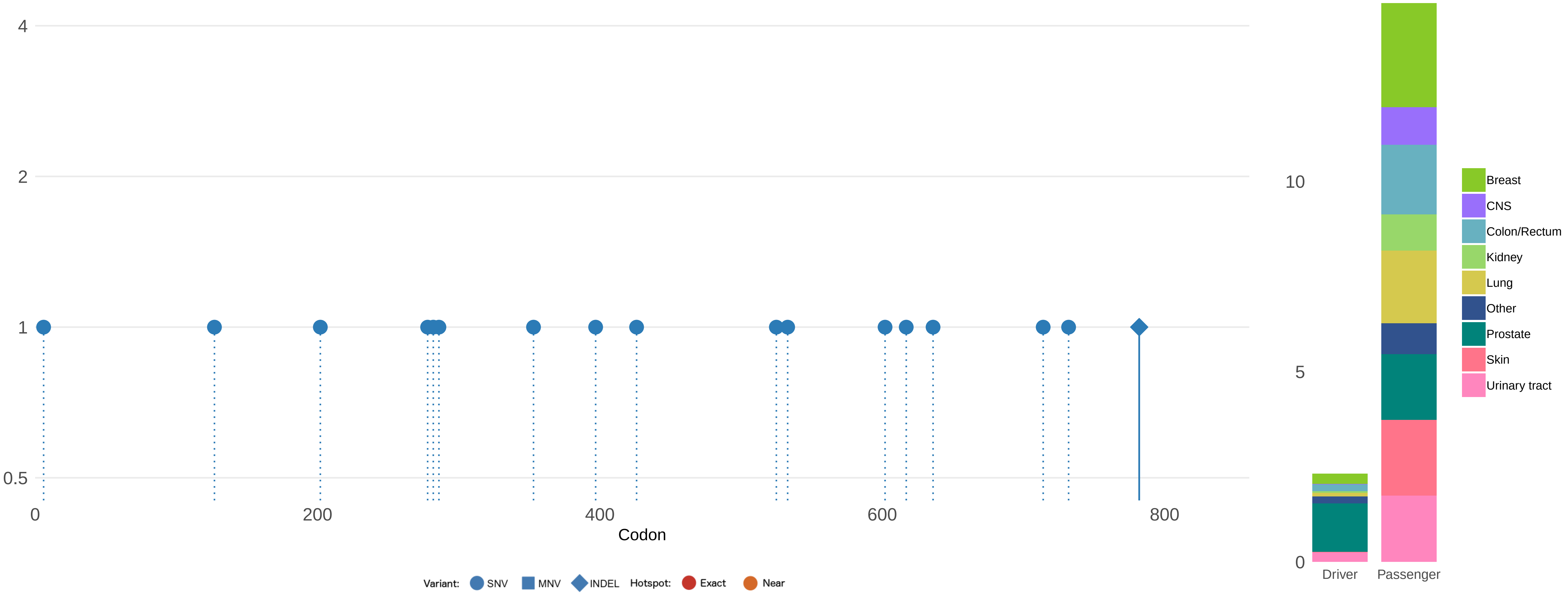

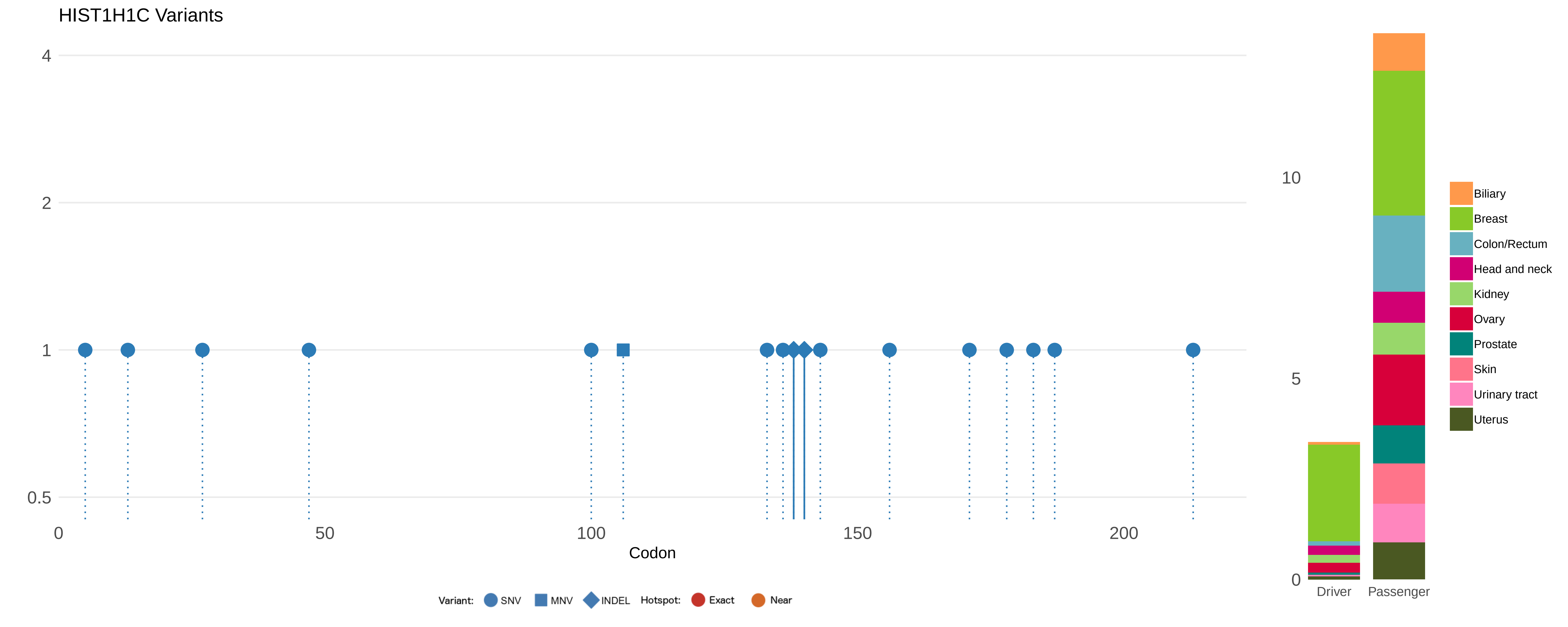
HIST1H1C Variants
4
2
1
0.5
0
50
100
150
200
Codon
Variant: SNV MNV INDEL Hotspot: Exact Near
10
5
0
Driver
Passenger
Biliary
Breast
Colon/Rectum
Head and neck
Kidney
Ovary
Prostate
Skin
Urinary tract
Uterus

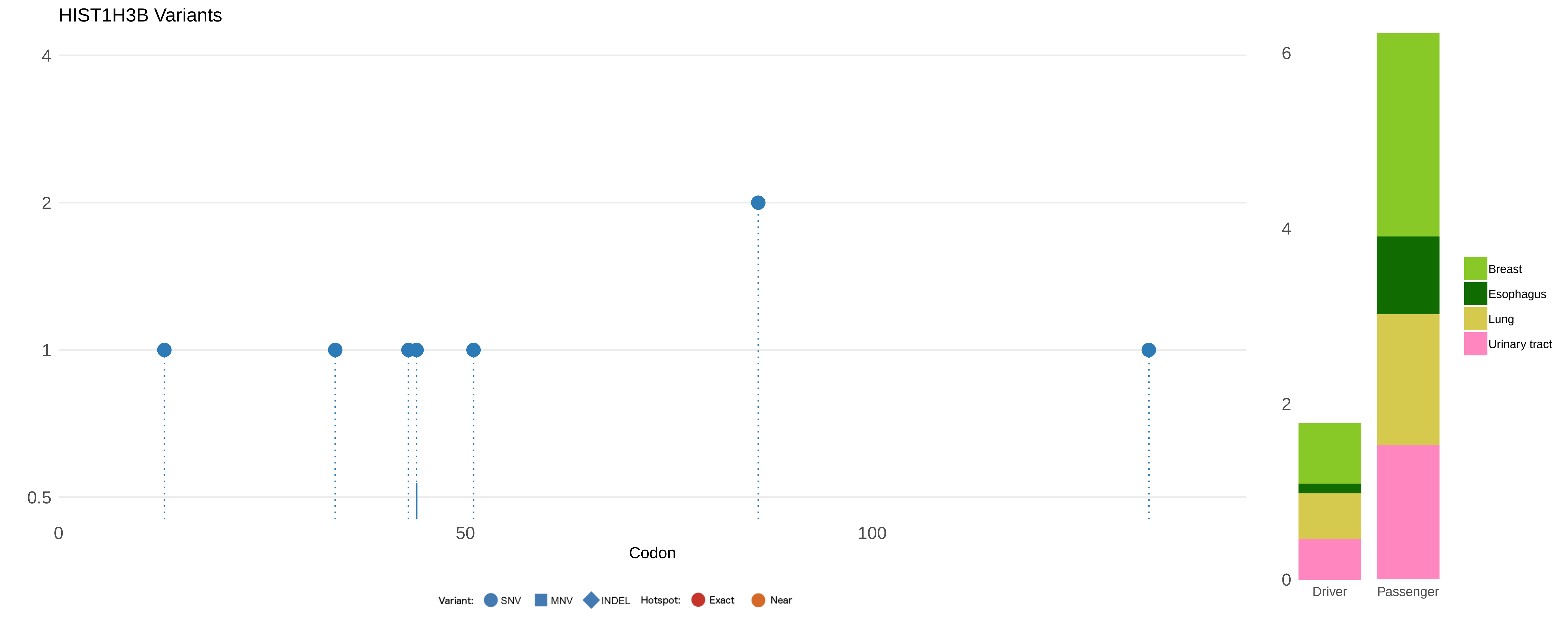

HIST1H3B Variants
4
2
1
0.5
0
50
100
Codon
Variant:
SNV
MNV
INDEL
Hotspot:
Exact
Near
6
4
2
0
Driver
Passenger
Breast
Esophagus
Lung
Urinary tract

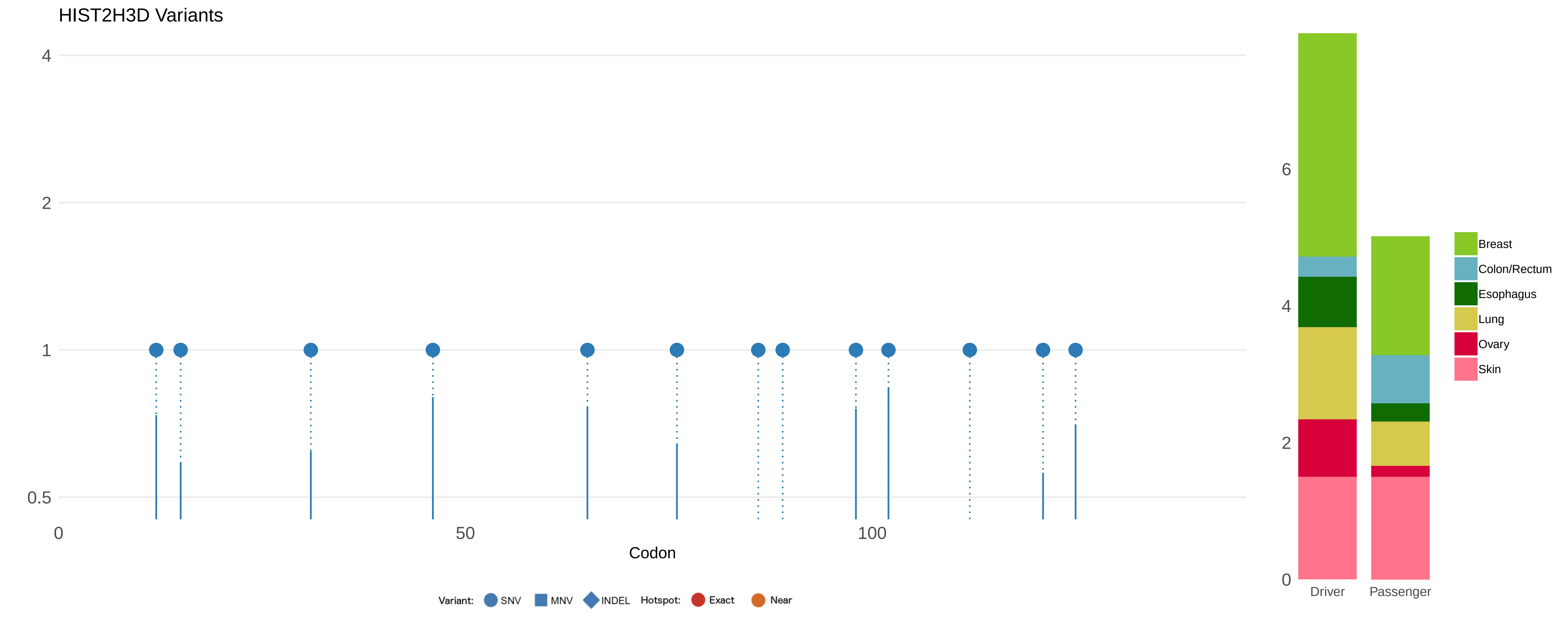

HIST2H3D Variants
4
2
1
0.5
0
50
100
Codon
Variant:
SNV
MNV
INDEL
Hotspot:
Exact
Near
6
4
2
0
Driver
Passenger
Breast
Colon/Rectum
Esophagus
Lung
Ovary
Skin

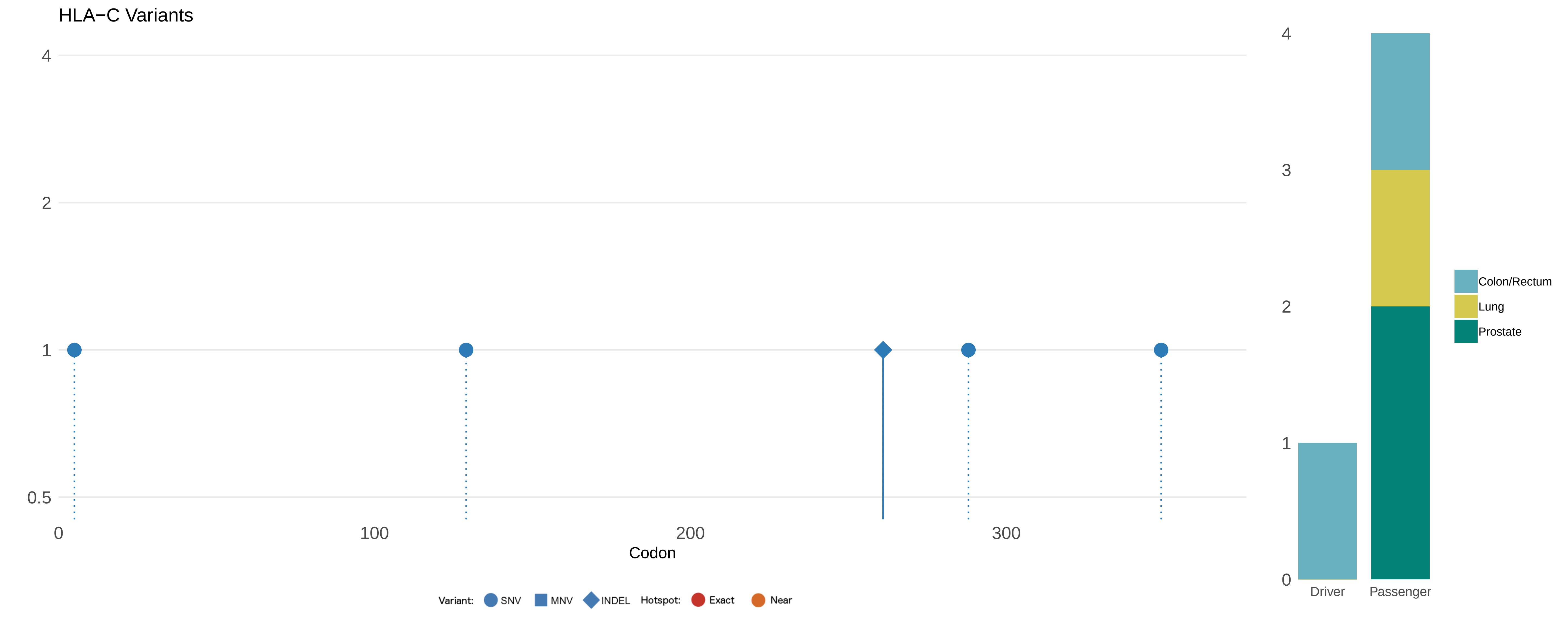
HLA−C Variants
4
2
1
0.5
0
100
200
300
Codon
Variant:
SNV
MNV
INDEL
Hotspot:
Exact
Near
4
3
2
1
0
Driver
Passenger
Colon/Rectum
Lung
Prostate

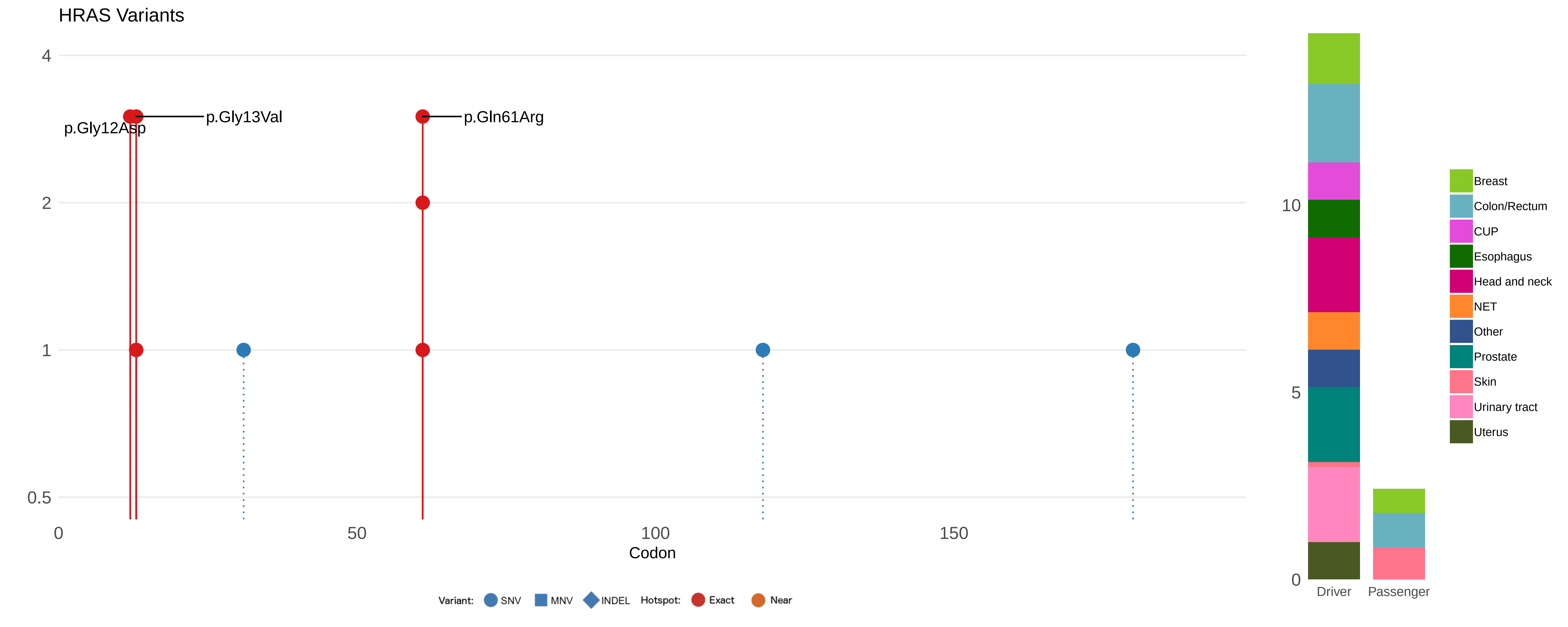
HRAS Variants
p.Gly12Asp
p.Gly13Val
p.Gln61Arg
4
2
1
0.5
0
50
100
150
Codon
Variant: SNV MNV INDEL Hotspot: Exact Near
10
5
0
Driver
Passenger
Breast
Colon/Rectum
CUP
Esophagus
Head and neck
NET
Other
Prostate
Skin
Urinary tract
Uterus

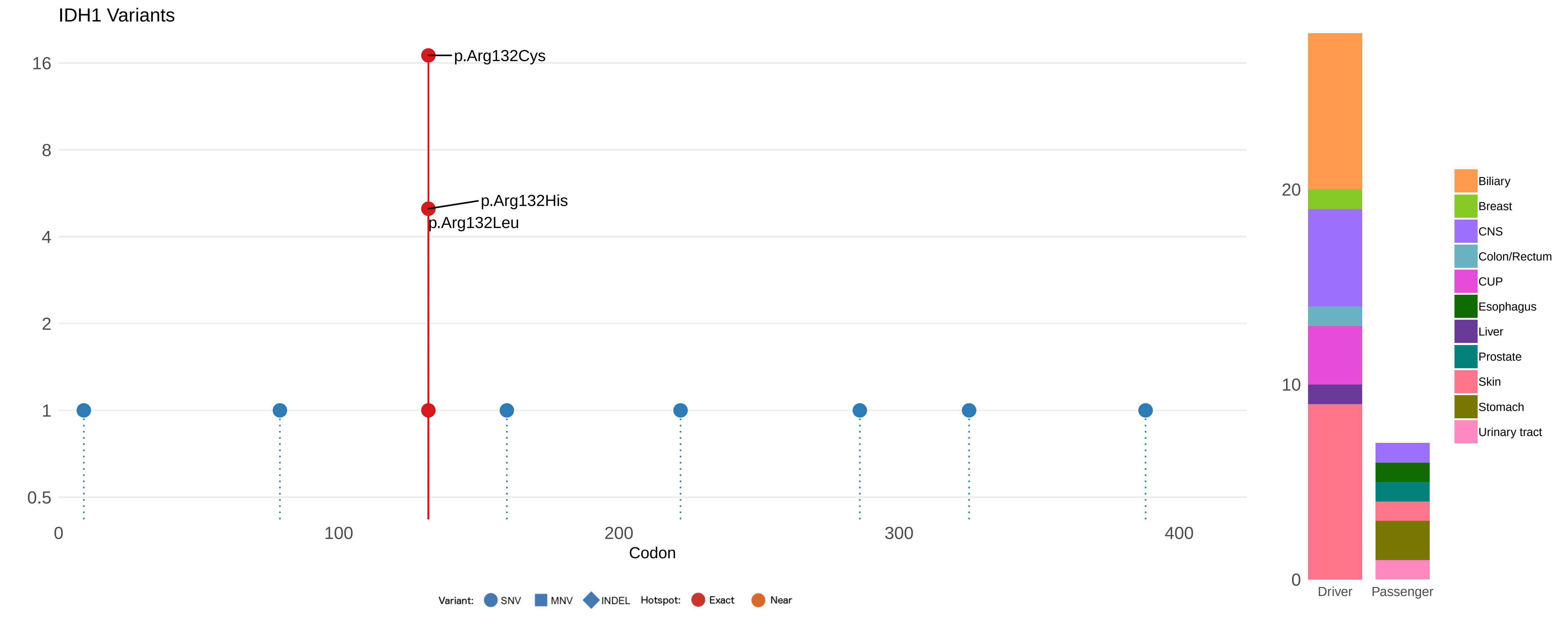

IDH1 Variants
p.Arg132Cys
p.Arg132His
p.Arg132Leu
16
8
4
2
1
0.5
0
100
200
300
400
Codon
Variant: SNV MNV INDEL Hotspot: Exact Near
20
10
0
Driver
Passenger
Biliary
Breast
CNS
Colon/Rectum
CUP
Esophagus
Liver
Prostate
Skin
Stomach
Urinary tract

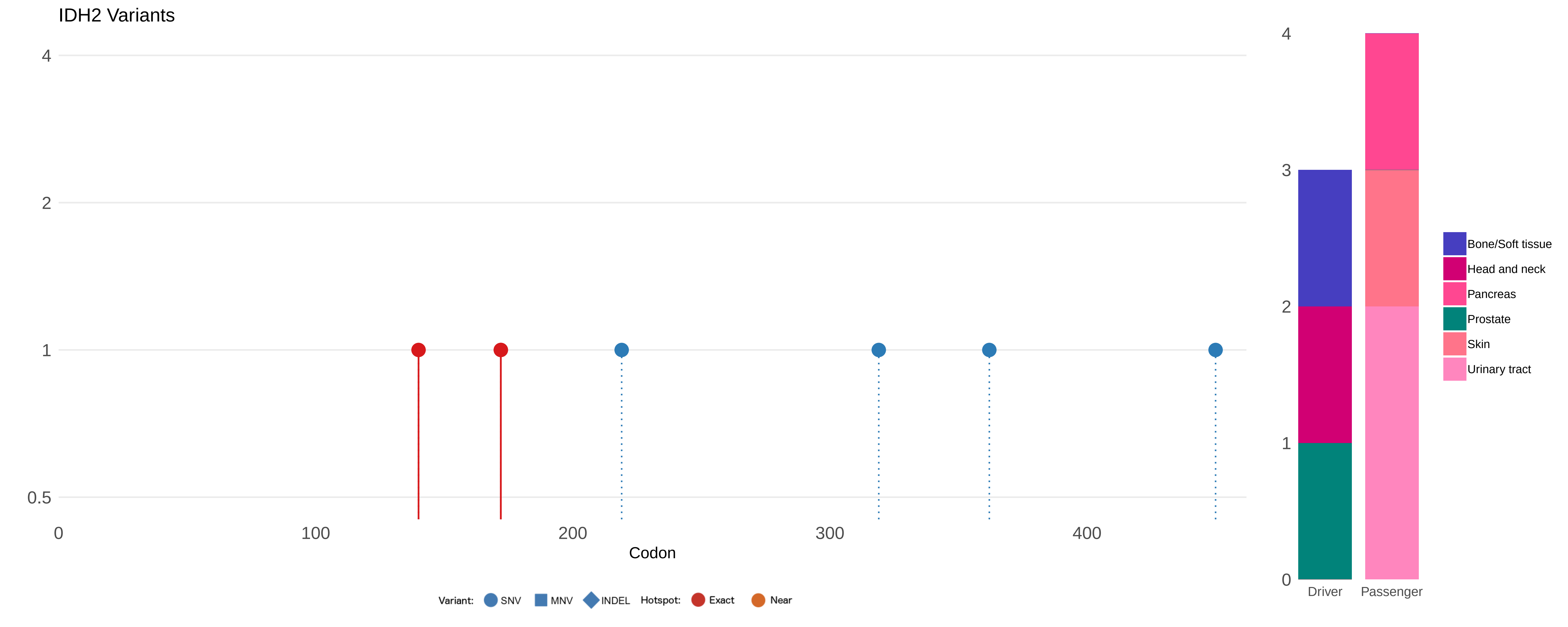
IDH2 Variants
4
2
1
0.5
0
100
200
300
400
Codon
Variant: SNV MNV INDEL Hotspot: Exact Near
4
3
2
1
0
Driver
Passenger
Bone/Soft tissue
Head and neck
Pancreas
Prostate
Skin
Urinary tract

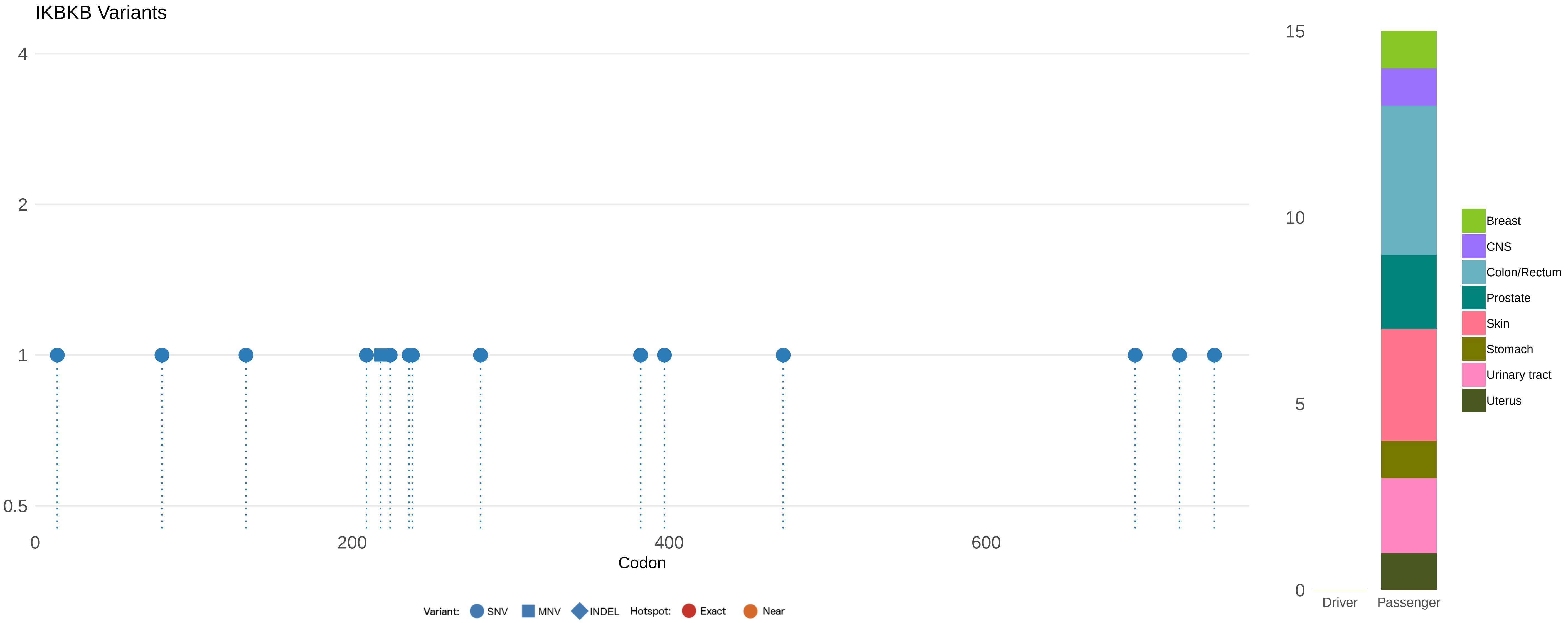

IKBKB Variants
4
2
1
0.5
0
200
400
600
Codon
Variant: SNV MNV INDEL Hotspot: Exact Near
15
10
5
0
Driver
Passenger
Breast
CNS
Colon/Rectum
Prostate
Skin
Stomach
Urinary tract
Uterus

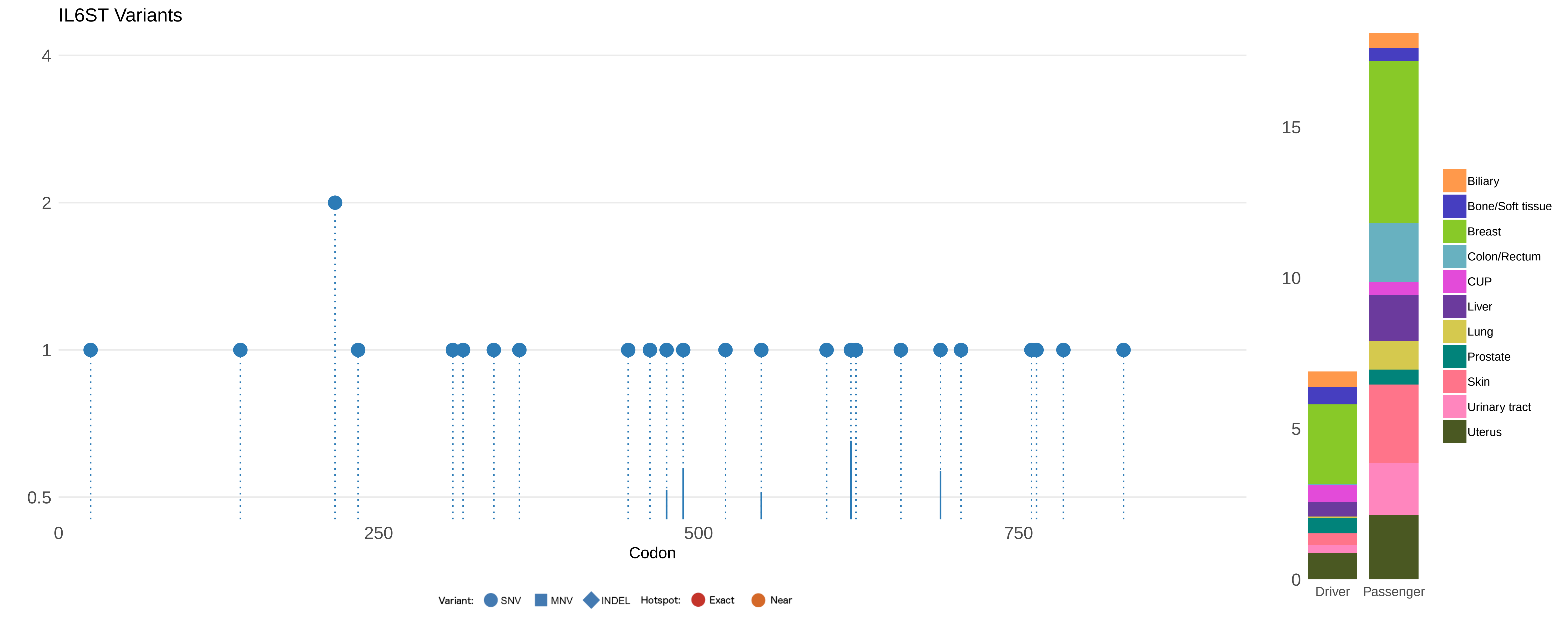
IL6ST Variants
4
2
1
0.5
0
250
500
750
Codon
Variant:
SNV
MNV
INDEL
Hotspot:
Exact
Near
15
10
5
0
Driver
Passenger
Biliary
Bone/Soft tissue
Breast
Colon/Rectum
CUP
Liver
Lung
Prostate
Skin
Urinary tract
Uterus

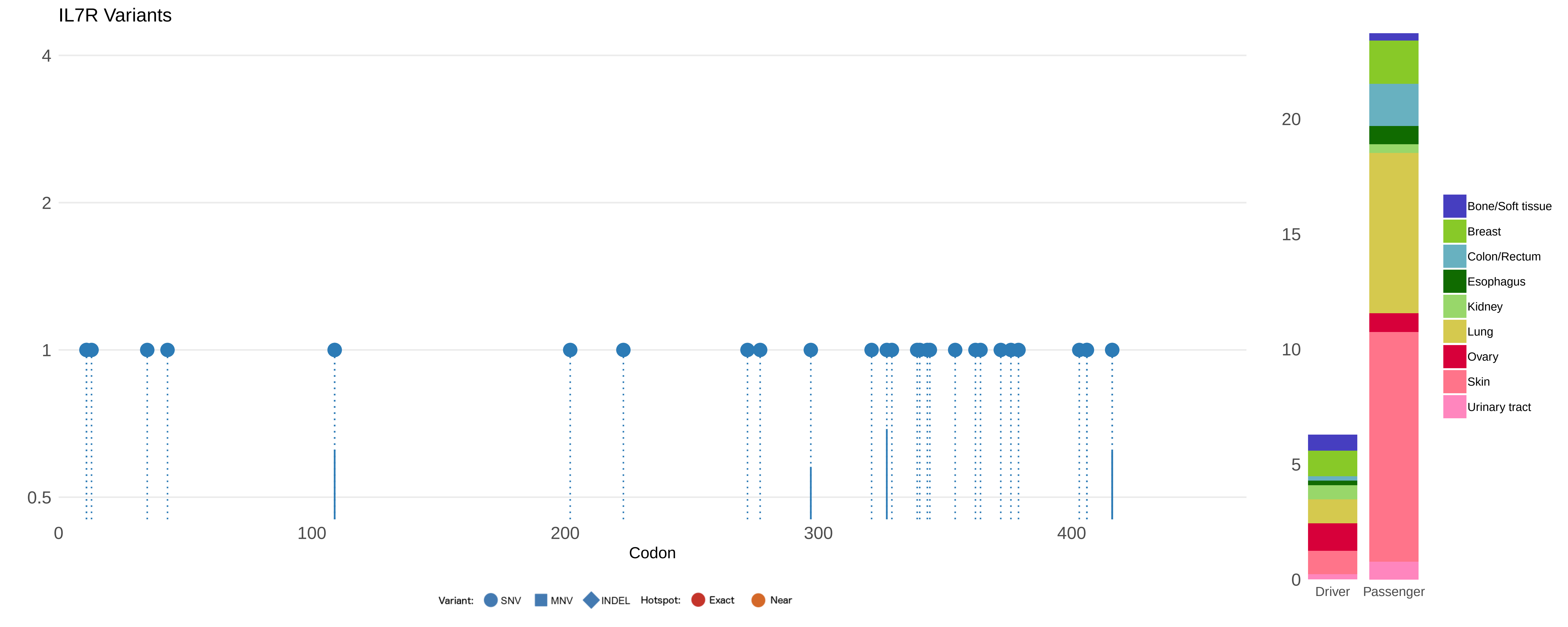

IL7R Variants
4
2
1
0.5
0
100
200
300
400
Codon
Variant:
SNV
MNV
INDEL
Hotspot:
Exact
Near
20
15
10
5
0
Driver
Passenger
Bone/Soft tissue
Breast
Colon/Rectum
Esophagus
Kidney
Lung
Ovary
Skin
Urinary tract

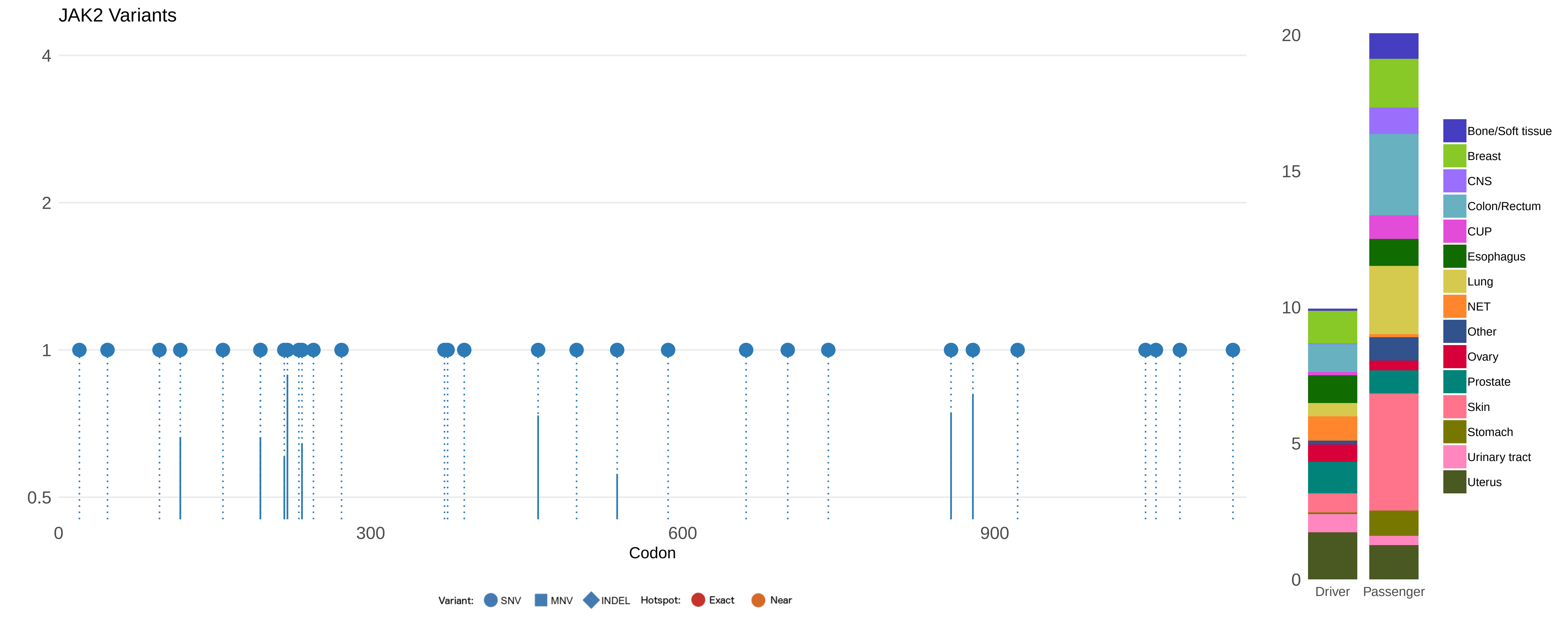

JAK2 Variants
4
2
1
0.5
0
300
600
900
Codon
Variant:
SNV
MNV
INDEL
Hotspot:
Exact
Near
20
15
10
5
0
Driver
Passenger
Bone/Soft tissue
Breast
CNS
Colon/Rectum
CUP
Esophagus
Lung
NET
Other
Ovary
Prostate
Skin
Stomach
Urinary tract
Uterus

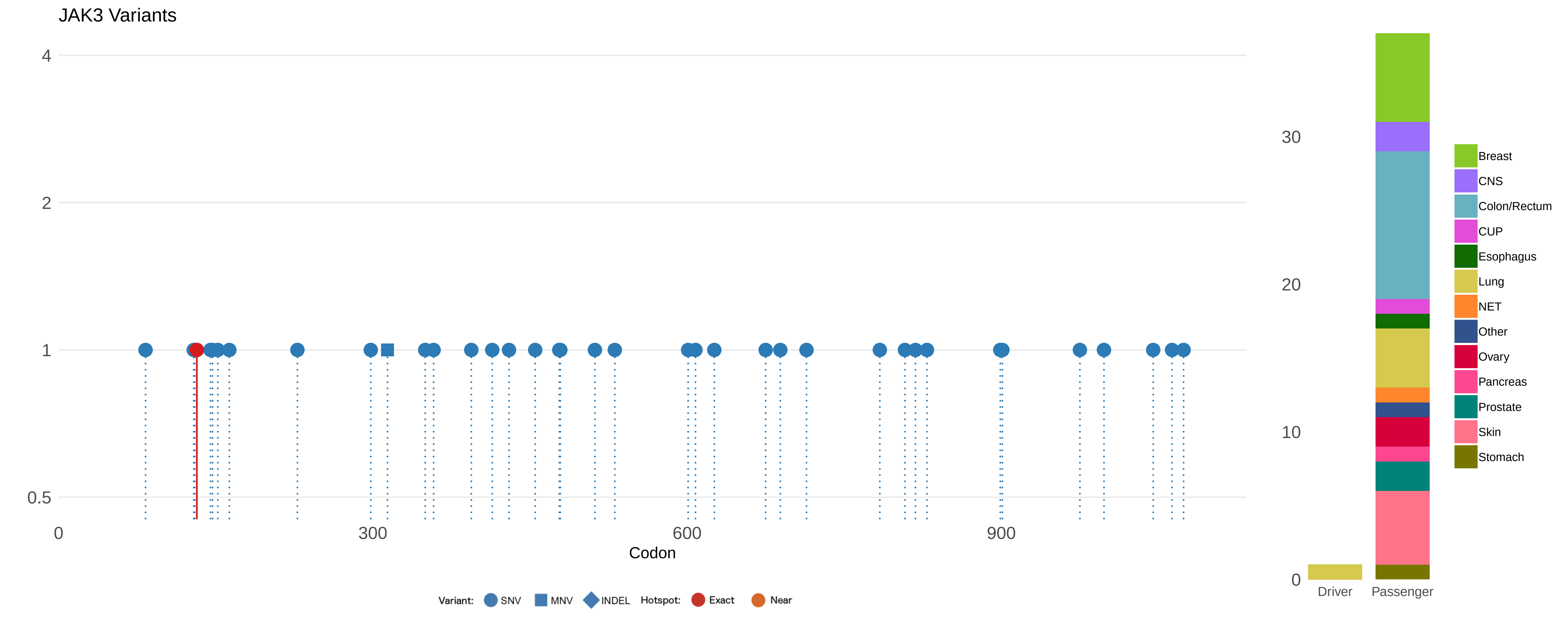

JAK3 Variants
4
2
1
0.5
0
300
600
900
Codon
Variant: SNV MNV INDEL Hotspot: Exact Near
30
20
10
0
Driver
Passenger
Breast
CNS
Colon/Rectum
CUP
Esophagus
Lung
NET
Other
Ovary
Pancreas
Prostate
Skin
Stomach

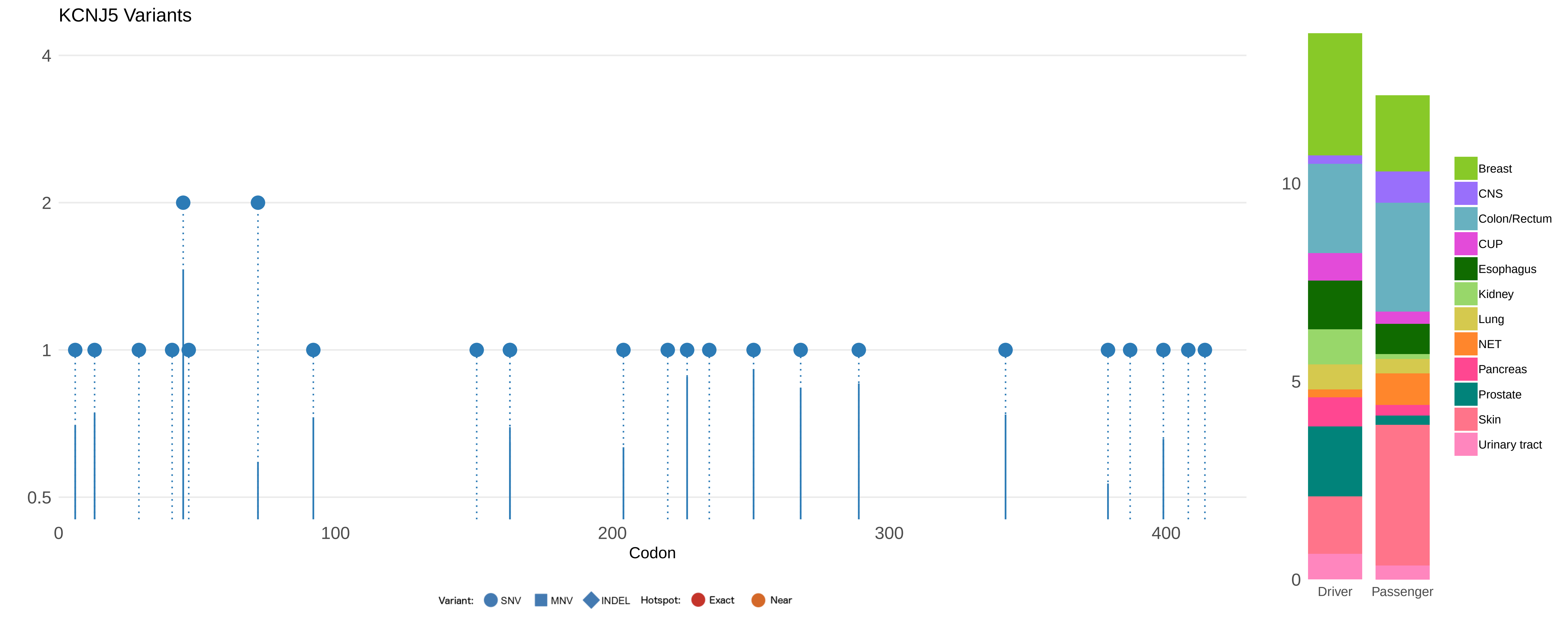

KCNJ5 Variants
4
2
1
0.5
0
100
200
300
400
Codon
Variant:
SNV
MNV
INDEL
Hotspot:
Exact
Near
10
5
0
Driver
Passenger
Breast
CNS
Colon/Rectum
CUP
Esophagus
Kidney
Lung
NET
Pancreas
Prostate
Skin
Urinary tract

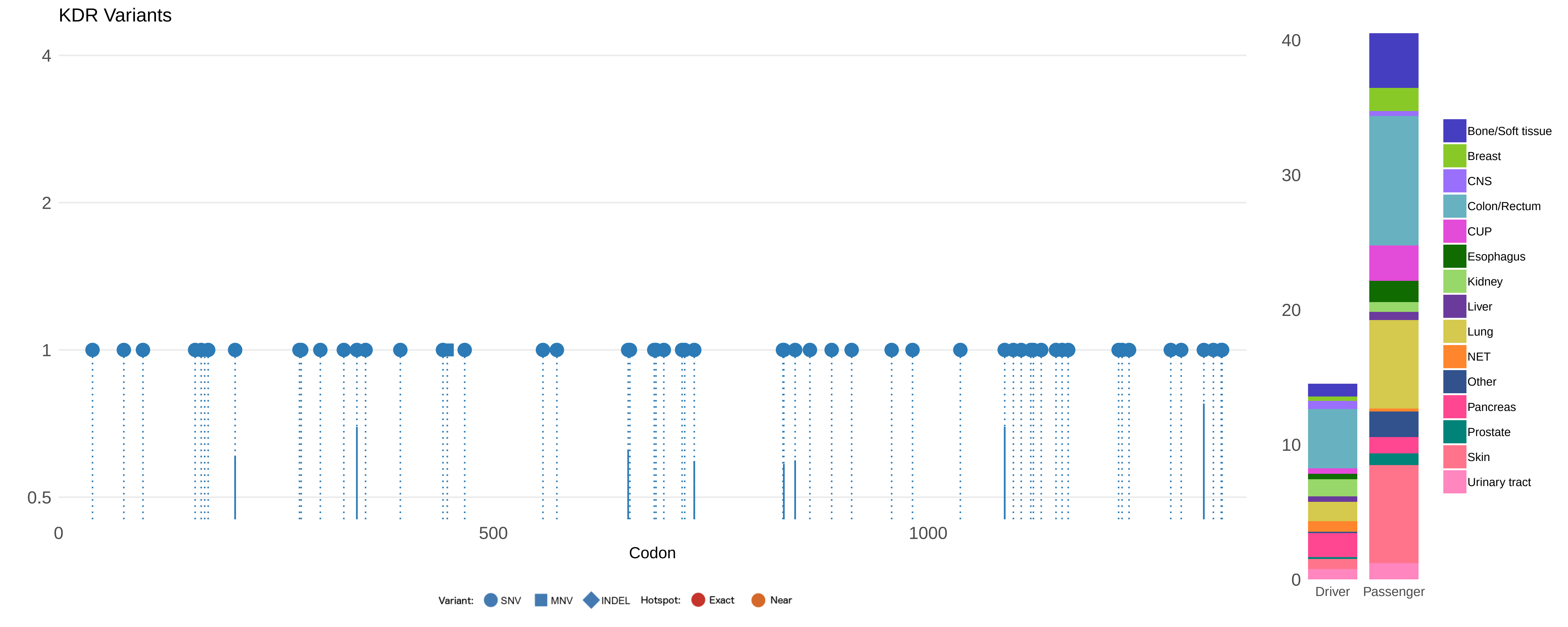

KDR Variants
4
2
1
0.5
0
500
1000
Codon
Variant:
SNV
MNV
INDEL
Hotspot:
Exact
Near
40
30
20
10
0
Driver
Passenger
Bone/Soft tissue
Breast
CNS
Colon/Rectum
CUP
Esophagus
Kidney
Liver
Lung
NET
Other
Pancreas
Prostate
Skin
Urinary tract

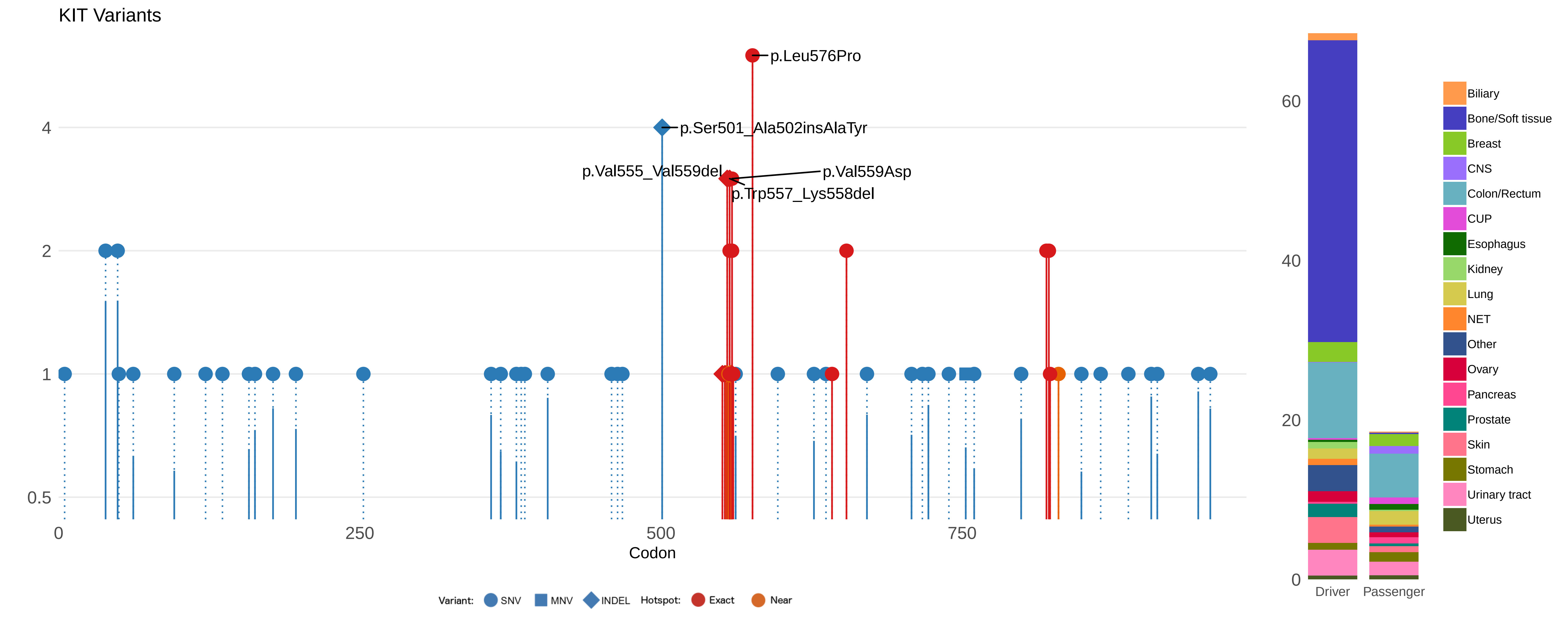

KIT Variants
p.Leu576Pro
p.Ser501_Ala502insAlaTyr
p.Val555_Val559del
p.Val559Asp
p.Trp557_Lys558del
4
2
1
0.5
0
250
500
750
Codon
Variant: SNV MNV INDEL Hotspot: Exact Near
60
40
20
0
Driver
Passenger
Biliary
Bone/Soft tissue
Breast
CNS
Colon/Rectum
CUP
Esophagus
Kidney
Lung
NET
Other
Ovary
Pancreas
Prostate
Skin
Stomach
Urinary tract
Uterus

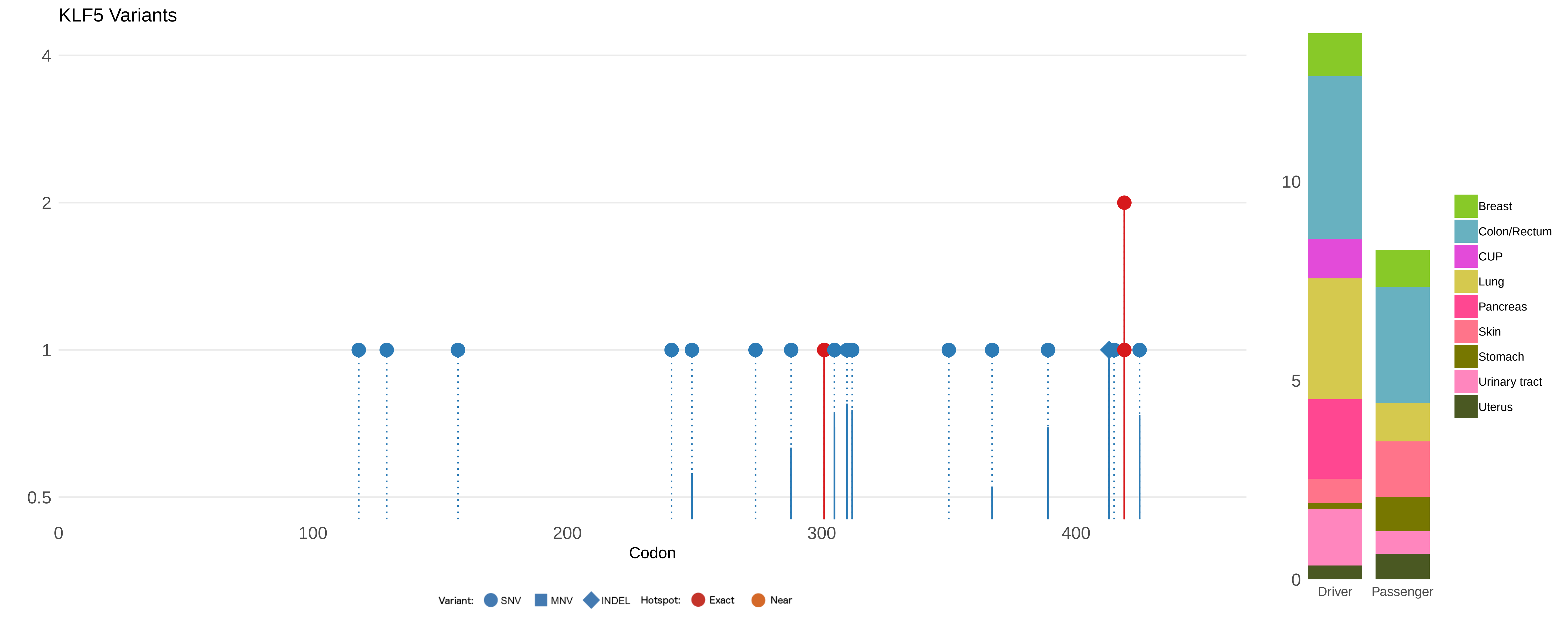

KLF5 Variants
4
2
1
0.5
0
100
200
300
400
Codon
Variant:
SNV
MNV
INDEL
Hotspot:
Exact
Near
10
5
0
Driver
Passenger
Breast
Colon/Rectum
CUP
Lung
Pancreas
Skin
Stomach
Urinary tract
Uterus

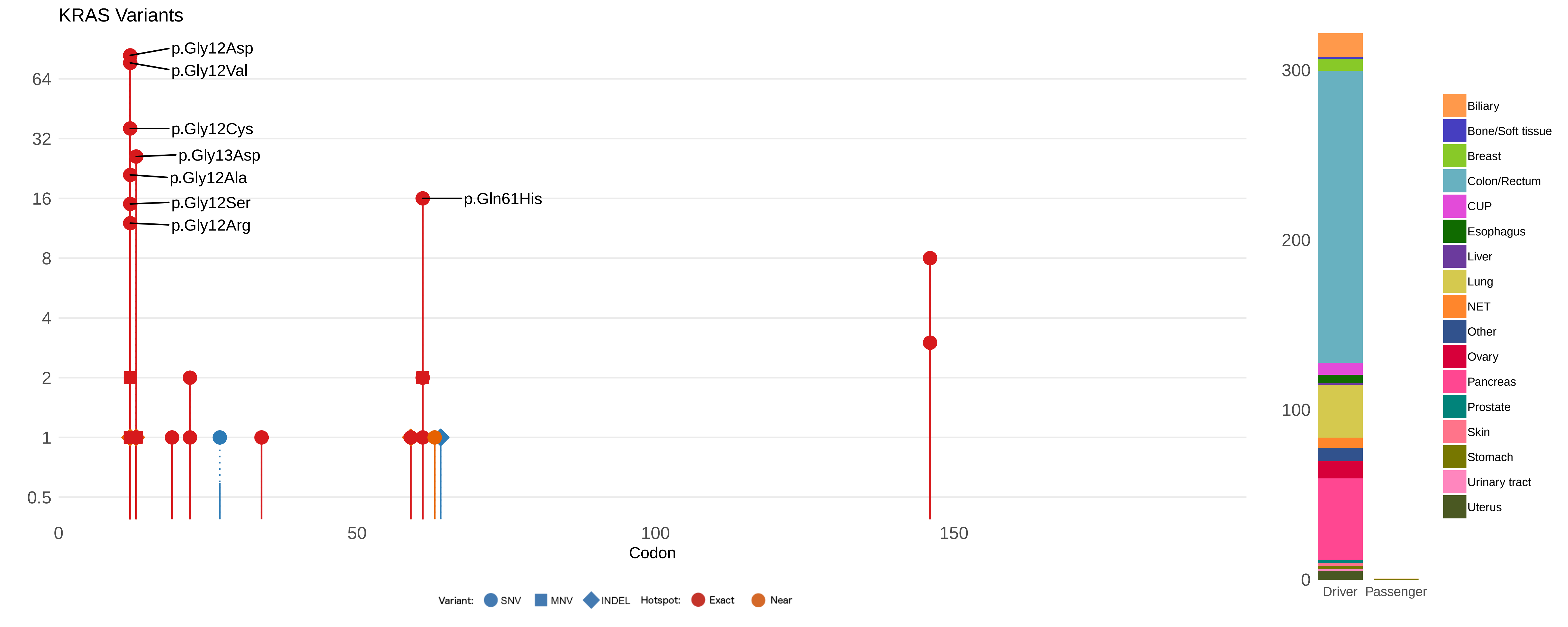

KRAS Variants
p.Gly12Asp
p.Gly12Val
p.Gly12Cys
p.Gly13Asp
p.Gly12Ala
p.Gly12Ser
p.Gly12Arg
p.Gln61His
64
32
16
8
4
2
1
0.5
0
50
100
150
Codon
Variant: SNV MNV INDEL Hotspot: Exact Near
300
200
100
0
Driver
Passenger
Biliary
Bone/Soft tissue
Breast
Colon/Rectum
CUP
Esophagus
Liver
Lung
NET
Other
Ovary
Pancreas
Prostate
Skin
Stomach
Urinary tract
Uterus

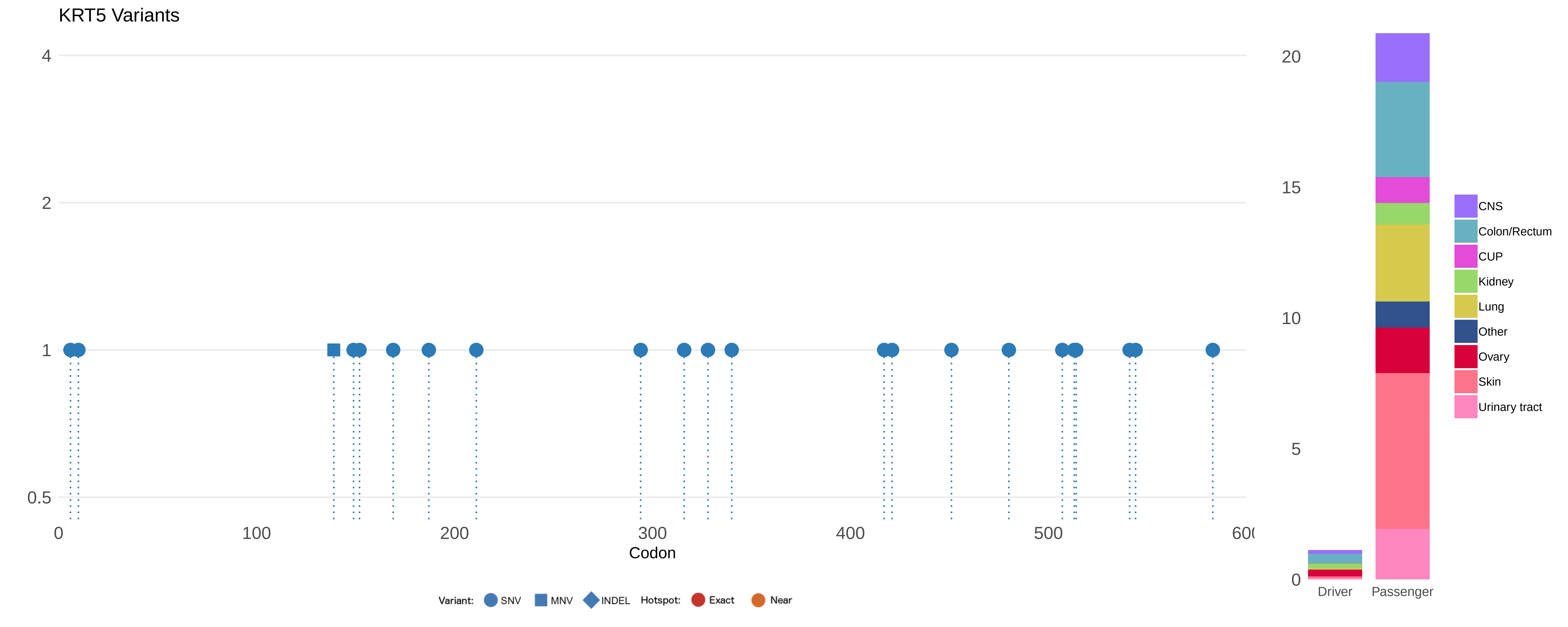

KRT5 Variants
4
2
1
0.5
0
100
200
300
400
500
600
Codon
Variant:
SNV
MNV
INDEL
Hotspot:
Exact
Near
20
15
10
5
0
Driver
Passenger
CNS
Colon/Rectum
CUP
Kidney
Lung
Other
Ovary
Skin
Urinary tract

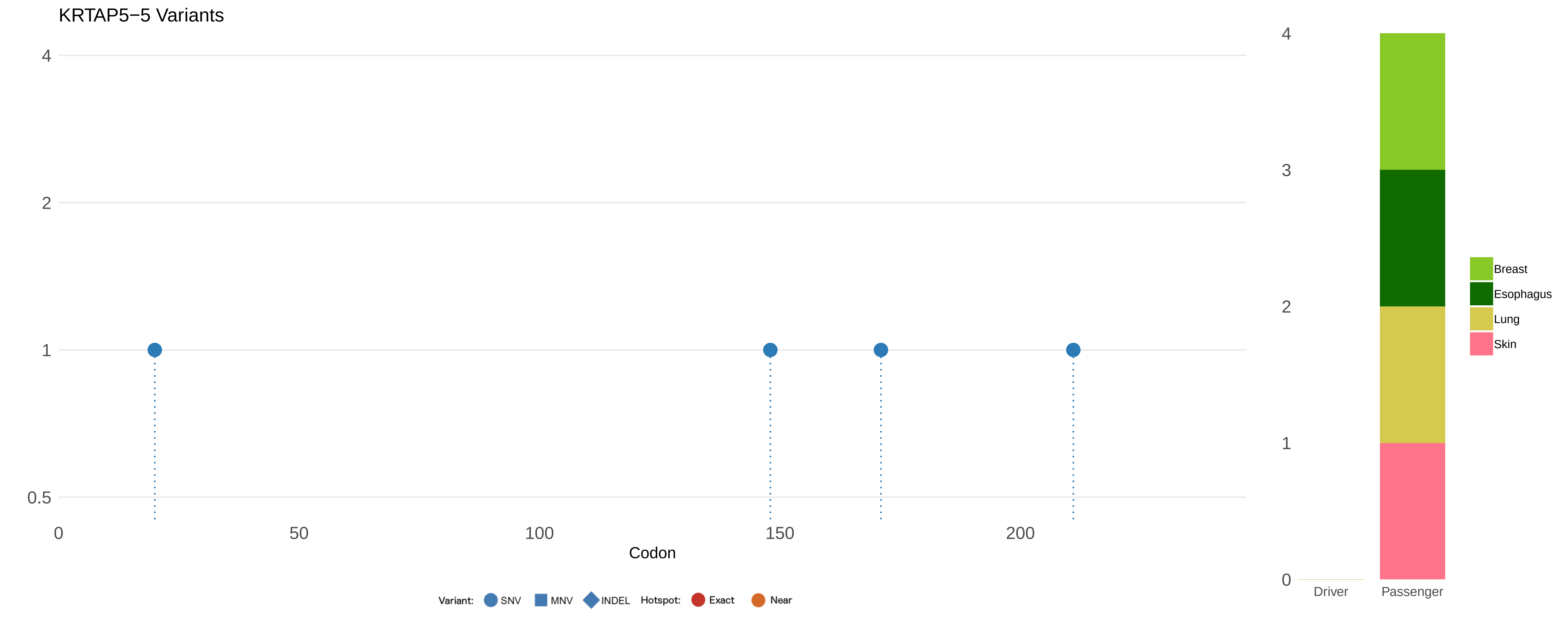

KRTAP5−5 Variants
4
2
1
0.5
0
50
100
150
200
Codon
Variant:
SNV
MNV
INDEL
Hotspot:
Exact
Near
4
3
2
1
0
Driver
Passenger
Breast
Esophagus
Lung
Skin

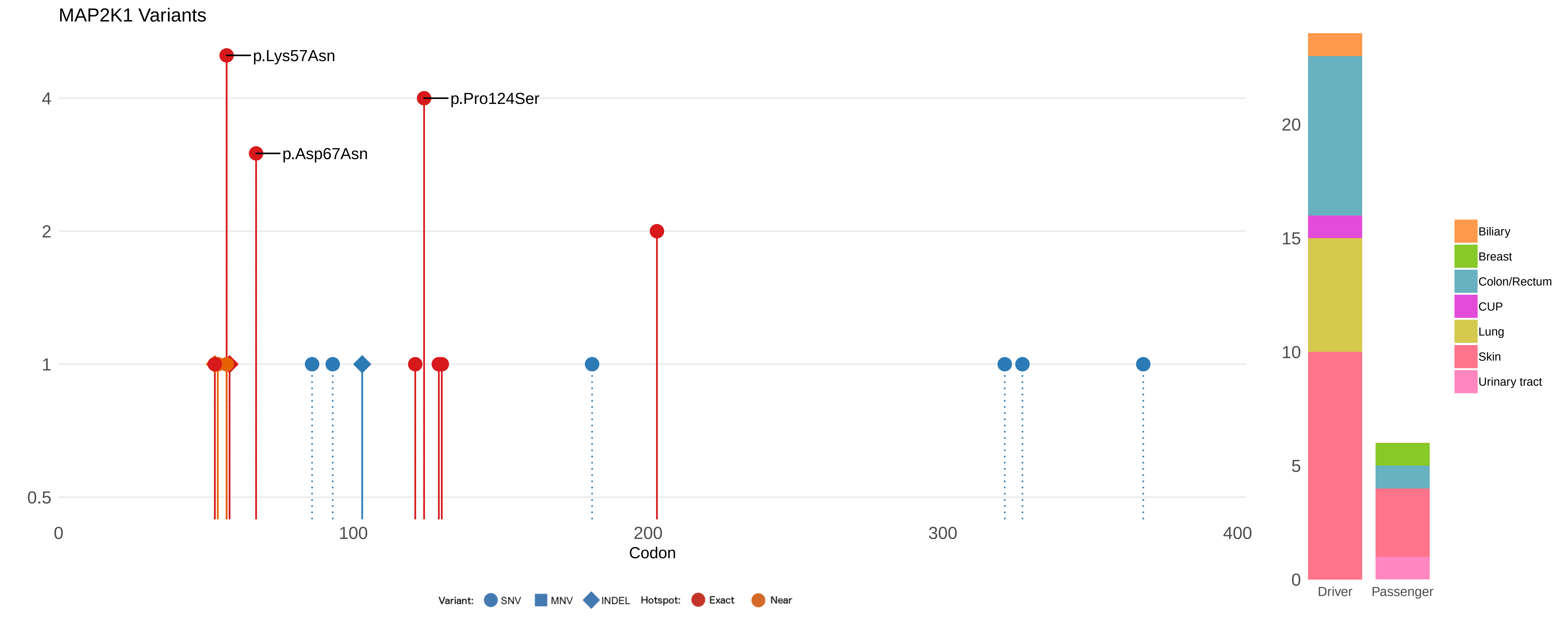

MAP2K1 Variants
p.Lys57Asn
p.Pro124Ser
p.Asp67Asn
4
2
1
0.5
0
100
200
300
400
Codon
Variant: SNV MNV INDEL Hotspot: Exact Near
20
15
10
5
0
Driver
Passenger
Biliary
Breast
Colon/Rectum
CUP
Lung
Skin
Urinary tract

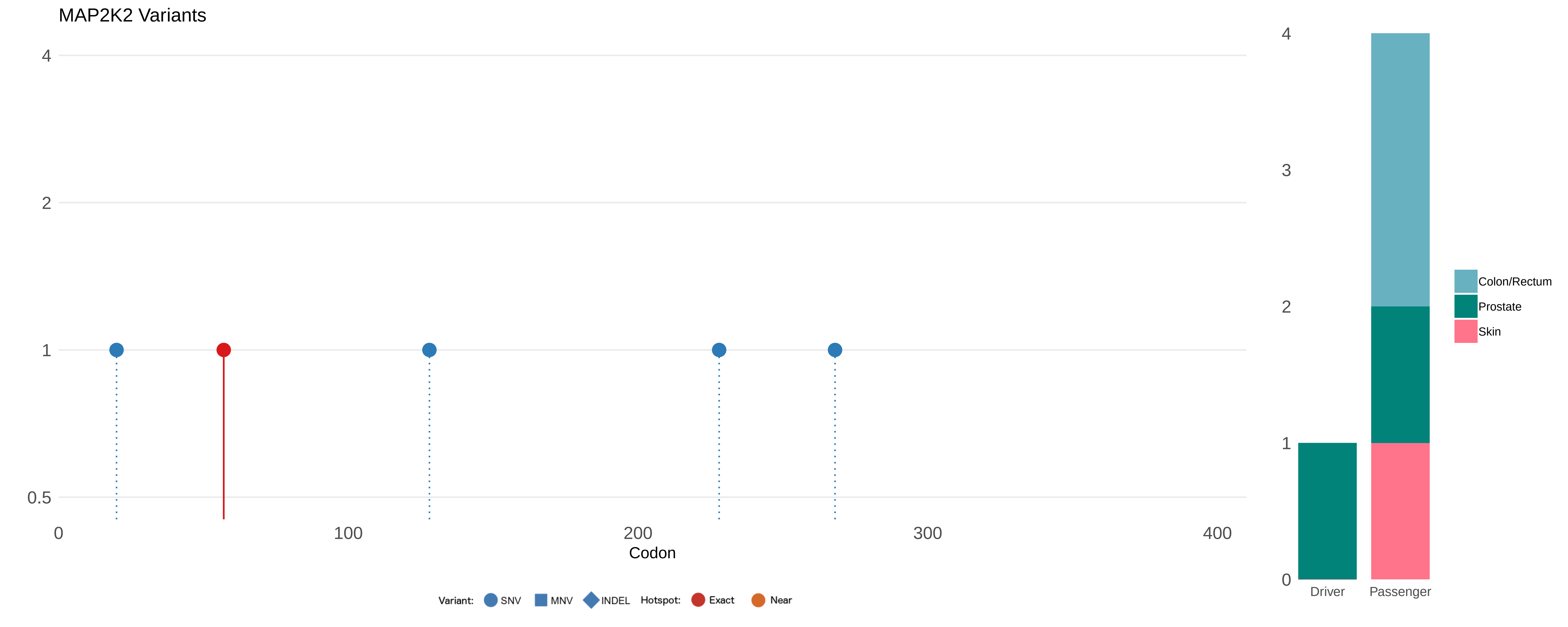

MAP2K2 Variants
4
2
1
0.5
0
100
200
300
400
Codon
Variant:
SNV
MNV
INDEL
Hotspot:
Exact
Near
4
3
2
1
0
Driver
Passenger
Colon/Rectum
Prostate
Skin

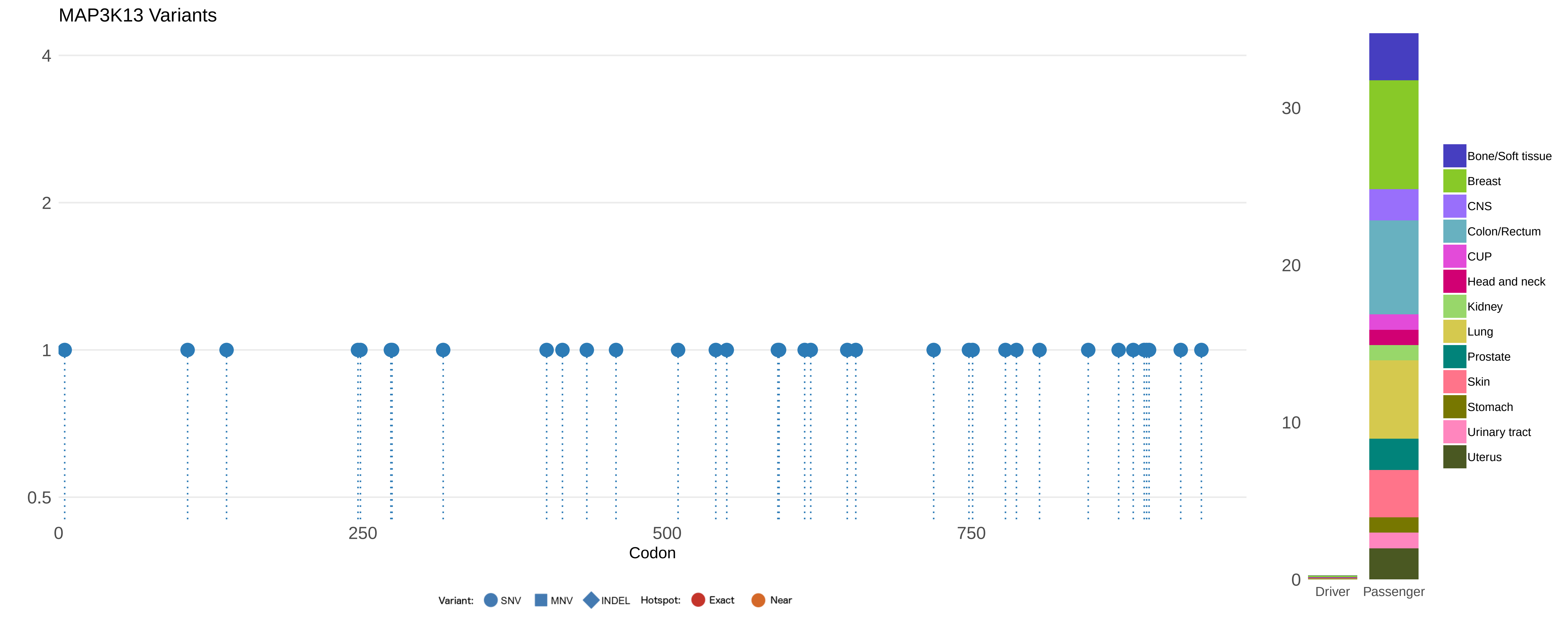

MAP3K13 Variants
4
2
1
0.5
0
250
500
750
Codon
Variant:
SNV
MNV
INDEL
Hotspot:
Exact
Near
30
20
10
0
Driver
Passenger
Bone/Soft tissue
Breast
CNS
Colon/Rectum
CUP
Head and neck
Kidney
Lung
Prostate
Skin
Stomach
Urinary tract
Uterus

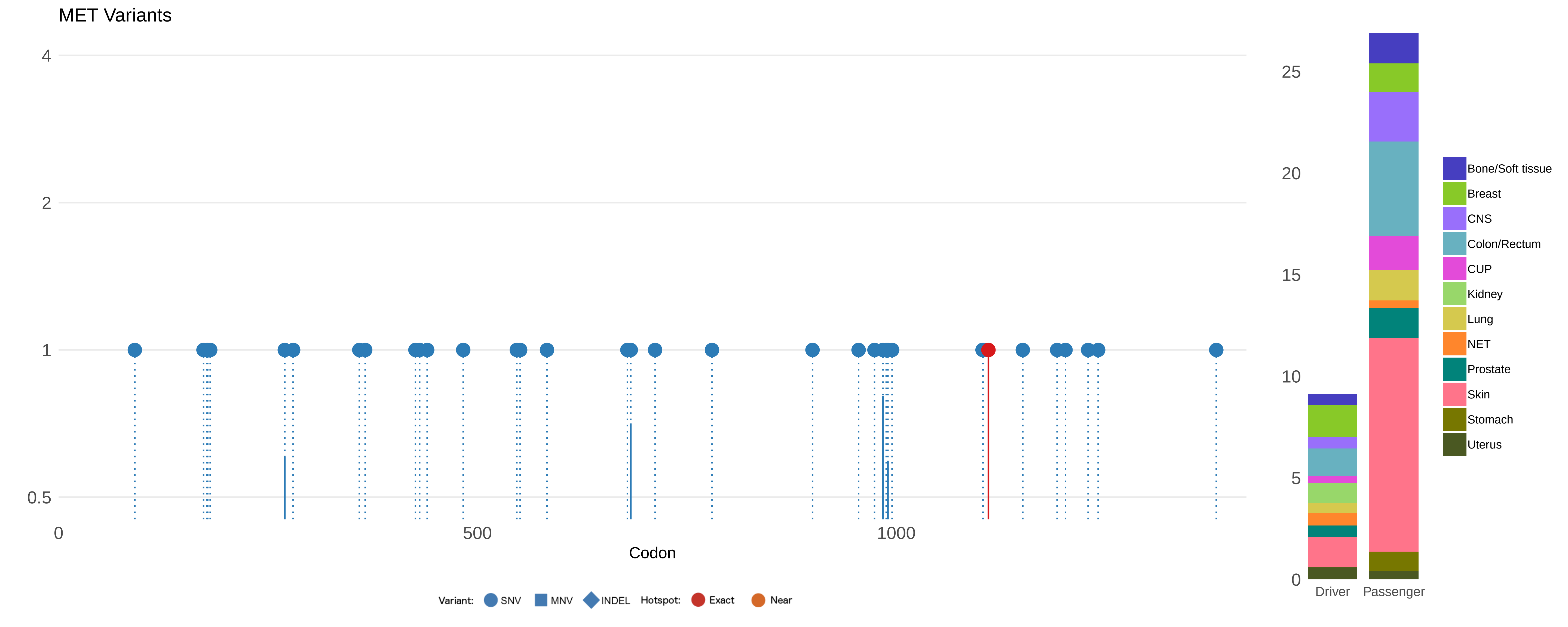

MET Variants
4
2
1
0.5
0
500
1000
Codon
Variant:
SNV
MNV
INDEL
Hotspot:
Exact
Near
25
20
15
10
5
0
Driver
Passenger
Bone/Soft tissue
Breast
CNS
Colon/Rectum
CUP
Kidney
Lung
NET
Prostate
Skin
Stomach
Uterus

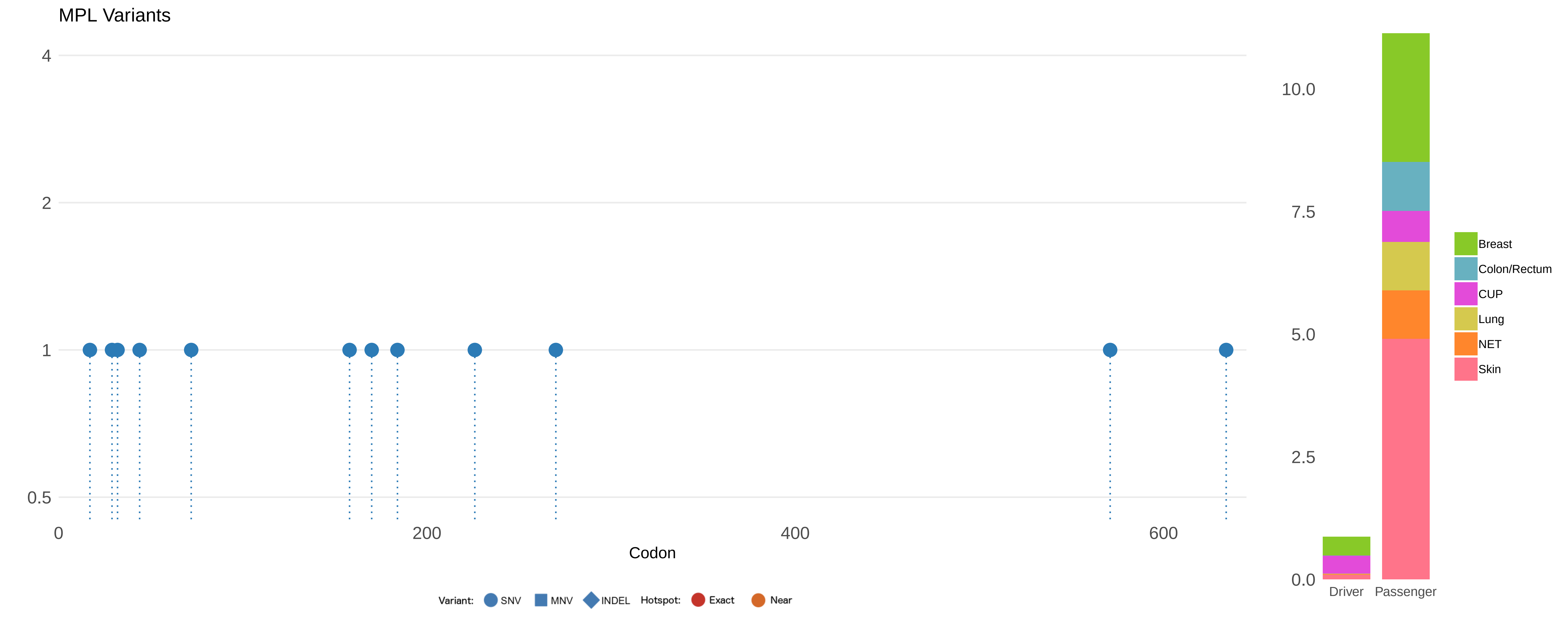

MPL Variants
4
2
1
0.5
0
200
400
600
Codon
Variant:
SNV
MNV
INDEL
Hotspot:
Exact
Near
10.0
7.5
5.0
2.5
0.0
Driver
Passenger
Breast
Colon/Rectum
CUP
Lung
NET
Skin

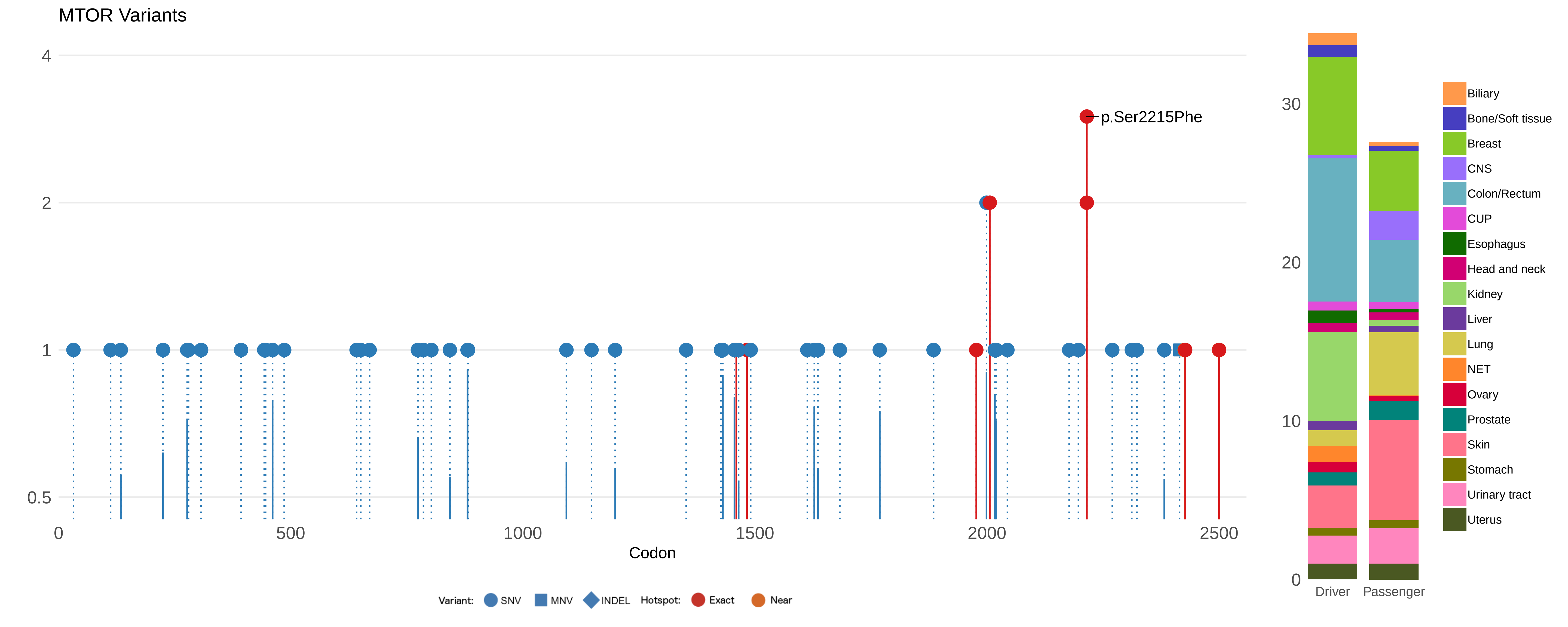

MTOR Variants
4
2
1
0.5
0
500
1000
1500
2000
2500
Codon
p.Ser2215Phe
Variant: SNV MNV INDEL Hotspot: Exact Near
30
20
10
0
Driver
Passenger
Biliary
Bone/Soft tissue
Breast
CNS
Colon/Rectum
CUP
Esophagus
Head and neck
Kidney
Liver
Lung
NET
Ovary
Prostate
Skin
Stomach
Urinary tract
Uterus

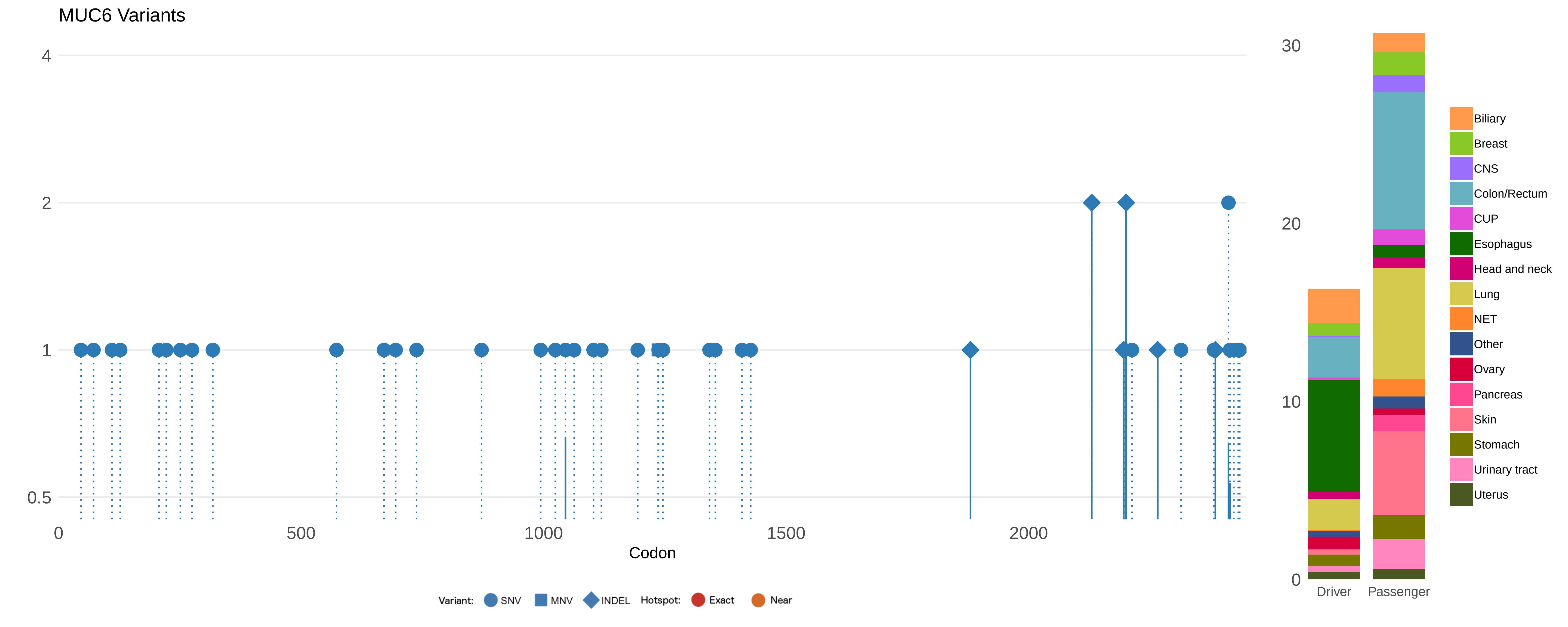
MUC6 Variants
4
2
1
0.5
0
500
1000
1500
2000
Codon
Variant:
SNV
MNV
INDEL
Hotspot:
Exact
Near
30
20
10
0
Driver
Passenger
Biliary
Breast
CNS
Colon/Rectum
CUP
Esophagus
Head and neck
Lung
NET
Other
Ovary
Pancreas
Skin
Stomach
Urinary tract
Uterus

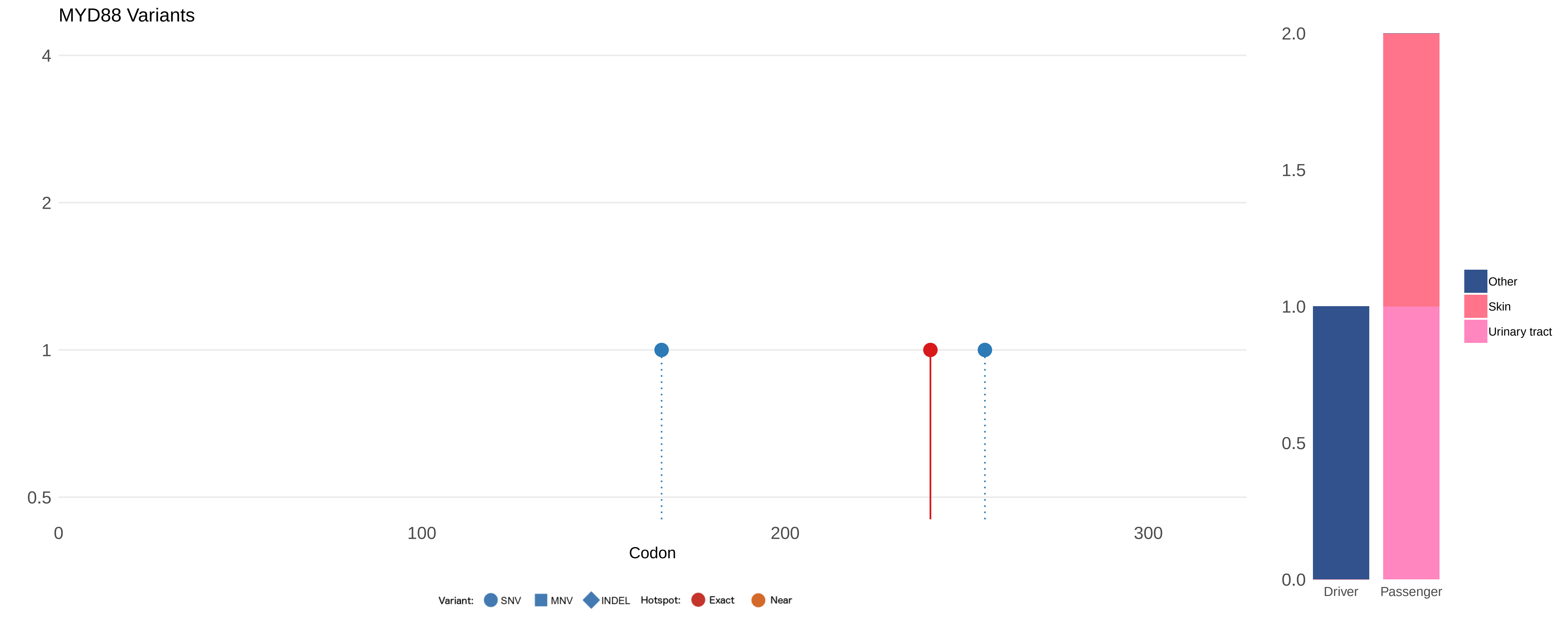

MYD88 Variants
4
2
1
0.5
0
100
200
300
Codon
Variant:
SNV
MNV
INDEL
Hotspot:
Exact
Near
2.0
1.5
1.0
0.5
0.0
Driver
Passenger
Other
Skin
Urinary tract

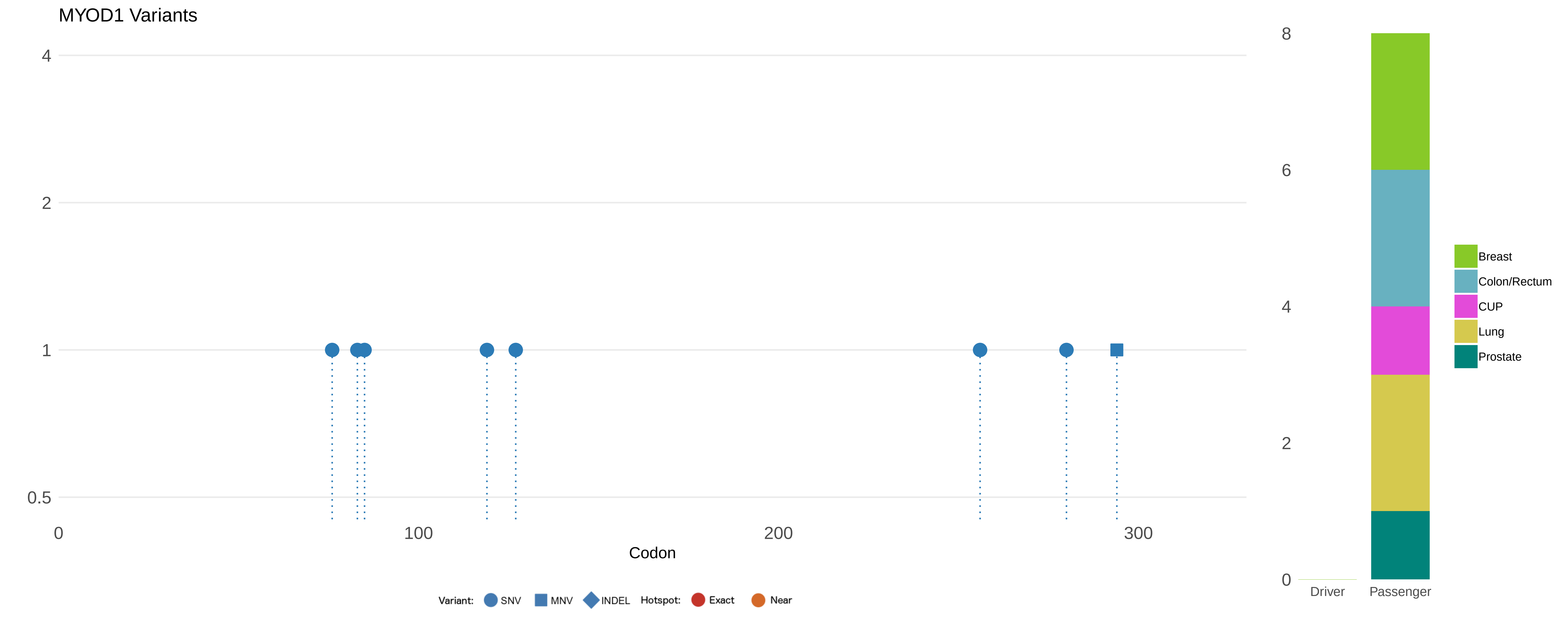
MYOD1 Variants
4
2
1
0.5
0
100
200
300
Codon
Variant: SNV MNV INDEL Hotspot: Exact Near
8
6
4
2
0
Driver
Passenger
Breast
Colon/Rectum
CUP
Lung
Prostate

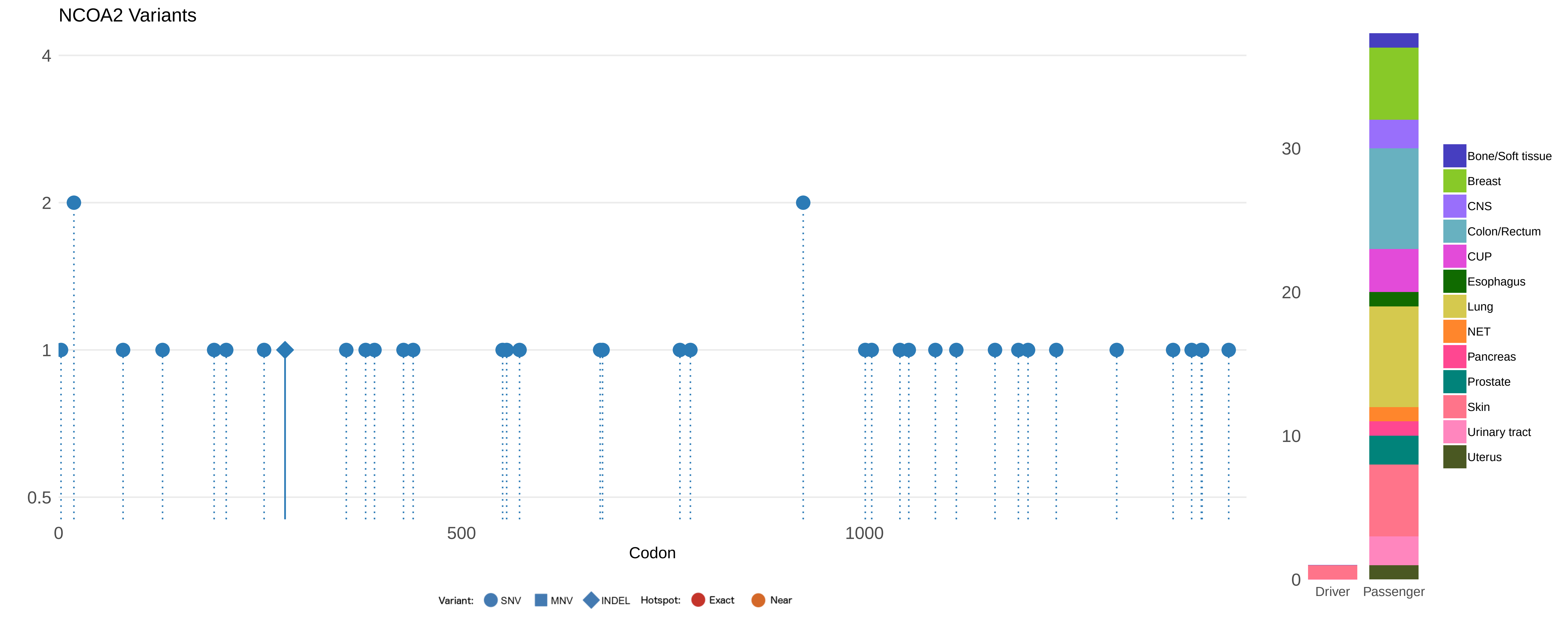

NCOA2 Variants
4
2
1
0.5
0
500
1000
Codon
Variant: SNV MNV INDEL Hotspot: Exact Near
30
20
10
0
Driver
Passenger
Bone/Soft tissue
Breast
CNS
Colon/Rectum
CUP
Esophagus
Lung
NET
Pancreas
Prostate
Skin
Urinary tract
Uterus

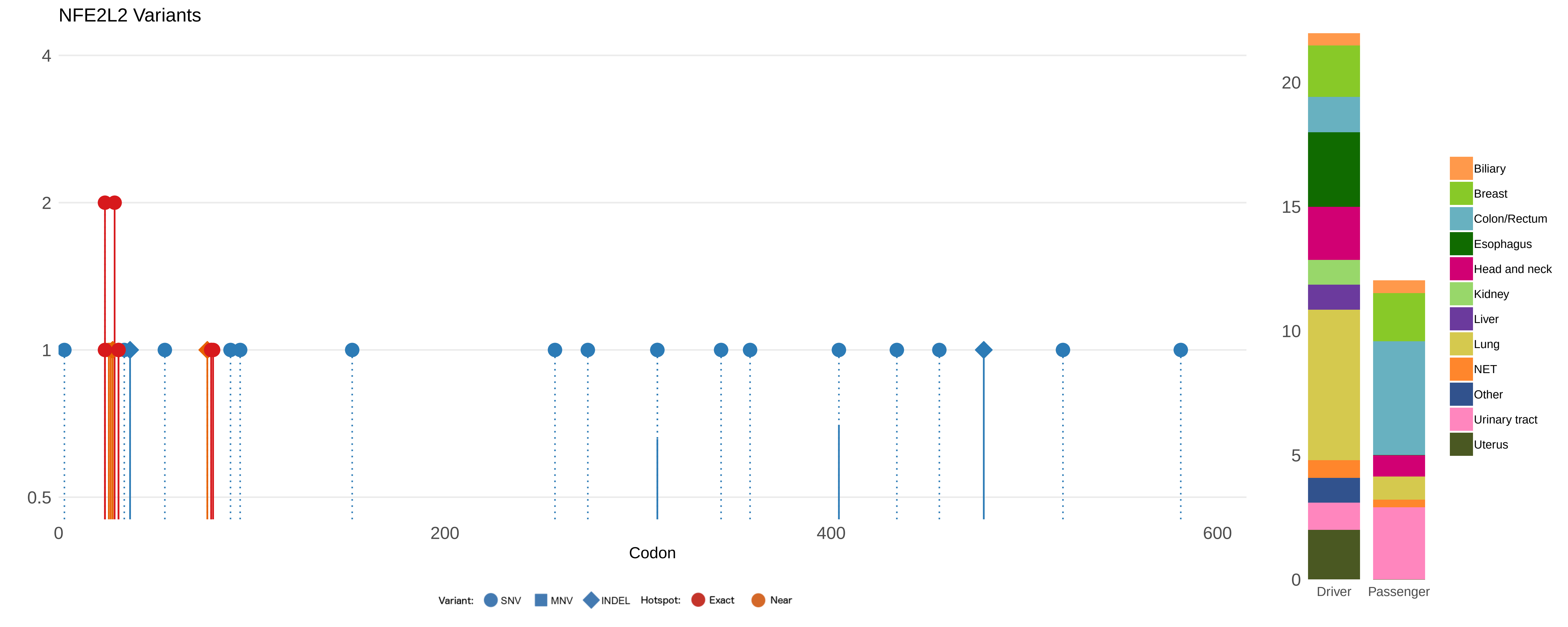

NFE2L2 Variants
4
2
1
0.5
0
200
400
600
Codon
Variant: SNV MNV INDEL Hotspot: Exact Near
20
15
10
5
0
Driver
Passenger
Biliary
Breast
Colon/Rectum
Esophagus
Head and neck
Kidney
Liver
Lung
NET
Other
Urinary tract
Uterus

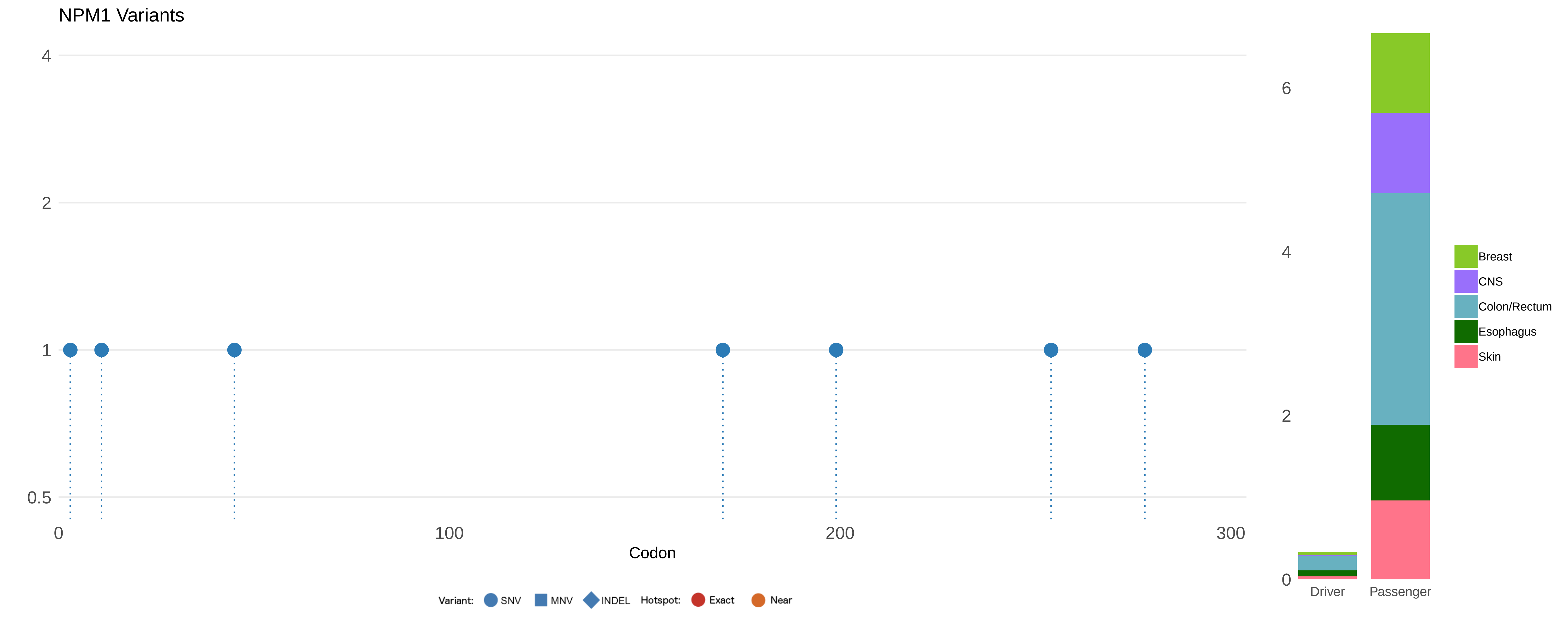

NPM1 Variants
4
2
1
0.5
0
100
200
300
Codon
Variant:
SNV
MNV
INDEL
Hotspot:
Exact
Near
6
4
2
0
Driver
Passenger
Breast
CNS
Colon/Rectum
Esophagus
Skin

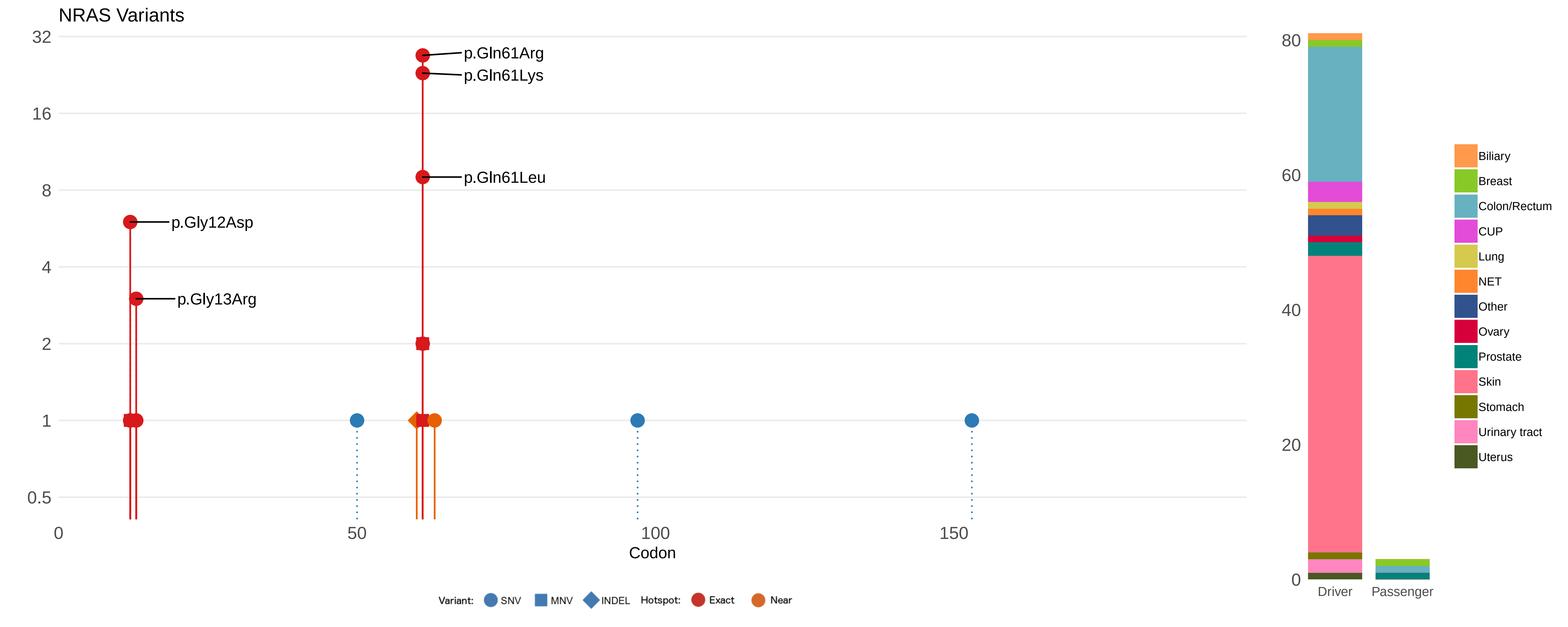

NRAS Variants
32
16
8
4
2
1
0.5
p.Gln61Arg
p.Gln61Lys
p.Gln61Leu
p.Gly12Asp
p.Gly13Arg
0
50
100
150
Codon
Variant: SNV MNV INDEL Hotspot: Exact Near
80
60
40
20
0
Driver
Passenger
Biliary
Breast
Colon/Rectum
CUP
Lung
NET
Other
Ovary
Prostate
Skin
Stomach
Urinary tract
Uterus

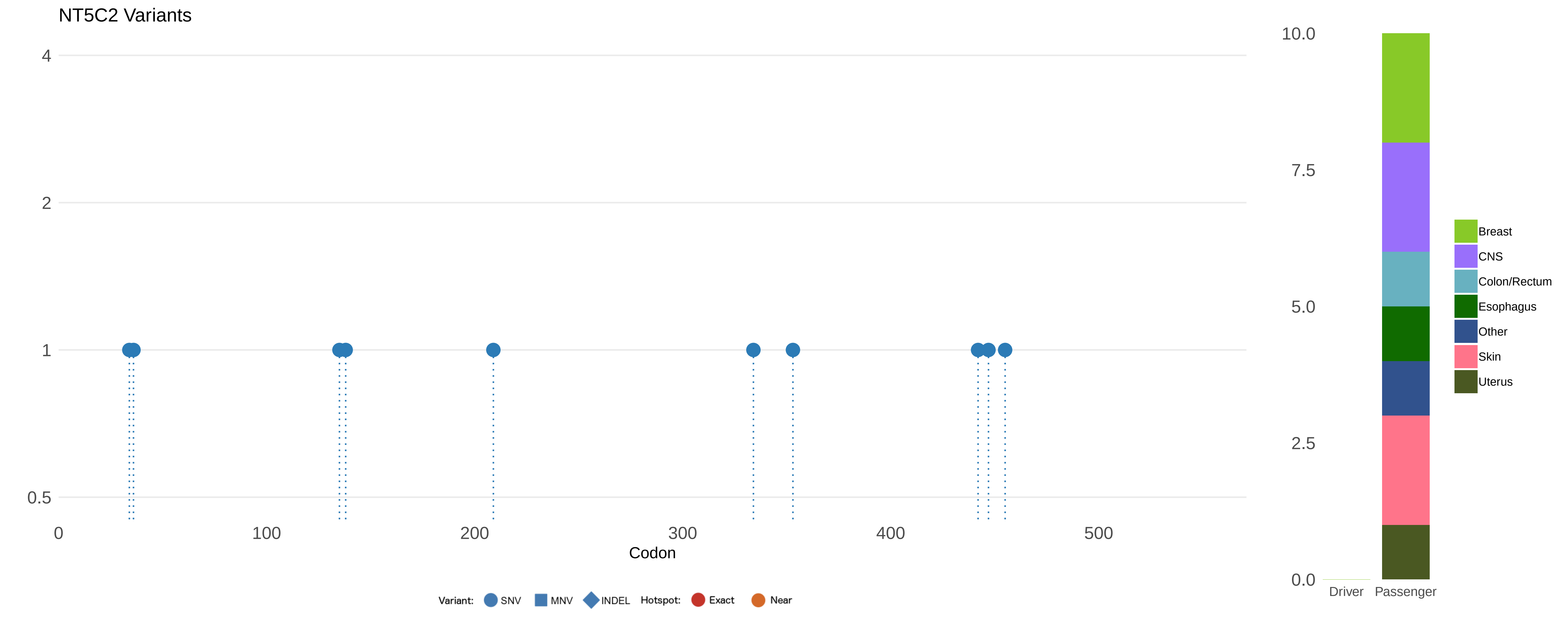

NT5C2 Variants
4
2
1
0.5
0
100
200
300
400
500
Codon
Variant:
SNV
MNV
INDEL
Hotspot:
Exact
Near
10.0
7.5
5.0
2.5
0.0
Driver
Passenger
Breast
CNS
Colon/Rectum
Esophagus
Other
Skin
Uterus

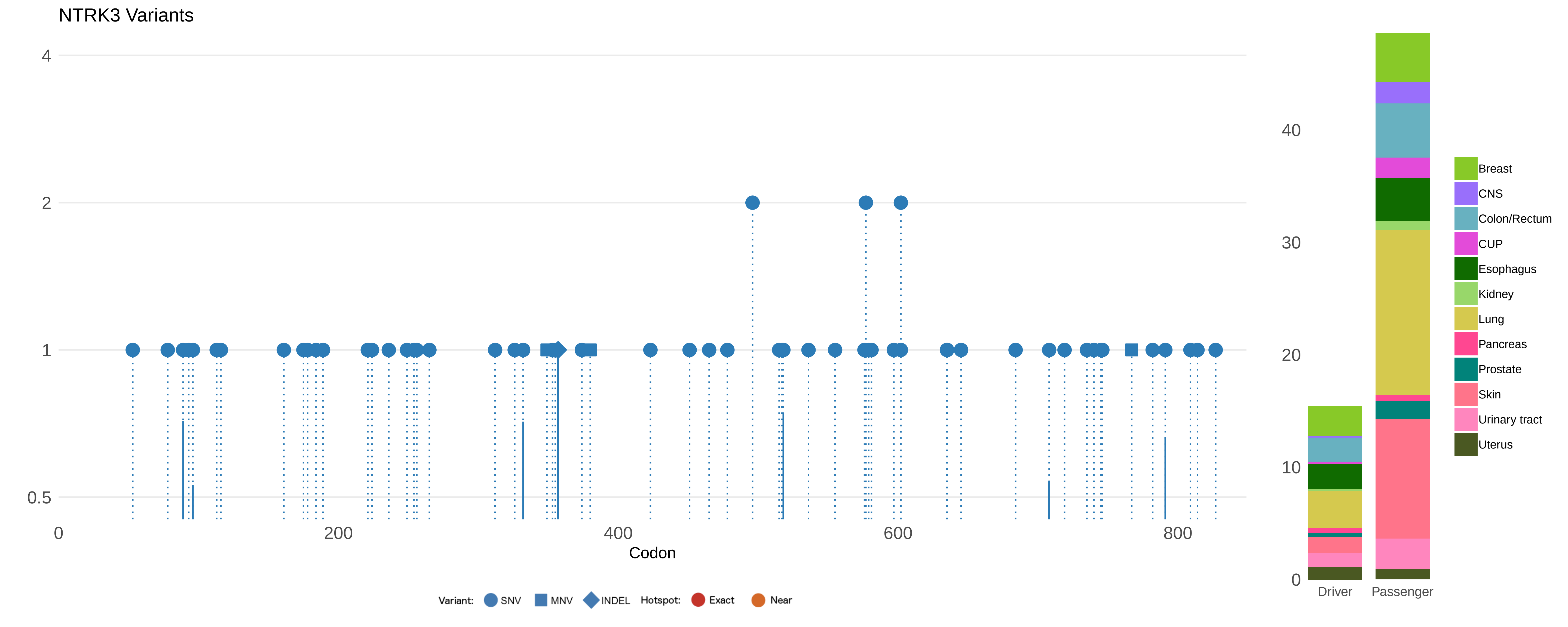

NTRK3 Variants
4
2
1
0.5
0
200
400
600
800
Codon
Variant: SNV MNV INDEL Hotspot: Exact Near
40
30
20
10
0
Driver
Passenger
Breast
CNS
Colon/Rectum
CUP
Esophagus
Kidney
Lung
Pancreas
Prostate
Skin
Urinary tract
Uterus

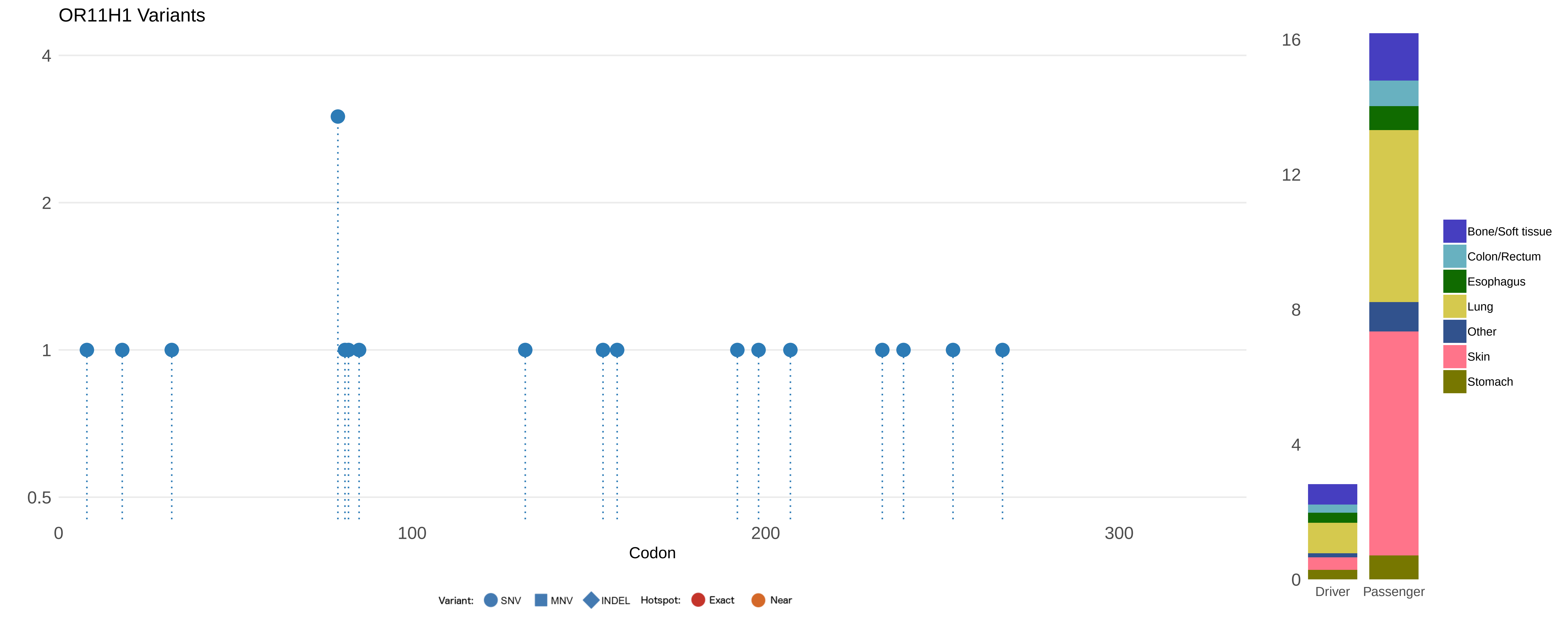
OR11H1 Variants
4
2
1
0.5
0
100
200
300
Codon
Variant:
SNV
MNV
INDEL
Hotspot:
Exact
Near
16
12
8
4
0
Driver
Passenger
Bone/Soft tissue
Colon/Rectum
Esophagus
Lung
Other
Skin
Stomach

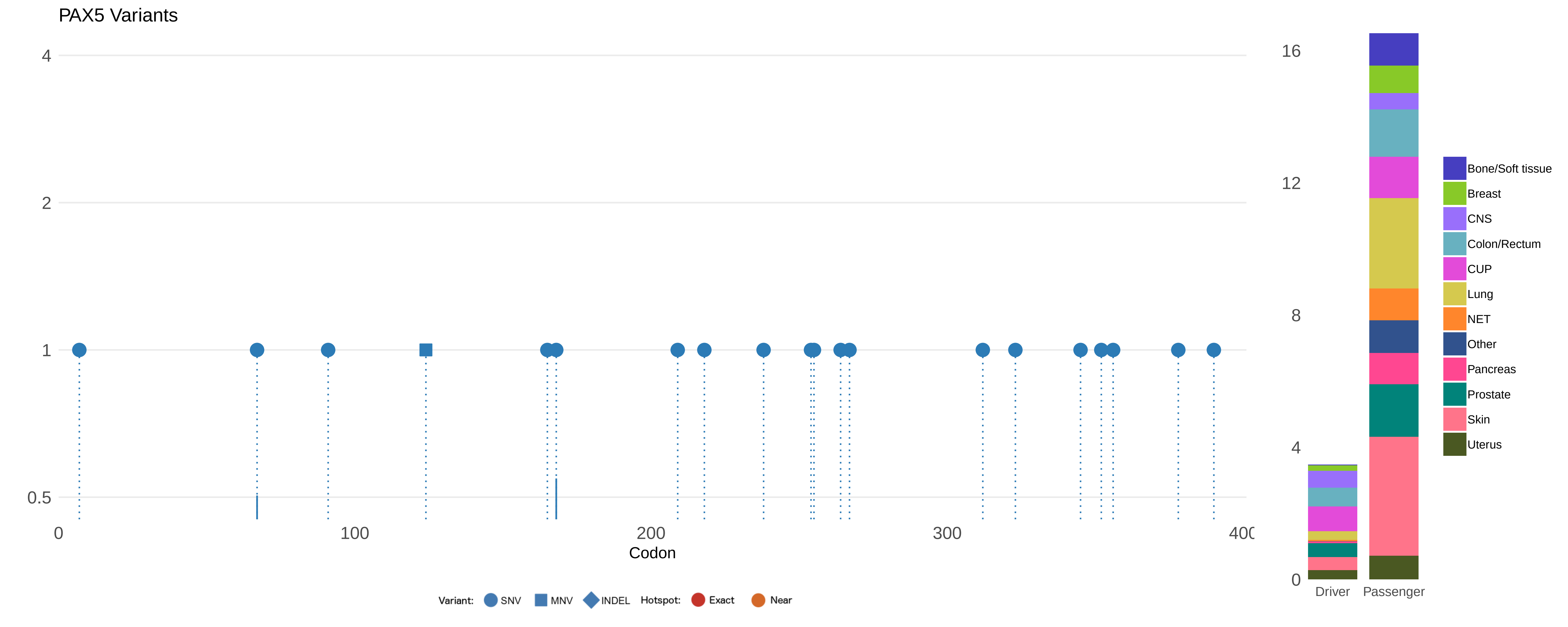
PAX5 Variants
4
2
1
0.5
0
100
200
300
400
Codon
Variant: SNV MNV INDEL Hotspot: Exact Near
16
12
8
4
0
Driver
Passenger
Bone/Soft tissue
Breast
CNS
Colon/Rectum
CUP
Lung
NET
Other
Pancreas
Prostate
Skin
Uterus

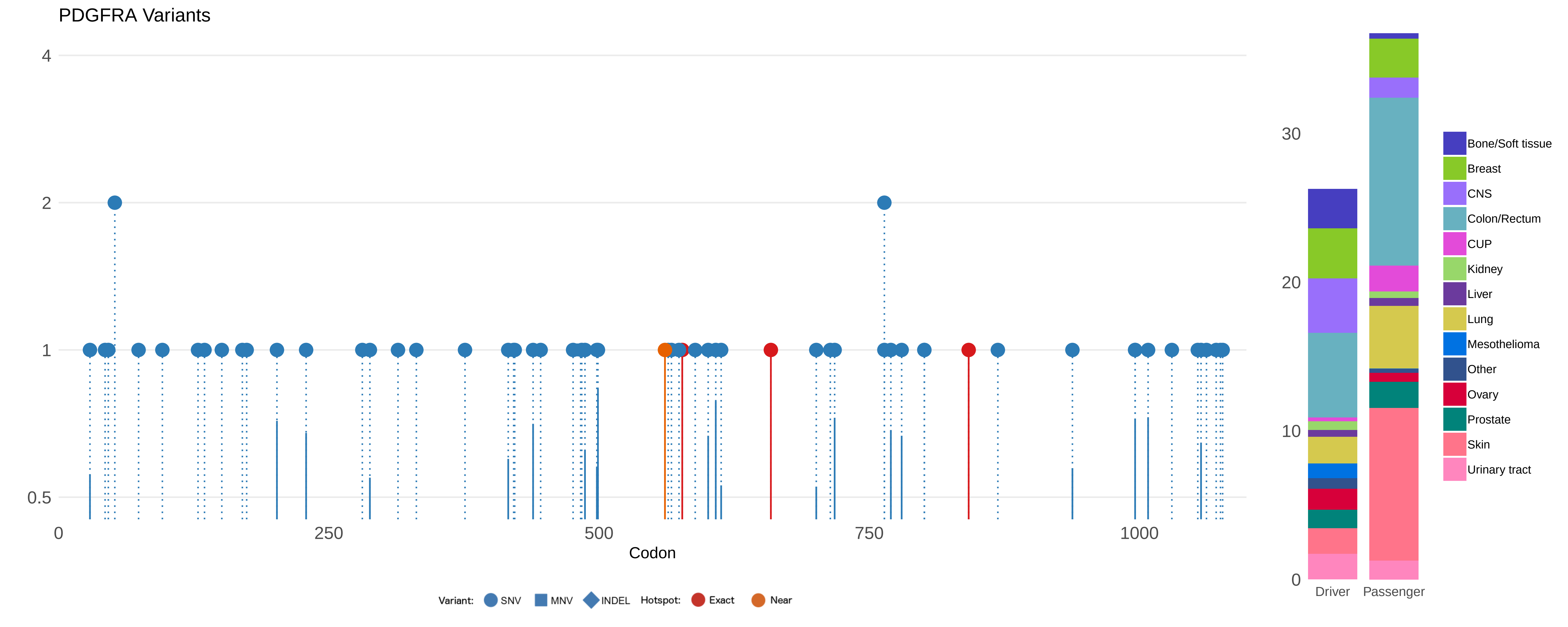

PDGFRA Variants
4
2
1
0.5
0
250
500
750
1000
Codon
Variant:
SNV
MNV
INDEL
Hotspot:
Exact
Near
30
20
10
0
Driver
Passenger
Bone/Soft tissue
Breast
CNS
Colon/Rectum
CUP
Kidney
Liver
Lung
Mesothelioma
Other
Ovary
Prostate
Skin
Urinary tract

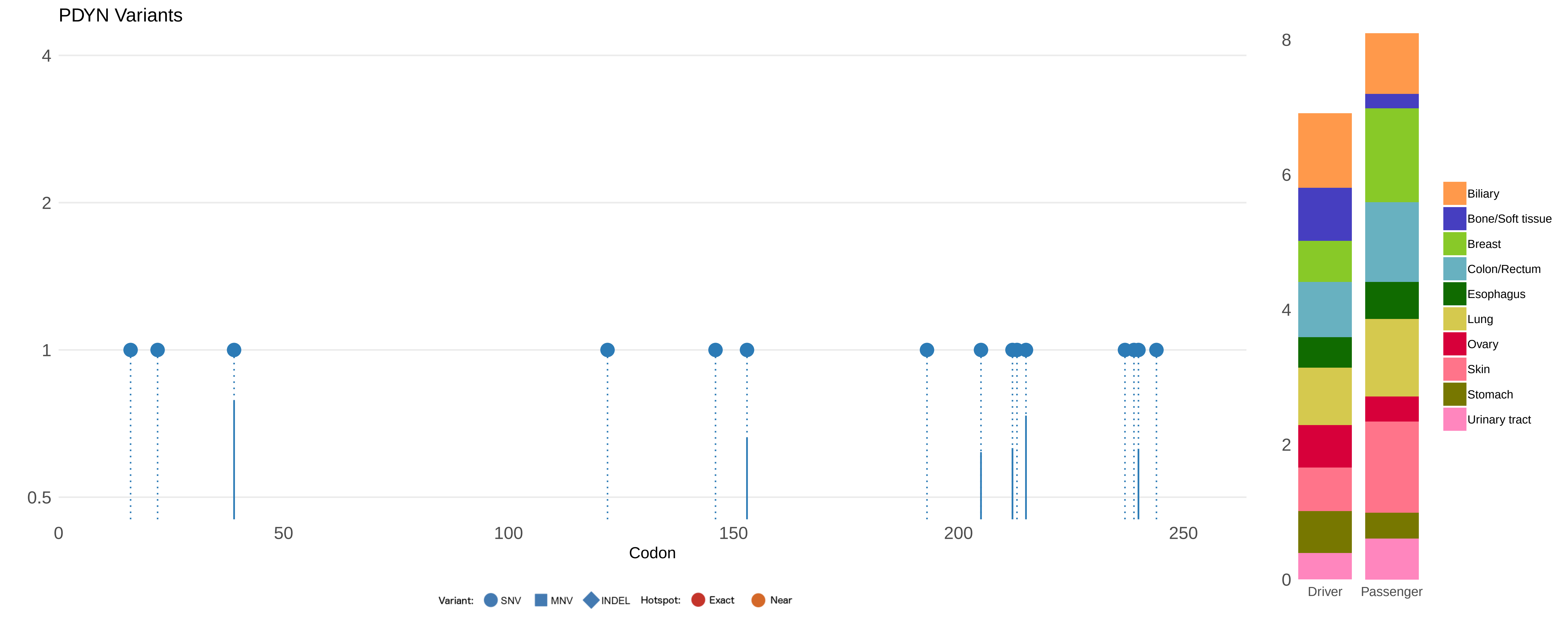

PDYN Variants
4
2
1
0.5
0
50
100
150
200
250
Codon
Variant: SNV MNV INDEL Hotspot: Exact Near
8
6
4
2
0
Driver
Passenger
Biliary
Bone/Soft tissue
Breast
Colon/Rectum
Esophagus
Lung
Ovary
Skin
Stomach
Urinary tract

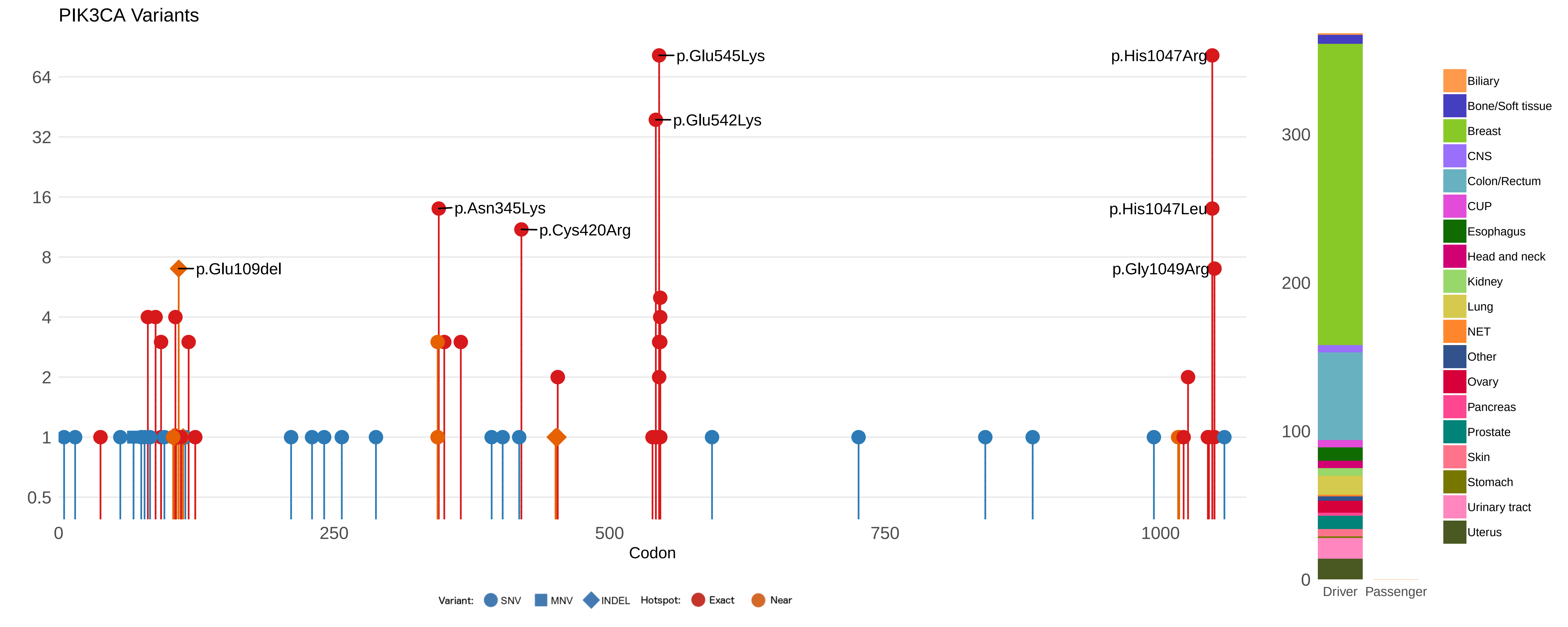
PIK3CA Variants
p.Glu545Lys
p.Glu542Lys
p.His1047Arg
p.His1047Leu
p.Gly1049Arg
p.Asn345Lys
p.Cys420Arg
p.Glu109del
64
32
16
8
4
2
1
0.5
0
250
500
750
1000
Codon
Variant: SNV MNV INDEL Hotspot: Exact Near
300
200
100
0
Driver
Passenger
Biliary
Bone/Soft tissue
Breast
CNS
Colon/Rectum
CUP
Esophagus
Head and neck
Kidney
Lung
NET
Other
Ovary
Pancreas
Prostate
Skin
Stomach
Urinary tract
Uterus

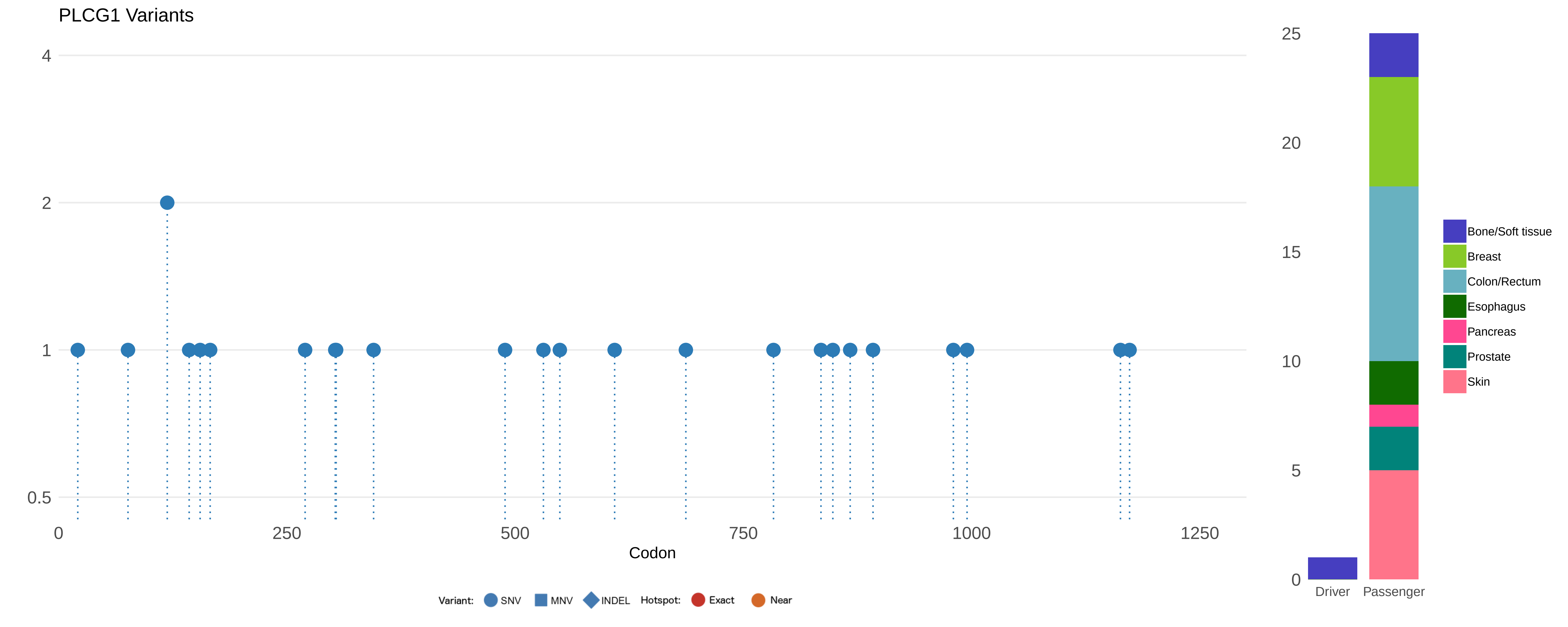
PLCG1 Variants
4
2
1
0.5
0
250
500
750
1000
1250
Codon
Variant: SNV MNV INDEL Hotspot: Exact Near
25
20
15
10
5
0
Driver
Passenger
Bone/Soft tissue
Breast
Colon/Rectum
Esophagus
Pancreas
Prostate
Skin

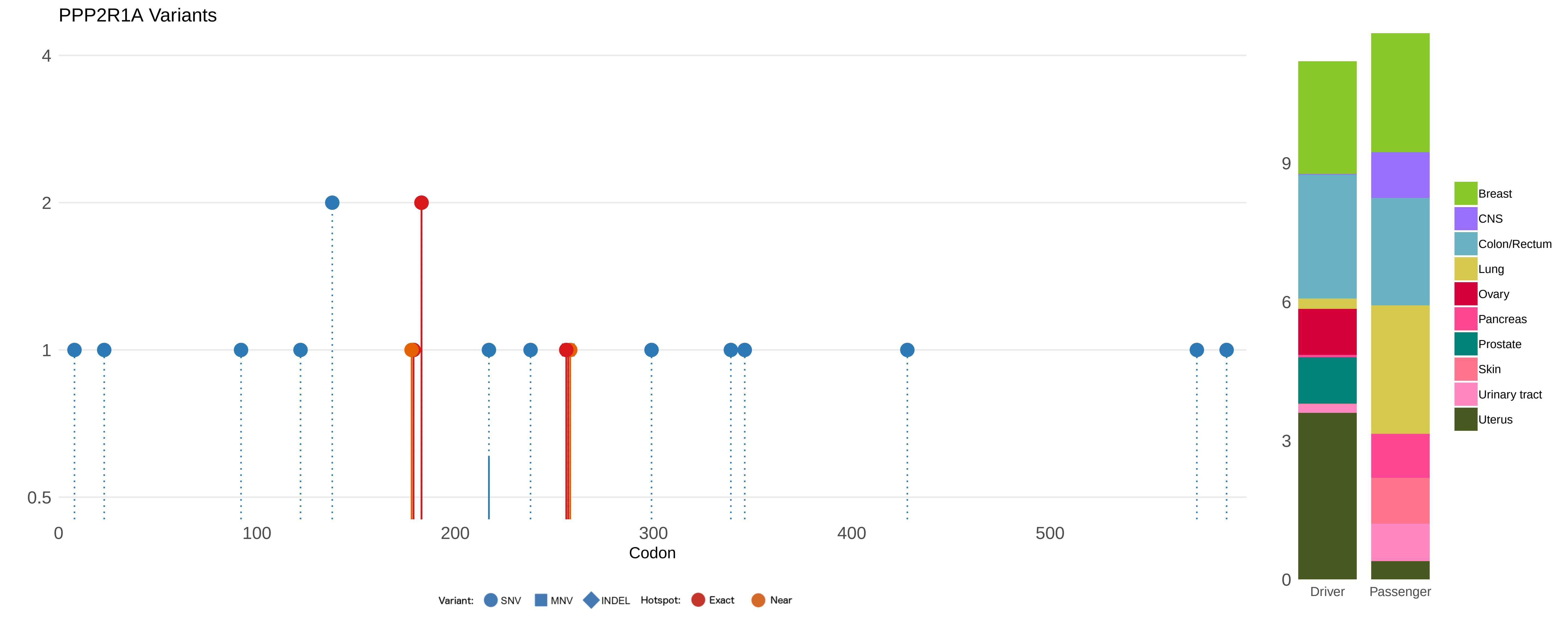
PPP2R1A Variants
4
2
1
0.5
0
100
200
300
400
500
Codon
Variant: SNV MNV INDEL Hotspot: Exact Near
9
6
3
0
Driver
Passenger
Breast
CNS
Colon/Rectum
Lung
Ovary
Pancreas
Prostate
Skin
Urinary tract
Uterus

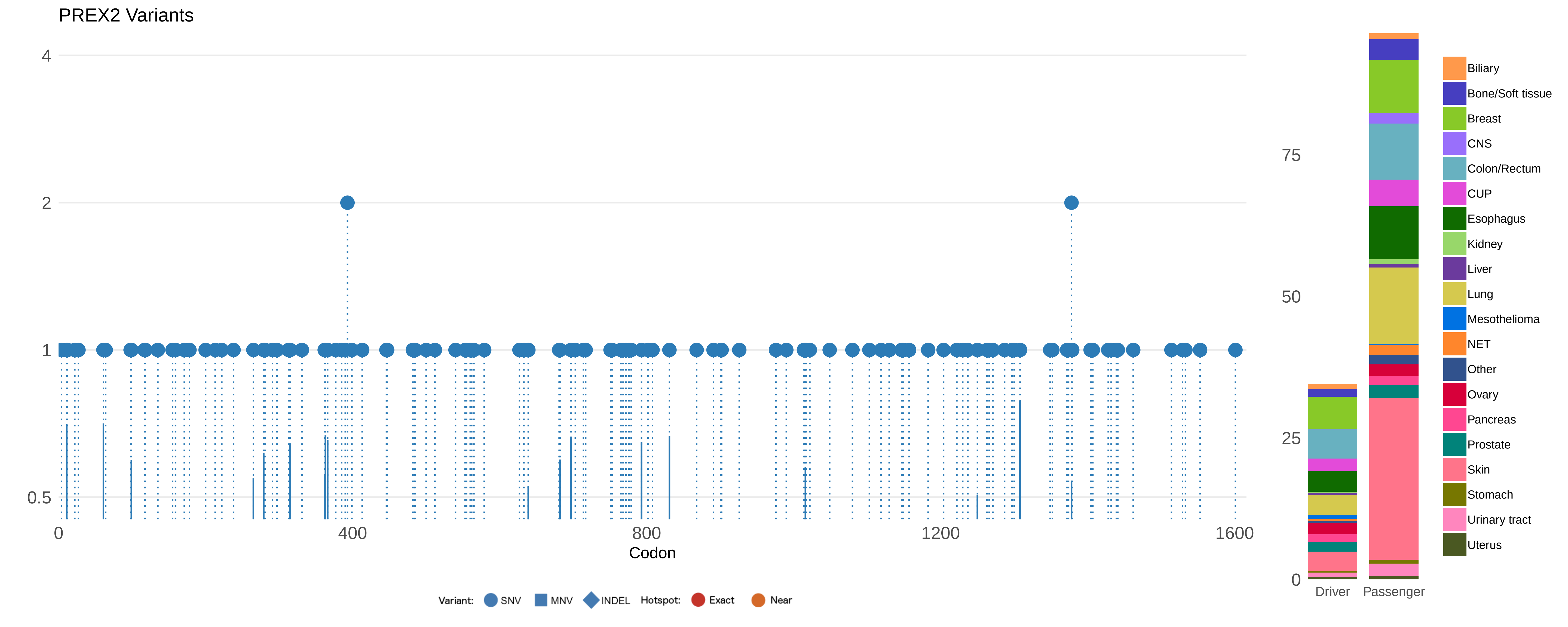

PREX2 Variants
4
2
1
0.5
0
400
800
1200
1600
Codon
Variant: SNV MNV INDEL Hotspot: Exact Near
75
50
25
0
Driver
Passenger
Biliary
Bone/Soft tissue
Breast
CNS
Colon/Rectum
CUP
Esophagus
Kidney
Liver
Lung
Mesothelioma
NET
Other
Ovary
Pancreas
Prostate
Skin
Stomach
Urinary tract
Uterus

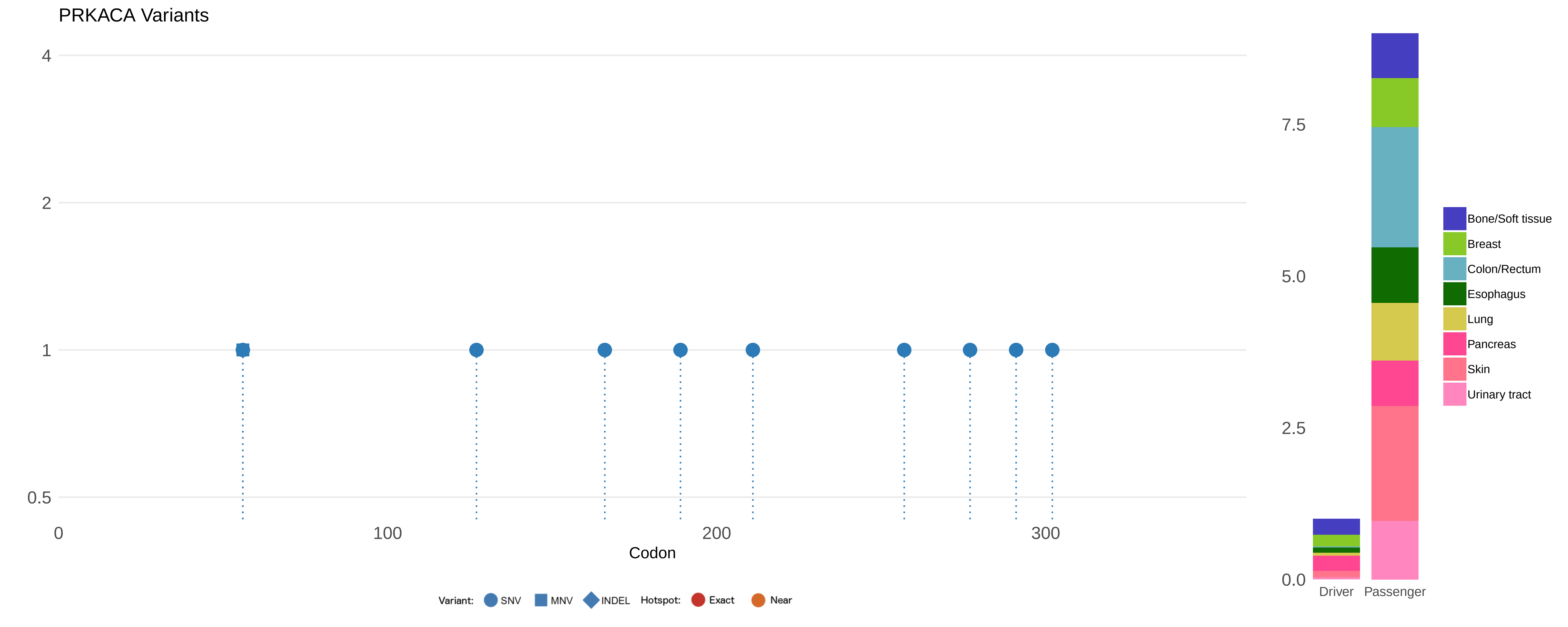

PRKACA Variants
4
2
1
0.5
0
100
200
300
Codon
Variant: SNV MNV INDEL Hotspot: Exact Near
7.5
5.0
2.5
0.0
Driver
Passenger
Bone/Soft tissue
Breast
Colon/Rectum
Esophagus
Lung
Pancreas
Skin
Urinary tract

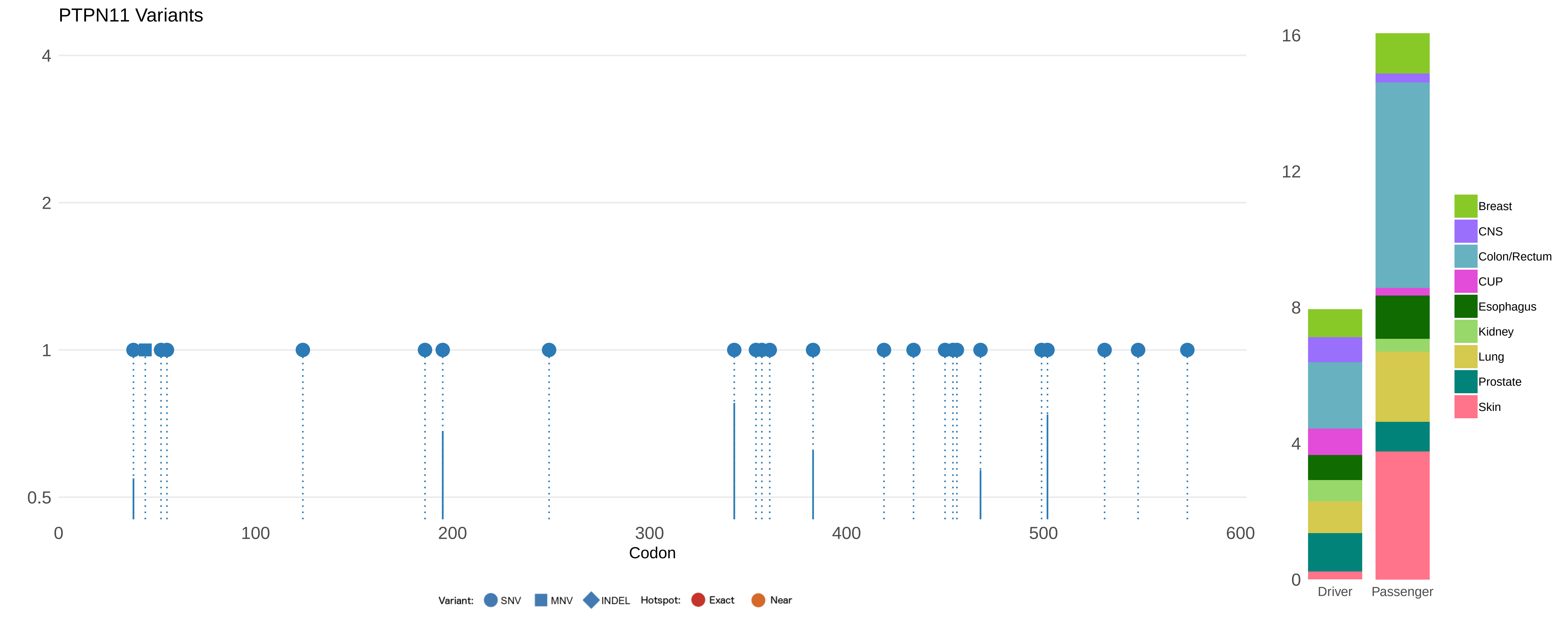

PTPN11 Variants
4
2
1
0.5
0
100
200
300
400
500
600
Codon
Variant: SNV MNV INDEL Hotspot: Exact Near
16
12
8
4
0
Driver
Passenger
Breast
CNS
Colon/Rectum
CUP
Esophagus
Kidney
Lung
Prostate
Skin

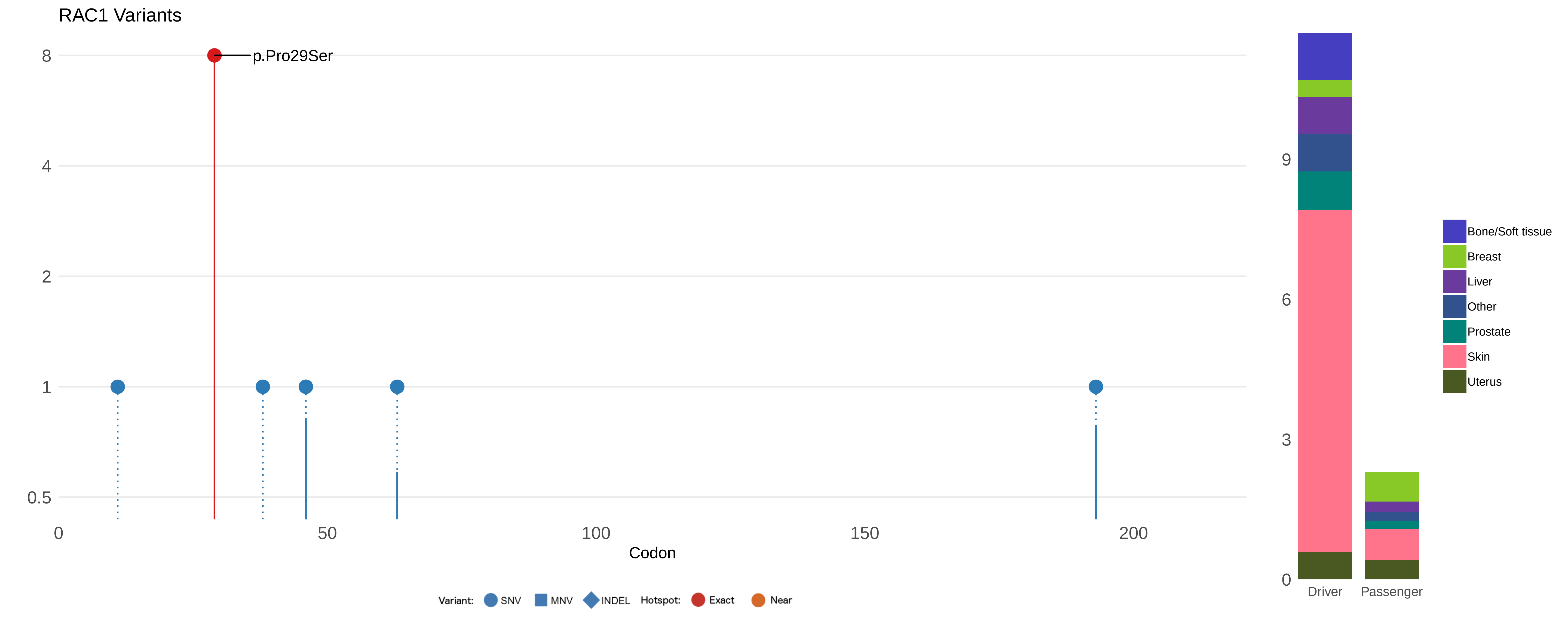

RAC1 Variants
p.Pro29Ser
8
4
2
1
0.5
0
50
100
150
200
Codon
Variant:
SNV
MNV
INDEL
Hotspot:
Exact
Near
9
6
3
0
Driver
Passenger
Bone/Soft tissue
Breast
Liver
Other
Prostate
Skin
Uterus

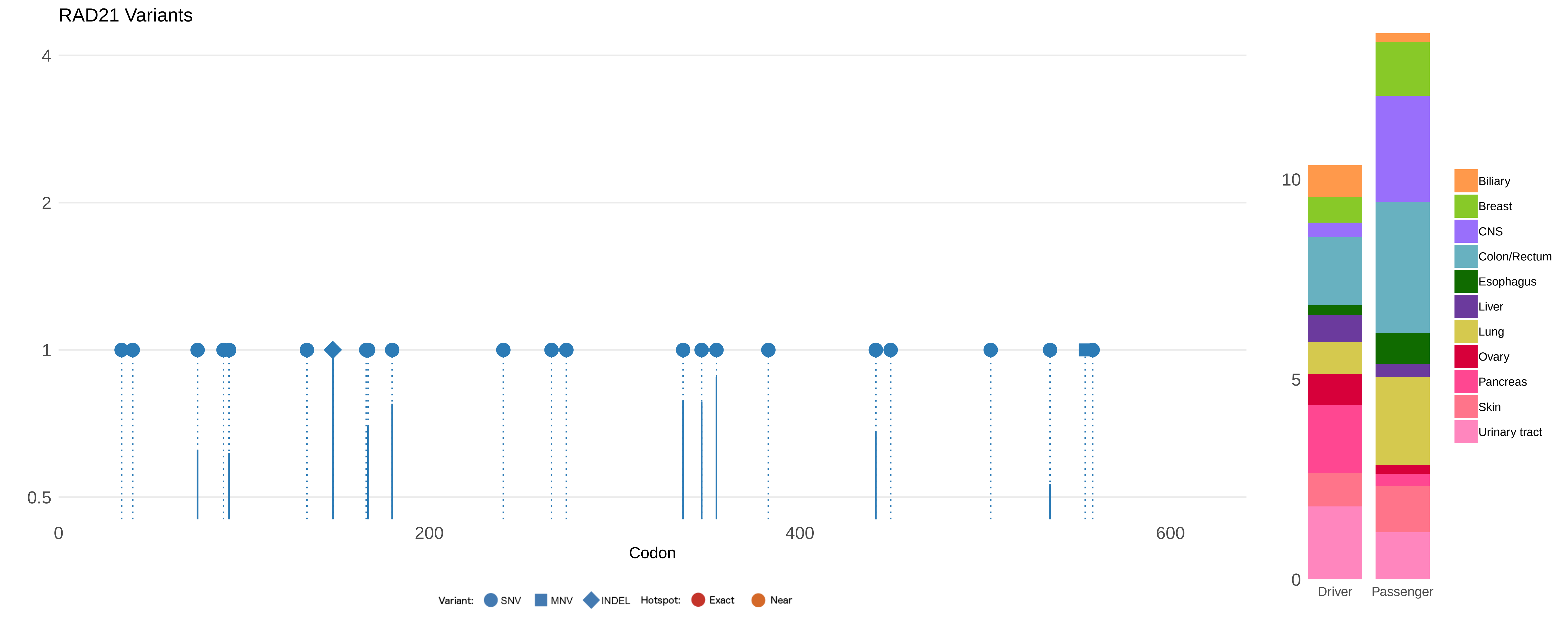
RAD21 Variants
4
2
1
0.5
0
200
400
600
Codon
Variant: SNV MNV INDEL Hotspot: Exact Near
10
5
0
Driver
Passenger
Biliary
Breast
CNS
Colon/Rectum
Esophagus
Liver
Lung
Ovary
Pancreas
Skin
Urinary tract

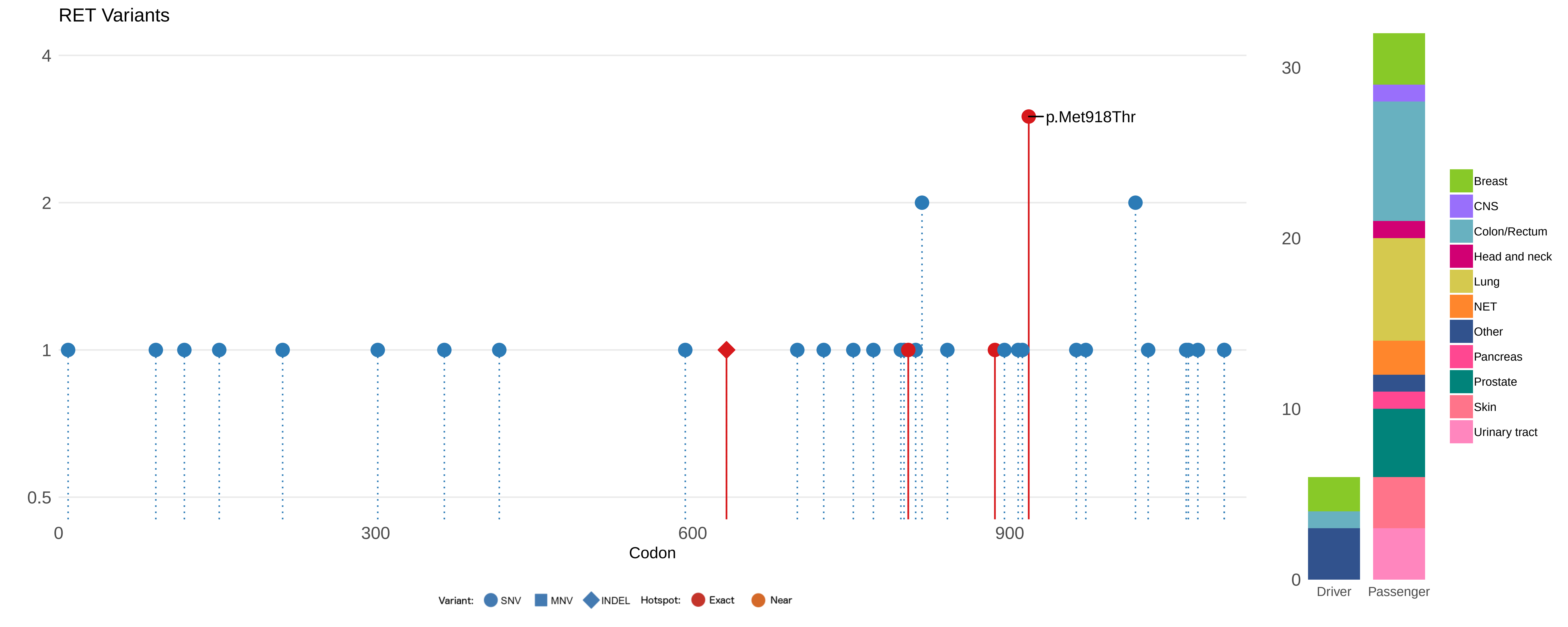

RET Variants
4
2
1
0.5
p.Met918Thr
0
300
600
900
Codon
Variant:
SNV
MNV
INDEL
Hotspot:
Exact
Near
30
20
10
0
Driver
Passenger
Breast
CNS
Colon/Rectum
Head and neck
Lung
NET
Other
Pancreas
Prostate
Skin
Urinary tract

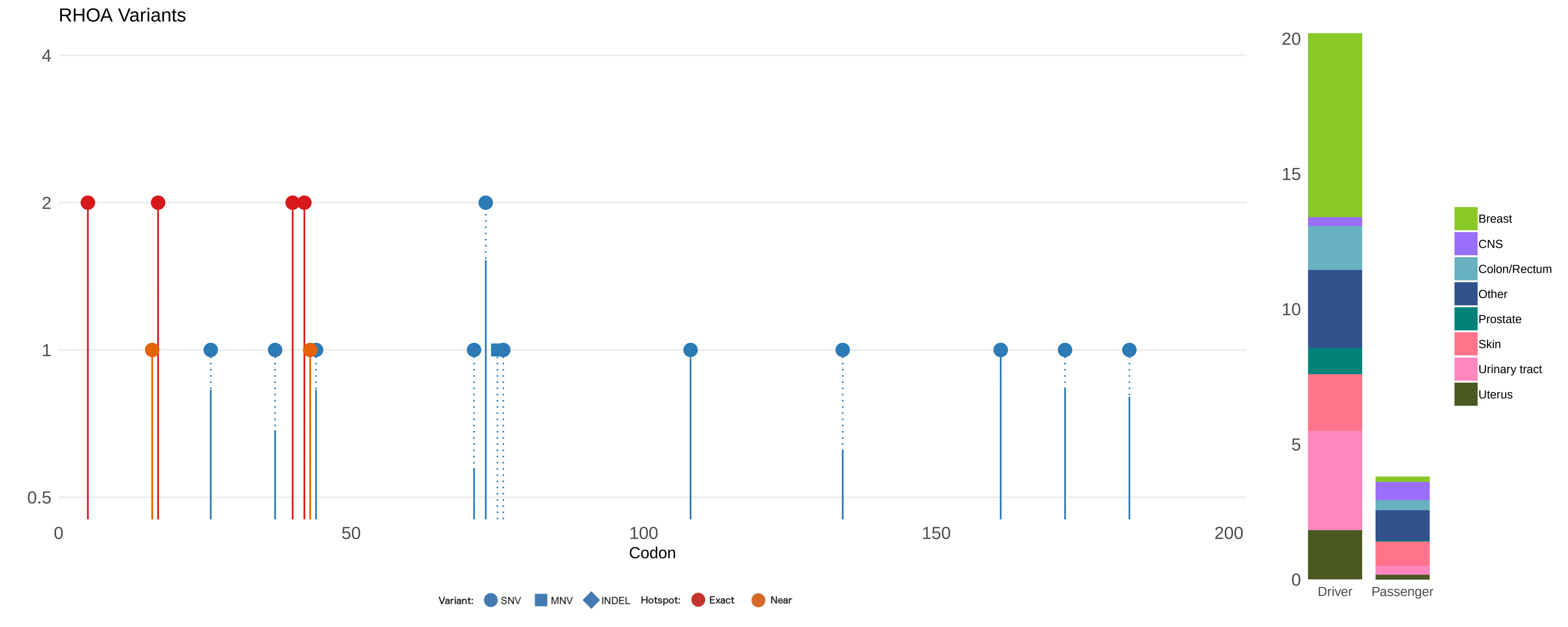

RHOA Variants
4
2
1
0.5
0
50
100
150
200
Codon
Variant:
SNV
MNV
INDEL
Hotspot:
Exact
Near
20
15
10
5
0
Driver
Passenger
Breast
CNS
Colon/Rectum
Other
Prostate
Skin
Urinary tract
Uterus

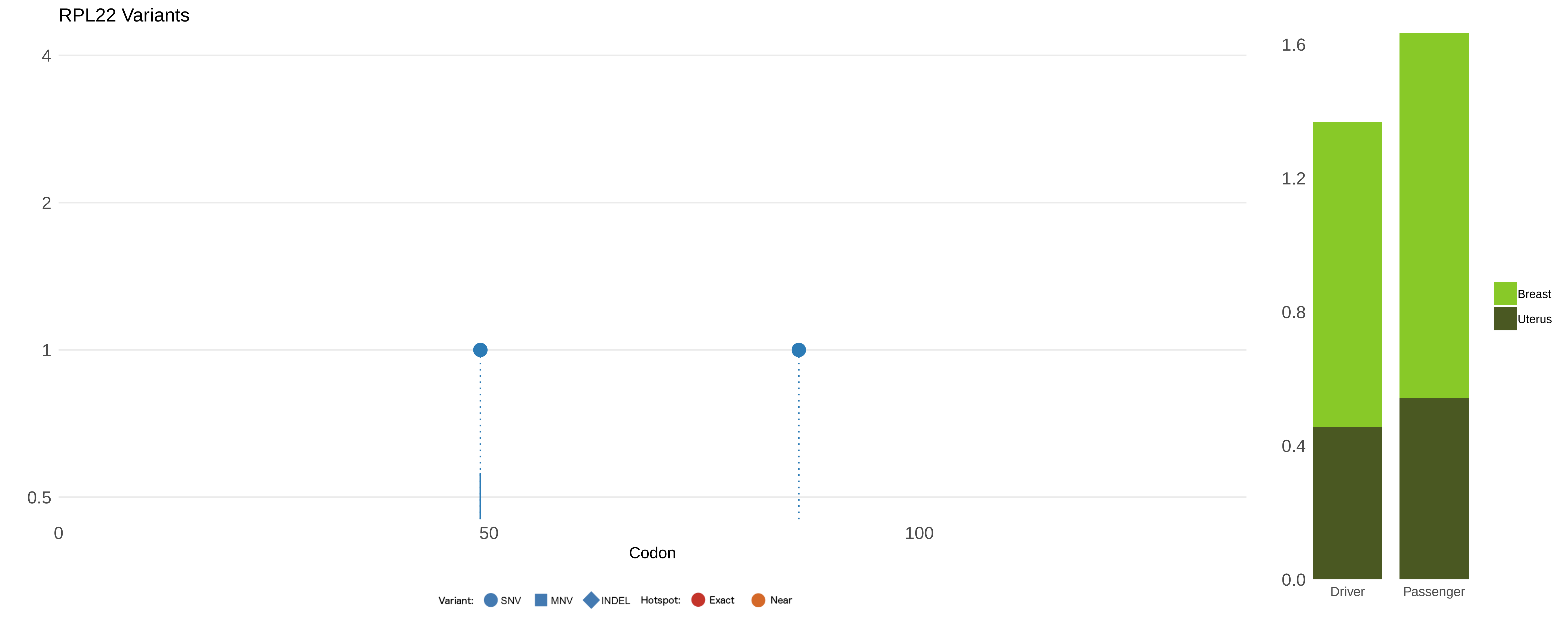

RPL22 Variants
4
2
1
0.5
0
50
100
Codon
Variant:
SNV
MNV
INDEL
Hotspot:
Exact
Near
1.6
1.2
0.8
0.4
0.0
Driver
Passenger
Breast
Uterus

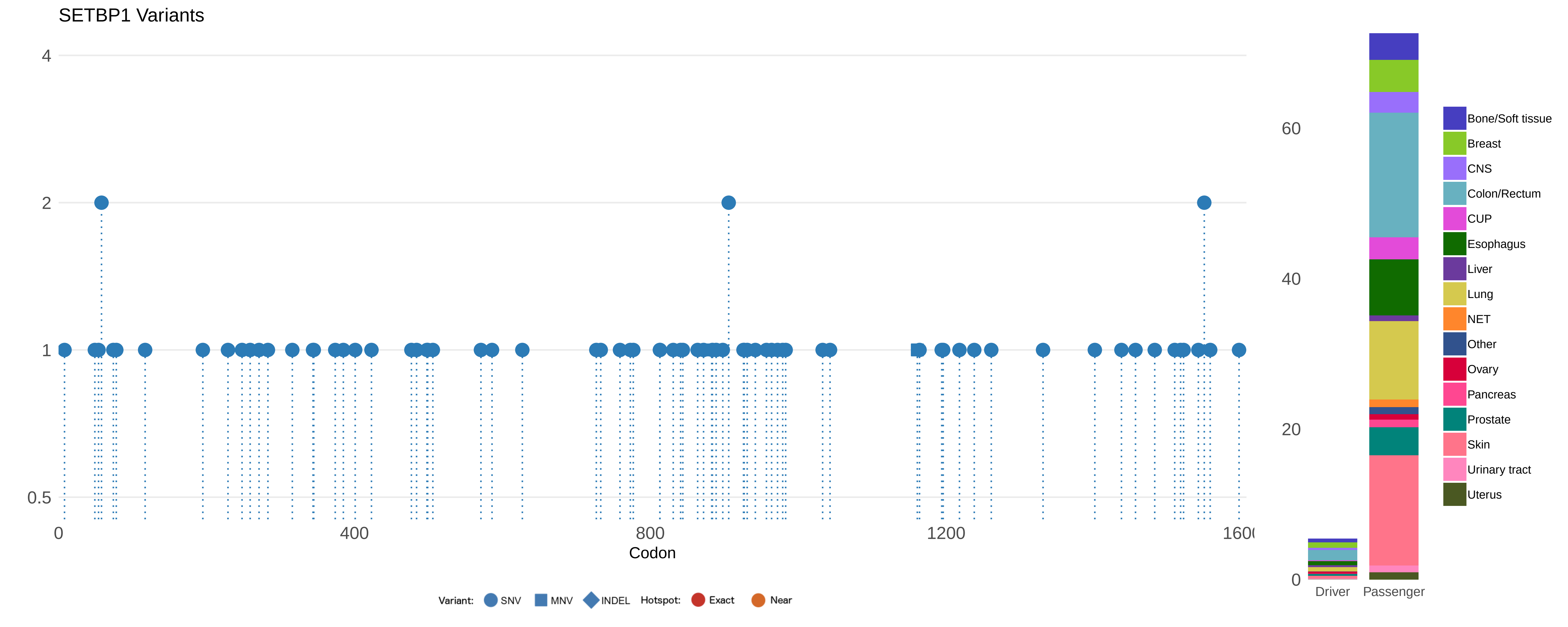

SETBP1 Variants
4
2
1
0.5
0
400
800
1200
1600
Codon
Variant: SNV MNV INDEL Hotspot: Exact Near
60
40
20
0
Driver
Passenger
Bone/Soft tissue
Breast
CNS
Colon/Rectum
CUP
Esophagus
Liver
Lung
NET
Other
Ovary
Pancreas
Prostate
Skin
Urinary tract
Uterus

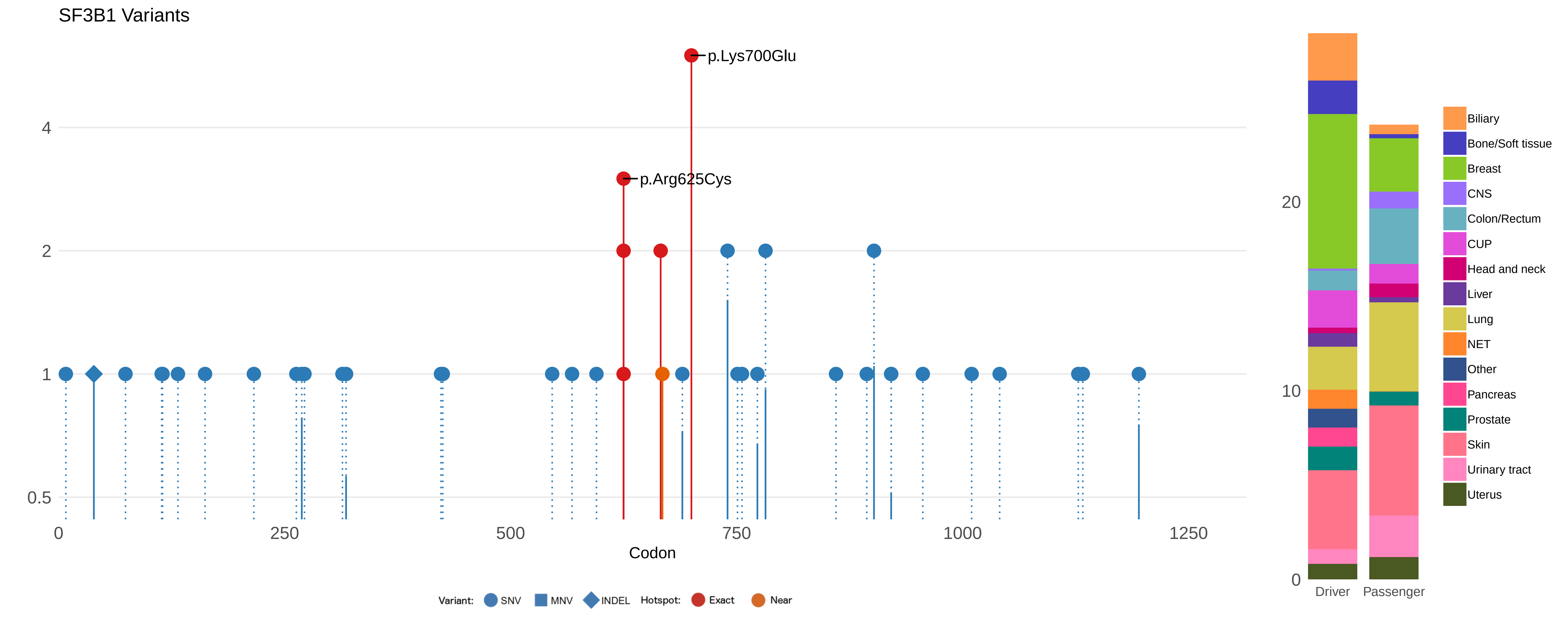
SF3B1 Variants
p.Lys700Glu
p.Arg625Cys
4
2
1
0.5
0
250
500
750
1000
1250
Codon
Variant: SNV MNV INDEL Hotspot: Exact Near
20
10
0
Driver
Passenger
Biliary
Bone/Soft tissue
Breast
CNS
Colon/Rectum
CUP
Head and neck
Liver
Lung
NET
Other
Pancreas
Prostate
Skin
Urinary tract
Uterus

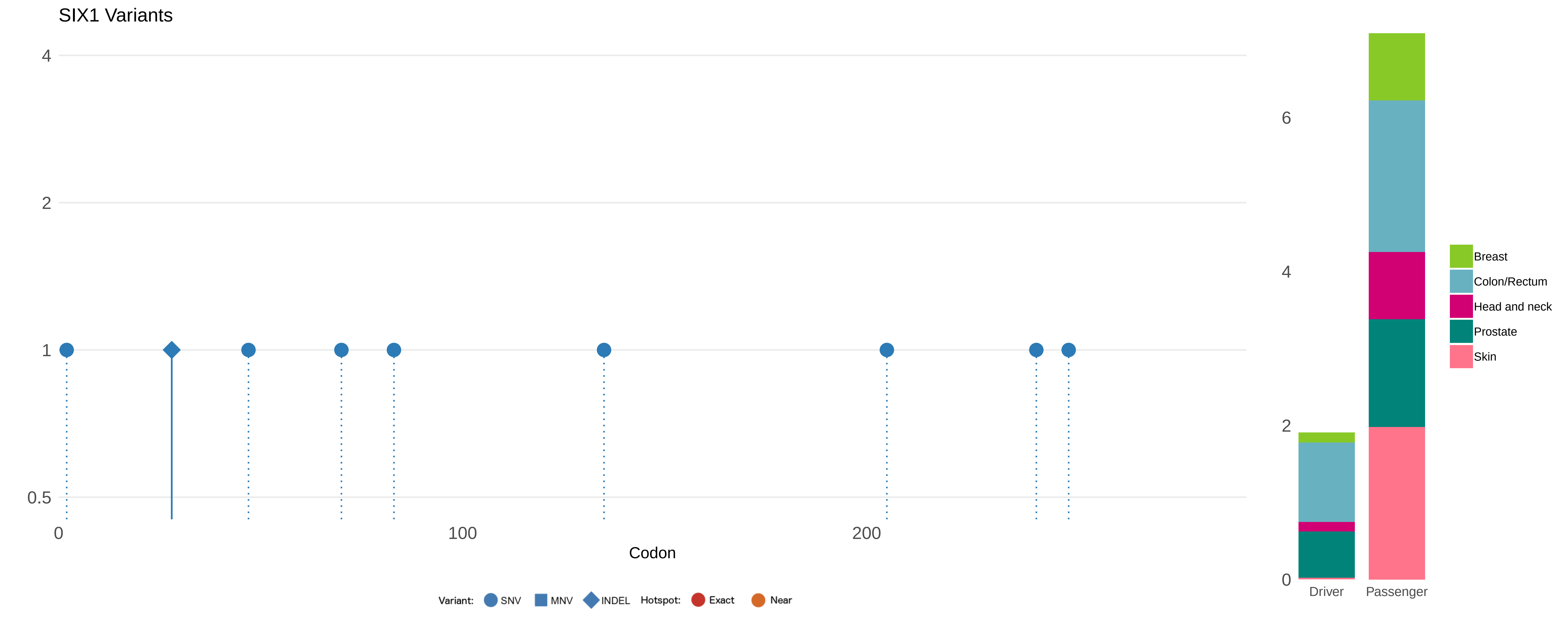

SIX1 Variants
4
2
1
0.5
0
100
200
Codon
Variant: SNV MNV INDEL Hotspot: Exact Near
6
4
2
0
Driver
Passenger
Breast
Colon/Rectum
Head and neck
Prostate
Skin

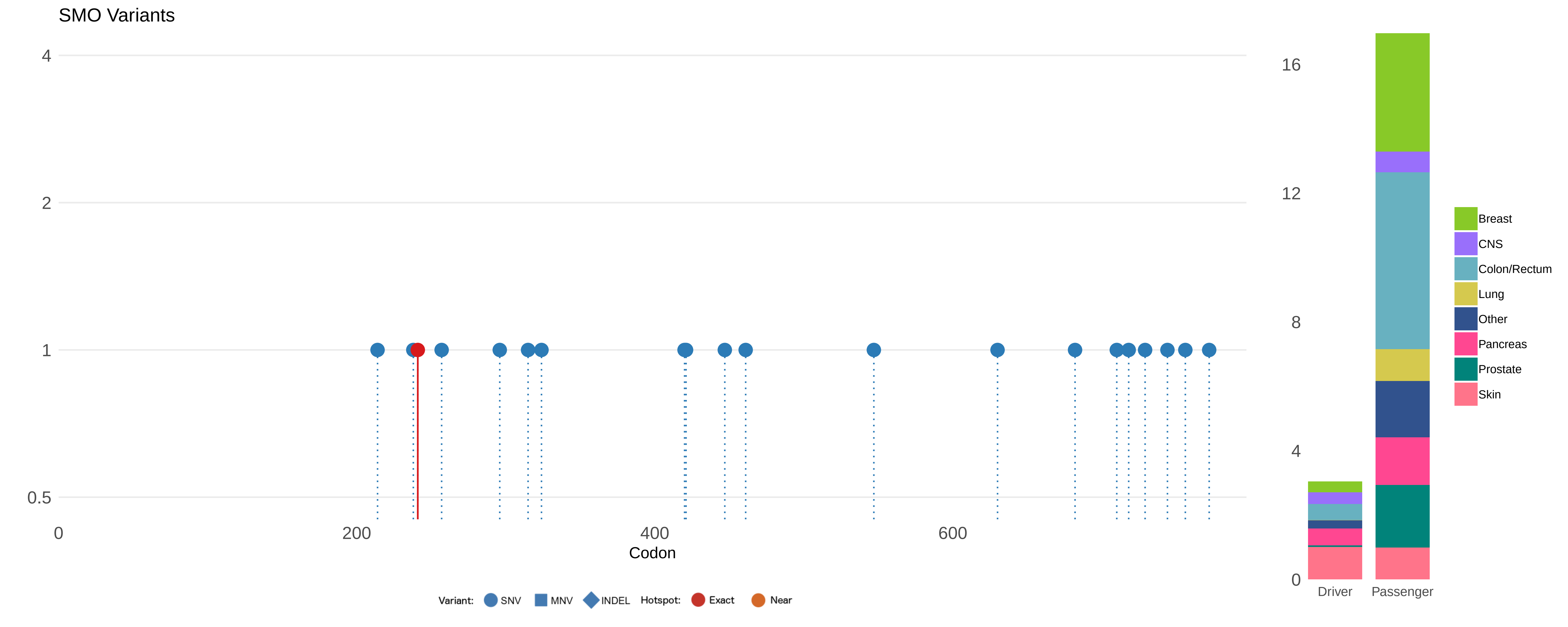
SMO Variants
4
2
1
0.5
0
200
400
600
Codon
Variant:
SNV
MNV
INDEL
Hotspot:
Exact
Near
16
12
8
4
0
Driver
Passenger
Breast
CNS
Colon/Rectum
Lung
Other
Pancreas
Prostate
Skin

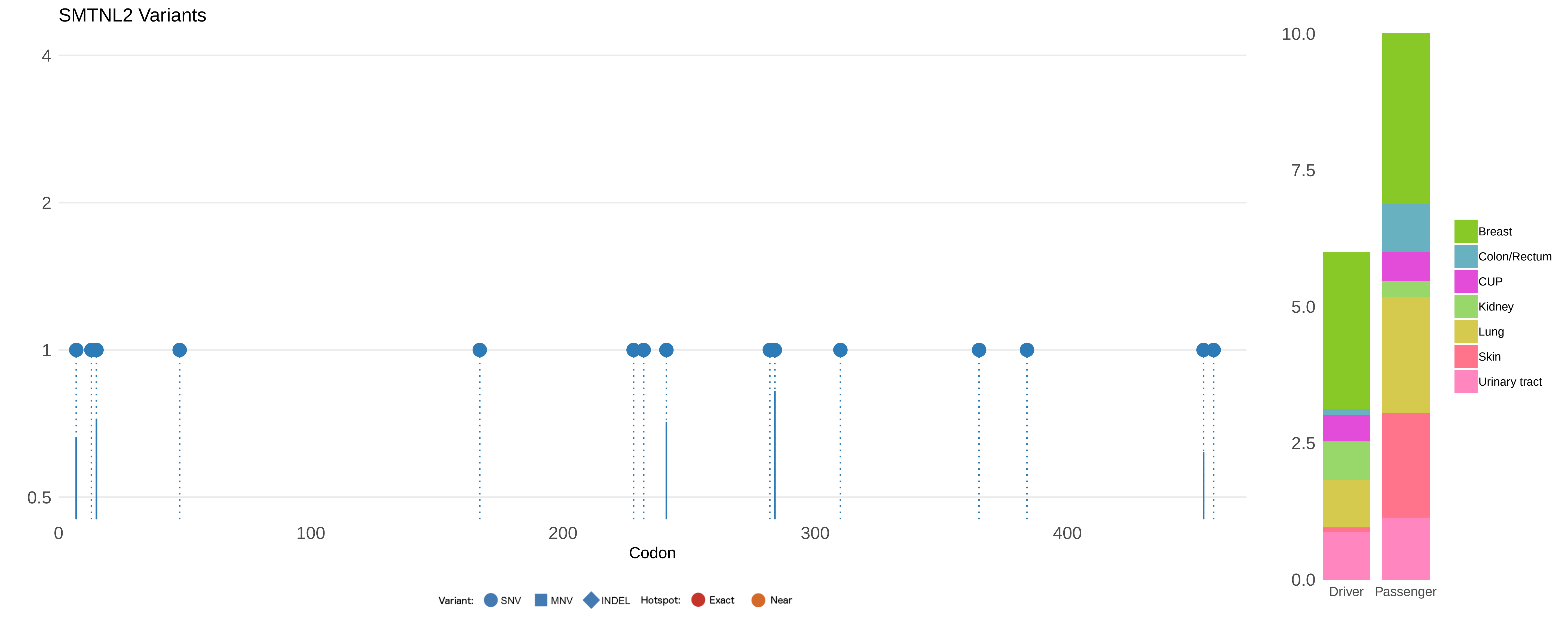

SMTNL2 Variants
4
2
1
0.5
0
100
200
300
400
Codon
Variant: SNV MNV INDEL Hotspot: Exact Near
10.0
7.5
5.0
2.5
0.0
Driver
Passenger
Breast
Colon/Rectum
CUP
Kidney
Lung
Skin
Urinary tract

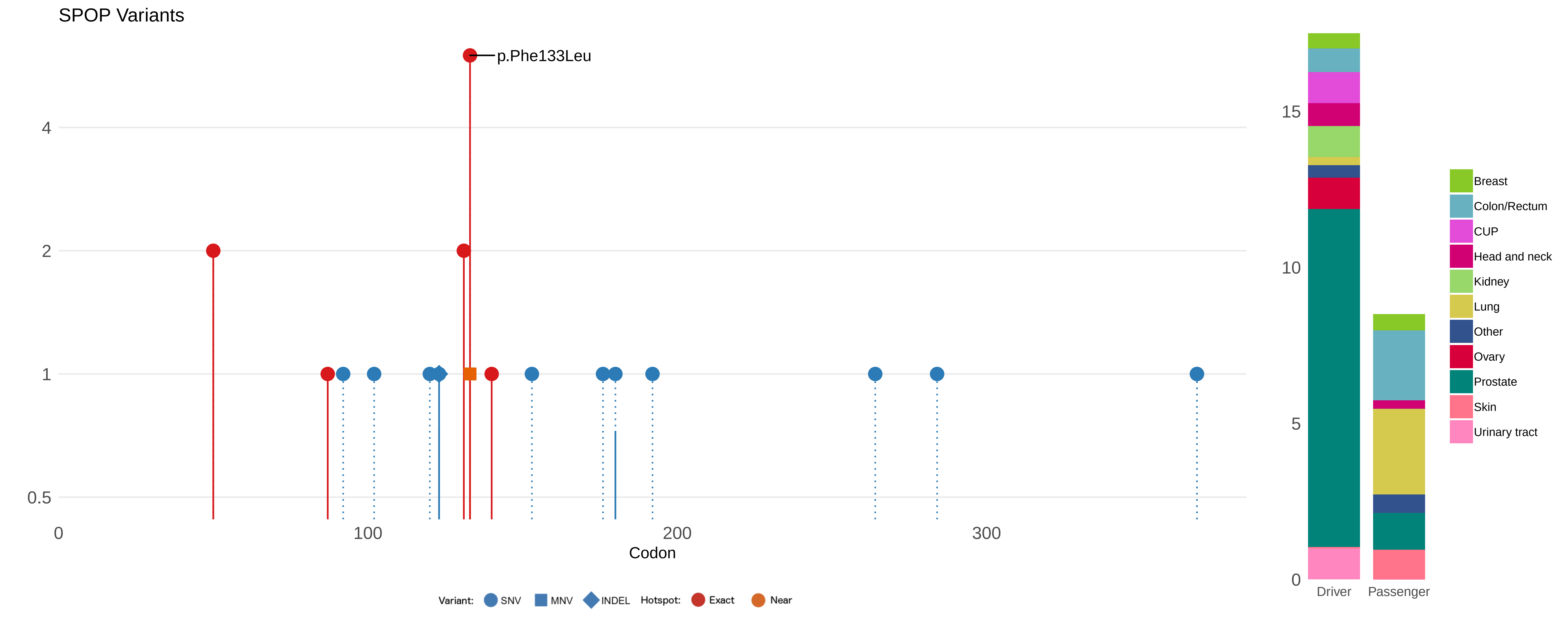
SPOP Variants
p.Phe133Leu
4
2
1
0.5
0
100
200
300
Codon
Variant: SNV MNV INDEL Hotspot: Exact Near
15
10
5
0
Driver
Passenger
Breast
Colon/Rectum
CUP
Head and neck
Kidney
Lung
Other
Ovary
Prostate
Skin
Urinary tract

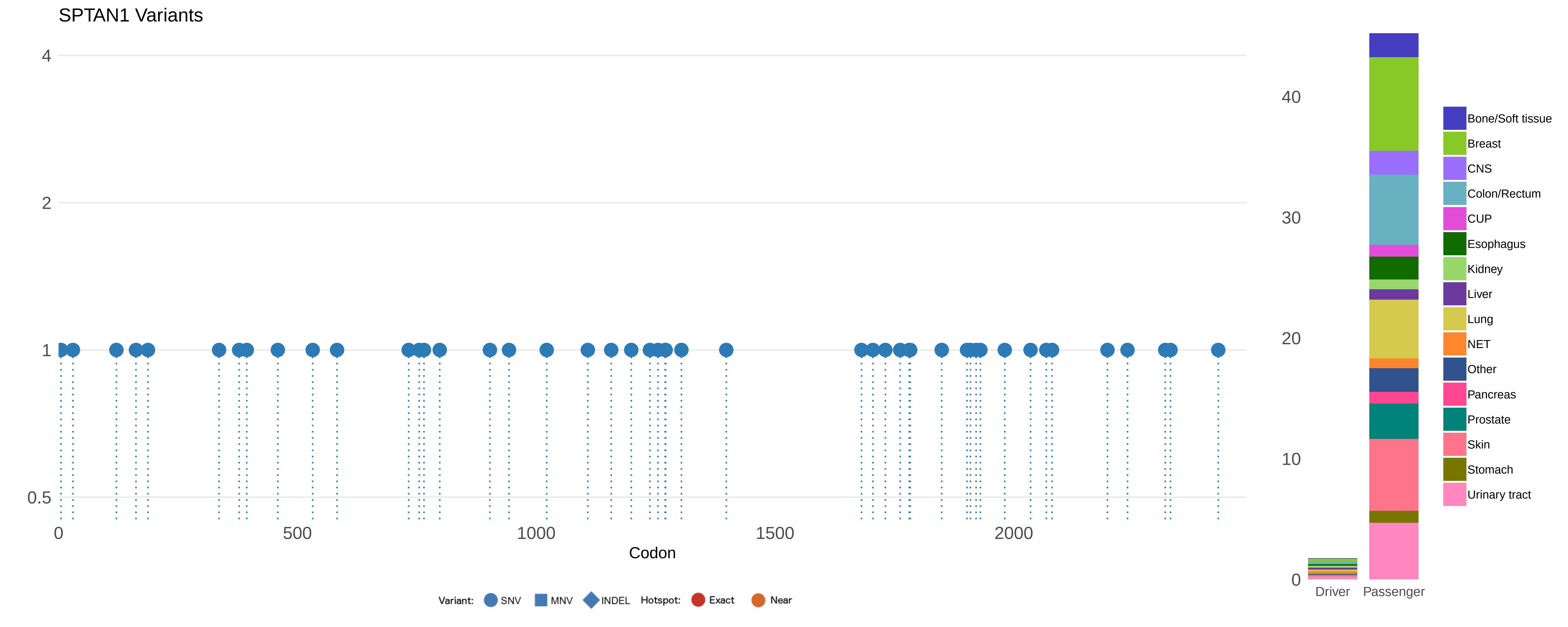
SPTAN1 Variants
4
2
1
0.5
0
500
1000
1500
2000
Codon
Variant: SNV MNV INDEL Hotspot: Exact Near
40
30
20
10
0
Driver
Passenger
Bone/Soft tissue
Breast
CNS
Colon/Rectum
CUP
Esophagus
Kidney
Liver
Lung
NET
Other
Pancreas
Prostate
Skin
Stomach
Urinary tract

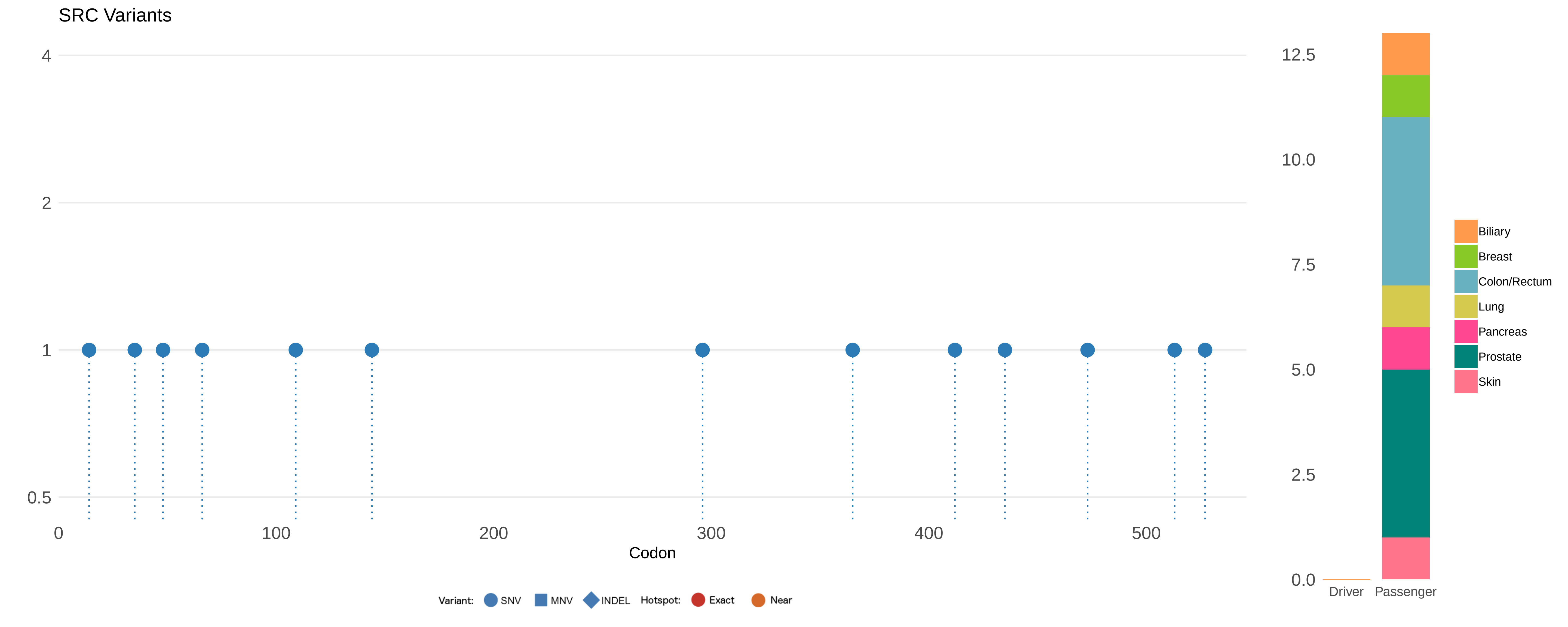

SRC Variants
4
2
1
0.5
0
100
200
300
400
500
Codon
Variant:
SNV
MNV
INDEL
Hotspot:
Exact
Near
12.5
10.0
7.5
5.0
2.5
0.0
Driver
Passenger
Biliary
Breast
Colon/Rectum
Lung
Pancreas
Prostate
Skin

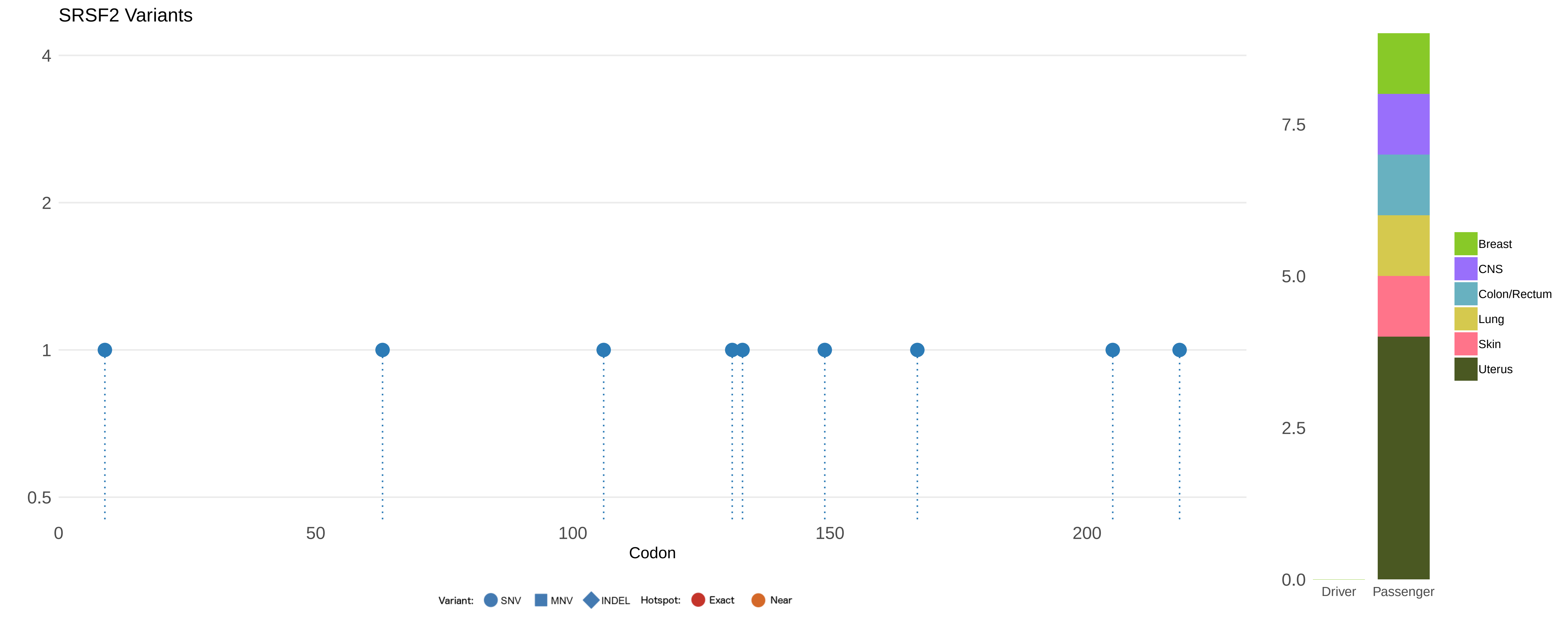
SRSF2 Variants
4
2
1
0.5
0
50
100
150
200
Codon
Variant:
SNV
MNV
INDEL
Hotspot:
Exact
Near
7.5
5.0
2.5
0.0
Driver
Passenger
Breast
CNS
Colon/Rectum
Lung
Skin
Uterus

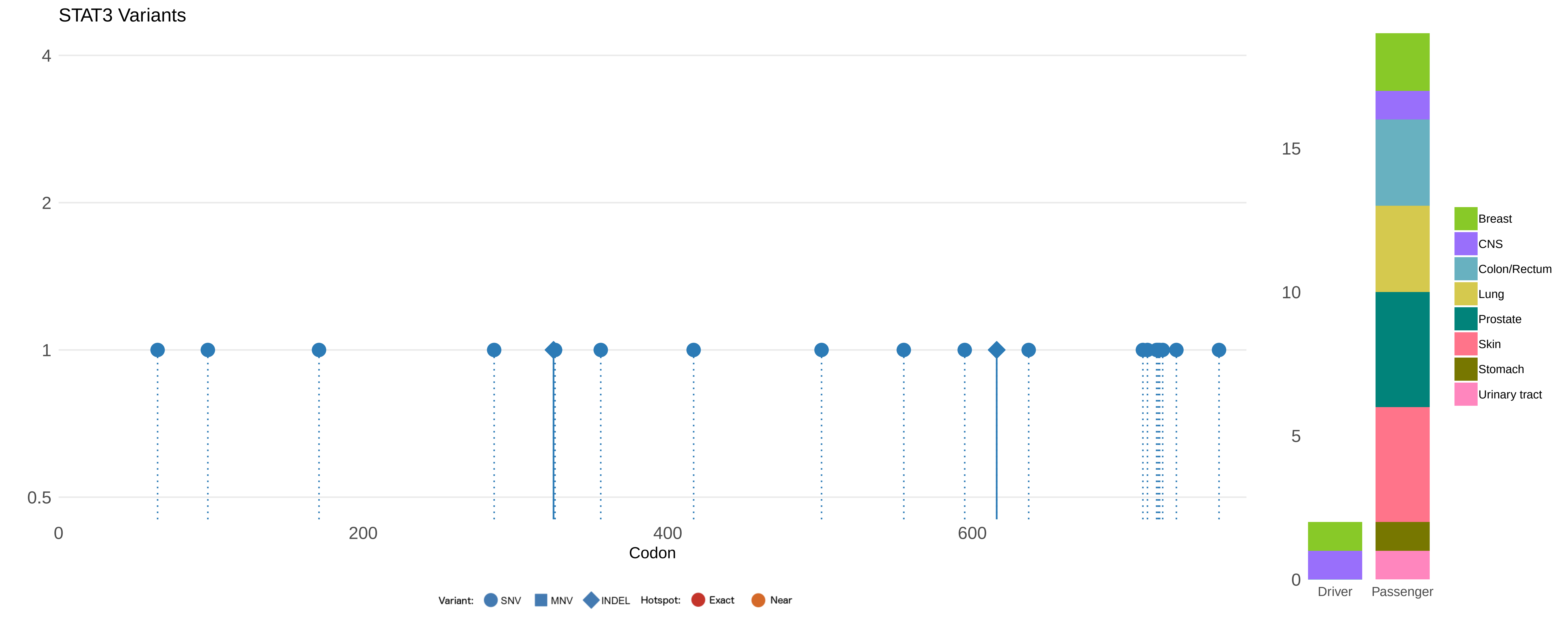
STAT3 Variants
4
2
1
0.5
0
200
400
600
Codon
Variant:
SNV
MNV
INDEL
Hotspot:
Exact
Near
15
10
5
0
Driver
Passenger
Breast
CNS
Colon/Rectum
Lung
Prostate
Skin
Stomach
Urinary tract

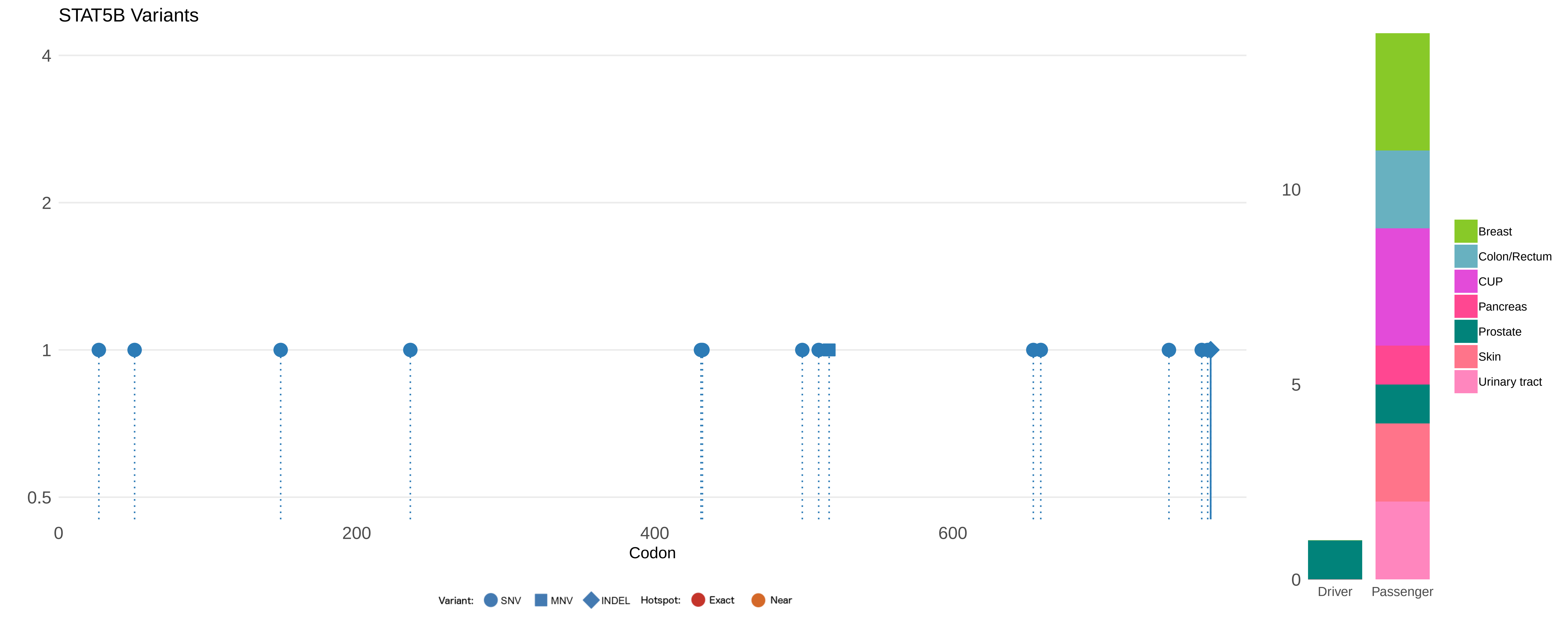
STAT5B Variants
4
2
1
0.5
0
200
400
600
Codon
Variant:
SNV
MNV
INDEL
Hotspot:
Exact
Near
10
5
0
Driver
Passenger
Breast
Colon/Rectum
CUP
Pancreas
Prostate
Skin
Urinary tract

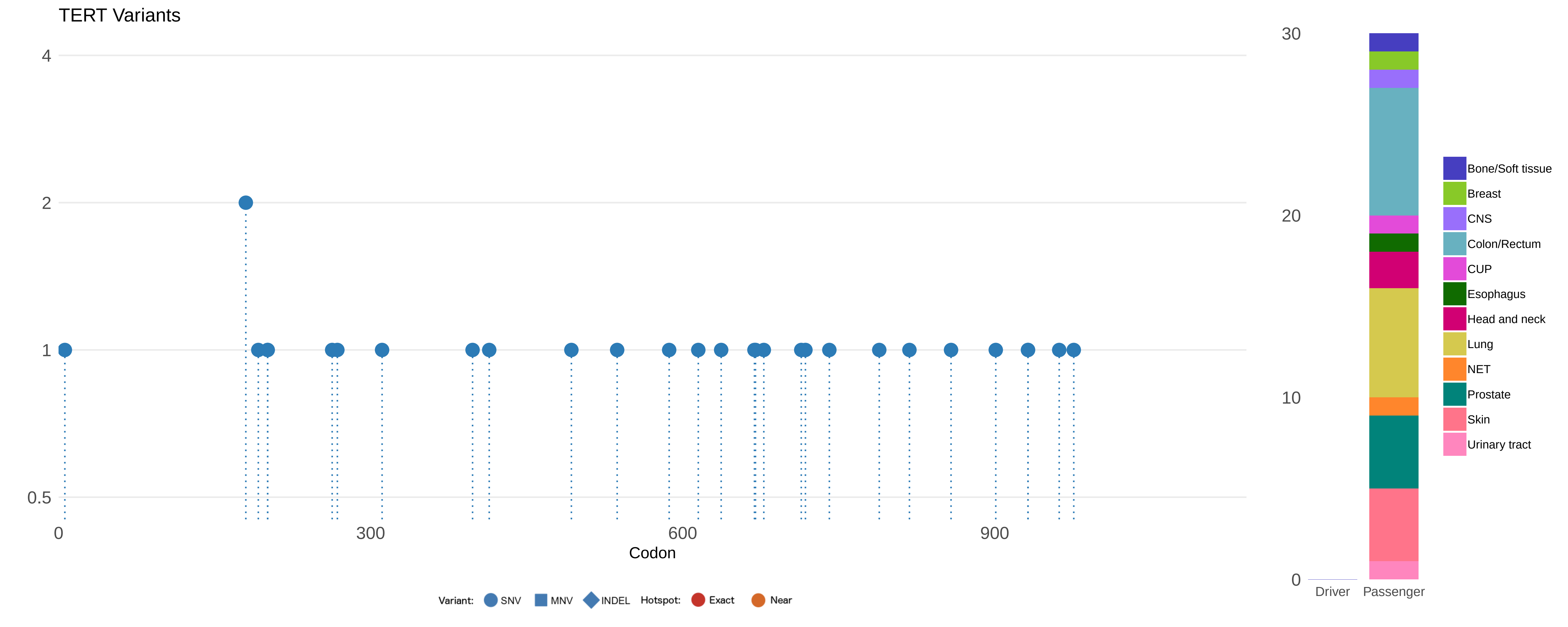

TERT Variants
4
2
1
0.5
0
300
600
900
Codon
Variant:
SNV
MNV
INDEL
Hotspot:
Exact
Near
30
20
10
0
Driver
Passenger
Bone/Soft tissue
Breast
CNS
Colon/Rectum
CUP
Esophagus
Head and neck
Lung
NET
Prostate
Skin
Urinary tract

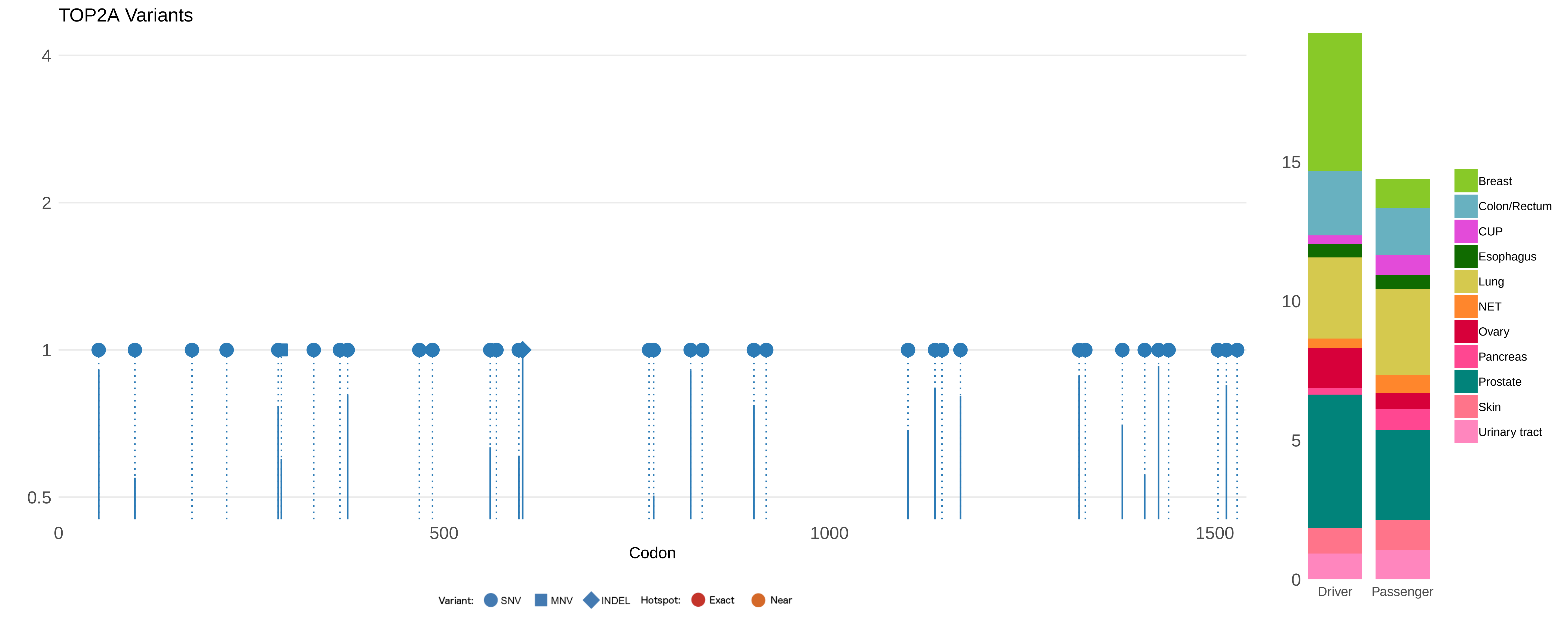

TOP2A Variants
4
2
1
0.5
0
500
1000
1500
Codon
Variant:
SNV
MNV
INDEL
Hotspot:
Exact
Near
15
10
5
0
Driver
Passenger
Breast
Colon/Rectum
CUP
Esophagus
Lung
NET
Ovary
Pancreas
Prostate
Skin
Urinary tract

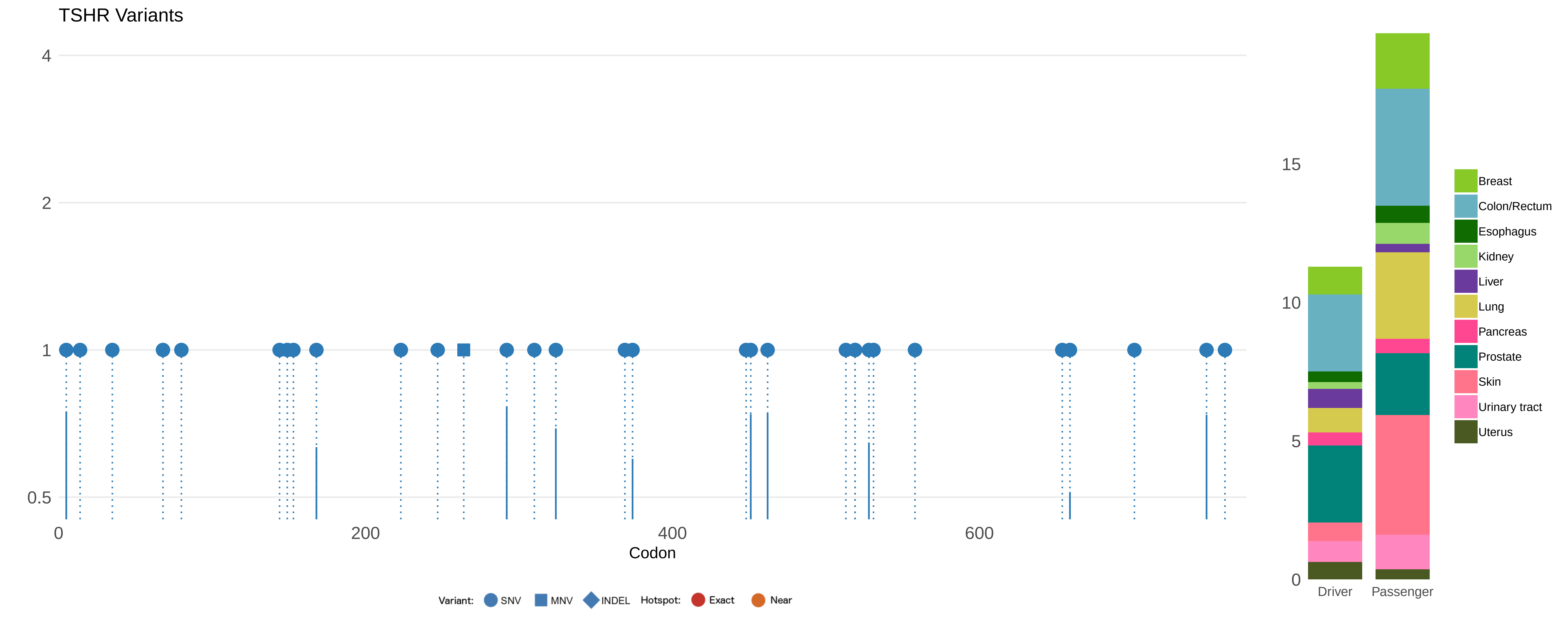
TSHR Variants
4
2
1
0.5
0
200
400
600
Codon
Variant: SNV MNV INDEL Hotspot: Exact Near
15
10
5
0
Driver
Passenger
Breast
Colon/Rectum
Esophagus
Kidney
Liver
Lung
Pancreas
Prostate
Skin
Urinary tract
Uterus

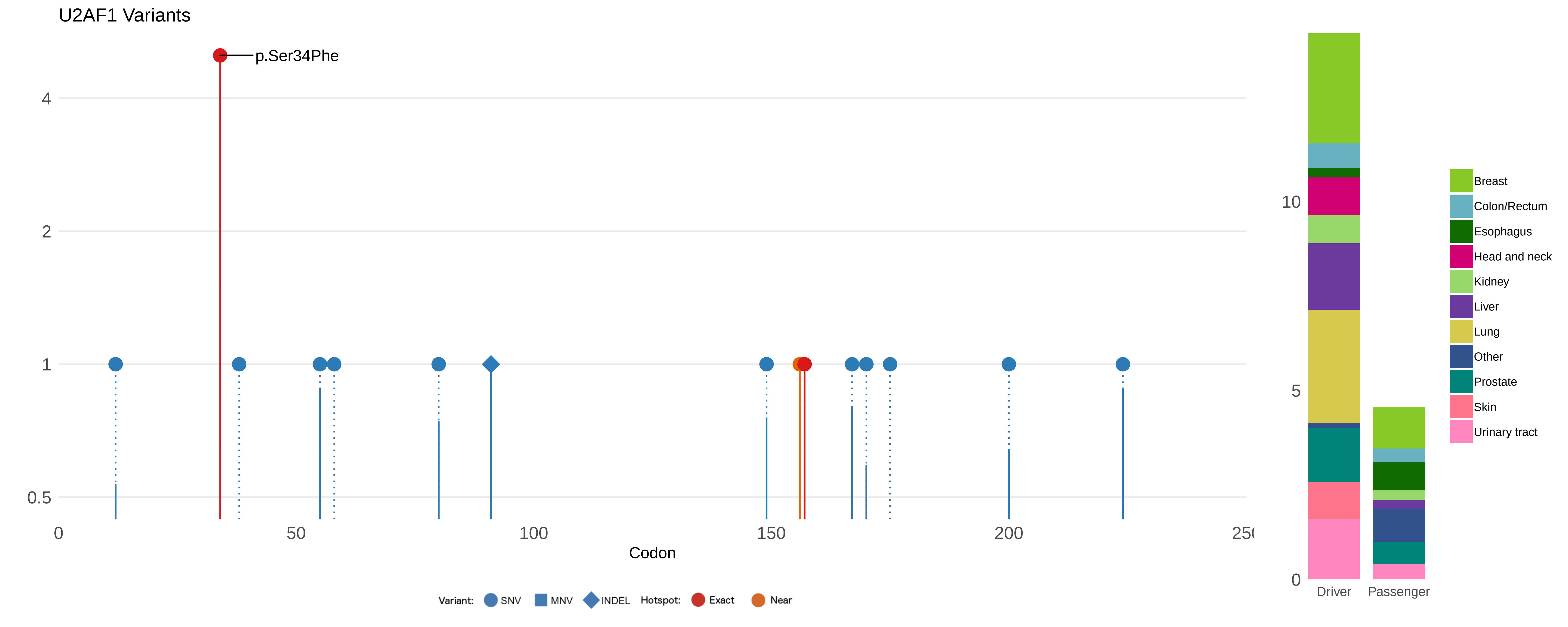

U2AF1 Variants
p.Ser34Phe
4
2
1
0.5
0
50
100
150
200
25(
Codon
Variant:
SNV
MNV
INDEL
Hotspot:
Exact
Near
10
5
0
Driver
Passenger
Breast
Colon/Rectum
Esophagus
Head and neck
Kidney
Liver
Lung
Other
Prostate
Skin
Urinary tract

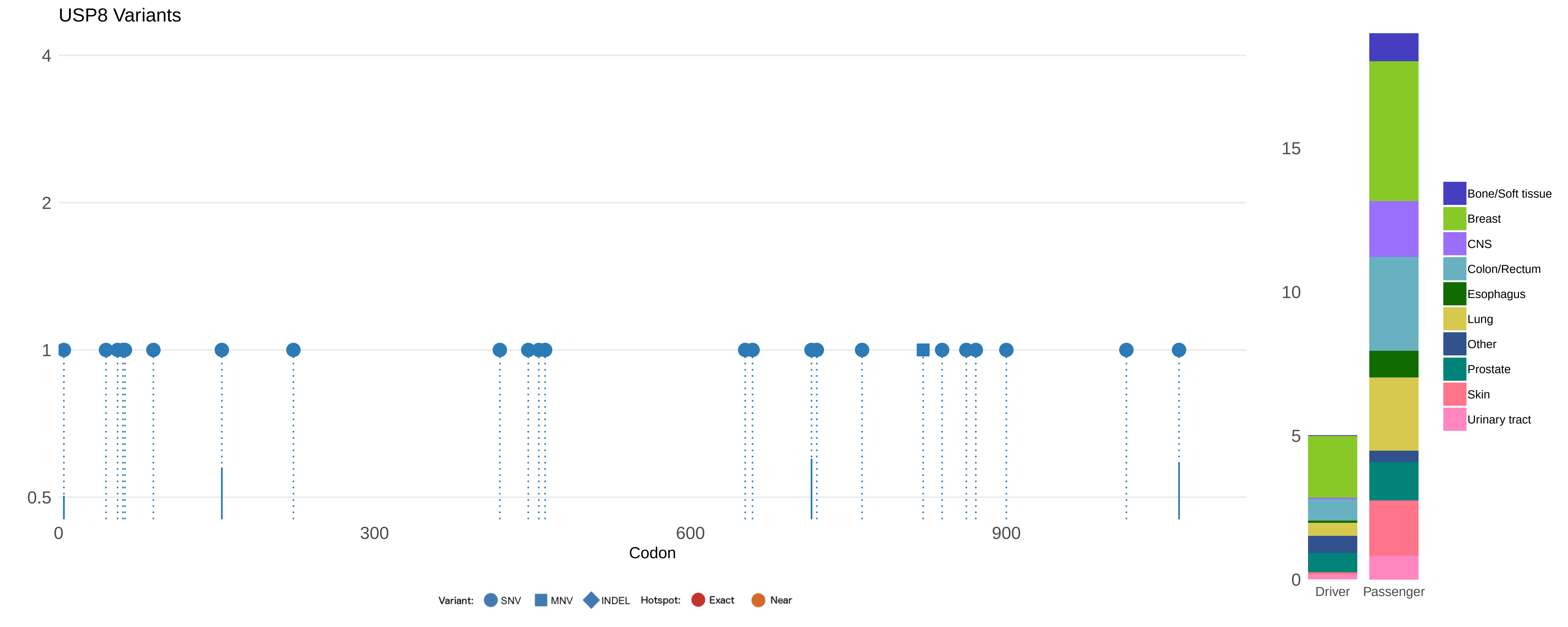
USP8 Variants
4
2
1
0.5
0
300
600
900
Codon
Variant: SNV MNV INDEL Hotspot: Exact Near
15
10
5
0
Driver
Passenger
Bone/Soft tissue
Breast
CNS
Colon/Rectum
Esophagus
Lung
Other
Prostate
Skin
Urinary tract

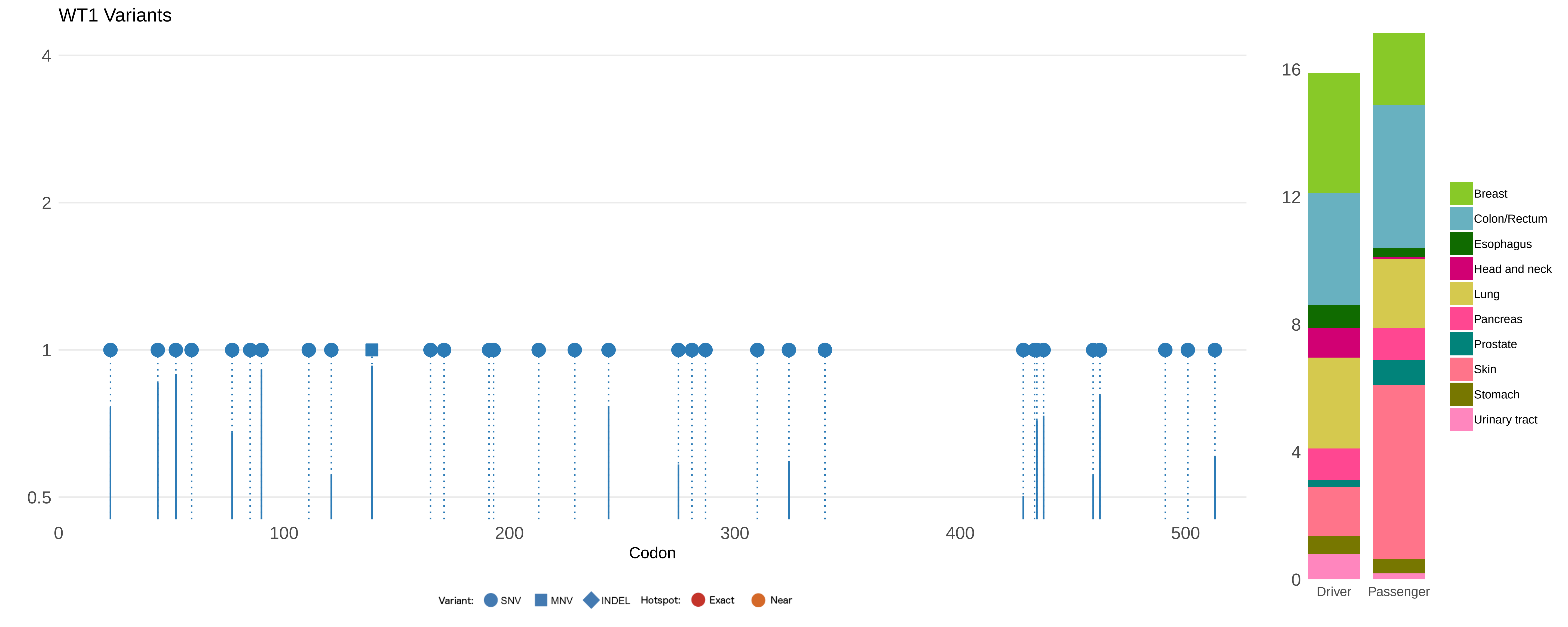

WT1 Variants
4
2
1
0.5
0
100
200
300
400
500
Codon
Variant:
SNV
MNV
INDEL
Hotspot:
Exact
Near
16
12
8
4
0
Driver
Passenger
Breast
Colon/Rectum
Esophagus
Head and neck
Lung
Pancreas
Prostate
Skin
Stomach
Urinary tract

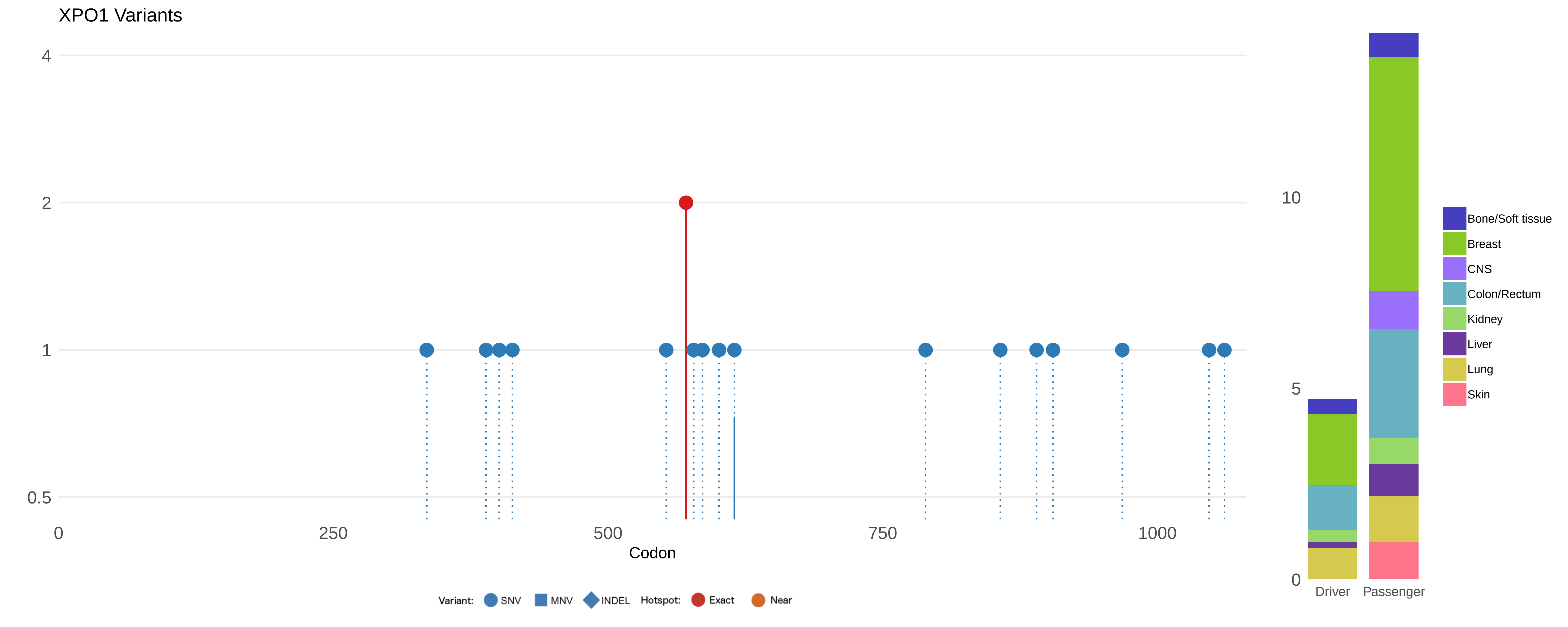
XPO1 Variants
4
2
1
0.5
0
250
500
750
1000
Codon
Variant: SNV MNV INDEL Hotspot: Exact Near
10
5
0
Driver
Passenger
Bone/Soft tissue
Breast
CNS
Colon/Rectum
Kidney
Liver
Lung
Skin

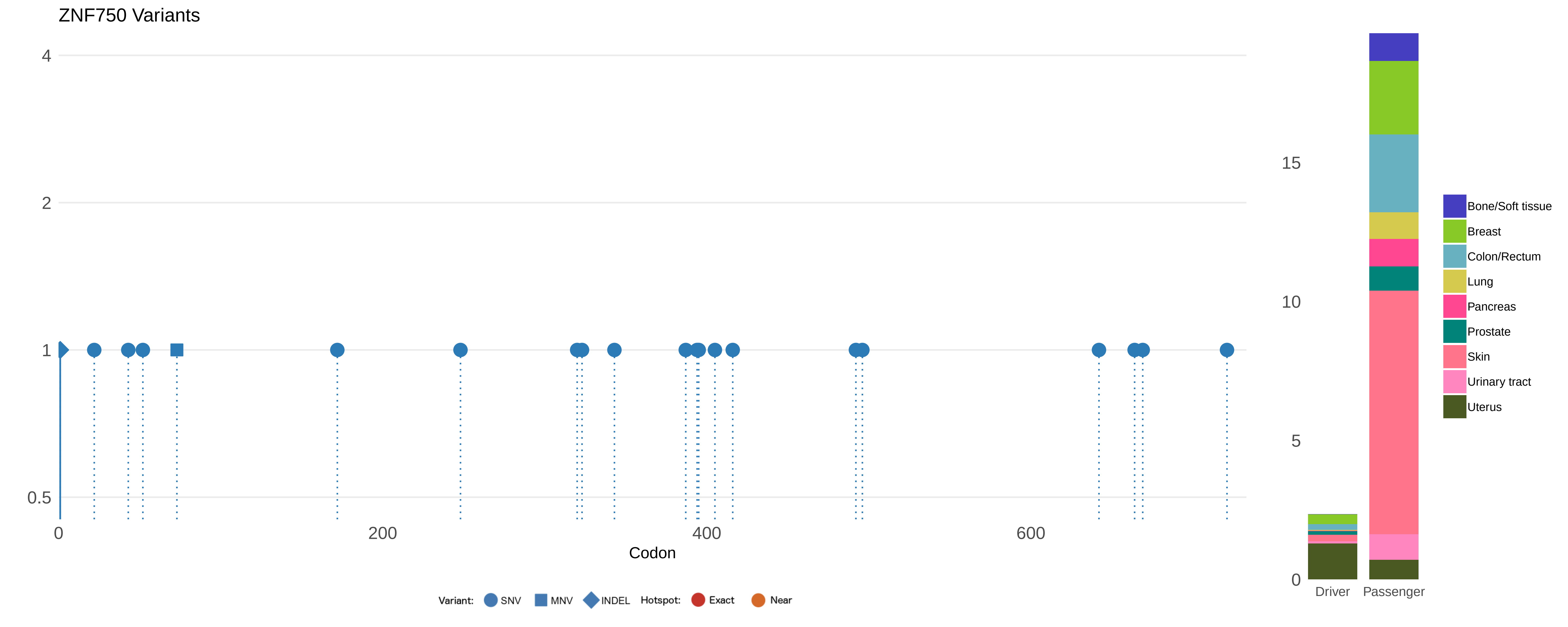

ZNF750 Variants
4
2
1
0.5
0
200
400
600
Codon
Variant:
SNV
MNV
INDEL
Hotspot:
Exact
Near
15
10
5
0
Driver
Passenger
Bone/Soft tissue
Breast
Colon/Rectum
Lung
Pancreas
Prostate
Skin
Urinary tract
Uterus
